# Developmental and Intergenerational Landscape of Human Circulatory Lipidome and its Association with Obesity Risk

**DOI:** 10.1101/2021.04.23.437677

**Authors:** Sartaj Ahmad Mir, Li Chen, Satvika Burugupalli, Bo Burla, Shanshan Ji, Adam Alexander T. Smith, Kothandaraman Narasimhan, Adaikalavan Ramasamy, Karen Mei-Ling Tan, Kevin Huynh, Corey Giles, Ding Mei, Gerard Wong, Fabian Yap, Kok Hian Tan, Fiona Collier, Richard Saffery, Peter Vuillermin, Anne K. Bendt, David Burgner, Anne-Louise Ponsonby, Yung Seng Lee, Yap Seng Chong, Peter D Gluckman, Johan G. Eriksson, Peter J. Meikle, Markus R. Wenk, Neerja Karnani

**Affiliations:** Department of Biochemistry, Yong Loo Lin School of Medicine, National University of Singapore, Singapore; Singapore Lipidomics Incubator, Life Sciences Institute, National University of Singapore, Singapore; Singapore Institute for Clinical Sciences, Agency for Science, Technology and Research, Singapore; Metabolomics Laboratory, Baker Heart and Diabetes Institute, Australia; KK Women’s and Children’s Hospital, Singapore; School of Medicine, Deakin University, Australia; Child Health Research Unit, Barwon Health, Australia; Murdoch Children’s Research Institute, University of Melbourne, Australia; The Florey Institute of Neuroscience and Mental Health, Australia; Department of Pediatrics, Yong Loo Lin School of Medicine, National University of Singapore, Singapore; Department of Obstetrics and Gynaecology, Yong Loo Lin School of Medicine, National University of Singapore, Singapore; Centre for Human Evolution, Adaptation and Disease, Liggins Institute, University of Auckland, Auckland, New Zealand; Folkhalsan Research Center, Helsinki, Finland; Department of General Practice and Primary Health Care, University of Helsinki, Helsinki, Finland; Bioinformatics Institute, Agency for Science, Technology and Research, Singapore

## Abstract

Lipids play a vital role in human health and development, but changes to their circulatory levels during gestation and in early life are poorly understood. Here we present the first developmental and intergenerational landscape of the human circulatory lipidome, derived by profiling of 480 lipid species representing 25 lipid classes, in mothers and their offspring (n=2491). Levels of 66% of the profiled lipids increased in maternal circulation during gestation, while cord blood had higher concentrations of acylcarnitines and lysophospholipids. The offspring lipidome at age six years revealed striking similarities with postnatal maternal lipidome (adult) in its lipid composition and concentrations. Comparison of lipids associated with child and maternal adiposity identified a 92% overlap, implying intergenerational similarities in the lipid signatures of obesity risk. We also catalogued lipid signatures linked with maternal adiposity during gestation and offspring birthweight, and validated (>70% overlap) the findings in an independent birth-cohort (n=1935).

## Introduction

Pregnancy is associated with changes to lipid metabolism, and an increase in the plasma lipid concentration as gestation progresses^1^. These changes are essential to maintain a constant supply of nutrients to the gestating mother and the growing fetus^2^. Maternal dyslipidemia has been associated with pregnancy-related complications and perinatal outcomes such as preeclampsia, gestational diabetes, preterm birth and macrosomia^3,4^. There is also mounting evidence that in obese glucose-tolerant mothers, lipids contribute more strongly to excess fetal fat accretion than glucose^5,6^. *In utero* nutrition plays a crucial role in fetal development. Sub-optimal *in utero* experience may not only influence the birth outcomes, but also alter the developmental programming to elevate the risk for chronic diseases later in life^7,8^. However, molecular mechanisms, whereby antenatal nutrition affects fetal development, remain underexplored. Population-based studies have increased our understanding of metabolism related to maternal and child health outcomes^9,10^. Recent studies have also provided evidence of changes in metabolic profiles across gestation^11,12^. However, there is a lack of population-level lipidomic studies to define longitudinal and intergenerational lipid profiles, as well as their association with adiposity. Here, we studied the changes to circulatory lipidome of women during pregnancy (antenatal vs postnatal plasma), and their offspring during development (cord blood and 6-year-old child plasma) in the GUSTO (Growing Up in Singapore Towards healthy Outcomes) birth cohort. We also delineated the associations of circulatory lipid profiles with pediatric and adult adiposity, and studied their overlap. We validated these findings in an independent birth cohort, the Barwon Infant Study (BIS), comprising mother-offspring dyads of different ethnic origin (Caucasian), recruited using an unselected antenatal sampling frame in the south east of Australia. This study provides a valuable resource for future developmental origins of health and disease (DOHaD) research.

## Results

### Overview of plasma lipidomics

A total of 480 lipid species representing 25 lipid classes were profiled across 2491 maternal and offspring plasma samples, covering 4-time points (antenatal and postnatal for mothers, and at birth and 6 years of age for their offspring) as illustrated in **Figure 1a**. Clinical characteristics of the study participants are provided in **Supplementary Table 1.** The combined unsupervised principal component analysis (PCA) identified two major components, with the first explaining 56.83% of the variance, and the second explaining 12.49% of the variance in the total study population. The PCA plot showed distinct clusters of mothers during gestation and post-pregnancy (antenatal vs postnatal), and also of their offspring at birth (cord blood) and 6 years of age **(Figure 1b).** The overlap in the clusters of postnatal mothers and their 6-year old offspring indicated the emerging similarities between the pediatric and adult lipidome. Subsequent analyses characterize and compare the antenatal and postnatal baseline lipids of mothers and their offspring using linear regression models and paired t-test (**Supplementary Table 2a-c)**. As both results were similar, only the linear regression results were discussed in this section.

**Figure 1.**
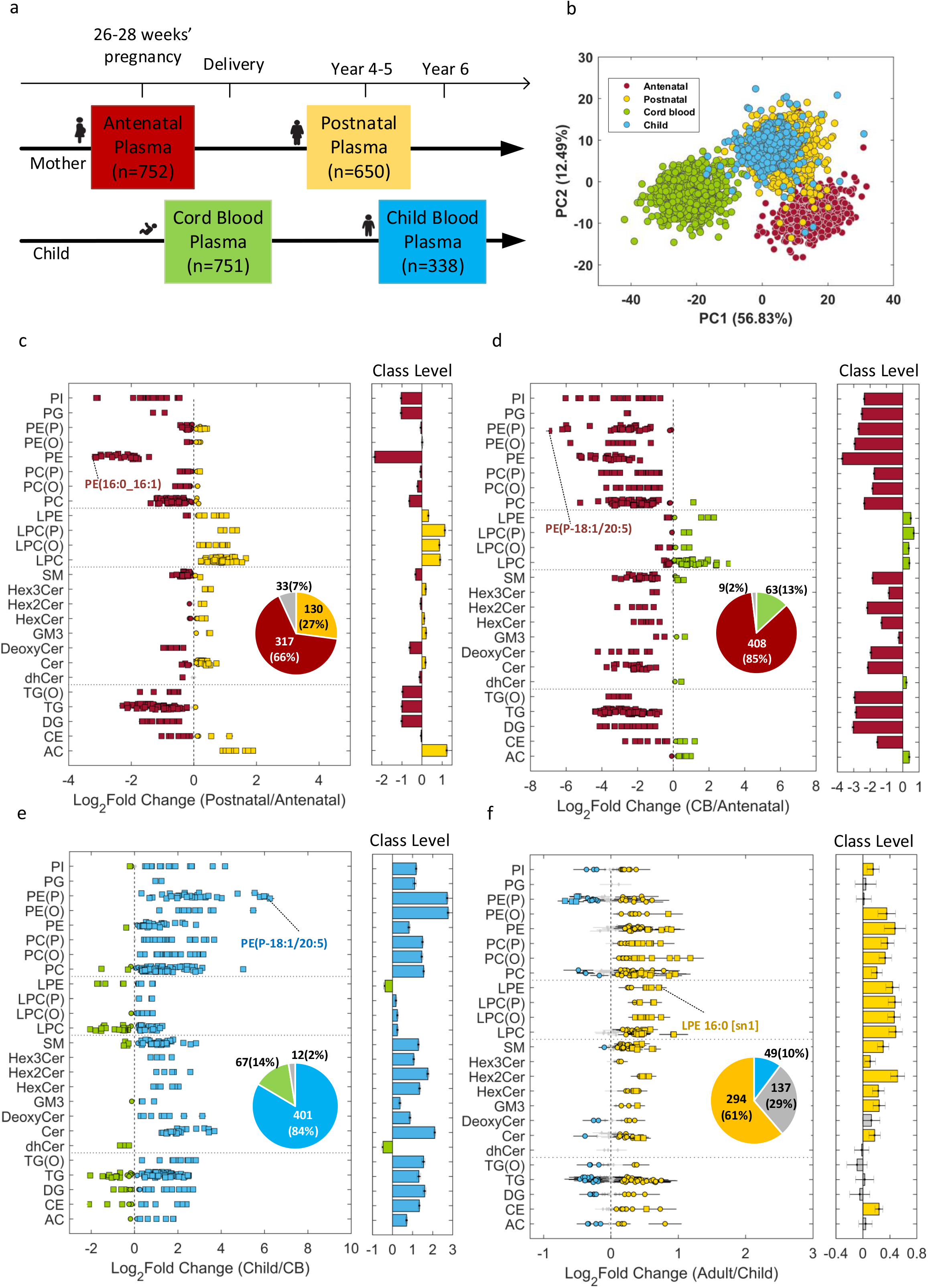
Temporal and developmental alterations to the baseline circulatory lipids in the GUSTO cohort: (a) Antenatal and postnatal plasma collection time points for mother-offspring dyads in GUSTO cohort. (b) PCA plot of lipidomics data (n=2491). (c) antenatal vs postnatal changes in maternal lipid profiles. (d) comparison of maternal antenatal plasma with cord blood (CB) lipid profiles. (e) changes in child lipid profile between birth and six years of age. (f) comparison of pediatric (six years old child) and adult (postnatal mothers) plasma lipidomes. The most significant lipid species based on adjusted p-values are shown in c-f. Effect size is shown as log_2_ of fold change. diamond – P_adj_ ≥ 0.05 (grey), circle – P_adj_ <0.05 and square – P_adj_ <1.00E-10.

#### Alterations to maternal circulatory lipidome during pregnancy

Alterations to the baseline circulatory lipids during pregnancy (antenatal vs postnatal) were assessed using linear regression models adjusted for the effects of mother’s ethnicity, pre-pregnancy BMI, age and education. 66% of the profiled plasma lipids were substantially elevated in pregnancy **(Figure 1c).** These primarily included phospholipids, such as phosphatidylcholine (PC), phosphatidylethanolamine (PE), phosphatidylinositol (PI) and phosphatidylglycerol (PG), with PE showing the most significant increase. PE (16:0_16:1) showed the greatest difference in circulation (log_2_FC=−3.16 and P_adj_<1.00E-30). Docosahexaenoic acid (DHA) containing lipid species like PE (16:0_22:6) and PE (18:0_22:6), as well as arachidonic acid (ARA) containing PE (16:0_20:4) and PE (18:0_20:4) were also elevated **(Supplementary Table 2b)**. The total PC to PE ratio was lower in pregnancy (**Supplementary Figure 1a**). However, the levels of LC-PUFA containing PC were higher than LC-PUFA containing PE. Additionally, ratios of DHA-PC to their corresponding DHA-PE species decreased by twofold or more in antenatal vs postnatal circulation **(Supplementary Figure 1b),** suggesting increased conversion of PE to PC by phosphatidylethanolamine N-methyltransferase (PEMT) in pregnancy as an important source for DHA-PC species in maternal plasma. Acylcarnitine (AC) species showed a general decrease in pregnancy. Lysophosphatidylcholine (LPC), lysophosphatidylethanolamine (LPE), alkyllysophosphatidylcholine (LPC (O)) and alkenyllysophosphatidylcholine (LPC (P)) were also low in pregnancy except saturated fatty acid containing species such as LPE (16:0). Although ether phospholipids including alkylphosphatidylcholine PC(O), alkenylphosphatidylcholine PC(P), alkenylphosphatidylethanolamine PE(P) and alkylphosphatidylethanolamine PE(O) were relatively unchanged, several polyunsaturated fatty acid (PUFA) containing plasmalogens such as PE (P-18:0/20:4), PE (P-18:0/20:5) and PE (P-18:1/20:4) were reduced in pregnancy. Amongst the sphingolipids, sphingomyelin (SM) were mostly elevated except SM(43:1), SM(44:1) and SM(44:2). Deoxyceramide (deoxyCer) species were higher in pregnancy whereas most ceramide (Cer) species containing very long chain fatty acid (VLCFA) were lower, dihydroceramide (dhCer) species such as dhCer(d18:0/22:0) and dhCer(d18:0/24:1) as well as several ceramide species containing saturated long chain fatty acid (LCFA) such as Cer(d18:1/16:0), Cer(d18:1/18:0), Cer(d18:1/20:0), and Cer(d18:2/16:0) were higher. GM3-ganglioside (GM3), monohexosylceramide (HexCer), dihexosylceramide (Hex2Cer) and trihexosylceramide (Hex3Cer) were low except HexCer(d18:1/18:0), HexCer(d18:1/20:0), Hex2Cer(d18:1/16:0) and Hex2Cer(d18:2/16:0) that showed an increase similar to that of the ceramide species. Neutral lipids including triacylglycerol (TG), alkyldiacylglycerol (TG(O)) and diacylglycerol (DG) were also substantially elevated in pregnancy. Cholesteryl esters (CE) species showed a divergent trend with a decrease in the levels of CE(18:0), CE(18:3), CE(20:4) and CE(20:5) in pregnancy **(Supplementary Table 2b)**.

#### Maternal-fetal lipid metabolism

To develop further insights into the maternal-fetal transfer of lipids, antenatal lipid profiles of mothers were compared to that of the offspring cord blood. In order to restrict this comparison to the baseline lipids, the linear regression models were adjusted for the effects of ethnicity, maternal education, pre-pregnancy BMI, maternal age, sex, gestational age and birth weight. Circulatory levels of 85% of the profiled lipids were substantially higher in maternal plasma than in cord blood **(Figure 1d)**. Concentration of all glycerophospholipid classes were lower in cord blood, while lysoglycerophospholipid classes were mostly higher with the exception of saturated lysophospahtidylcholine (LPC 16:0, LPC 18:0) and lysophosphatidylethanolamine (LPE 16:0, LPE 18:0), as well as several alkyllysophospatidylcholine and alkenyllysophospatidylcholine species. Levels of PUFA (ARA, dihomo-γ-linolenic acid (DGLA) and DHA) containing lipid species were elevated in cord blood. The highest fold change was observed in ARA containing lipids such as LPC 20:4 and LPE 20:4 (log_2_FC >2). Overall, PUFA containing LPC represented 27% of the total LPCs in cord blood, compared to 13% in antenatal maternal blood (**Supplementary Figure 1c**), indicating a higher fetal demand for these lipids to support its growth and development. Sphingolipid lipid species were mostly low in cord blood. Acylcarnitine species showed an overall increase in cord blood except AC(12:0) and AC(18:1). In summary, there was a generalized increase in acylcarnitines and lysophopholipids, but most of the complex lipids were present at lower levels in cord blood.

#### Alterations to circulatory lipids in early childhood, and a comparison with the adult lipidome

To interrogate the baseline alterations in the circulatory lipids in early childhood, we compared the plasma lipid profiles of the child at birth with that at 6 years of age. Linear regression models for this analysis were adjusted for ethnicity, sex, gestational age, birth weight and maternal education. A marked change (log_2_FC=−2.10 to 6.25) in the offspring lipidome was observed 6 years post-birth, with 84% of the lipids showing a relatively higher abundance, and 14% showing a lower abundance in circulation (**Figure 1e**). The lipid species that were present at higher concentrations in cord blood included PUFA containing lysoglycerophospholipid species such as LPC(20:4), LPC(22:6), LPE(20:4) and LPE(22:6) as well several diacylglycerol and triacylglycerol species such as DG(16:0_20:4), DG(16:0_22:6), TG(50:4) [NL-20:4], TG(56:8) [NL-22:6] and TG(58:8) [NL-22:6].

We also compared the 6-year-old pediatric and adult (postnatal mother: mean age 36 years) circulatory lipidome using linear regression models adjusted for sex, ethnicity, BMI and maternal education. There was a substantial overlap between the pediatric and adult plasma lipid profiles, and the differences were smaller in magnitude than those observed in pregnancy, and at birth. Levels of 61% lipid species in adults, and 10% in child, were marginally elevated (**Figure 1f**). A noticeable increase was observed in several DG and TG species including DG(16:0_16:1), DG(16:0_18:1), DG(18:0_18:1), TG(50:1) [NL-14:0], TG(50:1) [NL-16:0], TG(50:3) [NL-16:1], TG(52:4) [NL-16:1] and TG(54:1) [NL-18:1] that were relatively higher in child plasma. Although most of the glycerophosholipids including ether-linked phospholipids were decreased, several lipid species containing polyunsaturated omega-6 fatty acids such as PE(P-18:0/22:4), PE(P-18:1/22:4) and PE(P-16:0/22:4) were elevated in child plasma **(Supplementary Table 2b)**.

### Circulatory lipids associated with mother and child adiposity, and the intergenerational link

Adiposity is often associated with the changes in plasma lipid profiles, though less is known about the changes in physiological states such as pregnancy and during child development. To investigate this at depth, we studied the circulatory lipids associated with mother and child adiposity, and the intergenerational similarities between the two. For this four different regression models were implemented to identify the lipids associated with adiposity in each study group. Forest plots and bar plots were generated to visualize % change in the concentration of lipid species and lipid classes, respectively **(Figure 2a-d;** detailed in **Supplementary Table 3**.). The top 20 lipid species with the directionality of association (10 positive and 10 negative) with adiposity measures in each study group are shown through volcano plots, and the overlap between the subject groups is provided in **Figure 3** and **Supplementary Table 4**. Scatter plots comparing the effect sizes and adjusted p-values in the four association studies are shown in **Supplementary Figure 2**.

**Figure 2.**
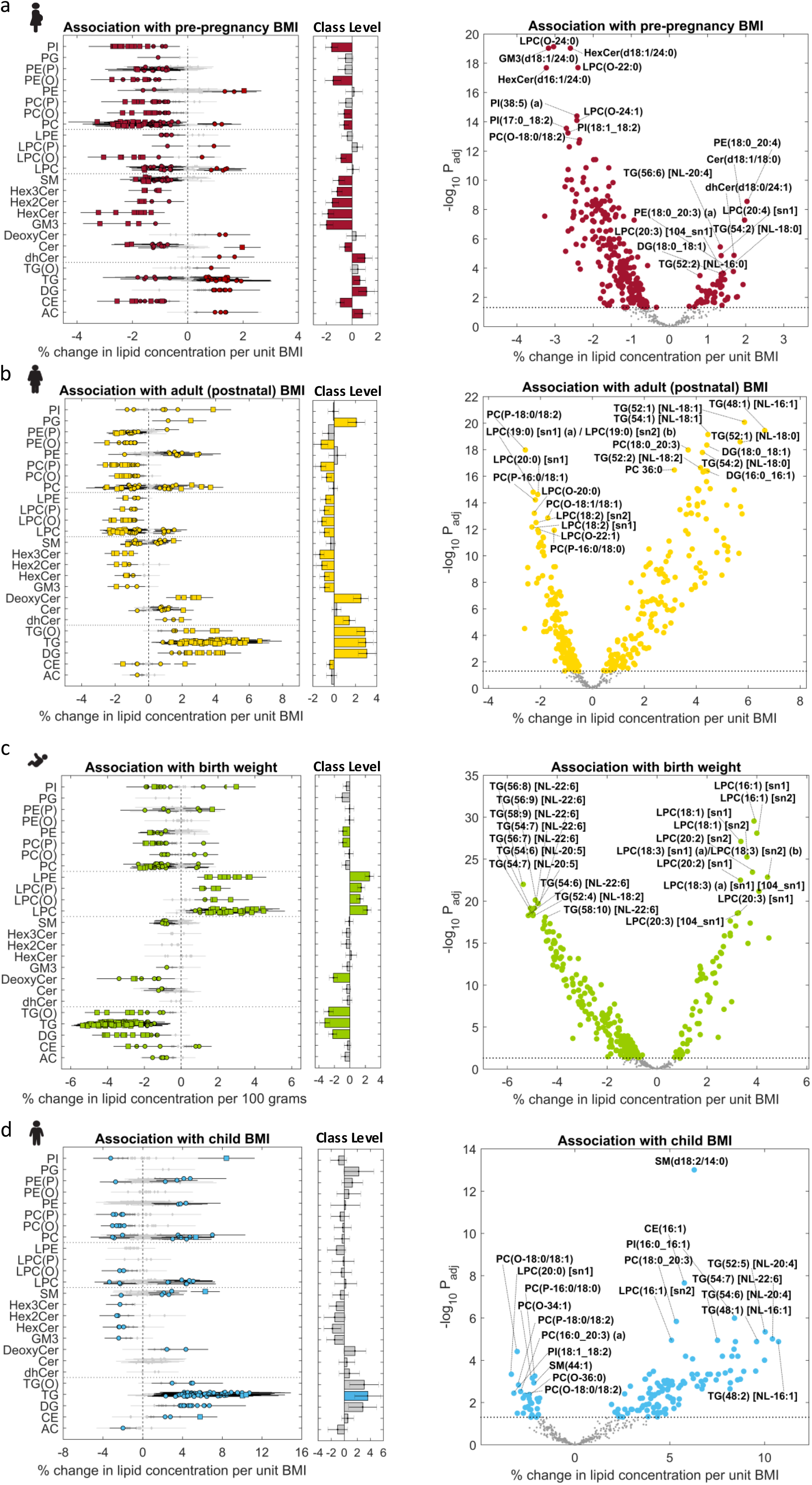
Association of maternal and child adiposity with circulatory lipids: (a) Association of pre-pregnancy BMI with antenatal plasma lipidome (pregnant state). (b) Association of maternal BMI with postnatal plasma lipidome (non-pregnant state). (c) Association of birth weight (BW) with cord blood plasma lipidome. (d) Association of child BMI with plasma lipidome at six years of age. The top 20 lipid species with the directionality of association (10 positive and 10 negative) with adiposity measures in each study group are shown through volcano plots. Effect sizes are shown as % change in lipid concentration per unit change in BMI, or per 100 grams change in birth weight. diamond – P_adj_ ≥ 0.05 (grey), circle – P_adj_ <0.05 and square – P_adj_ <1.00E-5.

**Figure 3.**
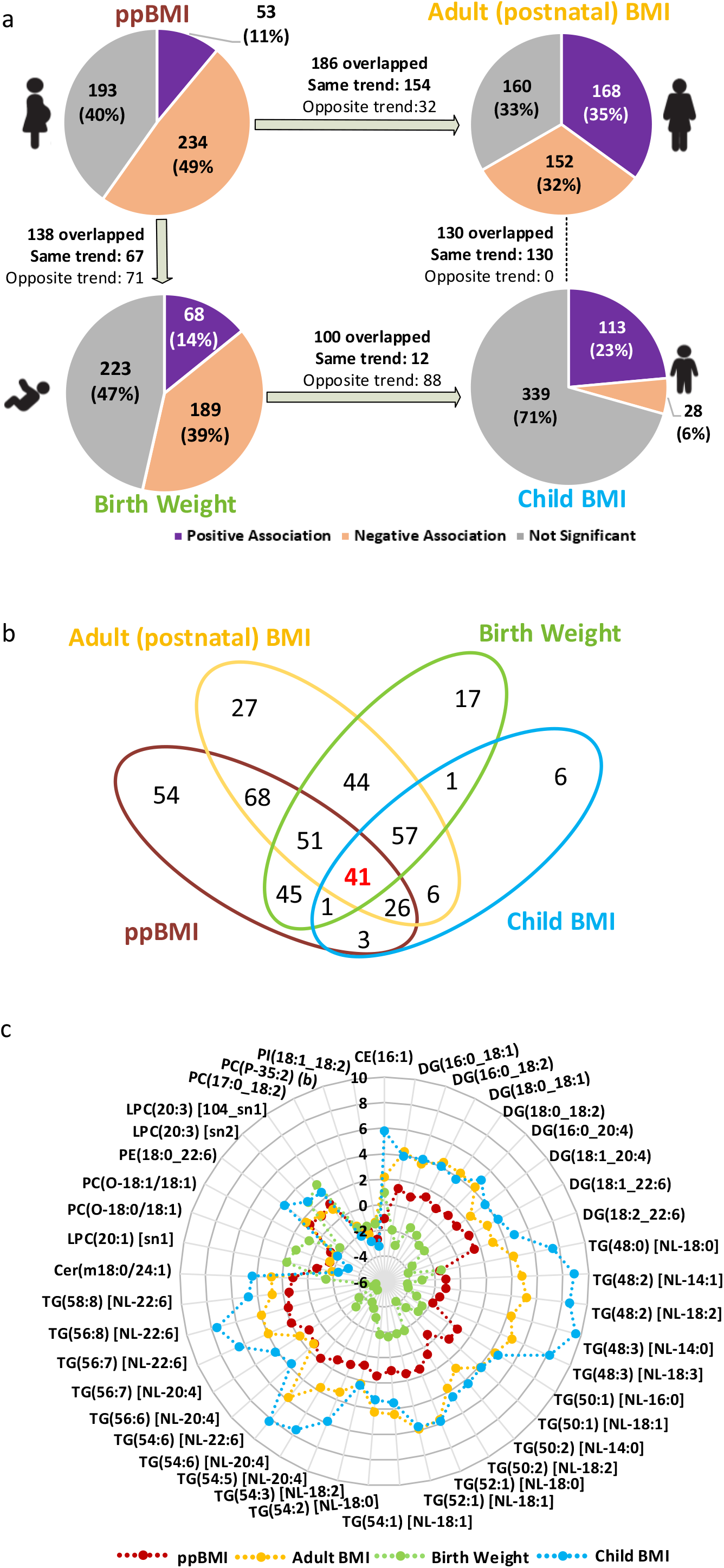
Comparison of circulatory lipid signatures associated with mother and child adiposity: (a) Pie chart comparing the percent overlap and directionality of association in the four studies. (b) Venn-diagram of lipids species that passed significance in the four association studies. (c) Effect sizes of 41 lipid species that overlapped between the four studies.

#### Maternal adiposity and circulatory lipids

The effects of pre-pregnancy BMI (ppBMI) on the antenatal lipid profile (pregnant state) were studied using linear regression models adjusted for maternal ethnicity, education, age and gestational weight gain at 26-28 weeks of pregnancy. 287 lipid species (60% of total) showed significant associations with ppBMI, of which 234 had positive and 53 had negative associations (**Figures 2a**). Most of the glycerophospholipids including ether-linked species showed a negative association with ppBMI, except for PUFA containing phosphatidylcholine species including PC(18:0_20:3) and PC(18:0_20:4), and phosphatidylethanolamine species such as PE(16:0_20:4), PE(18:0_20:4) and PE(18:0_22:6). Lysophosphatidylcholine species were negatively associated with maternal BMI except omega-6 fatty acid containing species including LPC(20:3) and LPC(20:4). Most of the triacylglycerol species were positively associated with ppBMI except those containing long chain fatty acids such as 14:0, 14:1, 16:1 and 18:3 that showed a negative association. Sphingolipids, including sphingomyelin, ceramide, mono-, di-, and trihexosylcermaide as well as GM3 gangliosides were negatively associated whereas deoxyceramide and dihydroceramide were positively associated with ppBMI. Acylcarnitine were positively and cholesteryl esters were negatively associated with ppBMI. Similar analysis post-pregnancy identified 320 lipid species (67%), with 168 positively associated and 152 negatively associated with postnatal BMI after accounting for the effects of ethnicity, education and age (**Figure 2b)**. Glycerophospholipids showed divergent trends in associations, lipid species containing polyunsaturated omega-6 fatty acids including LPC (20:3), PC(18:0_20:3), PC(18:1_20:3), PE(18:0_20:3), PI(16:0_20:4) and PE(18:0_22:4) were positively associated with BMI. Upon comparison, 186 BMI associated lipid species overlapped between the pregnant and non-pregnant states (**Supplementary Table 4a**). Among these, 154 lipid species (83%) showed consistent trends **(Supplementary Figure 3a)**, while the remaining 32 (17%) had opposite trends.

The lipid species that showed consistent trends belonged to ether-linked phospholipid classes including PC(O), PC(P), PE(O) and PE(P) as well as LPC(O) and LPC(P) that were negatively associated. Similarly, mono-, di- and trihexosylceramide and GM3 gangliosides were negatively associated whereas deoxyceramide and dihydroceramide showed positive association. Scatter plot comparing % change in the two studies showed a positive trend with R^2^=0.28 (**Supplementary Figure 2a-b**). Top 20 lipid species associated with BMI in pregnant and non-pregnant states are indicated in the volcano plots in **Figures 2a and 2b**, respectively.

#### Child adiposity and circulatory lipids

We next examined the relationship between plasma lipids and child adiposity at birth and 6 years of age. Since the inter-individual variation in BMI at birth is subtle and a less reliable measure, we instead used birth weight (BW) as a surrogate measure for adiposity. Association analysis with cord blood plasma lipids identified 257 (53%) lipid species to be significantly associated with BW by linear regression models adjusted for sex, ethnicity, maternal age, maternal education, pre-pregnancy BMI, total gestational weight gain, gestational age and parity (**Figure 2c and 3a**). Among these, 68 had positive and 189 had a negative association. LPC, LPC(O), LPC(P) and LPE were positively associated, while TG, TG(O) and DG were negatively associated with BW. TG species containing DHA showed strong negative association with BW, e.g.; (TG 56:8 [NL-22:6]: −5.36% change per 100 grams increase and P_adj_=1.02E-22) whereas several LPC and LPE species containing fatty acid 16:1 and 18:1 as well as omega-6 PUFA such as LPC 20:3, LPC 20:4 and LPE 20:4 were positively associated with BW. LPC 16:1 showed a strong positive association, with 3.89% increase in LPC 16:1 (sn1) (P_adj_=2.92E-30) and a 4.00% increase in LPC 16:1 (sn2) (P_adj_=8.10E-29) per 100 grams of BW.

To further compare the lipids linked with adiposity at birth and in early childhood, associations between lipid profiles of 6-year-old children and their BMI were compared with BW associated lipids. The linear regression models were adjusted for the effects of sex, ethnicity and maternal education. Plasma levels of 141 (29%, 113 high and 28 low) lipid species were substantially altered in children with higher BMI at age 6 (**Figure 2d and 3a**). The number of lipid species associated with BMI in children were lower across the lipid classes except that of TG species that showed positive associations. Glycerophospholipid classes showed divergent associations, several PUFA containing species PC(16:1_20:4), PC(18:0_20:3) and PE(18:0_20:3) as well as plasmalogen species PE(P-16:0/20:3), PE(P-16:0/20:5) and PE(P-18:0/20:3) were positively associated with BMI. Although few sphingolipid species were associated with BMI in children, SM (d18:2/14:0) showed strong positive association (6.29% change per unit BMI and P_adj_=9.93E-14). Furthermore, sex-stratified analysis identified several differences in association of lipid profiles with BW (**Supplementary Figure 4a-b)**, however, there was no difference in BMI associations between male and female offspring at 6 years (**Supplementary Figure 4c-d**). A higher number of lipid species were associated with BW in males (48%) as compared to females (31%). These differences were mainly driven by glycerophospholipid including ether-linked species, acylcarnitines, diacylglycerol and triacylglycerol species that showed a stronger association in males **(Supplementary Table 5)**. Comparison of BW and 6-year BMI associated lipids showed very different trends for most lipid classes. 88% of overlapping lipid species showed opposite trends and only 12 had a consistent trend **(Supplementary Figure 3c and Supplementary Table 4c)**. Scatter plot of effect sizes in two studies showed a negative trend with R^2^=0.31 (**Supplementary Figure 2e-f**). These findings indicate that the cord blood lipidome was associated with BW, however, these associations become weaker in early childhood, potentially due to postnatal exposures (**Supplementary Figure 2f**).

#### Intergenerational similarities in circulatory lipids associated with obesity risk

There is compelling evidence indicating that offspring of obese/overweight mothers bear a higher risk of having obesogenic growth trajectories and developing metabolic dysfunction/disease during their life-course^23^. Aligned with these reports, we found a significant correlation between the mothers and their offspring BMI. Having identified the circulatory lipids linked with mother and child obesity, we leveraged these findings to explore the intergenerational similarities through lipid signatures. Around 140 common lipid species were associated with ppBMI and BW, with 67 showing consistent trends **(Supplementary Figure 3b and Supplementary Table 4b)**. Comparison of child BMI-lipid associations with postnatal maternal-BMI lipids identified a 92% (130 of 141) overlap between the two, with consistent trends (**Figure 3a and Supplementary Table 4d**). Scatter plot of effect sizes confirmed a positive trend and strong correlation (R^2^=0.75) between the two (**Supplementary Figure 2g-h**). This observation not only confirmed the intergenerational link of obesity risk through lipid profiles but also indicated an early onset in childhood. To study the longitudinal and intergenerational obesogenic lipid signatures, we compared the adiposity associated lipids in all the 4 study groups (**Figure 3a**) and identified 41 overlapping lipid species (**Figure 3b-c and Supplementary Table 4e**). The directionality of association for most of the lipids showed opposite trend between BW and the other three studies, indicating that some lipids associated with adiposity in early childhood and adulthood may be beneficial for optimal fetal growth.

### Replication of obesity-linked lipid species in an ethnically diverse birth cohort

To investigate if the obesity-linked lipids were population-specific, or were independent of the ethnic effects and geographical location, we compared the findings from Asian-centric GUSTO cohort in Singapore with Caucasian-centric Barwon Infant Study (BIS) from Australia (**Figure 4a**). Due to the unavailability of the postnatal maternal and 6-year child plasma samples, we restricted the replication analysis to antenatal maternal blood (n=1042) and cord blood lipids (n=893). Similar to GUSTO, PCA plot in BIS also revealed distinct clusters for antenatal maternal and offspring cord blood samples (**Figure 4b)**. 464 of 480 lipid species profiled in GUSTO were also profiled in BIS and used for subsequent association analysis with maternal ppBMI and offspring BW (**Supplementary Table 6**).

**Figure 4.**
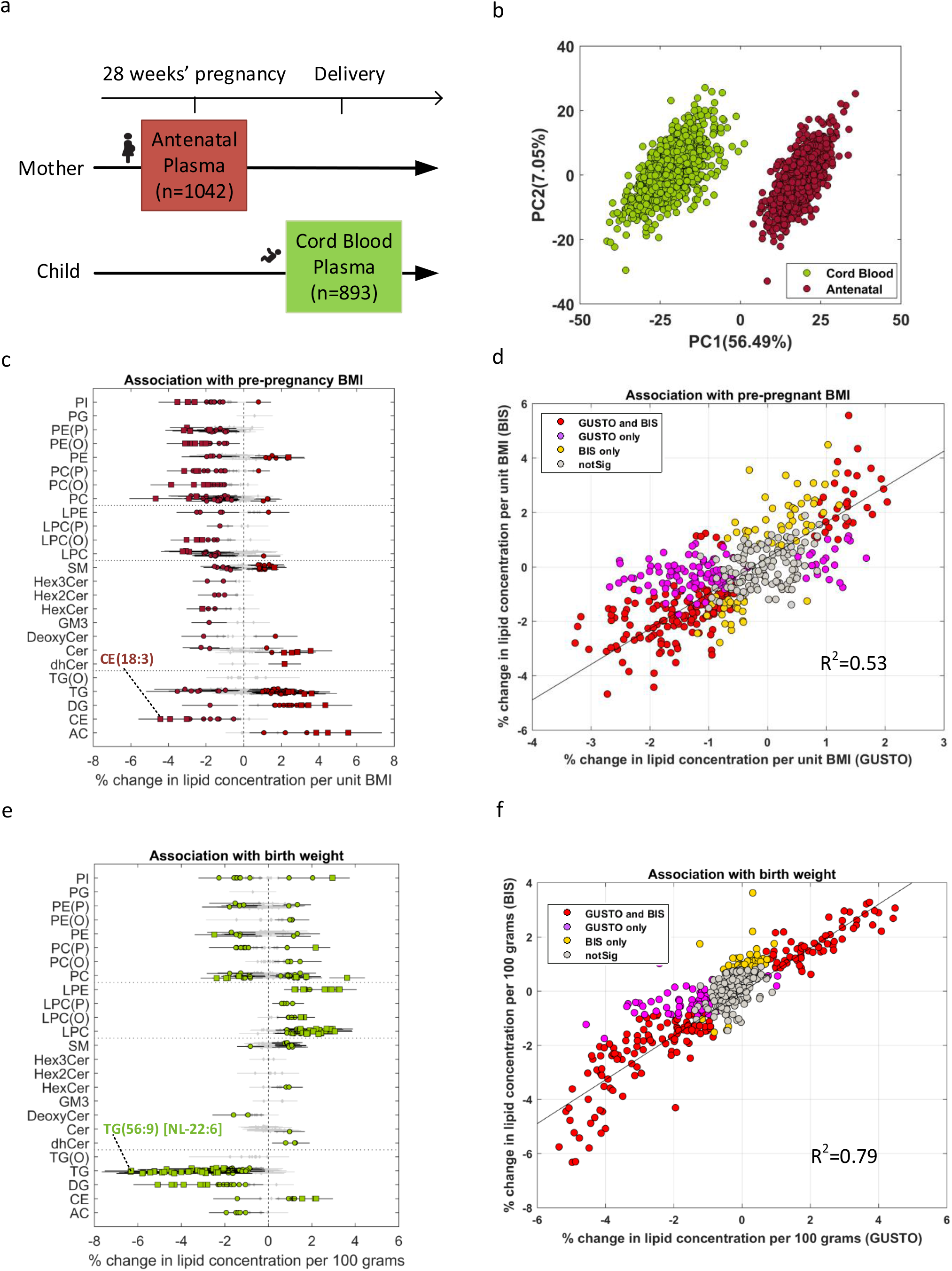
Replication of GUSTO identified ppBMI and BW lipid signatures in Barwon Infant Study (BIS). (a) Sample collection at two time points. (b) PCA plot of lipidomics data (n=1935). (c) Association of pre-pregnancy BMI (ppBMI) with antenatal plasma lipidome. (d) Scatter plot of effect sizes in GUSTO and BIS for ppBMI study. (e) Association of birth weight (BW) with cord blood plasma lipidome. (f) Scatter plot of effect sizes in GUSTO and BIS for BW study. Red-significant in both cohorts; purple – only significant in GUSTO; yellow – only significant in BIS; grey – not significant in both cohorts. The most significant lipid species based on adjusted p-values are labelled in c and e.

236 lipid species showed significant association with maternal BMI, of which 80 had positive and 156 had a negative association (**Figure 4c**). 170 of 236 (72%) lipid species overlapped with BMI-lipids identified in GUSTO mothers, and 80% of 464 lipid species had the same directionality of effect sizes in the two cohorts. Comparison of effect sizes in both the cohorts showed a significant positive trend with R^2^=0.53, though the effect sizes were generally larger in BIS subjects (**Figure 4d**). This could be due to a higher number of obese women or higher degree of metabolic dysregulation in the BIS cohort.

BW-lipid analysis identified 224 affected lipid species in BIS cord blood samples, of which 108 had positive and 116 had a negative association (**Figure 4e**). 176 of 224 (79%) lipids overlapped between the two cohorts. 81% of 464 lipid species showed the same directionality of effect sizes in the two cohorts, and their effect sizes were highly correlated (R^2^=0.79) (**Figure 4f**). Lipid species with discordant trends in the two cohorts can potentially be population-specific (Asian vs Australian). For example, SM showed negative association with ppBMI and BW in GUSTO, while it exhibited divergent trend with ppBMI and positive association with BW in BIS.

Meta-analysis in the two cohorts (**Supplementary Table 6**) identified 331 BMI and 295 BW associated lipid species, respectively (**Supplementary Figure 5a and 5c**). Top significant lipid species from the meta-analysis results are labelled in volcano plots (**Supplementary Figure 5b and 5d)**. 119 of 138 lipid species identified in ppBMI vs BW comparison in GUSTO (**Figure 3a and supplementary Table 4b)** were also significant in the meta-analysis, and these also included 35 of 41 lipid species identified in the 4 group intergenerational obesity risk analysis in GUSTO (**Figure 3c**).

## Discussion

Cross-sectional and longitudinal assessment of mother and child plasma lipidome in our study provided novel insights into the antenatal and postnatal circulating lipid species. It also identified similarities between child and adult lipidomes, and the intergenerational overlap in the circulatory lipids associated with adiposity. Characterization of baseline lipids not only confirmed the previous reports on the substantial increase in the levels of circulatory lipids in mid-late pregnancy^1,24,25^, but also provided the first in-depth profile of these changes covering 480 lipid species from 25 lipid classes. 66% of the profiled lipids increased in circulation in pregnancy, with phosphatidylethanolamine levels changing the most. In comparison to the gestating mother, cord blood showed lower concentration of most lipids, indicating selective transport and specific developmental needs of the growing fetus. A substantial number of proteins involved in lipid metabolism are expressed in fetal tissues^16^. However, the fetus is unable to independently support and maintain the circulatory lipid pools, which are complemented by maternal-fetal transport. The decrease in the levels of lysophospholipids, but elevated levels of phospholipids in maternal plasma is suggestive of the decreased activity of phospholipase A2 and lecithin-cholesterol acyltransferase^26^ in mid to late gestation, possibly to facilitate the placental uptake of intact phospholipid species for further metabolism and transport as lyso-PL into fetal circulation. The ratios of DHA-PC to DHA-PE decreased in pregnancy. Also, there was a significant decrease in DHA-LPC in antenatal maternal plasma as compared to both cord blood and maternal postnatal plasma. Taken together, these data suggest PC as a substantial contributor of DHA in maternal circulation and supply to the fetus^27,28^. Likewise, compared to cord blood and postnatal plasma, acylcarnitines were significantly lower in the antenatal plasma. These data indicate the potential role of lysophospholipids and acylcarnitines as molecular transducers in maternal-fetal lipid transport.

Post-birth, significant changes were evident in the offspring plasma lipidome by six years of age. The early childhood lipidome also showed a strong resemblance to the postnatal circulatory lipidome of the mother (adult), although most lipids were present at a higher concentration in the mother. However, there were higher levels of several polyunsaturated omega-6 containing glycerophospholipid species, including ether-linked phospholipids such as plasmalogens, particularly those with fatty acid 22:4. These lipid species may represent an essential pathway of ARA utilization to meet increasing demands for neuronal growth and development in early childhood^17–19^. Additionally, several triacylglycerol species containing long chain saturated and monounsaturated fatty acids were also higher in child plasma, suggesting an increase in *de novo* fatty acid synthesis with possible implications on metabolism in children^32,33^. As is evident from our analyses (birth to 6 years of age, and child to adult comparisons), levels of more than half of the profiled lipid species increased in circulation with age. The changes in lipids from birth to 6 years of age (log_2_FC=−2.10 to 6.25) were much higher in magnitude than those observed between pediatric and adult subjects (log_2_FC=−0.68 to 1.18) **(Figure 1e-f).** These findings indicate that lipid profiles are age-specific, and undergo a significant change in their levels in circulation to support growth and development during early childhood.

Our study also advances current understanding of the association between lipids and adiposity, especially in the context of the mother and child, and their intergenerational link. We observed an 83% overlap in the lipid species associated with BMI in pregnant vs non-pregnant states, suggesting a physiological state independent association of these lipids with adiposity. Among these, sphingolipids such as deoxyCer, and neutral lipids within DG and TG(O) classes showed a positive association with BMI, while ether-linked phospholipids and glycosphingolipids showed negative associations. These observation are also consistent with previous lipidomic studies of BMI in adult populations^34–36^. Notably, we also observed 17 % of the BMI associated lipids had divergent trends in pregnancy compared to the non-pregnant state. This was more apparent in 8 lipid classes (AC, CE, TG, PC, PE, PI, SM and Cer). For example, several saturated and monounsaturated LCFA containing TG showed negative association with BMI in pregnancy, but a positive association in mothers post-pregnancy. Divergent trends of lipid-BMI associations may arise due to the alterations in energy utilization and adaptive metabolism in obese women during pregnancy. Future work focusing on these lipids will provide deeper insights into the mechanism(s) involved in an obesogenic environment during gestation.

Comparison of lipids associated with child adiposity at birth vs 6 years of age showed opposite trends for most lipid classes, with only 12% of the lipids, belonging to 5 lipid classes (AC, CE, LPC, PC and PI; **Supplementary Figure 3c**) showing consistent trends and role in later adiposity. The divergent trends between BW-lipids and child BMI-lipids could arise due the effects of postnatal exposures on lipid metabolism during development. Notably, lipid species associated with child BMI were very similar to those associated with adult BMI. Several omega-6 fatty acid containing lipid species were significantly associated with ppBMI, birthweight and child BMI, suggesting these represent intergenerational signatures of obesity risk. Validation of ppBMI and BW findings in an independent birth cohort (BIS) identified lipid signatures that were both shared and unique to each cohort, although the majority of the signatures (~80%) remained consistent and had the same directionality in the two cohorts.

This study has a few limitations. Our analyses and interpretations of antenatal lipids are restricted to lipid levels in mid-pregnancy, as the blood samples were collected at only one time point in the cohort. Future studies in cohorts with multiple sampling may help profile longitudinal lipid trajectories from preconception to postnatal period, and also study causal relationship between lipids and obesity risk in women. The lack of postnatal maternal and offspring 6-year blood samples in BIS limited our ability to replicate some of the findings from the GUSTO cohort. Nevertheless, comparison of the antenatal and cord blood lipid profiles between the two cohorts identified a significant overlap between the baseline and obesity-linked lipid signatures. Pre-pregnancy maternal weight and maternal education level were self-reported during recruitment. Hence, bias may exist in the self-reported variables. However, self-reported ppBMI was highly correlated with BMI taken during the recruitment visit in early pregnancy **(Supplementary Figure 6)**.

In summary, and to the best of our knowledge, this is the first study to describe the population-based developmental and intergenerational landscape of the circulating lipidome. We characterized the longitudinal changes to the baseline lipidome, and also identified lipid signatures linked with obesity risk in women and children. Thus, our study provides an important resource for future research targeting early nutritional interventions to benefit maternal and child metabolic health.

## Acknowledgement

This work was supported by the A*STAR-NHMRC joint call funding (1711624031) available to NK, PJM, MRW, DB, ALP, RS, GW and PV. Translational Clinical Research (TCR) Flagship Program on Developmental Pathways to Metabolic Disease funded by the National Research Foundation (NRF) and administered by the National Medical Research Council (NMRC), Singapore - NMRC/TCR/004-NUS/2008 provided the funding support for subject recruitment and sample and data collection in GUSTO cohort study. Additional funds for data analysis in Singapore were supported by the Strategic Positioning Fund and IAFpp funds (H17/01/a0/005) available to NK through the Agency for Science, Technology and Research (A*STAR). The Singapore Lipidomics Incubator (SLING) is supported by grants from the National University of Singapore via the Life Sciences Institute, the National Research Foundation (NRF, NRFI2015-05 and NRFSBP-P4) and the NRF and A*STAR IAF-ICP I1901E0040. This work was also supported in part by the Victorian Government’s Operational Infrastructure Support Program. We wish to thank the study participants from GUSTO and BIS cohorts, for making this project possible as well as the staff at the respective institutions for assistance in project management.

## Author Contributions

NK, MRW and PJM conceived and supervised the study. YSL, JGE, YSC, NK PDG contributed to data and sample collection in GUSTO cohort. DB, ALP, RS, and PV contributed to data and sample collection in BIS cohort. SAM, BB, KN, DM, AR, KMLT, AKB, PJM, MRW and NK designed and performed experimental work. SAM, LC, BB and SJ performed the pre-processing and quality control, and LC carried out statistical analysis and association studies in GUSTO cohort. SB, AAT, KH and CG contributed to data analysis in BIS cohort. SAM, LC, PJM, MRW and NK interpreted the data and wrote the manuscript. All authors critically read and approved the manuscript content.

## Supplementary Figures

**Supplementary Figure 1**. Ratios of PC to PE levels and lysophospholipids (lysoPL): **(a)** Ratios of total PC to total PE levels in four sample groups. **(b)** Ratios of PC and PE lipid species containing DHA in antenatal and postnatal samples. **(c)** Percentage of lysoPL containing saturated fatty acid (SFA), monounsaturated fatty acid (MUFA) and polyunsaturated fatty acid (PUFA) in antenatal, postnatal, cord blood and child plasma.

**Supplementary Figure 2.** Scatter plots of effect sizes and adjusted p-values of circulatory lipids associated with mother and child adiposity: **(a-b)** Pre-pregnant BMI study vs. Adult BMI (postnatal) BMI study. **(c-d)** Pre-pregnant BMI study vs. BW study. **(e-f)** BW study vs. Child BMI study (6 years of age). **(g-h)** Child BMI study vs. Adult (postnatal) BMI study. Effect size is shown as % change in lipid concentration per unit BMI for BMI associations and per 100 grams for birth weight. The most significant lipid species are labelled in b, d,f and h. Related to Figure 2.

**Supplementary Figure 3.** Overlapping circulatory lipid signatures associated with mother and child adiposity: (a) Heat map of 154 overlapping lipid species with the same directionality of association between antenatal ppBMI and adult (postnatal) BMI study. (b) Heat map of 68 overlapping lipid species with the same association trend between antenatal ppBMI and cord blood BW study. (c) Heat map of 12 overlapping lipid species with the same association trend between cord blood BW study and child BMI study. Related to Supplementary Table 4.

**Supplementary Figure 4.** Sex difference in the birth weight and child BMI studies: **(a)** Association with birth weight in males. **(b)** Association with birth weight in females. **(c)** Association with child BMI in males. **(d)** Association with child BMI in females. diamond – P_adj_ ≥ 0.05 (grey), circle – P_adj_ <0.05 and square – P_adj_ <1.00E-5. Related to Supplementary Table 5.

**Supplementary Figure 5.** Forest plots and volcano plots of the meta-analysis results in GUSTO and BIS for ppBMI (**a and b**) and BW studies (**c and d**). diamond – P_adj_ ≥ 0.05 (grey), circle – P_adj_ <0.05 and square – P_adj_ <1.00E-5 in forest plots (a and c). Top 10 significant lipid species with positive and negative association are labelled in volcano plots (b and d, dotted line – P_adj_=0.05). Related to Supplementary Table 6.

**Supplementary Figure 6.** Pre-pregnancy and postnatal maternal body mass index (BMI): (**a**) Pre-pregnancy BMI vs booking BMI measured at the first clinical visit. (**b**) Pre-pregnancy BMI vs postnatal BMI (4-5 years after delivery).

## Supplementary Tables

**Supplementary Table 1**. Clinical characteristics of study participants in this study.

**Supplementary Table 2a**. Comparison of baseline lipid classes across four sample groups (antenatal, postnatal, cord blood and 6-year-old child) using linear regression model. Related to Figure 1.

**Supplementary Table 2b**. Comparison of baseline lipids species across four sample groups (antenatal, postnatal, cord blood and 6-year-old child) using linear regression model. Related to Figure 1.

**Supplementary Table 2c**. Comparison of baseline lipids species across four sample groups (antenatal, postnatal, cord blood and 6-year-old child) using paired t-test.

**Supplementary Table 3a**. Association of lipid classes with BMI and BW in four sample groups (antenatal, postnatal, cord blood and 6-year-old child) in terms of levels. Related to Figure 2.

**Supplementary Table 3b**. Association of lipid species with BMI and BW in four sample groups (antenatal, postnatal, cord blood and 6-year-old child) in terms of levels. Related to Figure 2.

**Supplementary Table 4a**. The overlapping lipid species between the ppBMI and postnatal BMI studies. Related to Figure 3.

**Supplementary Table 4b**. The overlapping lipid species between the ppBMI and birth weight studies. Related to Figure 3.

**Supplementary Table 4c**. The overlapping lipid species between the birth weight and 6-year-old child BMI studies. Related to Figure 3.

**Supplementary Table 4d**. The overlapping lipid species between the 6-year-old child BMI and adult (postnatal mother) BMI studies. Related to Figure 3.

**Supplementary Table 4e**. Overlapping lipid species in the four lipid-adiposity association studies. Related to Figure 3.

**Supplementary Table 5a.** Sex difference in the birth weight and child BMI studies in terms of lipid class levels. Related to Supplementary Figure 4.

**Supplementary Table 5b.** Sex difference in the birth weight and child BMI in terms of species levels. Related to Supplementary Figure 4.

**Supplementary Table 6**. Association studies of lipids with pre-pregnancy BMI and birth weight in Barwon Infant Study (BIS), and the meta-analysis results of GUSTO and BIS. Related to Figure 4 and Supplementary Figure 5.

## Methods

### Study participants and clinical characteristics

GUSTO (Growing Up in Singapore Towards healthy Outcomes) is a prospective mother–offspring cohort study in Singapore consisting of 1,247 women recruited between June 2009 and September 2010^39^. Study participants were of Chinese, Malay or Indian ethnicity, but with homogeneous parental ethnic background. Written informed consent was obtained at recruitment, and ethics approval for the study was granted by the Institute Review Board of KK Women's and Children’s Hospital (KKH) and National University Hospital (NUH). This study was registered under ClinicalTrials.gov on 1 July 2010 under the identifier NCT01174875 (http://www.clinicaltrials.gov/ct2/show/NCT01174875?term=GUSTO&rank=2).

Data on maternal age, ethnicity, education level, self-reported pre-pregnancy weights were collected from the participants during recruitment. Pre-pregnancy BMIs (ppBMI) were calculated as pre-pregnancy weights divided by height squared. Gestational weight gain (GWG) up to 26-28 weeks of pregnancy and total GWG were calculated by subtracting self-reported pre-pregnancy weights from weights measured at 26-28 weeks of gestation and weights measured at delivery respectively. For offspring born to the women enrolled in this study, information on gestational age, birth weight, sex and parity were retrieved from birth delivery reports. At postnatal follow-up visit, age, weight and height of mothers were collected at 4-5 years after delivery. Age, weight and height of children were collected at six-year-old. Adult BMI and child BMI were calculated accordingly.

The Barwon Infant Study (BIS) is a major birth cohort study being conducted by the Child Health Research Unit (CHRU) at Barwon Health in collaboration with the Murdoch Children’s Research Institute (MCRI) and Deakin University^40^. The association studies with birth weight and pre-pregnancy BMI were used for replication of findings in GUSTO cohort.

### Sample selection, preparation and experimental design

A total of 2,519 plasma samples from GUSTO cohort were selected for this lipidomics study. They consisted of antenatal maternal samples (n=763) collected at 26-28 weeks of pregnancy, cord blood samples (n=767) collected at delivery, postnatal maternal (n=651) collected at 4-5 years after delivery and child samples (n=338) collected at six years of age. All plasma samples were prepared in 10 aliquots for lipid extraction. A pooled quality control was prepared from a subset of the representative samples from the study. This pooled QC was then used to prepare a batch QCs (BQC) and a technical QCs (TQC). These QCs serve well defined roles of monitoring and correcting for biases introduced during sample preparation, LC-MS/MS-based measurements as well as for determining signal-to-noise ratios, coefficient of variation of individual lipid species, linearity and stability of the analytical platform. A stratified randomization strategy was used to allocate the samples into seven batches. At the same time, blanks, process blanks (PBLK) and aliquots of reference plasma samples from NIST-1950 were interspersed in between the samples. QCs and blanks were processed and analyzed along with the study samples within each batch.

### Lipid extraction and LC-MS/MS analysis

The samples were extracted according to the stratified randomization template and 438 samples (study samples as well as QCs) were extracted in one batch with a total of seven batches for the current study. Lipid extraction was carried out using butanol: methanol (extraction solvent) in a ratio of 1:1 containing 10 mM ammonium formate and deuterated or non-physiological internal standards as described previously^41^. 100μl of extraction solvent was added to each sample, vortexed for 10 seconds followed by sonication for 60 minutes with temperature maintained at 18-22 °C. The samples were centrifuged at speed of 13,000 rpm for 10 minutes. The supernatant (80μl) was collected in mass spectrometry compatible vials and stored at −80 °C for LC-MS/MS. These lipid extracts were analyzed by using Agilent 6490 QQQ mass spectrometer interfaced with an Agilent 1290 series HPLC system. Lipids were separated on a ZORBAX RRHD UHPLC C18 column (2.1×100mm 1.8mm, Agilent Technologies) with the thermostat set at 45 °C. Mass spectrometry analysis was performed in ESI positive ion mode with dynamic multiple reaction monitoring (dMRM). Mass spectrometry settings and MRM transitions for each lipid class, subclass and individual species were kept as described previously^35,42^. The following mass spectrometer parameters were used: gas temperature, 150 °C, gas flow rate 17L/min, nebuliser 20psi, Sheath gas temperature 200 °C, capillary voltage 3500V and sheath gas flow 10L/min. Isolation widths were set to unit resolution for both Q1 and Q2. TQCs were monitored for changes in peak area, width and retention time to determine the performance of the LC-MS/MS analysis. QC samples were analyzed along with the samples to monitor sample extraction efficiency as well as LC-MS performance and were subsequently used to do batch corrections.

### Data analysis

Peak integration was carried out in MassHunter Quantitative software (Agilent Technologies) to select area of each individual lipid species. Manual inspection was carried out to ensure that correct peaks were picked at specific retention time. Peak areas along with retention times were exported as .csv for further analysis. These data were filtered by using a cut-off for signal-to-noise ratio (S/N>10) and linear response (R^2^>0.8) of each lipid species. Peak areas of lipid species were normalized to their class specific or to a closely eluting relevant internal standard as described previously^35,42^. Batch QCs was used to correct signal drifts across the batches based on LOESS^43^ regression method (locally polynomial regression fitting, R version 3.5.1). Following that, lipid species were dropped if quality control coefficient of variation were greater than 20%. These stringent quality control steps enabled us to measure lipid species with precision (average CV = 9.8%). The approach applied here provides relative quantitation as the use of isotope-labelled internal standards required for absolute quantitation of each lipid species under investigation is not feasible. Finally, a total of 480 lipid species representing 25 lipid classes were used for the downstream data analysis. In addition, we removed 28 outlier samples based on principal component analysis (PCA) which resulted in 2,491 samples for further downstream analysis.

### Statistical Analysis

All lipidomics data were log_10_ transformed for the downstream analyses. The unsupervised principal component analysis of lipid species data from all the samples was applied to estimate overall differences between the four sample groups (antenatal mother, cord blood, postnatal mother and child). Linear regression methods were used for comparative lipidomics studies between antenatal vs. postnatal, antenatal vs. cord blood, cord blood vs. child and child vs. adult (postnatal mother) using lipid levels as outcomes and groups as predictors. In each study, potential confounders were identified by checking the associations with top ten principal components of studied samples and adjusted in the regression models accordingly. The regression coefficients (β) were converted to log_2_ fold change (FC=10^β^). In addition, paired t-test was also applied to compare group difference using paired samples with a much smaller sample size, and the results were compared to the linear regression results.

Similarly, association of lipids with BMI and birth weight were studied in the four sample groups using linear regression. Pre-pregnancy BMI was studied in antenatal lipidome. Child BMI and adult BMI were examined in child lipidome and postnatal mother lipidome respectively. Birth weight (BW) was investigated in cord blood lipidome as BW is widely used as a risk factor for overweight/obesity at later ages in pediatrics. Potential confounders in each study were detected using the same method as above and adjusted in the linear regression models accordingly. The regression coefficients (β) were converted to % change in lipid concentration per unit BMI (ppBMI, child BMI and adult BMI), or % change in lipid concentration per 100 grams of BW (% change = (10^β^– 1) × 100). The adjusted p-values were calculated by the Benjamini-Hochberg (BH) method in each study. The lipid species with the adjusted p-value (P_adj_) < 0.05 were considered to be significant.

The findings from GUSTO study were replicated in the Barwon Infant Study (BIS) cohort. As described before, linear regression was applied to investigate the association between ppBMI and antenatal lipid profile (pregnant state), and the association between BW and lipid profile of cord blood plasma. The ppBMI and BW results were compared between GUSTO and BIS. Scatter plots of effect sizes in two cohorts were used for showing the linear trend and the overlapping lipid species in the two cohorts. The overlapping percentage of associated lipid species was calculated in each study. Finally, meta-analysis was performed for ppBMI and BW studies in the two cohorts using inverse variance-weighted average method^44^. Meta-analysis results were shown in forest plot and volcano plot.

All the association analyses were implemented in MATLAB R2019b.

