## Supplementary Data for "Developmental and Intergenerational Landscape of Human Circulatory Lipidome and its Association with Obesity Risk"

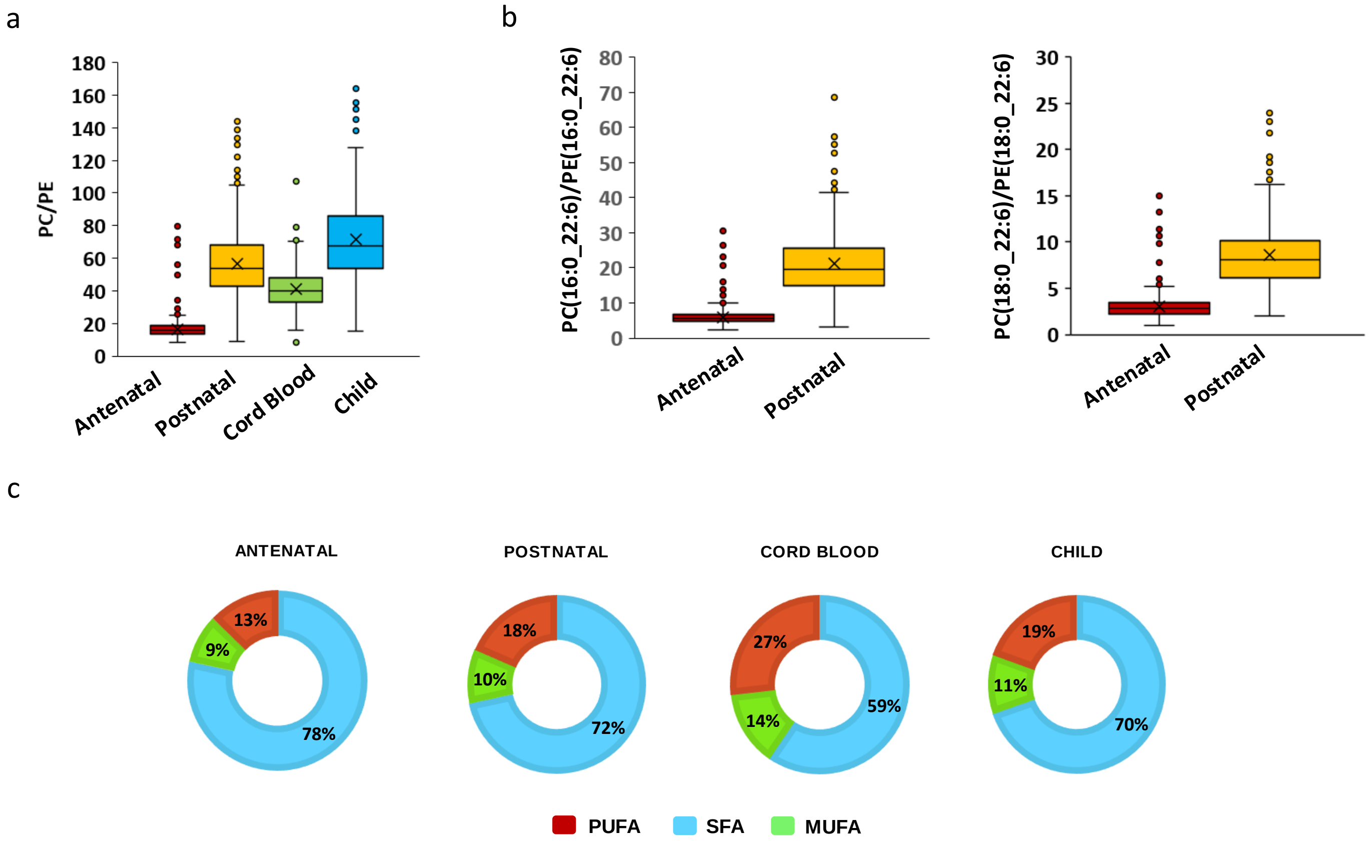

**Supplementary Figure 1**. Ratios of PC to PE levels and lysophospholipids (lysoPL): **(a)** Ratios of total PC to total PE levels in four sample groups. **(b)** Ratios of PC and PE lipid species containing DHA in antenatal and postnatal samples. **(c)** Percentage of lysoPL containing saturated fatty acid (SFA), monounsaturated fatty acid (MUFA) and polyunsaturated fatty acid (PUFA) in antenatal, postnatal, cord blood and child plasma.

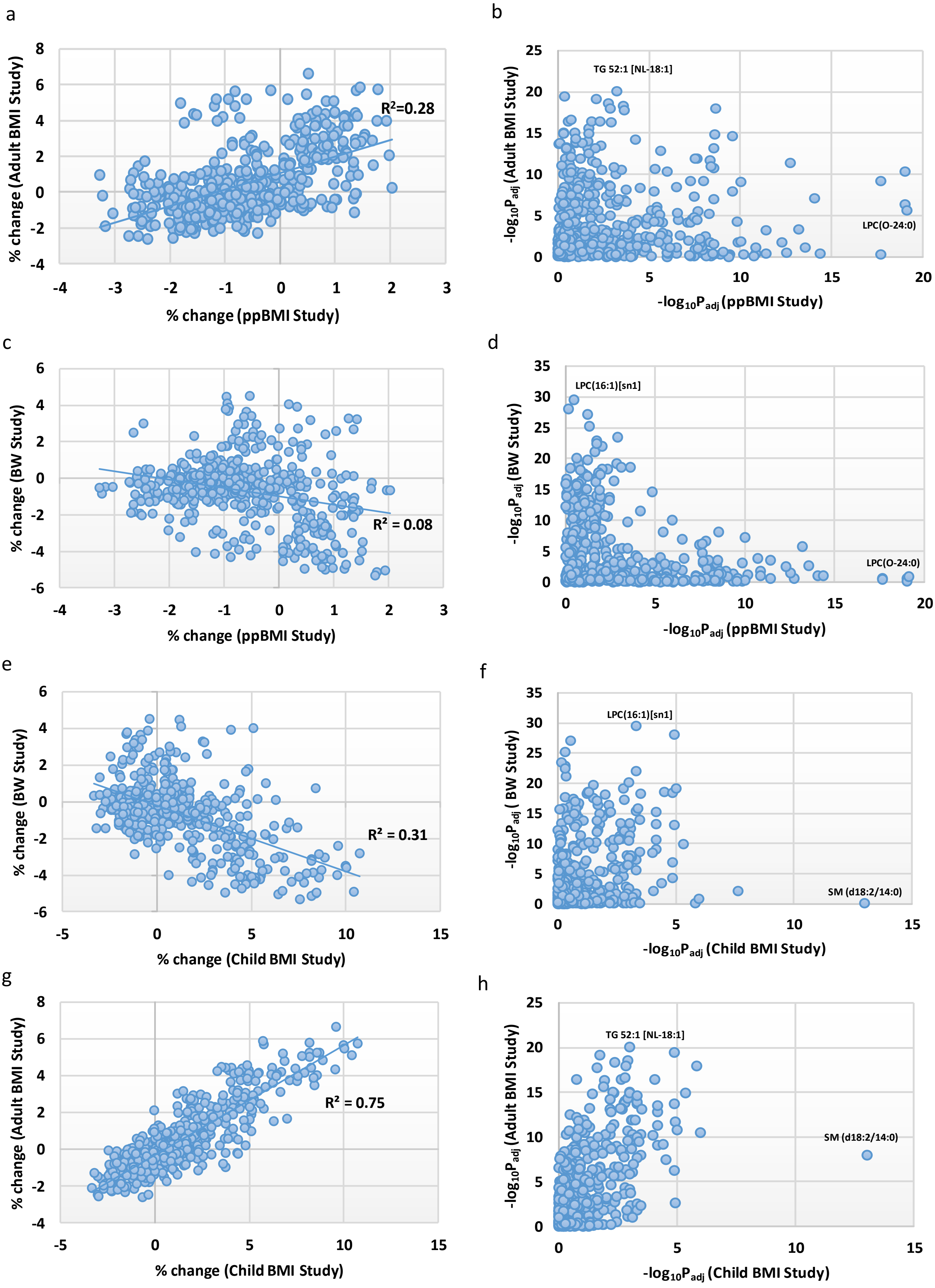

**Supplementary Figure 2.** Scatter plots of effect sizes and adjusted p-values of circulatory lipids associated with mother and child adiposity: **(a-b)** Pre-pregnant BMI study vs. Adult BMI (postnatal) BMI study. **(c-d)** Pre-pregnant BMI study vs. BW study. **(e-f)** BW study vs. Child BMI study (6 years of age). **(g-h)** Child BMI study vs. Adult (postnatal) BMI study. Effect size is shown as % change in lipid concentration per unit BMI for BMI associations and per 100 grams for birth weight. The most significant lipid species are labelled in b,d,f and h. Related to Figure 2.

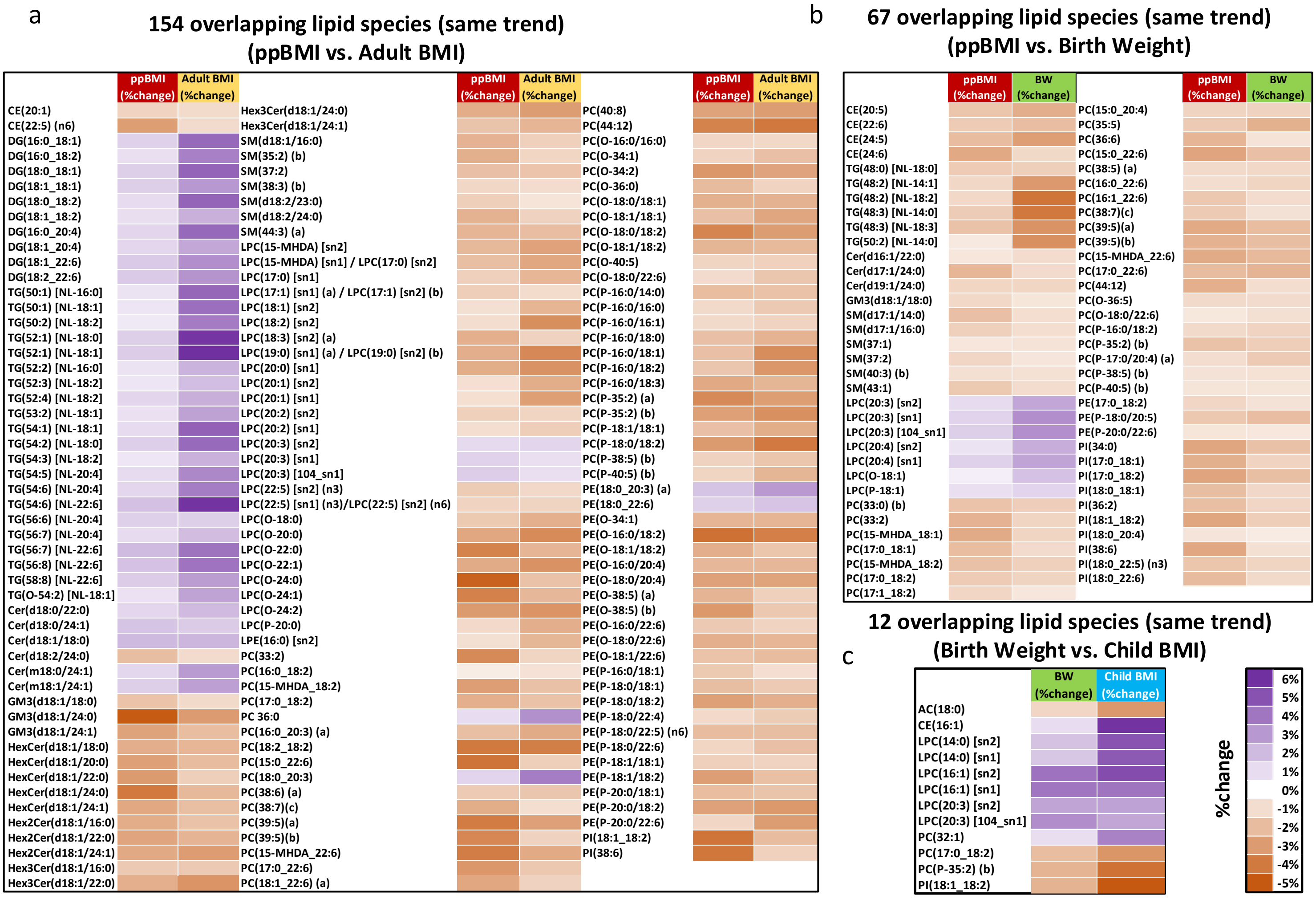

**Supplementary Figure 3.** Overlapping circulatory lipid signatures associated with mother and child adiposity: (a) Heat map of 154 overlapping lipid species with the same directionality of association between antenatal ppBMI and adult (postnatal) BMI study. (b) Heat map of 67 overlapping lipid species with the same association trend between antenatal ppBMI and cord blood BW study. (c) Heat map of 12 overlapping lipid species with the same association trend between cord blood BW study and child BMI study. Related to Supplementary Table 4.

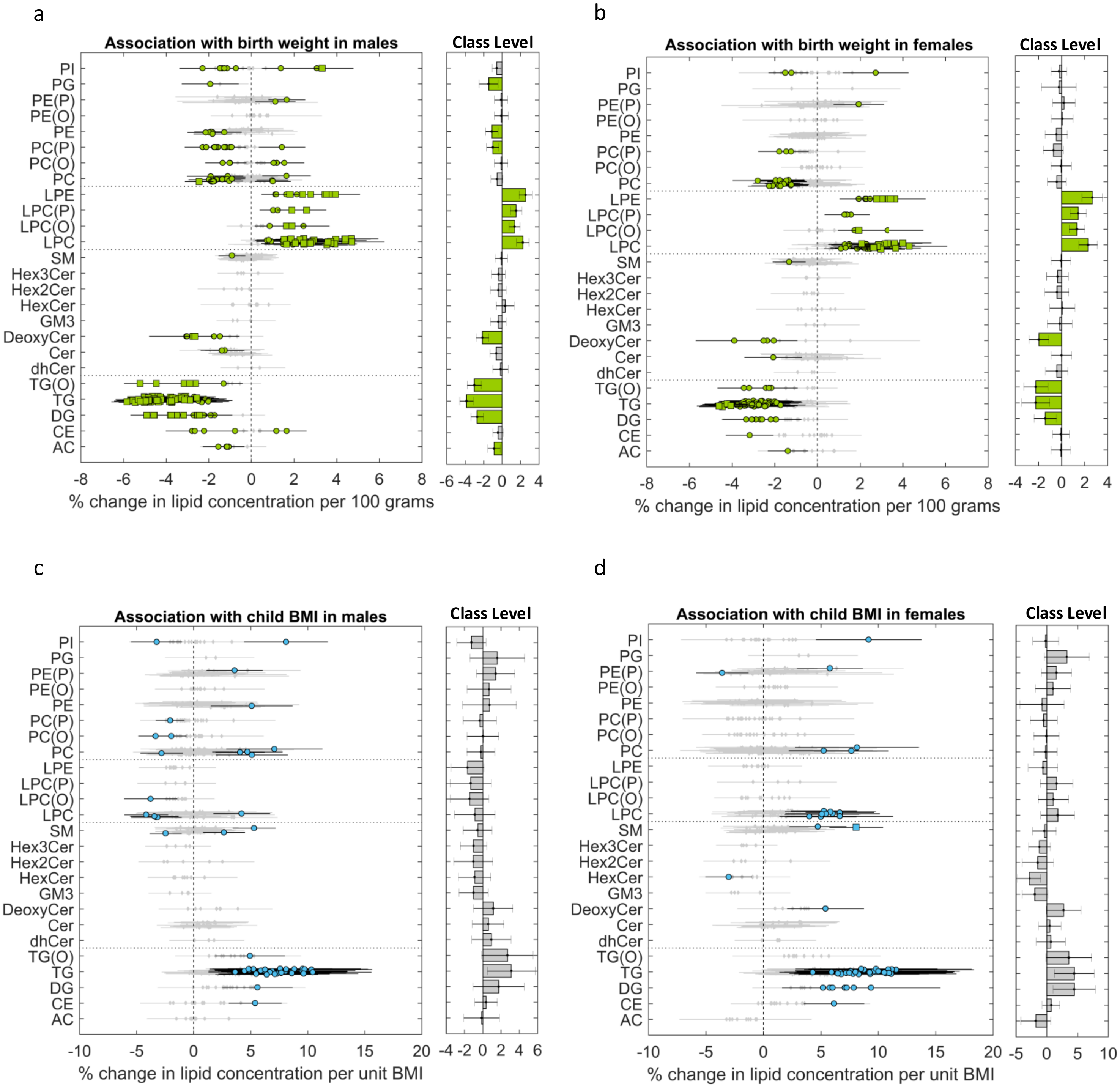

**Supplementary Figure 4.** Sex difference in the birth weight and child BMI studies: **(a)** Association with birth weight in males. **(b)** Association with birth weight in females. **(c)** Association with child BMI in males. **(d)** Association with child BMI in females. diamond – $P_{adj} \geq 0.05$ (grey), circle – $P_{adj}$ <0.05 and square – $P_{adj}$ <1.00E-5. Related to Supplementary Table 5.

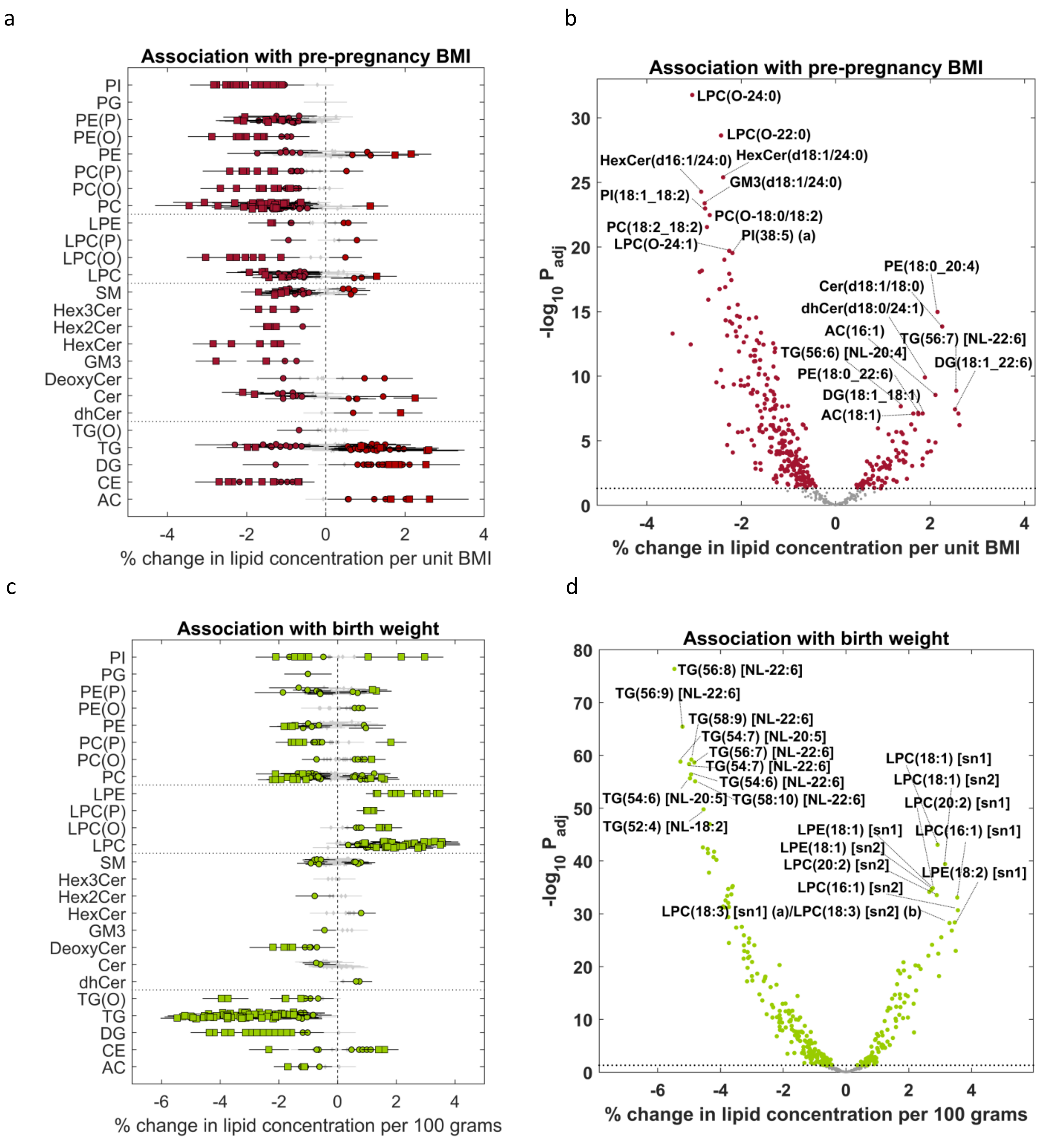

**Supplementary Figure 5.** Forest plots and volcano plots of the meta-analysis results in GUSTO and BIS for ppBMI (**a and b**) and BW studies (**c and d**). diamond – $P_{adj} \geq 0.05$ (grey), circle – $P_{adj} < 0.05$ and square – $P_{adj} < 1.00E\text{-}5$ in forest plots (a and c). Top 10 significant lipid species with positive and negative association are labelled in volcano plots (b and d, dotted line – $P_{adj}=0.05$). Related to Supplementary Table 6.

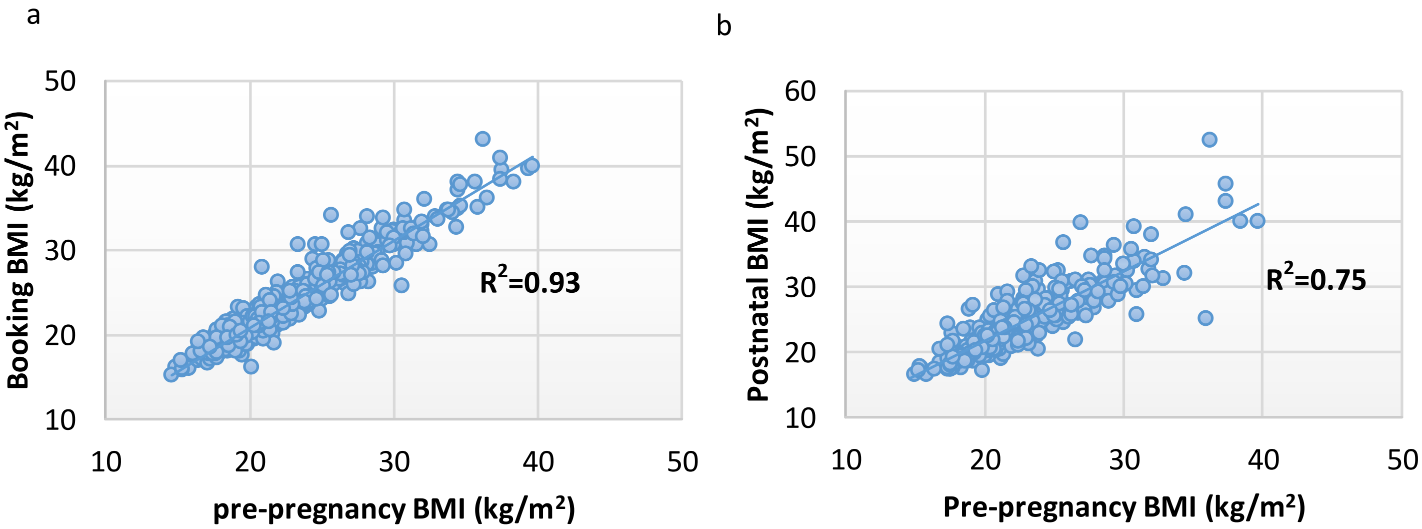

**Supplementary Figure 6.** Pre-pregnancy and postnatal maternal body mass index (BMI): (**a**) Pre-pregnancy BMI vs booking BMI measured at the first clinical visit. (**b**) Pre-pregnancy BMI vs postnatal BMI (4-5 years after delivery).

**Supplementary Table 1.** Clinical characteristics for participants in this study

| **Antenatal Mothers** (26-28 weeks' pregnancy) n=752 | | | **Postnatal Mothers** (4-5 years after delivery) n=650 | | |
|---|---|---|---|---|---|
| Variable | n | Mean (±STD) | Variable | n | Mean (±STD) |
| Ethnicity | | | Ethnicity | | |
| Chinese | 400 | - | Chinese | 375 | - |
| Malay | 198 | - | Malay | 160 | - |
| Indian | 154 | - | Indian | 115 | - |
| Maternal Education Level | | | Maternal Education Level | | |
| Secondary and below | 217 | - | Secondary and below | 194 | - |
| Post-secondary | 259 | - | Post-secondary | 223 | - |
| University | 270 | - | University | 228 | - |
| Maternal Age (years) | 752 | 30.68(±5.11) | Maternal Age (years) | 585 | 35.68(±5.15) |
| Pre-pregnancy BMI ($kg/m^2$) | 699 | 22.84(±4.33) | Pre-pregnancy BMI ($kg/m^2$) | 596 | 22.65(±4.34) |
| Gestational Weight Gain (kg) | 700 | 8.62(±4.31) | BMI ($kg/m^2$) | 541 | 24.55(±5.23) |

| **Newborns** n=751 | | | **Children** (Six-year-old) n=338 | | |
|---|---|---|---|---|---|
| Variable | n | Mean (±STD) | Variable | n | Mean (±STD) |
| Ethnicity | | | Ethnicity | | |
| Chinese | 368 | - | Chinese | 195 | - |
| Malay | 225 | - | Malay | 89 | - |
| Indian | 158 | - | Indian | 54 | - |
| Sex | | | Sex | | |
| Male | 401 | - | Male | 185 | - |
| Female | 350 | - | Female | 153 | - |
| Maternal Education Level | | | Maternal Education Level | | |
| Secondary and below | 251 | - | Secondary and below | 100 | - |
| Post-secondary | 284 | - | Post-secondary | 120 | - |
| University | 205 | - | University | 116 | - |
| Gestational Age (weeks) | 751 | 38.74(±1.37) | Gestational Age (weeks) | 338 | 38.79(±1.63) |
| Birth Weight (kg) | 751 | 3.10(±0.44) | Birth Weight (kg) | 338 | 3.10(±0.46) |
| Maternal Age (years) | 751 | 30.20(±5.20) | Child Age (years) | 330 | 6.06(±0.09) |
| Pre-pregnancy BMI ($kg/m^2$) | 679 | 22.97(±4.61) | Child BMI ($kg/m^2$) | 330 | 15.62(±2.33) |
| Total Gestational Weight Gain (kg) | 658 | 13.50(±5.76) | | | |
| Parity | | | | | |
| Primiparous | 330 | - | | | |
| Multiparous | 421 | - | | | |

**Supplementary Table 2a.** Comparison of baseline lipid classes across four sample groups (antenatal, postnatal, cord blood and 6-year-old child) using linear regression model. Related to Figure 1.

| | **From antenatal to postnatal** | | | | **From antenatal to cord blood** | | | | **From birth to child** | | | | **From child to adult (postnatal)** | | | |
|---|---|---|---|---|---|---|---|---|---|---|---|---|---|---|---|---|
| | adjusted for ethnicity, maternal age, maternal education level and pre-pregnancy BMI | | | | adjusted for ethnicity, maternal age, maternal education level, pre-pregnancy BMI, sex, gestational age and birth weight | | | | adjusted for sex, ethnicity, maternal education level, gestational age and birth weight | | | | adjusted for sex, ethnicity, BMI and maternal education level | | | |
| Lipid Class | $log_2FC$ (Postnatal/Antenatal) | 95%CI | p-value | p-value(BH) | $log_2FC$ (CordBlood/Antenatal) | 95%CI | p-value | p-value(BH) | $log_2FC$ (Child/CordBlood) | 95%CI | p-value | p-value(BH) | $log_2FC$ (Adult/Child) | 95%CI | p-value | p-value(BH) |
| AC | 1.23 | (1.19-1.28) | **2.31E-293** | **1.92E-292** | 0.40 | (0.34-0.46) | **9.89E-38** | **1.08E-37** | 0.70 | (0.64-0.76) | **1.82E-107** | **2.68E-107** | 0.04 | (-0.06-0.14) | 4.23E-01 | 5.28E-01 |
| CE | -0.05 | (-0.07--0.02) | **1.05E-03** | **1.19E-03** | -1.56 | (-1.6--1.52) | **0.00E+00** | **0.00E+00** | 1.34 | (1.3-1.38) | **0.00E+00** | **0.00E+00** | 0.24 | (0.18-0.3) | **5.16E-15** | **2.15E-14** |
| DG | -1.01 | (-1.08--0.95) | **1.67E-161** | **4.18E-161** | -3.04 | (-3.11--2.96) | **0.00E+00** | **0.00E+00** | 1.60 | (1.53-1.68) | **1.74E-236** | **3.62E-236** | -0.05 | (-0.19-0.1) | 5.50E-01 | 6.54E-01 |
| TG | -1.00 | (-1.06--0.95) | **5.58E-211** | **1.99E-210** | -2.87 | (-2.95--2.79) | **0.00E+00** | **0.00E+00** | 1.32 | (1.23-1.41) | **8.51E-134** | **1.42E-133** | 0.03 | (-0.11-0.16) | 6.98E-01 | 7.58E-01 |
| TG(O) | -0.97 | (-1.03--0.91) | **8.38E-160** | **1.91E-159** | -2.95 | (-3.03--2.87) | **0.00E+00** | **0.00E+00** | 1.55 | (1.47-1.63) | **1.27E-191** | **2.26E-191** | -0.09 | (-0.24-0.06) | 2.60E-01 | 3.42E-01 |
| dhCer | -0.12 | (-0.17--0.07) | **7.00E-06** | **8.75E-06** | 0.22 | ( 0.16- 0.28) | **8.89E-13** | **8.89E-13** | -0.48 | (-0.54--0.42) | **1.01E-50** | **1.27E-50** | -0.02 | (-0.13-0.09) | 7.40E-01 | 7.71E-01 |
| Cer | 0.18 | ( 0.14- 0.22) | **4.54E-19** | **6.67E-19** | -2.14 | (-2.19--2.09) | **0.00E+00** | **0.00E+00** | 2.10 | ( 2.05- 2.15) | **0.00E+00** | **0.00E+00** | 0.17 | ( 0.09-0.26) | **7.48E-05** | **1.25E-04** |
| DeoxyCer | -0.61 | (-0.67--0.55) | **1.63E-69** | **3.39E-69** | -1.96 | (-2.03--1.88) | **0.00E+00** | **0.00E+00** | 0.87 | (0.8-0.93) | **2.58E-118** | **4.03E-118** | 0.13 | (0-0.25) | 5.46E-02 | 7.58E-02 |
| GM3 | 0.20 | (0.16-0.25) | **7.95E-18** | **1.10E-17** | -0.25 | (-0.31--0.19) | **1.61E-15** | **1.67E-15** | 0.37 | (0.31-0.43) | **3.97E-32** | **4.51E-32** | 0.24 | (0.15-0.33) | **2.64E-07** | **5.49E-07** |
| HexCer | 0.12 | (0.08-0.16) | **1.31E-08** | **1.72E-08** | -1.30 | (-1.36--1.24) | **1.24E-247** | **1.83E-247** | 1.35 | (1.29-1.41) | **2.88E-238** | **6.54E-238** | 0.23 | (0.14-0.32) | **5.09E-07** | **9.78E-07** |
| Hex2Cer | -0.07 | (-0.12--0.02) | **3.61E-03** | **3.92E-03** | -2.17 | (-2.23--2.11) | **0.00E+00** | **0.00E+00** | 1.76 | (1.69-1.82) | **4.98E-304** | **1.38E-303** | 0.51 | (0.41-0.62) | **1.79E-20** | **2.24E-19** |
| Hex3Cer | 0.19 | (0.15-0.22) | **2.80E-22** | **4.38E-22** | -0.85 | (-0.9--0.8) | **1.89E-171** | **2.62E-171** | 1.06 | (1.01-1.11) | **3.87E-212** | **7.44E-212** | 0.11 | (0.03-0.19) | **7.60E-03** | **1.12E-02** |
| SM | -0.32 | (-0.36--0.28) | **1.64E-51** | **3.16E-51** | -1.85 | (-1.9--1.8) | **0.00E+00** | **0.00E+00** | 1.29 | (1.24-1.34) | **2.07E-298** | **5.17E-298** | 0.30 | (0.22-0.39) | **6.36E-12** | **1.77E-11** |
| LPC | 0.90 | (0.86-0.95) | **7.17E-246** | **2.99E-245** | 0.40 | (0.36-0.45) | **7.90E-60** | **8.98E-60** | 0.25 | (0.2-0.3) | **3.53E-21** | **3.67E-21** | 0.49 | (0.38-0.59) | **5.78E-20** | **3.61E-19** |
| LPC(O) | 0.87 | (0.83-0.9) | **8.75E-280** | **5.47E-279** | 0.37 | (0.33-0.41) | **1.82E-66** | **2.17E-66** | 0.25 | (0.2-0.29) | **7.27E-25** | **7.91E-25** | 0.46 | (0.37-0.56) | **8.65E-22** | **2.16E-20** |
| LPC(P) | 1.14 | (1.1-1.18) | **0.00E+00** | **0.00E+00** | 0.68 | (0.63-0.72) | **1.26E-163** | **1.66E-163** | 0.17 | (0.13-0.22) | **3.49E-12** | **3.49E-12** | 0.47 | (0.38-0.57) | **3.10E-20** | **2.59E-19** |
| LPE | 0.31 | (0.27-0.36) | **1.57E-46** | **2.80E-46** | 0.49 | (0.45-0.54) | **4.94E-88** | **6.17E-88** | -0.38 | (-0.43--0.33) | **1.79E-49** | **2.14E-49** | 0.44 | (0.34-0.54) | **2.79E-18** | **1.39E-17** |
| PC | -0.63 | (-0.67--0.6) | **5.16E-194** | **1.61E-193** | -2.35 | (-2.39--2.3) | **0.00E+00** | **0.00E+00** | 1.56 | (1.51-1.6) | **0.00E+00** | **0.00E+00** | 0.21 | (0.13-0.29) | **6.15E-07** | **1.10E-06** |
| PC(O) | -0.22 | (-0.27--0.18) | **4.62E-24** | **7.70E-24** | -1.85 | (-1.91--1.8) | **0.00E+00** | **0.00E+00** | 1.46 | (1.41-1.51) | **0.00E+00** | **0.00E+00** | 0.33 | (0.24-0.42) | **4.38E-12** | **1.37E-11** |
| PC(P) | -0.08 | (-0.13--0.04) | **5.40E-04** | **6.43E-04** | -1.76 | (-1.81--1.7) | **0.00E+00** | **0.00E+00** | 1.50 | (1.44-1.55) | **6.78E-307** | **2.12E-306** | 0.37 | (0.27-0.46) | **2.66E-13** | **9.49E-13** |
| PE | -2.36 | (-2.42--2.29) | **0.00E+00** | **0.00E+00** | -3.70 | (-3.76--3.64) | **0.00E+00** | **0.00E+00** | 0.82 | (0.75-0.88) | **4.98E-100** | **6.55E-100** | 0.48 | (0.32-0.63) | **3.67E-09** | **9.17E-09** |
| PE(O) | 0.02 | (-0.04-0.07) | 5.93E-01 | 5.93E-01 | -2.94 | (-3--2.87) | **0.00E+00** | **0.00E+00** | 2.77 | (2.71-2.83) | **0.00E+00** | **0.00E+00** | 0.35 | (0.23-0.48) | **7.57E-08** | **1.72E-07** |
| PE(P) | -0.06 | (-0.11--0.01) | **2.80E-02** | **2.92E-02** | -2.70 | (-2.76--2.64) | **0.00E+00** | **0.00E+00** | 2.73 | (2.67-2.78) | **0.00E+00** | **0.00E+00** | 0.01 | (-0.1-0.12) | 8.06E-01 | 8.06E-01 |
| PG | -1.04 | (-1.1--0.98) | **1.19E-173** | **3.31E-173** | -2.50 | (-2.58--2.42) | **0.00E+00** | **0.00E+00** | 1.11 | (1.02-1.2) | **5.87E-105** | **8.15E-105** | 0.04 | (-0.11-0.2) | 5.88E-01 | 6.68E-01 |
| PI | -1.04 | (-1.08--1) | **8.26E-279** | **4.13E-278** | -2.36 | (-2.4--2.31) | **0.00E+00** | **0.00E+00** | 1.18 | (1.14-1.22) | **0.00E+00** | **0.00E+00** | 0.15 | (0.06-0.25) | **1.21E-03** | **1.89E-03** |

**Supplementary Table 2b.** Comparison of baseline lipids species across four sample groups (antenatal, postnatal, cord blood and 6-year-old child) using linear regression model. Related to Figure 1.

| Studies | | From antenatal to postnatal | | | | From antenatal to cord blood | | | | From birth to child | | | | From child to adult (postnatal) | | | |
|---|---|---|---|---|---|---|---|---|---|---|---|---|---|---|---|---|---|
| covariates | | adjusted for ethnicity, maternal age, maternal education level and pre-pregnancy BMI | | | | adjusted for ethnicity, maternal age, maternal education level, pre-pregnancy BMI, sex, gestational age and birth weight | | | | adjusted for sex, ethnicity, maternal education level, gestational age and birth weight | | | | adjusted for sex, ethnicity, BMI and maternal education level | | | |
| No. | Lipid Species | $log_2FC$ (Postnatal/Antenatal) | 95%CI | p-value | p-value(BH) | $log_2FC$ (CordBlood/Antenatal) | 95%CI | p-value | p-value(BH) | $log_2FC$ (Child/CordBlood) | 95%CI | p-value | p-value(BH) | $log_2FC$ (Adult/Child) | 95%CI | p-value | p-value(BH) |
| 1 | AC(12:0) | 1.64 | (1.56-1.72) | 4.90E-214 | 3.02E-213 | -0.04 | (-0.13-0.04) | 3.28E-01 | 3.30E-01 | 1.80 | (1.72-1.87) | 7.53E-254 | 2.56E-253 | -0.22 | (-0.38--0.06) | 7.68E-03 | 1.22E-02 |
| 2 | AC(13:0) | 1.21 | (1.09-1.32) | 1.72E-81 | 3.69E-81 | 0.47 | (0.34-0.6) | 1.49E-12 | 1.60E-12 | -0.19 | (-0.31--0.06) | 3.21E-03 | 3.34E-03 | 0.80 | (0.56-1.05) | 1.98E-10 | 8.54E-10 |
| 3 | AC(14:0) | 1.10 | (1.05-1.16) | 2.16E-211 | 1.29E-210 | 0.84 | (0.77-0.9) | 8.03E-113 | 9.99E-113 | 0.27 | (0.2-0.33) | 1.04E-15 | 1.18E-15 | -0.15 | (-0.27--0.03) | 1.12E-02 | 1.74E-02 |
| 4 | AC(14:1) | 1.90 | (1.81-1.99) | 4.36E-237 | 3.13E-236 | 0.23 | (0.14-0.33) | 2.81E-06 | 2.97E-06 | 1.79 | (1.7-1.87) | 1.03E-228 | 3.18E-228 | -0.33 | (-0.51--0.16) | 1.69E-04 | 3.40E-04 |
| 5 | AC(14:2) | 1.74 | (1.65-1.83) | 1.03E-225 | 6.84E-225 | 0.50 | (0.41-0.59) | 1.71E-26 | 1.88E-26 | 1.45 | (1.37-1.52) | 2.19E-206 | 5.98E-206 | -0.34 | (-0.51--0.18) | 3.93E-05 | 9.03E-05 |
| 6 | AC(16:0) | 0.92 | (0.88-0.96) | 9.06E-251 | 7.63E-250 | 0.67 | (0.61-0.72) | 3.29E-112 | 4.07E-112 | 0.04 | (-0.01-0.09) | 1.15E-01 | 1.17E-01 | 0.16 | (0.08-0.24) | 8.22E-05 | 1.75E-04 |
| 7 | AC(16:1) | 1.36 | (1.29-1.42) | 3.38E-236 | 2.38E-235 | 0.98 | (0.9-1.05) | 1.05E-112 | 1.30E-112 | 0.13 | (0.06-0.21) | 2.88E-04 | 3.02E-04 | -0.02 | (-0.15-0.1) | 7.14E-01 | 7.54E-01 |
| 8 | AC(18:0) | 1.02 | (0.98-1.06) | 1.26E-271 | 1.35E-270 | 0.34 | (0.28-0.39) | 1.98E-33 | 2.19E-33 | 0.61 | (0.56-0.66) | 2.16E-93 | 3.51E-93 | 0.11 | (0.02-0.2) | 1.53E-02 | 2.28E-02 |
| 9 | AC(18:1) | 1.19 | (1.13-1.24) | 8.43E-255 | 7.23E-254 | -0.08 | (-0.15--0.01) | 2.03E-02 | 2.08E-02 | 1.02 | (0.95-1.08) | 1.82E-159 | 3.77E-159 | 0.17 | (0.07-0.27) | 5.91E-04 | 1.09E-03 |
| 10 | AC(18:2) | 1.12 | (1.07-1.17) | 3.00E-243 | 2.29E-242 | 0.73 | (0.67-0.8) | 1.26E-91 | 1.50E-91 | 0.11 | (0.06-0.17) | 1.21E-04 | 1.28E-04 | 0.20 | (0.1-0.29) | 4.58E-05 | 1.03E-04 |
| 11 | CE(14:0) | -0.49 | (-0.54--0.43) | 1.14E-68 | 2.28E-68 | -0.46 | (-0.51--0.4) | 7.57E-53 | 8.71E-53 | 0.02 | (-0.04-0.07) | 5.52E-01 | 5.55E-01 | -0.12 | (-0.23-0) | 4.39E-02 | 6.00E-02 |
| 12 | CE(16:1) | -0.79 | (-0.86--0.73) | 1.50E-106 | 3.99E-106 | 0.24 | (0.16-0.31) | 1.99E-10 | 2.13E-10 | -1.26 | (-1.32--1.2) | 2.07E-216 | 5.95E-216 | -0.06 | (-0.18-0.06) | 3.02E-01 | 3.57E-01 |
| 13 | CE(18:0) | 0.15 | (0.1-0.2) | 1.68E-10 | 2.12E-10 | 0.65 | (0.59-0.71) | 1.30E-77 | 1.51E-77 | -0.57 | (-0.64--0.5) | 2.27E-54 | 3.13E-54 | 0.16 | (0.05-0.27) | 4.11E-03 | 6.74E-03 |
| 14 | CE(18:1) | -0.29 | (-0.32--0.26) | 1.34E-86 | 3.00E-86 | -0.85 | (-0.89--0.81) | 6.17E-266 | 1.06E-265 | 0.51 | (0.47-0.55) | 6.21E-122 | 1.13E-121 | 0.08 | (0.03-0.14) | 3.91E-03 | 6.43E-03 |
| 15 | CE(18:2) | -0.12 | (-0.15--0.09) | 4.21E-15 | 5.63E-15 | -2.69 | (-2.73--2.65) | 0.00E+00 | 0.00E+00 | 2.41 | (2.36-2.45) | 0.00E+00 | 0.00E+00 | 0.22 | (0.15-0.28) | 5.10E-11 | 2.42E-10 |
| 16 | CE(18:3) | 0.11 | (0.06-0.17) | 5.13E-05 | 5.99E-05 | -2.13 | (-2.21--2.06) | 0.00E+00 | 0.00E+00 | 2.21 | (2.13-2.29) | 0.00E+00 | 0.00E+00 | 0.03 | (-0.1-0.15) | 6.70E-01 | 7.18E-01 |
| 17 | CE(20:1) | 0.00 | (-0.04-0.04) | 8.75E-01 | 8.79E-01 | 0.14 | (0.07-0.21) | 3.37E-05 | 3.53E-05 | -0.19 | (-0.26--0.12) | 8.39E-08 | 9.05E-08 | 0.23 | (0.14-0.32) | 1.00E-06 | 2.87E-06 |
| 18 | CE(20:2) | -0.48 | (-0.52--0.45) | 6.19E-130 | 1.92E-129 | -0.46 | (-0.51--0.41) | 4.52E-62 | 5.24E-62 | -0.13 | (-0.18--0.08) | 1.11E-06 | 1.18E-06 | 0.18 | (0.1-0.25) | 2.59E-06 | 7.02E-06 |
| 19 | CE(20:4) | 0.56 | (0.5-0.61) | 4.79E-84 | 1.05E-83 | 0.39 | (0.33-0.45) | 3.18E-34 | 3.54E-34 | 0.02 | (-0.04-0.08) | 4.73E-01 | 4.77E-01 | 0.32 | (0.21-0.42) | 2.74E-09 | 1.03E-08 |
| 20 | CE(20:5) | 1.14 | (1.02-1.25) | 3.01E-73 | 6.20E-73 | -1.44 | (-1.58--1.31) | 1.78E-86 | 2.09E-86 | 1.97 | (1.85-2.1) | 2.64E-148 | 5.23E-148 | 0.73 | (0.49-0.97) | 3.03E-09 | 1.12E-08 |
| 21 | CE(22:5) (n6) | -0.23 | (-0.28--0.19) | 9.65E-22 | 1.36E-21 | -0.44 | (-0.5--0.37) | 1.61E-35 | 1.80E-35 | 0.32 | (0.25-0.38) | 1.48E-21 | 1.76E-21 | -0.04 | (-0.13-0.05) | 4.18E-01 | 4.76E-01 |
| 22 | CE(22:6) | -0.33 | (-0.4--0.27) | 3.09E-23 | 4.41E-23 | -1.84 | (-1.93--1.76) | 7.97E-252 | 1.29E-251 | 0.97 | (0.88-1.06) | 2.31E-84 | 3.62E-84 | 0.53 | (0.38-0.67) | 2.09E-12 | 1.18E-11 |
| 23 | CE(24:5) | -1.03 | (-1.09--0.97) | 8.76E-169 | 3.86E-168 | 1.21 | (1.11-1.3) | 4.78E-111 | 5.88E-111 | -2.10 | (-2.2--2) | 1.29E-226 | 3.90E-226 | 0.01 | (-0.12-0.14) | 9.12E-01 | 9.27E-01 |
| 24 | CE(24:6) | -0.73 | (-0.79--0.66) | 4.32E-91 | 9.83E-91 | -0.34 | (-0.43--0.25) | 6.45E-13 | 6.97E-13 | -0.26 | (-0.35--0.16) | 4.64E-08 | 5.02E-08 | -0.03 | (-0.16-0.11) | 7.11E-01 | 7.54E-01 |
| 25 | DG(16:0_16:1) | -1.60 | (-1.69--1.51) | 3.19E-195 | 1.74E-194 | -2.14 | (-2.24--2.04) | 7.24E-248 | 1.16E-247 | 0.21 | (0.12-0.29) | 6.67E-06 | 7.09E-06 | -0.26 | (-0.43--0.08) | 4.83E-03 | 7.84E-03 |
| 26 | DG(14:0_18:2) | -1.65 | (-1.76--1.55) | 1.97E-159 | 7.76E-159 | -3.87 | (-4--3.75) | 0.00E+00 | 0.00E+00 | 1.73 | (1.59-1.86) | 2.18E-110 | 3.76E-110 | -0.20 | (-0.45-0.04) | 1.07E-01 | 1.38E-01 |
| 27 | DG(16:0_18:1) | -1.60 | (-1.67--1.52) | 5.56E-247 | 4.52E-246 | -3.28 | (-3.36--3.2) | 0.00E+00 | 0.00E+00 | 1.38 | (1.3-1.46) | 2.98E-160 | 6.24E-160 | -0.24 | (-0.42--0.07) | 5.59E-03 | 9.04E-03 |
| 28 | DG(16:1_18:1) | -1.11 | (-1.19--1.03) | 6.92E-126 | 2.08E-125 | -2.44 | (-2.53--2.35) | 0.00E+00 | 0.00E+00 | 0.79 | (0.7-0.88) | 1.86E-58 | 2.64E-58 | -0.09 | (-0.27-0.1) | 3.62E-01 | 4.20E-01 |
| 29 | DG(16:0_18:2) | -1.47 | (-1.54--1.4) | 1.78E-231 | 1.22E-230 | -3.72 | (-3.81--3.62) | 0.00E+00 | 0.00E+00 | 1.73 | (1.64-1.83) | 7.74E-183 | 1.83E-182 | -0.02 | (-0.18-0.15) | 8.50E-01 | 8.81E-01 |
| 30 | DG(18:0_18:1) | -0.97 | (-1.04--0.9) | 4.89E-130 | 1.53E-129 | -2.58 | (-2.66--2.51) | 0.00E+00 | 0.00E+00 | 1.36 | (1.28-1.44) | 3.99E-167 | 8.63E-167 | -0.30 | (-0.47--0.14) | 3.08E-04 | 5.91E-04 |
| 31 | DG(18:1_18:1) | -0.88 | (-0.94--0.81) | 2.33E-111 | 6.35E-111 | -3.29 | (-3.37--3.21) | 0.00E+00 | 0.00E+00 | 2.06 | (1.98-2.14) | 1.46E-289 | 5.99E-289 | -0.10 | (-0.25-0.06) | 2.36E-01 | 2.84E-01 |
| 32 | DG(18:0_18:2) | -1.06 | (-1.14--0.98) | 7.58E-129 | 2.32E-128 | -2.90 | (-2.99--2.8) | 0.00E+00 | 0.00E+00 | 1.49 | (1.39-1.59) | 1.67E-137 | 3.20E-137 | -0.22 | (-0.4--0.04) | 1.69E-02 | 2.49E-02 |
| 33 | DG(18:1_18:2) | -0.78 | (-0.85--0.71) | 6.54E-95 | 1.57E-94 | -3.81 | (-3.9--3.73) | 0.00E+00 | 0.00E+00 | 2.47 | (2.38-2.56) | 0.00E+00 | 0.00E+00 | 0.13 | (-0.03-0.28) | 1.07E-01 | 1.38E-01 |
| 34 | DG(18:2_18:2) | -0.65 | (-0.73--0.56) | 2.07E-48 | 3.63E-48 | -4.12 | (-4.23--4) | 0.00E+00 | 0.00E+00 | 2.70 | (2.59-2.82) | 9.97E-267 | 3.71E-266 | 0.34 | (0.16-0.52) | 2.80E-04 | 5.43E-04 |
| 35 | DG(18:1_18:3) | -0.47 | (-0.54--0.39) | 4.12E-32 | 6.32E-32 | -3.27 | (-3.37--3.18) | 0.00E+00 | 0.00E+00 | 2.30 | (2.2-2.4) | 1.50E-252 | 5.07E-252 | 0.08 | (-0.1-0.26) | 3.97E-01 | 4.55E-01 |
| 36 | DG(16:0_20:4) | -1.55 | (-1.63--1.46) | 4.30E-193 | 2.27E-192 | -1.43 | (-1.53--1.33) | 2.29E-142 | 3.00E-142 | -0.48 | (-0.57--0.38) | 7.21E-23 | 8.63E-23 | -0.13 | (-0.31-0.06) | 1.83E-01 | 2.24E-01 |
| 37 | DG(18:1_20:3) | -0.89 | (-0.96--0.82) | 4.14E-111 | 1.12E-110 | -1.13 | (-1.21--1.05) | 1.04E-136 | 1.36E-136 | 0.25 | (0.17-0.33) | 2.13E-09 | 2.33E-09 | -0.31 | (-0.47--0.15) | 1.60E-04 | 3.25E-04 |
| 38 | DG(18:1_20:4) | -0.70 | (-0.76--0.63) | 3.18E-83 | 6.87E-83 | -1.60 | (-1.68--1.52) | 6.80E-225 | 1.05E-224 | 0.50 | (0.42-0.58) | 1.28E-32 | 1.61E-32 | 0.06 | (-0.08-0.21) | 3.85E-01 | 4.44E-01 |
| 39 | DG(16:0_22:5) | -1.33 | (-1.41--1.25) | 1.70E-167 | 7.27E-167 | -1.07 | (-1.16--0.97) | 3.88E-91 | 4.61E-91 | -0.61 | (-0.7--0.52) | 2.28E-35 | 2.93E-35 | -0.01 | (-0.19-0.16) | 8.82E-01 | 9.06E-01 |
| 40 | DG(18:2_20:4) | -0.70 | (-0.77--0.62) | 1.01E-64 | 1.95E-64 | -1.94 | (-2.04--1.84) | 3.73E-213 | 5.59E-213 | 0.65 | (0.55-0.75) | 1.47E-34 | 1.88E-34 | 0.21 | (0.04-0.37) | 1.44E-02 | 2.17E-02 |
| 41 | DG(16:0_22:6) | -1.65 | (-1.75--1.54) | 6.60E-154 | 2.48E-153 | -1.37 | (-1.49--1.25) | 3.10E-97 | 3.73E-97 | -0.97 | (-1.08--0.85) | 3.97E-53 | 5.45E-53 | 0.06 | (-0.17-0.3) | 6.09E-01 | 6.67E-01 |
| 42 | DG(18:1_22:5) | -0.86 | (-0.93--0.78) | 1.51E-97 | 3.65E-97 | -1.00 | (-1.09--0.91) | 2.39E-89 | 2.83E-89 | -0.11 | (-0.2--0.02) | 1.85E-02 | 1.90E-02 | -0.04 | (-0.2-0.12) | 6.09E-01 | 6.67E-01 |
| 43 | DG(18:1_22:6) | -0.86 | (-0.96--0.76) | 5.80E-61 | 1.10E-60 | -1.29 | (-1.4--1.19) | 1.26E-104 | 1.53E-104 | -0.31 | (-0.41--0.21) | 2.05E-09 | 2.24E-09 | 0.26 | (0.05-0.47) | 1.37E-02 | 2.06E-02 |
| 44 | DG(18:2_22:6) | -0.85 | (-0.95--0.75) | 4.19E-57 | 7.74E-57 | -1.65 | (-1.77--1.52) | 2.59E-126 | 3.31E-126 | -0.20 | (-0.32--0.07) | 1.72E-03 | 1.80E-03 | 0.50 | (0.28-0.73) | 9.89E-06 | 2.40E-05 |
| 45 | TG(48:0) [NL-18:0] | -1.16 | (-1.26--1.05) | 4.85E-94 | 1.14E-93 | -1.72 | (-1.82--1.63) | 6.42E-192 | 9.26E-192 | 0.49 | (0.38-0.6) | 6.90E-18 | 8.02E-18 | -0.38 | (-0.63--0.13) | 3.09E-03 | 5.17E-03 |
| 46 | TG(48:1) [NL-16:1] | -2.28 | (-2.38--2.17) | 3.16E-238 | 2.30E-237 | -2.74 | (-2.86--2.62) | 9.15E-264 | 1.54E-263 | -0.12 | (-0.24-0) | 4.57E-02 | 4.68E-02 | -0.31 | (-0.55--0.07) | 1.08E-02 | 1.69E-02 |
| 47 | TG(48:1) [NL-18:1] | -1.89 | (-2--1.79) | 1.09E-189 | 5.70E-189 | -3.74 | (-3.85--3.63) | 0.00E+00 | 0.00E+00 | 1.36 | (1.23-1.49) | 2.02E-82 | 3.13E-82 | -0.22 | (-0.49-0.05) | 1.09E-01 | 1.40E-01 |
| 48 | TG(48:2) [NL-14:0] | -2.11 | (-2.22--1.99) | 6.65E-202 | 3.80E-201 | -3.96 | (-4.08--3.84) | 0.00E+00 | 0.00E+00 | 1.09 | (0.96-1.22) | 7.45E-57 | 1.04E-56 | -0.02 | (-0.29-0.26) | 9.01E-01 | 9.20E-01 |

| | | | | | | | | | | | | | | | | | |
|---|---|---|---|---|---|---|---|---|---|---|---|---|---|---|---|---|---|
| 49 | TG(48:2) [NL-14:1] | -1.79 | (-1.89--1.68) | **3.03E-167** | **1.29E-166** | -3.76 | (-3.87--3.65) | **0.00E+00** | **0.00E+00** | 1.55 | (1.43-1.67) | **5.38E-112** | **9.38E-112** | -0.29 | (-0.55--0.03) | **2.69E-02** | **3.87E-02** |
| 50 | TG(48:2) [NL-16:1] | -1.97 | (-2.09--1.85) | **4.27E-169** | **1.90E-168** | -2.64 | (-2.77--2.51) | **3.86E-232** | **5.99E-232** | -0.18 | (-0.3--0.06) | **3.87E-03** | **4.02E-03** | -0.03 | (-0.29-0.23) | 8.46E-01 | 8.78E-01 |
| 51 | TG(48:2) [NL-18:2] | -2.10 | (-2.21--1.98) | **8.74E-195** | **4.66E-194** | -4.38 | (-4.5--4.25) | **0.00E+00** | **0.00E+00** | 1.53 | (1.39-1.66) | **8.23E-92** | **1.32E-91** | -0.02 | (-0.3-0.26) | 8.85E-01 | 9.08E-01 |
| 52 | TG(48:3) [NL-14:0] | -1.70 | (-1.81--1.58) | **2.52E-143** | **8.65E-143** | -3.45 | (-3.57--3.32) | **0.00E+00** | **0.00E+00** | 0.88 | (0.75-1.01) | **5.32E-38** | **6.92E-38** | 0.13 | (-0.14-0.4) | 3.52E-01 | 4.10E-01 |
| 53 | TG(48:3) [NL-16:1] | -1.54 | (-1.64--1.43) | **9.23E-139** | **3.03E-138** | -2.38 | (-2.49--2.26) | **8.38E-238** | **1.33E-237** | 0.02 | (-0.09-0.12) | 7.76E-01 | 7.78E-01 | 0.12 | (-0.11-0.35) | 3.21E-01 | 3.76E-01 |
| 54 | TG(48:3) [NL-18:3] | -1.74 | (-1.87--1.61) | **4.90E-126** | **1.48E-125** | -3.44 | (-3.57--3.32) | **0.00E+00** | **0.00E+00** | 1.15 | (1.03-1.27) | **8.24E-67** | **1.21E-66** | -0.07 | (-0.36-0.23) | 6.60E-01 | 7.12E-01 |
| 55 | TG(49:1) [NL-17:1] | -1.70 | (-1.78--1.62) | **3.15E-246** | **2.52E-245** | -2.44 | (-2.53--2.35) | **0.00E+00** | **0.00E+00** | 0.34 | (0.26-0.42) | **2.24E-15** | **2.54E-15** | -0.18 | (-0.35--0.01) | **4.29E-02** | 5.90E-02 |
| 56 | TG(50:1) [NL-14:0] | -1.11 | (-1.2--1.02) | **3.27E-103** | **8.49E-103** | -1.95 | (-2.05--1.86) | **9.66E-233** | **1.51E-232** | 0.69 | (0.58-0.8) | **1.56E-32** | **1.96E-32** | -0.46 | (-0.69--0.23) | **9.70E-05** | **2.02E-04** |
| 57 | TG(50:1) [NL-16:0] | -1.61 | (-1.68--1.54) | **5.35E-266** | **5.35E-265** | -2.93 | (-3.01--2.84) | **0.00E+00** | **0.00E+00** | 0.96 | (0.86-1.06) | **5.68E-72** | **8.54E-72** | -0.20 | (-0.37--0.02) | **2.55E-02** | **3.68E-02** |
| 58 | TG(50:1) [NL-18:1] | -1.56 | (-1.63--1.49) | **3.39E-261** | **3.12E-260** | -2.90 | (-2.98--2.81) | **0.00E+00** | **0.00E+00** | 0.96 | (0.86-1.06) | **4.43E-73** | **6.73E-73** | -0.16 | (-0.33-0.01) | 6.96E-02 | 9.23E-02 |
| 59 | TG(50:2) [NL-14:0] | -0.96 | (-1.04--0.89) | **2.77E-114** | **7.82E-114** | -3.12 | (-3.22--3.03) | **0.00E+00** | **0.00E+00** | 1.70 | (1.59-1.82) | **3.03E-138** | **5.83E-138** | -0.06 | (-0.27-0.14) | 5.58E-01 | 6.20E-01 |
| 60 | TG(50:2) [NL-16:1] | -1.16 | (-1.23--1.08) | **5.65E-151** | **2.02E-150** | -2.43 | (-2.53--2.33) | **2.00E-286** | **3.61E-286** | 0.60 | (0.48-0.71) | **3.08E-24** | **3.73E-24** | -0.02 | (-0.21-0.16) | 8.02E-01 | 8.37E-01 |
| 61 | TG(50:2) [NL-18:1] | -1.03 | (-1.1--0.96) | **2.29E-139** | **7.63E-139** | -2.62 | (-2.72--2.52) | **0.00E+00** | **0.00E+00** | 0.95 | (0.83-1.06) | **4.65E-54** | **6.39E-54** | -0.01 | (-0.2-0.17) | 8.88E-01 | 9.09E-01 |
| 62 | TG(50:2) [NL-18:2] | -1.50 | (-1.56--1.43) | **3.11E-262** | **2.99E-261** | -3.51 | (-3.62--3.4) | **0.00E+00** | **0.00E+00** | 1.43 | (1.31-1.55) | **5.03E-100** | **8.26E-100** | 0.00 | (-0.17-0.16) | 9.74E-01 | 9.78E-01 |
| 63 | TG(50:3) [NL-14:0] | -1.30 | (-1.38--1.21) | **2.74E-144** | **9.46E-144** | -4.10 | (-4.21--3.99) | **0.00E+00** | **0.00E+00** | 2.00 | (1.87-2.12) | **1.06E-152** | **2.13E-152** | 0.17 | (-0.06-0.4) | 1.40E-01 | 1.75E-01 |
| 64 | TG(50:3) [NL-14:1] | -1.01 | (-1.08--0.94) | **2.14E-126** | **6.50E-126** | -3.12 | (-3.21--3.03) | **0.00E+00** | **0.00E+00** | 1.73 | (1.62-1.83) | **4.24E-168** | **9.21E-168** | -0.15 | (-0.33-0.04) | 1.24E-01 | 1.57E-01 |
| 65 | TG(50:3) [NL-16:1] | -1.38 | (-1.47--1.3) | **1.69E-160** | **6.69E-160** | -3.06 | (-3.17--2.96) | **0.00E+00** | **0.00E+00** | 0.63 | (0.52-0.75) | **1.06E-24** | **1.29E-24** | 0.24 | (0.03-0.45) | **2.82E-02** | **4.04E-02** |
| 66 | TG(50:3) [NL-18:2] | -1.28 | (-1.36--1.2) | **3.48E-157** | **1.34E-156** | -3.60 | (-3.71--3.49) | **0.00E+00** | **0.00E+00** | 1.30 | (1.18-1.42) | **3.94E-80** | **6.04E-80** | 0.26 | (0.05-0.47) | **1.37E-02** | **2.06E-02** |
| 67 | TG(50:3) [NL-18:3] | -1.71 | (-1.8--1.61) | **8.25E-186** | **4.08E-185** | -3.67 | (-3.78--3.56) | **0.00E+00** | **0.00E+00** | 1.25 | (1.13-1.37) | **1.73E-77** | **2.64E-77** | -0.01 | (-0.24-0.22) | 9.25E-01 | 9.39E-01 |
| 68 | TG(50:4) [NL-14:0] | -1.29 | (-1.39--1.18) | **1.02E-111** | **2.79E-111** | -3.88 | (-4--3.76) | **0.00E+00** | **0.00E+00** | 1.70 | (1.58-1.82) | **3.31E-125** | **6.15E-125** | 0.31 | (0.07-0.56) | **1.11E-02** | **1.72E-02** |
| 69 | TG(50:4) [NL-18:3] | -1.15 | (-1.25--1.04) | **8.31E-92** | **1.90E-91** | -2.94 | (-3.05--2.82) | **0.00E+00** | **0.00E+00** | 0.97 | (0.86-1.09) | **8.64E-57** | **1.20E-56** | 0.14 | (-0.09-0.37) | 2.43E-01 | 2.92E-01 |
| 70 | TG(50:4) [NL-20:4] | -2.14 | (-2.27--2) | **7.15E-163** | **2.94E-162** | -2.11 | (-2.25--1.96) | **4.42E-144** | **5.82E-144** | -0.67 | (-0.8--0.53) | **2.01E-21** | **2.36E-21** | -0.10 | (-0.39-0.19) | 5.03E-01 | 5.63E-01 |
| 71 | TG(51:1) [NL-17:0] | -1.46 | (-1.54--1.38) | **2.49E-198** | **1.37E-197** | -2.94 | (-3.03--2.84) | **0.00E+00** | **0.00E+00** | 1.14 | (1.04-1.23) | **4.56E-92** | **7.34E-92** | -0.34 | (-0.54--0.15) | **5.23E-04** | **9.77E-04** |
| 72 | TG(51:2) [NL-15:0] | -0.95 | (-1.02--0.88) | **1.33E-123** | **3.88E-123** | -3.63 | (-3.73--3.54) | **0.00E+00** | **0.00E+00** | 2.15 | (2.04-2.25) | **1.25E-208** | **3.42E-208** | 0.00 | (-0.18-0.18) | 9.73E-01 | 9.78E-01 |
| 73 | TG(51:2) [NL-17:0] | -1.35 | (-1.43--1.27) | **3.61E-187** | **1.81E-186** | -3.37 | (-3.46--3.27) | **0.00E+00** | **0.00E+00** | 1.36 | (1.25-1.46) | **4.38E-110** | **7.54E-110** | -0.04 | (-0.23-0.15) | 6.64E-01 | 7.13E-01 |
| 74 | TG(51:2) [NL-17:1] | -1.28 | (-1.35--1.2) | **1.75E-188** | **8.96E-188** | -3.00 | (-3.09--2.92) | **0.00E+00** | **0.00E+00** | 1.15 | (1.06-1.23) | **2.25E-109** | **3.83E-109** | -0.04 | (-0.22-0.13) | 6.29E-01 | 6.84E-01 |
| 75 | TG(52:1) [NL-18:0] | -1.39 | (-1.48--1.31) | **4.15E-163** | **1.72E-162** | -2.70 | (-2.8--2.6) | **0.00E+00** | **0.00E+00** | 1.06 | (0.95-1.17) | **7.70E-71** | **1.16E-70** | -0.41 | (-0.62--0.21) | **1.04E-04** | **2.16E-04** |
| 76 | TG(52:1) [NL-18:1] | -1.30 | (-1.39--1.22) | **4.99E-153** | **1.84E-152** | -2.59 | (-2.69--2.49) | **0.00E+00** | **0.00E+00** | 0.99 | (0.89-1.1) | **1.12E-64** | **1.64E-64** | -0.39 | (-0.59--0.18) | **1.88E-04** | **3.73E-04** |
| 77 | TG(52:2) [NL-16:0] | -0.62 | (-0.67--0.58) | **4.69E-125** | **1.39E-124** | -2.67 | (-2.74--2.6) | **0.00E+00** | **0.00E+00** | 1.69 | (1.6-1.77) | **5.95E-218** | **1.74E-217** | -0.04 | (-0.16-0.07) | 4.86E-01 | 5.45E-01 |
| 78 | TG(52:2) [NL-18:2] | -0.96 | (-1.03--0.89) | **3.61E-134** | **1.15E-133** | -2.30 | (-2.4--2.21) | **1.37E-280** | **2.42E-280** | 0.96 | (0.85-1.07) | **8.09E-62** | **1.17E-61** | -0.15 | (-0.31-0.02) | 8.12E-02 | 1.06E-01 |
| 79 | TG(52:3) [NL-16:1] | -0.43 | (-0.49--0.37) | **4.20E-44** | **7.12E-44** | -2.11 | (-2.21--2.02) | **9.79E-270** | **1.70E-269** | 1.05 | (0.95-1.16) | **8.71E-76** | **1.33E-75** | 0.12 | (-0.03-0.27) | 1.12E-01 | 1.43E-01 |
| 80 | TG(52:3) [NL-18:2] | -0.61 | (-0.66--0.56) | **6.80E-113** | **1.90E-112** | -3.57 | (-3.67--3.48) | **0.00E+00** | **0.00E+00** | 2.32 | (2.21-2.43) | **1.39E-227** | **4.22E-227** | 0.17 | (0.05-0.3) | **6.42E-03** | **1.03E-02** |
| 81 | TG(52:4) [NL-16:1] | -0.73 | (-0.8--0.65) | **1.54E-71** | **3.13E-71** | -3.21 | (-3.32--3.11) | **0.00E+00** | **0.00E+00** | 1.44 | (1.32-1.56) | **2.34E-101** | **3.87E-101** | 0.40 | (0.21-0.59) | **4.13E-05** | **9.35E-05** |
| 82 | TG(52:4) [NL-18:2] | -0.92 | (-1--0.85) | **1.03E-101** | **2.62E-101** | -4.34 | (-4.46--4.22) | **0.00E+00** | **0.00E+00** | 2.37 | (2.24-2.49) | **3.17E-186** | **7.68E-186** | 0.46 | (0.26-0.65) | **4.94E-06** | **1.26E-05** |
| 83 | TG(52:4) [NL-18:3] | -0.87 | (-0.94--0.79) | **5.45E-92** | **1.25E-91** | -3.90 | (-4.01--3.8) | **0.00E+00** | **0.00E+00** | 2.27 | (2.16-2.38) | **3.12E-210** | **8.71E-210** | 0.22 | (0.02-0.41) | **2.86E-02** | **4.06E-02** |
| 84 | TG(52:5) [NL-18:3] | -0.91 | (-1.01--0.82) | **3.44E-72** | **7.06E-72** | -3.82 | (-3.93--3.7) | **0.00E+00** | **0.00E+00** | 1.98 | (1.87-2.09) | **1.89E-175** | **4.36E-175** | 0.36 | (0.14-0.57) | **9.94E-04** | **1.78E-03** |
| 85 | TG(52:5) [NL-20:4] | -1.62 | (-1.73--1.51) | **9.17E-145** | **3.21E-144** | -1.32 | (-1.45--1.19) | **1.59E-78** | **1.86E-78** | -1.20 | (-1.32--1.07) | **9.50E-69** | **1.41E-68** | 0.10 | (-0.13-0.33) | 3.98E-01 | 4.56E-01 |
| 86 | TG(52:5) [NL-20:5] | -1.24 | (-1.36--1.12) | **1.42E-83** | **3.08E-83** | -2.16 | (-2.3--2.03) | **2.22E-165** | **3.08E-165** | -0.07 | (-0.2-0.05) | 2.39E-01 | 2.42E-01 | 0.37 | (0.13-0.61) | **2.59E-03** | **4.40E-03** |
| 87 | TG(53:2) [NL-18:1] | -0.75 | (-0.81--0.68) | **2.02E-94** | **4.82E-94** | -2.82 | (-2.9--2.74) | **0.00E+00** | **0.00E+00** | 1.76 | (1.67-1.84) | **4.64E-217** | **1.35E-216** | -0.19 | (-0.34--0.03) | **1.76E-02** | **2.58E-02** |
| 88 | TG(54:1) [NL-18:1] | -0.87 | (-0.94--0.8) | **4.70E-110** | **1.26E-109** | -2.47 | (-2.56--2.38) | **0.00E+00** | **0.00E+00** | 1.40 | (1.3-1.5) | **4.13E-123** | **7.59E-123** | -0.38 | (-0.55--0.22) | **5.29E-06** | **1.34E-05** |
| 89 | TG(54:2) [NL-18:0] | -0.72 | (-0.79--0.65) | **2.01E-78** | **4.24E-78** | -2.46 | (-2.55--2.37) | **0.00E+00** | **0.00E+00** | 1.42 | (1.33-1.52) | **5.62E-137** | **1.07E-136** | -0.26 | (-0.42--0.1) | **1.84E-03** | **3.18E-03** |
| 90 | TG(54:2) [NL-20:1] | -1.83 | (-1.9--1.75) | **7.34E-294** | **9.78E-293** | -4.08 | (-4.17--3.98) | **0.00E+00** | **0.00E+00** | 2.05 | (1.96-2.15) | **1.80E-239** | **5.82E-239** | -0.19 | (-0.36--0.02) | **2.46E-02** | **3.56E-02** |
| 91 | TG(54:3) [NL-18:1] | -0.22 | (-0.28--0.17) | **1.53E-16** | **2.07E-16** | -2.16 | (-2.24--2.09) | **0.00E+00** | **0.00E+00** | 1.71 | (1.63-1.79) | **9.12E-223** | **2.72E-222** | -0.08 | (-0.2-0.04) | 1.82E-01 | 2.23E-01 |
| 92 | TG(54:3) [NL-18:2] | -0.37 | (-0.42--0.31) | **1.28E-37** | **2.05E-37** | -2.52 | (-2.61--2.43) | **0.00E+00** | **0.00E+00** | 1.66 | (1.57-1.75) | **1.31E-187** | **3.18E-187** | 0.05 | (-0.07-0.18) | 4.00E-01 | 4.57E-01 |
| 93 | TG(54:4) [NL-18:2] | -0.23 | (-0.28--0.17) | **1.06E-13** | **1.39E-13** | -3.24 | (-3.34--3.14) | **0.00E+00** | **0.00E+00** | 2.50 | (2.39-2.61) | **7.34E-246** | **2.43E-245** | 0.14 | (0-0.28) | **4.84E-02** | 6.58E-02 |
| 94 | TG(54:4) [NL-20:3] | -1.19 | (-1.26--1.12) | **1.34E-176** | **6.24E-176** | -1.65 | (-1.75--1.55) | **5.80E-181** | **8.16E-181** | 0.12 | (0.02-0.23) | **1.83E-02** | **1.89E-02** | -0.22 | (-0.39--0.06) | **7.93E-03** | **1.25E-02** |
| 95 | TG(54:5) [NL-18:3] | -0.47 | (-0.55--0.39) | **5.20E-28** | **7.73E-28** | -3.63 | (-3.74--3.53) | **0.00E+00** | **0.00E+00** | 2.49 | (2.39-2.6) | **9.16E-255** | **3.14E-254** | 0.23 | (0.04-0.43) | **1.91E-02** | **2.80E-02** |
| 96 | TG(54:5) [NL-20:4] | -0.90 | (-0.96--0.83) | **4.74E-125** | **1.40E-124** | -1.71 | (-1.82--1.61) | **1.31E-166** | **1.82E-166** | 0.08 | (-0.03-0.2) | 1.42E-01 | 1.44E-01 | 0.16 | (0.01-0.32) | **4.03E-02** | 5.59E-02 |
| 97 | TG(54:6) [NL-18:3] | -0.59 | (-0.69--0.48) | **6.58E-28** | **9.71E-28** | -3.83 | (-3.95--3.71) | **0.00E+00** | **0.00E+00** | 2.47 | (2.35-2.58) | **1.04E-238** | **3.32E-238** | 0.41 | (0.19-0.63) | **3.19E-04** | **6.08E-04** |
| 98 | TG(54:6) [NL-20:4] | -1.15 | (-1.23--1.07) | **1.09E-137** | **3.53E-137** | -2.10 | (-2.21--1.99) | **4.80E-200** | **7.05E-200** | -0.10 | (-0.22-0.02) | 1.02E-01 | 1.04E-01 | 0.39 | (0.2-0.57) | **3.97E-05** | **9.06E-05** |
| 99 | TG(54:6) [NL-20:5] | -0.48 | (-0.59--0.38) | **2.65E-18** | **3.64E-18** | -2.43 | (-2.56--2.3) | **2.70E-200** | **3.98E-200** | 0.75 | (0.61-0.88) | **4.18E-27** | **5.15E-27** | 0.63 | (0.4-0.87) | **1.71E-07** | **5.19E-07** |
| 100 | TG(54:6) [NL-22:6] | -2.12 | (-2.24--2.01) | **3.98E-195** | **2.14E-194** | -1.67 | (-1.81--1.53) | **4.29E-104** | **5.21E-104** | -1.47 | (-1.61--1.34) | **2.01E-82** | **3.12E-82** | 0.19 | (-0.07-0.44) | 1.58E-01 | 1.96E-01 |
| 101 | TG(54:7) [NL-20:5] | -0.45 | (-0.56--0.34) | **1.76E-14** | **2.32E-14** | -2.18 | (-2.32--2.04) | **1.20E-158** | **1.64E-158** | 0.45 | (0.32-0.58) | **1.48E-11** | **1.64E-11** | 0.75 | (0.52-0.99) | **3.39E-10** | **1.44E-09** |
| 102 | TG(54:7) [NL-22:6] | -1.56 | (-1.68--1.44) | **4.06E-115** | **1.16E-114** | -0.77 | (-0.9--0.63) | **8.71E-27** | **9.61E-27** | -2.04 | (-2.17--1.91) | **4.94E-148** | **9.75E-148** | 0.38 | (0.12-0.63) | **4.50E-03** | **7.34E-03** |
| 103 | TG(56:6) [NL-20:4] | 0.06 | (0.01-0.1) | **1.70E-02** | **1.84E-02** | -1.21 | (-1.3--1.12) | **3.91E-123** | **4.95E-123** | 0.60 | (0.5-0.7) | **2.47E-31** | **3.08E-31** | 0.30 | (0.2-0.41) | **3.95E-08** | **1.30E-07** |

| | | | | | | | | | | | | | | | | | |
|---|---|---|---|---|---|---|---|---|---|---|---|---|---|---|---|---|---|
| 104 | TG(56:6) [NL-22:5] | -0.62 | (-0.68--0.55) | 2.10E-71 | 4.26E-71 | -1.12 | (-1.22--1.02) | 1.99E-94 | 2.37E-94 | -0.18 | (-0.28--0.08) | 6.22E-04 | 6.52E-04 | 0.20 | (0.05-0.35) | 8.38E-03 | 1.32E-02 |
| 105 | TG(56:7) [NL-20:4] | -0.52 | (-0.58--0.46) | 4.34E-55 | 7.92E-55 | -1.80 | (-1.9--1.69) | 1.20E-184 | 1.70E-184 | 0.48 | (0.37-0.59) | 1.36E-16 | 1.57E-16 | 0.36 | (0.21-0.51) | 2.24E-06 | 6.10E-06 |
| 106 | TG(56:7) [NL-20:5] | 0.07 | (-0.03-0.16) | 1.94E-01 | 2.04E-01 | -2.09 | (-2.21--1.97) | 3.52E-187 | 4.98E-187 | 1.10 | (0.99-1.22) | 2.15E-69 | 3.21E-69 | 0.60 | (0.39-0.81) | 2.23E-08 | 7.54E-08 |
| 107 | TG(56:7) [NL-22:5] | -0.96 | (-1.03--0.88) | 1.14E-112 | 3.13E-112 | -1.67 | (-1.77--1.56) | 1.51E-162 | 2.08E-162 | -0.33 | (-0.44--0.23) | 2.74E-09 | 2.98E-09 | 0.42 | (0.25-0.6) | 3.69E-06 | 9.63E-06 |
| 108 | TG(56:7) [NL-22:6] | -1.23 | (-1.33--1.12) | 6.53E-98 | 1.59E-97 | -1.74 | (-1.87--1.61) | 6.82E-125 | 8.67E-125 | -0.67 | (-0.8--0.54) | 1.07E-22 | 1.27E-22 | 0.48 | (0.24-0.72) | 1.03E-04 | 2.14E-04 |
| 109 | TG(56:8) [NL-20:4] | -0.67 | (-0.75--0.59) | 1.09E-55 | 2.01E-55 | -1.15 | (-1.26--1.05) | 1.40E-85 | 1.64E-85 | -0.25 | (-0.36--0.15) | 3.06E-06 | 3.25E-06 | 0.39 | (0.22-0.56) | 5.86E-06 | 1.46E-05 |
| 110 | TG(56:8) [NL-20:5] | -0.09 | (-0.19-0.02) | 1.17E-01 | 1.24E-01 | -1.97 | (-2.1--1.84) | 2.42E-155 | 3.27E-155 | 0.83 | (0.72-0.95) | 5.75E-42 | 7.59E-42 | 0.75 | (0.53-0.96) | 2.50E-11 | 1.24E-10 |
| 111 | TG(56:8) [NL-22:6] | -1.33 | (-1.44--1.22) | 7.02E-102 | 1.81E-101 | -1.72 | (-1.86--1.59) | 3.34E-112 | 4.12E-112 | -0.99 | (-1.12--0.86) | 6.66E-44 | 8.83E-44 | 0.65 | (0.4-0.89) | 2.57E-07 | 7.62E-07 |
| 112 | TG(56:9) [NL-22:6] | -0.97 | (-1.08--0.87) | 2.50E-66 | 4.90E-66 | -1.05 | (-1.18--0.92) | 2.66E-53 | 3.07E-53 | -1.17 | (-1.3--1.05) | 1.98E-67 | 2.91E-67 | 0.63 | (0.4-0.85) | 4.93E-08 | 1.60E-07 |
| 113 | TG(58:10) [NL-22:6] | -0.83 | (-0.93--0.72) | 2.99E-51 | 5.31E-51 | -0.92 | (-1.05--0.8) | 7.73E-45 | 8.79E-45 | -1.05 | (-1.16--0.93) | 5.09E-59 | 7.28E-59 | 0.66 | (0.44-0.88) | 6.07E-09 | 2.19E-08 |
| 114 | TG(58:8) [NL-22:6] | -0.89 | (-0.99--0.79) | 2.64E-60 | 4.96E-60 | -1.18 | (-1.3--1.06) | 2.46E-73 | 2.86E-73 | -0.82 | (-0.94--0.7) | 7.08E-37 | 9.16E-37 | 0.53 | (0.3-0.76) | 6.96E-06 | 1.71E-05 |
| 115 | TG(58:9) [NL-22:6] | -0.96 | (-1.06--0.85) | 1.92E-66 | 3.77E-66 | -1.30 | (-1.43--1.18) | 1.16E-81 | 1.37E-81 | -0.84 | (-0.97--0.72) | 1.77E-38 | 2.31E-38 | 0.67 | (0.44-0.89) | 1.34E-08 | 4.78E-08 |
| 116 | TG(O-50:1) [NL-16:0] | -0.61 | (-0.67--0.55) | 2.41E-77 | 5.02E-77 | -3.21 | (-3.28--3.14) | 0.00E+00 | 0.00E+00 | 2.64 | (2.57-2.72) | 0.00E+00 | 0.00E+00 | -0.11 | (-0.26-0.04) | 1.38E-01 | 1.73E-01 |
| 117 | TG(O-50:1) [NL-17:1] | -1.72 | (-1.8--1.64) | 3.59E-246 | 2.83E-245 | -2.48 | (-2.57--2.39) | 0.00E+00 | 0.00E+00 | 0.35 | (0.26-0.43) | 3.29E-15 | 3.72E-15 | -0.17 | (-0.35-0) | 4.92E-02 | 6.65E-02 |
| 118 | TG(O-50:1) [NL-18:1] | -0.49 | (-0.56--0.42) | 1.36E-40 | 2.23E-40 | -3.35 | (-3.43--3.28) | 0.00E+00 | 0.00E+00 | 2.83 | (2.75-2.91) | 0.00E+00 | 0.00E+00 | 0.04 | (-0.13-0.21) | 6.64E-01 | 7.13E-01 |
| 119 | TG(O-50:2) [NL-18:2] | -0.54 | (-0.6--0.48) | 3.60E-66 | 6.99E-66 | -2.67 | (-2.74--2.6) | 0.00E+00 | 0.00E+00 | 2.02 | (1.94-2.09) | 8.53E-306 | 3.94E-305 | -0.03 | (-0.17-0.11) | 7.10E-01 | 7.54E-01 |
| 120 | TG(O-52:2) [NL-16:0] | -0.77 | (-0.85--0.7) | 1.30E-74 | 2.69E-74 | -3.26 | (-3.35--3.16) | 0.00E+00 | 0.00E+00 | 2.45 | (2.35-2.55) | 5.33E-271 | 2.03E-270 | -0.31 | (-0.49--0.12) | 1.04E-03 | 1.85E-03 |
| 121 | TG(O-52:2) [NL-17:1] | -1.28 | (-1.35--1.21) | 6.56E-188 | 3.31E-187 | -3.03 | (-3.11--2.94) | 0.00E+00 | 0.00E+00 | 1.16 | (1.07-1.25) | 1.93E-109 | 3.30E-109 | -0.05 | (-0.22-0.13) | 6.09E-01 | 6.67E-01 |
| 122 | TG(O-52:2) [NL-18:1] | -0.68 | (-0.76--0.6) | 6.82E-55 | 1.24E-54 | -3.18 | (-3.28--3.08) | 0.00E+00 | 0.00E+00 | 2.49 | (2.38-2.59) | 4.19E-259 | 1.46E-258 | -0.32 | (-0.51--0.12) | 1.47E-03 | 2.56E-03 |
| 123 | TG(O-54:2) [NL-18:1] | -0.74 | (-0.81--0.68) | 3.63E-94 | 8.57E-94 | -2.82 | (-2.9--2.74) | 0.00E+00 | 0.00E+00 | 1.75 | (1.67-1.84) | 4.23E-216 | 1.21E-215 | -0.18 | (-0.33--0.02) | 2.40E-02 | 3.48E-02 |
| 124 | TG(O-54:4) [NL-17:1] | -0.65 | (-0.73--0.58) | 1.71E-61 | 3.24E-61 | -3.35 | (-3.45--3.24) | 0.00E+00 | 0.00E+00 | 1.80 | (1.69-1.91) | 7.38E-160 | 1.54E-159 | 0.35 | (0.18-0.52) | 8.08E-05 | 1.72E-04 |
| 125 | TG(O-54:4) [NL-18:2] | -0.62 | (-0.7--0.54) | 1.01E-50 | 1.80E-50 | -3.63 | (-3.74--3.51) | 0.00E+00 | 0.00E+00 | 2.14 | (2.02-2.26) | 2.50E-183 | 5.97E-183 | 0.38 | (0.2-0.57) | 5.90E-05 | 1.27E-04 |
| 126 | dhCer(d18:0/22:0) | -0.05 | (-0.11-0) | 6.91E-02 | 7.36E-02 | 0.09 | (0.03-0.16) | 4.40E-03 | 4.57E-03 | -0.25 | (-0.32--0.19) | 1.04E-14 | 1.17E-14 | -0.10 | (-0.22-0.02) | 1.07E-01 | 1.38E-01 |
| 127 | dhCer(d18:0/24:0) | 0.02 | (-0.03-0.08) | 3.88E-01 | 4.03E-01 | 0.46 | (0.39-0.52) | 1.07E-41 | 1.21E-41 | -0.50 | (-0.57--0.44) | 3.45E-50 | 4.65E-50 | -0.04 | (-0.15-0.07) | 4.72E-01 | 5.30E-01 |
| 128 | dhCer(d18:0/24:1) | -0.36 | (-0.42--0.3) | 4.60E-29 | 6.89E-29 | 0.03 | (-0.04-0.11) | 3.97E-01 | 3.98E-01 | -0.67 | (-0.75--0.6) | 4.21E-57 | 5.91E-57 | 0.10 | (-0.03-0.24) | 1.44E-01 | 1.79E-01 |
| 129 | Cer(d16:1/22:0) | 0.07 | (0.02-0.13) | 9.09E-03 | 9.93E-03 | -2.36 | (-2.43--2.29) | 0.00E+00 | 0.00E+00 | 2.22 | (2.15-2.3) | 0.00E+00 | 0.00E+00 | 0.00 | (-0.12-0.13) | 9.71E-01 | 9.78E-01 |
| 130 | Cer(d16:1/23:0) | 0.12 | (0.06-0.18) | 1.00E-04 | 1.16E-04 | -3.18 | (-3.28--3.08) | 0.00E+00 | 0.00E+00 | 3.23 | (3.11-3.34) | 0.00E+00 | 0.00E+00 | -0.14 | (-0.27-0) | 5.21E-02 | 7.00E-02 |
| 131 | Cer(d16:1/24:0) | 0.31 | (0.26-0.37) | 1.57E-26 | 2.28E-26 | -2.66 | (-2.73--2.59) | 0.00E+00 | 0.00E+00 | 2.85 | (2.77-2.92) | 0.00E+00 | 0.00E+00 | 0.07 | (-0.06-0.2) | 2.79E-01 | 3.31E-01 |
| 132 | Cer(d16:1/24:1) | 0.17 | (0.12-0.23) | 1.86E-09 | 2.33E-09 | -2.07 | (-2.15--2) | 0.00E+00 | 0.00E+00 | 2.01 | (1.93-2.09) | 1.53E-273 | 5.98E-273 | 0.13 | (0.01-0.26) | 3.09E-02 | 4.35E-02 |
| 133 | Cer(d17:1/22:0) | 0.24 | (0.18-0.3) | 1.03E-15 | 1.38E-15 | -1.87 | (-1.95--1.79) | 4.56E-264 | 7.70E-264 | 1.97 | (1.88-2.05) | 1.61E-248 | 5.37E-248 | -0.03 | (-0.15-0.1) | 7.02E-01 | 7.48E-01 |
| 134 | Cer(d17:1/23:0) | 0.23 | (0.17-0.29) | 1.49E-13 | 1.95E-13 | -3.32 | (-3.44--3.21) | 0.00E+00 | 0.00E+00 | 3.42 | (3.29-3.55) | 1.20E-292 | 4.96E-292 | -0.04 | (-0.18-0.1) | 5.93E-01 | 6.55E-01 |
| 135 | Cer(d17:1/24:0) | 0.43 | (0.38-0.48) | 2.44E-55 | 4.47E-55 | -2.04 | (-2.11--1.97) | 0.00E+00 | 0.00E+00 | 2.40 | (2.32-2.47) | 0.00E+00 | 0.00E+00 | 0.04 | (-0.08-0.16) | 4.69E-01 | 5.28E-01 |
| 136 | Cer(d17:1/24:1) | 0.26 | (0.2-0.31) | 2.71E-20 | 3.77E-20 | -2.20 | (-2.27--2.12) | 0.00E+00 | 0.00E+00 | 2.20 | (2.12-2.28) | 2.97E-300 | 1.31E-299 | 0.22 | (0.1-0.34) | 2.78E-04 | 5.39E-04 |
| 137 | Cer(d18:1/16:0) | -0.27 | (-0.31--0.23) | 5.49E-37 | 8.73E-37 | -2.08 | (-2.14--2.03) | 0.00E+00 | 0.00E+00 | 1.61 | (1.56-1.67) | 0.00E+00 | 0.00E+00 | 0.22 | (0.13-0.31) | 9.07E-07 | 2.61E-06 |
| 138 | Cer(d18:1/18:0) | -0.13 | (-0.19--0.08) | 4.60E-06 | 5.51E-06 | -1.62 | (-1.69--1.55) | 4.59E-280 | 8.10E-280 | 1.25 | (1.19-1.32) | 1.01E-194 | 2.54E-194 | -0.03 | (-0.15-0.09) | 6.36E-01 | 6.91E-01 |
| 139 | Cer(d18:1/20:0) | -0.19 | (-0.24--0.14) | 1.44E-14 | 1.91E-14 | -2.05 | (-2.11--1.99) | 0.00E+00 | 0.00E+00 | 1.53 | (1.47-1.59) | 2.78E-274 | 1.09E-273 | 0.21 | (0.11-0.31) | 8.77E-05 | 1.84E-04 |
| 140 | Cer(d18:1/22:0) | 0.14 | (0.1-0.18) | 1.56E-10 | 1.97E-10 | -2.24 | (-2.29--2.19) | 0.00E+00 | 0.00E+00 | 2.13 | (2.08-2.19) | 0.00E+00 | 0.00E+00 | 0.11 | (0.02-0.21) | 1.94E-02 | 2.83E-02 |
| 141 | Cer(d18:1/23:0) | 0.08 | (0.04-0.12) | 3.09E-04 | 3.54E-04 | -3.73 | (-3.8--3.67) | 0.00E+00 | 0.00E+00 | 3.69 | (3.62-3.76) | 0.00E+00 | 0.00E+00 | 0.05 | (-0.05-0.15) | 3.23E-01 | 3.78E-01 |
| 142 | Cer(d18:1/24:0) | 0.24 | (0.2-0.28) | 5.48E-28 | 8.12E-28 | -1.97 | (-2.03--1.92) | 0.00E+00 | 0.00E+00 | 2.05 | (1.99-2.1) | 0.00E+00 | 0.00E+00 | 0.13 | (0.04-0.22) | 4.54E-03 | 7.38E-03 |
| 143 | Cer(d18:1/24:1) | 0.01 | (-0.04-0.05) | 7.40E-01 | 7.46E-01 | -2.23 | (-2.29--2.16) | 0.00E+00 | 0.00E+00 | 1.91 | (1.85-1.97) | 0.00E+00 | 0.00E+00 | 0.32 | (0.23-0.42) | 9.38E-11 | 4.37E-10 |
| 144 | Cer(d18:2/16:0) | -0.39 | (-0.44--0.33) | 3.09E-40 | 5.00E-40 | -2.21 | (-2.28--2.13) | 0.00E+00 | 0.00E+00 | 1.32 | (1.24-1.4) | 2.55E-157 | 5.23E-157 | 0.44 | (0.32-0.56) | 1.64E-12 | 9.61E-12 |
| 145 | Cer(d18:2/22:0) | 0.43 | (0.38-0.48) | 3.15E-57 | 5.85E-57 | -1.43 | (-1.5--1.37) | 1.19E-258 | 1.97E-258 | 1.50 | (1.44-1.57) | 2.28E-250 | 7.67E-250 | 0.32 | (0.21-0.44) | 1.49E-08 | 5.27E-08 |
| 146 | Cer(d18:2/23:0) | 0.36 | (0.3-0.41) | 1.73E-38 | 2.77E-38 | -3.30 | (-3.39--3.22) | 0.00E+00 | 0.00E+00 | 3.43 | (3.35-3.52) | 0.00E+00 | 0.00E+00 | 0.23 | (0.12-0.35) | 7.37E-05 | 1.58E-04 |
| 147 | Cer(d18:2/24:0) | 0.52 | (0.47-0.57) | 2.50E-84 | 5.50E-84 | -1.73 | (-1.79--1.66) | 0.00E+00 | 0.00E+00 | 1.94 | (1.88-2.01) | 0.00E+00 | 0.00E+00 | 0.39 | (0.29-0.5) | 9.10E-13 | 5.39E-12 |
| 148 | Cer(d18:2/24:1) | 0.39 | (0.34-0.44) | 2.50E-48 | 4.37E-48 | -1.50 | (-1.57--1.43) | 6.62E-264 | 1.12E-263 | 1.48 | (1.41-1.55) | 3.19E-234 | 1.00E-233 | 0.48 | (0.37-0.59) | 1.05E-17 | 1.25E-16 |
| 149 | Cer(d19:1/24:0) | 0.72 | (0.65-0.8) | 3.91E-66 | 7.56E-66 | -0.95 | (-1.05--0.86) | 8.83E-81 | 1.03E-80 | 1.78 | (1.69-1.87) | 6.50E-197 | 1.67E-196 | -0.38 | (-0.56--0.21) | 2.07E-05 | 4.91E-05 |
| 150 | Cer(d19:1/24:1) | 0.47 | (0.38-0.56) | 1.28E-24 | 1.85E-24 | -1.30 | (-1.41--1.19) | 2.73E-102 | 3.30E-102 | 1.68 | (1.57-1.79) | 4.24E-145 | 8.28E-145 | -0.20 | (-0.4--0.01) | 4.33E-02 | 5.93E-02 |
| 151 | Cer(m18:0/22:0) | -0.90 | (-0.98--0.83) | 7.50E-114 | 2.10E-113 | -2.36 | (-2.45--2.27) | 0.00E+00 | 0.00E+00 | 1.25 | (1.16-1.34) | 4.66E-126 | 8.71E-126 | -0.19 | (-0.33--0.04) | 1.23E-02 | 1.90E-02 |
| 152 | Cer(m18:0/23:0) | -0.93 | (-1.01--0.85) | 1.54E-101 | 3.90E-101 | -4.22 | (-4.37--4.07) | 0.00E+00 | 0.00E+00 | 3.13 | (2.97-3.29) | 6.84E-196 | 1.74E-195 | -0.26 | (-0.42--0.09) | 2.43E-03 | 4.16E-03 |
| 153 | Cer(m18:0/24:0) | -0.72 | (-0.79--0.65) | 1.37E-83 | 3.00E-83 | -2.29 | (-2.37--2.2) | 0.00E+00 | 0.00E+00 | 1.37 | (1.29-1.44) | 1.48E-183 | 3.56E-183 | -0.09 | (-0.22-0.05) | 2.08E-01 | 2.53E-01 |
| 154 | Cer(m18:0/24:1) | -0.98 | (-1.05--0.9) | 9.17E-115 | 2.60E-114 | -3.03 | (-3.13--2.93) | 0.00E+00 | 0.00E+00 | 1.62 | (1.52-1.72) | 5.41E-153 | 1.09E-152 | 0.02 | (-0.14-0.17) | 8.10E-01 | 8.44E-01 |
| 155 | Cer(m18:1/22:0) | -0.53 | (-0.6--0.45) | 1.94E-40 | 3.16E-40 | -1.47 | (-1.56--1.38) | 4.62E-176 | 6.44E-176 | 0.28 | (0.2-0.36) | 3.46E-11 | 3.80E-11 | 0.24 | (0.08-0.4) | 3.74E-03 | 6.18E-03 |
| 156 | Cer(m18:1/23:0) | -0.55 | (-0.63--0.47) | 1.50E-38 | 2.42E-38 | -3.46 | (-3.61--3.32) | 8.74E-276 | 1.52E-275 | 2.28 | (2.12-2.44) | 4.73E-131 | 8.91E-131 | 0.16 | (-0.02-0.33) | 7.72E-02 | 1.01E-01 |
| 157 | Cer(m18:1/24:0) | -0.33 | (-0.4--0.26) | 1.09E-20 | 1.52E-20 | -1.31 | (-1.4--1.23) | 1.39E-159 | 1.90E-159 | 0.42 | (0.34-0.5) | 5.98E-24 | 7.23E-24 | 0.28 | (0.13-0.42) | 1.93E-04 | 3.82E-04 |
| 158 | Cer(m18:1/24:1) | -0.56 | (-0.63--0.48) | 1.31E-43 | 2.19E-43 | -2.06 | (-2.15--1.97) | 9.31E-256 | 1.52E-255 | 0.77 | (0.68-0.86) | 8.58E-57 | 1.20E-56 | 0.35 | (0.2-0.51) | 1.17E-05 | 2.80E-05 |

| | | | | | | | | | | | | | | | | | |
|---|---|---|---|---|---|---|---|---|---|---|---|---|---|---|---|---|---|
| 159 | GM3(d18:1/18:0) | 0.49 | (0.44-0.53) | **6.86E-81** | **1.46E-80** | 0.65 | (0.59-0.71) | **1.05E-89** | **1.24E-89** | -0.13 | (-0.18--0.07) | **1.76E-05** | **1.86E-05** | 0.07 | (-0.02-0.17) | 1.40E-01 | 1.75E-01 |
| 160 | GM3(d18:1/22:0) | 0.08 | (0.02-0.13) | **5.33E-03** | **5.90E-03** | -0.95 | (-1.03--0.87) | **6.35E-100** | **7.66E-100** | 0.90 | (0.82-0.99) | **9.00E-88** | **1.43E-87** | 0.15 | (0.04-0.27) | **6.89E-03** | **1.11E-02** |
| 161 | GM3(d18:1/24:0) | 0.52 | (0.47-0.57) | **6.00E-72** | **1.23E-71** | 0.19 | (0.11-0.26) | **4.75E-07** | **5.04E-07** | 0.34 | (0.27-0.41) | **1.63E-21** | **1.92E-21** | 0.27 | (0.16-0.37) | **6.14E-07** | **1.79E-06** |
| 162 | GM3(d18:1/24:1) | 0.03 | (-0.02-0.08) | 2.76E-01 | 2.89E-01 | -0.49 | (-0.56--0.42) | **3.80E-39** | **4.27E-39** | 0.37 | (0.3-0.44) | **1.11E-23** | **1.34E-23** | 0.39 | (0.28-0.5) | **1.05E-11** | **5.56E-11** |
| 163 | HexCer(d16:1/22:0) | 0.05 | (-0.01-0.11) | 7.71E-02 | 8.17E-02 | -2.22 | (-2.32--2.11) | **2.12E-235** | **3.35E-235** | 2.03 | (1.92-2.15) | **8.46E-174** | **1.93E-173** | 0.22 | (0.09-0.36) | **8.73E-04** | **1.59E-03** |
| 164 | HexCer(d16:1/24:0) | 0.37 | (0.31-0.43) | **1.78E-32** | **2.74E-32** | -1.70 | (-1.8--1.6) | **1.22E-179** | **1.71E-179** | 1.78 | (1.67-1.88) | **4.70E-164** | **1.00E-163** | 0.31 | (0.18-0.45) | **3.55E-06** | **9.37E-06** |
| 165 | HexCer(d18:1/16:0) | 0.08 | (0.04-0.12) | **1.81E-04** | **2.08E-04** | -1.70 | (-1.77--1.64) | **3.55E-307** | **6.65E-307** | 1.85 | (1.78-1.92) | **1.80E-298** | **7.79E-298** | 0.00 | (-0.09-0.09) | 9.84E-01 | 9.86E-01 |
| 166 | HexCer(d18:1/18:0) | -0.14 | (-0.19--0.08) | **1.50E-06** | **1.81E-06** | -1.30 | (-1.38--1.21) | **2.43E-158** | **3.30E-158** | 0.95 | (0.87-1.04) | **2.01E-88** | **3.21E-88** | 0.42 | (0.29-0.54) | **1.18E-10** | **5.31E-10** |
| 167 | HexCer(d18:1/20:0) | -0.10 | (-0.15--0.05) | **1.03E-04** | **1.19E-04** | -1.39 | (-1.46--1.31) | **4.54E-191** | **6.52E-191** | 1.11 | (1.03-1.19) | **1.54E-124** | **2.85E-124** | 0.34 | (0.24-0.45) | **5.96E-10** | **2.51E-09** |
| 168 | HexCer(d18:1/22:0) | 0.11 | (0.07-0.16) | **8.86E-07** | **1.07E-06** | -1.81 | (-1.87--1.75) | **0.00E+00** | **0.00E+00** | 1.74 | (1.68-1.8) | **4.88E-308** | **2.34E-307** | 0.35 | (0.25-0.45) | **7.19E-12** | **3.88E-11** |
| 169 | HexCer(d18:1/24:0) | 0.25 | (0.21-0.3) | **6.16E-27** | **8.95E-27** | -0.77 | (-0.83--0.7) | **8.61E-103** | **1.04E-102** | 0.87 | (0.81-0.94) | **1.45E-121** | **2.63E-121** | 0.37 | (0.28-0.47) | **1.77E-13** | **1.15E-12** |
| 170 | HexCer(d18:1/24:1) | 0.02 | (-0.03-0.07) | 4.82E-01 | 4.98E-01 | -1.22 | (-1.29--1.16) | **1.08E-204** | **1.59E-204** | 1.20 | (1.14-1.27) | **1.59E-190** | **3.94E-190** | 0.25 | (0.15-0.36) | **3.03E-06** | **8.12E-06** |
| 171 | Hex2Cer(d16:1/16:0) | 0.00 | (-0.05-0.06) | 9.37E-01 | 9.37E-01 | -3.13 | (-3.22--3.04) | **0.00E+00** | **0.00E+00** | 2.58 | (2.48-2.67) | **1.95E-298** | **8.35E-298** | 0.42 | (0.29-0.55) | **1.87E-10** | **8.15E-10** |
| 172 | Hex2Cer(d18:1/16:0) | -0.14 | (-0.19--0.09) | **1.01E-08** | **1.24E-08** | -2.31 | (-2.37--2.25) | **0.00E+00** | **0.00E+00** | 1.81 | (1.75-1.88) | **3.41E-302** | **1.53E-301** | 0.53 | (0.42-0.64) | **4.02E-20** | **6.23E-19** |
| 173 | Hex2Cer(d18:1/22:0) | 0.22 | (0.17-0.27) | **3.90E-17** | **5.34E-17** | -1.09 | (-1.17--1.02) | **5.16E-148** | **6.86E-148** | 1.01 | (0.94-1.09) | **5.50E-117** | **9.77E-117** | 0.46 | (0.34-0.57) | **6.11E-15** | **4.65E-14** |
| 174 | Hex2Cer(d18:1/24:1) | 0.34 | (0.28-0.39) | **5.63E-33** | **8.68E-33** | -1.57 | (-1.64--1.51) | **7.23E-265** | **1.23E-264** | 1.73 | (1.66-1.81) | **3.65E-264** | **1.33E-263** | 0.47 | (0.35-0.59) | **2.56E-14** | **1.89E-13** |
| 175 | Hex2Cer(d18:2/16:0) | -0.01 | (-0.06-0.03) | 5.56E-01 | 5.67E-01 | -1.93 | (-2--1.86) | **0.00E+00** | **0.00E+00** | 1.38 | (1.31-1.45) | **9.58E-196** | **2.42E-195** | 0.57 | (0.46-0.67) | **1.81E-23** | **4.34E-22** |
| 176 | Hex3Cer(d18:1/16:0) | 0.02 | (-0.02-0.05) | 3.68E-01 | 3.83E-01 | -1.06 | (-1.11--1.01) | **2.03E-233** | **3.19E-233** | 1.03 | (0.98-1.08) | **9.21E-206** | **2.50E-205** | 0.12 | (0.04-0.2) | **2.79E-03** | **4.72E-03** |
| 177 | Hex3Cer(d18:1/22:0) | 0.39 | (0.34-0.45) | **2.67E-43** | **4.46E-43** | -1.16 | (-1.24--1.08) | **4.62E-145** | **6.09E-145** | 1.72 | (1.64-1.8) | **1.19E-231** | **3.68E-231** | 0.05 | (-0.06-0.16) | 3.84E-01 | 4.44E-01 |
| 178 | Hex3Cer(d18:1/24:0) | 0.54 | (0.49-0.59) | **1.56E-85** | **3.46E-85** | -0.05 | (-0.12-0.02) | 1.88E-01 | 1.90E-01 | 0.74 | (0.67-0.81) | **2.44E-83** | **3.80E-83** | 0.07 | (-0.04-0.17) | 2.02E-01 | 2.46E-01 |
| 179 | Hex3Cer(d18:1/24:1) | 0.35 | (0.3-0.4) | **8.59E-44** | **1.45E-43** | -0.91 | (-0.98--0.85) | **2.62E-127** | **3.37E-127** | 1.27 | (1.2-1.34) | **2.34E-192** | **5.82E-192** | 0.15 | (0.05-0.25) | **2.46E-03** | **4.21E-03** |
| 180 | SM(d17:1/14:0) | -0.16 | (-0.22--0.11) | **2.52E-10** | **3.17E-10** | -1.62 | (-1.68--1.57) | **0.00E+00** | **0.00E+00** | 1.16 | (1.1-1.22) | **5.58E-201** | **1.48E-200** | 0.20 | (0.08-0.32) | **8.78E-04** | **1.59E-03** |
| 181 | SM(d18:0/14:0) | -0.23 | (-0.28--0.18) | **2.36E-19** | **3.26E-19** | 0.10 | (0.05-0.15) | **1.72E-04** | **1.80E-04** | -0.43 | (-0.48--0.38) | **1.69E-52** | **2.31E-52** | -0.02 | (-0.13-0.09) | 7.36E-01 | 7.75E-01 |
| 182 | SM(d18:1/14:0)/SM(d16:1/16:0) | -0.20 | (-0.24--0.16) | **1.15E-19** | **1.59E-19** | -1.88 | (-1.93--1.83) | **0.00E+00** | **0.00E+00** | 1.43 | (1.37-1.48) | **1.51E-307** | **7.09E-307** | 0.12 | (0.02-0.21) | **1.58E-02** | **2.34E-02** |
| 183 | SM(d18:2/14:0) | -0.18 | (-0.22--0.13) | **3.78E-14** | **4.97E-14** | -1.05 | (-1.1--1) | **6.38E-229** | **9.85E-229** | 0.33 | (0.27-0.38) | **2.50E-32** | **3.12E-32** | 0.26 | (0.16-0.35) | **1.85E-07** | **5.56E-07** |
| 184 | SM(d17:1/16:0) | -0.14 | (-0.18--0.1) | **2.73E-11** | **3.49E-11** | -1.88 | (-1.93--1.83) | **0.00E+00** | **0.00E+00** | 1.52 | (1.47-1.57) | **0.00E+00** | **0.00E+00** | 0.20 | (0.1-0.29) | **4.19E-05** | **9.45E-05** |
| 185 | SM(d18:1/16:0) | -0.32 | (-0.37--0.27) | **1.83E-37** | **2.92E-37** | -2.31 | (-2.36--2.25) | **0.00E+00** | **0.00E+00** | 1.70 | (1.65-1.76) | **0.00E+00** | **0.00E+00** | 0.42 | (0.32-0.53) | **7.69E-16** | **6.84E-15** |
| 186 | SM(d18:2/16:0) | -0.17 | (-0.2--0.13) | **2.98E-21** | **4.20E-21** | -1.16 | (-1.21--1.12) | **0.00E+00** | **0.00E+00** | 0.54 | (0.49-0.58) | **1.64E-98** | **2.67E-98** | 0.38 | (0.31-0.45) | **7.29E-24** | **2.06E-22** |
| 187 | SM(34:3) | -0.15 | (-0.19--0.11) | **7.39E-12** | **9.51E-12** | -1.16 | (-1.21--1.1) | **8.84E-268** | **1.52E-267** | 0.62 | (0.57-0.67) | **1.26E-92** | **2.04E-92** | 0.17 | (0.08-0.27) | **5.25E-04** | **9.77E-04** |
| 188 | SM(d18:2/17:0) | -0.10 | (-0.14--0.06) | **1.88E-06** | **2.27E-06** | -1.18 | (-1.23--1.13) | **3.42E-282** | **6.10E-282** | 0.70 | (0.65-0.75) | **2.06E-121** | **3.72E-121** | 0.28 | (0.19-0.36) | **7.18E-10** | **2.95E-09** |
| 189 | SM(35:2) (b) | -0.32 | (-0.37--0.27) | **2.35E-31** | **3.56E-31** | -2.48 | (-2.55--2.41) | **0.00E+00** | **0.00E+00** | 1.78 | (1.71-1.85) | **6.56E-279** | **2.60E-278** | 0.46 | (0.34-0.58) | **4.12E-14** | **2.99E-13** |
| 190 | SM(d18:1/18:0)/SM(d16:1/20:0) | -0.34 | (-0.38--0.29) | **8.03E-48** | **1.40E-47** | -1.04 | (-1.09--0.98) | **1.63E-213** | **2.45E-213** | 0.52 | (0.46-0.57) | **7.87E-70** | **1.18E-69** | 0.14 | (0.05-0.23) | **3.73E-03** | **6.17E-03** |
| 191 | SM(d18:2/18:1) | -0.24 | (-0.28--0.2) | **1.01E-31** | **1.55E-31** | -1.60 | (-1.65--1.55) | **0.00E+00** | **0.00E+00** | 1.11 | (1.06-1.17) | **7.91E-228** | **2.42E-227** | 0.14 | (0.05-0.22) | **1.21E-03** | **2.13E-03** |
| 192 | SM(37:1) | -0.33 | (-0.38--0.29) | **4.37E-41** | **7.23E-41** | -2.27 | (-2.33--2.21) | **0.00E+00** | **0.00E+00** | 1.65 | (1.58-1.71) | **1.85E-294** | **7.74E-294** | 0.25 | (0.14-0.35) | **3.04E-06** | **8.12E-06** |
| 193 | SM(37:2) | -0.39 | (-0.45--0.34) | **5.65E-46** | **9.72E-46** | -1.98 | (-2.04--1.92) | **0.00E+00** | **0.00E+00** | 1.05 | (0.99-1.11) | **2.54E-170** | **5.65E-170** | 0.63 | (0.52-0.74) | **2.77E-26** | **1.66E-24** |
| 194 | SM(d18:1/20:0)/SM(d16:1/22:0) | -0.34 | (-0.38--0.3) | **6.67E-68** | **1.32E-67** | -1.41 | (-1.46--1.37) | **0.00E+00** | **0.00E+00** | 0.91 | (0.86-0.96) | **2.42E-196** | **6.19E-196** | 0.13 | (0.05-0.21) | **1.01E-03** | **1.80E-03** |
| 195 | SM(d18:2/20:0) | -0.43 | (-0.47--0.38) | **1.10E-67** | **2.16E-67** | -0.92 | (-0.98--0.86) | **4.22E-150** | **5.65E-150** | 0.16 | (0.1-0.22) | **8.75E-07** | **9.35E-07** | 0.38 | (0.28-0.48) | **3.19E-13** | **1.99E-12** |
| 196 | SM(38:3) (a) | -0.43 | (-0.47--0.39) | **1.93E-86** | **4.28E-86** | -1.16 | (-1.21--1.11) | **1.03E-257** | **1.69E-257** | 0.34 | (0.29-0.39) | **6.09E-36** | **7.84E-36** | 0.26 | (0.17-0.34) | **1.87E-09** | **7.22E-09** |
| 197 | SM(38:3) (b) | -0.33 | (-0.39--0.28) | **1.74E-33** | **2.71E-33** | -2.09 | (-2.16--2.03) | **0.00E+00** | **0.00E+00** | 1.24 | (1.17-1.31) | **6.48E-186** | **1.56E-185** | 0.61 | (0.49-0.73) | **4.74E-22** | **9.10E-21** |
| 198 | SM(d16:1/23:0)/SM(d17:1/22:0) | -0.29 | (-0.33--0.25) | **1.33E-42** | **2.21E-42** | -2.61 | (-2.66--2.56) | **0.00E+00** | **0.00E+00** | 2.25 | (2.2-2.3) | **0.00E+00** | **0.00E+00** | -0.03 | (-0.12-0.07) | 5.67E-01 | 6.29E-01 |
| 199 | SM(d18:1/22:0)/SM(d16:1/24:0) | -0.35 | (-0.39--0.31) | **5.48E-59** | **1.03E-58** | -1.97 | (-2.02--1.92) | **0.00E+00** | **0.00E+00** | 1.46 | (1.41-1.51) | **0.00E+00** | **0.00E+00** | 0.10 | (0.01-0.18) | **2.85E-02** | **4.06E-02** |
| 200 | SM(d16:1/24:1) | -0.34 | (-0.39--0.3) | **3.76E-47** | **6.50E-47** | -2.11 | (-2.16--2.05) | **0.00E+00** | **0.00E+00** | 1.50 | (1.45-1.56) | **1.54E-297** | **6.53E-297** | 0.16 | (0.06-0.26) | **1.71E-03** | **2.97E-03** |
| 201 | SM(d18:2/22:0) | -0.12 | (-0.16--0.08) | **8.02E-09** | **9.95E-09** | -1.17 | (-1.22--1.12) | **2.24E-270** | **3.89E-270** | 0.76 | (0.7-0.81) | **1.16E-139** | **2.25E-139** | 0.32 | (0.23-0.4) | **1.28E-13** | **8.52E-13** |
| 202 | SM(40:3) (a) | -0.69 | (-0.74--0.65) | **5.43E-162** | **2.19E-161** | -1.53 | (-1.58--1.47) | **0.00E+00** | **0.00E+00** | 0.39 | (0.34-0.45) | **6.70E-41** | **8.81E-41** | 0.39 | (0.29-0.48) | **2.45E-15** | **2.03E-14** |
| 203 | SM(40:3) (b) | -0.44 | (-0.49--0.39) | **1.73E-63** | **3.31E-63** | -1.93 | (-1.99--1.87) | **0.00E+00** | **0.00E+00** | 1.23 | (1.16-1.29) | **1.48E-197** | **3.83E-197** | 0.26 | (0.15-0.37) | **6.31E-06** | **1.56E-05** |
| 204 | SM(41:0) | -0.46 | (-0.5--0.42) | **4.49E-98** | **1.10E-97** | -1.90 | (-1.97--1.83) | **0.00E+00** | **0.00E+00** | 1.47 | (1.39-1.55) | **6.63E-200** | **1.75E-199** | -0.10 | (-0.19--0.01) | **2.84E-02** | **4.06E-02** |
| 205 | SM(41:1) | -0.42 | (-0.46--0.38) | **1.03E-93** | **2.40E-93** | -3.27 | (-3.33--3.22) | **0.00E+00** | **0.00E+00** | 2.84 | (2.78-2.9) | **0.00E+00** | **0.00E+00** | -0.03 | (-0.12-0.05) | 4.38E-01 | 4.99E-01 |
| 206 | SM(d17:1/24:1) | -0.36 | (-0.4--0.32) | **3.19E-63** | **6.08E-63** | -1.78 | (-1.83--1.73) | **0.00E+00** | **0.00E+00** | 1.24 | (1.19-1.3) | **4.57E-262** | **1.65E-261** | 0.17 | (0.08-0.26) | **2.53E-04** | **4.95E-04** |
| 207 | SM(d18:2/23:0) | -0.19 | (-0.23--0.15) | **1.15E-23** | **1.65E-23** | -2.92 | (-2.98--2.87) | **0.00E+00** | **0.00E+00** | 2.59 | (2.53-2.65) | **0.00E+00** | **0.00E+00** | 0.18 | (0.11-0.26) | **5.79E-06** | **1.45E-05** |
| 208 | SM(d18:1/24:0) | -0.20 | (-0.24--0.16) | **1.86E-22** | **2.64E-22** | -1.22 | (-1.27--1.17) | **1.26E-281** | **2.24E-281** | 0.96 | (0.91-1.01) | **4.06E-198** | **1.05E-197** | 0.09 | (0-0.17) | **3.96E-02** | 5.51E-02 |
| 209 | SM(d18:1/24:1) | -0.55 | (-0.6--0.51) | **9.06E-102** | **2.32E-101** | -1.89 | (-1.95--1.84) | **0.00E+00** | **0.00E+00** | 1.09 | (1.04-1.15) | **5.54E-217** | **1.60E-216** | 0.30 | (0.21-0.4) | **1.29E-09** | **5.19E-09** |
| 210 | SM(d18:2/24:0) | -0.05 | (-0.09--0.02) | **2.79E-03** | **3.13E-03** | -1.02 | (-1.07--0.98) | **4.71E-243** | **7.49E-243** | 0.78 | (0.73-0.82) | **4.03E-157** | **8.22E-157** | 0.32 | (0.25-0.39) | **9.02E-17** | **9.21E-16** |
| 211 | SM(43:1) | 0.05 | (0.01-0.09) | **2.68E-02** | **2.89E-02** | -0.87 | (-0.94--0.81) | **4.04E-123** | **5.10E-123** | 1.03 | (0.96-1.1) | **2.25E-138** | **4.35E-138** | -0.09 | (-0.18-0) | 6.13E-02 | 8.18E-02 |
| 212 | SM(44:1) | 0.21 | (0.17-0.24) | **9.73E-28** | **1.43E-27** | 0.59 | (0.54-0.63) | **3.58E-109** | **4.40E-109** | -0.32 | (-0.37--0.28) | **3.64E-39** | **4.76E-39** | 0.04 | (-0.04-0.11) | 3.19E-01 | 3.76E-01 |
| 213 | SM(44:2) | 0.01 | (-0.03-0.06) | 5.36E-01 | 5.48E-01 | 0.19 | (0.14-0.24) | **1.71E-12** | **1.84E-12** | -0.28 | (-0.33--0.23) | **1.09E-28** | **1.35E-28** | 0.18 | (0.09-0.27) | **5.10E-05** | **1.12E-04** |

| | | | | | | | | | | | | | | | | | |
|---|---|---|---|---|---|---|---|---|---|---|---|---|---|---|---|---|---|
| 214 | SM(44:3) (a) | 0.00 | (-0.04-0.04) | 8.80E-01 | 8.81E-01 | 0.25 | (0.2-0.3) | **1.73E-22** | **1.88E-22** | -0.50 | (-0.54--0.45) | **1.31E-84** | **2.07E-84** | 0.39 | (0.31-0.47) | **4.89E-20** | **7.34E-19** |
| 215 | LPC(14:0) [sn2] | 0.36 | (0.29-0.43) | **6.81E-24** | **9.79E-24** | 0.44 | (0.37-0.5) | **1.22E-36** | **1.37E-36** | -0.31 | (-0.37--0.24) | **1.53E-21** | **1.80E-21** | 0.11 | (-0.05-0.26) | 1.74E-01 | 2.14E-01 |
| 216 | LPC(14:0) [sn1] | 0.39 | (0.32-0.45) | **7.58E-28** | **1.12E-27** | 0.33 | (0.27-0.4) | **8.14E-24** | **8.90E-24** | -0.15 | (-0.21--0.09) | **4.81E-07** | **5.15E-07** | 0.03 | (-0.12-0.18) | 7.05E-01 | 7.50E-01 |
| 217 | LPC(15:0) [sn2] | 0.59 | (0.54-0.65) | **1.37E-88** | **3.07E-88** | 0.06 | (0.01-0.12) | **3.01E-02** | **3.06E-02** | 0.26 | (0.2-0.32) | **5.78E-16** | **6.64E-16** | 0.25 | (0.12-0.38) | **1.67E-04** | **3.36E-04** |
| 218 | LPC(15:0) [sn1] | 0.60 | (0.56-0.65) | **6.72E-106** | **1.78E-105** | -0.07 | (-0.13--0.02) | **5.90E-03** | **6.10E-03** | 0.39 | (0.33-0.45) | **1.69E-36** | **2.18E-36** | 0.30 | (0.18-0.42) | **1.20E-06** | **3.37E-06** |
| 219 | LPC(16:0) [sn2] | 0.59 | (0.55-0.63) | **9.88E-151** | **3.51E-150** | 0.06 | (0.02-0.11) | **3.59E-03** | **3.73E-03** | 0.22 | (0.17-0.26) | **1.21E-18** | **1.41E-18** | 0.49 | (0.39-0.58) | **4.01E-23** | **9.17E-22** |
| 220 | LPC(16:0) [sn1] | 0.70 | (0.66-0.75) | **2.46E-151** | **8.94E-151** | -0.04 | (-0.09-0.02) | 1.68E-01 | 1.70E-01 | 0.36 | (0.3-0.42) | **6.75E-35** | **8.64E-35** | 0.59 | (0.48-0.7) | **4.60E-24** | **1.47E-22** |
| 221 | LPC(16:1) [sn2] | 0.83 | (0.77-0.89) | **1.31E-133** | **4.16E-133** | 1.93 | (1.86-1.99) | **0.00E+00** | **0.00E+00** | -1.43 | (-1.49--1.37) | **3.48E-261** | **1.24E-260** | 0.28 | (0.16-0.39) | **4.60E-06** | **1.18E-05** |
| 222 | LPC(16:1) [sn1] | 0.86 | (0.81-0.91) | **1.87E-169** | **8.54E-169** | 1.95 | (1.89-2.01) | **0.00E+00** | **0.00E+00** | -1.39 | (-1.45--1.34) | **5.15E-267** | **1.93E-266** | 0.25 | (0.14-0.36) | **1.16E-05** | **2.80E-05** |
| 223 | LPC(15-MHDA) [sn2] | 0.94 | (0.87-1) | **8.31E-130** | **2.56E-129** | 0.43 | (0.35-0.51) | **1.78E-25** | **1.96E-25** | 0.23 | (0.14-0.31) | **4.40E-07** | **4.73E-07** | 0.36 | (0.19-0.52) | **2.22E-05** | **5.25E-05** |
| 224 | LPC(15-MHDA) [sn1] / LPC(17:0) [sn2] | 0.93 | (0.87-0.99) | **1.62E-167** | **7.02E-167** | 0.00 | (-0.06-0.07) | 8.88E-01 | 8.88E-01 | 0.66 | (0.59-0.73) | **1.79E-66** | **2.62E-66** | 0.41 | (0.27-0.55) | **5.50E-09** | **2.00E-08** |
| 225 | LPC(17:0) [sn1] | 0.88 | (0.83-0.92) | **1.61E-213** | **9.81E-213** | -0.37 | (-0.42--0.33) | **5.70E-48** | **6.50E-48** | 0.97 | (0.92-1.03) | **3.05E-182** | **7.18E-182** | 0.46 | (0.35-0.57) | **2.88E-15** | **2.34E-14** |
| 226 | LPC(17:1) [sn1] (a) / LPC(17:1) [sn2] (b) | 0.85 | (0.8-0.9) | **8.19E-166** | **3.45E-165** | 1.09 | (1.03-1.15) | **4.73E-209** | **7.03E-209** | -0.53 | (-0.59--0.47) | **3.07E-58** | **4.34E-58** | 0.39 | (0.26-0.52) | **1.63E-09** | **6.40E-09** |
| 227 | LPC(18:0) [sn2] | 1.18 | (1.13-1.22) | **0.00E+00** | **0.00E+00** | 0.02 | (-0.02-0.07) | 2.93E-01 | 2.95E-01 | 0.97 | (0.92-1.02) | **6.74E-205** | **1.82E-204** | 0.31 | (0.21-0.42) | **5.11E-09** | **1.87E-08** |
| 228 | LPC(18:0) [sn1] | 1.20 | (1.16-1.24) | **0.00E+00** | **0.00E+00** | -0.03 | (-0.08-0.01) | 1.46E-01 | 1.48E-01 | 1.06 | (1.01-1.12) | **2.64E-210** | **7.42E-210** | 0.31 | (0.2-0.43) | **5.14E-08** | **1.64E-07** |
| 229 | LPC(18:1) [sn2] | 1.05 | (1-1.1) | **2.09E-240** | **1.54E-239** | 0.74 | (0.68-0.79) | **1.62E-125** | **2.07E-125** | 0.17 | (0.11-0.23) | **1.48E-08** | **1.60E-08** | 0.48 | (0.37-0.6) | **1.92E-15** | **1.61E-14** |
| 230 | LPC(18:1) [sn1] | 1.12 | (1.07-1.16) | **9.44E-308** | **1.30E-306** | 0.88 | (0.83-0.93) | **2.73E-192** | **3.95E-192** | 0.20 | (0.15-0.26) | **4.68E-14** | **5.25E-14** | 0.36 | (0.25-0.46) | **2.93E-11** | **1.44E-10** |
| 231 | LPC(18:2) [sn2] | 1.24 | (1.19-1.29) | **0.00E+00** | **0.00E+00** | 0.34 | (0.28-0.39) | **1.67E-33** | **1.86E-33** | 0.84 | (0.78-0.89) | **7.57E-137** | **1.44E-136** | 0.48 | (0.37-0.59) | **1.90E-17** | **2.17E-16** |
| 232 | LPC(18:2) [sn1] | 1.67 | (1.62-1.72) | **0.00E+00** | **0.00E+00** | 0.83 | (0.77-0.88) | **2.22E-139** | **2.90E-139** | 0.77 | (0.72-0.83) | **7.76E-114** | **1.37E-113** | 0.51 | (0.39-0.62) | **2.81E-16** | **2.60E-15** |
| 233 | LPC(18:3) [sn2] (a) | 0.71 | (0.65-0.78) | **9.31E-90** | **2.11E-89** | -0.52 | (-0.58--0.45) | **8.29E-51** | **9.50E-51** | 1.29 | (1.22-1.35) | **4.19E-210** | **1.16E-209** | 0.22 | (0.08-0.37) | **2.94E-03** | **4.93E-03** |
| 234 | LPC(18:3) [sn1] (a)/LPC(18:3) [sn2] (b) | 1.24 | (1.18-1.31) | **7.95E-205** | **4.65E-204** | 1.06 | (0.98-1.13) | **2.76E-140** | **3.62E-140** | 0.29 | (0.21-0.36) | **1.17E-13** | **1.30E-13** | 0.15 | (0-0.3) | 5.76E-02 | 7.73E-02 |
| 235 | LPC(18:3) (a) [sn1] [104_sn1] | 1.31 | (1.25-1.38) | **3.01E-214** | **1.88E-213** | 1.10 | (1.03-1.18) | **1.57E-146** | **2.07E-146** | 0.31 | (0.24-0.38) | **6.17E-16** | **7.05E-16** | 0.18 | (0.02-0.33) | **2.42E-02** | **3.51E-02** |
| 236 | LPC(19:0) [sn1] (a) / LPC(19:0) [sn2] (b) | 0.83 | (0.78-0.87) | **5.18E-199** | **2.89E-198** | -0.40 | (-0.45--0.34) | **1.31E-40** | **1.48E-40** | 1.04 | (0.97-1.1) | **5.08E-164** | **1.08E-163** | 0.45 | (0.34-0.56) | **4.00E-15** | **3.14E-14** |
| 237 | LPC(20:0) [sn1] | 0.96 | (0.92-1) | **1.17E-266** | **1.19E-265** | -0.32 | (-0.36--0.28) | **2.44E-44** | **2.77E-44** | 1.18 | (1.13-1.23) | **6.10E-283** | **2.48E-282** | 0.42 | (0.32-0.52) | **6.50E-16** | **5.89E-15** |
| 238 | LPC(20:1) [sn2] | 0.65 | (0.6-0.69) | **4.59E-139** | **1.52E-138** | -0.29 | (-0.34--0.25) | **4.01E-34** | **4.46E-34** | 0.79 | (0.74-0.84) | **1.31E-147** | **2.58E-147** | 0.42 | (0.31-0.53) | **6.68E-14** | **4.65E-13** |
| 239 | LPC(20:1) [sn1] | 0.77 | (0.73-0.82) | **8.69E-177** | **4.13E-176** | -0.23 | (-0.28--0.19) | **2.18E-23** | **2.37E-23** | 0.86 | (0.81-0.91) | **4.35E-178** | **1.01E-177** | 0.47 | (0.36-0.57) | **6.60E-17** | **7.04E-16** |
| 240 | LPC(20:2) [sn2] | 0.24 | (0.19-0.28) | **3.38E-24** | **4.87E-24** | 0.45 | (0.4-0.51) | **1.24E-51** | **1.43E-51** | -0.31 | (-0.37--0.25) | **1.72E-24** | **2.09E-24** | 0.38 | (0.28-0.49) | **3.77E-13** | **2.32E-12** |
| 241 | LPC(20:2) [sn1] | 0.60 | (0.56-0.64) | **1.12E-132** | **3.54E-132** | 0.93 | (0.88-0.98) | **1.02E-192** | **1.48E-192** | -0.42 | (-0.47--0.36) | **7.42E-45** | **9.89E-45** | 0.38 | (0.28-0.48) | **1.64E-13** | **1.08E-12** |
| 242 | LPC(20:3) [sn2] | 0.37 | (0.31-0.42) | **4.77E-34** | **7.45E-34** | 1.65 | (1.59-1.71) | **0.00E+00** | **0.00E+00** | -1.33 | (-1.39--1.27) | **2.58E-237** | **8.21E-237** | 0.05 | (-0.08-0.17) | 4.68E-01 | 5.28E-01 |
| 243 | LPC(20:3) [sn1] | 0.88 | (0.82-0.94) | **1.48E-142** | **4.99E-142** | 2.40 | (2.33-2.46) | **0.00E+00** | **0.00E+00** | -1.54 | (-1.6--1.47) | **1.91E-264** | **7.07E-264** | 0.08 | (-0.05-0.21) | 2.35E-01 | 2.83E-01 |
| 244 | LPC(20:3) [104_sn1] | 0.93 | (0.87-0.99) | **3.80E-151** | **1.37E-150** | 2.43 | (2.37-2.5) | **0.00E+00** | **0.00E+00** | -1.53 | (-1.59--1.46) | **2.10E-260** | **7.35E-260** | 0.07 | (-0.06-0.2) | 2.76E-01 | 3.28E-01 |
| 245 | LPC(20:4) [sn2] | 0.87 | (0.83-0.92) | **8.47E-205** | **4.90E-204** | 2.43 | (2.37-2.48) | **0.00E+00** | **0.00E+00** | -1.79 | (-1.85--1.74) | **0.00E+00** | **0.00E+00** | 0.41 | (0.3-0.51) | **4.58E-14** | **3.23E-13** |
| 246 | LPC(20:4) [sn1] | 1.34 | (1.29-1.39) | **0.00E+00** | **0.00E+00** | 3.13 | (3.08-3.19) | **0.00E+00** | **0.00E+00** | -2.05 | (-2.11--1.99) | **0.00E+00** | **0.00E+00** | 0.49 | (0.37-0.6) | **1.64E-16** | **1.61E-15** |
| 247 | LPC(20:5) [sn2] | 1.19 | (1.1-1.29) | **4.81E-105** | **1.27E-104** | 0.72 | (0.61-0.83) | **5.81E-37** | **6.52E-37** | -0.28 | (-0.38--0.17) | **1.67E-07** | **1.80E-07** | 0.93 | (0.72-1.14) | **1.35E-17** | **1.58E-16** |
| 248 | LPC(20:5) [sn1] | 1.54 | (1.44-1.64) | **1.67E-153** | **6.23E-153** | 1.24 | (1.13-1.35) | **6.00E-95** | **7.18E-95** | -0.45 | (-0.55--0.35) | **6.00E-17** | **6.94E-17** | 0.93 | (0.72-1.15) | **3.27E-17** | **3.65E-16** |
| 249 | LPC(22:4) [sn2] | 0.23 | (0.17-0.28) | **3.40E-16** | **4.58E-16** | 0.95 | (0.89-1.02) | **1.65E-155** | **2.23E-155** | -0.47 | (-0.53--0.41) | **1.28E-48** | **1.72E-48** | -0.11 | (-0.23-0.01) | 7.31E-02 | 9.64E-02 |
| 250 | LPC(22:4) [sn1] | 0.63 | (0.58-0.68) | **9.98E-99** | **2.46E-98** | 1.76 | (1.7-1.82) | **0.00E+00** | **0.00E+00** | -0.91 | (-0.97--0.85) | **4.93E-147** | **9.65E-147** | -0.04 | (-0.16-0.09) | 5.58E-01 | 6.20E-01 |
| 251 | LPC(22:5) [sn2] (n3) | 0.90 | (0.84-0.95) | **1.46E-167** | **6.39E-167** | 0.49 | (0.43-0.56) | **1.81E-43** | **2.05E-43** | 0.26 | (0.19-0.32) | **5.60E-14** | **6.25E-14** | 0.34 | (0.22-0.46) | **2.17E-08** | **7.39E-08** |
| 252 | LPC(22:5) [sn1] (n3)/LPC(22:5) [sn2] (n6) | 1.07 | (1.02-1.13) | **1.05E-226** | **7.13E-226** | 0.79 | (0.73-0.85) | **1.50E-118** | **1.88E-118** | 0.28 | (0.22-0.34) | **5.56E-20** | **6.51E-20** | 0.22 | (0.11-0.34) | **2.19E-04** | **4.29E-04** |
| 253 | LPC(22:5) [sn1] (n6) | 0.67 | (0.6-0.73) | **7.86E-78** | **1.65E-77** | 1.54 | (1.46-1.63) | **2.54E-206** | **3.76E-206** | -0.51 | (-0.59--0.43) | **8.39E-33** | **1.06E-32** | -0.10 | (-0.24-0.04) | 1.63E-01 | 2.01E-01 |
| 254 | LPC(22:5) (n3) [sn1] [104_sn1] | 1.37 | (1.31-1.42) | **1.35E-268** | **1.41E-267** | 0.79 | (0.72-0.86) | **2.21E-95** | **2.66E-95** | 0.48 | (0.41-0.54) | **2.73E-42** | **3.61E-42** | 0.34 | (0.21-0.46) | **1.36E-07** | **4.17E-07** |
| 255 | LPC(22:6) [sn2] | 0.44 | (0.38-0.5) | **2.36E-50** | **4.15E-50** | 0.74 | (0.68-0.8) | **8.25E-106** | **1.01E-105** | -0.53 | (-0.6--0.47) | **1.06E-56** | **1.47E-56** | 0.46 | (0.33-0.58) | **2.08E-12** | **1.18E-11** |
| 256 | LPC(22:6) [sn1] | 0.92 | (0.86-0.98) | **1.99E-159** | **7.78E-159** | 1.43 | (1.36-1.49) | **8.90E-262** | **1.48E-261** | -0.71 | (-0.77--0.64) | **2.11E-86** | **3.34E-86** | 0.46 | (0.33-0.59) | **1.15E-11** | **5.95E-11** |
| 257 | LPC(O-16:0) | 1.07 | (1.03-1.11) | **0.00E+00** | **0.00E+00** | 0.91 | (0.86-0.95) | **9.99E-233** | **1.56E-232** | -0.15 | (-0.2--0.1) | **1.17E-08** | **1.27E-08** | 0.53 | (0.43-0.64) | **2.18E-22** | **4.76E-21** |
| 258 | LPC(O-18:0) | 0.90 | (0.86-0.95) | **3.94E-200** | **2.22E-199** | -0.20 | (-0.26--0.15) | **8.06E-13** | **8.70E-13** | 0.83 | (0.77-0.89) | **1.88E-123** | **3.47E-123** | 0.44 | (0.32-0.56) | **1.92E-12** | **1.11E-11** |
| 259 | LPC(O-18:1) | 0.98 | (0.94-1.02) | **4.01E-285** | **4.70E-284** | 0.60 | (0.55-0.64) | **5.89E-123** | **7.43E-123** | 0.20 | (0.15-0.25) | **9.84E-14** | **1.10E-13** | 0.36 | (0.26-0.46) | **1.07E-11** | **5.58E-11** |
| 260 | LPC(O-20:0) | 0.94 | (0.89-0.98) | **7.14E-218** | **4.57E-217** | -0.09 | (-0.16--0.02) | **1.21E-02** | **1.24E-02** | 0.80 | (0.73-0.88) | **8.89E-82** | **1.37E-81** | 0.47 | (0.35-0.59) | **4.06E-15** | **3.14E-14** |
| 261 | LPC(O-20:1) | 0.92 | (0.85-0.99) | **6.51E-132** | **2.04E-131** | 0.10 | (0.02-0.19) | **1.41E-02** | **1.44E-02** | 0.34 | (0.26-0.42) | **8.46E-17** | **9.76E-17** | 0.74 | (0.61-0.88) | **5.17E-26** | **2.76E-24** |
| 262 | LPC(O-22:0) | 0.47 | (0.43-0.52) | **9.97E-94** | **2.32E-93** | -0.13 | (-0.17--0.08) | **6.07E-07** | **6.42E-07** | 0.40 | (0.36-0.45) | **7.91E-63** | **1.14E-62** | 0.44 | (0.36-0.53) | **5.49E-24** | **1.65E-22** |
| 263 | LPC(O-22:1) | 0.70 | (0.65-0.75) | **6.78E-138** | **2.21E-137** | -0.09 | (-0.14--0.03) | **1.31E-03** | **1.37E-03** | 0.59 | (0.53-0.64) | **2.31E-82** | **3.57E-82** | 0.56 | (0.45-0.67) | **3.09E-22** | **6.45E-21** |
| 264 | LPC(O-24:0) | 0.17 | (0.13-0.22) | **2.48E-12** | **3.21E-12** | -0.86 | (-0.91--0.8) | **1.74E-147** | **2.31E-147** | 0.83 | (0.78-0.88) | **4.34E-169** | **9.61E-169** | 0.43 | (0.34-0.52) | **1.07E-20** | **1.71E-19** |
| 265 | LPC(O-24:1) | 0.29 | (0.24-0.33) | **2.52E-31** | **3.80E-31** | -0.74 | (-0.8--0.69) | **1.01E-127** | **1.31E-127** | 0.84 | (0.79-0.89) | **1.83E-168** | **4.01E-168** | 0.44 | (0.35-0.54) | **1.31E-19** | **1.84E-18** |
| 266 | LPC(O-24:2) | 0.48 | (0.43-0.53) | **2.81E-66** | **5.48E-66** | 0.13 | (0.08-0.19) | **3.54E-06** | **3.73E-06** | 0.07 | (0.02-0.12) | **9.02E-03** | **9.35E-03** | 0.61 | (0.5-0.72) | **7.06E-26** | **3.39E-24** |
| 267 | LPC(P-16:0) | 1.38 | (1.34-1.42) | **0.00E+00** | **0.00E+00** | 0.90 | (0.85-0.94) | **2.45E-221** | **3.75E-221** | 0.05 | (0-0.1) | 6.64E-02 | 6.79E-02 | 0.63 | (0.52-0.73) | **1.44E-30** | **1.39E-28** |
| 268 | LPC(P-18:0) | 0.98 | (0.94-1.02) | **1.40E-281** | **1.61E-280** | 0.59 | (0.54-0.63) | **1.81E-119** | **2.28E-119** | 0.21 | (0.16-0.26) | **3.68E-15** | **4.14E-15** | 0.35 | (0.25-0.45) | **2.20E-11** | **1.10E-10** |

| | | | | | | | | | | | | | | | | | |
|---|---|---|---|---|---|---|---|---|---|---|---|---|---|---|---|---|---|
| 269 | LPC(P-18:1) | 1.23 | (1.18-1.28) | 5.32E-291 | 6.73E-290 | -0.07 | (-0.12--0.01) | 2.53E-02 | 2.59E-02 | 0.79 | (0.72-0.85) | 7.64E-106 | 1.29E-105 | 0.64 | (0.52-0.76) | 3.04E-25 | 1.22E-23 |
| 270 | LPC(P-20:0) | 0.86 | (0.81-0.91) | 3.90E-169 | 1.75E-168 | -0.07 | (-0.13--0.01) | 1.35E-02 | 1.39E-02 | 0.81 | (0.75-0.87) | 2.92E-117 | 5.23E-117 | 0.43 | (0.31-0.55) | 7.83E-12 | 4.17E-11 |
| 271 | LPE(16:0) [sn2] | -0.19 | (-0.24--0.14) | 9.60E-15 | 1.28E-14 | -0.05 | (-0.1-0) | 6.50E-02 | 6.61E-02 | -0.53 | (-0.58--0.47) | 2.70E-69 | 4.01E-69 | 0.72 | (0.6-0.83) | 9.17E-34 | 2.20E-31 |
| 272 | LPE(16:0) [sn1] | -0.26 | (-0.3--0.21) | 2.90E-27 | 4.23E-27 | -0.14 | (-0.19--0.09) | 4.47E-08 | 4.77E-08 | -0.51 | (-0.56--0.46) | 2.19E-72 | 3.32E-72 | 0.71 | (0.6-0.82) | 1.55E-36 | 7.46E-34 |
| 273 | LPE(18:0) [sn2] | 0.18 | (0.14-0.22) | 9.26E-17 | 1.26E-16 | -0.26 | (-0.3--0.21) | 2.06E-25 | 2.26E-25 | 0.15 | (0.1-0.19) | 2.37E-09 | 2.59E-09 | 0.51 | (0.42-0.61) | 2.15E-24 | 7.36E-23 |
| 274 | LPE(18:0) [sn1] | 0.13 | (0.08-0.17) | 3.87E-09 | 4.82E-09 | -0.36 | (-0.4--0.31) | 7.99E-50 | 9.13E-50 | 0.19 | (0.14-0.24) | 4.76E-14 | 5.32E-14 | 0.53 | (0.43-0.63) | 1.66E-24 | 6.14E-23 |
| 275 | LPE(18:1) [sn2] | 0.01 | (-0.04-0.07) | 6.18E-01 | 6.27E-01 | -0.23 | (-0.28--0.17) | 1.40E-15 | 1.51E-15 | 0.21 | (0.15-0.27) | 2.95E-11 | 3.26E-11 | 0.25 | (0.12-0.38) | 1.84E-04 | 3.67E-04 |
| 276 | LPE(18:1) [sn1] | 0.05 | (-0.01-0.1) | 7.61E-02 | 8.08E-02 | -0.28 | (-0.34--0.23) | 8.21E-24 | 8.96E-24 | 0.31 | (0.25-0.37) | 8.46E-23 | 1.01E-22 | 0.25 | (0.11-0.38) | 2.98E-04 | 5.75E-04 |
| 277 | LPE(18:2) [sn2] | 0.72 | (0.66-0.78) | 6.14E-100 | 1.53E-99 | -0.16 | (-0.22--0.09) | 1.09E-05 | 1.15E-05 | 0.85 | (0.78-0.92) | 2.37E-99 | 3.87E-99 | 0.29 | (0.15-0.43) | 3.35E-05 | 7.78E-05 |
| 278 | LPE(18:2) [sn1] | 0.90 | (0.84-0.96) | 6.95E-145 | 2.45E-144 | 0.08 | (0.01-0.14) | 2.74E-02 | 2.80E-02 | 0.82 | (0.75-0.89) | 8.28E-100 | 1.36E-99 | 0.27 | (0.13-0.4) | 1.11E-04 | 2.27E-04 |
| 279 | LPE(20:4) [sn1] | 1.04 | (1-1.09) | 2.02E-258 | 1.80E-257 | 2.28 | (2.22-2.34) | 0.00E+00 | 0.00E+00 | -1.35 | (-1.4--1.29) | 1.83E-270 | 6.92E-270 | 0.31 | (0.21-0.4) | 2.45E-09 | 9.27E-09 |
| 280 | LPE(22:6) [sn2] | 0.34 | (0.29-0.4) | 2.66E-35 | 4.20E-35 | 1.57 | (1.51-1.63) | 0.00E+00 | 0.00E+00 | -1.66 | (-1.72--1.6) | 7.89E-308 | 3.75E-307 | 0.57 | (0.45-0.69) | 5.75E-20 | 8.37E-19 |
| 281 | LPE(22:6) [sn1] | 0.64 | (0.58-0.69) | 3.21E-100 | 8.03E-100 | 1.93 | (1.87-1.99) | 0.00E+00 | 0.00E+00 | -1.72 | (-1.78--1.66) | 0.00E+00 | 0.00E+00 | 0.60 | (0.49-0.72) | 1.26E-23 | 3.36E-22 |
| 282 | PC(28:0) | -0.83 | (-0.93--0.73) | 1.54E-54 | 2.79E-54 | -1.90 | (-1.99--1.81) | 1.20E-229 | 1.86E-229 | 1.03 | (0.96-1.11) | 7.60E-118 | 1.37E-117 | 0.02 | (-0.19-0.23) | 8.68E-01 | 8.96E-01 |
| 283 | PC(14:0_16:0) | -0.99 | (-1.06--0.93) | 1.11E-151 | 4.05E-151 | -1.70 | (-1.77--1.63) | 3.48E-279 | 6.09E-279 | 0.58 | (0.52-0.65) | 5.72E-59 | 8.15E-59 | 0.18 | (0.04-0.31) | 1.16E-02 | 1.80E-02 |
| 284 | PC(31:0) (a) | -0.87 | (-0.97--0.78) | 3.10E-64 | 5.95E-64 | -1.94 | (-2.04--1.84) | 1.04E-210 | 1.56E-210 | 1.26 | (1.16-1.37) | 1.02E-100 | 1.68E-100 | -0.48 | (-0.71--0.26) | 2.31E-05 | 5.44E-05 |
| 285 | PC(31:0) (b) | -0.65 | (-0.71--0.6) | 3.44E-94 | 8.17E-94 | -1.27 | (-1.35--1.19) | 1.06E-158 | 1.44E-158 | 0.56 | (0.47-0.64) | 2.19E-33 | 2.78E-33 | -0.01 | (-0.15-0.13) | 8.75E-01 | 9.01E-01 |
| 286 | PC(31:1) | -1.10 | (-1.18--1.02) | 8.99E-126 | 2.68E-125 | -1.66 | (-1.74--1.57) | 2.76E-209 | 4.11E-209 | 0.35 | (0.27-0.44) | 5.83E-16 | 6.68E-16 | -0.07 | (-0.24-0.11) | 4.47E-01 | 5.07E-01 |
| 287 | PC(16:0_16:0) | -0.71 | (-0.76--0.67) | 1.73E-162 | 7.02E-162 | -0.95 | (-1.01--0.89) | 5.49E-160 | 7.54E-160 | 0.06 | (0-0.12) | 3.55E-02 | 3.65E-02 | 0.27 | (0.18-0.36) | 1.67E-08 | 5.81E-08 |
| 288 | PC(32:1) | -1.19 | (-1.26--1.12) | 4.60E-164 | 1.92E-163 | -1.12 | (-1.21--1.04) | 2.96E-127 | 3.80E-127 | -0.17 | (-0.24--0.09) | 1.57E-05 | 1.66E-05 | -0.14 | (-0.29-0.01) | 6.74E-02 | 8.96E-02 |
| 289 | PC(32:2) | -1.07 | (-1.14--1.01) | 3.14E-159 | 1.21E-158 | -3.48 | (-3.55--3.42) | 0.00E+00 | 0.00E+00 | 2.29 | (2.23-2.36) | 0.00E+00 | 0.00E+00 | 0.02 | (-0.13-0.18) | 7.56E-01 | 7.94E-01 |
| 290 | PC(33:0) (a) | -0.80 | (-0.87--0.73) | 6.49E-94 | 1.52E-93 | -1.18 | (-1.26--1.09) | 3.97E-127 | 5.09E-127 | 0.17 | (0.08-0.27) | 2.79E-04 | 2.94E-04 | -0.01 | (-0.18-0.16) | 9.47E-01 | 9.57E-01 |
| 291 | PC(33:0) (b) | -0.37 | (-0.42--0.32) | 1.09E-40 | 1.78E-40 | -1.54 | (-1.61--1.46) | 7.08E-248 | 1.14E-247 | 0.97 | (0.9-1.05) | 1.84E-109 | 3.16E-109 | 0.17 | (0.05-0.29) | 7.28E-03 | 1.16E-02 |
| 292 | PC(33:1) | -1.16 | (-1.22--1.09) | 4.15E-189 | 2.14E-188 | -2.32 | (-2.39--2.25) | 0.00E+00 | 0.00E+00 | 1.08 | (1.01-1.15) | 4.74E-153 | 9.59E-153 | 0.02 | (-0.12-0.16) | 7.73E-01 | 8.10E-01 |
| 293 | PC(33:2) | -0.83 | (-0.88--0.77) | 2.18E-140 | 7.32E-140 | -4.20 | (-4.27--4.14) | 0.00E+00 | 0.00E+00 | 3.13 | (3.06-3.2) | 0.00E+00 | 0.00E+00 | 0.28 | (0.15-0.41) | 1.67E-05 | 3.99E-05 |
| 294 | PC(16:0_18:0) | 0.10 | (0.06-0.14) | 5.36E-07 | 6.53E-07 | -0.81 | (-0.87--0.76) | 6.94E-163 | 9.58E-163 | 0.76 | (0.71-0.81) | 2.58E-136 | 4.88E-136 | 0.23 | (0.15-0.32) | 1.34E-07 | 4.11E-07 |
| 295 | PC(16:0_18:1) | -1.07 | (-1.12--1.01) | 2.88E-216 | 1.82E-215 | -2.34 | (-2.41--2.28) | 0.00E+00 | 0.00E+00 | 1.24 | (1.17-1.3) | 3.40E-199 | 8.92E-199 | 0.10 | (-0.03-0.22) | 1.25E-01 | 1.59E-01 |
| 296 | PC(16:0_18:2) | -0.57 | (-0.6--0.53) | 3.11E-180 | 1.49E-179 | -3.50 | (-3.56--3.43) | 0.00E+00 | 0.00E+00 | 2.79 | (2.72-2.86) | 0.00E+00 | 0.00E+00 | 0.24 | (0.16-0.33) | 2.70E-08 | 9.06E-08 |
| 297 | PC(16:1_18:2) | -0.69 | (-0.75--0.64) | 2.22E-110 | 5.98E-110 | -2.96 | (-3.03--2.9) | 0.00E+00 | 0.00E+00 | 1.93 | (1.87-2) | 0.00E+00 | 0.00E+00 | 0.31 | (0.19-0.43) | 1.83E-07 | 5.52E-07 |
| 298 | PC(16:0_18:3) (a) | -1.06 | (-1.12--1) | 2.61E-184 | 1.28E-183 | -3.64 | (-3.71--3.56) | 0.00E+00 | 0.00E+00 | 2.52 | (2.45-2.6) | 0.00E+00 | 0.00E+00 | 0.13 | (0-0.27) | 4.89E-02 | 6.63E-02 |
| 299 | PC(14:0_20:4) | -0.97 | (-1.04--0.9) | 1.64E-119 | 4.70E-119 | -2.97 | (-3.05--2.9) | 0.00E+00 | 0.00E+00 | 1.76 | (1.69-1.84) | 3.94E-273 | 1.51E-272 | 0.11 | (-0.05-0.28) | 1.78E-01 | 2.18E-01 |
| 300 | PC(34:5) | -0.53 | (-0.64--0.43) | 4.22E-21 | 5.92E-21 | -3.47 | (-3.58--3.36) | 0.00E+00 | 0.00E+00 | 2.21 | (2.11-2.32) | 5.03E-234 | 1.57E-233 | 0.62 | (0.38-0.85) | 3.28E-07 | 9.66E-07 |
| 301 | PC(15-MHDA_18:1) | -0.53 | (-0.6--0.46) | 8.41E-45 | 1.43E-44 | -2.05 | (-2.14--1.97) | 4.30E-295 | 7.85E-295 | 1.51 | (1.43-1.6) | 3.03E-174 | 6.92E-174 | -0.01 | (-0.17-0.16) | 9.27E-01 | 9.39E-01 |
| 302 | PC(17:0_18:1) | -0.57 | (-0.62--0.52) | 9.17E-104 | 2.39E-103 | -1.91 | (-1.97--1.86) | 0.00E+00 | 0.00E+00 | 1.30 | (1.25-1.36) | 5.03E-258 | 1.74E-257 | 0.06 | (-0.05-0.17) | 2.65E-01 | 3.17E-01 |
| 303 | PC(15-MHDA_18:2) | -0.68 | (-0.75--0.61) | 1.49E-68 | 2.96E-68 | -3.32 | (-3.4--3.24) | 0.00E+00 | 0.00E+00 | 2.51 | (2.43-2.6) | 0.00E+00 | 0.00E+00 | 0.20 | (0.04-0.36) | 1.35E-02 | 2.05E-02 |
| 304 | PC(17:0_18:2) | -0.46 | (-0.51--0.41) | 1.27E-69 | 2.54E-69 | -3.32 | (-3.38--3.26) | 0.00E+00 | 0.00E+00 | 2.65 | (2.58-2.71) | 0.00E+00 | 0.00E+00 | 0.33 | (0.22-0.44) | 1.89E-09 | 7.27E-09 |
| 305 | PC(17:1_18:2) | -0.76 | (-0.81--0.71) | 2.55E-144 | 8.86E-144 | -2.63 | (-2.69--2.57) | 0.00E+00 | 0.00E+00 | 1.66 | (1.6-1.72) | 0.00E+00 | 0.00E+00 | 0.12 | (0-0.24) | 4.18E-02 | 5.78E-02 |
| 306 | PC(15:0_20:3) | -0.32 | (-0.35--0.3) | 4.66E-101 | 1.18E-100 | -2.84 | (-2.91--2.77) | 0.00E+00 | 0.00E+00 | 2.42 | (2.34-2.5) | 0.00E+00 | 0.00E+00 | 0.14 | (0.07-0.21) | 5.45E-05 | 1.19E-04 |
| 307 | PC(15:0_20:4) | -0.74 | (-0.8--0.68) | 6.63E-105 | 1.74E-104 | -2.48 | (-2.55--2.41) | 0.00E+00 | 0.00E+00 | 1.28 | (1.21-1.34) | 1.15E-187 | 2.82E-187 | 0.46 | (0.32-0.59) | 1.12E-10 | 5.06E-10 |
| 308 | PC(35:5) | -0.29 | (-0.41--0.18) | 2.40E-07 | 2.94E-07 | -3.24 | (-3.36--3.12) | 0.00E+00 | 0.00E+00 | 2.10 | (1.99-2.22) | 4.45E-190 | 1.09E-189 | 0.78 | (0.55-1.02) | 1.74E-10 | 7.68E-10 |
| 309 | PC 36:0 | -0.46 | (-0.52--0.4) | 2.19E-50 | 3.87E-50 | 1.13 | (1.07-1.19) | 9.14E-218 | 1.38E-217 | -1.52 | (-1.58--1.47) | 4.58E-307 | 2.14E-306 | -0.38 | (-0.51--0.26) | 2.87E-09 | 1.07E-08 |
| 310 | PC(18:0_18:1) | -0.34 | (-0.39--0.28) | 4.23E-34 | 6.63E-34 | -1.24 | (-1.3--1.19) | 1.16E-257 | 1.90E-257 | 0.95 | (0.9-1.01) | 4.74E-179 | 1.10E-178 | -0.10 | (-0.21-0.01) | 8.80E-02 | 1.15E-01 |
| 311 | PC(18:0_18:2) | -0.06 | (-0.11--0.01) | 1.29E-02 | 1.41E-02 | -3.14 | (-3.2--3.08) | 0.00E+00 | 0.00E+00 | 3.02 | (2.96-3.08) | 0.00E+00 | 0.00E+00 | 0.10 | (0-0.21) | 6.13E-02 | 8.18E-02 |
| 312 | PC(16:0_20:3) (a) | -0.55 | (-0.6--0.49) | 1.79E-80 | 3.80E-80 | -3.41 | (-3.48--3.35) | 0.00E+00 | 0.00E+00 | 2.72 | (2.65-2.78) | 0.00E+00 | 0.00E+00 | 0.42 | (0.31-0.53) | 4.48E-13 | 2.69E-12 |
| 313 | PC(16:0_20:3) (b) | -1.38 | (-1.44--1.31) | 2.72E-241 | 2.04E-240 | -1.92 | (-1.99--1.85) | 0.00E+00 | 0.00E+00 | 0.39 | (0.33-0.45) | 2.77E-34 | 3.53E-34 | 0.00 | (-0.13-0.13) | 9.88E-01 | 9.88E-01 |
| 314 | PC(18:2_18:2) | -0.15 | (-0.22--0.08) | 8.13E-06 | 9.68E-06 | -5.22 | (-5.3--5.13) | 0.00E+00 | 0.00E+00 | 5.02 | (4.93-5.1) | 0.00E+00 | 0.00E+00 | 0.52 | (0.37-0.68) | 1.64E-11 | 8.36E-11 |
| 315 | PC(16:1_20:4) | -0.61 | (-0.66--0.55) | 6.60E-80 | 1.40E-79 | -1.43 | (-1.5--1.37) | 1.55E-247 | 2.47E-247 | 0.43 | (0.37-0.49) | 1.71E-37 | 2.22E-37 | 0.34 | (0.22-0.47) | 4.74E-08 | 1.55E-07 |
| 316 | PC(16:0_20:5) | -0.38 | (-0.48--0.27) | 3.50E-12 | 4.52E-12 | -2.58 | (-2.69--2.46) | 2.34E-265 | 3.98E-265 | 1.34 | (1.23-1.44) | 4.91E-109 | 8.33E-109 | 0.90 | (0.68-1.12) | 1.64E-15 | 1.40E-14 |
| 317 | PC(36:6) | -1.15 | (-1.23--1.07) | 6.42E-143 | 2.18E-142 | -3.30 | (-3.38--3.22) | 0.00E+00 | 0.00E+00 | 1.77 | (1.7-1.85) | 2.92E-261 | 1.04E-260 | 0.30 | (0.12-0.47) | 9.02E-04 | 1.62E-03 |
| 318 | PC(15-MHDA_20:4) | -0.82 | (-0.89--0.75) | 8.14E-101 | 2.05E-100 | -1.78 | (-1.86--1.71) | 1.75E-265 | 2.98E-265 | 0.67 | (0.58-0.75) | 2.08E-52 | 2.83E-52 | 0.27 | (0.12-0.43) | 6.20E-04 | 1.14E-03 |
| 319 | PC(17:0_20:4) | -0.41 | (-0.46--0.35) | 6.77E-44 | 1.14E-43 | -1.10 | (-1.16--1.04) | 5.85E-188 | 8.36E-188 | 0.29 | (0.23-0.36) | 6.06E-20 | 7.07E-20 | 0.41 | (0.29-0.53) | 7.65E-11 | 3.60E-10 |
| 320 | PC(15:0_22:6) | -0.83 | (-0.9--0.77) | 7.04E-113 | 1.95E-112 | -2.79 | (-2.87--2.72) | 0.00E+00 | 0.00E+00 | 1.51 | (1.44-1.59) | 5.42E-215 | 1.54E-214 | 0.43 | (0.28-0.57) | 1.67E-08 | 5.81E-08 |
| 321 | PC(38:2) | -0.55 | (-0.59--0.51) | 1.50E-119 | 4.33E-119 | -1.24 | (-1.29--1.19) | 2.55E-297 | 4.68E-297 | 0.59 | (0.54-0.64) | 1.13E-101 | 1.88E-101 | 0.10 | (0.01-0.18) | 3.09E-02 | 4.35E-02 |
| 322 | PC(18:0_20:3) | -0.46 | (-0.53--0.4) | 1.76E-40 | 2.88E-40 | -0.21 | (-0.27--0.14) | 2.76E-10 | 2.95E-10 | -0.30 | (-0.36--0.24) | 1.70E-22 | 2.02E-22 | -0.37 | (-0.51--0.24) | 8.97E-08 | 2.85E-07 |
| 323 | PC(18:1_20:3) | -0.84 | (-0.9--0.79) | 7.38E-157 | 2.81E-156 | -1.75 | (-1.81--1.69) | 0.00E+00 | 0.00E+00 | 0.72 | (0.66-0.78) | 2.67E-105 | 4.48E-105 | 0.10 | (-0.01-0.21) | 7.72E-02 | 1.01E-01 |

| | | | | | | | | | | | | | | | | | |
|---|---|---|---|---|---|---|---|---|---|---|---|---|---|---|---|---|---|
| 324 | PC(38:4) (b) | -1.09 | (-1.16--1.03) | 3.24E-182 | 1.57E-181 | -1.82 | (-1.89--1.75) | 0.00E+00 | 0.00E+00 | 0.76 | (0.69-0.83) | 2.59E-86 | 4.09E-86 | -0.17 | (-0.31--0.04) | 1.28E-02 | 1.94E-02 |
| 325 | PC(18:0_20:4) | 0.08 | (0.02-0.13) | 6.69E-03 | 7.36E-03 | -0.17 | (-0.23--0.1) | 2.55E-07 | 2.71E-07 | -0.03 | (-0.09-0.03) | 3.60E-01 | 3.63E-01 | 0.18 | (0.06-0.29) | 3.20E-03 | 5.32E-03 |
| 326 | PC(38:5) (a) | -0.38 | (-0.43--0.34) | 2.14E-57 | 3.98E-57 | -2.25 | (-2.3--2.19) | 0.00E+00 | 0.00E+00 | 1.56 | (1.5-1.62) | 0.00E+00 | 0.00E+00 | 0.35 | (0.25-0.45) | 1.67E-11 | 8.44E-11 |
| 327 | PC(38:5) (b) | -0.75 | (-0.81--0.68) | 1.06E-89 | 2.39E-89 | -1.89 | (-1.96--1.82) | 0.00E+00 | 0.00E+00 | 0.88 | (0.81-0.95) | 8.88E-115 | 1.57E-114 | 0.22 | (0.07-0.37) | 3.13E-03 | 5.22E-03 |
| 328 | PC(38:6) (a) | -0.09 | (-0.15--0.04) | 7.41E-04 | 8.41E-04 | -2.88 | (-2.95--2.82) | 0.00E+00 | 0.00E+00 | 2.42 | (2.35-2.48) | 0.00E+00 | 0.00E+00 | 0.60 | (0.48-0.72) | 3.53E-22 | 7.07E-21 |
| 329 | PC(16:0_22:6) | -0.86 | (-0.92--0.8) | 1.05E-134 | 3.38E-134 | -1.64 | (-1.71--1.57) | 1.44E-269 | 2.49E-269 | 0.34 | (0.26-0.41) | 1.95E-17 | 2.27E-17 | 0.47 | (0.33-0.61) | 1.04E-10 | 4.77E-10 |
| 330 | PC(18:2_20:5) | 0.15 | (0.06-0.24) | 7.21E-04 | 8.20E-04 | -3.25 | (-3.34--3.16) | 0.00E+00 | 0.00E+00 | 2.81 | (2.72-2.9) | 0.00E+00 | 0.00E+00 | 0.86 | (0.67-1.05) | 9.17E-18 | 1.16E-16 |
| 331 | PC(16:1_22:6) | -0.71 | (-0.77--0.65) | 1.23E-96 | 2.96E-96 | -1.73 | (-1.79--1.66) | 0.00E+00 | 0.00E+00 | 0.55 | (0.49-0.61) | 1.56E-57 | 2.20E-57 | 0.50 | (0.37-0.64) | 4.27E-13 | 2.60E-12 |
| 332 | PC(38:7)(c) | -0.84 | (-0.89--0.79) | 3.32E-161 | 1.33E-160 | -1.90 | (-1.96--1.85) | 0.00E+00 | 0.00E+00 | 0.67 | (0.62-0.73) | 1.37E-102 | 2.29E-102 | 0.44 | (0.33-0.56) | 7.56E-14 | 5.11E-13 |
| 333 | PC(39:5)(a) | -0.54 | (-0.62--0.47) | 2.59E-45 | 4.43E-45 | -2.18 | (-2.26--2.09) | 0.00E+00 | 0.00E+00 | 1.25 | (1.17-1.34) | 5.89E-137 | 1.12E-136 | 0.47 | (0.31-0.63) | 1.97E-08 | 6.82E-08 |
| 334 | PC(39:5)(b) | -0.67 | (-0.74--0.6) | 7.58E-77 | 1.58E-76 | -2.24 | (-2.32--2.17) | 0.00E+00 | 0.00E+00 | 1.24 | (1.16-1.32) | 2.06E-149 | 4.13E-149 | 0.36 | (0.21-0.52) | 3.72E-06 | 9.66E-06 |
| 335 | PC(15-MHDA_22:6) | -0.46 | (-0.53--0.39) | 5.16E-36 | 8.18E-36 | -2.17 | (-2.25--2.08) | 0.00E+00 | 0.00E+00 | 1.42 | (1.34-1.5) | 3.56E-166 | 7.66E-166 | 0.38 | (0.22-0.53) | 1.56E-06 | 4.33E-06 |
| 336 | PC(17:0_22:6) | -0.79 | (-0.85--0.72) | 5.46E-99 | 1.35E-98 | -2.21 | (-2.28--2.14) | 0.00E+00 | 0.00E+00 | 0.98 | (0.9-1.05) | 9.36E-111 | 1.63E-110 | 0.48 | (0.32-0.63) | 1.54E-09 | 6.12E-09 |
| 337 | PC(18:0_22:5) (n6) | -0.23 | (-0.3--0.16) | 1.02E-10 | 1.29E-10 | -1.78 | (-1.88--1.67) | 2.15E-176 | 3.01E-176 | 1.51 | (1.4-1.62) | 5.45E-123 | 9.98E-123 | -0.11 | (-0.26-0.03) | 1.30E-01 | 1.64E-01 |
| 338 | PC(18:0_22:6) | -0.53 | (-0.6--0.46) | 2.24E-51 | 3.99E-51 | -1.30 | (-1.37--1.23) | 2.04E-187 | 2.90E-187 | 0.41 | (0.34-0.49) | 1.16E-25 | 1.42E-25 | 0.16 | (0.01-0.31) | 3.26E-02 | 4.57E-02 |
| 339 | PC(40:7) (a) | -0.48 | (-0.54--0.42) | 4.15E-53 | 7.47E-53 | -1.15 | (-1.23--1.08) | 3.90E-149 | 5.20E-149 | 0.47 | (0.39-0.56) | 6.30E-29 | 7.81E-29 | 0.33 | (0.2-0.46) | 1.32E-06 | 3.67E-06 |
| 340 | PC(18:1_22:6) (a) | -0.84 | (-0.89--0.79) | 2.27E-169 | 1.03E-168 | -1.52 | (-1.58--1.46) | 1.11E-285 | 1.99E-285 | 0.36 | (0.29-0.43) | 2.24E-25 | 2.75E-25 | 0.38 | (0.27-0.5) | 1.32E-10 | 5.89E-10 |
| 341 | PC(40:8) | -0.29 | (-0.34--0.24) | 7.54E-32 | 1.15E-31 | -1.51 | (-1.57--1.45) | 3.78E-308 | 7.12E-308 | 0.97 | (0.91-1.03) | 8.28E-162 | 1.75E-161 | 0.58 | (0.48-0.69) | 5.20E-27 | 3.57E-25 |
| 342 | PC(44:12) | -0.71 | (-0.81--0.61) | 6.98E-41 | 1.15E-40 | -1.77 | (-1.86--1.67) | 3.52E-195 | 5.14E-195 | 0.48 | (0.39-0.58) | 2.03E-23 | 2.44E-23 | 0.96 | (0.73-1.18) | 2.11E-16 | 2.02E-15 |
| 343 | PC(O-16:0/16:0) | -0.22 | (-0.26--0.18) | 6.69E-29 | 9.97E-29 | -0.74 | (-0.79--0.69) | 1.35E-131 | 1.75E-131 | 0.47 | (0.42-0.52) | 3.83E-59 | 5.49E-59 | 0.22 | (0.14-0.3) | 1.00E-07 | 3.15E-07 |
| 344 | PC(O-32:1) | -0.43 | (-0.48--0.38) | 8.01E-59 | 1.50E-58 | -0.97 | (-1.03--0.91) | 8.24E-154 | 1.11E-153 | 0.25 | (0.19-0.32) | 1.53E-15 | 1.73E-15 | 0.41 | (0.31-0.52) | 4.38E-14 | 3.14E-13 |
| 345 | PC(O-34:1) | -0.23 | (-0.27--0.2) | 1.82E-31 | 2.77E-31 | -1.34 | (-1.39--1.29) | 9.28E-303 | 1.72E-302 | 1.08 | (1.02-1.13) | 1.57E-211 | 4.43E-211 | 0.25 | (0.17-0.33) | 1.48E-09 | 5.94E-09 |
| 346 | PC(O-34:2) | -0.31 | (-0.37--0.26) | 2.84E-27 | 4.16E-27 | -3.75 | (-3.81--3.68) | 0.00E+00 | 0.00E+00 | 3.18 | (3.11-3.25) | 0.00E+00 | 0.00E+00 | 0.61 | (0.49-0.73) | 1.12E-21 | 2.07E-20 |
| 347 | PC(O-34:4) | 0.03 | (-0.06-0.12) | 5.39E-01 | 5.51E-01 | -2.50 | (-2.59--2.41) | 0.00E+00 | 0.00E+00 | 1.35 | (1.26-1.43) | 1.43E-160 | 3.00E-160 | 1.18 | (0.98-1.38) | 6.55E-29 | 5.24E-27 |
| 348 | PC(O-35:4) | 0.06 | (-0.02-0.14) | 1.70E-01 | 1.79E-01 | -2.95 | (-3.03--2.87) | 0.00E+00 | 0.00E+00 | 2.14 | (2.06-2.21) | 9.61E-299 | 4.19E-298 | 0.98 | (0.79-1.17) | 1.47E-23 | 3.72E-22 |
| 349 | PC(O-36:0) | -0.27 | (-0.31--0.22) | 1.86E-30 | 2.79E-30 | -1.47 | (-1.53--1.4) | 1.41E-260 | 2.35E-260 | 1.17 | (1.1-1.24) | 2.62E-172 | 5.88E-172 | 0.17 | (0.07-0.26) | 7.04E-04 | 1.29E-03 |
| 350 | PC(O-18:0/18:1) | -0.13 | (-0.18--0.09) | 3.79E-10 | 4.75E-10 | -1.39 | (-1.44--1.33) | 2.54E-304 | 4.72E-304 | 1.29 | (1.24-1.35) | 2.14E-264 | 7.83E-264 | 0.21 | (0.13-0.3) | 1.15E-06 | 3.26E-06 |
| 351 | PC(O-18:1/18:1) | -0.11 | (-0.15--0.07) | 2.29E-07 | 2.82E-07 | -1.85 | (-1.9--1.79) | 0.00E+00 | 0.00E+00 | 1.63 | (1.58-1.68) | 0.00E+00 | 0.00E+00 | 0.35 | (0.27-0.43) | 7.87E-17 | 8.21E-16 |
| 352 | PC(O-18:0/18:2) | -0.19 | (-0.24--0.14) | 2.28E-13 | 2.98E-13 | -3.22 | (-3.28--3.16) | 0.00E+00 | 0.00E+00 | 2.91 | (2.85-2.97) | 0.00E+00 | 0.00E+00 | 0.43 | (0.33-0.53) | 2.15E-16 | 2.02E-15 |
| 353 | PC(O-18:1/18:2) | -0.25 | (-0.3--0.2) | 7.58E-23 | 1.08E-22 | -3.20 | (-3.26--3.14) | 0.00E+00 | 0.00E+00 | 2.76 | (2.7-2.82) | 0.00E+00 | 0.00E+00 | 0.46 | (0.36-0.56) | 9.17E-18 | 1.16E-16 |
| 354 | PC(O-16:0/20:3) | -0.31 | (-0.35--0.26) | 9.20E-35 | 1.45E-34 | -1.88 | (-1.93--1.82) | 0.00E+00 | 0.00E+00 | 1.48 | (1.42-1.54) | 1.79E-273 | 6.91E-273 | 0.16 | (0.06-0.27) | 2.20E-03 | 3.79E-03 |
| 355 | PC(O-16:0/20:4) | -0.16 | (-0.21--0.11) | 2.52E-11 | 3.24E-11 | -1.48 | (-1.54--1.43) | 0.00E+00 | 0.00E+00 | 1.16 | (1.11-1.22) | 3.97E-223 | 1.19E-222 | 0.28 | (0.18-0.38) | 1.28E-07 | 3.96E-07 |
| 356 | PC(O-36:5) | 0.12 | (0.03-0.21) | 9.10E-03 | 9.93E-03 | -3.21 | (-3.3--3.12) | 0.00E+00 | 0.00E+00 | 2.57 | (2.49-2.66) | 0.00E+00 | 0.00E+00 | 0.88 | (0.67-1.08) | 4.92E-17 | 5.37E-16 |
| 357 | PC(O-18:0/20:4) | -0.02 | (-0.07-0.02) | 3.17E-01 | 3.31E-01 | -1.05 | (-1.11--1) | 1.39E-220 | 2.12E-220 | 1.06 | (1-1.11) | 8.12E-209 | 2.24E-208 | 0.12 | (0.02-0.21) | 1.54E-02 | 2.28E-02 |
| 358 | PC(O-38:5) | -0.17 | (-0.22--0.13) | 3.06E-15 | 4.11E-15 | -2.11 | (-2.16--2.05) | 0.00E+00 | 0.00E+00 | 1.87 | (1.81-1.93) | 0.00E+00 | 0.00E+00 | 0.21 | (0.11-0.3) | 2.85E-05 | 6.63E-05 |
| 359 | PC(O-16:0/22:6) | -0.59 | (-0.66--0.53) | 9.17E-70 | 1.85E-69 | -1.97 | (-2.03--1.9) | 0.00E+00 | 0.00E+00 | 1.02 | (0.95-1.08) | 2.91E-149 | 5.80E-149 | 0.42 | (0.28-0.55) | 1.92E-09 | 7.31E-09 |
| 360 | PC(O-40:5) | -0.26 | (-0.3--0.22) | 4.35E-31 | 6.54E-31 | -1.86 | (-1.91--1.81) | 0.00E+00 | 0.00E+00 | 1.57 | (1.52-1.62) | 0.00E+00 | 0.00E+00 | 0.18 | (0.09-0.27) | 8.69E-05 | 1.83E-04 |
| 361 | PC(O-18:0/22:6) | -0.24 | (-0.29--0.19) | 1.62E-20 | 2.26E-20 | -1.41 | (-1.46--1.35) | 1.11E-301 | 2.04E-301 | 1.06 | (1.01-1.12) | 2.81E-204 | 7.49E-204 | 0.23 | (0.12-0.34) | 2.33E-05 | 5.45E-05 |
| 362 | PC(O-40:7) (a) | -0.44 | (-0.5--0.38) | 2.68E-47 | 4.65E-47 | -2.21 | (-2.27--2.15) | 0.00E+00 | 0.00E+00 | 1.50 | (1.43-1.56) | 4.78E-261 | 1.69E-260 | 0.41 | (0.29-0.53) | 1.01E-10 | 4.68E-10 |
| 363 | PC(P-16:0/14:0) | -0.24 | (-0.3--0.19) | 1.37E-17 | 1.88E-17 | -0.93 | (-1--0.86) | 2.72E-131 | 3.52E-131 | 0.44 | (0.37-0.51) | 5.12E-34 | 6.50E-34 | 0.34 | (0.22-0.47) | 9.59E-08 | 3.03E-07 |
| 364 | PC(P-16:0/16:0) | -0.03 | (-0.08-0.01) | 1.69E-01 | 1.78E-01 | -0.76 | (-0.82--0.7) | 8.53E-115 | 1.06E-114 | 0.53 | (0.47-0.59) | 1.17E-60 | 1.69E-60 | 0.40 | (0.3-0.5) | 1.20E-15 | 1.05E-14 |
| 365 | PC(P-16:0/16:1) | -0.33 | (-0.39--0.28) | 7.48E-34 | 1.17E-33 | -0.83 | (-0.9--0.77) | 1.91E-115 | 2.39E-115 | 0.01 | (-0.05-0.08) | 6.54E-01 | 6.57E-01 | 0.59 | (0.49-0.7) | 2.98E-25 | 1.22E-23 |
| 366 | PC(P-16:0/18:0) | -0.01 | (-0.05-0.02) | 5.18E-01 | 5.33E-01 | -0.84 | (-0.89--0.79) | 1.83E-190 | 2.62E-190 | 0.75 | (0.71-0.8) | 1.69E-157 | 3.48E-157 | 0.31 | (0.24-0.38) | 9.77E-17 | 9.77E-16 |
| 367 | PC(P-16:0/18:1) | 0.02 | (-0.03-0.06) | 4.94E-01 | 5.09E-01 | -0.93 | (-0.99--0.87) | 2.02E-164 | 2.79E-164 | 0.66 | (0.6-0.71) | 1.19E-104 | 1.98E-104 | 0.59 | (0.49-0.68) | 1.53E-31 | 2.44E-29 |
| 368 | PC(P-16:0/18:2) | -0.09 | (-0.14--0.04) | 4.96E-04 | 5.67E-04 | -3.52 | (-3.59--3.46) | 0.00E+00 | 0.00E+00 | 3.27 | (3.2-3.33) | 0.00E+00 | 0.00E+00 | 0.42 | (0.31-0.53) | 7.54E-14 | 5.11E-13 |
| 369 | PC(P-16:0/18:3) | 0.01 | (-0.04-0.07) | 6.07E-01 | 6.18E-01 | -3.03 | (-3.09--2.97) | 0.00E+00 | 0.00E+00 | 2.81 | (2.76-2.87) | 0.00E+00 | 0.00E+00 | 0.40 | (0.28-0.52) | 3.29E-11 | 1.60E-10 |
| 370 | PC(P-35:2) (a) | -0.20 | (-0.28--0.13) | 3.58E-07 | 4.37E-07 | -3.72 | (-3.8--3.63) | 0.00E+00 | 0.00E+00 | 3.32 | (3.24-3.4) | 0.00E+00 | 0.00E+00 | 0.41 | (0.23-0.58) | 5.24E-06 | 1.33E-05 |
| 371 | PC(P-35:2) (b) | -0.11 | (-0.17--0.04) | 1.00E-03 | 1.13E-03 | -4.04 | (-4.11--3.96) | 0.00E+00 | 0.00E+00 | 3.68 | (3.61-3.75) | 0.00E+00 | 0.00E+00 | 0.57 | (0.43-0.71) | 4.06E-15 | 3.14E-14 |
| 372 | PC(P-15:0/20:4) (b) | 0.09 | (0.03-0.16) | 4.04E-03 | 4.50E-03 | -1.60 | (-1.68--1.53) | 3.95E-255 | 6.42E-255 | 1.45 | (1.37-1.52) | 5.33E-199 | 1.39E-198 | 0.25 | (0.1-0.4) | 1.04E-03 | 1.85E-03 |
| 373 | PC(P-18:1/18:1) | -0.06 | (-0.11--0.02) | 8.60E-03 | 9.45E-03 | -1.13 | (-1.2--1.07) | 7.07E-188 | 1.01E-187 | 0.84 | (0.78-0.9) | 4.16E-117 | 7.43E-117 | 0.47 | (0.37-0.56) | 1.53E-21 | 2.71E-20 |
| 374 | PC(P-18:0/18:2) | -0.02 | (-0.07-0.03) | 4.30E-01 | 4.45E-01 | -3.29 | (-3.35--3.23) | 0.00E+00 | 0.00E+00 | 3.20 | (3.15-3.26) | 0.00E+00 | 0.00E+00 | 0.40 | (0.29-0.5) | 1.96E-13 | 1.26E-12 |
| 375 | PC(P-16:0/20:4) | -0.12 | (-0.18--0.07) | 1.98E-06 | 2.37E-06 | -1.22 | (-1.28--1.16) | 7.29E-221 | 1.11E-220 | 1.00 | (0.94-1.06) | 4.31E-166 | 9.24E-166 | 0.22 | (0.11-0.33) | 4.94E-05 | 1.09E-04 |
| 376 | PC(P-16:0/20:5) | 0.19 | (0.09-0.28) | 9.83E-05 | 1.14E-04 | -2.98 | (-3.08--2.88) | 0.00E+00 | 0.00E+00 | 2.55 | (2.46-2.64) | 0.00E+00 | 0.00E+00 | 0.74 | (0.53-0.94) | 5.19E-12 | 2.86E-11 |
| 377 | PC(P-17:0/20:4) (a) | -0.12 | (-0.19--0.05) | 7.93E-04 | 8.98E-04 | -1.95 | (-2.03--1.87) | 2.55E-296 | 4.68E-296 | 1.73 | (1.65-1.81) | 3.43E-222 | 1.02E-221 | 0.21 | (0.05-0.38) | 1.25E-02 | 1.92E-02 |
| 378 | PC(P-17:0/20:4) (b) | 0.10 | (0.03-0.16) | 2.56E-03 | 2.87E-03 | -1.72 | (-1.79--1.64) | 9.54E-280 | 1.68E-279 | 1.59 | (1.52-1.66) | 1.15E-236 | 3.64E-236 | 0.37 | (0.23-0.52) | 2.25E-07 | 6.72E-07 |

| | | | | | | | | | | | | | | | | | |
|---|---|---|---|---|---|---|---|---|---|---|---|---|---|---|---|---|---|
| 379 | PC(P-18:0/20:4) | 0.19 | (0.14-0.24) | 3.64E-14 | 4.80E-14 | -1.06 | (-1.12--1) | 1.24E-199 | 1.82E-199 | 1.28 | (1.22-1.33) | 1.18E-244 | 3.89E-244 | 0.14 | (0.03-0.24) | 1.29E-02 | 1.96E-02 |
| 380 | PC(P-38:5) (a) | -0.08 | (-0.13--0.03) | 3.08E-03 | 3.44E-03 | -2.01 | (-2.07--1.94) | 0.00E+00 | 0.00E+00 | 1.68 | (1.62-1.74) | 0.00E+00 | 0.00E+00 | 0.33 | (0.22-0.44) | 1.78E-09 | 6.95E-09 |
| 381 | PC(P-38:5) (b) | -0.21 | (-0.26--0.16) | 4.77E-15 | 6.36E-15 | -1.38 | (-1.45--1.32) | 1.77E-250 | 2.84E-250 | 1.12 | (1.05-1.18) | 1.92E-171 | 4.28E-171 | 0.16 | (0.04-0.27) | 7.20E-03 | 1.15E-02 |
| 382 | PC(P-16:0/22:6) | -0.45 | (-0.51--0.39) | 3.89E-42 | 6.47E-42 | -1.88 | (-1.95--1.81) | 0.00E+00 | 0.00E+00 | 1.17 | (1.1-1.24) | 1.05E-173 | 2.39E-173 | 0.38 | (0.25-0.52) | 3.12E-08 | 1.03E-07 |
| 383 | PC(P-20:0/20:4) | -0.18 | (-0.23--0.13) | 1.57E-12 | 2.04E-12 | -0.96 | (-1.02--0.9) | 5.29E-155 | 7.11E-155 | 0.97 | (0.91-1.03) | 1.31E-157 | 2.72E-157 | 0.03 | (-0.08-0.13) | 6.27E-01 | 6.84E-01 |
| 384 | PC(P-40:5) (b) | -0.05 | (-0.1-0) | 6.14E-02 | 6.55E-02 | -1.24 | (-1.3--1.19) | 5.33E-259 | 8.82E-259 | 1.17 | (1.12-1.23) | 7.56E-243 | 2.47E-242 | 0.21 | (0.11-0.31) | 4.11E-05 | 9.34E-05 |
| 385 | PC(P-18:0/22:6) | -0.01 | (-0.07-0.05) | 7.22E-01 | 7.29E-01 | -1.80 | (-1.87--1.74) | 0.00E+00 | 0.00E+00 | 1.73 | (1.67-1.79) | 1.80E-305 | 8.23E-305 | 0.26 | (0.13-0.39) | 1.39E-04 | 2.83E-04 |
| 386 | PE(16:0_16:1) | -3.16 | (-3.27--3.05) | 0.00E+00 | 0.00E+00 | -3.71 | (-3.82--3.6) | 0.00E+00 | 0.00E+00 | 0.13 | (0.02-0.23) | 1.90E-02 | 1.96E-02 | 0.11 | (-0.13-0.35) | 3.86E-01 | 4.44E-01 |
| 387 | PE(16:0_18:1) | -2.76 | (-2.83--2.68) | 0.00E+00 | 0.00E+00 | -4.34 | (-4.42--4.27) | 0.00E+00 | 0.00E+00 | 1.25 | (1.17-1.33) | 1.59E-153 | 3.24E-153 | 0.24 | (0.06-0.42) | 7.61E-03 | 1.21E-02 |
| 388 | PE(16:0_18:2) | -2.98 | (-3.06--2.89) | 0.00E+00 | 0.00E+00 | -5.25 | (-5.34--5.17) | 0.00E+00 | 0.00E+00 | 1.88 | (1.79-1.96) | 2.99E-239 | 9.62E-239 | 0.40 | (0.21-0.6) | 4.67E-05 | 1.04E-04 |
| 389 | PE(16:1_18:2) | -2.41 | (-2.5--2.33) | 0.00E+00 | 0.00E+00 | -3.23 | (-3.33--3.14) | 0.00E+00 | 0.00E+00 | 0.53 | (0.46-0.61) | 2.56E-40 | 3.36E-40 | 0.27 | (0.11-0.44) | 1.41E-03 | 2.47E-03 |
| 390 | PE(16:0_18:3) (a) | -2.95 | (-3.04--2.86) | 0.00E+00 | 0.00E+00 | -4.63 | (-4.72--4.55) | 0.00E+00 | 0.00E+00 | 1.48 | (1.4-1.57) | 1.35E-173 | 3.05E-173 | 0.18 | (-0.02-0.37) | 7.25E-02 | 9.58E-02 |
| 391 | PE(16:0_18:3) (b) | -2.00 | (-2.09--1.91) | 6.60E-260 | 5.98E-259 | -2.95 | (-3.04--2.86) | 0.00E+00 | 0.00E+00 | 0.85 | (0.76-0.94) | 1.43E-63 | 2.07E-63 | -0.05 | (-0.26-0.16) | 6.25E-01 | 6.83E-01 |
| 392 | PE(17:0_18:2) | -2.10 | (-2.16--2.03) | 0.00E+00 | 0.00E+00 | -3.73 | (-3.79--3.66) | 0.00E+00 | 0.00E+00 | 1.36 | (1.3-1.43) | 7.17E-222 | 2.11E-221 | 0.26 | (0.1-0.41) | 1.42E-03 | 2.49E-03 |
| 393 | PE(18:0_18:1) | -2.12 | (-2.19--2.05) | 0.00E+00 | 0.00E+00 | -4.63 | (-4.7--4.56) | 0.00E+00 | 0.00E+00 | 2.21 | (2.13-2.29) | 0.00E+00 | 0.00E+00 | 0.09 | (-0.08-0.26) | 2.76E-01 | 3.28E-01 |
| 394 | PE(18:1_18:1) | -1.88 | (-1.95--1.82) | 0.00E+00 | 0.00E+00 | -3.26 | (-3.34--3.17) | 0.00E+00 | 0.00E+00 | 1.24 | (1.15-1.34) | 4.58E-113 | 8.05E-113 | 0.11 | (-0.05-0.26) | 1.76E-01 | 2.16E-01 |
| 395 | PE(18:0_18:2) | -1.92 | (-1.99--1.85) | 0.00E+00 | 0.00E+00 | -5.13 | (-5.2--5.06) | 0.00E+00 | 0.00E+00 | 2.84 | (2.76-2.91) | 0.00E+00 | 0.00E+00 | 0.21 | (0.04-0.38) | 1.61E-02 | 2.38E-02 |
| 396 | PE(18:1_18:2) | -2.56 | (-2.64--2.48) | 0.00E+00 | 0.00E+00 | -5.16 | (-5.24--5.08) | 0.00E+00 | 0.00E+00 | 2.34 | (2.25-2.42) | 0.00E+00 | 0.00E+00 | 0.39 | (0.21-0.58) | 3.89E-05 | 8.98E-05 |
| 397 | PE(16:0_20:3) | -2.70 | (-2.78--2.62) | 0.00E+00 | 0.00E+00 | -3.65 | (-3.73--3.56) | 0.00E+00 | 0.00E+00 | 0.68 | (0.6-0.76) | 4.27E-53 | 5.84E-53 | 0.19 | (0.01-0.38) | 4.19E-02 | 5.78E-02 |
| 398 | PE(16:0_20:4) | -2.49 | (-2.56--2.43) | 0.00E+00 | 0.00E+00 | -2.89 | (-2.96--2.82) | 0.00E+00 | 0.00E+00 | 0.00 | (-0.08-0.07) | 9.25E-01 | 9.25E-01 | 0.46 | (0.3-0.63) | 5.04E-08 | 1.62E-07 |
| 399 | PE(16:1_20:4) | -1.96 | (-2.03--1.88) | 0.00E+00 | 0.00E+00 | -1.86 | (-1.94--1.77) | 1.33E-254 | 2.15E-254 | -0.38 | (-0.45--0.31) | 2.17E-23 | 2.60E-23 | 0.30 | (0.16-0.45) | 4.86E-05 | 1.08E-04 |
| 400 | PE(16:0_20:5) | -1.94 | (-2.03--1.85) | 8.70E-233 | 6.05E-232 | -3.28 | (-3.37--3.18) | 0.00E+00 | 0.00E+00 | 0.65 | (0.56-0.73) | 1.25E-45 | 1.67E-45 | 0.75 | (0.55-0.95) | 2.55E-13 | 1.61E-12 |
| 401 | PE(17:0_20:4) | -1.81 | (-1.88--1.75) | 0.00E+00 | 0.00E+00 | -2.77 | (-2.84--2.7) | 0.00E+00 | 0.00E+00 | 0.58 | (0.51-0.65) | 1.70E-50 | 2.31E-50 | 0.36 | (0.21-0.51) | 3.55E-06 | 9.37E-06 |
| 402 | PE(18:0_20:3) (a) | -1.88 | (-1.96--1.8) | 1.97E-292 | 2.56E-291 | -2.47 | (-2.55--2.4) | 0.00E+00 | 0.00E+00 | 0.25 | (0.18-0.32) | 3.15E-11 | 3.47E-11 | -0.04 | (-0.21-0.13) | 6.43E-01 | 6.97E-01 |
| 403 | PE(18:0_20:4) | -1.43 | (-1.49--1.37) | 1.38E-288 | 1.65E-287 | -2.29 | (-2.35--2.23) | 0.00E+00 | 0.00E+00 | 0.40 | (0.34-0.47) | 6.36E-32 | 7.93E-32 | 0.35 | (0.22-0.49) | 7.02E-07 | 2.03E-06 |
| 404 | PE(38:5) (a) | -2.33 | (-2.4--2.26) | 0.00E+00 | 0.00E+00 | -3.31 | (-3.38--3.24) | 0.00E+00 | 0.00E+00 | 0.54 | (0.47-0.61) | 1.97E-44 | 2.62E-44 | 0.53 | (0.37-0.69) | 2.90E-10 | 1.24E-09 |
| 405 | PE(38:5) (b) | -2.22 | (-2.3--2.14) | 0.00E+00 | 0.00E+00 | -3.76 | (-3.85--3.67) | 0.00E+00 | 0.00E+00 | 1.10 | (1-1.2) | 9.71E-90 | 1.55E-89 | 0.44 | (0.25-0.62) | 4.32E-06 | 1.12E-05 |
| 406 | PE(16:0_22:6) | -2.61 | (-2.69--2.54) | 0.00E+00 | 0.00E+00 | -3.78 | (-3.86--3.71) | 0.00E+00 | 0.00E+00 | 0.35 | (0.27-0.44) | 6.39E-16 | 7.29E-16 | 0.84 | (0.65-1.03) | 1.03E-17 | 1.25E-16 |
| 407 | PE(17:0_22:6) | -1.87 | (-1.94--1.8) | 0.00E+00 | 0.00E+00 | -2.84 | (-2.91--2.76) | 0.00E+00 | 0.00E+00 | 0.40 | (0.32-0.47) | 2.59E-26 | 3.19E-26 | 0.55 | (0.39-0.7) | 6.97E-12 | 3.80E-11 |
| 408 | PE(18:0_22:4) | -1.76 | (-1.83--1.68) | 7.99E-281 | 8.92E-280 | -2.46 | (-2.54--2.37) | 0.00E+00 | 0.00E+00 | 0.61 | (0.53-0.68) | 2.33E-50 | 3.16E-50 | -0.16 | (-0.31--0.01) | 3.94E-02 | 5.50E-02 |
| 409 | PE(18:0_22:5) (n3) | -1.78 | (-1.85--1.71) | 0.00E+00 | 0.00E+00 | -3.49 | (-3.57--3.41) | 0.00E+00 | 0.00E+00 | 1.18 | (1.09-1.26) | 3.64E-129 | 6.83E-129 | 0.28 | (0.11-0.44) | 8.26E-04 | 1.51E-03 |
| 410 | PE(18:0_22:5) (n6) | -1.87 | (-1.95--1.78) | 7.23E-262 | 6.80E-261 | -3.05 | (-3.14--2.95) | 0.00E+00 | 0.00E+00 | 0.93 | (0.84-1.03) | 4.94E-72 | 7.46E-72 | 0.07 | (-0.11-0.25) | 4.50E-01 | 5.09E-01 |
| 411 | PE(18:0_22:6) | -2.01 | (-2.09--1.94) | 0.00E+00 | 0.00E+00 | -3.62 | (-3.7--3.53) | 0.00E+00 | 0.00E+00 | 0.68 | (0.59-0.77) | 5.40E-46 | 7.24E-46 | 0.58 | (0.4-0.76) | 6.69E-10 | 2.77E-09 |
| 412 | PE(40:7) | -3.13 | (-3.21--3.05) | 0.00E+00 | 0.00E+00 | -4.36 | (-4.44--4.28) | 0.00E+00 | 0.00E+00 | 0.44 | (0.35-0.52) | 8.83E-23 | 1.05E-22 | 0.90 | (0.7-1.09) | 1.56E-18 | 2.14E-17 |
| 413 | PE(O-34:1) | -0.09 | (-0.14--0.03) | 1.42E-03 | 1.60E-03 | -3.47 | (-3.56--3.39) | 0.00E+00 | 0.00E+00 | 3.52 | (3.43-3.61) | 0.00E+00 | 0.00E+00 | 0.15 | (0.03-0.27) | 1.27E-02 | 1.94E-02 |
| 414 | PE(O-16:0/18:2) | -0.16 | (-0.23--0.08) | 2.18E-05 | 2.58E-05 | -5.76 | (-5.9--5.62) | 0.00E+00 | 0.00E+00 | 5.48 | (5.33-5.63) | 0.00E+00 | 0.00E+00 | 0.54 | (0.38-0.69) | 3.62E-11 | 1.74E-10 |
| 415 | PE(O-18:1/18:2) | 0.06 | (0-0.13) | 5.95E-02 | 6.36E-02 | -3.00 | (-3.09--2.91) | 0.00E+00 | 0.00E+00 | 2.90 | (2.82-2.99) | 0.00E+00 | 0.00E+00 | 0.32 | (0.18-0.46) | 8.78E-06 | 2.15E-05 |
| 416 | PE(O-16:0/20:4) | 0.14 | (0.07-0.21) | 4.57E-05 | 5.37E-05 | -3.07 | (-3.15--2.99) | 0.00E+00 | 0.00E+00 | 3.02 | (2.93-3.1) | 0.00E+00 | 0.00E+00 | 0.35 | (0.2-0.5) | 5.37E-06 | 1.35E-05 |
| 417 | PE(O-36:5) | -0.22 | (-0.32--0.12) | 2.76E-05 | 3.24E-05 | -3.40 | (-3.5--3.31) | 0.00E+00 | 0.00E+00 | 2.50 | (2.42-2.58) | 0.00E+00 | 0.00E+00 | 0.85 | (0.64-1.07) | 1.58E-14 | 1.18E-13 |
| 418 | PE(O-16:0/22:4) | -0.09 | (-0.14--0.04) | 8.50E-04 | 9.60E-04 | -1.25 | (-1.32--1.19) | 8.24E-219 | 1.25E-218 | 1.15 | (1.08-1.21) | 3.93E-181 | 9.20E-181 | 0.10 | (-0.02-0.21) | 9.50E-02 | 1.24E-01 |
| 419 | PE(O-18:0/20:4) | 0.22 | (0.14-0.29) | 6.21E-09 | 7.72E-09 | -2.78 | (-2.87--2.7) | 0.00E+00 | 0.00E+00 | 2.80 | (2.71-2.88) | 0.00E+00 | 0.00E+00 | 0.33 | (0.17-0.5) | 5.81E-05 | 1.26E-04 |
| 420 | PE(O-38:5) (a) | 0.16 | (0.1-0.23) | 6.00E-07 | 7.30E-07 | -3.65 | (-3.72--3.57) | 0.00E+00 | 0.00E+00 | 3.64 | (3.57-3.71) | 0.00E+00 | 0.00E+00 | 0.29 | (0.14-0.43) | 1.07E-04 | 2.20E-04 |
| 421 | PE(O-38:5) (b) | -0.07 | (-0.14--0.01) | 3.08E-02 | 3.31E-02 | -2.25 | (-2.33--2.17) | 0.00E+00 | 0.00E+00 | 2.02 | (1.94-2.1) | 3.65E-280 | 1.46E-279 | 0.34 | (0.2-0.49) | 3.61E-06 | 9.47E-06 |
| 422 | PE(O-16:0/22:6) | -0.22 | (-0.28--0.16) | 4.71E-13 | 6.12E-13 | -2.92 | (-2.99--2.85) | 0.00E+00 | 0.00E+00 | 2.48 | (2.41-2.55) | 0.00E+00 | 0.00E+00 | 0.42 | (0.28-0.55) | 1.18E-09 | 4.79E-09 |
| 423 | PE(O-18:0/22:6) | 0.08 | (0.02-0.13) | 8.79E-03 | 9.64E-03 | -2.28 | (-2.34--2.22) | 0.00E+00 | 0.00E+00 | 2.21 | (2.15-2.27) | 0.00E+00 | 0.00E+00 | 0.31 | (0.19-0.44) | 1.05E-06 | 2.97E-06 |
| 424 | PE(O-18:1/22:6) | -0.11 | (-0.16--0.05) | 9.61E-05 | 1.12E-04 | -2.57 | (-2.63--2.51) | 0.00E+00 | 0.00E+00 | 2.29 | (2.23-2.35) | 0.00E+00 | 0.00E+00 | 0.36 | (0.24-0.49) | 1.06E-08 | 3.79E-08 |
| 425 | PE(P-16:0/18:1) | -0.08 | (-0.13--0.04) | 5.41E-04 | 6.17E-04 | -1.29 | (-1.35--1.23) | 7.45E-263 | 1.25E-262 | 1.63 | (1.58-1.69) | 0.00E+00 | 0.00E+00 | -0.27 | (-0.37--0.17) | 1.20E-07 | 3.73E-07 |
| 426 | PE(P-16:0/18:2) | -0.17 | (-0.23--0.11) | 8.40E-08 | 1.03E-07 | -4.68 | (-4.75--4.6) | 0.00E+00 | 0.00E+00 | 4.76 | (4.68-4.84) | 0.00E+00 | 0.00E+00 | -0.13 | (-0.26-0) | 5.15E-02 | 6.94E-02 |
| 427 | PE(P-16:0/20:3) | -0.08 | (-0.14--0.01) | 1.70E-02 | 1.84E-02 | -2.07 | (-2.15--2) | 0.00E+00 | 0.00E+00 | 2.32 | (2.25-2.4) | 0.00E+00 | 0.00E+00 | -0.48 | (-0.62--0.35) | 3.27E-12 | 1.82E-11 |
| 428 | PE(P-16:0/20:4) | -0.01 | (-0.07-0.06) | 7.69E-01 | 7.73E-01 | -2.82 | (-2.89--2.74) | 0.00E+00 | 0.00E+00 | 2.91 | (2.84-2.99) | 0.00E+00 | 0.00E+00 | -0.12 | (-0.26-0.02) | 1.05E-01 | 1.37E-01 |
| 429 | PE(P-16:0/20:5) | -0.04 | (-0.16-0.08) | 5.20E-01 | 5.33E-01 | -6.00 | (-6.16--5.84) | 0.00E+00 | 0.00E+00 | 5.55 | (5.38-5.72) | 0.00E+00 | 0.00E+00 | 0.46 | (0.2-0.71) | 3.95E-04 | 7.46E-04 |
| 430 | PE(P-16:0/22:4) | -0.23 | (-0.28--0.18) | 1.31E-19 | 1.82E-19 | 0.01 | (-0.05-0.07) | 7.03E-01 | 7.04E-01 | 0.32 | (0.26-0.38) | 8.21E-25 | 1.00E-24 | -0.51 | (-0.61--0.4) | 6.69E-21 | 1.11E-19 |
| 431 | PE(P-16:0/22:5) (n3) | -0.05 | (-0.1-0) | 4.44E-02 | 4.77E-02 | -2.95 | (-3.01--2.88) | 0.00E+00 | 0.00E+00 | 2.98 | (2.92-3.05) | 0.00E+00 | 0.00E+00 | -0.09 | (-0.21-0.02) | 1.10E-01 | 1.41E-01 |
| 432 | PE(P-16:0/22:5) (n6) | -0.38 | (-0.43--0.32) | 1.15E-38 | 1.85E-38 | -1.26 | (-1.33--1.19) | 4.86E-195 | 7.06E-195 | 1.27 | (1.2-1.34) | 3.45E-183 | 8.21E-183 | -0.35 | (-0.47--0.22) | 2.90E-08 | 9.67E-08 |
| 433 | PE(P-16:0/22:6) | -0.43 | (-0.48--0.38) | 7.08E-53 | 1.27E-52 | -2.16 | (-2.22--2.1) | 0.00E+00 | 0.00E+00 | 1.63 | (1.57-1.69) | 1.15E-294 | 4.83E-294 | 0.18 | (0.06-0.29) | 2.49E-03 | 4.24E-03 |

| | | | | | | | | | | | | | | | | | |
|---|---|---|---|---|---|---|---|---|---|---|---|---|---|---|---|---|---|
| 434 | PE(P-17:0/20:4) (a) | 0.17 | (0.09-0.25) | **2.47E-05** | **2.91E-05** | -2.11 | (-2.19--2.04) | **0.00E+00** | **0.00E+00** | 2.04 | (1.97-2.11) | **0.00E+00** | **0.00E+00** | 0.22 | (0.05-0.39) | **1.23E-02** | **1.90E-02** |
| 435 | PE(P-17:0/20:4) (b) | 0.20 | (0.13-0.27) | **3.57E-08** | **4.41E-08** | -2.39 | (-2.47--2.32) | **0.00E+00** | **0.00E+00** | 2.30 | (2.23-2.37) | **0.00E+00** | **0.00E+00** | 0.35 | (0.19-0.5) | **9.42E-06** | **2.30E-05** |
| 436 | PE(P-17:0/22:6) (a) | -0.17 | (-0.24--0.1) | **4.70E-06** | **5.61E-06** | -1.94 | (-2.02--1.86) | **6.55E-292** | **1.19E-291** | 1.47 | (1.4-1.55) | **1.75E-192** | **4.38E-192** | 0.34 | (0.18-0.5) | **4.82E-05** | **1.07E-04** |
| 437 | PE(P-17:0/22:6) (b) | -0.13 | (-0.2--0.07) | **1.12E-04** | **1.29E-04** | -2.45 | (-2.52--2.37) | **0.00E+00** | **0.00E+00** | 1.98 | (1.91-2.05) | **0.00E+00** | **0.00E+00** | 0.47 | (0.32-0.62) | **6.51E-10** | **2.72E-09** |
| 438 | PE(P-18:0/18:1) | -0.02 | (-0.08-0.03) | 4.12E-01 | 4.27E-01 | -2.96 | (-3.03--2.89) | **0.00E+00** | **0.00E+00** | 3.27 | (3.21-3.34) | **0.00E+00** | **0.00E+00** | -0.19 | (-0.31--0.08) | **8.92E-04** | **1.61E-03** |
| 439 | PE(P-18:0/18:2) | -0.07 | (-0.13--0.01) | **1.37E-02** | **1.49E-02** | -5.89 | (-5.97--5.81) | **0.00E+00** | **0.00E+00** | 6.06 | (5.98-6.14) | **0.00E+00** | **0.00E+00** | -0.03 | (-0.16-0.1) | 6.48E-01 | 7.01E-01 |
| 440 | PE(P-18:0/20:3) | -0.03 | (-0.09-0.03) | 2.87E-01 | 3.00E-01 | -2.33 | (-2.39--2.27) | **0.00E+00** | **0.00E+00** | 2.52 | (2.47-2.58) | **0.00E+00** | **0.00E+00** | -0.28 | (-0.4--0.17) | **1.71E-06** | **4.72E-06** |
| 441 | PE(P-18:0/20:4) | 0.32 | (0.26-0.39) | **7.50E-22** | **1.06E-21** | -2.97 | (-3.05--2.89) | **0.00E+00** | **0.00E+00** | 3.29 | (3.22-3.36) | **0.00E+00** | **0.00E+00** | 0.07 | (-0.07-0.21) | 3.26E-01 | 3.80E-01 |
| 442 | PE(P-18:0/20:5) | 0.17 | (0.05-0.29) | **6.48E-03** | **7.15E-03** | -6.21 | (-6.37--6.06) | **0.00E+00** | **0.00E+00** | 5.80 | (5.64-5.95) | **0.00E+00** | **0.00E+00** | 0.61 | (0.36-0.86) | **2.65E-06** | **7.14E-06** |
| 443 | PE(P-18:0/22:4) | -0.05 | (-0.11-0) | **4.71E-02** | 5.05E-02 | -0.10 | (-0.17--0.03) | **5.50E-03** | **5.70E-03** | 0.83 | (0.76-0.89) | **1.84E-105** | **3.09E-105** | -0.68 | (-0.79--0.57) | **3.65E-31** | **4.37E-29** |
| 444 | PE(P-18:0/22:5) (n3) | 0.22 | (0.17-0.27) | **6.61E-17** | **9.01E-17** | -2.43 | (-2.5--2.37) | **0.00E+00** | **0.00E+00** | 2.79 | (2.73-2.85) | **0.00E+00** | **0.00E+00** | -0.06 | (-0.16-0.05) | 3.15E-01 | 3.71E-01 |
| 445 | PE(P-18:0/22:5) (n6) | -0.12 | (-0.17--0.06) | **1.22E-05** | **1.45E-05** | -1.46 | (-1.52--1.4) | **3.82E-283** | **6.85E-283** | 1.69 | (1.63-1.76) | **3.92E-301** | **1.74E-300** | -0.22 | (-0.34--0.1) | **3.91E-04** | **7.41E-04** |
| 446 | PE(P-18:0/22:6) | 0.01 | (-0.05-0.07) | 7.16E-01 | 7.25E-01 | -2.50 | (-2.57--2.44) | **0.00E+00** | **0.00E+00** | 2.53 | (2.47-2.59) | **0.00E+00** | **0.00E+00** | 0.14 | (0.01-0.27) | **2.89E-02** | **4.09E-02** |
| 447 | PE(P-18:1/18:1) | -0.17 | (-0.21--0.12) | **6.16E-11** | **7.82E-11** | -2.39 | (-2.45--2.33) | **0.00E+00** | **0.00E+00** | 2.64 | (2.58-2.69) | **0.00E+00** | **0.00E+00** | -0.26 | (-0.36--0.15) | **1.81E-06** | **4.96E-06** |
| 448 | PE(P-18:1/18:2) | -0.22 | (-0.28--0.15) | **3.59E-11** | **4.57E-11** | -5.92 | (-6.01--5.82) | **0.00E+00** | **0.00E+00** | 5.98 | (5.88-6.08) | **0.00E+00** | **0.00E+00** | -0.06 | (-0.2-0.07) | 3.55E-01 | 4.12E-01 |
| 449 | PE(P-18:1/20:4) | 0.13 | (0.06-0.19) | **9.45E-05** | **1.10E-04** | -3.18 | (-3.26--3.1) | **0.00E+00** | **0.00E+00** | 3.42 | (3.34-3.49) | **0.00E+00** | **0.00E+00** | -0.04 | (-0.18-0.1) | 5.72E-01 | 6.33E-01 |
| 450 | PE(P-18:1/20:5) | -0.26 | (-0.38--0.14) | **1.73E-05** | **2.05E-05** | -6.95 | (-7.13--6.76) | **0.00E+00** | **0.00E+00** | 6.25 | (6.06-6.45) | **0.00E+00** | **0.00E+00** | 0.52 | (0.27-0.78) | **5.75E-05** | **1.25E-04** |
| 451 | PE(P-18:1/22:4) | -0.20 | (-0.25--0.14) | **3.07E-11** | **3.92E-11** | -0.20 | (-0.27--0.12) | **2.70E-07** | **2.86E-07** | 0.69 | (0.62-0.76) | **4.72E-68** | **6.94E-68** | -0.58 | (-0.7--0.46) | **2.22E-21** | **3.80E-20** |
| 452 | PE(P-18:1/22:5) (a) | 0.08 | (0.02-0.13) | **4.92E-03** | **5.45E-03** | -2.50 | (-2.56--2.43) | **0.00E+00** | **0.00E+00** | 2.66 | (2.6-2.72) | **0.00E+00** | **0.00E+00** | 0.02 | (-0.09-0.14) | 6.82E-01 | 7.29E-01 |
| 453 | PE(P-18:1/22:6) (a) | -0.37 | (-0.43--0.31) | **1.95E-33** | **3.02E-33** | -2.96 | (-3.03--2.89) | **0.00E+00** | **0.00E+00** | 2.56 | (2.49-2.62) | **0.00E+00** | **0.00E+00** | 0.17 | (0.04-0.3) | **9.78E-03** | **1.53E-02** |
| 454 | PE(P-20:0/18:1) | -0.11 | (-0.17--0.06) | **4.69E-05** | **5.49E-05** | -3.50 | (-3.62--3.38) | **0.00E+00** | **0.00E+00** | 4.02 | (3.89-4.15) | **0.00E+00** | **0.00E+00** | -0.52 | (-0.63--0.4) | **6.06E-18** | **8.09E-17** |
| 455 | PE(P-20:0/18:2) | -0.07 | (-0.12--0.02) | **4.13E-03** | **4.58E-03** | -3.29 | (-3.34--3.24) | **0.00E+00** | **0.00E+00** | 3.81 | (3.76-3.86) | **0.00E+00** | **0.00E+00** | -0.31 | (-0.42--0.2) | **2.13E-08** | **7.31E-08** |
| 456 | PE(P-20:0/20:4) | 0.42 | (0.37-0.47) | **2.45E-54** | **4.42E-54** | -1.57 | (-1.65--1.5) | **4.72E-235** | **7.42E-235** | 2.41 | (2.33-2.49) | **0.00E+00** | **0.00E+00** | -0.28 | (-0.39--0.17) | **4.21E-07** | **1.23E-06** |
| 457 | PE(P-20:0/22:6) | -0.12 | (-0.16--0.07) | **9.70E-07** | **1.17E-06** | -1.87 | (-1.92--1.81) | **0.00E+00** | **0.00E+00** | 1.97 | (1.92-2.03) | **0.00E+00** | **0.00E+00** | -0.01 | (-0.11-0.09) | 8.53E-01 | 8.82E-01 |
| 458 | PE(P-20:1/20:4) | 0.35 | (0.27-0.44) | **1.53E-16** | **2.07E-16** | -2.86 | (-2.96--2.75) | **0.00E+00** | **0.00E+00** | 3.04 | (2.94-3.14) | **0.00E+00** | **0.00E+00** | 0.22 | (0.04-0.4) | **1.52E-02** | **2.27E-02** |
| 459 | PG(34:2) | -1.30 | (-1.38--1.23) | **5.76E-175** | **2.66E-174** | -2.53 | (-2.64--2.43) | **9.18E-287** | **1.66E-286** | 0.97 | (0.85-1.08) | **2.13E-54** | **2.95E-54** | 0.12 | (-0.07-0.3) | 2.16E-01 | 2.62E-01 |
| 460 | PG(36:1) | -0.93 | (-1--0.86) | **8.11E-121** | **2.36E-120** | -2.60 | (-2.7--2.5) | **6.20E-305** | **1.16E-304** | 1.25 | (1.13-1.36) | **3.36E-84** | **5.25E-84** | 0.01 | (-0.17-0.19) | 9.07E-01 | 9.25E-01 |
| 461 | PI(16:0/16:0) | -3.08 | (-3.17--2.99) | **0.00E+00** | **0.00E+00** | -4.55 | (-4.65--4.46) | **0.00E+00** | **0.00E+00** | 1.23 | (1.14-1.33) | **1.01E-118** | **1.83E-118** | 0.11 | (-0.08-0.3) | 2.67E-01 | 3.19E-01 |
| 462 | PI(16:0_16:1) | -1.94 | (-2.04--1.84) | **4.93E-220** | **3.24E-219** | -2.64 | (-2.74--2.54) | **0.00E+00** | **0.00E+00** | 0.54 | (0.45-0.62) | **4.54E-33** | **5.74E-33** | -0.36 | (-0.56--0.17) | **3.11E-04** | **5.95E-04** |
| 463 | PI(34:0) | -3.12 | (-3.21--3.03) | **0.00E+00** | **0.00E+00** | -5.31 | (-5.44--5.19) | **0.00E+00** | **0.00E+00** | 1.64 | (1.52-1.77) | **4.45E-112** | **7.80E-112** | 0.37 | (0.17-0.58) | **4.66E-04** | **8.77E-04** |
| 464 | PI(34:1) | -1.63 | (-1.7--1.57) | **0.00E+00** | **0.00E+00** | -4.31 | (-4.38--4.25) | **0.00E+00** | **0.00E+00** | 2.57 | (2.51-2.63) | **0.00E+00** | **0.00E+00** | 0.09 | (-0.05-0.22) | 2.20E-01 | 2.65E-01 |
| 465 | PI(17:0_18:1) | -1.46 | (-1.52--1.4) | **9.61E-290** | **1.18E-288** | -4.28 | (-4.37--4.19) | **0.00E+00** | **0.00E+00** | 2.65 | (2.55-2.75) | **6.49E-305** | **2.94E-304** | 0.03 | (-0.12-0.17) | 7.24E-01 | 7.63E-01 |
| 466 | PI(17:0_18:2) | -1.44 | (-1.51--1.38) | **2.02E-277** | **2.21E-276** | -4.39 | (-4.48--4.29) | **0.00E+00** | **0.00E+00** | 2.64 | (2.53-2.74) | **8.74E-283** | **3.53E-282** | 0.11 | (-0.03-0.25) | 1.16E-01 | 1.48E-01 |
| 467 | PI(18:0_18:1) | -1.69 | (-1.75--1.62) | **0.00E+00** | **6.40E-308** | -6.03 | (-6.1--5.95) | **0.00E+00** | **0.00E+00** | 4.18 | (4.11-4.26) | **0.00E+00** | **0.00E+00** | 0.28 | (0.14-0.43) | **1.73E-04** | **3.46E-04** |
| 468 | PI(36:2) | -1.41 | (-1.46--1.36) | **0.00E+00** | **0.00E+00** | -5.19 | (-5.25--5.13) | **0.00E+00** | **0.00E+00** | 3.61 | (3.55-3.67) | **0.00E+00** | **0.00E+00** | 0.20 | (0.09-0.31) | **6.07E-04** | **1.12E-03** |
| 469 | PI(18:1_18:2) | -1.17 | (-1.23--1.12) | **7.59E-219** | **4.93E-218** | -2.99 | (-3.06--2.93) | **0.00E+00** | **0.00E+00** | 1.74 | (1.68-1.8) | **0.00E+00** | **0.00E+00** | 0.26 | (0.13-0.4) | **8.43E-05** | **1.78E-04** |
| 470 | PI(16:0_20:3) (a) | -1.44 | (-1.5--1.38) | **1.12E-265** | **1.10E-264** | -1.94 | (-2--1.88) | **0.00E+00** | **0.00E+00** | 0.70 | (0.65-0.75) | **1.92E-110** | **3.33E-110** | -0.26 | (-0.39--0.13) | **1.27E-04** | **2.60E-04** |
| 471 | PI(16:0_20:3) (b) | -1.04 | (-1.11--0.97) | **3.19E-156** | **1.21E-155** | -0.72 | (-0.8--0.64) | **1.03E-61** | **1.19E-61** | -0.13 | (-0.21--0.06) | **6.75E-04** | **7.06E-04** | -0.21 | (-0.35--0.07) | **2.83E-03** | **4.76E-03** |
| 472 | PI(16:0_20:4) | -1.05 | (-1.1--0.99) | **3.13E-208** | **1.85E-207** | -1.11 | (-1.17--1.05) | **4.67E-214** | **7.05E-214** | 0.05 | (-0.01-0.1) | 8.69E-02 | 8.88E-02 | -0.09 | (-0.21-0.03) | 1.43E-01 | 1.78E-01 |
| 473 | PI(18:0_20:2) | -1.72 | (-1.79--1.66) | **0.00E+00** | **0.00E+00** | -3.34 | (-3.43--3.25) | **0.00E+00** | **0.00E+00** | 1.58 | (1.49-1.67) | **2.93E-174** | **6.73E-174** | 0.15 | (0.01-0.29) | **3.40E-02** | **4.76E-02** |
| 474 | PI(18:0_20:3) (a) | -0.52 | (-0.56--0.47) | **1.39E-93** | **3.21E-93** | -1.47 | (-1.52--1.42) | **0.00E+00** | **0.00E+00** | 0.80 | (0.75-0.85) | **1.75E-168** | **3.85E-168** | 0.17 | (0.07-0.27) | **5.16E-04** | **9.68E-04** |
| 475 | PI(18:0_20:4) | -0.48 | (-0.52--0.43) | **1.31E-81** | **2.83E-81** | -1.42 | (-1.47--1.38) | **0.00E+00** | **0.00E+00** | 0.78 | (0.73-0.82) | **3.79E-168** | **8.26E-168** | 0.18 | (0.09-0.28) | **2.11E-04** | **4.15E-04** |
| 476 | PI(38:5) (a) | -0.86 | (-0.91--0.81) | **9.79E-177** | **4.61E-176** | -0.73 | (-0.79--0.67) | **1.66E-105** | **2.02E-105** | -0.22 | (-0.27--0.16) | **3.82E-13** | **4.24E-13** | 0.11 | (0-0.22) | **4.52E-02** | 6.17E-02 |
| 477 | PI(38:6) | -1.34 | (-1.4--1.28) | **4.72E-257** | **4.12E-256** | -1.79 | (-1.85--1.73) | **0.00E+00** | **0.00E+00** | 0.50 | (0.44-0.55) | **1.29E-57** | **1.82E-57** | 0.04 | (-0.09-0.17) | 5.35E-01 | 5.98E-01 |
| 478 | PI(18:0_22:4) | -1.26 | (-1.32--1.2) | **1.54E-250** | **1.27E-249** | -1.82 | (-1.88--1.75) | **0.00E+00** | **0.00E+00** | 0.57 | (0.51-0.63) | **2.36E-68** | **3.49E-68** | -0.02 | (-0.14-0.11) | 7.94E-01 | 8.31E-01 |
| 479 | PI(18:0_22:5) (n3) | -0.46 | (-0.52--0.39) | **1.11E-45** | **1.91E-45** | -2.65 | (-2.73--2.58) | **0.00E+00** | **0.00E+00** | 1.88 | (1.81-1.95) | **0.00E+00** | **0.00E+00** | 0.20 | (0.07-0.33) | **1.95E-03** | **3.37E-03** |
| 480 | PI(18:0_22:6) | -1.29 | (-1.35--1.23) | **1.85E-243** | **1.44E-242** | -2.53 | (-2.6--2.47) | **0.00E+00** | **0.00E+00** | 1.18 | (1.12-1.24) | **1.69E-204** | **4.55E-204** | 0.10 | (-0.03-0.23) | 1.33E-01 | 1.67E-01 |

**Supplementary Table 2c.** Comparison of baseline lipids species across four sample groups (antenatal, postnatal, cord blood and 6-year-old child) using paired t-test.

| Studies | | From antenatal to postnatal (381 pairs) | | | | From antenatal to cord blood (445 pairs) | | | | From birth to child (237 pairs) | | | | From child to adult (postnatal) (272 pairs) | | | |
|---|---|---|---|---|---|---|---|---|---|---|---|---|---|---|---|---|---|
| No. | Lipid Species | $log_2FC$ (Postnatal/Antenatal) | 95%CI | p-value | p-value(BH) | $log_2FC$ (CordBlood/Antenatal) | 95%CI | p-value | p-value(BH) | $log_2FC$ (Child/CordBlood) | 95%CI | p-value | p-value(BH) | $log_2FC$ (Adult/Child) | 95%CI | p-value | p-value(BH) |
| 1 | AC(12:0) | 1.56 | (1.46,1.66) | **8.65E-108** | **4.66E-107** | 0.00 | (-0.08,0.08) | 9.62E-01 | 9.64E-01 | 1.81 | (1.69,1.92) | **1.61E-85** | **4.72E-85** | -0.13 | (-0.24,-0.02) | **2.13E-02** | **2.55E-02** |
| 2 | AC(13:0) | 1.12 | (1,1.24) | **2.67E-57** | **5.22E-57** | 0.57 | (0.48,0.66) | **7.33E-32** | **8.10E-32** | -0.24 | (-0.42,-0.07) | **5.58E-03** | **5.84E-03** | 0.85 | (0.68,1.02) | **4.38E-20** | **2.15E-19** |
| 3 | AC(14:0) | 1.06 | (0.99,1.13) | **6.39E-105** | **3.07E-104** | 0.86 | (0.81,0.92) | **9.23E-111** | **1.19E-110** | 0.29 | (0.2,0.38) | **2.05E-09** | **2.37E-09** | -0.03 | (-0.11,0.06) | 5.25E-01 | 5.54E-01 |
| 4 | AC(14:1) | 1.81 | (1.7,1.92) | **8.37E-110** | **4.62E-109** | 0.29 | (0.21,0.38) | **2.22E-11** | **2.36E-11** | 1.82 | (1.7,1.95) | **1.11E-79** | **2.89E-79** | -0.15 | (-0.27,-0.04) | **1.12E-02** | **1.37E-02** |
| 5 | AC(14:2) | 1.65 | (1.54,1.75) | **1.19E-106** | **6.00E-106** | 0.50 | (0.42,0.58) | **1.34E-30** | **1.48E-30** | 1.45 | (1.35,1.56) | **2.58E-73** | **6.06E-73** | -0.20 | (-0.31,-0.08) | **5.88E-04** | **8.09E-04** |
| 6 | AC(16:0) | 0.90 | (0.86,0.95) | **1.93E-129** | **1.54E-128** | 0.67 | (0.63,0.71) | **1.55E-109** | **1.99E-109** | 0.07 | (0,0.13) | **4.36E-02** | **4.53E-02** | 0.19 | (0.14,0.25) | **3.38E-11** | **8.69E-11** |
| 7 | AC(16:1) | 1.31 | (1.23,1.38) | **3.65E-113** | **2.22E-112** | 1.03 | (0.97,1.1) | **4.53E-113** | **5.88E-113** | 0.17 | (0.08,0.27) | **6.52E-04** | **6.89E-04** | 0.20 | (0.11,0.29) | **1.38E-05** | **2.14E-05** |
| 8 | AC(18:0) | 1.00 | (0.95,1.06) | **1.14E-134** | **1.05E-133** | 0.32 | (0.27,0.36) | **4.65E-37** | **5.19E-37** | 0.61 | (0.54,0.67) | **3.01E-44** | **4.88E-44** | 0.07 | (0.01,0.14) | **2.00E-02** | **2.40E-02** |
| 9 | AC(18:1) | 1.15 | (1.09,1.22) | **4.29E-118** | **2.90E-117** | -0.06 | (-0.12,0) | 6.95E-02 | 7.11E-02 | 1.06 | (0.98,1.14) | **9.58E-72** | **2.19E-71** | 0.24 | (0.17,0.3) | **4.75E-12** | **1.33E-11** |
| 10 | AC(18:2) | 1.08 | (1.02,1.14) | **1.61E-118** | **1.10E-117** | 0.76 | (0.7,0.81) | **2.48E-91** | **3.03E-91** | 0.15 | (0.08,0.22) | **6.27E-05** | **6.75E-05** | 0.23 | (0.17,0.3) | **3.63E-12** | **1.03E-11** |
| 11 | CE(14:0) | -0.49 | (-0.55,-0.43) | **7.19E-45** | **1.25E-44** | -0.49 | (-0.54,-0.44) | **2.65E-57** | **3.06E-57** | 0.01 | (-0.08,0.09) | 8.78E-01 | 8.78E-01 | -0.02 | (-0.1,0.06) | 6.22E-01 | 6.42E-01 |
| 12 | CE(16:1) | -0.82 | (-0.9,-0.75) | **2.64E-71** | **6.44E-71** | 0.16 | (0.09,0.23) | **1.41E-05** | **1.47E-05** | -1.21 | (-1.3,-1.12) | **5.04E-75** | **1.23E-74** | 0.26 | (0.18,0.34) | **5.49E-10** | **1.26E-09** |
| 13 | CE(18:0) | 0.17 | (0.13,0.21) | **1.80E-13** | **2.35E-13** | 0.54 | (0.48,0.6) | **2.50E-59** | **2.90E-59** | -0.49 | (-0.57,-0.41) | **9.54E-28** | **1.32E-27** | 0.15 | (0.08,0.22) | **6.24E-05** | **9.29E-05** |
| 14 | CE(18:1) | -0.31 | (-0.33,-0.28) | **2.04E-76** | **5.43E-76** | -0.91 | (-0.94,-0.87) | **1.53E-183** | **2.60E-183** | 0.53 | (0.49,0.58) | **1.05E-59** | **1.98E-59** | 0.08 | (0.04,0.11) | **1.36E-05** | **2.12E-05** |
| 15 | CE(18:2) | -0.15 | (-0.18,-0.12) | **2.75E-21** | **3.87E-21** | -2.75 | (-2.78,-2.71) | **0.00E+00** | **0.00E+00** | 2.41 | (2.36,2.46) | **4.55E-189** | **4.13E-187** | 0.20 | (0.15,0.24) | **1.18E-14** | **4.06E-14** |
| 16 | CE(18:3) | 0.05 | (-0.02,0.11) | 1.41E-01 | 1.50E-01 | -2.21 | (-2.28,-2.14) | **2.80E-224** | **6.81E-224** | 2.22 | (2.13,2.31) | **7.62E-123** | **4.69E-122** | 0.08 | (-0.01,0.16) | 9.53E-02 | 1.08E-01 |
| 17 | CE(20:1) | 0.00 | (-0.05,0.04) | 8.32E-01 | 8.39E-01 | 0.06 | (-0.01,0.12) | 7.74E-02 | 7.91E-02 | -0.15 | (-0.24,-0.07) | **4.92E-04** | **5.23E-04** | 0.14 | (0.08,0.21) | **2.43E-05** | **3.70E-05** |
| 18 | CE(20:2) | -0.50 | (-0.54,-0.47) | **3.38E-95** | **1.31E-94** | -0.54 | (-0.59,-0.49) | **6.59E-72** | **7.77E-72** | -0.14 | (-0.21,-0.08) | **2.66E-05** | **2.87E-05** | 0.18 | (0.13,0.23) | **3.39E-12** | **9.75E-12** |
| 19 | CE(20:4) | 0.50 | (0.45,0.55) | **1.96E-64** | **4.33E-64** | 0.33 | (0.28,0.38) | **2.48E-30** | **2.72E-30** | -0.03 | (-0.11,0.04) | 3.45E-01 | 3.50E-01 | 0.14 | (0.07,0.21) | **8.96E-05** | **1.32E-04** |
| 20 | CE(20:5) | 1.06 | (0.95,1.17) | **1.15E-56** | **2.25E-56** | -1.49 | (-1.59,-1.4) | **9.14E-117** | **1.21E-116** | 1.91 | (1.75,2.07) | **2.41E-64** | **4.82E-64** | 0.60 | (0.45,0.76) | **1.79E-13** | **5.70E-13** |
| 21 | CE(22:5) (n6) | -0.26 | (-0.31,-0.21) | **9.85E-25** | **1.44E-24** | -0.49 | (-0.55,-0.44) | **1.16E-48** | **1.32E-48** | 0.30 | (0.22,0.38) | **3.75E-12** | **4.48E-12** | -0.09 | (-0.15,-0.03) | **2.03E-03** | **2.64E-03** |
| 22 | CE(22:6) | -0.36 | (-0.42,-0.3) | **2.45E-26** | **3.59E-26** | -1.87 | (-1.94,-1.81) | **1.67E-204** | **3.25E-204** | 0.89 | (0.78,1.01) | **2.08E-39** | **3.23E-39** | 0.52 | (0.43,0.61) | **8.74E-25** | **6.66E-24** |
| 23 | CE(24:5) | -1.07 | (-1.13,-1) | **1.69E-111** | **9.87E-111** | 1.11 | (1.03,1.19) | **2.11E-94** | **2.63E-94** | -2.08 | (-2.21,-1.95) | **8.80E-88** | **2.69E-87** | -0.07 | (-0.16,0.02) | 1.16E-01 | 1.29E-01 |
| 24 | CE(24:6) | -0.83 | (-0.9,-0.76) | **1.52E-76** | **4.08E-76** | -0.47 | (-0.55,-0.39) | **6.60E-26** | **7.21E-26** | -0.21 | (-0.33,-0.09) | **5.12E-04** | **5.42E-04** | -0.12 | (-0.21,-0.02) | **1.40E-02** | **1.71E-02** |
| 25 | DG(16:0_16:1) | -1.57 | (-1.67,-1.46) | **1.88E-102** | **8.67E-102** | -2.08 | (-2.17,-1.99) | **1.08E-165** | **1.68E-165** | 0.24 | (0.13,0.35) | **3.04E-05** | **3.28E-05** | 0.31 | (0.18,0.43) | **1.23E-06** | **2.10E-06** |
| 26 | DG(14:0_18:2) | -1.65 | (-1.78,-1.53) | **4.04E-89** | **1.33E-88** | -3.76 | (-3.88,-3.64) | **1.31E-222** | **3.09E-222** | 1.74 | (1.56,1.91) | **9.07E-51** | **1.59E-50** | 0.40 | (0.22,0.58) | **1.52E-05** | **2.34E-05** |
| 27 | DG(16:0_18:1) | -1.60 | (-1.68,-1.52) | **1.25E-138** | **1.25E-137** | -3.20 | (-3.28,-3.12) | **2.00E-269** | **8.75E-269** | 1.42 | (1.31,1.54) | **2.40E-66** | **4.96E-66** | 0.27 | (0.15,0.39) | **1.29E-05** | **2.01E-05** |
| 28 | DG(16:1_18:1) | -1.13 | (-1.22,-1.04) | **1.77E-77** | **4.84E-77** | -2.34 | (-2.43,-2.26) | **5.82E-195** | **1.06E-194** | 0.84 | (0.71,0.97) | **9.89E-29** | **1.39E-28** | 0.45 | (0.32,0.58) | **6.19E-11** | **1.56E-10** |
| 29 | DG(16:0_18:2) | -1.48 | (-1.55,-1.4) | **3.77E-136** | **3.62E-135** | -3.65 | (-3.74,-3.56) | **4.47E-266** | **1.83E-265** | 1.75 | (1.63,1.88) | **5.73E-76** | **1.44E-75** | 0.47 | (0.35,0.58) | **3.00E-14** | **1.00E-13** |
| 30 | DG(18:0_18:1) | -0.99 | (-1.07,-0.92) | **2.51E-90** | **8.80E-90** | -2.55 | (-2.62,-2.47) | **6.14E-238** | **1.78E-237** | 1.38 | (1.27,1.49) | **1.26E-66** | **2.62E-66** | 0.25 | (0.13,0.36) | **3.62E-05** | **5.47E-05** |
| 31 | DG(18:1_18:1) | -0.92 | (-0.99,-0.84) | **2.19E-81** | **6.40E-81** | -3.20 | (-3.28,-3.13) | **7.35E-274** | **3.43E-273** | 2.10 | (1.99,2.21) | **8.18E-103** | **3.22E-102** | 0.27 | (0.16,0.38) | **1.65E-06** | **2.81E-06** |
| 32 | DG(18:0_18:2) | -1.09 | (-1.17,-1.01) | **1.66E-93** | **6.21E-93** | -2.88 | (-2.97,-2.78) | **8.60E-213** | **1.81E-212** | 1.51 | (1.38,1.64) | **9.88E-60** | **1.87E-59** | 0.37 | (0.25,0.5) | **1.55E-08** | **3.12E-08** |
| 33 | DG(18:1_18:2) | -0.83 | (-0.9,-0.76) | **5.07E-75** | **1.31E-74** | -3.71 | (-3.8,-3.63) | **2.12E-276** | **1.03E-275** | 2.51 | (2.39,2.63) | **4.26E-112** | **2.07E-111** | 0.48 | (0.37,0.58) | **5.76E-17** | **2.47E-16** |
| 34 | DG(18:2_18:2) | -0.72 | (-0.8,-0.63) | **2.70E-49** | **4.91E-49** | -4.05 | (-4.16,-3.94) | **1.32E-248** | **4.42E-248** | 2.75 | (2.61,2.89) | **1.76E-103** | **7.05E-103** | 0.70 | (0.57,0.82) | **5.67E-23** | **3.63E-22** |
| 35 | DG(18:1_18:3) | -0.50 | (-0.58,-0.42) | **1.18E-30** | **1.83E-30** | -3.17 | (-3.26,-3.08) | **3.43E-238** | **9.97E-238** | 2.32 | (2.18,2.46) | **1.75E-90** | **5.63E-90** | 0.43 | (0.31,0.56) | **2.54E-11** | **6.60E-11** |
| 36 | DG(16:0_20:4) | -1.57 | (-1.66,-1.47) | **1.95E-116** | **1.28E-115** | -1.40 | (-1.49,-1.32) | **1.44E-117** | **1.92E-117** | -0.54 | (-0.66,-0.41) | **1.29E-15** | **1.59E-15** | 0.33 | (0.21,0.46) | **2.60E-07** | **4.74E-07** |
| 37 | DG(18:1_20:3) | -0.93 | (-1,-0.85) | **8.58E-84** | **2.66E-83** | -1.05 | (-1.12,-0.97) | **4.12E-97** | **5.16E-97** | 0.22 | (0.11,0.33) | **1.06E-04** | **1.14E-04** | -0.06 | (-0.17,0.04) | 2.40E-01 | 2.59E-01 |
| 38 | DG(18:1_20:4) | -0.75 | (-0.82,-0.69) | **1.22E-75** | **3.22E-75** | -1.57 | (-1.64,-1.49) | **2.89E-159** | **4.39E-159** | 0.45 | (0.34,0.55) | **1.65E-15** | **2.03E-15** | 0.34 | (0.24,0.43) | **1.69E-11** | **4.41E-11** |
| 39 | DG(16:0_22:5) | -1.35 | (-1.43,-1.26) | **1.73E-103** | **8.15E-103** | -0.97 | (-1.06,-0.88) | **2.23E-71** | **2.62E-71** | -0.69 | (-0.81,-0.57) | **5.43E-23** | **7.18E-23** | 0.28 | (0.16,0.41) | **1.07E-05** | **1.69E-05** |
| 40 | DG(18:2_20:4) | -0.75 | (-0.82,-0.68) | **1.10E-64** | **2.44E-64** | -1.89 | (-1.98,-1.8) | **2.38E-153** | **3.57E-153** | 0.63 | (0.49,0.76) | **2.99E-17** | **3.75E-17** | 0.53 | (0.42,0.64) | **1.81E-18** | **8.43E-18** |
| 41 | DG(16:0_22:6) | -1.68 | (-1.79,-1.58) | **1.73E-107** | **9.12E-107** | -1.31 | (-1.41,-1.22) | **1.09E-94** | **1.36E-94** | -1.00 | (-1.15,-0.84) | **4.87E-28** | **6.78E-28** | 0.69 | (0.53,0.85) | **1.10E-15** | **4.27E-15** |
| 42 | DG(18:1_22:5) | -0.92 | (-1,-0.84) | **2.68E-71** | **6.50E-71** | -0.93 | (-1.01,-0.85) | **1.04E-72** | **1.23E-72** | -0.16 | (-0.3,-0.03) | **1.42E-02** | **1.48E-02** | 0.18 | (0.08,0.29) | **1.02E-03** | **1.38E-03** |
| 43 | DG(18:1_22:6) | -0.92 | (-1.01,-0.82) | **4.96E-58** | **9.84E-58** | -1.24 | (-1.33,-1.15) | **1.66E-99** | **2.09E-99** | -0.33 | (-0.46,-0.2) | **7.23E-07** | **8.02E-07** | 0.71 | (0.57,0.85) | **1.48E-20** | **7.62E-20** |
| 44 | DG(18:2_22:6) | -0.92 | (-1.02,-0.82) | **8.85E-52** | **1.66E-51** | -1.59 | (-1.69,-1.48) | **5.74E-105** | **7.30E-105** | -0.25 | (-0.4,-0.11) | **7.49E-04** | **7.90E-04** | 0.91 | (0.76,1.06) | **8.68E-26** | **7.57E-25** |
| 45 | TG(48:0) [NL-18:0] | -1.12 | (-1.24,-0.99) | **1.32E-51** | **2.47E-51** | -1.73 | (-1.83,-1.64) | **2.53E-135** | **3.51E-135** | 0.54 | (0.38,0.7) | **1.60E-10** | **1.87E-10** | 0.12 | (-0.06,0.3) | 1.77E-01 | 1.94E-01 |
| 46 | TG(48:1) [NL-16:1] | -2.19 | (-2.32,-2.06) | **3.03E-114** | **1.89E-113** | -2.58 | (-2.69,-2.46) | **1.31E-163** | **2.02E-163** | -0.05 | (-0.22,0.13) | 6.01E-01 | 6.03E-01 | 0.49 | (0.32,0.66) | **5.12E-08** | **9.87E-08** |
| 47 | TG(48:1) [NL-18:1] | -1.86 | (-1.99,-1.74) | **1.23E-98** | **5.01E-98** | -3.57 | (-3.68,-3.47) | **5.89E-231** | **1.56E-230** | 1.41 | (1.22,1.59) | **4.97E-36** | **7.51E-36** | 0.36 | (0.17,0.56) | **3.06E-04** | **4.32E-04** |
| 48 | TG(48:2) [NL-14:0] | -2.08 | (-2.21,-1.94) | **5.89E-105** | **2.86E-104** | -3.78 | (-3.89,-3.66) | **3.72E-226** | **9.41E-226** | 1.15 | (0.97,1.34) | **1.53E-26** | **2.09E-26** | 0.64 | (0.44,0.85) | **1.42E-09** | **3.12E-09** |
| 49 | TG(48:2) [NL-14:1] | -1.75 | (-1.88,-1.62) | **1.26E-90** | **4.48E-90** | -3.62 | (-3.74,-3.51) | **6.78E-224** | **1.64E-223** | 1.62 | (1.44,1.79) | **3.79E-47** | **6.42E-47** | 0.34 | (0.16,0.52) | **3.04E-04** | **4.31E-04** |
| 50 | TG(48:2) [NL-16:1] | -1.92 | (-2.06,-1.78) | **4.29E-90** | **1.47E-89** | -2.46 | (-2.58,-2.33) | **2.48E-143** | **3.54E-143** | -0.11 | (-0.29,0.07) | 2.16E-01 | 2.20E-01 | 0.71 | (0.53,0.9) | **1.96E-13** | **6.18E-13** |
| 51 | TG(48:2) [NL-18:2] | -2.09 | (-2.22,-1.96) | **3.85E-105** | **1.89E-104** | -4.23 | (-4.35,-4.11) | **2.98E-238** | **8.73E-238** | 1.58 | (1.39,1.77) | **2.67E-40** | **4.20E-40** | 0.66 | (0.45,0.86) | **1.07E-09** | **2.38E-09** |

| | | | | | | | | | | | | | | | | | |
|---|---|---|---|---|---|---|---|---|---|---|---|---|---|---|---|---|---|
| 52 | TG(48:3) [NL-14:0] | -1.67 | (-1.8,-1.54) | 3.16E-81 | 9.19E-81 | -3.25 | (-3.37,-3.13) | 1.11E-190 | 1.94E-190 | 0.96 | (0.77,1.15) | 2.28E-19 | 2.95E-19 | 0.74 | (0.55,0.94) | 1.24E-12 | 3.69E-12 |
| 53 | TG(48:3) [NL-16:1] | -1.51 | (-1.63,-1.39) | 1.39E-80 | 4.00E-80 | -2.19 | (-2.31,-2.08) | 1.33E-141 | 1.87E-141 | 0.10 | (-0.06,0.25) | 2.12E-01 | 2.16E-01 | 0.69 | (0.53,0.85) | 3.13E-15 | 1.15E-14 |
| 54 | TG(48:3) [NL-18:3] | -1.70 | (-1.85,-1.55) | 2.15E-71 | 5.28E-71 | -3.30 | (-3.42,-3.18) | 1.31E-197 | 2.43E-197 | 1.20 | (1.02,1.38) | 2.93E-30 | 4.19E-30 | 0.47 | (0.26,0.68) | 1.86E-05 | 2.85E-05 |
| 55 | TG(49:1) [NL-17:1] | -1.66 | (-1.75,-1.57) | 3.70E-127 | 2.87E-126 | -2.31 | (-2.4,-2.23) | 1.56E-198 | 2.90E-198 | 0.36 | (0.25,0.47) | 5.94E-10 | 6.93E-10 | 0.33 | (0.21,0.44) | 1.11E-07 | 2.09E-07 |
| 56 | TG(50:1) [NL-14:0] | -1.10 | (-1.21,-0.98) | 3.54E-58 | 7.13E-58 | -1.89 | (-1.98,-1.8) | 2.54E-149 | 3.72E-149 | 0.72 | (0.57,0.87) | 1.82E-17 | 2.29E-17 | 0.16 | (-0.01,0.32) | 6.14E-02 | 7.02E-02 |
| 57 | TG(50:1) [NL-16:0] | -1.59 | (-1.67,-1.51) | 4.00E-132 | 3.31E-131 | -2.80 | (-2.88,-2.71) | 4.08E-233 | 1.10E-232 | 0.99 | (0.86,1.12) | 2.04E-35 | 3.05E-35 | 0.27 | (0.15,0.38) | 1.04E-05 | 1.65E-05 |
| 58 | TG(50:1) [NL-18:1] | -1.55 | (-1.63,-1.47) | 1.07E-130 | 8.68E-130 | -2.80 | (-2.88,-2.72) | 2.94E-237 | 8.45E-237 | 1.00 | (0.87,1.13) | 3.94E-37 | 5.98E-37 | 0.30 | (0.18,0.41) | 5.63E-07 | 9.93E-07 |
| 59 | TG(50:2) [NL-14:0] | -0.94 | (-1.04,-0.85) | 9.76E-64 | 2.13E-63 | -2.91 | (-3.01,-2.82) | 4.97E-219 | 1.14E-218 | 1.78 | (1.62,1.93) | 1.74E-61 | 3.38E-61 | 0.30 | (0.15,0.44) | 7.21E-05 | 1.07E-04 |
| 60 | TG(50:2) [NL-16:1] | -1.16 | (-1.25,-1.07) | 2.31E-82 | 6.87E-82 | -2.22 | (-2.32,-2.12) | 6.11E-163 | 9.33E-163 | 0.65 | (0.5,0.81) | 1.49E-14 | 1.81E-14 | 0.49 | (0.36,0.62) | 1.54E-12 | 4.57E-12 |
| 61 | TG(50:2) [NL-18:1] | -1.03 | (-1.12,-0.95) | 1.63E-77 | 4.47E-77 | -2.41 | (-2.51,-2.31) | 1.84E-179 | 3.04E-179 | 1.01 | (0.85,1.17) | 2.19E-28 | 3.05E-28 | 0.48 | (0.35,0.61) | 3.97E-12 | 1.12E-11 |
| 62 | TG(50:2) [NL-18:2] | -1.49 | (-1.56,-1.41) | 2.63E-132 | 2.21E-131 | -3.35 | (-3.46,-3.25) | 6.61E-222 | 1.55E-221 | 1.46 | (1.31,1.61) | 4.70E-48 | 8.03E-48 | 0.48 | (0.37,0.6) | 5.25E-15 | 1.88E-14 |
| 63 | TG(50:3) [NL-14:0] | -1.31 | (-1.41,-1.21) | 8.08E-85 | 2.52E-84 | -3.90 | (-4.01,-3.79) | 5.39E-244 | 1.72E-243 | 2.06 | (1.88,2.23) | 3.41E-62 | 6.69E-62 | 0.66 | (0.5,0.83) | 7.45E-14 | 2.43E-13 |
| 64 | TG(50:3) [NL-14:1] | -1.00 | (-1.09,-0.91) | 3.68E-72 | 9.19E-72 | -2.95 | (-3.03,-2.86) | 3.00E-236 | 8.41E-236 | 1.80 | (1.66,1.94) | 1.48E-67 | 3.14E-67 | 0.26 | (0.13,0.39) | 1.07E-04 | 1.56E-04 |
| 65 | TG(50:3) [NL-16:1] | -1.38 | (-1.48,-1.28) | 1.16E-89 | 3.90E-89 | -2.87 | (-2.97,-2.76) | 6.06E-193 | 1.08E-192 | 0.71 | (0.54,0.88) | 8.56E-15 | 1.04E-14 | 0.89 | (0.74,1.04) | 3.02E-26 | 2.78E-25 |
| 66 | TG(50:3) [NL-18:2] | -1.30 | (-1.39,-1.21) | 3.03E-91 | 1.10E-90 | -3.40 | (-3.51,-3.29) | 4.95E-217 | 1.10E-216 | 1.36 | (1.19,1.53) | 9.86E-39 | 1.51E-38 | 0.85 | (0.7,1) | 5.05E-25 | 3.97E-24 |
| 67 | TG(50:3) [NL-18:3] | -1.69 | (-1.8,-1.58) | 3.97E-103 | 1.85E-102 | -3.48 | (-3.58,-3.37) | 3.71E-222 | 8.73E-222 | 1.27 | (1.11,1.43) | 7.01E-37 | 1.06E-36 | 0.60 | (0.43,0.76) | 7.46E-12 | 2.03E-11 |
| 68 | TG(50:4) [NL-14:0] | -1.29 | (-1.4,-1.18) | 1.11E-76 | 3.02E-76 | -3.69 | (-3.81,-3.58) | 1.03E-219 | 2.37E-219 | 1.72 | (1.54,1.9) | 2.94E-50 | 5.15E-50 | 0.78 | (0.61,0.95) | 3.21E-17 | 1.41E-16 |
| 69 | TG(50:4) [NL-18:3] | -1.15 | (-1.26,-1.03) | 1.78E-59 | 3.65E-59 | -2.75 | (-2.86,-2.63) | 5.08E-177 | 8.29E-177 | 1.01 | (0.84,1.17) | 9.55E-27 | 1.31E-26 | 0.69 | (0.52,0.85) | 3.91E-15 | 1.42E-14 |
| 70 | TG(50:4) [NL-20:4] | -2.14 | (-2.29,-1.98) | 1.99E-92 | 7.35E-92 | -1.96 | (-2.09,-1.82) | 1.09E-102 | 1.37E-102 | -0.71 | (-0.9,-0.52) | 2.84E-12 | 3.41E-12 | 0.49 | (0.28,0.7) | 5.47E-06 | 8.91E-06 |
| 71 | TG(51:1) [NL-17:0] | -1.44 | (-1.53,-1.35) | 1.12E-108 | 6.12E-108 | -2.75 | (-2.84,-2.66) | 4.57E-216 | 1.01E-215 | 1.16 | (1.02,1.3) | 2.79E-40 | 4.37E-40 | 0.22 | (0.09,0.36) | 1.37E-03 | 1.83E-03 |
| 72 | TG(51:2) [NL-15:0] | -0.97 | (-1.05,-0.9) | 1.69E-83 | 5.17E-83 | -3.41 | (-3.5,-3.32) | 7.89E-253 | 2.77E-252 | 2.16 | (2.02,2.31) | 1.99E-80 | 5.24E-80 | 0.36 | (0.23,0.49) | 6.36E-08 | 1.22E-07 |
| 73 | TG(51:2) [NL-17:0] | -1.35 | (-1.43,-1.26) | 1.56E-106 | 7.71E-106 | -3.12 | (-3.22,-3.03) | 7.23E-230 | 1.90E-229 | 1.37 | (1.22,1.52) | 8.35E-48 | 1.42E-47 | 0.49 | (0.35,0.62) | 1.30E-11 | 3.43E-11 |
| 74 | TG(51:2) [NL-17:1] | -1.27 | (-1.35,-1.19) | 1.69E-107 | 9.04E-107 | -2.82 | (-2.9,-2.74) | 8.48E-239 | 2.50E-238 | 1.16 | (1.03,1.29) | 6.62E-47 | 1.12E-46 | 0.45 | (0.33,0.58) | 6.09E-12 | 1.68E-11 |
| 75 | TG(52:1) [NL-18:0] | -1.39 | (-1.49,-1.3) | 6.66E-102 | 2.99E-101 | -2.67 | (-2.76,-2.57) | 8.29E-201 | 1.57E-200 | 1.11 | (0.95,1.26) | 4.36E-33 | 6.46E-33 | 0.26 | (0.12,0.4) | 4.21E-04 | 5.87E-04 |
| 76 | TG(52:1) [NL-18:1] | -1.29 | (-1.38,-1.2) | 1.02E-93 | 3.85E-93 | -2.54 | (-2.64,-2.45) | 1.28E-195 | 2.35E-195 | 1.04 | (0.89,1.19) | 3.67E-31 | 5.32E-31 | 0.31 | (0.17,0.45) | 2.20E-05 | 3.37E-05 |
| 77 | TG(52:2) [NL-16:0] | -0.64 | (-0.69,-0.59) | 8.03E-83 | 2.42E-82 | -2.53 | (-2.6,-2.46) | 7.54E-240 | 2.26E-239 | 1.73 | (1.63,1.84) | 1.68E-88 | 5.25E-88 | 0.24 | (0.17,0.32) | 1.86E-09 | 4.03E-09 |
| 78 | TG(52:2) [NL-18:2] | -0.96 | (-1.03,-0.89) | 1.72E-85 | 5.39E-85 | -2.26 | (-2.35,-2.17) | 3.21E-176 | 5.21E-176 | 0.99 | (0.85,1.13) | 1.65E-31 | 2.40E-31 | 0.40 | (0.29,0.51) | 3.90E-11 | 9.97E-11 |
| 79 | TG(52:3) [NL-16:1] | -0.43 | (-0.5,-0.36) | 3.45E-29 | 5.28E-29 | -1.89 | (-1.98,-1.8) | 3.00E-152 | 4.49E-152 | 1.13 | (0.99,1.27) | 6.21E-39 | 9.56E-39 | 0.43 | (0.33,0.53) | 6.46E-15 | 2.30E-14 |
| 80 | TG(52:3) [NL-18:2] | -0.65 | (-0.7,-0.6) | 2.54E-80 | 7.26E-80 | -3.37 | (-3.47,-3.27) | 1.76E-233 | 4.80E-233 | 2.35 | (2.21,2.5) | 2.99E-89 | 9.51E-89 | 0.48 | (0.39,0.56) | 4.55E-24 | 3.31E-23 |
| 81 | TG(52:4) [NL-16:1] | -0.79 | (-0.87,-0.7) | 1.01E-54 | 1.95E-54 | -2.99 | (-3.1,-2.88) | 4.12E-201 | 7.81E-201 | 1.51 | (1.34,1.67) | 1.37E-45 | 2.27E-45 | 0.84 | (0.7,0.97) | 2.13E-28 | 2.33E-27 |
| 82 | TG(52:4) [NL-18:2] | -0.96 | (-1.04,-0.88) | 1.30E-76 | 3.52E-76 | -4.16 | (-4.28,-4.04) | 5.22E-239 | 1.55E-238 | 2.42 | (2.25,2.6) | 2.85E-74 | 6.83E-74 | 0.89 | (0.76,1.02) | 2.49E-31 | 4.43E-30 |
| 83 | TG(52:4) [NL-18:3] | -0.90 | (-0.98,-0.82) | 3.92E-69 | 9.18E-69 | -3.71 | (-3.81,-3.61) | 3.24E-249 | 1.10E-248 | 2.30 | (2.14,2.45) | 3.13E-80 | 8.21E-80 | 0.61 | (0.48,0.74) | 3.52E-17 | 1.54E-16 |
| 84 | TG(52:5) [NL-18:3] | -0.93 | (-1.03,-0.84) | 4.83E-58 | 9.62E-58 | -3.61 | (-3.72,-3.5) | 1.71E-226 | 4.35E-226 | 2.02 | (1.86,2.18) | 6.33E-68 | 1.34E-67 | 0.79 | (0.64,0.93) | 1.71E-22 | 1.03E-21 |
| 85 | TG(52:5) [NL-20:4] | -1.62 | (-1.74,-1.5) | 6.23E-88 | 2.02E-87 | -1.14 | (-1.27,-1.02) | 2.91E-56 | 3.35E-56 | -1.23 | (-1.41,-1.06) | 1.16E-32 | 1.72E-32 | 0.74 | (0.58,0.9) | 5.81E-17 | 2.47E-16 |
| 86 | TG(52:5) [NL-20:5] | -1.24 | (-1.35,-1.12) | 1.66E-65 | 3.72E-65 | -2.06 | (-2.17,-1.95) | 7.10E-138 | 9.90E-138 | -0.12 | (-0.29,0.04) | 1.30E-01 | 1.34E-01 | 0.94 | (0.77,1.1) | 4.24E-23 | 2.79E-22 |
| 87 | TG(53:2) [NL-18:1] | -0.76 | (-0.83,-0.69) | 4.13E-67 | 9.48E-67 | -2.65 | (-2.73,-2.57) | 3.81E-225 | 9.47E-225 | 1.78 | (1.66,1.9) | 7.56E-81 | 2.03E-80 | 0.20 | (0.09,0.31) | 4.85E-04 | 6.73E-04 |
| 88 | TG(54:1) [NL-18:1] | -0.86 | (-0.93,-0.78) | 1.04E-71 | 2.59E-71 | -2.43 | (-2.52,-2.34) | 1.52E-203 | 2.91E-203 | 1.42 | (1.28,1.57) | 8.25E-52 | 1.47E-51 | 0.20 | (0.09,0.32) | 4.58E-04 | 6.38E-04 |
| 89 | TG(54:2) [NL-18:0] | -0.72 | (-0.79,-0.65) | 1.90E-59 | 3.89E-59 | -2.41 | (-2.5,-2.33) | 1.71E-202 | 3.27E-202 | 1.47 | (1.33,1.6) | 2.40E-56 | 4.46E-56 | 0.32 | (0.21,0.44) | 2.10E-08 | 4.18E-08 |
| 90 | TG(54:2) [NL-20:1] | -1.83 | (-1.91,-1.76) | 1.72E-156 | 2.58E-155 | -3.98 | (-4.06,-3.89) | 1.09E-291 | 7.60E-291 | 2.10 | (1.97,2.22) | 8.66E-91 | 2.83E-90 | 0.12 | (0.01,0.24) | 3.27E-02 | 3.87E-02 |
| 91 | TG(54:3) [NL-18:1] | -0.22 | (-0.28,-0.16) | 1.17E-12 | 1.50E-12 | -2.02 | (-2.1,-1.95) | 1.91E-194 | 3.48E-194 | 1.77 | (1.67,1.88) | 1.71E-88 | 5.30E-88 | 0.12 | (0.04,0.2) | 5.27E-03 | 6.62E-03 |
| 92 | TG(54:3) [NL-18:2] | -0.39 | (-0.45,-0.34) | 7.37E-36 | 1.19E-35 | -2.45 | (-2.53,-2.36) | 9.98E-208 | 2.00E-207 | 1.69 | (1.57,1.81) | 1.22E-75 | 3.03E-75 | 0.44 | (0.35,0.52) | 6.76E-20 | 3.28E-19 |
| 93 | TG(54:4) [NL-18:2] | -0.26 | (-0.31,-0.2) | 1.93E-16 | 2.61E-16 | -3.06 | (-3.16,-2.96) | 4.07E-215 | 8.88E-215 | 2.57 | (2.43,2.72) | 7.04E-95 | 2.45E-94 | 0.35 | (0.26,0.45) | 5.54E-12 | 1.54E-11 |
| 94 | TG(54:4) [NL-20:3] | -1.22 | (-1.29,-1.14) | 7.36E-112 | 4.36E-111 | -1.51 | (-1.6,-1.42) | 7.19E-117 | 9.56E-117 | 0.12 | (-0.02,0.26) | 8.57E-02 | 8.82E-02 | 0.18 | (0.07,0.29) | 1.42E-03 | 1.88E-03 |
| 95 | TG(54:5) [NL-18:3] | -0.51 | (-0.6,-0.42) | 1.80E-26 | 2.65E-26 | -3.44 | (-3.54,-3.34) | 4.31E-237 | 1.22E-236 | 2.53 | (2.38,2.68) | 2.84E-91 | 9.42E-91 | 0.53 | (0.4,0.66) | 1.21E-13 | 3.87E-13 |
| 96 | TG(54:5) [NL-20:4] | -0.94 | (-1.01,-0.87) | 8.89E-90 | 3.00E-89 | -1.54 | (-1.64,-1.44) | 2.69E-109 | 3.45E-109 | 0.07 | (-0.09,0.22) | 3.99E-01 | 4.04E-01 | 0.57 | (0.46,0.67) | 3.94E-23 | 2.63E-22 |
| 97 | TG(54:6) [NL-18:3] | -0.61 | (-0.71,-0.51) | 2.15E-27 | 3.25E-27 | -3.66 | (-3.78,-3.55) | 2.03E-223 | 4.90E-223 | 2.50 | (2.35,2.66) | 1.01E-85 | 2.99E-85 | 0.67 | (0.52,0.82) | 2.65E-16 | 1.06E-15 |
| 98 | TG(54:6) [NL-20:4] | -1.21 | (-1.29,-1.13) | 2.67E-102 | 1.22E-101 | -1.93 | (-2.04,-1.82) | 3.01E-129 | 4.13E-129 | -0.15 | (-0.32,0.01) | 6.74E-02 | 6.97E-02 | 0.87 | (0.75,0.99) | 9.35E-35 | 2.81E-33 |
| 99 | TG(54:6) [NL-20:5] | -0.54 | (-0.65,-0.44) | 1.28E-21 | 1.81E-21 | -2.32 | (-2.43,-2.21) | 4.51E-152 | 6.73E-152 | 0.69 | (0.51,0.87) | 3.92E-13 | 4.72E-13 | 1.08 | (0.92,1.25) | 6.44E-30 | 8.59E-29 |
| 100 | TG(54:6) [NL-22:6] | -2.12 | (-2.23,-2) | 1.13E-125 | 8.31E-125 | -1.54 | (-1.66,-1.43) | 1.49E-92 | 1.83E-92 | -1.51 | (-1.68,-1.33) | 2.02E-43 | 3.25E-43 | 0.93 | (0.76,1.11) | 1.93E-21 | 1.10E-20 |
| 101 | TG(54:7) [NL-20:5] | -0.51 | (-0.62,-0.4) | 3.02E-18 | 4.14E-18 | -2.09 | (-2.21,-1.98) | 4.67E-131 | 6.44E-131 | 0.41 | (0.23,0.59) | 9.28E-06 | 1.01E-05 | 1.20 | (1.03,1.37) | 1.09E-32 | 2.28E-31 |
| 102 | TG(54:7) [NL-22:6] | -1.58 | (-1.71,-1.46) | 2.59E-82 | 7.66E-82 | -0.62 | (-0.74,-0.51) | 2.05E-23 | 2.23E-23 | -2.05 | (-2.21,-1.88) | 6.70E-66 | 1.37E-65 | 1.14 | (0.96,1.32) | 6.03E-28 | 6.30E-27 |
| 103 | TG(56:6) [NL-20:4] | -0.01 | (-0.05,0.04) | 7.88E-01 | 7.97E-01 | -1.04 | (-1.13,-0.96) | 2.21E-79 | 2.64E-79 | 0.57 | (0.44,0.7) | 3.64E-16 | 4.50E-16 | 0.50 | (0.43,0.56) | 3.60E-35 | 1.24E-33 |
| 104 | TG(56:6) [NL-22:5] | -0.66 | (-0.73,-0.6) | 3.30E-64 | 7.27E-64 | -0.91 | (-1,-0.81) | 2.59E-58 | 3.01E-58 | -0.23 | (-0.37,-0.08) | 1.93E-03 | 2.03E-03 | 0.51 | (0.41,0.61) | 1.86E-20 | 9.50E-20 |
| 105 | TG(56:7) [NL-20:4] | -0.57 | (-0.63,-0.51) | 2.53E-54 | 4.85E-54 | -1.64 | (-1.73,-1.54) | 6.77E-124 | 9.15E-124 | 0.42 | (0.27,0.56) | 1.11E-07 | 1.25E-07 | 0.63 | (0.54,0.73) | 2.51E-30 | 3.54E-29 |
| 106 | TG(56:7) [NL-20:5] | 0.01 | (-0.09,0.11) | 8.47E-01 | 8.52E-01 | -2.00 | (-2.1,-1.9) | 3.43E-149 | 5.01E-149 | 1.07 | (0.91,1.23) | 1.61E-29 | 2.30E-29 | 0.93 | (0.78,1.07) | 1.44E-29 | 1.78E-28 |

| | | | | | | | | | | | | | | | | | |
|---|---|---|---|---|---|---|---|---|---|---|---|---|---|---|---|---|---|
| 107 | TG(56:7) [NL-22:5] | -1.00 | (-1.07,-0.92) | **1.64E-90** | **5.80E-90** | -1.48 | (-1.58,-1.38) | **5.39E-101** | **6.79E-101** | -0.35 | (-0.51,-0.19) | **1.84E-05** | **1.99E-05** | 0.87 | (0.74,1) | **6.37E-32** | **1.27E-30** |
| 108 | TG(56:7) [NL-22:6] | -1.27 | (-1.37,-1.17) | **3.25E-83** | **9.87E-83** | -1.60 | (-1.7,-1.49) | **2.05E-104** | **2.60E-104** | -0.70 | (-0.86,-0.53) | **2.51E-14** | **3.04E-14** | 1.05 | (0.89,1.21) | **7.91E-31** | **1.27E-29** |
| 109 | TG(56:8) [NL-20:4] | -0.73 | (-0.81,-0.66) | **4.09E-59** | **8.32E-59** | -1.06 | (-1.16,-0.96) | **2.00E-70** | **2.34E-70** | -0.35 | (-0.49,-0.21) | **2.88E-06** | **3.16E-06** | 0.65 | (0.54,0.76) | **3.32E-26** | **3.01E-25** |
| 110 | TG(56:8) [NL-20:5] | -0.14 | (-0.24,-0.04) | **7.57E-03** | **8.43E-03** | -1.88 | (-1.98,-1.77) | **6.99E-128** | **9.53E-128** | 0.77 | (0.61,0.93) | **7.65E-18** | **9.69E-18** | 0.97 | (0.82,1.12) | **3.09E-29** | **3.71E-28** |
| 111 | TG(56:8) [NL-22:6] | -1.39 | (-1.49,-1.28) | **5.78E-86** | **1.83E-85** | -1.60 | (-1.72,-1.48) | **6.56E-94** | **8.14E-94** | -1.02 | (-1.2,-0.85) | **3.96E-24** | **5.26E-24** | 1.29 | (1.13,1.46) | **8.66E-39** | **4.63E-37** |
| 112 | TG(56:9) [NL-22:6] | -1.03 | (-1.13,-0.93) | **3.88E-61** | **8.17E-61** | -0.92 | (-1.03,-0.81) | **9.97E-50** | **1.14E-49** | -1.22 | (-1.39,-1.06) | **1.16E-34** | **1.73E-34** | 1.14 | (0.99,1.3) | **7.51E-35** | **2.40E-33** |
| 113 | TG(58:10) [NL-22:6] | -0.87 | (-0.97,-0.77) | **1.72E-50** | **3.18E-50** | -0.81 | (-0.91,-0.7) | **9.21E-43** | **1.04E-42** | -1.12 | (-1.28,-0.96) | **3.00E-31** | **4.37E-31** | 1.08 | (0.92,1.23) | **5.95E-33** | **1.43E-31** |
| 114 | TG(58:8) [NL-22:6] | -0.93 | (-1.03,-0.84) | **5.28E-56** | **1.03E-55** | -1.03 | (-1.13,-0.93) | **7.33E-67** | **8.56E-67** | -0.86 | (-1.02,-0.7) | **5.35E-22** | **7.04E-22** | 1.00 | (0.84,1.15) | **6.19E-30** | **8.49E-29** |
| 115 | TG(58:9) [NL-22:6] | -1.02 | (-1.12,-0.92) | **9.93E-62** | **2.12E-61** | -1.17 | (-1.28,-1.07) | **2.13E-71** | **2.50E-71** | -0.88 | (-1.05,-0.71) | **4.64E-21** | **6.07E-21** | 1.09 | (0.93,1.24) | **7.67E-33** | **1.75E-31** |
| 116 | TG(O-50:1) [NL-16:0] | -0.60 | (-0.67,-0.54) | **2.05E-55** | **3.96E-55** | -3.19 | (-3.25,-3.12) | **1.89E-302** | **1.82E-301** | 2.67 | (2.57,2.78) | **1.63E-128** | **1.24E-127** | -0.06 | (-0.15,0.03) | 1.96E-01 | 2.14E-01 |
| 117 | TG(O-50:1) [NL-17:1] | -1.68 | (-1.77,-1.59) | **1.04E-126** | **7.92E-126** | -2.35 | (-2.43,-2.26) | **2.25E-197** | **4.15E-197** | 0.37 | (0.26,0.48) | **4.23E-10** | **4.95E-10** | 0.33 | (0.21,0.45) | **1.21E-07** | **2.27E-07** |
| 118 | TG(O-50:1) [NL-18:1] | -0.48 | (-0.55,-0.41) | **7.81E-33** | **1.23E-32** | -3.37 | (-3.44,-3.3) | **2.40E-296** | **2.02E-295** | 2.85 | (2.74,2.97) | **6.17E-128** | **4.36E-127** | 0.08 | (-0.02,0.18) | 1.35E-01 | 1.50E-01 |
| 119 | TG(O-50:2) [NL-18:2] | -0.57 | (-0.63,-0.51) | **4.23E-51** | **7.87E-51** | -2.65 | (-2.72,-2.59) | **2.70E-273** | **1.22E-272** | 2.10 | (2,2.2) | **3.78E-110** | **1.71E-109** | 0.10 | (0.01,0.19) | **2.48E-02** | **2.94E-02** |
| 120 | TG(O-52:2) [NL-16:0] | -0.78 | (-0.86,-0.7) | **7.10E-60** | **1.48E-59** | -3.16 | (-3.26,-3.06) | **2.77E-225** | **6.92E-225** | 2.49 | (2.34,2.64) | **2.36E-89** | **7.57E-89** | -0.01 | (-0.13,0.11) | 8.65E-01 | 8.80E-01 |
| 121 | TG(O-52:2) [NL-17:1] | -1.28 | (-1.36,-1.2) | **6.19E-107** | **3.19E-106** | -2.84 | (-2.92,-2.76) | **4.06E-237** | **1.15E-236** | 1.18 | (1.05,1.3) | **4.39E-47** | **7.42E-47** | 0.45 | (0.33,0.58) | **8.57E-12** | **2.28E-11** |
| 122 | TG(O-52:2) [NL-18:1] | -0.66 | (-0.74,-0.58) | **6.29E-45** | **1.10E-44** | -3.10 | (-3.2,-3) | **1.17E-217** | **2.63E-217** | 2.52 | (2.36,2.67) | **1.73E-87** | **5.26E-87** | 0.00 | (-0.11,0.12) | 9.40E-01 | 9.42E-01 |
| 123 | TG(O-54:2) [NL-18:1] | -0.76 | (-0.83,-0.69) | **6.08E-67** | **1.38E-66** | -2.65 | (-2.73,-2.57) | **8.64E-225** | **2.11E-224** | 1.78 | (1.66,1.9) | **1.84E-80** | **4.88E-80** | 0.20 | (0.09,0.31) | **3.43E-04** | **4.82E-04** |
| 124 | TG(O-54:4) [NL-17:1] | -0.70 | (-0.77,-0.62) | **6.88E-51** | **1.27E-50** | -3.07 | (-3.17,-2.96) | **1.93E-209** | **3.91E-209** | 1.80 | (1.65,1.94) | **5.98E-66** | **1.23E-65** | 0.70 | (0.58,0.82) | **2.98E-24** | **2.20E-23** |
| 125 | TG(O-54:4) [NL-18:2] | -0.67 | (-0.75,-0.6) | **1.38E-48** | **2.49E-48** | -3.39 | (-3.51,-3.28) | **1.94E-213** | **4.15E-213** | 2.16 | (1.99,2.32) | **1.64E-70** | **3.63E-70** | 0.69 | (0.56,0.82) | **2.85E-21** | **1.59E-20** |
| 126 | dhCer(d18:0/22:0) | -0.01 | (-0.07,0.05) | 6.90E-01 | 7.05E-01 | 0.02 | (-0.04,0.07) | 6.12E-01 | 6.18E-01 | -0.26 | (-0.34,-0.17) | **2.28E-08** | **2.61E-08** | 0.20 | (0.12,0.27) | **6.86E-07** | **1.20E-06** |
| 127 | dhCer(d18:0/24:0) | 0.05 | (0,0.11) | 5.11E-02 | 5.54E-02 | 0.41 | (0.36,0.47) | **7.53E-38** | **8.43E-38** | -0.48 | (-0.56,-0.4) | **4.01E-26** | **5.47E-26** | 0.09 | (0.02,0.16) | **1.41E-02** | **1.72E-02** |
| 128 | dhCer(d18:0/24:1) | -0.32 | (-0.38,-0.26) | **1.44E-22** | **2.05E-22** | -0.01 | (-0.08,0.07) | 8.42E-01 | 8.46E-01 | -0.65 | (-0.75,-0.55) | **3.24E-27** | **4.47E-27** | 0.33 | (0.24,0.41) | **2.94E-12** | **8.57E-12** |
| 129 | Cer(d16:1/22:0) | 0.08 | (0.03,0.14) | **1.97E-03** | **2.21E-03** | -2.34 | (-2.4,-2.28) | **1.79E-250** | **6.15E-250** | 2.23 | (2.14,2.33) | **1.21E-122** | **7.38E-122** | 0.21 | (0.12,0.29) | **2.66E-06** | **4.47E-06** |
| 130 | Cer(d16:1/23:0) | 0.09 | (0.03,0.15) | **4.57E-03** | **5.12E-03** | -3.18 | (-3.27,-3.08) | **3.92E-228** | **1.02E-227** | 3.27 | (3.12,3.42) | **4.68E-114** | **2.44E-113** | 0.06 | (-0.03,0.16) | 2.00E-01 | 2.17E-01 |
| 131 | Cer(d16:1/24:0) | 0.31 | (0.25,0.36) | **3.72E-24** | **5.37E-24** | -2.68 | (-2.75,-2.62) | **4.04E-274** | **1.90E-273** | 2.84 | (2.75,2.93) | **1.12E-148** | **1.45E-147** | 0.15 | (0.06,0.24) | **8.03E-04** | **1.10E-03** |
| 132 | Cer(d16:1/24:1) | 0.19 | (0.14,0.24) | **3.72E-13** | **4.81E-13** | -2.07 | (-2.13,-2) | **3.17E-213** | **6.77E-213** | 2.02 | (1.93,2.12) | **2.46E-107** | **1.04E-106** | 0.27 | (0.19,0.35) | **7.05E-10** | **1.59E-09** |
| 133 | Cer(d17:1/22:0) | 0.24 | (0.18,0.3) | **8.04E-14** | **1.05E-13** | -1.88 | (-1.95,-1.8) | **4.06E-179** | **6.68E-179** | 1.94 | (1.83,2.05) | **4.13E-95** | **1.45E-94** | 0.13 | (0.05,0.22) | **2.45E-03** | **3.17E-03** |
| 134 | Cer(d17:1/23:0) | 0.23 | (0.17,0.29) | **2.75E-13** | **3.56E-13** | -3.21 | (-3.32,-3.11) | **2.68E-210** | **5.45E-210** | 3.40 | (3.23,3.56) | **7.83E-109** | **3.42E-108** | 0.06 | (-0.03,0.15) | 1.90E-01 | 2.08E-01 |
| 135 | Cer(d17:1/24:0) | 0.43 | (0.38,0.48) | **1.13E-49** | **2.07E-49** | -2.04 | (-2.1,-1.97) | **8.51E-225** | **2.09E-224** | 2.39 | (2.3,2.48) | **5.93E-130** | **4.99E-129** | 0.08 | (0.01,0.16) | **3.67E-02** | **4.30E-02** |
| 136 | Cer(d17:1/24:1) | 0.25 | (0.2,0.3) | **9.42E-21** | **1.32E-20** | -2.16 | (-2.23,-2.09) | **2.63E-212** | **5.50E-212** | 2.20 | (2.1,2.31) | **5.11E-113** | **2.58E-112** | 0.26 | (0.18,0.34) | **1.49E-09** | **3.23E-09** |
| 137 | Cer(d18:1/16:0) | -0.26 | (-0.3,-0.22) | **3.91E-32** | **6.13E-32** | -2.08 | (-2.12,-2.03) | **3.60E-283** | **1.99E-282** | 1.62 | (1.55,1.68) | **3.20E-124** | **2.08E-123** | 0.18 | (0.12,0.24) | **2.52E-09** | **5.42E-09** |
| 138 | Cer(d18:1/18:0) | -0.12 | (-0.17,-0.07) | **1.11E-05** | **1.32E-05** | -1.61 | (-1.67,-1.54) | **2.31E-182** | **3.87E-182** | 1.23 | (1.15,1.32) | **8.50E-77** | **2.16E-76** | 0.23 | (0.15,0.31) | **3.14E-08** | **6.18E-08** |
| 139 | Cer(d18:1/20:0) | -0.18 | (-0.23,-0.13) | **1.55E-13** | **2.02E-13** | -2.02 | (-2.07,-1.96) | **2.42E-253** | **8.60E-253** | 1.54 | (1.47,1.62) | **8.02E-108** | **3.41E-107** | 0.31 | (0.24,0.38) | **3.47E-16** | **1.38E-15** |
| 140 | Cer(d18:1/22:0) | 0.15 | (0.11,0.19) | **3.55E-12** | **4.50E-12** | -2.27 | (-2.31,-2.22) | **9.11E-296** | **7.29E-295** | 2.13 | (2.07,2.2) | **2.49E-149** | **3.32E-148** | 0.28 | (0.22,0.34) | **1.65E-16** | **6.76E-16** |
| 141 | Cer(d18:1/23:0) | 0.08 | (0.04,0.12) | **2.74E-04** | **3.15E-04** | -3.78 | (-3.84,-3.72) | **0.00E+00** | **0.00E+00** | 3.72 | (3.64,3.8) | **2.94E-187** | **2.02E-185** | 0.17 | (0.11,0.24) | **5.66E-07** | **9.96E-07** |
| 142 | Cer(d18:1/24:0) | 0.23 | (0.19,0.27) | **1.57E-24** | **2.28E-24** | -2.02 | (-2.07,-1.97) | **2.26E-264** | **8.88E-264** | 2.04 | (1.97,2.1) | **1.42E-147** | **1.70E-146** | 0.17 | (0.11,0.23) | **7.51E-08** | **1.42E-07** |
| 143 | Cer(d18:1/24:1) | 0.03 | (-0.02,0.07) | 2.08E-01 | 2.19E-01 | -2.24 | (-2.3,-2.19) | **6.68E-263** | **2.59E-262** | 1.91 | (1.83,1.99) | **2.27E-124** | **1.50E-123** | 0.34 | (0.28,0.4) | **8.98E-23** | **5.60E-22** |
| 144 | Cer(d18:2/16:0) | -0.37 | (-0.43,-0.31) | **4.51E-31** | **7.02E-31** | -2.21 | (-2.28,-2.14) | **4.97E-220** | **1.15E-219** | 1.29 | (1.18,1.4) | **1.62E-61** | **3.16E-61** | 0.48 | (0.4,0.56) | **2.44E-25** | **2.02E-24** |
| 145 | Cer(d18:2/22:0) | 0.44 | (0.39,0.49) | **1.07E-53** | **2.03E-53** | -1.48 | (-1.53,-1.42) | **3.06E-193** | **5.51E-193** | 1.50 | (1.42,1.58) | **2.21E-99** | **8.43E-99** | 0.42 | (0.35,0.49) | **5.29E-25** | **4.09E-24** |
| 146 | Cer(d18:2/23:0) | 0.36 | (0.31,0.41) | **1.05E-39** | **1.74E-39** | -3.34 | (-3.41,-3.26) | **1.75E-281** | **9.57E-281** | 3.45 | (3.35,3.55) | **3.25E-156** | **5.20E-155** | 0.29 | (0.22,0.37) | **1.24E-12** | **3.69E-12** |
| 147 | Cer(d18:2/24:0) | 0.51 | (0.46,0.56) | **2.10E-70** | **5.04E-70** | -1.78 | (-1.84,-1.72) | **1.40E-212** | **2.94E-212** | 1.93 | (1.85,2.01) | **7.46E-123** | **4.65E-122** | 0.34 | (0.27,0.41) | **2.41E-19** | **1.15E-18** |
| 148 | Cer(d18:2/24:1) | 0.40 | (0.35,0.44) | **9.94E-46** | **1.75E-45** | -1.55 | (-1.61,-1.49) | **2.32E-180** | **3.85E-180** | 1.50 | (1.41,1.58) | **1.27E-94** | **4.37E-94** | 0.47 | (0.4,0.54) | **9.26E-33** | **2.02E-31** |
| 149 | Cer(d19:1/24:0) | 0.72 | (0.65,0.8) | **7.44E-58** | **1.46E-57** | -0.94 | (-1.02,-0.87) | **5.14E-79** | **6.12E-79** | 1.73 | (1.6,1.86) | **3.14E-73** | **7.34E-73** | -0.13 | (-0.25,-0.01) | **3.29E-02** | **3.89E-02** |
| 150 | Cer(d19:1/24:1) | 0.47 | (0.38,0.55) | **1.46E-24** | **2.13E-24** | -1.27 | (-1.37,-1.18) | **9.71E-93** | **1.20E-92** | 1.64 | (1.5,1.78) | **2.74E-61** | **5.28E-61** | 0.08 | (-0.05,0.21) | 2.45E-01 | 2.63E-01 |
| 151 | Cer(m18:0/22:0) | -0.90 | (-0.97,-0.82) | **1.08E-71** | **2.67E-71** | -2.35 | (-2.43,-2.27) | **2.38E-205** | **4.72E-205** | 1.28 | (1.17,1.39) | **6.27E-60** | **1.19E-59** | 0.26 | (0.16,0.35) | **1.58E-07** | **2.94E-07** |
| 152 | Cer(m18:0/23:0) | -0.96 | (-1.04,-0.87) | **8.86E-71** | **2.14E-70** | -4.13 | (-4.27,-4) | **4.57E-216** | **1.01E-215** | 3.26 | (3.05,3.47) | **1.53E-84** | **4.38E-84** | 0.12 | (0,0.24) | **4.55E-02** | 5.27E-02 |
| 153 | Cer(m18:0/24:0) | -0.72 | (-0.79,-0.65) | **1.32E-62** | **2.87E-62** | -2.31 | (-2.38,-2.25) | **1.09E-239** | **3.26E-239** | 1.37 | (1.28,1.47) | **1.34E-80** | **3.57E-80** | 0.19 | (0.09,0.28) | **9.04E-05** | **1.32E-04** |
| 154 | Cer(m18:0/24:1) | -0.93 | (-1.01,-0.85) | **3.35E-72** | **8.43E-72** | -2.98 | (-3.07,-2.89) | **1.70E-233** | **4.66E-233** | 1.62 | (1.5,1.75) | **7.03E-70** | **1.54E-69** | 0.41 | (0.31,0.51) | **2.35E-13** | **7.33E-13** |
| 155 | Cer(m18:1/22:0) | -0.54 | (-0.61,-0.47) | **3.71E-39** | **6.11E-39** | -1.43 | (-1.5,-1.36) | **8.20E-152** | **1.21E-151** | 0.33 | (0.22,0.43) | **3.35E-09** | **3.86E-09** | 0.59 | (0.48,0.7) | **1.35E-22** | **8.22E-22** |
| 156 | Cer(m18:1/23:0) | -0.54 | (-0.62,-0.46) | **3.84E-33** | **6.08E-33** | -3.36 | (-3.49,-3.22) | **8.10E-183** | **1.36E-182** | 2.34 | (2.15,2.53) | **3.19E-67** | **6.72E-67** | 0.53 | (0.41,0.66) | **2.02E-15** | **7.62E-15** |
| 157 | Cer(m18:1/24:0) | -0.33 | (-0.4,-0.26) | **3.55E-20** | **4.96E-20** | -1.33 | (-1.4,-1.26) | **3.36E-143** | **4.77E-143** | 0.41 | (0.31,0.51) | **1.59E-13** | **1.91E-13** | 0.51 | (0.41,0.61) | **2.85E-20** | **1.44E-19** |
| 158 | Cer(m18:1/24:1) | -0.56 | (-0.63,-0.49) | **3.20E-43** | **5.44E-43** | -2.08 | (-2.16,-2) | **4.14E-186** | **7.05E-186** | 0.80 | (0.68,0.93) | **7.96E-29** | **1.12E-28** | 0.68 | (0.57,0.78) | **1.41E-29** | **1.78E-28** |
| 159 | GM3(d18:1/18:0) | 0.53 | (0.48,0.57) | **1.14E-73** | **2.90E-73** | 0.66 | (0.61,0.72) | **9.71E-80** | **1.16E-79** | -0.13 | (-0.2,-0.06) | **2.42E-04** | **2.58E-04** | -0.08 | (-0.15,-0.02) | **1.62E-02** | **1.96E-02** |
| 160 | GM3(d18:1/22:0) | 0.05 | (0,0.11) | 6.58E-02 | 7.10E-02 | -0.98 | (-1.05,-0.9) | **5.36E-89** | **6.49E-89** | 0.88 | (0.79,0.97) | **3.53E-50** | **6.15E-50** | 0.12 | (0.05,0.19) | **9.07E-04** | **1.23E-03** |
| 161 | GM3(d18:1/24:0) | 0.48 | (0.42,0.54) | **6.03E-46** | **1.07E-45** | 0.13 | (0.06,0.19) | **2.30E-04** | **2.39E-04** | 0.34 | (0.26,0.42) | **2.12E-14** | **2.57E-14** | -0.03 | (-0.1,0.03) | 3.25E-01 | 3.47E-01 |

| | | | | | | | | | | | | | | | | | |
|---|---|---|---|---|---|---|---|---|---|---|---|---|---|---|---|---|---|
| 162 | GM3(d18:1/24:1) | 0.03 | (-0.02,0.09) | 2.57E-01 | 2.68E-01 | -0.52 | (-0.59,-0.45) | **1.26E-42** | **1.41E-42** | 0.37 | (0.29,0.46) | **1.85E-15** | **2.26E-15** | 0.12 | (0.05,0.2) | **1.23E-03** | **1.65E-03** |
| 163 | HexCer(d16:1/22:0) | 0.06 | (0.01,0.12) | **3.18E-02** | **3.48E-02** | -2.18 | (-2.27,-2.08) | **1.12E-164** | **1.73E-164** | 2.04 | (1.91,2.17) | **3.98E-86** | **1.18E-85** | 0.24 | (0.16,0.32) | **4.93E-09** | **1.04E-08** |
| 164 | HexCer(d16:1/24:0) | 0.34 | (0.29,0.4) | **4.06E-27** | **6.09E-27** | -1.75 | (-1.84,-1.66) | **2.61E-143** | **3.71E-143** | 1.74 | (1.62,1.86) | **1.30E-75** | **3.21E-75** | 0.27 | (0.19,0.34) | **9.05E-11** | **2.22E-10** |
| 165 | HexCer(d18:1/16:0) | 0.08 | (0.04,0.12) | **1.16E-04** | **1.35E-04** | -1.82 | (-1.88,-1.76) | **3.76E-212** | **7.82E-212** | 1.86 | (1.79,1.93) | **1.00E-127** | **6.88E-127** | -0.04 | (-0.11,0.02) | 1.56E-01 | 1.72E-01 |
| 166 | HexCer(d18:1/18:0) | -0.11 | (-0.17,-0.05) | **1.41E-04** | **1.63E-04** | -1.33 | (-1.4,-1.26) | **1.76E-133** | **2.43E-133** | 0.96 | (0.86,1.06) | **4.24E-48** | **7.26E-48** | 0.22 | (0.14,0.3) | **8.84E-08** | **1.67E-07** |
| 167 | HexCer(d18:1/20:0) | -0.09 | (-0.14,-0.05) | **1.33E-04** | **1.54E-04** | -1.40 | (-1.47,-1.33) | **1.47E-150** | **2.17E-150** | 1.14 | (1.05,1.22) | **2.52E-71** | **5.65E-71** | 0.17 | (0.1,0.25) | **5.83E-06** | **9.45E-06** |
| 168 | HexCer(d18:1/22:0) | 0.10 | (0.06,0.14) | **2.72E-07** | **3.33E-07** | -1.89 | (-1.94,-1.83) | **5.07E-232** | **1.36E-231** | 1.77 | (1.7,1.84) | **6.98E-127** | **4.72E-126** | 0.20 | (0.14,0.26) | **1.25E-09** | **2.76E-09** |
| 169 | HexCer(d18:1/24:0) | 0.24 | (0.2,0.28) | **1.18E-26** | **1.76E-26** | -0.85 | (-0.9,-0.79) | **5.63E-108** | **7.21E-108** | 0.89 | (0.82,0.96) | **7.42E-71** | **1.66E-70** | 0.16 | (0.1,0.22) | **1.31E-06** | **2.24E-06** |
| 170 | HexCer(d18:1/24:1) | 0.02 | (-0.02,0.07) | 2.87E-01 | 2.99E-01 | -1.30 | (-1.35,-1.24) | **1.65E-169** | **2.62E-169** | 1.22 | (1.15,1.29) | **4.63E-93** | **1.56E-92** | 0.05 | (-0.02,0.12) | 1.40E-01 | 1.55E-01 |
| 171 | Hex2Cer(d16:1/16:0) | 0.03 | (-0.02,0.08) | 2.89E-01 | 3.00E-01 | -3.15 | (-3.22,-3.07) | **2.81E-265** | **1.14E-264** | 2.59 | (2.48,2.7) | **1.29E-122** | **7.72E-122** | 0.59 | (0.51,0.67) | **1.68E-35** | **6.21E-34** |
| 172 | Hex2Cer(d18:1/16:0) | -0.13 | (-0.18,-0.09) | **9.14E-09** | **1.14E-08** | -2.34 | (-2.39,-2.28) | **1.11E-271** | **4.95E-271** | 1.82 | (1.75,1.89) | **2.45E-132** | **2.18E-131** | 0.37 | (0.3,0.43) | **6.83E-24** | **4.82E-23** |
| 173 | Hex2Cer(d18:1/22:0) | 0.23 | (0.18,0.28) | **1.10E-18** | **1.52E-18** | -1.09 | (-1.16,-1.02) | **1.97E-118** | **2.63E-118** | 1.01 | (0.93,1.09) | **1.16E-69** | **2.52E-69** | 0.27 | (0.2,0.34) | **8.93E-13** | **2.70E-12** |
| 174 | Hex2Cer(d18:1/24:1) | 0.37 | (0.33,0.42) | **2.02E-45** | **3.55E-45** | -1.62 | (-1.68,-1.56) | **2.27E-199** | **4.26E-199** | 1.75 | (1.68,1.82) | **2.84E-128** | **2.09E-127** | 0.20 | (0.13,0.27) | **1.27E-08** | **2.60E-08** |
| 175 | Hex2Cer(d18:2/16:0) | 0.02 | (-0.03,0.06) | 4.68E-01 | 4.83E-01 | -1.99 | (-2.05,-1.93) | **2.02E-225** | **5.08E-225** | 1.39 | (1.32,1.47) | **5.54E-100** | **2.13E-99** | 0.57 | (0.51,0.64) | **9.06E-45** | **8.70E-43** |
| 176 | Hex3Cer(d18:1/16:0) | 0.00 | (-0.04,0.03) | 8.86E-01 | 8.89E-01 | -1.09 | (-1.13,-1.04) | **3.57E-175** | **5.77E-175** | 1.02 | (0.96,1.08) | **1.11E-93** | **3.79E-93** | 0.00 | (-0.06,0.05) | 9.23E-01 | 9.27E-01 |
| 177 | Hex3Cer(d18:1/22:0) | 0.43 | (0.38,0.49) | **4.17E-41** | **7.00E-41** | -1.19 | (-1.26,-1.12) | **5.31E-115** | **6.98E-115** | 1.73 | (1.64,1.83) | **4.61E-99** | **1.74E-98** | -0.15 | (-0.22,-0.07) | **2.26E-04** | **3.24E-04** |
| 178 | Hex3Cer(d18:1/24:0) | 0.53 | (0.48,0.58) | **7.29E-66** | **1.64E-65** | -0.10 | (-0.16,-0.04) | **1.44E-03** | **1.49E-03** | 0.78 | (0.7,0.86) | **2.14E-49** | **3.69E-49** | -0.18 | (-0.25,-0.11) | **2.84E-07** | **5.14E-07** |
| 179 | Hex3Cer(d18:1/24:1) | 0.35 | (0.3,0.4) | **1.95E-37** | **3.19E-37** | -0.95 | (-1.01,-0.9) | **3.88E-115** | **5.12E-115** | 1.27 | (1.19,1.35) | **2.83E-85** | **8.18E-85** | -0.03 | (-0.1,0.03) | 3.25E-01 | 3.47E-01 |
| 180 | SM(d17:1/14:0) | -0.18 | (-0.22,-0.13) | **2.46E-14** | **3.24E-14** | -1.63 | (-1.68,-1.59) | **1.14E-240** | **3.45E-240** | 1.15 | (1.07,1.23) | **2.29E-75** | **5.61E-75** | 0.28 | (0.2,0.36) | **2.80E-11** | **7.24E-11** |
| 181 | SM(d18:0/14:0) | -0.21 | (-0.26,-0.16) | **6.88E-17** | **9.30E-17** | 0.05 | (0,0.1) | **3.13E-02** | **3.21E-02** | -0.42 | (-0.5,-0.35) | **1.45E-25** | **1.97E-25** | 0.13 | (0.06,0.2) | **5.48E-04** | **7.56E-04** |
| 182 | SM(d18:1/14:0)/SM(d16:1/16:0) | -0.19 | (-0.23,-0.16) | **2.17E-22** | **3.09E-22** | -1.92 | (-1.96,-1.88) | **3.24E-291** | **2.13E-290** | 1.43 | (1.36,1.5) | **1.46E-112** | **7.14E-112** | 0.26 | (0.2,0.32) | **4.34E-15** | **1.57E-14** |
| 183 | SM(d18:2/14:0) | -0.18 | (-0.22,-0.14) | **3.36E-16** | **4.50E-16** | -1.12 | (-1.17,-1.08) | **7.67E-181** | **1.28E-180** | 0.33 | (0.26,0.41) | **8.37E-18** | **1.06E-17** | 0.57 | (0.5,0.64) | **4.36E-41** | **3.49E-39** |
| 184 | SM(d17:1/16:0) | -0.15 | (-0.19,-0.12) | **4.55E-17** | **6.17E-17** | -1.91 | (-1.95,-1.87) | **4.05E-292** | **2.90E-291** | 1.50 | (1.43,1.57) | **1.18E-115** | **6.36E-115** | 0.21 | (0.14,0.27) | **3.81E-10** | **8.84E-10** |
| 185 | SM(d18:1/16:0) | -0.33 | (-0.37,-0.28) | **1.02E-35** | **1.65E-35** | -2.37 | (-2.42,-2.32) | **2.40E-291** | **1.62E-290** | 1.71 | (1.63,1.78) | **3.05E-123** | **1.95E-122** | 0.30 | (0.23,0.37) | **1.22E-16** | **5.08E-16** |
| 186 | SM(d18:2/16:0) | -0.17 | (-0.2,-0.14) | **7.00E-25** | **1.02E-24** | -1.25 | (-1.29,-1.21) | **1.34E-230** | **3.53E-230** | 0.55 | (0.49,0.6) | **1.13E-54** | **2.06E-54** | 0.50 | (0.45,0.55) | **4.29E-57** | **1.03E-54** |
| 187 | SM(34:3) | -0.15 | (-0.19,-0.11) | **2.66E-12** | **3.39E-12** | -1.19 | (-1.23,-1.14) | **1.59E-193** | **2.87E-193** | 0.62 | (0.55,0.69) | **2.71E-43** | **4.33E-43** | 0.37 | (0.3,0.44) | **8.29E-22** | **4.79E-21** |
| 188 | SM(d18:2/17:0) | -0.10 | (-0.14,-0.07) | **1.79E-08** | **2.21E-08** | -1.18 | (-1.23,-1.14) | **1.01E-204** | **1.97E-204** | 0.68 | (0.61,0.74) | **2.37E-54** | **4.30E-54** | 0.37 | (0.31,0.43) | **8.01E-28** | **8.18E-27** |
| 189 | SM(35:2) (b) | -0.34 | (-0.39,-0.29) | **3.42E-34** | **5.49E-34** | -2.57 | (-2.62,-2.51) | **1.88E-278** | **9.80E-278** | 1.80 | (1.71,1.89) | **1.86E-106** | **7.69E-106** | 0.37 | (0.29,0.45) | **1.34E-17** | **6.03E-17** |
| 190 | SM(d18:1/18:0)/SM(d16:1/20:0) | -0.32 | (-0.36,-0.28) | **1.64E-40** | **2.74E-40** | -1.07 | (-1.12,-1.03) | **4.94E-163** | **7.58E-163** | 0.52 | (0.45,0.58) | **9.34E-36** | **1.40E-35** | 0.20 | (0.14,0.26) | **2.42E-10** | **5.74E-10** |
| 191 | SM(d18:2/18:1) | -0.22 | (-0.26,-0.18) | **6.66E-27** | **9.96E-27** | -1.61 | (-1.66,-1.57) | **2.07E-241** | **6.31E-241** | 1.13 | (1.06,1.19) | **1.04E-92** | **3.45E-92** | 0.23 | (0.17,0.28) | **3.81E-14** | **1.26E-13** |
| 192 | SM(37:1) | -0.34 | (-0.39,-0.3) | **8.88E-43** | **1.51E-42** | -2.24 | (-2.29,-2.19) | **1.07E-276** | **5.35E-276** | 1.62 | (1.55,1.7) | **1.10E-112** | **5.44E-112** | 0.25 | (0.19,0.32) | **1.58E-12** | **4.67E-12** |
| 193 | SM(37:2) | -0.41 | (-0.46,-0.36) | **1.30E-42** | **2.19E-42** | -2.00 | (-2.06,-1.94) | **6.15E-240** | **1.86E-239** | 1.03 | (0.95,1.11) | **1.39E-69** | **3.02E-69** | 0.50 | (0.43,0.58) | **1.99E-31** | **3.68E-30** |
| 194 | SM(d18:1/20:0)/SM(d16:1/22:0) | -0.34 | (-0.37,-0.3) | **5.44E-59** | **1.10E-58** | -1.43 | (-1.47,-1.39) | **4.26E-242** | **1.32E-241** | 0.90 | (0.84,0.96) | **4.37E-81** | **1.18E-80** | 0.17 | (0.12,0.22) | **1.84E-10** | **4.43E-10** |
| 195 | SM(d18:2/20:0) | -0.43 | (-0.48,-0.39) | **1.16E-61** | **2.45E-61** | -1.01 | (-1.06,-0.95) | **1.02E-141** | **1.44E-141** | 0.15 | (0.08,0.23) | **1.20E-04** | **1.29E-04** | 0.38 | (0.32,0.45) | **5.14E-26** | **4.57E-25** |
| 196 | SM(38:3) (a) | -0.43 | (-0.47,-0.38) | **6.49E-64** | **1.42E-63** | -1.20 | (-1.25,-1.16) | **3.59E-188** | **6.22E-188** | 0.32 | (0.25,0.38) | **5.92E-18** | **7.53E-18** | 0.38 | (0.32,0.44) | **8.99E-31** | **1.35E-29** |
| 197 | SM(38:3) (b) | -0.36 | (-0.42,-0.31) | **6.93E-31** | **1.08E-30** | -2.14 | (-2.2,-2.09) | **6.12E-253** | **2.16E-252** | 1.23 | (1.14,1.31) | **3.99E-74** | **9.54E-74** | 0.47 | (0.39,0.55) | **2.47E-26** | **2.33E-25** |
| 198 | SM(d16:1/23:0)/SM(d17:1/22:0) | -0.30 | (-0.33,-0.26) | **1.19E-43** | **2.05E-43** | -2.63 | (-2.68,-2.59) | **0.00E+00** | **0.00E+00** | 2.23 | (2.16,2.3) | **7.33E-152** | **1.07E-150** | 0.07 | (0.01,0.13) | **2.52E-02** | **2.98E-02** |
| 199 | SM(d18:1/22:0)/SM(d16:1/24:0) | -0.35 | (-0.39,-0.31) | **2.03E-50** | **3.74E-50** | -2.02 | (-2.07,-1.98) | **1.81E-288** | **1.09E-287** | 1.45 | (1.39,1.51) | **1.12E-119** | **6.45E-119** | 0.17 | (0.12,0.23) | **8.46E-10** | **1.90E-09** |
| 200 | SM(d16:1/24:1) | -0.33 | (-0.37,-0.29) | **5.44E-47** | **9.70E-47** | -2.12 | (-2.16,-2.07) | **2.38E-286** | **1.39E-285** | 1.49 | (1.42,1.56) | **6.58E-108** | **2.82E-107** | 0.26 | (0.2,0.32) | **8.33E-15** | **2.92E-14** |
| 201 | SM(d18:2/22:0) | -0.13 | (-0.17,-0.1) | **1.23E-12** | **1.57E-12** | -1.25 | (-1.29,-1.2) | **1.05E-204** | **2.05E-204** | 0.75 | (0.69,0.81) | **3.54E-64** | **7.04E-64** | 0.33 | (0.27,0.38) | **3.13E-27** | **3.07E-26** |
| 202 | SM(40:3) (a) | -0.69 | (-0.74,-0.65) | **2.74E-107** | **1.43E-106** | -1.57 | (-1.62,-1.52) | **7.42E-229** | **1.94E-228** | 0.37 | (0.29,0.44) | **2.21E-20** | **2.87E-20** | 0.44 | (0.38,0.5) | **1.28E-37** | **5.58E-36** |
| 203 | SM(40:3) (b) | -0.46 | (-0.5,-0.41) | **3.55E-58** | **7.13E-58** | -1.98 | (-2.02,-1.93) | **3.55E-270** | **1.56E-269** | 1.21 | (1.13,1.3) | **3.66E-79** | **9.38E-79** | 0.24 | (0.16,0.31) | **1.11E-09** | **2.46E-09** |
| 204 | SM(41:0) | -0.45 | (-0.49,-0.42) | **2.96E-72** | **7.48E-72** | -1.96 | (-2.02,-1.9) | **8.09E-218** | **1.83E-217** | 1.45 | (1.36,1.55) | **9.75E-84** | **2.73E-83** | -0.02 | (-0.08,0.04) | 5.33E-01 | 5.61E-01 |
| 205 | SM(41:1) | -0.43 | (-0.46,-0.39) | **6.27E-73** | **1.59E-72** | -3.33 | (-3.38,-3.28) | **0.00E+00** | **0.00E+00** | 2.83 | (2.77,2.9) | **4.94E-176** | **1.83E-174** | 0.03 | (-0.03,0.08) | 3.25E-01 | 3.47E-01 |
| 206 | SM(d17:1/24:1) | -0.36 | (-0.4,-0.33) | **5.29E-60** | **1.11E-59** | -1.81 | (-1.85,-1.76) | **1.47E-268** | **6.25E-268** | 1.23 | (1.16,1.29) | **2.30E-97** | **8.57E-97** | 0.16 | (0.11,0.22) | **6.72E-08** | **1.28E-07** |
| 207 | SM(d18:2/23:0) | -0.21 | (-0.24,-0.17) | **2.64E-28** | **4.03E-28** | -3.00 | (-3.05,-2.95) | **0.00E+00** | **0.00E+00** | 2.59 | (2.52,2.66) | **6.11E-164** | **1.33E-162** | 0.16 | (0.11,0.22) | **7.05E-09** | **1.46E-08** |
| 208 | SM(d18:1/24:0) | -0.20 | (-0.24,-0.17) | **2.72E-22** | **3.86E-22** | -1.25 | (-1.3,-1.2) | **7.07E-199** | **1.32E-198** | 0.95 | (0.89,1.01) | **3.82E-83** | **1.06E-82** | 0.05 | (0,0.1) | **3.68E-02** | **4.30E-02** |
| 209 | SM(d18:1/24:1) | -0.55 | (-0.6,-0.5) | **1.53E-75** | **4.02E-75** | -1.94 | (-1.99,-1.89) | **2.31E-255** | **8.34E-255** | 1.09 | (1.02,1.16) | **2.98E-84** | **8.45E-84** | 0.25 | (0.19,0.31) | **2.20E-15** | **8.24E-15** |
| 210 | SM(d18:2/24:0) | -0.06 | (-0.1,-0.03) | **2.15E-04** | **2.47E-04** | -1.07 | (-1.11,-1.03) | **1.03E-186** | **1.77E-186** | 0.76 | (0.7,0.81) | **1.08E-71** | **2.45E-71** | 0.20 | (0.15,0.24) | **9.46E-15** | **3.29E-14** |
| 211 | SM(43:1) | 0.04 | (0,0.08) | **3.40E-02** | **3.72E-02** | -0.92 | (-0.98,-0.86) | **4.34E-113** | **5.64E-113** | 1.00 | (0.92,1.09) | **2.69E-60** | **5.14E-60** | -0.12 | (-0.19,-0.06) | **1.36E-04** | **1.97E-04** |
| 212 | SM(44:1) | 0.20 | (0.17,0.24) | **1.50E-26** | **2.22E-26** | 0.54 | (0.5,0.58) | **1.07E-91** | **1.31E-91** | -0.33 | (-0.39,-0.28) | **3.48E-26** | **4.75E-26** | -0.06 | (-0.11,-0.01) | **2.37E-02** | **2.82E-02** |
| 213 | SM(44:2) | 0.01 | (-0.03,0.05) | 7.21E-01 | 7.30E-01 | 0.16 | (0.12,0.21) | **1.63E-12** | **1.73E-12** | -0.30 | (-0.36,-0.24) | **7.48E-19** | **9.63E-19** | 0.09 | (0.04,0.15) | **1.39E-03** | **1.85E-03** |
| 214 | SM(44:3) (a) | -0.02 | (-0.06,0.02) | 2.39E-01 | 2.51E-01 | 0.20 | (0.16,0.24) | **1.12E-18** | **1.20E-18** | -0.52 | (-0.58,-0.46) | **2.07E-45** | **3.41E-45** | 0.23 | (0.18,0.29) | **7.79E-16** | **3.07E-15** |
| 215 | LPC(14:0) [sn2] | 0.39 | (0.31,0.47) | **1.21E-19** | **1.68E-19** | 0.39 | (0.32,0.45) | **4.09E-28** | **4.48E-28** | -0.32 | (-0.42,-0.22) | **4.65E-10** | **5.43E-10** | 0.27 | (0.15,0.38) | **5.32E-06** | **8.72E-06** |
| 216 | LPC(14:0) [sn1] | 0.40 | (0.32,0.48) | **8.04E-22** | **1.14E-21** | 0.28 | (0.22,0.35) | **8.91E-18** | **9.59E-18** | -0.17 | (-0.26,-0.08) | **2.91E-04** | **3.10E-04** | 0.26 | (0.15,0.37) | **5.35E-06** | **8.73E-06** |

| | | | | | | | | | | | | | | | | | |
|---|---|---|---|---|---|---|---|---|---|---|---|---|---|---|---|---|---|
| 217 | LPC(15:0) [sn2] | 0.60 | (0.54,0.66) | **9.20E-60** | **1.90E-59** | 0.11 | (0.06,0.17) | **2.16E-05** | **2.25E-05** | 0.23 | (0.15,0.32) | **3.17E-07** | **3.55E-07** | 0.25 | (0.16,0.34) | **2.10E-07** | **3.87E-07** |
| 218 | LPC(15:0) [sn1] | 0.61 | (0.55,0.66) | **2.15E-70** | **5.15E-70** | -0.03 | (-0.08,0.02) | 1.84E-01 | 1.87E-01 | 0.36 | (0.28,0.44) | **2.52E-15** | **3.08E-15** | 0.28 | (0.2,0.37) | **3.62E-10** | **8.44E-10** |
| 219 | LPC(16:0) [sn2] | 0.60 | (0.56,0.65) | **4.03E-90** | **1.39E-89** | 0.00 | (-0.04,0.04) | 9.64E-01 | 9.64E-01 | 0.22 | (0.15,0.29) | **3.52E-09** | **4.04E-09** | 0.38 | (0.31,0.44) | **1.18E-25** | **1.01E-24** |
| 220 | LPC(16:0) [sn1] | 0.71 | (0.66,0.76) | **4.53E-90** | **1.54E-89** | -0.09 | (-0.14,-0.04) | **1.36E-04** | **1.42E-04** | 0.36 | (0.28,0.44) | **3.16E-16** | **3.93E-16** | 0.45 | (0.37,0.52) | **1.47E-26** | **1.41E-25** |
| 221 | LPC(16:1) [sn2] | 0.84 | (0.77,0.9) | **1.79E-82** | **5.37E-82** | 1.84 | (1.77,1.9) | **1.25E-193** | **2.26E-193** | -1.42 | (-1.51,-1.33) | **3.32E-86** | **9.89E-86** | 0.41 | (0.33,0.48) | **3.61E-22** | **2.14E-21** |
| 222 | LPC(16:1) [sn1] | 0.88 | (0.82,0.94) | **3.62E-99** | **1.51E-98** | 1.90 | (1.83,1.96) | **7.98E-214** | **1.72E-213** | -1.38 | (-1.47,-1.3) | **2.38E-87** | **7.19E-87** | 0.36 | (0.29,0.44) | **4.62E-19** | **2.17E-18** |
| 223 | LPC(15-MHDA) [sn2] | 0.95 | (0.88,1.02) | **1.13E-87** | **3.64E-87** | 0.57 | (0.5,0.64) | **5.93E-44** | **6.71E-44** | 0.19 | (0.07,0.31) | **1.99E-03** | **2.09E-03** | 0.14 | (0.03,0.26) | **1.06E-02** | **1.31E-02** |
| 224 | LPC(15-MHDA) [sn1] / LPC(17:0) [sn2] | 0.93 | (0.87,0.99) | **3.12E-100** | **1.34E-99** | 0.08 | (0.02,0.13) | **6.98E-03** | **7.19E-03** | 0.63 | (0.53,0.73) | **1.92E-28** | **2.69E-28** | 0.20 | (0.11,0.29) | **3.04E-05** | **4.60E-05** |
| 225 | LPC(17:0) [sn1] | 0.86 | (0.81,0.92) | **8.57E-114** | **5.27E-113** | -0.35 | (-0.39,-0.31) | **4.27E-48** | **4.86E-48** | 0.94 | (0.86,1.02) | **2.90E-65** | **5.92E-65** | 0.27 | (0.2,0.35) | **1.60E-11** | **4.19E-11** |
| 226 | LPC(17:1) [sn1] (a) / LPC(17:1) [sn2] (b) | 0.85 | (0.78,0.91) | **8.22E-92** | **3.01E-91** | 1.14 | (1.09,1.2) | **6.84E-152** | **1.01E-151** | -0.56 | (-0.64,-0.47) | **4.29E-29** | **6.10E-29** | 0.28 | (0.19,0.37) | **1.46E-09** | **3.19E-09** |
| 227 | LPC(18:0) [sn2] | 1.18 | (1.13,1.23) | **8.38E-158** | **1.39E-156** | -0.01 | (-0.05,0.02) | 4.59E-01 | 4.64E-01 | 0.98 | (0.91,1.05) | **6.27E-75** | **1.52E-74** | 0.23 | (0.16,0.3) | **3.43E-10** | **8.06E-10** |
| 228 | LPC(18:0) [sn1] | 1.20 | (1.14,1.25) | **6.08E-156** | **8.84E-155** | -0.08 | (-0.12,-0.04) | **2.49E-04** | **2.58E-04** | 1.06 | (0.98,1.13) | **5.87E-76** | **1.47E-75** | 0.23 | (0.15,0.3) | **8.45E-09** | **1.75E-08** |
| 229 | LPC(18:1) [sn2] | 1.05 | (0.99,1.11) | **1.46E-118** | **1.02E-117** | 0.65 | (0.6,0.71) | **3.56E-78** | **4.23E-78** | 0.17 | (0.09,0.26) | **8.92E-05** | **9.58E-05** | 0.22 | (0.14,0.29) | **5.64E-08** | **1.08E-07** |
| 230 | LPC(18:1) [sn1] | 1.11 | (1.06,1.17) | **3.21E-141** | **3.35E-140** | 0.80 | (0.75,0.85) | **7.16E-115** | **9.39E-115** | 0.21 | (0.13,0.28) | **2.25E-07** | **2.53E-07** | 0.11 | (0.04,0.18) | **1.78E-03** | **2.33E-03** |
| 231 | LPC(18:2) [sn2] | 1.24 | (1.18,1.31) | **7.23E-138** | **7.09E-137** | 0.23 | (0.18,0.28) | **3.43E-17** | **3.68E-17** | 0.81 | (0.73,0.89) | **3.50E-52** | **6.27E-52** | 0.14 | (0.07,0.22) | **2.50E-04** | **3.56E-04** |
| 232 | LPC(18:2) [sn1] | 1.68 | (1.61,1.75) | **1.29E-162** | **2.48E-161** | 0.73 | (0.68,0.79) | **4.15E-92** | **5.09E-92** | 0.74 | (0.66,0.82) | **6.78E-44** | **1.09E-43** | 0.14 | (0.06,0.23) | **7.75E-04** | **1.06E-03** |
| 233 | LPC(18:3) [sn2] (a) | 0.71 | (0.63,0.78) | **2.95E-52** | **5.58E-52** | -0.59 | (-0.65,-0.53) | **5.39E-57** | **6.21E-57** | 1.26 | (1.16,1.36) | **4.68E-67** | **9.80E-67** | 0.03 | (-0.07,0.13) | 5.60E-01 | 5.86E-01 |
| 234 | LPC(18:3) [sn1] (a)/LPC(18:3) [sn2] (b) | 1.25 | (1.17,1.33) | **1.33E-99** | **5.66E-99** | 0.99 | (0.91,1.06) | **1.09E-90** | **1.32E-90** | 0.28 | (0.17,0.39) | **1.34E-06** | **1.48E-06** | -0.01 | (-0.11,0.1) | 8.72E-01 | 8.81E-01 |
| 235 | LPC(18:3) (a) [sn1] [104_sn1] | 1.31 | (1.22,1.4) | **2.73E-100** | **1.18E-99** | 1.02 | (0.94,1.1) | **4.27E-92** | **5.23E-92** | 0.30 | (0.19,0.41) | **3.87E-07** | **4.32E-07** | -0.01 | (-0.11,0.1) | 8.71E-01 | 8.81E-01 |
| 236 | LPC(19:0) [sn1] (a) / LPC(19:0) [sn2] (b) | 0.81 | (0.76,0.86) | **3.73E-112** | **2.24E-111** | -0.34 | (-0.39,-0.29) | **8.55E-34** | **9.48E-34** | 1.04 | (0.96,1.12) | **5.50E-69** | **1.18E-68** | 0.12 | (0.04,0.19) | **2.22E-03** | **2.88E-03** |
| 237 | LPC(20:0) [sn1] | 0.95 | (0.9,1) | **2.89E-133** | **2.47E-132** | -0.35 | (-0.39,-0.31) | **9.29E-50** | **1.06E-49** | 1.19 | (1.12,1.25) | **1.48E-96** | **5.37E-96** | 0.11 | (0.05,0.18) | **1.05E-03** | **1.42E-03** |
| 238 | LPC(20:1) [sn2] | 0.66 | (0.61,0.72) | **2.65E-76** | **7.03E-76** | -0.29 | (-0.34,-0.25) | **1.28E-34** | **1.42E-34** | 0.78 | (0.71,0.85) | **3.55E-56** | **6.56E-56** | 0.18 | (0.1,0.26) | **5.85E-06** | **9.45E-06** |
| 239 | LPC(20:1) [sn1] | 0.79 | (0.73,0.84) | **8.74E-99** | **3.58E-98** | -0.24 | (-0.28,-0.2) | **1.63E-25** | **1.77E-25** | 0.85 | (0.78,0.92) | **1.51E-62** | **2.98E-62** | 0.18 | (0.11,0.26) | **4.94E-06** | **8.12E-06** |
| 240 | LPC(20:2) [sn2] | 0.23 | (0.18,0.28) | **6.89E-18** | **9.42E-18** | 0.37 | (0.31,0.42) | **6.39E-34** | **7.10E-34** | -0.36 | (-0.43,-0.28) | **8.03E-17** | **1.00E-16** | 0.20 | (0.13,0.26) | **3.59E-08** | **7.04E-08** |
| 241 | LPC(20:2) [sn1] | 0.60 | (0.55,0.65) | **7.06E-78** | **1.98E-77** | 0.85 | (0.8,0.9) | **7.89E-122** | **1.06E-121** | -0.46 | (-0.54,-0.38) | **1.47E-24** | **1.99E-24** | 0.18 | (0.11,0.25) | **2.91E-07** | **5.25E-07** |
| 242 | LPC(20:3) [sn2] | 0.37 | (0.31,0.43) | **7.73E-30** | **1.19E-29** | 1.52 | (1.46,1.58) | **1.56E-192** | **2.76E-192** | -1.34 | (-1.43,-1.26) | **2.70E-85** | **7.85E-85** | 0.13 | (0.06,0.21) | **5.42E-04** | **7.50E-04** |
| 243 | LPC(20:3) [sn1] | 0.89 | (0.83,0.96) | **3.40E-91** | **1.23E-90** | 2.28 | (2.22,2.34) | **5.13E-244** | **1.65E-243** | -1.54 | (-1.64,-1.45) | **1.05E-90** | **3.39E-90** | 0.12 | (0.04,0.2) | **4.60E-03** | **5.82E-03** |
| 244 | LPC(20:3) [104_sn1] | 0.94 | (0.88,1.01) | **8.24E-95** | **3.16E-94** | 2.32 | (2.26,2.39) | **1.39E-243** | **4.42E-243** | -1.54 | (-1.64,-1.45) | **1.17E-88** | **3.68E-88** | 0.11 | (0.03,0.19) | **7.49E-03** | **9.34E-03** |
| 245 | LPC(20:4) [sn2] | 0.87 | (0.82,0.92) | **2.69E-117** | **1.79E-116** | 2.34 | (2.29,2.39) | **2.50E-289** | **1.56E-288** | -1.83 | (-1.9,-1.75) | **1.66E-124** | **1.11E-123** | 0.28 | (0.22,0.35) | **7.93E-16** | **3.09E-15** |
| 246 | LPC(20:4) [sn1] | 1.32 | (1.26,1.38) | **5.07E-152** | **6.76E-151** | 3.06 | (3.01,3.11) | **0.00E+00** | **0.00E+00** | -2.09 | (-2.17,-2.01) | **7.12E-130** | **5.79E-129** | 0.28 | (0.21,0.35) | **2.27E-13** | **7.13E-13** |
| 247 | LPC(20:5) [sn2] | 1.16 | (1.06,1.25) | **6.68E-74** | **1.71E-73** | 0.59 | (0.52,0.66) | **3.31E-44** | **3.75E-44** | -0.34 | (-0.48,-0.2) | **3.05E-06** | **3.35E-06** | 0.83 | (0.69,0.97) | **1.46E-25** | **1.23E-24** |
| 248 | LPC(20:5) [sn1] | 1.51 | (1.41,1.62) | **1.93E-98** | **7.79E-98** | 1.13 | (1.06,1.21) | **2.63E-106** | **3.36E-106** | -0.52 | (-0.66,-0.38) | **1.00E-11** | **1.20E-11** | 0.84 | (0.69,0.98) | **2.69E-25** | **2.19E-24** |
| 249 | LPC(22:4) [sn2] | 0.22 | (0.16,0.28) | **2.55E-13** | **3.31E-13** | 0.88 | (0.82,0.93) | **1.66E-114** | **2.17E-114** | -0.50 | (-0.58,-0.42) | **8.64E-28** | **1.20E-27** | -0.20 | (-0.27,-0.13) | **1.45E-07** | **2.70E-07** |
| 250 | LPC(22:4) [sn1] | 0.61 | (0.56,0.67) | **5.36E-67** | **1.23E-66** | 1.69 | (1.64,1.75) | **1.93E-217** | **4.32E-217** | -0.92 | (-1,-0.84) | **5.42E-61** | **1.04E-60** | -0.17 | (-0.25,-0.1) | **1.25E-05** | **1.95E-05** |
| 251 | LPC(22:5) [sn2] (n3) | 0.88 | (0.82,0.94) | **1.65E-95** | **6.48E-95** | 0.42 | (0.36,0.47) | **4.27E-38** | **4.80E-38** | 0.22 | (0.13,0.31) | **3.63E-06** | **3.97E-06** | 0.20 | (0.13,0.28) | **1.12E-07** | **2.11E-07** |
| 252 | LPC(22:5) [sn1] (n3)/LPC(22:5) [sn2] (n6) | 1.04 | (0.98,1.1) | **3.77E-120** | **2.66E-119** | 0.72 | (0.67,0.78) | **8.65E-86** | **1.04E-85** | 0.21 | (0.13,0.29) | **4.17E-07** | **4.64E-07** | 0.07 | (-0.01,0.14) | 8.30E-02 | 9.42E-02 |
| 253 | LPC(22:5) [sn1] (n6) | 0.68 | (0.61,0.75) | **5.96E-58** | **1.18E-57** | 1.48 | (1.4,1.55) | **5.10E-140** | **7.14E-140** | -0.53 | (-0.63,-0.42) | **1.64E-18** | **2.10E-18** | -0.30 | (-0.39,-0.21) | **7.76E-11** | **1.94E-10** |
| 254 | LPC(22:5) (n3) [sn1] [104_sn1] | 1.35 | (1.28,1.42) | **2.39E-134** | **2.17E-133** | 0.72 | (0.66,0.78) | **1.34E-85** | **1.61E-85** | 0.40 | (0.31,0.5) | **3.21E-16** | **3.98E-16** | 0.16 | (0.08,0.24) | **1.34E-04** | **1.94E-04** |
| 255 | LPC(22:6) [sn2] | 0.45 | (0.39,0.5) | **1.20E-43** | **2.05E-43** | 0.67 | (0.61,0.72) | **1.71E-87** | **2.06E-87** | -0.60 | (-0.69,-0.52) | **4.76E-31** | **6.88E-31** | 0.31 | (0.23,0.39) | **9.79E-14** | **3.15E-13** |
| 256 | LPC(22:6) [sn1] | 0.92 | (0.86,0.98) | **2.77E-102** | **1.26E-101** | 1.35 | (1.3,1.41) | **2.57E-179** | **4.23E-179** | -0.78 | (-0.87,-0.69) | **6.85E-42** | **1.09E-41** | 0.28 | (0.2,0.36) | **1.04E-10** | **2.54E-10** |
| 257 | LPC(O-16:0) | 1.08 | (1.03,1.12) | **9.33E-157** | **1.44E-155** | 0.86 | (0.82,0.9) | **1.99E-158** | **3.01E-158** | -0.16 | (-0.23,-0.09) | **1.05E-05** | **1.13E-05** | 0.36 | (0.29,0.42) | **2.70E-21** | **1.53E-20** |
| 258 | LPC(O-18:0) | 0.89 | (0.84,0.95) | **1.14E-110** | **6.53E-110** | -0.15 | (-0.19,-0.1) | **5.15E-09** | **5.43E-09** | 0.80 | (0.72,0.88) | **1.96E-48** | **3.37E-48** | 0.23 | (0.15,0.32) | **4.46E-08** | **8.67E-08** |
| 259 | LPC(O-18:1) | 0.99 | (0.94,1.04) | **3.86E-142** | **4.21E-141** | 0.56 | (0.52,0.6) | **1.60E-96** | **2.01E-96** | 0.19 | (0.11,0.26) | **9.16E-07** | **1.01E-06** | 0.23 | (0.16,0.29) | **8.51E-11** | **2.11E-10** |
| 260 | LPC(O-20:0) | 0.91 | (0.86,0.96) | **8.82E-116** | **5.72E-115** | -0.03 | (-0.09,0.04) | 4.26E-01 | 4.31E-01 | 0.81 | (0.71,0.91) | **2.32E-39** | **3.59E-39** | 0.14 | (0.06,0.22) | **4.06E-04** | **5.68E-04** |
| 261 | LPC(O-20:1) | 0.97 | (0.89,1.04) | **1.73E-81** | **5.09E-81** | 0.01 | (-0.06,0.09) | 7.09E-01 | 7.14E-01 | 0.31 | (0.2,0.42) | **3.74E-08** | **4.25E-08** | 0.59 | (0.5,0.67) | **8.39E-31** | **1.30E-29** |
| 262 | LPC(O-22:0) | 0.47 | (0.42,0.51) | **2.15E-71** | **5.28E-71** | -0.17 | (-0.21,-0.13) | **1.27E-13** | **1.36E-13** | 0.37 | (0.32,0.42) | **4.71E-32** | **6.91E-32** | 0.22 | (0.17,0.28) | **5.11E-14** | **1.68E-13** |
| 263 | LPC(O-22:1) | 0.70 | (0.65,0.75) | **1.16E-97** | **4.66E-97** | -0.15 | (-0.19,-0.1) | **3.34E-10** | **3.53E-10** | 0.58 | (0.51,0.65) | **2.00E-39** | **3.12E-39** | 0.25 | (0.17,0.32) | **5.43E-10** | **1.25E-09** |
| 264 | LPC(O-24:0) | 0.14 | (0.09,0.18) | **3.11E-09** | **3.91E-09** | -0.90 | (-0.95,-0.85) | **1.04E-130** | **1.43E-130** | 0.79 | (0.74,0.85) | **5.47E-74** | **1.30E-73** | 0.21 | (0.15,0.27) | **6.54E-12** | **1.79E-11** |
| 265 | LPC(O-24:1) | 0.26 | (0.22,0.3) | **1.71E-28** | **2.62E-28** | -0.81 | (-0.86,-0.76) | **6.29E-120** | **8.46E-120** | 0.81 | (0.75,0.88) | **3.23E-70** | **7.10E-70** | 0.22 | (0.16,0.28) | **8.25E-12** | **2.24E-11** |
| 266 | LPC(O-24:2) | 0.45 | (0.4,0.5) | **1.19E-53** | **2.27E-53** | 0.06 | (0.01,0.1) | **1.70E-02** | **1.75E-02** | 0.02 | (-0.05,0.09) | 5.97E-01 | 6.01E-01 | 0.34 | (0.26,0.41) | **5.74E-17** | **2.47E-16** |
| 267 | LPC(P-16:0) | 1.37 | (1.32,1.42) | **2.24E-180** | **9.77E-179** | 0.86 | (0.82,0.9) | **4.71E-161** | **7.18E-161** | 0.05 | (-0.02,0.12) | 1.81E-01 | 1.85E-01 | 0.44 | (0.37,0.51) | **4.38E-31** | **7.51E-30** |
| 268 | LPC(P-18:0) | 0.99 | (0.94,1.03) | **2.00E-141** | **2.14E-140** | 0.56 | (0.52,0.6) | **1.07E-93** | **1.32E-93** | 0.21 | (0.13,0.28) | **7.28E-08** | **8.23E-08** | 0.23 | (0.16,0.29) | **2.41E-10** | **5.74E-10** |
| 269 | LPC(P-18:1) | 1.24 | (1.18,1.29) | **1.82E-144** | **2.13E-143** | -0.10 | (-0.15,-0.05) | **1.97E-04** | **2.05E-04** | 0.78 | (0.69,0.87) | **2.80E-44** | **4.55E-44** | 0.52 | (0.44,0.6) | **7.00E-29** | **8.00E-28** |
| 270 | LPC(P-20:0) | 0.87 | (0.81,0.92) | **3.13E-104** | **1.49E-103** | -0.12 | (-0.17,-0.07) | **1.37E-06** | **1.44E-06** | 0.81 | (0.73,0.9) | **4.55E-52** | **8.11E-52** | 0.18 | (0.1,0.26) | **8.73E-06** | **1.39E-05** |
| 271 | LPE(16:0) [sn2] | -0.16 | (-0.22,-0.11) | **1.02E-08** | **1.27E-08** | -0.13 | (-0.18,-0.08) | **2.29E-07** | **2.41E-07** | -0.53 | (-0.61,-0.45) | **1.94E-30** | **2.79E-30** | 0.47 | (0.4,0.54) | **1.28E-31** | **2.46E-30** |

| | | | | | | | | | | | | | | | | | |
|---|---|---|---|---|---|---|---|---|---|---|---|---|---|---|---|---|---|
| 272 | LPE(16:0) [sn1] | -0.24 | (-0.29,-0.19) | 2.66E-18 | 3.66E-18 | -0.23 | (-0.28,-0.18) | 1.77E-19 | 1.91E-19 | -0.51 | (-0.58,-0.43) | 6.59E-30 | 9.42E-30 | 0.48 | (0.41,0.55) | 3.39E-34 | 9.05E-33 |
| 273 | LPE(18:0) [sn2] | 0.22 | (0.17,0.27) | 1.89E-17 | 2.58E-17 | -0.32 | (-0.37,-0.28) | 4.59E-38 | 5.15E-38 | 0.17 | (0.1,0.24) | 2.00E-06 | 2.20E-06 | 0.36 | (0.3,0.42) | 2.59E-24 | 1.95E-23 |
| 274 | LPE(18:0) [sn1] | 0.13 | (0.08,0.18) | 5.51E-08 | 6.79E-08 | -0.43 | (-0.47,-0.39) | 4.96E-62 | 5.78E-62 | 0.20 | (0.13,0.27) | 2.42E-08 | 2.76E-08 | 0.37 | (0.31,0.43) | 3.93E-25 | 3.14E-24 |
| 275 | LPE(18:1) [sn2] | 0.05 | (-0.02,0.12) | 1.47E-01 | 1.55E-01 | -0.29 | (-0.34,-0.24) | 7.85E-25 | 8.54E-25 | 0.20 | (0.11,0.29) | 1.02E-05 | 1.12E-05 | 0.09 | (0,0.18) | 4.08E-02 | 4.75E-02 |
| 276 | LPE(18:1) [sn1] | 0.07 | (0.01,0.13) | 3.44E-02 | 3.75E-02 | -0.33 | (-0.39,-0.28) | 1.63E-30 | 1.80E-30 | 0.31 | (0.22,0.4) | 5.22E-11 | 6.15E-11 | 0.09 | (0,0.18) | 4.82E-02 | 5.55E-02 |
| 277 | LPE(18:2) [sn2] | 0.73 | (0.66,0.81) | 5.08E-62 | 1.09E-61 | -0.26 | (-0.33,-0.2) | 1.39E-14 | 1.49E-14 | 0.86 | (0.76,0.96) | 5.86E-44 | 9.47E-44 | 0.13 | (0.05,0.22) | 3.09E-03 | 3.98E-03 |
| 278 | LPE(18:2) [sn1] | 0.92 | (0.84,0.99) | 1.21E-79 | 3.42E-79 | -0.03 | (-0.09,0.03) | 3.17E-01 | 3.22E-01 | 0.84 | (0.74,0.94) | 3.43E-43 | 5.47E-43 | 0.12 | (0.04,0.21) | 5.45E-03 | 6.83E-03 |
| 279 | LPE(20:4) [sn1] | 1.03 | (0.97,1.08) | 1.48E-124 | 1.07E-123 | 2.19 | (2.14,2.25) | 7.71E-263 | 2.96E-262 | -1.35 | (-1.42,-1.27) | 6.01E-97 | 2.20E-96 | 0.18 | (0.12,0.24) | 1.88E-08 | 3.77E-08 |
| 280 | LPE(22:6) [sn2] | 0.36 | (0.3,0.42) | 4.81E-28 | 7.30E-28 | 1.53 | (1.47,1.58) | 8.90E-203 | 1.70E-202 | -1.68 | (-1.77,-1.6) | 9.92E-105 | 4.04E-104 | 0.51 | (0.43,0.59) | 1.38E-29 | 1.78E-28 |
| 281 | LPE(22:6) [sn1] | 0.62 | (0.56,0.68) | 2.03E-62 | 4.40E-62 | 1.87 | (1.81,1.92) | 2.09E-233 | 5.66E-233 | -1.76 | (-1.84,-1.68) | 8.29E-112 | 3.98E-111 | 0.50 | (0.42,0.57) | 5.79E-31 | 9.59E-30 |
| 282 | PC(28:0) | -0.79 | (-0.9,-0.67) | 2.16E-34 | 3.47E-34 | -1.92 | (-2.01,-1.84) | 4.21E-159 | 6.37E-159 | 1.06 | (0.95,1.18) | 3.75E-46 | 6.25E-46 | 0.04 | (-0.11,0.18) | 6.16E-01 | 6.37E-01 |
| 283 | PC(14:0_16:0) | -0.97 | (-1.05,-0.9) | 2.56E-86 | 8.20E-86 | -1.74 | (-1.81,-1.67) | 1.33E-182 | 2.23E-182 | 0.59 | (0.5,0.68) | 5.35E-29 | 7.57E-29 | 0.15 | (0.06,0.23) | 1.25E-03 | 1.67E-03 |
| 284 | PC(31:0) (a) | -0.83 | (-0.93,-0.73) | 2.10E-45 | 3.68E-45 | -1.79 | (-1.88,-1.7) | 5.90E-145 | 8.48E-145 | 1.21 | (1.06,1.36) | 5.26E-39 | 8.12E-39 | -0.28 | (-0.44,-0.13) | 2.92E-04 | 4.14E-04 |
| 285 | PC(31:0) (b) | -0.64 | (-0.7,-0.58) | 2.23E-67 | 5.17E-67 | -1.19 | (-1.27,-1.12) | 3.06E-115 | 4.04E-115 | 0.53 | (0.43,0.64) | 1.50E-20 | 1.95E-20 | -0.01 | (-0.09,0.08) | 9.10E-01 | 9.16E-01 |
| 286 | PC(31:1) | -1.09 | (-1.18,-1) | 8.11E-78 | 2.26E-77 | -1.61 | (-1.69,-1.53) | 5.63E-142 | 7.97E-142 | 0.37 | (0.25,0.48) | 2.67E-09 | 3.08E-09 | 0.15 | (0.03,0.27) | 1.64E-02 | 1.99E-02 |
| 287 | PC(16:0_16:0) | -0.71 | (-0.76,-0.66) | 3.93E-99 | 1.63E-98 | -1.00 | (-1.05,-0.94) | 1.05E-128 | 1.43E-128 | 0.07 | (0,0.14) | 6.63E-02 | 6.87E-02 | 0.19 | (0.13,0.25) | 3.48E-10 | 8.14E-10 |
| 288 | PC(32:1) | -1.20 | (-1.29,-1.12) | 1.98E-94 | 7.53E-94 | -1.18 | (-1.26,-1.1) | 1.39E-103 | 1.76E-103 | -0.13 | (-0.24,-0.03) | 1.17E-02 | 1.23E-02 | 0.12 | (0.03,0.21) | 1.00E-02 | 1.24E-02 |
| 289 | PC(32:2) | -1.05 | (-1.13,-0.97) | 1.28E-89 | 4.26E-89 | -3.52 | (-3.59,-3.46) | 0.00E+00 | 0.00E+00 | 2.30 | (2.2,2.39) | 5.13E-119 | 2.90E-118 | 0.16 | (0.06,0.27) | 2.53E-03 | 3.28E-03 |
| 290 | PC(33:0) (a) | -0.81 | (-0.88,-0.73) | 1.06E-68 | 2.47E-68 | -1.00 | (-1.08,-0.92) | 1.55E-87 | 1.87E-87 | 0.15 | (0.02,0.28) | 2.18E-02 | 2.27E-02 | -0.01 | (-0.12,0.11) | 9.04E-01 | 9.12E-01 |
| 291 | PC(33:0) (b) | -0.38 | (-0.43,-0.32) | 2.54E-36 | 4.15E-36 | -1.45 | (-1.51,-1.39) | 4.13E-166 | 6.47E-166 | 0.96 | (0.86,1.05) | 2.34E-52 | 4.21E-52 | 0.11 | (0.04,0.19) | 4.61E-03 | 5.82E-03 |
| 292 | PC(33:1) | -1.17 | (-1.24,-1.1) | 6.03E-111 | 3.49E-110 | -2.30 | (-2.36,-2.23) | 8.93E-242 | 2.75E-241 | 1.05 | (0.96,1.15) | 2.28E-58 | 4.28E-58 | 0.08 | (-0.01,0.18) | 8.69E-02 | 9.84E-02 |
| 293 | PC(33:2) | -0.83 | (-0.88,-0.77) | 1.16E-96 | 4.59E-96 | -4.20 | (-4.26,-4.15) | 0.00E+00 | 0.00E+00 | 3.10 | (3.01,3.19) | 6.57E-157 | 1.17E-155 | 0.24 | (0.16,0.33) | 4.87E-08 | 9.42E-08 |
| 294 | PC(16:0_18:0) | 0.08 | (0.05,0.12) | 2.42E-05 | 2.85E-05 | -0.86 | (-0.91,-0.82) | 2.12E-135 | 2.95E-135 | 0.77 | (0.71,0.83) | 1.28E-66 | 2.65E-66 | 0.18 | (0.12,0.23) | 4.66E-09 | 9.90E-09 |
| 295 | PC(16:0_18:1) | -1.07 | (-1.13,-1.01) | 7.65E-122 | 5.48E-121 | -2.39 | (-2.45,-2.33) | 2.30E-263 | 8.98E-263 | 1.27 | (1.18,1.35) | 5.88E-81 | 1.58E-80 | 0.07 | (0,0.15) | 5.10E-02 | 5.86E-02 |
| 296 | PC(16:0_18:2) | -0.55 | (-0.59,-0.52) | 1.55E-106 | 7.71E-106 | -3.55 | (-3.6,-3.49) | 0.00E+00 | 0.00E+00 | 2.80 | (2.73,2.88) | 3.80E-167 | 1.01E-165 | 0.18 | (0.13,0.23) | 7.52E-11 | 1.89E-10 |
| 297 | PC(16:1_18:2) | -0.70 | (-0.75,-0.64) | 9.18E-78 | 2.55E-77 | -3.01 | (-3.06,-2.95) | 0.00E+00 | 0.00E+00 | 1.95 | (1.86,2.03) | 7.01E-119 | 3.91E-118 | 0.37 | (0.3,0.44) | 1.23E-20 | 6.48E-20 |
| 298 | PC(16:0_18:3) (a) | -1.04 | (-1.11,-0.97) | 3.63E-101 | 1.58E-100 | -3.68 | (-3.75,-3.61) | 0.00E+00 | 0.00E+00 | 2.51 | (2.42,2.6) | 1.84E-133 | 1.67E-132 | 0.13 | (0.05,0.22) | 1.88E-03 | 2.46E-03 |
| 299 | PC(14:0_20:4) | -0.97 | (-1.05,-0.89) | 1.07E-77 | 2.94E-77 | -3.02 | (-3.09,-2.95) | 1.38E-280 | 7.37E-280 | 1.76 | (1.65,1.86) | 8.48E-89 | 2.68E-88 | 0.28 | (0.16,0.4) | 7.89E-06 | 1.27E-05 |
| 300 | PC(34:5) | -0.55 | (-0.66,-0.43) | 3.15E-19 | 4.35E-19 | -3.53 | (-3.63,-3.44) | 1.85E-251 | 6.38E-251 | 2.16 | (2.01,2.31) | 3.36E-79 | 8.67E-79 | 0.81 | (0.64,0.98) | 4.07E-18 | 1.88E-17 |
| 301 | PC(15-MHDA_18:1) | -0.54 | (-0.61,-0.46) | 6.96E-40 | 1.16E-39 | -1.93 | (-2,-1.86) | 2.75E-191 | 4.83E-191 | 1.47 | (1.35,1.59) | 4.32E-65 | 8.76E-65 | -0.09 | (-0.2,0.02) | 1.01E-01 | 1.14E-01 |
| 302 | PC(17:0_18:1) | -0.59 | (-0.64,-0.54) | 3.15E-75 | 8.17E-75 | -1.88 | (-1.92,-1.83) | 3.81E-260 | 1.41E-259 | 1.29 | (1.22,1.37) | 2.04E-94 | 6.99E-94 | 0.01 | (-0.06,0.08) | 7.53E-01 | 7.69E-01 |
| 303 | PC(15-MHDA_18:2) | -0.66 | (-0.72,-0.59) | 3.71E-58 | 7.43E-58 | -3.26 | (-3.33,-3.2) | 2.60E-299 | 2.31E-298 | 2.48 | (2.37,2.6) | 4.64E-113 | 2.37E-112 | 0.08 | (-0.03,0.19) | 1.36E-01 | 1.51E-01 |
| 304 | PC(17:0_18:2) | -0.47 | (-0.51,-0.42) | 5.84E-60 | 1.22E-59 | -3.29 | (-3.33,-3.24) | 0.00E+00 | 0.00E+00 | 2.63 | (2.55,2.71) | 2.39E-154 | 3.70E-153 | 0.18 | (0.1,0.25) | 3.91E-06 | 6.47E-06 |
| 305 | PC(17:1_18:2) | -0.77 | (-0.82,-0.72) | 2.23E-95 | 8.70E-95 | -2.61 | (-2.66,-2.56) | 0.00E+00 | 0.00E+00 | 1.63 | (1.55,1.71) | 6.28E-103 | 2.49E-102 | 0.22 | (0.14,0.3) | 2.06E-07 | 3.81E-07 |
| 306 | PC(15:0_20:3) | -0.30 | (-0.33,-0.28) | 7.50E-65 | 1.67E-64 | -2.84 | (-2.9,-2.77) | 8.21E-281 | 4.43E-280 | 2.42 | (2.33,2.5) | 3.91E-141 | 4.17E-140 | 0.12 | (0.08,0.16) | 2.09E-08 | 4.18E-08 |
| 307 | PC(15:0_20:4) | -0.78 | (-0.84,-0.72) | 3.33E-90 | 1.16E-89 | -2.46 | (-2.52,-2.4) | 5.47E-269 | 2.36E-268 | 1.23 | (1.13,1.33) | 3.58E-65 | 7.29E-65 | 0.43 | (0.34,0.52) | 1.56E-17 | 6.94E-17 |
| 308 | PC(35:5) | -0.34 | (-0.45,-0.23) | 8.03E-10 | 1.01E-09 | -3.25 | (-3.34,-3.16) | 1.49E-247 | 4.96E-247 | 2.05 | (1.89,2.21) | 5.72E-69 | 1.22E-68 | 0.87 | (0.7,1.04) | 1.34E-20 | 7.01E-20 |
| 309 | PC 36:0 | -0.43 | (-0.48,-0.37) | 1.94E-39 | 3.21E-39 | 1.06 | (1.01,1.12) | 2.47E-148 | 3.60E-148 | -1.53 | (-1.61,-1.46) | 5.78E-109 | 2.54E-108 | -0.01 | (-0.09,0.07) | 8.15E-01 | 8.31E-01 |
| 310 | PC(18:0_18:1) | -0.35 | (-0.41,-0.29) | 6.71E-27 | 1.00E-26 | -1.28 | (-1.34,-1.23) | 2.43E-172 | 3.88E-172 | 0.98 | (0.91,1.06) | 2.26E-71 | 5.09E-71 | -0.01 | (-0.08,0.06) | 7.13E-01 | 7.32E-01 |
| 311 | PC(18:0_18:2) | -0.07 | (-0.12,-0.01) | 1.17E-02 | 1.30E-02 | -3.20 | (-3.25,-3.15) | 0.00E+00 | 0.00E+00 | 3.05 | (2.97,3.13) | 7.85E-172 | 2.51E-170 | 0.10 | (0.03,0.17) | 6.64E-03 | 8.30E-03 |
| 312 | PC(16:0_20:3) (a) | -0.58 | (-0.63,-0.53) | 5.90E-67 | 1.34E-66 | -3.44 | (-3.5,-3.38) | 0.00E+00 | 0.00E+00 | 2.71 | (2.62,2.8) | 5.55E-143 | 6.20E-142 | 0.19 | (0.12,0.27) | 2.28E-06 | 3.85E-06 |
| 313 | PC(16:0_20:3) (b) | -1.39 | (-1.46,-1.32) | 3.50E-135 | 3.29E-134 | -2.00 | (-2.07,-1.93) | 4.78E-214 | 1.04E-213 | 0.39 | (0.31,0.48) | 6.35E-18 | 8.06E-18 | 0.21 | (0.13,0.3) | 4.37E-07 | 7.83E-07 |
| 314 | PC(18:2_18:2) | -0.16 | (-0.23,-0.09) | 1.08E-05 | 1.29E-05 | -5.35 | (-5.42,-5.28) | 0.00E+00 | 0.00E+00 | 5.00 | (4.88,5.12) | 4.60E-177 | 1.84E-175 | 0.16 | (0.05,0.26) | 4.91E-03 | 6.19E-03 |
| 315 | PC(16:1_20:4) | -0.61 | (-0.67,-0.55) | 1.45E-61 | 3.07E-61 | -1.48 | (-1.54,-1.42) | 6.95E-174 | 1.12E-173 | 0.43 | (0.33,0.52) | 1.24E-17 | 1.56E-17 | 0.43 | (0.35,0.5) | 4.29E-23 | 2.79E-22 |
| 316 | PC(16:0_20:5) | -0.42 | (-0.52,-0.32) | 3.42E-15 | 4.52E-15 | -2.66 | (-2.75,-2.58) | 1.23E-227 | 3.17E-227 | 1.30 | (1.16,1.44) | 9.68E-47 | 1.62E-46 | 0.93 | (0.78,1.08) | 5.25E-28 | 5.60E-27 |
| 317 | PC(36:6) | -1.15 | (-1.23,-1.07) | 6.48E-93 | 2.41E-92 | -3.32 | (-3.39,-3.25) | 3.29E-300 | 2.98E-299 | 1.71 | (1.61,1.82) | 8.74E-87 | 2.62E-86 | 0.42 | (0.3,0.55) | 1.60E-10 | 3.89E-10 |
| 318 | PC(15-MHDA_20:4) | -0.84 | (-0.9,-0.78) | 2.63E-86 | 8.35E-86 | -1.74 | (-1.81,-1.68) | 1.36E-187 | 2.34E-187 | 0.61 | (0.49,0.73) | 2.96E-20 | 3.84E-20 | 0.25 | (0.14,0.35) | 6.40E-06 | 1.03E-05 |
| 319 | PC(17:0_20:4) | -0.45 | (-0.49,-0.4) | 1.37E-54 | 2.63E-54 | -1.06 | (-1.12,-1.01) | 1.05E-149 | 1.54E-149 | 0.26 | (0.17,0.35) | 2.53E-08 | 2.88E-08 | 0.33 | (0.26,0.41) | 1.60E-15 | 6.11E-15 |
| 320 | PC(15:0_22:6) | -0.85 | (-0.91,-0.79) | 3.27E-89 | 1.08E-88 | -2.74 | (-2.8,-2.68) | 1.78E-288 | 1.08E-287 | 1.43 | (1.32,1.54) | 2.39E-72 | 5.53E-72 | 0.41 | (0.31,0.51) | 2.62E-14 | 8.79E-14 |
| 321 | PC(38:2) | -0.56 | (-0.6,-0.51) | 9.10E-81 | 2.63E-80 | -1.29 | (-1.33,-1.24) | 8.52E-204 | 1.64E-203 | 0.57 | (0.51,0.63) | 1.70E-46 | 2.85E-46 | 0.17 | (0.11,0.22) | 1.54E-08 | 3.12E-08 |
| 322 | PC(18:0_20:3) | -0.47 | (-0.53,-0.4) | 2.35E-33 | 3.75E-33 | -0.29 | (-0.35,-0.23) | 4.92E-20 | 5.33E-20 | -0.28 | (-0.36,-0.19) | 8.47E-10 | 9.84E-10 | 0.10 | (0.01,0.19) | 3.38E-02 | 3.98E-02 |
| 323 | PC(18:1_20:3) | -0.86 | (-0.92,-0.8) | 1.76E-101 | 7.75E-101 | -1.78 | (-1.84,-1.73) | 1.26E-216 | 2.80E-216 | 0.73 | (0.65,0.81) | 7.10E-46 | 1.18E-45 | 0.20 | (0.12,0.27) | 2.20E-07 | 4.02E-07 |
| 324 | PC(38:4) (b) | -1.10 | (-1.17,-1.03) | 6.30E-107 | 3.22E-106 | -1.78 | (-1.85,-1.71) | 7.38E-188 | 1.27E-187 | 0.75 | (0.67,0.84) | 5.38E-43 | 8.55E-43 | -0.04 | (-0.12,0.05) | 3.91E-01 | 4.14E-01 |
| 325 | PC(18:0_20:4) | 0.03 | (-0.02,0.08) | 2.19E-01 | 2.30E-01 | -0.20 | (-0.25,-0.15) | 4.67E-13 | 4.98E-13 | -0.03 | (-0.11,0.05) | 3.95E-01 | 4.01E-01 | 0.25 | (0.18,0.32) | 8.54E-12 | 2.28E-11 |
| 326 | PC(38:5) (a) | -0.41 | (-0.46,-0.37) | 6.44E-52 | 1.21E-51 | -2.26 | (-2.31,-2.21) | 1.67E-291 | 1.14E-290 | 1.54 | (1.47,1.62) | 4.76E-109 | 2.12E-108 | 0.31 | (0.24,0.38) | 4.75E-18 | 2.17E-17 |

| | | | | | | | | | | | | | | | | | |
|---|---|---|---|---|---|---|---|---|---|---|---|---|---|---|---|---|---|
| 327 | PC(38:5) (b) | -0.78 | (-0.85,-0.71) | **2.34E-70** | **5.56E-70** | -1.94 | (-2,-1.87) | **2.00E-207** | **3.97E-207** | 0.86 | (0.76,0.95) | **8.91E-45** | **1.46E-44** | 0.30 | (0.19,0.4) | **2.87E-08** | **5.70E-08** |
| 328 | PC(38:6) (a) | -0.13 | (-0.18,-0.07) | **4.12E-06** | **4.94E-06** | -2.94 | (-3,-2.88) | **2.89E-295** | **2.24E-294** | 2.38 | (2.29,2.47) | **8.17E-128** | **5.68E-127** | 0.43 | (0.35,0.51) | **7.19E-22** | **4.21E-21** |
| 329 | PC(16:0_22:6) | -0.88 | (-0.94,-0.82) | **1.05E-101** | **4.66E-101** | -1.66 | (-1.71,-1.6) | **5.33E-213** | **1.13E-212** | 0.28 | (0.19,0.37) | **1.59E-09** | **1.85E-09** | 0.41 | (0.33,0.49) | **2.92E-20** | **1.46E-19** |
| 330 | PC(18:2_20:5) | 0.12 | (0.03,0.21) | **1.04E-02** | **1.15E-02** | -3.30 | (-3.38,-3.22) | **2.13E-276** | **1.03E-275** | 2.76 | (2.64,2.89) | **3.20E-113** | **1.65E-112** | 0.68 | (0.55,0.81) | **1.23E-20** | **6.48E-20** |
| 331 | PC(16:1_22:6) | -0.72 | (-0.78,-0.67) | **6.63E-79** | **1.87E-78** | -1.72 | (-1.77,-1.66) | **1.08E-218** | **2.47E-218** | 0.50 | (0.41,0.59) | **8.17E-24** | **1.08E-23** | 0.47 | (0.38,0.56) | **1.10E-20** | **5.94E-20** |
| 332 | PC(38:7)(c) | -0.85 | (-0.9,-0.8) | **3.27E-115** | **2.09E-114** | -1.93 | (-1.97,-1.88) | **1.24E-268** | **5.32E-268** | 0.63 | (0.55,0.7) | **6.13E-40** | **9.58E-40** | 0.39 | (0.31,0.46) | **4.00E-21** | **2.21E-20** |
| 333 | PC(39:5)(a) | -0.58 | (-0.64,-0.51) | **1.03E-49** | **1.89E-49** | -2.08 | (-2.15,-2.01) | **6.44E-216** | **1.41E-215** | 1.16 | (1.05,1.28) | **4.54E-51** | **8.01E-51** | 0.28 | (0.17,0.39) | **7.38E-07** | **1.28E-06** |
| 334 | PC(39:5)(b) | -0.70 | (-0.77,-0.64) | **3.50E-67** | **8.08E-67** | -2.15 | (-2.22,-2.09) | **8.22E-232** | **2.19E-231** | 1.15 | (1.05,1.26) | **2.53E-56** | **4.69E-56** | 0.25 | (0.14,0.35) | **3.67E-06** | **6.11E-06** |
| 335 | PC(15-MHDA_22:6) | -0.48 | (-0.54,-0.42) | **7.01E-40** | **1.16E-39** | -2.08 | (-2.15,-2.01) | **1.87E-211** | **3.81E-211** | 1.35 | (1.24,1.47) | **2.44E-62** | **4.80E-62** | 0.20 | (0.09,0.3) | **2.04E-04** | **2.93E-04** |
| 336 | PC(17:0_22:6) | -0.79 | (-0.85,-0.73) | **1.37E-83** | **4.22E-83** | -2.13 | (-2.19,-2.07) | **1.96E-247** | **6.48E-247** | 0.88 | (0.77,0.99) | **2.44E-38** | **3.73E-38** | 0.38 | (0.28,0.48) | **3.38E-12** | **9.75E-12** |
| 337 | PC(18:0_22:5) (n6) | -0.24 | (-0.31,-0.17) | **5.42E-12** | **6.86E-12** | -1.83 | (-1.92,-1.74) | **3.38E-147** | **4.91E-147** | 1.42 | (1.28,1.57) | **3.43E-50** | **5.99E-50** | 0.04 | (-0.05,0.13) | 3.78E-01 | 4.02E-01 |
| 338 | PC(18:0_22:6) | -0.55 | (-0.61,-0.49) | **8.13E-48** | **1.46E-47** | -1.30 | (-1.36,-1.24) | **5.90E-165** | **9.16E-165** | 0.36 | (0.26,0.46) | **1.02E-11** | **1.22E-11** | 0.34 | (0.25,0.43) | **2.19E-12** | **6.40E-12** |
| 339 | PC(40:7) (a) | -0.51 | (-0.57,-0.45) | **1.38E-48** | **2.49E-48** | -1.30 | (-1.37,-1.22) | **2.64E-125** | **3.59E-125** | 0.44 | (0.33,0.56) | **1.22E-12** | **1.47E-12** | 0.31 | (0.23,0.39) | **2.78E-13** | **8.61E-13** |
| 340 | PC(18:1_22:6) (a) | -0.86 | (-0.91,-0.81) | **1.01E-114** | **6.40E-114** | -1.47 | (-1.53,-1.42) | **8.38E-193** | **1.49E-192** | 0.35 | (0.27,0.43) | **1.72E-15** | **2.11E-15** | 0.26 | (0.19,0.33) | **3.62E-12** | **1.03E-11** |
| 341 | PC(40:8) | -0.31 | (-0.36,-0.26) | **1.61E-32** | **2.54E-32** | -1.57 | (-1.62,-1.51) | **3.15E-196** | **5.79E-196** | 0.94 | (0.86,1.02) | **3.73E-62** | **7.29E-62** | 0.30 | (0.23,0.37) | **3.04E-15** | **1.12E-14** |
| 342 | PC(44:12) | -0.71 | (-0.81,-0.6) | **3.01E-33** | **4.79E-33** | -1.75 | (-1.83,-1.67) | **4.22E-164** | **6.52E-164** | 0.41 | (0.27,0.54) | **2.86E-08** | **3.25E-08** | 0.60 | (0.45,0.74) | **1.80E-14** | **6.12E-14** |
| 343 | PC(O-16:0/16:0) | -0.24 | (-0.27,-0.2) | **2.56E-31** | **4.00E-31** | -0.82 | (-0.87,-0.77) | **1.39E-126** | **1.89E-126** | 0.47 | (0.41,0.54) | **1.17E-35** | **1.76E-35** | 0.09 | (0.04,0.14) | **6.63E-04** | **9.09E-04** |
| 344 | PC(O-32:1) | -0.46 | (-0.51,-0.41) | **1.41E-59** | **2.90E-59** | -1.05 | (-1.11,-1) | **5.94E-141** | **8.34E-141** | 0.27 | (0.2,0.35) | **1.23E-11** | **1.46E-11** | 0.33 | (0.27,0.39) | **4.27E-21** | **2.33E-20** |
| 345 | PC(O-34:1) | -0.25 | (-0.29,-0.21) | **6.63E-34** | **1.06E-33** | -1.42 | (-1.46,-1.37) | **4.72E-213** | **1.00E-212** | 1.09 | (1.03,1.15) | **2.23E-96** | **7.97E-96** | 0.06 | (0.01,0.11) | **1.67E-02** | **2.02E-02** |
| 346 | PC(O-34:2) | -0.31 | (-0.37,-0.25) | **5.46E-23** | **7.83E-23** | -3.80 | (-3.86,-3.75) | **0.00E+00** | **0.00E+00** | 3.17 | (3.09,3.26) | **3.21E-166** | **8.12E-165** | 0.29 | (0.21,0.38) | **5.38E-11** | **1.37E-10** |
| 347 | PC(O-34:4) | -0.03 | (-0.12,0.07) | 5.61E-01 | 5.78E-01 | -2.39 | (-2.47,-2.3) | **1.01E-200** | **1.90E-200** | 1.34 | (1.22,1.46) | **1.17E-58** | **2.20E-58** | 1.01 | (0.88,1.14) | **3.09E-38** | **1.48E-36** |
| 348 | PC(O-35:4) | -0.02 | (-0.1,0.06) | 6.60E-01 | 6.77E-01 | -2.92 | (-3,-2.85) | **2.96E-259** | **1.08E-258** | 2.13 | (2.01,2.25) | **1.12E-95** | **3.96E-95** | 0.78 | (0.66,0.9) | **8.39E-29** | **9.37E-28** |
| 349 | PC(O-36:0) | -0.27 | (-0.31,-0.22) | **3.61E-27** | **5.43E-27** | -1.51 | (-1.57,-1.45) | **3.94E-186** | **6.73E-186** | 1.17 | (1.09,1.25) | **8.09E-76** | **2.01E-75** | 0.04 | (-0.02,0.1) | 1.87E-01 | 2.05E-01 |
| 350 | PC(O-18:0/18:1) | -0.15 | (-0.19,-0.11) | **2.13E-13** | **2.77E-13** | -1.51 | (-1.56,-1.46) | **9.29E-221** | **2.17E-220** | 1.32 | (1.26,1.38) | **2.47E-111** | **1.16E-110** | 0.00 | (-0.05,0.06) | 8.67E-01 | 8.80E-01 |
| 351 | PC(O-18:1/18:1) | -0.12 | (-0.15,-0.08) | **6.94E-09** | **8.68E-09** | -1.90 | (-1.94,-1.86) | **1.58E-278** | **8.32E-278** | 1.64 | (1.58,1.7) | **1.61E-136** | **1.55E-135** | 0.12 | (0.07,0.18) | **8.93E-06** | **1.42E-05** |
| 352 | PC(O-18:0/18:2) | -0.19 | (-0.24,-0.14) | **2.21E-12** | **2.82E-12** | -3.33 | (-3.38,-3.27) | **0.00E+00** | **0.00E+00** | 2.92 | (2.84,2.99) | **4.24E-172** | **1.46E-170** | 0.15 | (0.08,0.23) | **2.70E-05** | **4.10E-05** |
| 353 | PC(O-18:1/18:2) | -0.24 | (-0.29,-0.19) | **1.86E-19** | **2.58E-19** | -3.24 | (-3.29,-3.19) | **0.00E+00** | **0.00E+00** | 2.75 | (2.67,2.82) | **7.14E-164** | **1.49E-162** | 0.23 | (0.16,0.31) | **6.63E-10** | **1.50E-09** |
| 354 | PC(O-16:0/20:3) | -0.30 | (-0.35,-0.25) | **1.47E-26** | **2.19E-26** | -1.93 | (-1.98,-1.88) | **7.92E-252** | **2.75E-251** | 1.48 | (1.41,1.55) | **5.96E-110** | **2.67E-109** | 0.14 | (0.07,0.2) | **7.66E-05** | **1.13E-04** |
| 355 | PC(O-16:0/20:4) | -0.17 | (-0.22,-0.13) | **3.79E-13** | **4.88E-13** | -1.55 | (-1.59,-1.5) | **7.04E-225** | **1.74E-224** | 1.15 | (1.08,1.22) | **7.00E-91** | **2.30E-90** | 0.17 | (0.1,0.23) | **4.97E-07** | **8.84E-07** |
| 356 | PC(O-36:5) | 0.08 | (-0.01,0.17) | 7.49E-02 | 8.06E-02 | -3.26 | (-3.33,-3.2) | **2.45E-295** | **1.93E-294** | 2.50 | (2.39,2.62) | **9.98E-113** | **4.99E-112** | 0.79 | (0.67,0.92) | **5.51E-29** | **6.45E-28** |
| 357 | PC(O-18:0/20:4) | -0.04 | (-0.08,0) | 5.34E-02 | 5.77E-02 | -1.12 | (-1.17,-1.08) | **2.80E-183** | **4.74E-183** | 1.05 | (0.98,1.11) | **4.20E-84** | **1.19E-83** | -0.02 | (-0.08,0.05) | 6.09E-01 | 6.32E-01 |
| 358 | PC(O-38:5) | -0.18 | (-0.22,-0.13) | **2.90E-15** | **3.85E-15** | -2.15 | (-2.2,-2.1) | **3.96E-278** | **2.02E-277** | 1.86 | (1.79,1.93) | **3.19E-131** | **2.78E-130** | 0.08 | (0.02,0.14) | **9.44E-03** | **1.17E-02** |
| 359 | PC(O-16:0/22:6) | -0.59 | (-0.65,-0.53) | **3.25E-62** | **6.99E-62** | -2.02 | (-2.07,-1.97) | **3.43E-267** | **1.43E-266** | 0.97 | (0.89,1.05) | **4.82E-65** | **9.71E-65** | 0.36 | (0.28,0.44) | **1.01E-17** | **4.58E-17** |
| 360 | PC(O-40:5) | -0.27 | (-0.31,-0.23) | **3.32E-36** | **5.40E-36** | -1.92 | (-1.96,-1.88) | **7.51E-285** | **4.29E-284** | 1.56 | (1.49,1.62) | **3.40E-128** | **2.47E-127** | 0.05 | (-0.01,0.11) | 1.09E-01 | 1.22E-01 |
| 361 | PC(O-18:0/22:6) | -0.25 | (-0.29,-0.2) | **4.21E-23** | **6.05E-23** | -1.45 | (-1.49,-1.41) | **8.22E-234** | **2.28E-233** | 1.03 | (0.96,1.1) | **1.86E-81** | **5.07E-81** | 0.09 | (0.03,0.16) | **4.28E-03** | **5.45E-03** |
| 362 | PC(O-40:7) (a) | -0.44 | (-0.49,-0.38) | **4.95E-44** | **8.52E-44** | -2.25 | (-2.3,-2.2) | **6.99E-290** | **4.42E-289** | 1.46 | (1.38,1.54) | **2.18E-96** | **7.88E-96** | 0.29 | (0.22,0.36) | **1.90E-14** | **6.41E-14** |
| 363 | PC(P-16:0/14:0) | -0.25 | (-0.3,-0.2) | **4.38E-20** | **6.11E-20** | -0.96 | (-1.02,-0.91) | **2.65E-119** | **3.55E-119** | 0.45 | (0.37,0.54) | **3.20E-21** | **4.19E-21** | 0.23 | (0.15,0.3) | **1.37E-08** | **2.79E-08** |
| 364 | PC(P-16:0/16:0) | -0.06 | (-0.1,-0.01) | **9.05E-03** | **1.01E-02** | -0.84 | (-0.89,-0.79) | **2.17E-111** | **2.81E-111** | 0.54 | (0.47,0.61) | **3.82E-37** | **5.82E-37** | 0.22 | (0.16,0.27) | **6.10E-12** | **1.68E-11** |
| 365 | PC(P-16:0/16:1) | -0.37 | (-0.42,-0.32) | **9.86E-45** | **1.71E-44** | -0.91 | (-0.96,-0.85) | **2.01E-114** | **2.62E-114** | 0.03 | (-0.05,0.1) | 4.58E-01 | 4.63E-01 | 0.50 | (0.43,0.57) | **4.79E-36** | **1.92E-34** |
| 366 | PC(P-16:0/18:0) | -0.02 | (-0.06,0.01) | 1.59E-01 | 1.68E-01 | -0.89 | (-0.94,-0.85) | **5.20E-154** | **7.83E-154** | 0.77 | (0.71,0.82) | **1.44E-73** | **3.40E-73** | 0.08 | (0.03,0.12) | **1.75E-03** | **2.30E-03** |
| 367 | PC(P-16:0/18:1) | -0.01 | (-0.05,0.03) | 7.15E-01 | 7.26E-01 | -0.98 | (-1.03,-0.93) | **1.89E-144** | **2.71E-144** | 0.67 | (0.61,0.74) | **6.87E-56** | **1.26E-55** | 0.26 | (0.2,0.33) | **1.37E-15** | **5.25E-15** |
| 368 | PC(P-16:0/18:2) | -0.11 | (-0.15,-0.06) | **2.72E-05** | **3.20E-05** | -3.54 | (-3.59,-3.49) | **0.00E+00** | **0.00E+00** | 3.28 | (3.2,3.35) | **6.74E-181** | **2.94E-179** | 0.12 | (0.04,0.19) | **3.35E-03** | **4.30E-03** |
| 369 | PC(P-16:0/18:3) | -0.01 | (-0.07,0.04) | 6.70E-01 | 6.86E-01 | -3.05 | (-3.1,-3) | **0.00E+00** | **0.00E+00** | 2.82 | (2.74,2.89) | **4.45E-164** | **1.02E-162** | 0.20 | (0.12,0.28) | **1.11E-06** | **1.92E-06** |
| 370 | PC(P-35:2) (a) | -0.22 | (-0.3,-0.14) | **4.55E-08** | **5.62E-08** | -3.80 | (-3.87,-3.72) | **9.39E-306** | **9.39E-305** | 3.31 | (3.21,3.42) | **1.18E-147** | **1.45E-146** | 0.14 | (0.03,0.25) | **1.21E-02** | **1.49E-02** |
| 371 | PC(P-35:2) (b) | -0.13 | (-0.19,-0.07) | **6.92E-05** | **8.12E-05** | -4.02 | (-4.09,-3.96) | **0.00E+00** | **0.00E+00** | 3.64 | (3.55,3.73) | **1.88E-170** | **5.64E-169** | 0.22 | (0.13,0.31) | **2.72E-06** | **4.55E-06** |
| 372 | PC(P-15:0/20:4) (b) | 0.07 | (0,0.13) | **3.86E-02** | **4.20E-02** | -1.56 | (-1.63,-1.5) | **1.03E-173** | **1.65E-173** | 1.44 | (1.35,1.53) | **6.14E-85** | **1.77E-84** | 0.15 | (0.06,0.24) | **8.97E-04** | **1.22E-03** |
| 373 | PC(P-18:1/18:1) | -0.07 | (-0.11,-0.03) | **8.48E-04** | **9.66E-04** | -1.27 | (-1.33,-1.21) | **9.41E-146** | **1.36E-145** | 0.87 | (0.79,0.95) | **2.91E-55** | **5.33E-55** | 0.25 | (0.19,0.31) | **1.63E-14** | **5.57E-14** |
| 374 | PC(P-18:0/18:2) | -0.03 | (-0.07,0.02) | 2.90E-01 | 3.00E-01 | -3.34 | (-3.39,-3.29) | **0.00E+00** | **0.00E+00** | 3.20 | (3.13,3.28) | **1.69E-181** | **8.13E-180** | 0.03 | (-0.04,0.11) | 3.80E-01 | 4.03E-01 |
| 375 | PC(P-16:0/20:4) | -0.14 | (-0.19,-0.09) | **3.64E-09** | **4.56E-09** | -1.28 | (-1.33,-1.22) | **5.77E-177** | **9.39E-177** | 0.98 | (0.91,1.05) | **4.77E-72** | **1.09E-71** | 0.07 | (0,0.14) | **3.61E-02** | **4.23E-02** |
| 376 | PC(P-16:0/20:5) | 0.17 | (0.09,0.26) | **9.27E-05** | **1.09E-04** | -3.03 | (-3.1,-2.96) | **2.74E-285** | **1.58E-284** | 2.51 | (2.39,2.63) | **8.65E-111** | **3.96E-110** | 0.63 | (0.5,0.75) | **1.13E-19** | **5.41E-19** |
| 377 | PC(P-17:0/20:4) (a) | -0.16 | (-0.23,-0.1) | **5.36E-07** | **6.52E-07** | -2.00 | (-2.06,-1.94) | **4.59E-223** | **1.10E-222** | 1.68 | (1.58,1.79) | **9.79E-84** | **2.73E-83** | 0.06 | (-0.04,0.17) | 2.40E-01 | 2.59E-01 |
| 378 | PC(P-17:0/20:4) (b) | 0.05 | (-0.01,0.11) | 8.32E-02 | 8.89E-02 | -1.76 | (-1.82,-1.7) | **2.76E-205** | **5.45E-205** | 1.54 | (1.45,1.64) | **7.51E-88** | **2.31E-87** | 0.18 | (0.09,0.27) | **9.83E-05** | **1.43E-04** |
| 379 | PC(P-18:0/20:4) | 0.17 | (0.12,0.21) | **5.66E-14** | **7.45E-14** | -1.11 | (-1.16,-1.06) | **1.55E-165** | **2.41E-165** | 1.26 | (1.18,1.33) | **7.68E-93** | **2.58E-92** | -0.05 | (-0.12,0.01) | 1.13E-01 | 1.27E-01 |
| 380 | PC(P-38:5) (a) | -0.08 | (-0.12,-0.03) | **7.19E-04** | **8.22E-04** | -2.06 | (-2.11,-2.01) | **3.29E-265** | **1.33E-264** | 1.67 | (1.6,1.75) | **2.68E-118** | **1.48E-117** | 0.23 | (0.16,0.3) | **8.00E-11** | **1.99E-10** |
| 381 | PC(P-38:5) (b) | -0.22 | (-0.27,-0.17) | **1.31E-17** | **1.79E-17** | -1.34 | (-1.4,-1.28) | **7.22E-167** | **1.14E-166** | 1.12 | (1.03,1.2) | **1.60E-71** | **3.63E-71** | -0.02 | (-0.09,0.05) | 5.58E-01 | 5.85E-01 |

| | | | | | | | | | | | | | | | | | |
|---|---|---|---|---|---|---|---|---|---|---|---|---|---|---|---|---|---|
| 382 | PC(P-16:0/22:6) | -0.45 | (-0.5,-0.39) | **2.39E-44** | **4.12E-44** | -1.91 | (-1.96,-1.86) | **2.02E-244** | **6.61E-244** | 1.14 | (1.05,1.22) | **7.00E-73** | **1.63E-72** | 0.23 | (0.16,0.31) | **1.20E-08** | **2.46E-08** |
| 383 | PC(P-20:0/20:4) | -0.20 | (-0.24,-0.15) | **7.06E-16** | **9.41E-16** | -1.02 | (-1.07,-0.97) | **5.48E-145** | **7.90E-145** | 0.93 | (0.86,1) | **3.39E-69** | **7.29E-69** | -0.18 | (-0.25,-0.11) | **3.68E-07** | **6.61E-07** |
| 384 | PC(P-40:5) (b) | -0.05 | (-0.09,-0.01) | **1.49E-02** | **1.64E-02** | -1.28 | (-1.33,-1.24) | **1.29E-211** | **2.65E-211** | 1.16 | (1.09,1.22) | **1.16E-97** | **4.37E-97** | 0.01 | (-0.05,0.07) | 7.27E-01 | 7.44E-01 |
| 385 | PC(P-18:0/22:6) | -0.01 | (-0.06,0.04) | 6.95E-01 | 7.08E-01 | -1.81 | (-1.86,-1.76) | **6.46E-246** | **2.12E-245** | 1.69 | (1.6,1.77) | **1.65E-105** | **6.75E-105** | 0.00 | (-0.08,0.08) | 9.42E-01 | 9.42E-01 |
| 386 | PE(16:0_16:1) | -3.13 | (-3.26,-3) | **9.54E-161** | **1.70E-159** | -3.76 | (-3.86,-3.66) | **1.33E-253** | **4.76E-253** | 0.11 | (-0.04,0.27) | 1.40E-01 | 1.44E-01 | 0.45 | (0.31,0.6) | **4.26E-09** | **9.08E-09** |
| 387 | PE(16:0_18:1) | -2.76 | (-2.85,-2.67) | **1.40E-199** | **1.68E-197** | -4.35 | (-4.42,-4.29) | **0.00E+00** | **0.00E+00** | 1.26 | (1.15,1.36) | **5.78E-65** | **1.16E-64** | 0.34 | (0.23,0.45) | **5.35E-09** | **1.12E-08** |
| 388 | PE(16:0_18:2) | -2.97 | (-3.06,-2.87) | **3.88E-201** | **6.21E-199** | -5.30 | (-5.37,-5.23) | **0.00E+00** | **0.00E+00** | 1.89 | (1.79,1.99) | **1.12E-101** | **4.37E-101** | 0.45 | (0.33,0.56) | **3.61E-13** | **1.10E-12** |
| 389 | PE(16:1_18:2) | -2.38 | (-2.48,-2.28) | **1.45E-164** | **3.02E-163** | -3.24 | (-3.32,-3.16) | **3.29E-261** | **1.23E-260** | 0.57 | (0.47,0.67) | **1.57E-24** | **2.11E-24** | 0.32 | (0.21,0.43) | **1.02E-08** | **2.10E-08** |
| 390 | PE(16:0_18:3) (a) | -2.92 | (-3.02,-2.82) | **8.51E-189** | **5.84E-187** | -4.60 | (-4.68,-4.53) | **0.00E+00** | **0.00E+00** | 1.51 | (1.4,1.63) | **1.59E-70** | **3.53E-70** | 0.25 | (0.13,0.37) | **6.25E-05** | **9.29E-05** |
| 391 | PE(16:0_18:3) (b) | -1.96 | (-2.07,-1.86) | **1.57E-126** | **1.18E-125** | -2.97 | (-3.06,-2.89) | **3.13E-244** | **1.02E-243** | 0.88 | (0.75,1) | **2.51E-32** | **3.69E-32** | 0.09 | (-0.05,0.23) | 2.00E-01 | 2.17E-01 |
| 392 | PE(17:0_18:2) | -2.10 | (-2.18,-2.02) | **5.48E-178** | **2.02E-176** | -3.72 | (-3.77,-3.66) | **0.00E+00** | **0.00E+00** | 1.37 | (1.28,1.46) | **7.14E-83** | **1.97E-82** | 0.29 | (0.19,0.39) | **3.01E-08** | **5.94E-08** |
| 393 | PE(18:0_18:1) | -2.12 | (-2.2,-2.04) | **2.10E-175** | **6.73E-174** | -4.60 | (-4.67,-4.54) | **0.00E+00** | **0.00E+00** | 2.23 | (2.13,2.33) | **1.61E-114** | **8.57E-114** | 0.31 | (0.2,0.42) | **4.20E-08** | **8.19E-08** |
| 394 | PE(18:1_18:1) | -1.89 | (-1.97,-1.82) | **7.99E-167** | **1.83E-165** | -3.30 | (-3.38,-3.22) | **5.72E-265** | **2.29E-264** | 1.26 | (1.17,1.35) | **1.79E-76** | **4.51E-76** | 0.16 | (0.07,0.24) | **2.42E-04** | **3.46E-04** |
| 395 | PE(18:0_18:2) | -1.91 | (-1.99,-1.84) | **5.67E-168** | **1.36E-166** | -5.16 | (-5.22,-5.1) | **0.00E+00** | **0.00E+00** | 2.89 | (2.79,2.98) | **1.04E-146** | **1.22E-145** | 0.42 | (0.32,0.53) | **7.56E-14** | **2.45E-13** |
| 396 | PE(18:1_18:2) | -2.53 | (-2.62,-2.44) | **2.74E-183** | **1.46E-181** | -5.12 | (-5.19,-5.06) | **0.00E+00** | **0.00E+00** | 2.37 | (2.27,2.47) | **1.20E-117** | **6.53E-117** | 0.30 | (0.17,0.42) | **2.53E-06** | **4.27E-06** |
| 397 | PE(16:0_20:3) | -2.68 | (-2.77,-2.58) | **7.78E-187** | **4.67E-185** | -3.67 | (-3.74,-3.6) | **6.22E-306** | **6.35E-305** | 0.68 | (0.58,0.79) | **2.07E-30** | **2.98E-30** | 0.30 | (0.19,0.41) | **4.39E-07** | **7.83E-07** |
| 398 | PE(16:0_20:4) | -2.51 | (-2.59,-2.43) | **3.67E-203** | **8.80E-201** | -2.91 | (-2.97,-2.85) | **8.26E-294** | **6.20E-293** | -0.02 | (-0.1,0.07) | 7.07E-01 | 7.08E-01 | 0.41 | (0.31,0.51) | **2.77E-15** | **1.03E-14** |
| 399 | PE(16:1_20:4) | -1.97 | (-2.06,-1.89) | **1.59E-158** | **2.73E-157** | -1.90 | (-1.97,-1.82) | **8.06E-179** | **1.32E-178** | -0.46 | (-0.55,-0.37) | **7.64E-19** | **9.80E-19** | 0.34 | (0.25,0.42) | **3.21E-13** | **9.89E-13** |
| 400 | PE(16:0_20:5) | -1.95 | (-2.04,-1.85) | **4.44E-140** | **4.54E-139** | -3.29 | (-3.36,-3.21) | **1.57E-273** | **7.25E-273** | 0.64 | (0.52,0.75) | **1.29E-22** | **1.71E-22** | 0.73 | (0.59,0.86) | **1.12E-22** | **6.92E-22** |
| 401 | PE(17:0_20:4) | -1.82 | (-1.89,-1.75) | **6.12E-178** | **2.10E-176** | -2.75 | (-2.81,-2.69) | **6.43E-291** | **4.17E-290** | 0.55 | (0.46,0.65) | **3.15E-24** | **4.21E-24** | 0.33 | (0.24,0.43) | **8.73E-11** | **2.15E-10** |
| 402 | PE(18:0_20:3) (a) | -1.87 | (-1.96,-1.79) | **4.02E-152** | **5.52E-151** | -2.50 | (-2.57,-2.43) | **1.49E-243** | **4.71E-243** | 0.25 | (0.16,0.35) | **5.53E-07** | **6.14E-07** | 0.35 | (0.24,0.46) | **3.38E-10** | **7.99E-10** |
| 403 | PE(18:0_20:4) | -1.46 | (-1.52,-1.4) | **4.07E-163** | **8.15E-162** | -2.30 | (-2.36,-2.24) | **7.46E-265** | **2.96E-264** | 0.42 | (0.33,0.5) | **1.71E-19** | **2.21E-19** | 0.47 | (0.38,0.55) | **3.06E-23** | **2.07E-22** |
| 404 | PE(38:5) (a) | -2.34 | (-2.41,-2.26) | **8.54E-194** | **6.83E-192** | -3.31 | (-3.37,-3.25) | **0.00E+00** | **0.00E+00** | 0.55 | (0.45,0.64) | **1.73E-24** | **2.32E-24** | 0.45 | (0.35,0.55) | **1.47E-16** | **6.07E-16** |
| 405 | PE(38:5) (b) | -2.23 | (-2.32,-2.14) | **7.15E-169** | **1.81E-167** | -3.76 | (-3.84,-3.68) | **2.82E-291** | **1.88E-290** | 1.10 | (0.97,1.23) | **8.24E-41** | **1.30E-40** | 0.44 | (0.32,0.57) | **9.51E-12** | **2.52E-11** |
| 406 | PE(16:0_22:6) | -2.64 | (-2.73,-2.55) | **1.49E-195** | **1.43E-193** | -3.80 | (-3.86,-3.73) | **0.00E+00** | **0.00E+00** | 0.31 | (0.2,0.41) | **6.36E-08** | **7.20E-08** | 0.82 | (0.71,0.94) | **1.26E-33** | **3.17E-32** |
| 407 | PE(17:0_22:6) | -1.91 | (-1.98,-1.83) | **4.06E-173** | **1.22E-171** | -2.79 | (-2.86,-2.73) | **1.99E-272** | **8.91E-272** | 0.36 | (0.26,0.46) | **4.75E-11** | **5.62E-11** | 0.59 | (0.48,0.69) | **5.19E-24** | **3.72E-23** |
| 408 | PE(18:0_22:4) | -1.77 | (-1.85,-1.69) | **2.80E-147** | **3.44E-146** | -2.46 | (-2.53,-2.38) | **2.09E-223** | **5.01E-223** | 0.61 | (0.51,0.71) | **4.81E-26** | **6.54E-26** | 0.09 | (0,0.18) | **4.11E-02** | **4.78E-02** |
| 409 | PE(18:0_22:5) (n3) | -1.80 | (-1.87,-1.73) | **5.09E-165** | **1.11E-163** | -3.48 | (-3.55,-3.4) | **9.37E-294** | **6.92E-293** | 1.18 | (1.06,1.29) | **4.13E-53** | **7.45E-53** | 0.57 | (0.47,0.67) | **9.59E-24** | **6.67E-23** |
| 410 | PE(18:0_22:5) (n6) | -1.88 | (-1.97,-1.79) | **3.23E-148** | **4.19E-147** | -3.06 | (-3.14,-2.97) | **5.50E-250** | **1.87E-249** | 0.95 | (0.82,1.08) | **4.18E-33** | **6.21E-33** | 0.23 | (0.12,0.35) | **5.22E-05** | **7.83E-05** |
| 411 | PE(18:0_22:6) | -2.04 | (-2.12,-1.96) | **4.85E-172** | **1.37E-170** | -3.61 | (-3.68,-3.53) | **2.56E-296** | **2.12E-295** | 0.67 | (0.55,0.79) | **2.30E-24** | **3.08E-24** | 0.90 | (0.79,1.02) | **8.68E-39** | **4.63E-37** |
| 412 | PE(40:7) | -3.12 | (-3.22,-3.03) | **7.29E-208** | **3.50E-205** | -4.30 | (-4.38,-4.23) | **0.00E+00** | **0.00E+00** | 0.40 | (0.29,0.52) | **8.48E-11** | **9.98E-11** | 0.79 | (0.66,0.92) | **2.34E-27** | **2.34E-26** |
| 413 | PE(O-34:1) | -0.07 | (-0.13,-0.01) | **1.51E-02** | **1.66E-02** | -3.50 | (-3.58,-3.42) | **5.78E-275** | **2.74E-274** | 3.56 | (3.45,3.67) | **2.26E-151** | **3.19E-150** | -0.13 | (-0.21,-0.04) | **3.08E-03** | **3.97E-03** |
| 414 | PE(O-16:0/18:2) | -0.14 | (-0.21,-0.06) | **5.15E-04** | **5.89E-04** | -5.84 | (-5.97,-5.7) | **1.91E-276** | **9.45E-276** | 5.56 | (5.39,5.73) | **7.94E-150** | **1.09E-148** | 0.14 | (0.03,0.25) | **1.26E-02** | **1.54E-02** |
| 415 | PE(O-18:1/18:2) | 0.10 | (0.03,0.16) | **4.97E-03** | **5.56E-03** | -2.91 | (-2.99,-2.83) | **2.74E-242** | **8.56E-242** | 2.89 | (2.79,3) | **4.28E-134** | **3.95E-133** | 0.06 | (-0.03,0.16) | 1.76E-01 | 1.94E-01 |
| 416 | PE(O-16:0/20:4) | 0.14 | (0.07,0.21) | **1.67E-04** | **1.92E-04** | -3.05 | (-3.13,-2.97) | **1.36E-262** | **5.18E-262** | 2.99 | (2.89,3.1) | **7.52E-137** | **7.36E-136** | 0.10 | (0,0.2) | **4.61E-02** | 5.32E-02 |
| 417 | PE(O-36:5) | -0.25 | (-0.35,-0.14) | **3.88E-06** | **4.65E-06** | -3.35 | (-3.44,-3.25) | **3.52E-237** | **1.00E-236** | 2.47 | (2.35,2.59) | **1.47E-108** | **6.37E-108** | 0.63 | (0.49,0.77) | **8.35E-17** | **3.51E-16** |
| 418 | PE(O-16:0/22:4) | -0.11 | (-0.17,-0.05) | **1.28E-04** | **1.48E-04** | -1.25 | (-1.3,-1.2) | **1.57E-168** | **2.48E-168** | 1.18 | (1.1,1.27) | **2.67E-77** | **6.81E-77** | -0.02 | (-0.1,0.06) | 5.68E-01 | 5.91E-01 |
| 419 | PE(O-18:0/20:4) | 0.22 | (0.14,0.3) | **7.50E-08** | **9.20E-08** | -2.77 | (-2.85,-2.69) | **1.46E-233** | **4.02E-233** | 2.79 | (2.67,2.9) | **1.38E-121** | **8.06E-121** | 0.10 | (-0.02,0.21) | 9.90E-02 | 1.12E-01 |
| 420 | PE(O-38:5) (a) | 0.17 | (0.11,0.24) | **1.15E-06** | **1.39E-06** | -3.66 | (-3.73,-3.59) | **0.00E+00** | **0.00E+00** | 3.64 | (3.54,3.74) | **9.74E-163** | **1.95E-161** | 0.11 | (0,0.21) | **4.46E-02** | 5.18E-02 |
| 421 | PE(O-38:5) (b) | -0.06 | (-0.13,0.01) | 1.12E-01 | 1.19E-01 | -2.26 | (-2.34,-2.18) | **6.76E-208** | **1.36E-207** | 1.99 | (1.88,2.1) | **5.22E-97** | **1.93E-96** | 0.14 | (0.04,0.24) | **4.61E-03** | **5.82E-03** |
| 422 | PE(O-16:0/22:6) | -0.21 | (-0.27,-0.15) | **6.06E-12** | **7.65E-12** | -3.01 | (-3.07,-2.95) | **5.37E-297** | **4.69E-296** | 2.46 | (2.38,2.55) | **2.49E-140** | **2.54E-139** | 0.25 | (0.17,0.33) | **3.18E-09** | **6.80E-09** |
| 423 | PE(O-18:0/22:6) | 0.09 | (0.03,0.15) | **2.53E-03** | **2.85E-03** | -2.31 | (-2.37,-2.25) | **3.69E-257** | **1.34E-256** | 2.20 | (2.12,2.29) | **2.32E-130** | **1.99E-129** | 0.13 | (0.04,0.21) | **3.36E-03** | **4.30E-03** |
| 424 | PE(O-18:1/22:6) | -0.09 | (-0.15,-0.04) | **1.41E-03** | **1.60E-03** | -2.59 | (-2.64,-2.53) | **1.40E-293** | **1.02E-292** | 2.26 | (2.17,2.35) | **6.74E-130** | **5.58E-129** | 0.18 | (0.11,0.26) | **4.89E-06** | **8.06E-06** |
| 425 | PE(P-16:0/18:1) | -0.04 | (-0.09,0) | 7.63E-02 | 8.19E-02 | -1.37 | (-1.42,-1.31) | **4.16E-192** | **7.33E-192** | 1.65 | (1.58,1.72) | **3.05E-122** | **1.81E-121** | -0.37 | (-0.44,-0.31) | **2.82E-23** | **1.93E-22** |
| 426 | PE(P-16:0/18:2) | -0.10 | (-0.17,-0.04) | **1.95E-03** | **2.20E-03** | -4.72 | (-4.79,-4.65) | **0.00E+00** | **0.00E+00** | 4.76 | (4.67,4.86) | **6.49E-196** | **1.04E-193** | -0.24 | (-0.34,-0.15) | **7.07E-07** | **1.23E-06** |
| 427 | PE(P-16:0/20:3) | -0.02 | (-0.09,0.05) | 5.72E-01 | 5.88E-01 | -2.10 | (-2.17,-2.02) | **5.08E-208** | **1.02E-207** | 2.32 | (2.23,2.41) | **1.50E-128** | **1.16E-127** | -0.37 | (-0.45,-0.28) | **8.18E-15** | **2.89E-14** |
| 428 | PE(P-16:0/20:4) | 0.03 | (-0.03,0.09) | 3.59E-01 | 3.71E-01 | -2.86 | (-2.93,-2.79) | **8.17E-277** | **4.13E-276** | 2.92 | (2.82,3.01) | **7.70E-144** | **8.80E-143** | -0.12 | (-0.21,-0.03) | **1.05E-02** | **1.29E-02** |
| 429 | PE(P-16:0/20:5) | -0.02 | (-0.13,0.09) | 6.96E-01 | 7.08E-01 | -6.11 | (-6.25,-5.97) | **1.23E-275** | **5.91E-275** | 5.48 | (5.27,5.69) | **9.08E-130** | **7.26E-129** | 0.48 | (0.32,0.63) | **5.04E-09** | **1.06E-08** |
| 430 | PE(P-16:0/22:4) | -0.22 | (-0.26,-0.17) | **3.17E-16** | **4.26E-16** | -0.04 | (-0.09,0.01) | 1.12E-01 | 1.14E-01 | 0.34 | (0.27,0.42) | **3.21E-18** | **4.10E-18** | -0.59 | (-0.65,-0.52) | **2.95E-47** | **3.54E-45** |
| 431 | PE(P-16:0/22:5) (n3) | -0.05 | (-0.1,0.01) | 7.73E-02 | 8.28E-02 | -2.97 | (-3.02,-2.91) | **0.00E+00** | **0.00E+00** | 2.97 | (2.89,3.06) | **1.50E-159** | **2.87E-158** | -0.09 | (-0.17,-0.01) | **2.13E-02** | **2.55E-02** |
| 432 | PE(P-16:0/22:5) (n6) | -0.35 | (-0.41,-0.29) | **2.75E-27** | **4.15E-27** | -1.29 | (-1.35,-1.23) | **1.50E-168** | **2.38E-168** | 1.33 | (1.23,1.42) | **8.39E-75** | **2.02E-74** | -0.42 | (-0.5,-0.33) | **3.36E-20** | **1.66E-19** |
| 433 | PE(P-16:0/22:6) | -0.41 | (-0.46,-0.36) | **6.84E-47** | **1.22E-46** | -2.25 | (-2.29,-2.2) | **2.24E-287** | **1.33E-286** | 1.63 | (1.55,1.7) | **7.02E-111** | **3.24E-110** | 0.15 | (0.09,0.22) | **1.08E-05** | **1.70E-05** |
| 434 | PE(P-17:0/20:4) (a) | 0.14 | (0.06,0.22) | **1.06E-03** | **1.20E-03** | -2.13 | (-2.2,-2.06) | **9.14E-212** | **1.88E-211** | 2.02 | (1.92,2.12) | **5.44E-104** | **2.20E-103** | 0.18 | (0.07,0.29) | **1.57E-03** | **2.07E-03** |
| 435 | PE(P-17:0/20:4) (b) | 0.16 | (0.09,0.24) | **1.11E-05** | **1.31E-05** | -2.38 | (-2.45,-2.31) | **1.64E-227** | **4.18E-227** | 2.29 | (2.19,2.39) | **2.82E-114** | **1.49E-113** | 0.23 | (0.13,0.33) | **1.44E-05** | **2.22E-05** |
| 436 | PE(P-17:0/22:6) (a) | -0.19 | (-0.26,-0.12) | **2.89E-07** | **3.53E-07** | -1.94 | (-2,-1.87) | **8.50E-205** | **1.67E-204** | 1.46 | (1.35,1.57) | **3.16E-72** | **7.28E-72** | 0.28 | (0.17,0.38) | **5.36E-07** | **9.50E-07** |

| | | | | | | | | | | | | | | | | | |
|---|---|---|---|---|---|---|---|---|---|---|---|---|---|---|---|---|---|
| 437 | PE(P-17:0/22:6) (b) | -0.15 | (-0.22,-0.09) | **6.64E-06** | **7.93E-06** | -2.44 | (-2.5,-2.38) | **9.12E-267** | **3.77E-266** | 1.98 | (1.88,2.08) | **1.82E-106** | **7.61E-106** | 0.30 | (0.21,0.4) | **5.55E-10** | **1.26E-09** |
| 438 | PE(P-18:0/18:1) | 0.00 | (-0.05,0.06) | 9.11E-01 | 9.11E-01 | -2.99 | (-3.05,-2.93) | **6.93E-301** | **6.52E-300** | 3.27 | (3.19,3.34) | **5.12E-183** | **2.73E-181** | -0.35 | (-0.43,-0.27) | **2.44E-16** | **9.84E-16** |
| 439 | PE(P-18:0/18:2) | -0.05 | (-0.11,0.02) | 1.44E-01 | 1.53E-01 | -5.95 | (-6.03,-5.88) | **0.00E+00** | **0.00E+00** | 6.08 | (5.98,6.18) | **8.68E-217** | **4.17E-214** | -0.22 | (-0.31,-0.13) | **3.82E-06** | **6.34E-06** |
| 440 | PE(P-18:0/20:3) | 0.00 | (-0.06,0.06) | 9.05E-01 | 9.06E-01 | -2.32 | (-2.38,-2.26) | **1.55E-262** | **5.85E-262** | 2.55 | (2.47,2.62) | **2.66E-156** | **4.41E-155** | -0.29 | (-0.37,-0.22) | **3.40E-13** | **1.04E-12** |
| 441 | PE(P-18:0/20:4) | 0.33 | (0.27,0.39) | **1.13E-23** | **1.63E-23** | -3.00 | (-3.07,-2.93) | **3.70E-278** | **1.91E-277** | 3.28 | (3.18,3.37) | **7.52E-157** | **1.29E-155** | -0.05 | (-0.14,0.04) | 3.17E-01 | 3.40E-01 |
| 442 | PE(P-18:0/20:5) | 0.15 | (0.03,0.27) | **1.11E-02** | **1.23E-02** | -6.17 | (-6.3,-6.03) | **8.16E-284** | **4.61E-283** | 5.80 | (5.6,6) | **8.73E-141** | **9.11E-140** | 0.49 | (0.34,0.65) | **1.42E-09** | **3.12E-09** |
| 443 | PE(P-18:0/22:4) | -0.06 | (-0.11,0) | **4.72E-02** | 5.13E-02 | -0.16 | (-0.22,-0.1) | **6.56E-08** | **6.91E-08** | 0.84 | (0.76,0.91) | **2.28E-57** | **4.25E-57** | -0.79 | (-0.86,-0.72) | **8.23E-62** | **3.95E-59** |
| 444 | PE(P-18:0/22:5) (n3) | 0.23 | (0.18,0.29) | **5.18E-16** | **6.93E-16** | -2.45 | (-2.5,-2.39) | **9.29E-297** | **7.96E-296** | 2.79 | (2.71,2.86) | **3.12E-168** | **8.82E-167** | -0.14 | (-0.21,-0.06) | **3.85E-04** | **5.40E-04** |
| 445 | PE(P-18:0/22:5) (n6) | -0.11 | (-0.17,-0.06) | **1.27E-04** | **1.48E-04** | -1.47 | (-1.53,-1.41) | **2.86E-190** | **4.99E-190** | 1.72 | (1.63,1.82) | **4.01E-96** | **1.42E-95** | -0.39 | (-0.48,-0.3) | **1.86E-16** | **7.55E-16** |
| 446 | PE(P-18:0/22:6) | 0.03 | (-0.02,0.08) | 2.44E-01 | 2.55E-01 | -2.55 | (-2.6,-2.5) | **6.87E-296** | **5.59E-295** | 2.51 | (2.43,2.6) | **9.32E-143** | **1.02E-141** | -0.01 | (-0.09,0.06) | 7.13E-01 | 7.32E-01 |
| 447 | PE(P-18:1/18:1) | -0.13 | (-0.18,-0.08) | **5.20E-07** | **6.33E-07** | -2.42 | (-2.47,-2.37) | **4.07E-290** | **2.61E-289** | 2.65 | (2.58,2.72) | **8.11E-166** | **1.95E-164** | -0.40 | (-0.47,-0.32) | **7.29E-23** | **4.60E-22** |
| 448 | PE(P-18:1/18:2) | -0.17 | (-0.23,-0.1) | **7.14E-07** | **8.65E-07** | -5.96 | (-6.05,-5.87) | **0.00E+00** | **0.00E+00** | 5.97 | (5.84,6.09) | **5.17E-189** | **4.13E-187** | -0.25 | (-0.35,-0.15) | **1.04E-06** | **1.80E-06** |
| 449 | PE(P-18:1/20:4) | 0.16 | (0.09,0.22) | **1.17E-06** | **1.41E-06** | -3.19 | (-3.26,-3.12) | **2.97E-289** | **1.83E-288** | 3.41 | (3.32,3.51) | **5.02E-158** | **9.27E-157** | -0.14 | (-0.23,-0.05) | **3.38E-03** | **4.31E-03** |
| 450 | PE(P-18:1/20:5) | -0.25 | (-0.36,-0.14) | **1.19E-05** | **1.41E-05** | -6.96 | (-7.13,-6.79) | **2.99E-268** | **1.26E-267** | 6.18 | (5.94,6.43) | **5.22E-128** | **3.74E-127** | 0.43 | (0.27,0.59) | **2.16E-07** | **3.95E-07** |
| 451 | PE(P-18:1/22:4) | -0.18 | (-0.24,-0.12) | **1.14E-08** | **1.41E-08** | -0.23 | (-0.3,-0.17) | **6.11E-12** | **6.48E-12** | 0.70 | (0.62,0.78) | **1.94E-44** | **3.17E-44** | -0.72 | (-0.8,-0.65) | **5.75E-53** | **9.19E-51** |
| 452 | PE(P-18:1/22:5) (a) | 0.08 | (0.02,0.13) | **5.28E-03** | **5.90E-03** | -2.50 | (-2.56,-2.45) | **9.15E-292** | **6.46E-291** | 2.65 | (2.57,2.73) | **2.58E-152** | **3.86E-151** | -0.07 | (-0.15,0.01) | 8.30E-02 | 9.42E-02 |
| 453 | PE(P-18:1/22:6) (a) | -0.34 | (-0.4,-0.29) | **5.67E-30** | **8.75E-30** | -3.02 | (-3.07,-2.96) | **0.00E+00** | **0.00E+00** | 2.55 | (2.46,2.64) | **2.08E-138** | **2.08E-137** | 0.06 | (-0.02,0.14) | 1.24E-01 | 1.38E-01 |
| 454 | PE(P-20:0/18:1) | -0.10 | (-0.15,-0.04) | **1.17E-03** | **1.33E-03** | -3.52 | (-3.63,-3.4) | **6.92E-212** | **1.43E-211** | 4.04 | (3.89,4.2) | **2.25E-128** | **1.69E-127** | -0.65 | (-0.73,-0.57) | **6.49E-40** | **4.45E-38** |
| 455 | PE(P-20:0/18:2) | -0.03 | (-0.09,0.02) | 1.89E-01 | 1.99E-01 | -3.32 | (-3.37,-3.27) | **0.00E+00** | **0.00E+00** | 3.81 | (3.75,3.88) | **1.39E-210** | **3.35E-208** | -0.56 | (-0.64,-0.49) | **1.20E-34** | **3.40E-33** |
| 456 | PE(P-20:0/20:4) | 0.42 | (0.37,0.47) | **2.15E-47** | **3.85E-47** | -1.58 | (-1.65,-1.51) | **4.76E-174** | **7.67E-174** | 2.41 | (2.32,2.49) | **4.61E-135** | **4.34E-134** | -0.48 | (-0.55,-0.41) | **1.58E-30** | **2.30E-29** |
| 457 | PE(P-20:0/22:6) | -0.11 | (-0.15,-0.06) | **2.84E-06** | **3.42E-06** | -1.92 | (-1.97,-1.88) | **1.89E-273** | **8.64E-273** | 1.96 | (1.89,2.04) | **1.40E-129** | **1.10E-128** | -0.22 | (-0.29,-0.16) | **1.93E-10** | **4.64E-10** |
| 458 | PE(P-20:1/20:4) | 0.36 | (0.27,0.44) | **2.73E-15** | **3.63E-15** | -2.81 | (-2.9,-2.72) | **1.74E-218** | **3.96E-218** | 3.02 | (2.89,3.15) | **2.77E-119** | **1.58E-118** | 0.08 | (-0.04,0.2) | 1.87E-01 | 2.05E-01 |
| 459 | PG(34:2) | -1.24 | (-1.33,-1.15) | **9.59E-91** | **3.44E-90** | -2.54 | (-2.64,-2.44) | **1.70E-188** | **2.96E-188** | 0.95 | (0.79,1.11) | **7.98E-26** | **1.08E-25** | 0.24 | (0.13,0.36) | **5.65E-05** | **8.45E-05** |
| 460 | PG(36:1) | -0.89 | (-0.98,-0.81) | **3.85E-69** | **9.07E-69** | -2.47 | (-2.56,-2.38) | **4.18E-193** | **7.48E-193** | 1.35 | (1.2,1.5) | **3.14E-45** | **5.17E-45** | 0.36 | (0.24,0.48) | **6.49E-09** | **1.35E-08** |
| 461 | PI(16:0/16:0) | -3.06 | (-3.17,-2.95) | **2.26E-178** | **9.04E-177** | -4.54 | (-4.63,-4.45) | **1.75E-302** | **1.72E-301** | 1.18 | (1.06,1.3) | **1.35E-51** | **2.39E-51** | 0.27 | (0.15,0.39) | **1.33E-05** | **2.07E-05** |
| 462 | PI(16:0_16:1) | -1.93 | (-2.05,-1.82) | **1.48E-110** | **8.35E-110** | -2.60 | (-2.69,-2.51) | **3.80E-204** | **7.35E-204** | 0.54 | (0.43,0.66) | **5.15E-17** | **6.44E-17** | 0.12 | (-0.01,0.25) | 6.18E-02 | 7.04E-02 |
| 463 | PI(34:0) | -3.14 | (-3.25,-3.02) | **8.72E-183** | **4.19E-181** | -5.20 | (-5.31,-5.09) | **3.49E-294** | **2.66E-293** | 1.64 | (1.48,1.8) | **1.21E-54** | **2.20E-54** | 0.46 | (0.33,0.59) | **8.51E-12** | **2.28E-11** |
| 464 | PI(34:1) | -1.65 | (-1.73,-1.58) | **4.07E-154** | **5.75E-153** | -4.29 | (-4.35,-4.23) | **0.00E+00** | **0.00E+00** | 2.58 | (2.5,2.66) | **2.55E-148** | **3.23E-147** | 0.08 | (0,0.17) | 5.67E-02 | 6.50E-02 |
| 465 | PI(17:0_18:1) | -1.46 | (-1.53,-1.39) | **4.85E-144** | **5.54E-143** | -4.11 | (-4.18,-4.03) | **0.00E+00** | **0.00E+00** | 2.63 | (2.52,2.74) | **4.63E-123** | **2.92E-122** | 0.10 | (0.01,0.19) | **2.31E-02** | **2.75E-02** |
| 466 | PI(17:0_18:2) | -1.45 | (-1.52,-1.38) | **3.58E-143** | **4.00E-142** | -4.27 | (-4.36,-4.19) | **2.73E-300** | **2.52E-299** | 2.65 | (2.51,2.79) | **3.96E-101** | **1.53E-100** | 0.23 | (0.14,0.33) | **1.18E-06** | **2.04E-06** |
| 467 | PI(18:0_18:1) | -1.70 | (-1.77,-1.62) | **3.29E-157** | **5.26E-156** | -5.98 | (-6.04,-5.91) | **0.00E+00** | **0.00E+00** | 4.17 | (4.08,4.26) | **3.36E-187** | **2.02E-185** | 0.16 | (0.07,0.25) | **9.65E-04** | **1.30E-03** |
| 468 | PI(36:2) | -1.43 | (-1.48,-1.37) | **3.39E-171** | **9.03E-170** | -5.15 | (-5.21,-5.1) | **0.00E+00** | **0.00E+00** | 3.61 | (3.54,3.68) | **2.26E-193** | **2.72E-191** | 0.13 | (0.06,0.2) | **1.70E-04** | **2.45E-04** |
| 469 | PI(18:1_18:2) | -1.18 | (-1.25,-1.12) | **3.68E-127** | **2.87E-126** | -2.96 | (-3.01,-2.9) | **0.00E+00** | **0.00E+00** | 1.74 | (1.66,1.82) | **1.37E-111** | **6.52E-111** | 0.03 | (-0.06,0.12) | 4.74E-01 | 5.01E-01 |
| 470 | PI(16:0_20:3) (a) | -1.43 | (-1.5,-1.36) | **1.50E-133** | **1.31E-132** | -1.96 | (-2.01,-1.9) | **2.67E-242** | **8.39E-242** | 0.70 | (0.63,0.78) | **1.61E-49** | **2.80E-49** | -0.19 | (-0.27,-0.11) | **8.10E-06** | **1.30E-05** |
| 471 | PI(16:0_20:3) (b) | -1.09 | (-1.17,-1.01) | **7.08E-89** | **2.31E-88** | -0.75 | (-0.84,-0.66) | **4.59E-50** | **5.27E-50** | -0.09 | (-0.19,0.01) | 7.36E-02 | 7.60E-02 | -0.18 | (-0.27,-0.09) | **6.33E-05** | **9.37E-05** |
| 472 | PI(16:0_20:4) | -1.06 | (-1.12,-0.99) | **2.63E-110** | **1.47E-109** | -1.09 | (-1.15,-1.04) | **5.57E-152** | **8.27E-152** | 0.03 | (-0.05,0.1) | 5.05E-01 | 5.09E-01 | -0.02 | (-0.09,0.06) | 6.35E-01 | 6.54E-01 |
| 473 | PI(18:0_20:2) | -1.67 | (-1.74,-1.61) | **1.81E-161** | **3.35E-160** | -3.33 | (-3.42,-3.25) | **1.13E-261** | **4.23E-261** | 1.53 | (1.42,1.63) | **3.18E-79** | **8.26E-79** | 0.14 | (0.05,0.22) | **1.44E-03** | **1.90E-03** |
| 474 | PI(18:0_20:3) (a) | -0.52 | (-0.56,-0.47) | **3.13E-75** | **8.17E-75** | -1.46 | (-1.5,-1.42) | **7.88E-228** | **2.03E-227** | 0.79 | (0.73,0.85) | **1.81E-69** | **3.91E-69** | 0.13 | (0.07,0.19) | **4.46E-05** | **6.71E-05** |
| 475 | PI(18:0_20:4) | -0.48 | (-0.53,-0.44) | **1.32E-69** | **3.12E-69** | -1.41 | (-1.46,-1.37) | **5.39E-223** | **1.28E-222** | 0.77 | (0.7,0.83) | **4.73E-64** | **9.39E-64** | 0.15 | (0.09,0.21) | **1.86E-06** | **3.16E-06** |
| 476 | PI(38:5) (a) | -0.86 | (-0.92,-0.8) | **2.64E-99** | **1.11E-98** | -0.66 | (-0.72,-0.61) | **1.44E-77** | **1.70E-77** | -0.22 | (-0.3,-0.14) | **2.67E-07** | **3.00E-07** | 0.02 | (-0.05,0.09) | 5.65E-01 | 5.90E-01 |
| 477 | PI(38:6) | -1.36 | (-1.42,-1.29) | **4.66E-146** | **5.59E-145** | -1.79 | (-1.84,-1.73) | **5.06E-235** | **1.41E-234** | 0.47 | (0.39,0.54) | **7.90E-29** | **1.12E-28** | -0.10 | (-0.18,-0.02) | **1.04E-02** | **1.29E-02** |
| 478 | PI(18:0_22:4) | -1.24 | (-1.3,-1.18) | **1.08E-133** | **9.62E-133** | -1.83 | (-1.89,-1.77) | **6.13E-214** | **1.33E-213** | 0.59 | (0.5,0.67) | **1.12E-31** | **1.63E-31** | -0.02 | (-0.09,0.05) | 5.92E-01 | 6.15E-01 |
| 479 | PI(18:0_22:5) (n3) | -0.49 | (-0.55,-0.43) | **6.84E-42** | **1.15E-41** | -2.63 | (-2.69,-2.57) | **1.69E-283** | **9.45E-283** | 1.86 | (1.77,1.95) | **3.94E-111** | **1.84E-110** | 0.26 | (0.18,0.34) | **4.18E-10** | **9.64E-10** |
| 480 | PI(18:0_22:6) | -1.31 | (-1.37,-1.25) | **1.88E-147** | **2.37E-146** | -2.50 | (-2.55,-2.45) | **0.00E+00** | **0.00E+00** | 1.18 | (1.1,1.25) | **1.45E-82** | **3.98E-82** | 0.02 | (-0.05,0.1) | 5.51E-01 | 5.79E-01 |

**Supplementary Table 3a.** Association of lipid classes with BMI and BW in four sample groups (antenatal, postnatal, cord blood and 6-year-old child) in terms of levels. Related to Figure 2.

| | **ppBMI (antenatal)** | | | | **Adult BMI (postnatal)** | | | | **Birth Weight (cord blood)** | | | | **Child BMI (six-year-old)** | | | |
|---|---|---|---|---|---|---|---|---|---|---|---|---|---|---|---|---|
| | adjusted for ethnicity, maternal age, maternal education level and gestational weight gain at 26-28 weeks of gestation | | | | adjusted for ethnicity, maternal age and maternal education level | | | | adjusted for sex, ethnicity, maternal age, maternal education level, pre-pregnancy BMI, total gestational weight gain, gestational age and parity | | | | adjusted for sex, ethnicity and maternal education level | | | |
| Lipid Class | %change in lipid concentration per unit BMI | 95%CI | p-value | p-value(BH) | %change in lipid concentration per unit BMI | 95%CI | p-value | p-value(BH) | %change in lipid concentration per 100 grams | 95%CI | p-value | p-value(BH) | %change in lipid concentration per unit BMI | 95%CI | p-value | p-value(BH) |
| AC | 0.81 | (0.18-1.44) | **1.13E-02** | **1.76E-02** | -0.24 | (-0.72-0.25) | 3.38E-01 | 3.84E-01 | -0.62 | (-1.14--0.06) | **3.00E-02** | 6.26E-02 | -1.00 | (-2.51-0.54) | 2.00E-01 | 3.84E-01 |
| CE | -0.93 | (-1.26--0.6) | **4.94E-08** | **2.29E-07** | -0.43 | (-0.71--0.14) | **3.26E-03** | **4.80E-03** | -0.30 | (-0.74-0.17) | 2.06E-01 | 2.86E-01 | 0.54 | (-0.38-1.47) | 2.50E-01 | 4.47E-01 |
| DG | 1.12 | (0.48-1.76) | **5.79E-04** | **1.45E-03** | 3.05 | (2.26-3.85) | **8.01E-14** | **1.00E-12** | -2.21 | (-2.77--1.61) | **9.98E-11** | **4.99E-10** | 2.83 | (0.69-5.03) | **9.66E-03** | 6.04E-02 |
| TG | 0.59 | (0.15-1.04) | **8.50E-03** | **1.63E-02** | 2.95 | (2.24-3.66) | **8.03E-16** | **2.01E-14** | -3.26 | (-3.92--2.53) | **2.12E-13** | **1.76E-12** | 3.62 | (1.61-5.67) | **4.09E-04** | **1.02E-02** |
| TG(O) | 0.44 | (-0.14-1.02) | 1.35E-01 | 1.53E-01 | 2.88 | (2.09-3.67) | **1.48E-12** | **1.24E-11** | -2.74 | (-3.37--2.05) | **9.36E-12** | **5.85E-11** | 3.05 | (0.82-5.32) | **7.22E-03** | 6.01E-02 |
| dhCer | 0.99 | ( 0.41- 1.57) | **7.81E-04** | **1.63E-03** | 1.42 | ( 0.85- 1.99) | **9.62E-07** | **2.67E-06** | -0.32 | (-0.95- 0.35) | 3.35E-01 | 4.22E-01 | 0.80 | (-0.77- 2.39) | 3.18E-01 | 5.29E-01 |
| Cer | -0.56 | (-0.99--0.13) | **1.10E-02** | **1.76E-02** | 0.22 | (-0.22- 0.66) | 3.19E-01 | 3.80E-01 | -0.41 | (-0.95- 0.16) | 1.54E-01 | 2.57E-01 | 0.41 | (-0.83- 1.68) | 5.16E-01 | 6.79E-01 |
| DeoxyCer | 0.31 | (-0.43-1.05) | 4.14E-01 | 4.50E-01 | 2.50 | (1.79-3.21) | **5.39E-12** | **3.37E-11** | -2.12 | (-2.66--1.54) | **1.22E-10** | **5.10E-10** | 1.65 | (-0.04-3.36) | 5.57E-02 | 1.76E-01 |
| GM3 | -1.98 | (-2.5--1.47) | **1.59E-13** | **1.98E-12** | -0.89 | (-1.36--0.42) | **2.46E-04** | **4.38E-04** | -0.34 | (-1.01-0.37) | 3.37E-01 | 4.22E-01 | -1.46 | (-2.71--0.19) | **2.41E-02** | 1.20E-01 |
| HexCer | -1.86 | (-2.32--1.4) | **1.61E-14** | **4.03E-13** | -0.90 | (-1.35--0.45) | **9.66E-05** | **2.01E-04** | 0.19 | (-0.53-0.97) | 6.16E-01 | 7.33E-01 | -1.81 | (-3.09--0.5) | **6.98E-03** | 6.01E-02 |
| Hex2Cer | -1.53 | (-2.04--1.02) | **6.86E-09** | **4.29E-08** | -1.16 | (-1.67--0.64) | **1.49E-05** | **3.72E-05** | -0.46 | (-1.12-0.25) | 1.95E-01 | 2.86E-01 | -1.46 | (-3.09-0.21) | 8.61E-02 | 2.39E-01 |
| Hex3Cer | -1.17 | (-1.58--0.75) | **5.51E-08** | **2.29E-07** | -1.34 | (-1.74--0.94) | **1.52E-10** | **7.60E-10** | -0.44 | (-1.02-0.19) | 1.65E-01 | 2.57E-01 | -1.20 | (-2.32--0.07) | **3.69E-02** | 1.54E-01 |
| SM | -1.04 | (-1.49--0.58) | **1.18E-05** | **3.70E-05** | -0.32 | (-0.75-0.13) | 1.60E-01 | 2.00E-01 | -0.10 | (-0.63-0.46) | 7.17E-01 | 7.79E-01 | -0.58 | (-1.78-0.65) | 3.53E-01 | 5.51E-01 |
| LPC | 0.07 | (-0.4-0.55) | 7.62E-01 | 7.62E-01 | -0.91 | (-1.38--0.45) | **1.28E-04** | **2.45E-04** | 2.25 | (1.73-2.79) | **8.80E-19** | **1.10E-17** | 0.24 | (-1.48-1.98) | 7.88E-01 | 8.56E-01 |
| LPC(O) | -0.87 | (-1.26--0.47) | **1.96E-05** | **5.44E-05** | -1.15 | (-1.56--0.74) | **6.12E-08** | **2.55E-07** | 1.31 | (0.86-1.77) | **2.73E-09** | **8.53E-09** | -0.45 | (-2.04-1.17) | 5.83E-01 | 6.94E-01 |
| LPC(P) | 0.40 | (-0.04-0.83) | 7.32E-02 | 9.64E-02 | -0.92 | (-1.36--0.48) | **4.97E-05** | **1.13E-04** | 1.47 | (1-1.96) | **1.85E-10** | **6.62E-10** | -0.14 | (-1.84-1.59) | 8.71E-01 | 8.77E-01 |
| LPE | -0.37 | (-0.81-0.07) | 1.01E-01 | 1.20E-01 | -0.79 | (-1.28--0.31) | **1.45E-03** | **2.26E-03** | 2.55 | (1.99-3.13) | **4.07E-21** | **1.02E-19** | -1.18 | (-2.61-0.27) | 1.09E-01 | 2.74E-01 |
| PC | -0.62 | (-0.98--0.26) | **7.21E-04** | **1.63E-03** | -0.08 | (-0.49-0.33) | 7.04E-01 | 7.33E-01 | -0.53 | (-1.03-0) | 5.04E-02 | 9.01E-02 | -0.34 | (-1.52-0.85) | 5.70E-01 | 6.94E-01 |
| PC(O) | -0.61 | (-1.09--0.14) | **1.13E-02** | **1.76E-02** | -0.71 | (-1.18--0.23) | **3.52E-03** | **4.88E-03** | -0.11 | (-0.68-0.49) | 7.15E-01 | 7.79E-01 | -0.23 | (-1.57-1.12) | 7.36E-01 | 8.36E-01 |
| PC(P) | -0.49 | (-1.03-0.05) | 7.27E-02 | 9.64E-02 | -1.25 | (-1.74--0.76) | **6.73E-07** | **2.10E-06** | -0.92 | (-1.45--0.35) | **1.83E-03** | **5.07E-03** | -0.61 | (-2.02-0.81) | 3.96E-01 | 5.82E-01 |
| PE | 0.15 | (-0.5-0.8) | 6.49E-01 | 6.76E-01 | 0.30 | (-0.51-1.12) | 4.64E-01 | 5.04E-01 | -0.91 | (-1.47--0.31) | **3.63E-03** | **9.09E-03** | 0.18 | (-2.07-2.48) | 8.77E-01 | 8.77E-01 |
| PE(O) | -1.47 | (-2.09--0.85) | **4.80E-06** | **1.72E-05** | -1.22 | (-1.87--0.57) | **2.76E-04** | **4.60E-04** | -0.06 | (-0.65-0.58) | 8.60E-01 | 8.96E-01 | 0.69 | (-1.14-2.56) | 4.60E-01 | 6.40E-01 |
| PE(P) | -0.54 | (-1.11-0.05) | 7.07E-02 | 9.64E-02 | -0.54 | (-1.11-0.03) | 6.24E-02 | 8.21E-02 | -0.04 | (-0.63-0.59) | 9.00E-01 | 9.00E-01 | 1.24 | (-0.35-2.85) | 1.28E-01 | 2.90E-01 |
| PG | -0.51 | (-1.09-0.07) | 8.44E-02 | 1.06E-01 | 2.08 | (1.27-2.89) | **4.82E-07** | **1.72E-06** | -1.00 | (-1.87--0.04) | **4.14E-02** | 7.96E-02 | 2.23 | (-0.06-4.58) | 5.64E-02 | 1.76E-01 |
| PI | -1.59 | (-2.07--1.1) | **2.95E-10** | **2.46E-09** | -0.07 | (-0.56-0.42) | 7.73E-01 | 7.73E-01 | -0.47 | (-0.87--0.05) | **2.97E-02** | 6.26E-02 | -0.85 | (-2.1-0.41) | 1.85E-01 | 3.84E-01 |

**Supplementary Table 3b.** Association of lipid species with BMI and BW in four sample groups (antenatal, postnatal, cord blood and 6y child) in terms of levels. Related to Figure 2.

| Studies | | **ppBMI (antenatal)** | | | | **Adult BMI (postnatal)** | | | | **Birth Weight (cord blood)** | | | | **Child BMI (six-year-old)** | | | |
|---|---|---|---|---|---|---|---|---|---|---|---|---|---|---|---|---|---|
| Covariates | | adjusted for ethnicity, maternal age, maternal education level and gestational weight gain at 26-28 weeks of gestation | | | | adjusted for ethnicity, maternal age and maternal education level | | | | adjusted for sex, ethnicity, maternal age, maternal education level, pre-pregnancy BMI, total gestational weight gain, gestational age and parity | | | | adjusted for sex, ethnicity and maternal education level | | | |
| No. | Lipid Species | %change in lipid concentration per unit BMI | 95%CI | p-value | p-value(BH) | %change in lipid concentration per unit BMI | 95%CI | p-value | p-value(BH) | %change in lipid concentration per 100 grams | 95%CI | p-value | p-value(BH) | %change in lipid concentration per unit BMI | 95%CI | p-value | p-value(BH) |
| 1 | AC(12:0) | 0.31 | (-0.75-1.38) | 5.65E-01 | 6.18E-01 | -0.66 | (-1.46-0.15) | 1.10E-01 | 1.47E-01 | -1.55 | (-2.14--0.93) | **4.02E-06** | **1.32E-05** | -1.49 | (-3.93-1.02) | 2.42E-01 | 3.94E-01 |
| 2 | AC(13:0) | 1.33 | (-0.01-2.68) | 5.12E-02 | 7.95E-02 | -0.35 | (-1.59-0.91) | 5.84E-01 | 6.40E-01 | -1.16 | (-2.22-0.03) | 5.65E-02 | 9.86E-02 | 0.87 | (-2.62-4.49) | 6.28E-01 | 7.41E-01 |
| 3 | AC(14:0) | 0.33 | (-0.37-1.04) | 3.52E-01 | 4.17E-01 | 0.01 | (-0.56-0.59) | 9.69E-01 | 9.73E-01 | -1.03 | (-1.59--0.43) | **1.07E-03** | **2.58E-03** | -1.31 | (-3.15-0.58) | 1.72E-01 | 3.06E-01 |
| 4 | AC(14:1) | 1.38 | (0.22-2.56) | **1.98E-02** | **3.50E-02** | -0.11 | (-0.96-0.75) | 8.03E-01 | 8.39E-01 | -1.08 | (-1.73--0.37) | **3.63E-03** | **8.13E-03** | -2.04 | (-4.66-0.65) | 1.36E-01 | 2.51E-01 |
| 5 | AC(14:2) | 1.03 | (-0.08-2.15) | 6.79E-02 | 1.00E-01 | -0.44 | (-1.25-0.37) | 2.84E-01 | 3.39E-01 | -0.94 | (-1.49--0.36) | **1.99E-03** | **4.67E-03** | -1.62 | (-4.07-0.9) | 2.05E-01 | 3.43E-01 |
| 6 | AC(16:0) | 0.33 | (-0.19-0.85) | 2.12E-01 | 2.71E-01 | 0.00 | (-0.42-0.42) | 9.90E-01 | 9.92E-01 | -0.47 | (-1-0.09) | 9.81E-02 | 1.58E-01 | -0.44 | (-1.62-0.76) | 4.71E-01 | 6.14E-01 |
| 7 | AC(16:1) | 1.19 | (0.37-2.02) | **4.39E-03** | **9.66E-03** | 0.35 | (-0.27-0.98) | 2.67E-01 | 3.22E-01 | -1.00 | (-1.64--0.31) | **5.17E-03** | **1.12E-02** | -0.23 | (-2.24-1.82) | 8.25E-01 | 8.94E-01 |
| 8 | AC(18:0) | -0.08 | (-0.6-0.45) | 7.74E-01 | 8.06E-01 | -0.68 | (-1.13--0.23) | **3.16E-03** | **6.00E-03** | -0.78 | (-1.3--0.24) | **5.68E-03** | **1.22E-02** | -1.98 | (-3.33--0.62) | **4.60E-03** | **1.97E-02** |
| 9 | AC(18:1) | 1.36 | (0.66-2.06) | **1.34E-04** | **4.24E-04** | -0.12 | (-0.62-0.37) | 6.29E-01 | 6.83E-01 | 0.10 | (-0.58-0.83) | 7.86E-01 | 8.50E-01 | -0.64 | (-2.04-0.79) | 3.80E-01 | 5.29E-01 |
| 10 | AC(18:2) | 0.99 | (0.31-1.67) | **4.42E-03** | **9.69E-03** | -0.41 | (-0.89-0.08) | 9.84E-02 | 1.33E-01 | -0.43 | (-1.04-0.23) | 1.98E-01 | 2.86E-01 | -0.30 | (-1.64-1.06) | 6.67E-01 | 7.58E-01 |
| 11 | CE(14:0) | -1.93 | (-2.48--1.38) | **2.08E-11** | **3.99E-10** | 0.21 | (-0.36-0.8) | 4.69E-01 | 5.28E-01 | -0.50 | (-1.02-0.05) | 7.17E-02 | 1.22E-01 | 2.26 | (0.55-3.99) | **9.50E-03** | **3.47E-02** |
| 12 | CE(16:1) | -1.06 | (-1.82--0.29) | **7.23E-03** | **1.46E-02** | 2.17 | (1.53-2.82) | **6.89E-11** | **4.60E-10** | 0.98 | (0.31-1.69) | **3.55E-03** | **8.00E-03** | 5.77 | (4.04-7.53) | **9.25E-11** | **2.22E-08** |
| 13 | CE(18:0) | -1.13 | (-1.64--0.61) | **2.13E-05** | **8.04E-05** | -0.05 | (-0.57-0.47) | 8.47E-01 | 8.74E-01 | 0.27 | (-0.5-1.1) | 5.02E-01 | 5.92E-01 | 0.82 | (-0.9-2.56) | 3.51E-01 | 5.06E-01 |
| 14 | CE(18:1) | -0.92 | (-1.23--0.62) | **7.33E-09** | **5.29E-08** | 0.06 | (-0.22-0.34) | 6.75E-01 | 7.21E-01 | 0.15 | (-0.28-0.6) | 4.96E-01 | 5.86E-01 | 0.45 | (-0.36-1.27) | 2.78E-01 | 4.39E-01 |
| 15 | CE(18:2) | -0.83 | (-1.18--0.48) | **4.08E-06** | **1.78E-05** | -0.32 | (-0.64-0) | **4.82E-02** | 7.03E-02 | -0.41 | (-0.9-0.12) | 1.27E-01 | 1.98E-01 | 0.36 | (-0.64-1.38) | 4.80E-01 | 6.23E-01 |
| 16 | CE(18:3) | -1.93 | (-2.54--1.32) | **1.28E-09** | **1.25E-08** | 0.72 | (0.11-1.32) | **2.04E-02** | **3.21E-02** | 0.31 | (-0.55-1.25) | 4.89E-01 | 5.80E-01 | 2.78 | (0.7-4.91) | **8.76E-03** | **3.24E-02** |
| 17 | CE(20:1) | -0.83 | (-1.29--0.37) | **4.07E-04** | **1.15E-03** | -0.70 | (-1.17--0.22) | **4.06E-03** | **7.58E-03** | 0.56 | (-0.26-1.44) | 1.87E-01 | 2.71E-01 | -1.24 | (-2.51-0.06) | 6.17E-02 | 1.45E-01 |
| 18 | CE(20:2) | -1.33 | (-1.74--0.91) | **7.59E-10** | **7.92E-09** | 0.10 | (-0.26-0.46) | 5.84E-01 | 6.40E-01 | 0.79 | (0.16-1.46) | **1.36E-02** | **2.71E-02** | 0.46 | (-0.64-1.56) | 4.14E-01 | 5.58E-01 |
| 19 | CE(20:4) | -0.66 | (-1.3--0.02) | **4.41E-02** | 7.05E-02 | -1.54 | (-2.06--1.02) | **1.28E-08** | **5.79E-08** | -0.17 | (-0.73-0.43) | 5.78E-01 | 6.63E-01 | 0.49 | (-1.02-2.02) | 5.25E-01 | 6.51E-01 |
| 20 | CE(20:5) | -1.71 | (-3.04--0.36) | **1.31E-02** | **2.41E-02** | -0.98 | (-2.2-0.25) | 1.18E-01 | 1.56E-01 | -2.42 | (-3.39--1.3) | **9.07E-05** | **2.59E-04** | 3.76 | (0.43-7.2) | **2.68E-02** | 7.82E-02 |
| 21 | CE(22:5) (n6) | -1.88 | (-2.46--1.29) | **7.49E-10** | **7.92E-09** | -0.68 | (-1.1--0.26) | **1.77E-03** | **3.56E-03** | 0.43 | (-0.33-1.25) | 2.74E-01 | 3.66E-01 | 1.10 | (-0.31-2.52) | 1.25E-01 | 2.34E-01 |
| 22 | CE(22:6) | -1.55 | (-2.27--0.82) | **3.29E-05** | **1.18E-04** | -0.52 | (-1.23-0.2) | 1.56E-01 | 2.01E-01 | -1.97 | (-2.73--1.12) | **1.95E-05** | **6.12E-05** | 2.24 | (0.07-4.46) | **4.26E-02** | 1.10E-01 |
| 23 | CE(24:5) | -2.01 | (-2.72--1.3) | **4.73E-08** | **3.03E-07** | -0.34 | (-1-0.33) | 3.22E-01 | 3.80E-01 | -2.88 | (-3.68--1.98) | **3.40E-08** | **1.41E-07** | -1.19 | (-3.09-0.75) | 2.27E-01 | 3.72E-01 |
| 24 | CE(24:6) | -2.55 | (-3.28--1.82) | **2.50E-11** | **4.44E-10** | -0.16 | (-0.84-0.53) | 6.43E-01 | 6.92E-01 | -1.14 | (-2.05--0.14) | **2.66E-02** | **4.98E-02** | 0.84 | (-1.15-2.87) | 4.10E-01 | 5.58E-01 |
| 25 | DG(16:0_16:1) | 0.64 | (-0.39-1.68) | 2.23E-01 | 2.80E-01 | 4.43 | (3.46-5.4) | **9.58E-19** | **4.12E-17** | -1.32 | (-2.11--0.45) | **3.64E-03** | **8.13E-03** | 6.21 | (3.58-8.89) | **3.25E-06** | **8.67E-05** |
| 26 | DG(14:0_18:2) | -1.03 | (-2.03--0.02) | **4.55E-02** | 7.18E-02 | 4.19 | (2.86-5.54) | **7.61E-10** | **4.11E-09** | -3.08 | (-4.04--1.96) | **1.68E-06** | **5.70E-06** | 6.70 | (3.02-10.51) | **3.23E-04** | **2.28E-03** |
| 27 | DG(16:0_18:1) | 1.33 | (0.59-2.07) | **4.36E-04** | **1.22E-03** | 4.26 | (3.33-5.21) | **1.32E-18** | **4.86E-17** | -1.91 | (-2.57--1.19) | **1.35E-06** | **4.62E-06** | 3.90 | (1.38-6.49) | **2.40E-03** | **1.18E-02** |
| 28 | DG(16:1_18:1) | 0.78 | (-0.08-1.65) | 7.61E-02 | 1.10E-01 | 4.04 | (3.07-5.03) | **1.05E-15** | **1.80E-14** | -2.08 | (-2.77--1.32) | **7.91E-07** | **2.84E-06** | 6.17 | (3.26-9.17) | **2.92E-05** | **4.64E-04** |
| 29 | DG(16:0_18:2) | 0.92 | (0.21-1.65) | **1.15E-02** | **2.16E-02** | 3.62 | (2.72-4.52) | **5.53E-15** | **7.59E-14** | -3.07 | (-3.76--2.31) | **1.09E-11** | **6.22E-11** | 4.00 | (1.57-6.48) | **1.20E-03** | **6.49E-03** |
| 30 | DG(18:0_18:1) | 1.36 | (0.69-2.04) | **7.91E-05** | **2.60E-04** | 4.41 | (3.51-5.31) | **4.70E-21** | **4.51E-19** | -1.29 | (-1.99--0.53) | **1.23E-03** | **2.94E-03** | 4.06 | (1.71-6.46) | **6.87E-04** | **4.23E-03** |
| 31 | DG(18:1_18:1) | 1.37 | (0.64-2.1) | **2.23E-04** | **6.69E-04** | 2.93 | (2.08-3.77) | **1.44E-11** | **1.03E-10** | -1.85 | (-2.5--1.16) | **1.20E-06** | **4.16E-06** | 2.05 | (-0.25-4.4) | 8.10E-02 | 1.74E-01 |
| 32 | DG(18:0_18:2) | 0.99 | (0.23-1.76) | **1.04E-02** | **1.97E-02** | 4.40 | (3.41-5.39) | **7.51E-18** | **2.57E-16** | -1.62 | (-2.52--0.61) | **2.43E-03** | **5.62E-03** | 3.76 | (1.3-6.29) | **2.68E-03** | **1.27E-02** |
| 33 | DG(18:1_18:2) | 0.95 | (0.22-1.68) | **1.02E-02** | **1.97E-02** | 2.26 | (1.45-3.08) | **5.91E-08** | **2.42E-07** | -3.15 | (-3.76--2.48) | **8.45E-15** | **7.12E-14** | 1.89 | (-0.33-4.15) | 9.51E-02 | 1.93E-01 |
| 34 | DG(18:2_18:2) | 0.88 | (-0.05-1.82) | 6.31E-02 | 9.47E-02 | 1.84 | (0.87-2.81) | **1.86E-04** | **4.53E-04** | -4.16 | (-4.84--3.39) | **9.59E-17** | **1.10E-15** | 2.33 | (-0.23-4.96) | 7.47E-02 | 1.63E-01 |
| 35 | DG(18:1_18:3) | 0.37 | (-0.41-1.15) | 3.56E-01 | 4.21E-01 | 2.29 | (1.37-3.21) | **1.02E-06** | **3.33E-06** | -2.63 | (-3.39--1.77) | **6.81E-08** | **2.73E-07** | 3.15 | (0.38-6.01) | **2.58E-02** | 7.63E-02 |
| 36 | DG(16:0_20:4) | 1.19 | (0.26-2.12) | **1.22E-02** | **2.27E-02** | 4.27 | (3.25-5.3) | **4.79E-16** | **9.59E-15** | -2.06 | (-2.83--1.21) | **9.20E-06** | **2.95E-05** | 4.97 | (2.34-7.66) | **1.98E-04** | **1.61E-03** |
| 37 | DG(18:1_20:3) | 0.43 | (-0.33-1.19) | 2.68E-01 | 3.33E-01 | 2.33 | (1.5-3.17) | **4.24E-08** | **1.82E-07** | -0.30 | (-1.09-0.55) | 4.72E-01 | 5.69E-01 | 1.17 | (-1.22-3.63) | 3.39E-01 | 4.95E-01 |
| 38 | DG(18:1_20:4) | 1.14 | (0.4-1.88) | **2.42E-03** | **5.53E-03** | 2.47 | (1.71-3.22) | **1.81E-10** | **1.10E-09** | -2.06 | (-2.71--1.36) | **1.08E-07** | **4.14E-07** | 3.90 | (1.68-6.16) | **5.47E-04** | **3.60E-03** |
| 39 | DG(16:0_22:5) | 0.58 | (-0.3-1.46) | 2.00E-01 | 2.56E-01 | 2.42 | (1.46-3.38) | **7.01E-07** | **2.39E-06** | -1.95 | (-2.72--1.09) | **2.99E-05** | **9.02E-05** | 4.60 | (2.03-7.23) | **4.22E-04** | **2.89E-03** |
| 40 | DG(18:2_20:4) | 0.81 | (-0.04-1.66) | 6.22E-02 | 9.35E-02 | 2.42 | (1.55-3.3) | **5.34E-08** | **2.23E-07** | -3.14 | (-3.87--2.33) | **7.74E-11** | **4.08E-10** | 4.23 | (1.78-6.73) | **6.78E-04** | **4.23E-03** |
| 41 | DG(16:0_22:6) | 1.16 | (0.04-2.3) | **4.29E-02** | 6.89E-02 | 4.15 | (2.87-5.45) | **2.35E-10** | **1.41E-09** | -4.09 | (-4.76--3.33) | **6.80E-17** | **8.37E-16** | 5.50 | (2.13-8.99) | **1.31E-03** | **6.84E-03** |
| 42 | DG(18:1_22:5) | 0.51 | (-0.32-1.35) | 2.30E-01 | 2.89E-01 | 1.38 | (0.54-2.23) | **1.34E-03** | **2.78E-03** | -1.90 | (-2.66--1.06) | **2.96E-05** | **9.00E-05** | 1.63 | (-0.72-4.04) | 1.75E-01 | 3.07E-01 |
| 43 | DG(18:1_22:6) | 1.53 | (0.47-2.61) | **4.63E-03** | **9.97E-03** | 3.18 | (2.04-4.34) | **5.26E-08** | **2.22E-07** | -3.49 | (-4.15--2.76) | **7.44E-15** | **6.49E-14** | 4.14 | (1.32-7.04) | **3.88E-03** | **1.73E-02** |
| 44 | DG(18:2_22:6) | 1.52 | (0.42-2.63) | **6.55E-03** | **1.35E-02** | 3.05 | (1.87-4.24) | **3.60E-07** | **1.28E-06** | -4.00 | (-4.74--3.16) | **9.00E-14** | **6.75E-13** | 4.79 | (1.45-8.24) | **4.80E-03** | **2.04E-02** |

| | | | | | | | | | | | | | | | | | |
|---|---|---|---|---|---|---|---|---|---|---|---|---|---|---|---|---|---|
| 45 | TG(48:0) [NL-18:0] | -1.56 | (-2.43--0.68) | **5.76E-04** | **1.53E-03** | 4.34 | (2.99-5.71) | **3.54E-10** | **2.07E-09** | -1.44 | (-2.28--0.51) | **3.14E-03** | **7.14E-03** | 7.45 | (3.74-11.29) | **7.09E-05** | **8.54E-04** |
| 46 | TG(48:1) [NL-16:1] | 0.51 | (-0.62-1.65) | 3.76E-01 | 4.39E-01 | 6.64 | (5.32-7.97) | **1.51E-22** | **3.62E-20** | -2.43 | (-3.32--1.41) | **1.61E-05** | **5.11E-05** | 9.58 | (5.88-13.41) | **2.99E-07** | **1.32E-05** |
| 47 | TG(48:1) [NL-18:1] | -0.61 | (-1.49-0.27) | 1.75E-01 | 2.30E-01 | 5.21 | (3.76-6.69) | **2.44E-12** | **2.13E-11** | -3.15 | (-3.94--2.26) | **2.16E-09** | **9.86E-09** | 8.20 | (3.97-12.59) | **1.20E-04** | **1.17E-03** |
| 48 | TG(48:2) [NL-14:0] | -0.94 | (-1.97-0.1) | 7.69E-02 | 1.11E-01 | 5.21 | (3.73-6.72) | **5.81E-12** | **4.73E-11** | -3.76 | (-4.46--2.97) | **4.13E-14** | **3.31E-13** | 8.73 | (4.41-13.22) | **6.10E-05** | **7.91E-04** |
| 49 | TG(48:2) [NL-14:1] | -1.20 | (-2.25--0.13) | **2.78E-02** | **4.70E-02** | 4.79 | (3.42-6.18) | **7.51E-12** | **5.72E-11** | -3.06 | (-3.81--2.23) | **4.50E-10** | **2.23E-09** | 8.90 | (4.81-13.16) | **1.61E-05** | **3.36E-04** |
| 50 | TG(48:2) [NL-16:1] | -0.17 | (-1.47-1.15) | 8.03E-01 | 8.32E-01 | 5.71 | (4.32-7.12) | **1.20E-15** | **1.99E-14** | -2.85 | (-3.65--1.96) | **3.48E-08** | **1.43E-07** | 10.74 | (6.57-15.07) | **3.03E-07** | **1.32E-05** |
| 51 | TG(48:2) [NL-18:2] | -1.16 | (-2.2--0.12) | **2.91E-02** | **4.87E-02** | 5.21 | (3.68-6.77) | **2.87E-11** | **2.03E-10** | -4.14 | (-4.86--3.32) | **5.42E-15** | **4.91E-14** | 8.58 | (4.3-13.04) | **7.11E-05** | **8.54E-04** |
| 52 | TG(48:3) [NL-14:0] | -1.51 | (-2.64--0.37) | **9.55E-03** | **1.85E-02** | 4.32 | (2.9-5.76) | **2.31E-09** | **1.14E-08** | -4.04 | (-4.73--3.26) | **1.02E-15** | **1.04E-14** | 9.53 | (5.17-14.08) | **1.43E-05** | **3.28E-04** |
| 53 | TG(48:3) [NL-16:1] | -1.14 | (-2.29-0.02) | 5.44E-02 | 8.37E-02 | 4.00 | (2.79-5.24) | **1.30E-10** | **8.10E-10** | -2.89 | (-3.59--2.12) | **1.90E-10** | **9.60E-10** | 8.59 | (4.99-12.31) | **2.28E-06** | **6.49E-05** |
| 54 | TG(48:3) [NL-18:3] | -1.81 | (-3.07--0.53) | **5.76E-03** | **1.21E-02** | 4.99 | (3.41-6.59) | **5.23E-10** | **2.92E-09** | -3.23 | (-3.91--2.48) | **8.75E-13** | **5.83E-12** | 8.18 | (3.69-12.86) | **3.09E-04** | **2.27E-03** |
| 55 | TG(49:1) [NL-17:1] | 0.33 | (-0.53-1.21) | 4.51E-01 | 5.10E-01 | 4.04 | (3.11-4.97) | **2.94E-17** | **8.82E-16** | -2.19 | (-2.84--1.47) | **5.29E-08** | **2.14E-07** | 4.94 | (2.47-7.47) | **8.54E-05** | **9.54E-04** |
| 56 | TG(50:1) [NL-14:0] | -0.81 | (-1.57--0.05) | **3.75E-02** | 6.17E-02 | 4.78 | (3.52-6.05) | **1.11E-13** | **1.24E-12** | -1.46 | (-2.35--0.47) | **4.76E-03** | **1.03E-02** | 6.89 | (3.47-10.42) | **7.00E-05** | **8.54E-04** |
| 57 | TG(50:1) [NL-16:0] | 0.83 | (0.22-1.43) | **7.27E-03** | **1.46E-02** | 4.34 | (3.39-5.3) | **1.03E-18** | **4.12E-17** | -2.49 | (-3.25--1.64) | **2.17E-07** | **8.15E-07** | 4.63 | (2.1-7.22) | **3.13E-04** | **2.27E-03** |
| 58 | TG(50:1) [NL-18:1] | 0.76 | (0.18-1.34) | **1.03E-02** | **1.97E-02** | 4.10 | (3.16-5.05) | **4.06E-17** | **1.08E-15** | -2.47 | (-3.24--1.62) | **2.61E-07** | **9.63E-07** | 4.17 | (1.73-6.68) | **7.99E-04** | **4.74E-03** |
| 59 | TG(50:2) [NL-14:0] | -0.65 | (-1.21--0.07) | **2.69E-02** | **4.57E-02** | 2.80 | (1.76-3.86) | **1.50E-07** | **5.54E-07** | -3.34 | (-4.1--2.48) | **1.16E-10** | **6.01E-10** | 4.36 | (1.14-7.69) | **7.83E-03** | **3.01E-02** |
| 60 | TG(50:2) [NL-16:1] | 0.65 | (-0.07-1.38) | 7.62E-02 | 1.10E-01 | 4.04 | (3.07-5.02) | **6.67E-16** | **1.28E-14** | -3.01 | (-3.81--2.11) | **1.08E-08** | **4.67E-08** | 6.45 | (3.42-9.58) | **2.79E-05** | **4.62E-04** |
| 61 | TG(50:2) [NL-18:1] | 0.32 | (-0.31-0.95) | 3.19E-01 | 3.85E-01 | 3.59 | (2.65-4.54) | **1.19E-13** | **1.30E-12** | -3.12 | (-3.91--2.21) | **4.54E-09** | **2.00E-08** | 6.24 | (3.2-9.38) | **5.20E-05** | **7.33E-04** |
| 62 | TG(50:2) [NL-18:2] | 0.66 | (0.08-1.24) | **2.58E-02** | **4.41E-02** | 3.72 | (2.82-4.62) | **8.45E-16** | **1.53E-14** | -4.17 | (-4.92--3.31) | **5.26E-14** | **4.14E-13** | 4.53 | (2.11-7) | **2.30E-04** | **1.78E-03** |
| 63 | TG(50:3) [NL-14:0] | -0.51 | (-1.24-0.22) | 1.70E-01 | 2.25E-01 | 3.13 | (1.93-4.34) | **2.88E-07** | **1.04E-06** | -4.25 | (-4.95--3.45) | **4.04E-16** | **4.31E-15** | 5.47 | (1.88-9.18) | **2.69E-03** | **1.27E-02** |
| 64 | TG(50:3) [NL-14:1] | -0.64 | (-1.29-0.02) | 5.67E-02 | 8.67E-02 | 2.66 | (1.69-3.63) | **7.85E-08** | **3.14E-07** | -2.39 | (-3.17--1.51) | **1.11E-06** | **3.90E-06** | 4.17 | (1.33-7.09) | **3.92E-03** | **1.73E-02** |
| 65 | TG(50:3) [NL-16:1] | 0.58 | (-0.25-1.43) | 1.70E-01 | 2.25E-01 | 4.38 | (3.28-5.5) | **1.32E-14** | **1.67E-13** | -3.88 | (-4.58--3.09) | **8.11E-15** | **6.95E-14** | 8.28 | (4.83-11.85) | **2.15E-06** | **6.49E-05** |
| 66 | TG(50:3) [NL-18:2] | 0.28 | (-0.45-1.02) | 4.51E-01 | 5.10E-01 | 3.72 | (2.65-4.81) | **1.38E-11** | **1.00E-10** | -4.37 | (-5.06--3.59) | **3.53E-17** | **4.70E-16** | 7.14 | (3.78-10.61) | **2.72E-05** | **4.62E-04** |
| 67 | TG(50:3) [NL-18:3] | 0.14 | (-0.78-1.07) | 7.65E-01 | 8.02E-01 | 5.18 | (3.92-6.44) | **8.59E-16** | **1.53E-14** | -3.81 | (-4.54--2.98) | **2.31E-13** | **1.65E-12** | 6.76 | (3.2-10.44) | **1.78E-04** | **1.52E-03** |
| 68 | TG(50:4) [NL-14:0] | -0.87 | (-1.89-0.15) | 9.35E-02 | 1.31E-01 | 2.75 | (1.47-4.04) | **2.32E-05** | **6.36E-05** | -4.31 | (-4.97--3.57) | **2.52E-18** | **4.32E-17** | 5.22 | (1.61-8.96) | **4.43E-03** | **1.92E-02** |
| 69 | TG(50:4) [NL-18:3] | -0.72 | (-1.76-0.33) | 1.76E-01 | 2.31E-01 | 4.14 | (2.91-5.38) | **5.20E-11** | **3.51E-10** | -3.93 | (-4.56--3.22) | **9.72E-18** | **1.37E-16** | 7.93 | (4.25-11.74) | **2.01E-05** | **4.02E-04** |
| 70 | TG(50:4) [NL-20:4] | -0.81 | (-2.19-0.58) | 2.50E-01 | 3.11E-01 | 5.64 | (4.02-7.29) | **8.80E-12** | **6.60E-11** | -3.52 | (-4.34--2.57) | **6.23E-10** | **3.05E-09** | 10.00 | (5.69-14.48) | **3.96E-06** | **1.00E-04** |
| 71 | TG(51:1) [NL-17:0] | 0.67 | (-0.08-1.42) | 7.92E-02 | 1.13E-01 | 4.54 | (3.5-5.59) | **2.82E-17** | **8.82E-16** | -2.72 | (-3.44--1.91) | **4.45E-09** | **1.98E-08** | 4.90 | (2-7.89) | **8.93E-04** | **5.23E-03** |
| 72 | TG(51:2) [NL-15:0] | -0.52 | (-1.18-0.13) | 1.19E-01 | 1.65E-01 | 2.32 | (1.4-3.24) | **7.50E-07** | **2.50E-06** | -3.37 | (-4.11--2.54) | **1.77E-11** | **9.75E-11** | 2.95 | (0.1-5.89) | **4.21E-02** | 1.10E-01 |
| 73 | TG(51:2) [NL-17:0] | 0.37 | (-0.32-1.06) | 2.96E-01 | 3.59E-01 | 4.07 | (3.06-5.09) | **6.13E-15** | **8.18E-14** | -3.40 | (-4.11--2.6) | **2.54E-12** | **1.63E-11** | 5.48 | (2.52-8.52) | **2.67E-04** | **2.03E-03** |
| 74 | TG(51:2) [NL-17:1] | 0.36 | (-0.31-1.02) | 2.93E-01 | 3.56E-01 | 3.80 | (2.86-4.74) | **3.86E-15** | **5.98E-14** | -2.89 | (-3.53--2.19) | **2.44E-12** | **1.58E-11** | 4.90 | (2.22-7.65) | **3.17E-04** | **2.27E-03** |
| 75 | TG(52:1) [NL-18:0] | 1.38 | (0.59-2.18) | **6.58E-04** | **1.73E-03** | 5.69 | (4.53-6.86) | **2.15E-21** | **2.58E-19** | -1.86 | (-2.76--0.83) | **6.81E-04** | **1.69E-03** | 5.79 | (2.8-8.86) | **1.32E-04** | **1.22E-03** |
| 76 | TG(52:1) [NL-18:1] | 1.45 | (0.69-2.23) | **2.04E-04** | **6.17E-04** | 5.86 | (4.73-7) | **1.79E-23** | **8.60E-21** | -1.80 | (-2.71--0.77) | **9.96E-04** | **2.41E-03** | 5.71 | (2.83-8.68) | **9.59E-05** | **1.00E-03** |
| 77 | TG(52:2) [NL-16:0] | 0.80 | (0.4-1.21) | **1.01E-04** | **3.29E-04** | 2.18 | (1.57-2.79) | **3.69E-12** | **3.11E-11** | -2.79 | (-3.44--2.07) | **3.12E-11** | **1.68E-10** | 1.93 | (0.21-3.67) | **2.76E-02** | 7.97E-02 |
| 78 | TG(52:2) [NL-18:2] | 0.46 | (-0.15-1.07) | 1.38E-01 | 1.88E-01 | 4.17 | (3.27-5.07) | **4.36E-19** | **2.33E-17** | -2.75 | (-3.58--1.81) | **3.20E-07** | **1.17E-06** | 4.45 | (2.1-6.85) | **1.98E-04** | **1.61E-03** |
| 79 | TG(52:3) [NL-16:1] | 0.39 | (-0.18-0.97) | 1.79E-01 | 2.35E-01 | 1.94 | (1.21-2.66) | **1.73E-07** | **6.33E-07** | -3.37 | (-4.1--2.56) | **1.00E-11** | **5.79E-11** | 3.39 | (0.97-5.88) | **5.95E-03** | **2.36E-02** |
| 80 | TG(52:3) [NL-18:2] | 0.76 | (0.32-1.2) | **8.08E-04** | **2.08E-03** | 1.84 | (1.22-2.48) | **1.11E-08** | **5.14E-08** | -4.40 | (-5.07--3.64) | **3.04E-18** | **4.87E-17** | 1.85 | (-0.08-3.81) | 6.03E-02 | 1.43E-01 |
| 81 | TG(52:4) [NL-16:1] | 0.68 | (-0.05-1.41) | 6.77E-02 | 9.99E-02 | 2.22 | (1.27-3.18) | **4.67E-06** | **1.43E-05** | -4.52 | (-5.17--3.78) | **1.14E-19** | **2.48E-18** | 4.16 | (1.08-7.34) | **7.91E-03** | **3.01E-02** |
| 82 | TG(52:4) [NL-18:2] | 1.20 | (0.44-1.96) | **1.92E-03** | **4.52E-03** | 2.29 | (1.29-3.3) | **7.06E-06** | **2.12E-05** | -4.97 | (-5.63--4.21) | **2.07E-20** | **5.23E-19** | 3.62 | (0.64-6.69) | **1.72E-02** | 5.52E-02 |
| 83 | TG(52:4) [NL-18:3] | 0.47 | (-0.24-1.18) | 1.97E-01 | 2.54E-01 | 3.19 | (2.17-4.21) | **8.03E-10** | **4.24E-09** | -4.11 | (-4.78--3.35) | **9.43E-17** | **1.10E-15** | 4.40 | (1.43-7.46) | **3.60E-03** | **1.63E-02** |
| 84 | TG(52:5) [NL-18:3] | 0.10 | (-0.9-1.11) | 8.49E-01 | 8.72E-01 | 2.89 | (1.78-4.01) | **3.35E-07** | **1.20E-06** | -4.16 | (-4.81--3.43) | **2.87E-18** | **4.75E-17** | 5.19 | (2-8.47) | **1.36E-03** | **6.99E-03** |
| 85 | TG(52:5) [NL-20:4] | 0.48 | (-0.71-1.69) | 4.28E-01 | 4.86E-01 | 5.46 | (4.2-6.74) | **3.97E-17** | **1.08E-15** | -3.63 | (-4.4--2.74) | **2.19E-11** | **1.19E-10** | 10.03 | (6.39-13.8) | **4.75E-08** | **4.56E-06** |
| 86 | TG(52:5) [NL-20:5] | 0.85 | (-0.43-2.14) | 1.95E-01 | 2.52E-01 | 5.06 | (3.69-6.44) | **4.80E-13** | **4.70E-12** | -4.43 | (-5.13--3.64) | **4.66E-17** | **5.88E-16** | 7.78 | (4.53-11.13) | **2.30E-06** | **6.49E-05** |
| 87 | TG(53:2) [NL-18:1] | 0.88 | (0.2-1.56) | **1.08E-02** | **2.03E-02** | 2.66 | (1.85-3.47) | **1.63E-10** | **1.00E-09** | -2.69 | (-3.33--1.98) | **5.26E-11** | **2.81E-10** | 1.90 | (-0.42-4.27) | 1.10E-01 | 2.14E-01 |
| 88 | TG(54:1) [NL-18:1] | 0.99 | (0.32-1.66) | **3.57E-03** | **8.00E-03** | 4.45 | (3.57-5.34) | **4.54E-22** | **7.26E-20** | -1.64 | (-2.56--0.59) | **2.89E-03** | **6.61E-03** | 3.54 | (1.13-6.02) | **3.96E-03** | **1.73E-02** |
| 89 | TG(54:2) [NL-18:0] | 1.41 | (0.72-2.1) | **6.55E-05** | **2.23E-04** | 4.24 | (3.35-5.13) | **2.72E-20** | **1.63E-18** | -1.85 | (-2.69--0.91) | **2.40E-04** | **6.39E-04** | 3.30 | (0.94-5.71) | **6.02E-03** | **2.36E-02** |
| 90 | TG(54:2) [NL-20:1] | 0.43 | (-0.34-1.21) | 2.74E-01 | 3.39E-01 | 2.74 | (1.85-3.64) | **1.93E-09** | **9.85E-09** | -1.84 | (-2.64--0.95) | **1.25E-04** | **3.49E-04** | 1.66 | (-0.73-4.12) | 1.74E-01 | 3.07E-01 |
| 91 | TG(54:3) [NL-18:1] | 0.57 | (0.01-1.13) | **4.50E-02** | 7.13E-02 | 1.26 | (0.64-1.88) | **7.52E-05** | **1.93E-04** | -2.18 | (-2.89--1.41) | **4.18E-07** | **1.52E-06** | 0.18 | (-1.55-1.95) | 8.36E-01 | 8.99E-01 |
| 92 | TG(54:3) [NL-18:2] | 0.72 | (0.14-1.3) | **1.43E-02** | **2.59E-02** | 2.23 | (1.57-2.89) | **5.13E-11** | **3.51E-10** | -3.06 | (-3.75--2.3) | **1.28E-11** | **7.23E-11** | 2.32 | (0.48-4.19) | **1.32E-02** | **4.65E-02** |
| 93 | TG(54:4) [NL-18:2] | 0.35 | (-0.28-0.98) | 2.76E-01 | 3.40E-01 | 0.85 | (0.16-1.55) | **1.61E-02** | **2.62E-02** | -4.03 | (-4.74--3.23) | **5.43E-15** | **4.91E-14** | 0.62 | (-1.51-2.8) | 5.68E-01 | 6.95E-01 |
| 94 | TG(54:4) [NL-20:3] | 0.64 | (-0.05-1.33) | 6.76E-02 | 9.99E-02 | 3.70 | (2.83-4.58) | **2.18E-16** | **4.75E-15** | -2.73 | (-3.52--1.84) | **8.04E-08** | **3.16E-07** | 4.94 | (2.41-7.54) | **1.24E-04** | **1.19E-03** |
| 95 | TG(54:5) [NL-18:3] | 0.07 | (-0.8-0.94) | 8.78E-01 | 8.96E-01 | 1.87 | (0.89-2.85) | **1.77E-04** | **4.33E-04** | -3.77 | (-4.42--3.03) | **5.03E-16** | **5.25E-15** | 2.95 | (-0.03-6.02) | 5.22E-02 | 1.29E-01 |
| 96 | TG(54:5) [NL-20:4] | 1.04 | (0.38-1.71) | **1.91E-03** | **4.51E-03** | 3.37 | (2.55-4.2) | **2.12E-15** | **3.39E-14** | -3.73 | (-4.49--2.86) | **4.40E-12** | **2.64E-11** | 5.84 | (3.43-8.3) | **1.90E-06** | **6.49E-05** |

| | | | | | | | | | | | | | | | | | |
|---|---|---|---|---|---|---|---|---|---|---|---|---|---|---|---|---|---|
| 97 | TG(54:6) [NL-18:3] | -0.40 | (-1.56-0.76) | 4.95E-01 | 5.50E-01 | 1.37 | (0.22-2.54) | **1.91E-02** | **3.03E-02** | -3.86 | (-4.52--3.12) | **3.29E-16** | **3.59E-15** | 2.95 | (-0.25-6.26) | 7.12E-02 | 1.58E-01 |
| 98 | TG(54:6) [NL-20:4] | 1.16 | (0.33-2) | **6.24E-03** | **1.29E-02** | 3.69 | (2.71-4.68) | **2.17E-13** | **2.27E-12** | -4.11 | (-4.83--3.28) | **1.19E-14** | **9.81E-14** | 7.52 | (4.68-10.44) | **1.91E-07** | **1.15E-05** |
| 99 | TG(54:6) [NL-20:5] | 0.85 | (-0.29-1.99) | 1.44E-01 | 1.96E-01 | 3.82 | (2.52-5.13) | **7.53E-09** | **3.51E-08** | -5.04 | (-5.69--4.28) | **7.74E-21** | **2.50E-19** | 8.39 | (5.04-11.85) | **7.54E-07** | **3.02E-05** |
| 100 | TG(54:6) [NL-22:6] | 1.79 | (0.57-3.02) | **4.10E-03** | **9.06E-03** | 5.76 | (4.33-7.22) | **4.96E-15** | **7.00E-14** | -5.16 | (-5.83--4.38) | **1.98E-20** | **5.23E-19** | 8.20 | (4.48-12.06) | **1.29E-05** | **3.10E-04** |
| 101 | TG(54:7) [NL-20:5] | 0.47 | (-0.81-1.77) | 4.74E-01 | 5.30E-01 | 3.45 | (2.17-4.75) | **1.37E-07** | **5.08E-07** | -4.97 | (-5.63--4.22) | **1.23E-20** | **3.49E-19** | 8.46 | (5.22-11.8) | **2.49E-07** | **1.32E-05** |
| 102 | TG(54:7) [NL-22:6] | 0.98 | (-0.34-2.32) | 1.48E-01 | 1.99E-01 | 5.09 | (3.68-6.52) | **1.79E-12** | **1.62E-11** | -4.92 | (-5.56--4.18) | **1.93E-21** | **7.12E-20** | 10.42 | (6.51-14.47) | **1.23E-07** | **9.83E-06** |
| 103 | TG(56:6) [NL-20:4] | 1.34 | (0.81-1.87) | **7.12E-07** | **3.60E-06** | 1.36 | (0.85-1.88) | **2.38E-07** | **8.65E-07** | -3.00 | (-3.77--2.15) | **1.89E-09** | **8.92E-09** | 3.72 | (1.91-5.57) | **5.78E-05** | **7.86E-04** |
| 104 | TG(56:6) [NL-22:5] | 0.62 | (-0.03-1.28) | 6.20E-02 | 9.35E-02 | 1.99 | (1.2-2.77) | **7.33E-07** | **2.46E-06** | -3.83 | (-4.52--3.06) | **4.63E-15** | **4.36E-14** | 3.96 | (1.82-6.14) | **2.87E-04** | **2.15E-03** |
| 105 | TG(56:7) [NL-20:4] | 1.02 | (0.35-1.7) | **2.84E-03** | **6.40E-03** | 1.88 | (1.14-2.62) | **6.63E-07** | **2.29E-06** | -3.66 | (-4.41--2.81) | **3.38E-12** | **2.08E-11** | 4.16 | (1.73-6.64) | **7.60E-04** | **4.56E-03** |
| 106 | TG(56:7) [NL-20:5] | 0.39 | (-0.72-1.5) | 4.93E-01 | 5.50E-01 | 2.47 | (1.35-3.61) | **1.56E-05** | **4.46E-05** | -3.79 | (-4.51--2.97) | **1.22E-13** | **9.04E-13** | 5.58 | (2.64-8.6) | **1.82E-04** | **1.54E-03** |
| 107 | TG(56:7) [NL-22:5] | 0.79 | (0.02-1.56) | **4.44E-02** | 7.06E-02 | 2.55 | (1.61-3.5) | **1.14E-07** | **4.45E-07** | -4.29 | (-4.93--3.57) | **2.43E-19** | **4.87E-18** | 5.15 | (2.46-7.91) | **1.64E-04** | **1.46E-03** |
| 108 | TG(56:7) [NL-22:6] | 1.93 | (0.84-3.04) | **4.97E-04** | **1.35E-03** | 3.96 | (2.65-5.28) | **3.33E-09** | **1.61E-08** | -5.07 | (-5.72--4.33) | **1.85E-21** | **7.12E-20** | 6.45 | (3.01-10) | **2.16E-04** | **1.70E-03** |
| 109 | TG(56:8) [NL-20:4] | 0.80 | (-0.13-1.74) | 9.22E-02 | 1.30E-01 | 1.88 | (1-2.77) | **2.85E-05** | **7.72E-05** | -2.95 | (-3.72--2.08) | **5.67E-09** | **2.47E-08** | 5.12 | (2.54-7.75) | **9.23E-05** | **1.00E-03** |
| 110 | TG(56:8) [NL-20:5] | 0.18 | (-1.06-1.45) | 7.72E-01 | 8.06E-01 | 1.59 | (0.43-2.77) | **7.20E-03** | **1.26E-02** | -4.13 | (-4.79--3.37) | **3.84E-17** | **4.98E-16** | 5.03 | (2.01-8.13) | **1.03E-03** | **5.94E-03** |
| 111 | TG(56:8) [NL-22:6] | 1.76 | (0.54-2.99) | **4.49E-03** | **9.76E-03** | 3.97 | (2.64-5.31) | **4.96E-09** | **2.38E-08** | -5.36 | (-5.97--4.65) | **1.69E-24** | **1.02E-22** | 7.57 | (4.02-11.24) | **2.51E-05** | **4.62E-04** |
| 112 | TG(56:9) [NL-22:6] | 0.79 | (-0.4-1.99) | 1.94E-01 | 2.50E-01 | 3.07 | (1.89-4.27) | **3.82E-07** | **1.35E-06** | -4.88 | (-5.5--4.17) | **1.57E-22** | **7.54E-21** | 6.66 | (3.29-10.14) | **9.49E-05** | **1.00E-03** |
| 113 | TG(58:10) [NL-22:6] | 0.95 | (-0.23-2.15) | 1.13E-01 | 1.58E-01 | 2.08 | (0.92-3.25) | **4.34E-04** | **9.92E-04** | -4.48 | (-5.11--3.76) | **3.35E-20** | **8.05E-19** | 5.17 | (1.89-8.56) | **1.90E-03** | **9.52E-03** |
| 114 | TG(58:8) [NL-22:6] | 1.46 | (0.38-2.56) | **7.98E-03** | **1.58E-02** | 2.80 | (1.58-4.04) | **7.21E-06** | **2.15E-05** | -4.50 | (-5.16--3.76) | **3.67E-19** | **7.05E-18** | 4.73 | (1.39-8.18) | **5.34E-03** | **2.16E-02** |
| 115 | TG(58:9) [NL-22:6] | 1.22 | (0.07-2.38) | **3.70E-02** | 6.10E-02 | 2.17 | (0.97-3.38) | **3.75E-04** | **8.69E-04** | -4.76 | (-5.39--4.06) | **4.43E-22** | **1.93E-20** | 4.51 | (1.17-7.97) | **7.98E-03** | **3.02E-02** |
| 116 | TG(O-50:1) [NL-16:0] | -0.20 | (-0.8-0.39) | 5.04E-01 | 5.56E-01 | 0.74 | (-0.02-1.51) | 5.73E-02 | 8.14E-02 | -0.59 | (-1.3-0.18) | 1.29E-01 | 2.00E-01 | 1.88 | (-0.28-4.09) | 8.85E-02 | 1.85E-01 |
| 117 | TG(O-50:1) [NL-17:1] | 0.33 | (-0.54-1.21) | 4.53E-01 | 5.11E-01 | 4.08 | (3.14-5.03) | **4.97E-17** | **1.26E-15** | -2.21 | (-2.88--1.47) | **9.38E-08** | **3.66E-07** | 4.99 | (2.47-7.57) | **1.01E-04** | **1.03E-03** |
| 118 | TG(O-50:1) [NL-18:1] | -0.61 | (-1.27-0.05) | 6.87E-02 | 1.01E-01 | 0.57 | (-0.34-1.49) | 2.18E-01 | 2.74E-01 | -0.98 | (-1.68--0.22) | **1.24E-02** | **2.49E-02** | 2.75 | (0.37-5.19) | **2.36E-02** | 7.16E-02 |
| 119 | TG(O-50:2) [NL-18:2] | -0.07 | (-0.64-0.51) | 8.21E-01 | 8.50E-01 | 0.77 | (0.05-1.5) | **3.73E-02** | 5.56E-02 | -0.45 | (-1.21-0.36) | 2.64E-01 | 3.60E-01 | 2.97 | (0.89-5.09) | **5.11E-03** | **2.10E-02** |
| 120 | TG(O-52:2) [NL-16:0] | -0.21 | (-1.02-0.6) | 6.04E-01 | 6.52E-01 | 2.25 | (1.31-3.2) | **2.59E-06** | **8.08E-06** | -1.15 | (-2.06--0.15) | **2.56E-02** | **4.81E-02** | 3.32 | (0.48-6.25) | **2.20E-02** | 6.79E-02 |
| 121 | TG(O-52:2) [NL-17:1] | 0.36 | (-0.31-1.03) | 2.93E-01 | 3.56E-01 | 3.82 | (2.87-4.77) | **4.43E-15** | **6.45E-14** | -2.91 | (-3.55--2.2) | **4.20E-12** | **2.55E-11** | 4.89 | (2.19-7.66) | **3.63E-04** | **2.52E-03** |
| 122 | TG(O-52:2) [NL-18:1] | -0.45 | (-1.28-0.38) | 2.85E-01 | 3.50E-01 | 2.26 | (1.27-3.27) | **8.29E-06** | **2.44E-05** | -1.84 | (-2.72--0.85) | **5.37E-04** | **1.36E-03** | 5.04 | (2-8.16) | **1.08E-03** | **6.15E-03** |
| 123 | TG(O-54:2) [NL-18:1] | 0.85 | (0.18-1.53) | **1.33E-02** | **2.43E-02** | 2.63 | (1.82-3.44) | **2.48E-10** | **1.47E-09** | -2.66 | (-3.31--1.95) | **1.06E-10** | **5.53E-10** | 1.86 | (-0.46-4.23) | 1.16E-01 | 2.20E-01 |
| 124 | TG(O-54:4) [NL-17:1] | 0.08 | (-0.68-0.85) | 8.36E-01 | 8.61E-01 | 1.61 | (0.72-2.51) | **4.02E-04** | **9.24E-04** | -4.03 | (-4.73--3.24) | **1.59E-15** | **1.56E-14** | 2.88 | (0.29-5.53) | **2.92E-02** | 8.30E-02 |
| 125 | TG(O-54:4) [NL-18:2] | 0.07 | (-0.73-0.88) | 8.68E-01 | 8.88E-01 | 1.49 | (0.53-2.46) | **2.38E-03** | **4.69E-03** | -4.57 | (-5.24--3.8) | **1.07E-18** | **1.90E-17** | 2.73 | (-0.04-5.59) | 5.37E-02 | 1.31E-01 |
| 126 | dhCer(d18:0/22:0) | 1.14 | (0.5-1.79) | **4.77E-04** | **1.32E-03** | 1.96 | (1.35-2.58) | **4.98E-10** | **2.81E-09** | -0.06 | (-0.73-0.66) | 8.74E-01 | 9.06E-01 | 1.23 | (-0.54-3.02) | 1.74E-01 | 3.07E-01 |
| 127 | dhCer(d18:0/24:0) | 0.21 | (-0.38-0.8) | 4.92E-01 | 5.49E-01 | 0.98 | (0.38-1.57) | **1.25E-03** | **2.63E-03** | -0.54 | (-1.2-0.18) | 1.36E-01 | 2.10E-01 | -0.03 | (-1.63-1.6) | 9.71E-01 | 9.87E-01 |
| 128 | dhCer(d18:0/24:1) | 1.70 | (0.99-2.41) | **3.01E-06** | **1.36E-05** | 1.49 | (0.78-2.2) | **4.22E-05** | **1.11E-04** | -0.27 | (-1.09-0.63) | 5.47E-01 | 6.37E-01 | 1.59 | (-0.43-3.66) | 1.24E-01 | 2.33E-01 |
| 129 | Cer(d16:1/22:0) | -0.73 | (-1.33--0.13) | **1.79E-02** | **3.18E-02** | 1.20 | (0.56-1.84) | **2.31E-04** | **5.51E-04** | -1.03 | (-1.75--0.25) | **1.07E-02** | **2.19E-02** | 2.30 | (0.33-4.31) | **2.24E-02** | 6.84E-02 |
| 130 | Cer(d16:1/23:0) | -0.99 | (-1.66--0.32) | **3.93E-03** | **8.74E-03** | 0.76 | (0.1-1.42) | **2.38E-02** | **3.69E-02** | -1.07 | (-2.23-0.26) | 1.10E-01 | 1.74E-01 | 2.15 | (-0.09-4.44) | 6.04E-02 | 1.43E-01 |
| 131 | Cer(d16:1/24:0) | -2.03 | (-2.65--1.4) | **5.17E-10** | **5.77E-09** | 0.28 | (-0.34-0.91) | 3.74E-01 | 4.29E-01 | -0.79 | (-1.51--0.01) | **4.60E-02** | 8.21E-02 | 1.80 | (-0.29-3.93) | 9.09E-02 | 1.87E-01 |
| 132 | Cer(d16:1/24:1) | -0.40 | (-1.03-0.23) | 2.11E-01 | 2.70E-01 | 0.69 | (0.07-1.32) | **2.90E-02** | **4.44E-02** | -0.52 | (-1.39-0.44) | 2.75E-01 | 3.66E-01 | 1.85 | (0.02-3.71) | **4.75E-02** | 1.20E-01 |
| 133 | Cer(d17:1/22:0) | -1.27 | (-1.92--0.61) | **1.62E-04** | **5.04E-04** | 0.68 | (0.02-1.35) | **4.48E-02** | 6.57E-02 | -0.82 | (-1.72-0.18) | 1.04E-01 | 1.66E-01 | 0.95 | (-0.94-2.87) | 3.27E-01 | 4.87E-01 |
| 134 | Cer(d17:1/23:0) | -1.28 | (-1.96--0.6) | **2.35E-04** | **6.97E-04** | 0.02 | (-0.67-0.72) | 9.57E-01 | 9.69E-01 | -1.29 | (-2.61-0.26) | 9.78E-02 | 1.58E-01 | 1.83 | (-0.39-4.1) | 1.06E-01 | 2.10E-01 |
| 135 | Cer(d17:1/24:0) | -2.19 | (-2.76--1.61) | **4.35E-13** | **1.39E-11** | -0.28 | (-0.85-0.3) | 3.45E-01 | 4.00E-01 | -1.07 | (-1.78--0.28) | **8.53E-03** | **1.79E-02** | 0.10 | (-1.77-2.02) | 9.13E-01 | 9.57E-01 |
| 136 | Cer(d17:1/24:1) | -1.05 | (-1.67--0.43) | **8.90E-04** | **2.26E-03** | -0.29 | (-0.87-0.29) | 3.24E-01 | 3.81E-01 | -0.91 | (-1.72--0.03) | **4.31E-02** | 7.72E-02 | 1.03 | (-0.88-2.98) | 2.92E-01 | 4.50E-01 |
| 137 | Cer(d18:1/16:0) | -0.87 | (-1.33--0.4) | **2.78E-04** | **8.08E-04** | -0.36 | (-0.81-0.09) | 1.17E-01 | 1.56E-01 | -0.52 | (-1.09-0.1) | 9.85E-02 | 1.58E-01 | -0.45 | (-1.74-0.85) | 4.93E-01 | 6.31E-01 |
| 138 | Cer(d18:1/18:0) | 1.97 | (1.31-2.64) | **7.34E-09** | **5.29E-08** | 2.07 | (1.45-2.7) | **1.07E-10** | **6.87E-10** | -0.68 | (-1.34-0.03) | 6.05E-02 | 1.04E-01 | 1.92 | (0.19-3.69) | **2.95E-02** | 8.32E-02 |
| 139 | Cer(d18:1/20:0) | 0.23 | (-0.34-0.8) | 4.26E-01 | 4.86E-01 | 0.79 | (0.25-1.33) | **3.92E-03** | **7.41E-03** | -0.73 | (-1.36--0.05) | **3.51E-02** | 6.41E-02 | 0.96 | (-0.59-2.54) | 2.26E-01 | 3.71E-01 |
| 140 | Cer(d18:1/22:0) | -0.08 | (-0.56-0.4) | 7.34E-01 | 7.74E-01 | 0.97 | (0.48-1.47) | **1.24E-04** | **3.06E-04** | -0.09 | (-0.69-0.55) | 7.78E-01 | 8.43E-01 | 0.93 | (-0.49-2.36) | 2.00E-01 | 3.37E-01 |
| 141 | Cer(d18:1/23:0) | -0.40 | (-0.88-0.09) | 1.07E-01 | 1.50E-01 | 0.49 | (0-0.99) | **4.97E-02** | 7.21E-02 | -0.41 | (-1.17-0.41) | 3.11E-01 | 4.09E-01 | 1.11 | (-0.37-2.61) | 1.43E-01 | 2.63E-01 |
| 142 | Cer(d18:1/24:0) | -0.95 | (-1.41--0.48) | **7.18E-05** | **2.39E-04** | 0.09 | (-0.39-0.56) | 7.23E-01 | 7.62E-01 | -0.39 | (-1-0.25) | 2.23E-01 | 3.11E-01 | 0.03 | (-1.3-1.38) | 9.63E-01 | 9.85E-01 |
| 143 | Cer(d18:1/24:1) | 0.09 | (-0.44-0.62) | 7.49E-01 | 7.88E-01 | 0.06 | (-0.44-0.56) | 8.23E-01 | 8.59E-01 | -0.31 | (-0.96-0.38) | 3.67E-01 | 4.68E-01 | 0.62 | (-0.76-2.03) | 3.78E-01 | 5.29E-01 |
| 144 | Cer(d18:2/16:0) | -0.40 | (-1.02-0.23) | 2.17E-01 | 2.76E-01 | 0.17 | (-0.43-0.79) | 5.75E-01 | 6.33E-01 | 0.15 | (-0.78-1.18) | 7.60E-01 | 8.29E-01 | 0.35 | (-1.47-2.2) | 7.08E-01 | 7.93E-01 |
| 145 | Cer(d18:2/22:0) | -0.35 | (-0.91-0.21) | 2.22E-01 | 2.80E-01 | 0.39 | (-0.18-0.97) | 1.77E-01 | 2.27E-01 | -0.26 | (-0.97-0.5) | 4.89E-01 | 5.80E-01 | 0.35 | (-1.27-1.99) | 6.76E-01 | 7.65E-01 |
| 146 | Cer(d18:2/23:0) | -0.56 | (-1.16-0.03) | 6.45E-02 | 9.61E-02 | -0.11 | (-0.69-0.48) | 7.09E-01 | 7.53E-01 | -0.40 | (-1.35-0.64) | 4.35E-01 | 5.35E-01 | 1.17 | (-0.56-2.92) | 1.85E-01 | 3.18E-01 |
| 147 | Cer(d18:2/24:0) | -1.25 | (-1.79--0.69) | **1.15E-05** | **4.61E-05** | -0.68 | (-1.22--0.14) | **1.42E-02** | **2.32E-02** | -0.83 | (-1.51--0.08) | **3.00E-02** | 5.52E-02 | -0.65 | (-2.18-0.91) | 4.11E-01 | 5.58E-01 |
| 148 | Cer(d18:2/24:1) | -0.04 | (-0.61-0.54) | 8.96E-01 | 9.10E-01 | -0.32 | (-0.88-0.24) | 2.57E-01 | 3.10E-01 | -0.02 | (-0.75-0.78) | 9.64E-01 | 9.75E-01 | 0.54 | (-1.01-2.11) | 4.96E-01 | 6.33E-01 |

| | | | | | | | | | | | | | | | | | |
|---|---|---|---|---|---|---|---|---|---|---|---|---|---|---|---|---|---|
| 149 | Cer(d19:1/24:0) | -1.22 | (-2.12--0.32) | **8.01E-03** | **1.58E-02** | 1.08 | (0.23-1.93) | **1.29E-02** | **2.13E-02** | -1.23 | (-2.03--0.34) | **7.85E-03** | **1.67E-02** | 0.08 | (-2.67-2.91) | 9.55E-01 | 9.81E-01 |
| 150 | Cer(d19:1/24:1) | -0.31 | (-1.35-0.74) | 5.63E-01 | 6.17E-01 | 1.17 | (0.21-2.14) | **1.64E-02** | **2.66E-02** | -1.22 | (-2.19--0.13) | **2.94E-02** | 5.42E-02 | 1.08 | (-2.03-4.28) | 5.00E-01 | 6.34E-01 |
| 151 | Cer(m18:0/22:0) | 0.25 | (-0.53-1.04) | 5.30E-01 | 5.82E-01 | 2.95 | (2.15-3.76) | **8.19E-13** | **7.86E-12** | -1.25 | (-2.16--0.23) | **1.75E-02** | **3.42E-02** | 2.43 | (0.5-4.39) | **1.34E-02** | **4.68E-02** |
| 152 | Cer(m18:0/23:0) | 0.41 | (-0.46-1.28) | 3.58E-01 | 4.22E-01 | 2.81 | (1.92-3.7) | **6.59E-10** | **3.60E-09** | -1.33 | (-3.02-0.76) | 1.95E-01 | 2.82E-01 | 2.72 | (0.35-5.15) | **2.45E-02** | 7.30E-02 |
| 153 | Cer(m18:0/24:0) | -0.62 | (-1.4-0.17) | 1.22E-01 | 1.69E-01 | 1.77 | (1.04-2.5) | **1.94E-06** | **6.16E-06** | -1.21 | (-1.95--0.4) | **4.18E-03** | **9.24E-03** | 2.12 | (0.28-3.98) | **2.38E-02** | 7.17E-02 |
| 154 | Cer(m18:0/24:1) | 1.15 | (0.26-2.04) | **1.11E-02** | **2.08E-02** | 2.84 | (2-3.68) | **3.94E-11** | **2.74E-10** | -1.46 | (-2.42--0.37) | **9.81E-03** | **2.02E-02** | 4.35 | (2.12-6.63) | **1.28E-04** | **1.20E-03** |
| 155 | Cer(m18:1/22:0) | 0.63 | (-0.22-1.49) | 1.44E-01 | 1.96E-01 | 3.00 | (2.15-3.86) | **6.87E-12** | **5.32E-11** | -2.51 | (-3.17--1.79) | **9.94E-10** | **4.73E-09** | 1.19 | (-1.06-3.48) | 3.01E-01 | 4.58E-01 |
| 156 | Cer(m18:1/23:0) | 0.17 | (-0.73-1.08) | 7.11E-01 | 7.52E-01 | 2.64 | (1.71-3.57) | **2.81E-08** | **1.23E-07** | -3.37 | (-4.59--1.88) | **7.84E-05** | **2.27E-04** | 2.08 | (-0.45-4.68) | 1.07E-01 | 2.12E-01 |
| 157 | Cer(m18:1/24:0) | -0.42 | (-1.2-0.36) | 2.91E-01 | 3.56E-01 | 2.12 | (1.35-2.89) | **7.27E-08** | **2.96E-07** | -2.59 | (-3.23--1.88) | **1.87E-10** | **9.55E-10** | -0.03 | (-2.04-2.03) | 9.80E-01 | 9.92E-01 |
| 158 | Cer(m18:1/24:1) | 1.37 | (0.47-2.27) | **2.70E-03** | **6.11E-03** | 2.73 | (1.88-3.59) | **4.23E-10** | **2.42E-09** | -2.16 | (-2.91--1.32) | **3.30E-06** | **1.10E-05** | 1.88 | (-0.3-4.1) | 9.05E-02 | 1.87E-01 |
| 159 | GM3(d18:1/18:0) | -1.16 | (-1.7--0.62) | **2.65E-05** | **9.76E-05** | -0.70 | (-1.2--0.2) | **5.94E-03** | **1.06E-02** | -0.78 | (-1.37--0.16) | **1.53E-02** | **3.01E-02** | -1.22 | (-2.62-0.19) | 8.96E-02 | 1.86E-01 |
| 160 | GM3(d18:1/22:0) | -2.16 | (-2.77--1.54) | **2.44E-11** | **4.44E-10** | 0.01 | (-0.56-0.59) | 9.60E-01 | 9.71E-01 | 0.09 | (-0.88-1.16) | 8.64E-01 | 9.01E-01 | -0.55 | (-2.13-1.07) | 5.03E-01 | 6.36E-01 |
| 161 | GM3(d18:1/24:0) | -3.17 | (-3.78--2.56) | **5.91E-22** | **9.47E-20** | -1.92 | (-2.46--1.39) | **6.41E-12** | **5.13E-11** | -0.48 | (-1.25-0.35) | 2.45E-01 | 3.39E-01 | -2.41 | (-3.83--0.96) | **1.23E-03** | **6.55E-03** |
| 162 | GM3(d18:1/24:1) | -1.77 | (-2.36--1.17) | **1.05E-08** | **7.23E-08** | -1.28 | (-1.87--0.68) | **2.89E-05** | **7.79E-05** | -0.08 | (-0.88-0.79) | 8.48E-01 | 8.99E-01 | -1.68 | (-3.15--0.18) | **2.86E-02** | 8.18E-02 |
| 163 | HexCer(d16:1/22:0) | -1.73 | (-2.36--1.09) | **1.69E-07** | **9.90E-07** | 0.01 | (-0.64-0.67) | 9.69E-01 | 9.73E-01 | -0.73 | (-1.99-0.73) | 3.11E-01 | 4.09E-01 | -0.95 | (-2.95-1.08) | 3.54E-01 | 5.07E-01 |
| 164 | HexCer(d16:1/24:0) | -3.23 | (-3.88--2.57) | **2.16E-20** | **2.07E-18** | -0.23 | (-0.88-0.42) | 4.87E-01 | 5.45E-01 | -0.86 | (-1.98-0.41) | 1.73E-01 | 2.55E-01 | 0.12 | (-1.92-2.2) | 9.09E-01 | 9.57E-01 |
| 165 | HexCer(d18:1/16:0) | -1.32 | (-1.8--0.83) | **1.66E-07** | **9.85E-07** | -0.27 | (-0.74-0.19) | 2.49E-01 | 3.05E-01 | -0.03 | (-0.84-0.84) | 9.41E-01 | 9.57E-01 | -1.63 | (-3.01--0.24) | **2.21E-02** | 6.79E-02 |
| 166 | HexCer(d18:1/18:0) | -1.56 | (-2.15--0.97) | **3.24E-07** | **1.80E-06** | -1.40 | (-2.04--0.75) | **2.70E-05** | **7.38E-05** | 0.29 | (-0.73-1.41) | 5.96E-01 | 6.81E-01 | -1.86 | (-3.59--0.11) | **3.74E-02** | 1.00E-01 |
| 167 | HexCer(d18:1/20:0) | -1.84 | (-2.41--1.27) | **4.40E-10** | **5.02E-09** | -1.29 | (-1.81--0.76) | **2.18E-06** | **6.88E-06** | 0.06 | (-0.87-1.08) | 9.06E-01 | 9.31E-01 | -2.39 | (-4.02--0.73) | **5.05E-03** | **2.10E-02** |
| 168 | HexCer(d18:1/22:0) | -1.93 | (-2.43--1.44) | **1.05E-13** | **3.87E-12** | -0.92 | (-1.41--0.43) | **2.61E-04** | **6.17E-04** | 0.50 | (-0.26-1.31) | 2.01E-01 | 2.88E-01 | -2.35 | (-3.8--0.88) | **1.85E-03** | **9.34E-03** |
| 169 | HexCer(d18:1/24:0) | -2.60 | (-3.1--2.09) | **5.92E-22** | **9.47E-20** | -1.34 | (-1.83--0.85) | **1.19E-07** | **4.57E-07** | 0.07 | (-0.67-0.87) | 8.63E-01 | 9.01E-01 | -1.92 | (-3.37--0.46) | **1.03E-02** | **3.73E-02** |
| 170 | HexCer(d18:1/24:1) | -1.68 | (-2.24--1.13) | **5.30E-09** | **4.04E-08** | -1.23 | (-1.76--0.69) | **8.47E-06** | **2.48E-05** | 0.56 | (-0.2-1.39) | 1.50E-01 | 2.28E-01 | -1.42 | (-2.9-0.08) | 6.31E-02 | 1.48E-01 |
| 171 | Hex2Cer(d16:1/16:0) | -1.30 | (-1.92--0.67) | **5.63E-05** | **1.93E-04** | 0.64 | (0.03-1.26) | **3.93E-02** | 5.84E-02 | -1.04 | (-2.01-0.05) | 6.05E-02 | 1.04E-01 | 2.20 | (0.06-4.39) | **4.43E-02** | 1.14E-01 |
| 172 | Hex2Cer(d18:1/16:0) | -1.58 | (-2.1--1.05) | **6.78E-09** | **5.09E-08** | -1.17 | (-1.7--0.63) | **2.30E-05** | **6.34E-05** | -0.53 | (-1.21-0.2) | 1.48E-01 | 2.26E-01 | -1.45 | (-3.16-0.3) | 1.03E-01 | 2.07E-01 |
| 173 | Hex2Cer(d18:1/22:0) | -1.73 | (-2.29--1.17) | **2.30E-09** | **2.05E-08** | -1.43 | (-2--0.87) | **8.84E-07** | **2.91E-06** | -0.20 | (-1.06-0.73) | 6.55E-01 | 7.37E-01 | -2.48 | (-4.09--0.83) | **3.43E-03** | **1.57E-02** |
| 174 | Hex2Cer(d18:1/24:1) | -1.66 | (-2.25--1.07) | **4.96E-08** | **3.13E-07** | -1.91 | (-2.5--1.32) | **5.82E-10** | **3.21E-09** | -0.14 | (-0.92-0.71) | 7.40E-01 | 8.15E-01 | -2.62 | (-4.35--0.85) | **3.91E-03** | **1.73E-02** |
| 175 | Hex2Cer(d18:2/16:0) | -0.67 | (-1.21--0.12) | **1.65E-02** | **2.97E-02** | -0.31 | (-0.85-0.22) | 2.50E-01 | 3.06E-01 | -0.49 | (-1.27-0.35) | 2.44E-01 | 3.38E-01 | 0.71 | (-0.97-2.41) | 4.08E-01 | 5.58E-01 |
| 176 | Hex3Cer(d18:1/16:0) | -1.04 | (-1.45--0.62) | **1.58E-06** | **7.44E-06** | -1.03 | (-1.43--0.63) | **6.36E-07** | **2.21E-06** | -0.49 | (-1.07-0.13) | 1.21E-01 | 1.90E-01 | -0.80 | (-1.93-0.33) | 1.65E-01 | 2.97E-01 |
| 177 | Hex3Cer(d18:1/22:0) | -1.54 | (-2.14--0.93) | **9.28E-07** | **4.59E-06** | -2.07 | (-2.63--1.51) | **2.59E-12** | **2.22E-11** | 0.15 | (-0.79-1.19) | 7.65E-01 | 8.33E-01 | -1.67 | (-3.22--0.11) | **3.65E-02** | 9.91E-02 |
| 178 | Hex3Cer(d18:1/24:0) | -1.57 | (-2.15--0.99) | **1.64E-07** | **9.85E-07** | -1.90 | (-2.42--1.37) | **6.58E-12** | **5.18E-11** | -0.46 | (-1.22-0.37) | 2.68E-01 | 3.62E-01 | -2.28 | (-3.7--0.83) | **2.21E-03** | **1.09E-02** |
| 179 | Hex3Cer(d18:1/24:1) | -1.14 | (-1.69--0.58) | **6.61E-05** | **2.23E-04** | -1.42 | (-1.92--0.93) | **3.28E-08** | **1.42E-07** | -0.55 | (-1.29-0.25) | 1.73E-01 | 2.55E-01 | -1.03 | (-2.44-0.41) | 1.59E-01 | 2.90E-01 |
| 180 | SM(d17:1/14:0) | -1.88 | (-2.45--1.3) | **3.06E-10** | **3.86E-09** | -0.04 | (-0.58-0.51) | 8.94E-01 | 9.17E-01 | -0.98 | (-1.49--0.44) | **5.42E-04** | **1.36E-03** | 2.34 | (0.42-4.3) | **1.70E-02** | 5.51E-02 |
| 181 | SM(d18:0/14:0) | -0.89 | (-1.42--0.36) | **1.08E-03** | **2.70E-03** | 0.73 | (0.18-1.28) | **9.62E-03** | **1.64E-02** | -0.26 | (-0.77-0.28) | 3.36E-01 | 4.34E-01 | 1.63 | (0.05-3.24) | **4.26E-02** | 1.10E-01 |
| 182 | SM(d18:1/14:0)/SM(d16:1/16:0) | -1.03 | (-1.5--0.55) | **2.66E-05** | **9.76E-05** | 0.72 | (0.27-1.18) | **1.93E-03** | **3.86E-03** | -0.56 | (-1.05--0.03) | **3.86E-02** | 6.97E-02 | 2.11 | (0.57-3.68) | **7.37E-03** | **2.85E-02** |
| 183 | SM(d18:2/14:0) | 0.18 | (-0.33-0.69) | 4.96E-01 | 5.50E-01 | 1.49 | (1.01-1.98) | **2.12E-09** | **1.07E-08** | 0.07 | (-0.46-0.63) | 7.91E-01 | 8.51E-01 | 6.29 | (4.83-7.77) | **2.07E-16** | **9.93E-14** |
| 184 | SM(d17:1/16:0) | -1.40 | (-1.85--0.95) | **2.57E-09** | **2.25E-08** | -0.22 | (-0.65-0.22) | 3.29E-01 | 3.84E-01 | -0.98 | (-1.47--0.47) | **2.86E-04** | **7.59E-04** | -0.38 | (-1.9-1.16) | 6.27E-01 | 7.41E-01 |
| 185 | SM(d18:1/16:0) | -1.46 | (-1.99--0.92) | **1.89E-07** | **1.08E-06** | -0.89 | (-1.41--0.37) | **9.07E-04** | **1.95E-03** | -0.34 | (-0.91-0.28) | 2.74E-01 | 3.66E-01 | -1.60 | (-3.02--0.16) | **3.01E-02** | 8.45E-02 |
| 186 | SM(d18:2/16:0) | 0.08 | (-0.32-0.48) | 7.06E-01 | 7.48E-01 | 0.65 | (0.28-1.02) | **5.74E-04** | **1.26E-03** | 0.06 | (-0.44-0.6) | 8.06E-01 | 8.64E-01 | 1.21 | (0.14-2.29) | **2.69E-02** | 7.82E-02 |
| 187 | SM(34:3) | 0.20 | (-0.28-0.68) | 4.10E-01 | 4.70E-01 | 0.97 | (0.49-1.46) | **8.92E-05** | **2.27E-04** | 0.13 | (-0.42-0.72) | 6.47E-01 | 7.30E-01 | 2.92 | (1.38-4.49) | **2.12E-04** | **1.70E-03** |
| 188 | SM(d18:2/17:0) | -0.37 | (-0.83-0.1) | 1.23E-01 | 1.70E-01 | 0.32 | (-0.1-0.74) | 1.30E-01 | 1.71E-01 | -0.91 | (-1.41--0.38) | **9.15E-04** | **2.25E-03** | 1.15 | (-0.24-2.56) | 1.04E-01 | 2.08E-01 |
| 189 | SM(35:2) (b) | -1.53 | (-2.12--0.94) | **4.86E-07** | **2.56E-06** | -0.79 | (-1.37--0.2) | **8.48E-03** | **1.46E-02** | -0.68 | (-1.39-0.1) | 8.62E-02 | 1.42E-01 | -1.19 | (-2.94-0.58) | 1.85E-01 | 3.18E-01 |
| 190 | SM(d18:1/18:0)/SM(d16:1/20:0) | 0.25 | (-0.26-0.76) | 3.41E-01 | 4.06E-01 | 0.68 | (0.2-1.16) | **5.56E-03** | **1.00E-02** | -0.05 | (-0.63-0.56) | 8.71E-01 | 9.06E-01 | 0.38 | (-1.02-1.81) | 5.93E-01 | 7.19E-01 |
| 191 | SM(d18:2/18:1) | 0.37 | (-0.11-0.84) | 1.30E-01 | 1.79E-01 | 0.35 | (-0.07-0.77) | 1.01E-01 | 1.36E-01 | -0.01 | (-0.59-0.61) | 9.77E-01 | 9.79E-01 | 1.30 | (0.08-2.53) | **3.65E-02** | 9.91E-02 |
| 192 | SM(37:1) | -0.75 | (-1.3--0.2) | **7.20E-03** | **1.46E-02** | -0.09 | (-0.6-0.41) | 7.20E-01 | 7.61E-01 | -0.82 | (-1.45--0.15) | **1.76E-02** | **3.42E-02** | 0.13 | (-1.47-1.76) | 8.71E-01 | 9.29E-01 |
| 193 | SM(37:2) | -1.21 | (-1.81--0.6) | **1.07E-04** | **3.46E-04** | -1.13 | (-1.68--0.57) | **7.90E-05** | **2.02E-04** | -0.76 | (-1.35--0.12) | **2.06E-02** | **3.93E-02** | -0.55 | (-2.23-1.16) | 5.25E-01 | 6.51E-01 |
| 194 | SM(d18:1/20:0)/SM(d16:1/22:0) | -0.75 | (-1.17--0.33) | **4.82E-04** | **1.32E-03** | 0.43 | (0.04-0.83) | **2.93E-02** | **4.47E-02** | 0.28 | (-0.27-0.85) | 3.27E-01 | 4.25E-01 | 0.57 | (-0.67-1.83) | 3.67E-01 | 5.21E-01 |
| 195 | SM(d18:2/20:0) | -0.36 | (-0.86-0.14) | 1.60E-01 | 2.13E-01 | 0.11 | (-0.41-0.62) | 6.89E-01 | 7.33E-01 | 0.31 | (-0.42-1.1) | 4.15E-01 | 5.15E-01 | 0.44 | (-1.07-1.98) | 5.66E-01 | 6.94E-01 |
| 196 | SM(38:3) (a) | 0.49 | (0-0.98) | **4.88E-02** | 7.63E-02 | 0.56 | (0.14-0.98) | **8.55E-03** | **1.47E-02** | -0.52 | (-1.05-0.06) | 7.63E-02 | 1.28E-01 | 1.93 | (0.68-3.2) | **2.56E-03** | **1.24E-02** |
| 197 | SM(38:3) (b) | -0.77 | (-1.33--0.22) | **6.71E-03** | **1.37E-02** | -0.67 | (-1.28--0.07) | **3.00E-02** | **4.55E-02** | -0.07 | (-0.77-0.69) | 8.53E-01 | 9.00E-01 | 0.47 | (-1.3-2.27) | 6.02E-01 | 7.24E-01 |
| 198 | SM(d16:1/23:0)/SM(d17:1/22:0) | -1.08 | (-1.55--0.62) | **6.16E-06** | **2.62E-05** | 0.38 | (-0.05-0.81) | 8.35E-02 | 1.15E-01 | -0.37 | (-0.89-0.18) | 1.79E-01 | 2.62E-01 | 1.38 | (-0.17-2.96) | 8.16E-02 | 1.74E-01 |
| 199 | SM(d18:1/22:0)/SM(d16:1/24:0) | -0.59 | (-1.06--0.13) | **1.28E-02** | **2.38E-02** | 0.71 | (0.26-1.16) | **2.08E-03** | **4.13E-03** | 0.40 | (-0.17-1) | 1.76E-01 | 2.58E-01 | 0.43 | (-0.84-1.71) | 5.12E-01 | 6.45E-01 |
| 200 | SM(d16:1/24:1) | -0.70 | (-1.21--0.19) | **7.53E-03** | **1.51E-02** | 0.55 | (0.05-1.05) | **3.15E-02** | **4.75E-02** | -0.12 | (-0.71-0.51) | 7.00E-01 | 7.76E-01 | 2.62 | (1.12-4.15) | **6.11E-04** | **3.91E-03** |

| | | | | | | | | | | | | | | | | | |
|---|---|---|---|---|---|---|---|---|---|---|---|---|---|---|---|---|---|
| 201 | SM(d18:2/22:0) | -0.79 | (-1.25--0.33) | **7.92E-04** | **2.04E-03** | 0.01 | (-0.41-0.43) | 9.67E-01 | 9.73E-01 | 0.38 | (-0.22-1.02) | 2.17E-01 | 3.06E-01 | 0.42 | (-0.78-1.64) | 4.91E-01 | 6.31E-01 |
| 202 | SM(40:3) (a) | -0.13 | (-0.62-0.36) | 5.98E-01 | 6.48E-01 | 0.29 | (-0.19-0.77) | 2.39E-01 | 2.94E-01 | -0.44 | (-1.01-0.17) | 1.54E-01 | 2.32E-01 | 1.68 | (0.3-3.08) | **1.72E-02** | 5.52E-02 |
| 203 | SM(40:3) (b) | -0.91 | (-1.46--0.36) | **1.23E-03** | **3.06E-03** | -0.26 | (-0.79-0.27) | 3.35E-01 | 3.91E-01 | -0.89 | (-1.5--0.24) | **8.17E-03** | **1.72E-02** | 0.93 | (-0.86-2.74) | 3.11E-01 | 4.70E-01 |
| 204 | SM(41:0) | -0.42 | (-0.85-0.01) | 5.33E-02 | 8.22E-02 | 0.74 | (0.28-1.19) | **1.45E-03** | **2.96E-03** | -0.51 | (-1.39-0.45) | 2.86E-01 | 3.79E-01 | 0.37 | (-0.9-1.65) | 5.73E-01 | 7.00E-01 |
| 205 | SM(41:1) | -0.70 | (-1.12--0.28) | **1.15E-03** | **2.85E-03** | 0.39 | (-0.02-0.81) | 6.15E-02 | 8.66E-02 | -0.21 | (-0.85-0.47) | 5.36E-01 | 6.25E-01 | 0.49 | (-0.79-1.78) | 4.55E-01 | 6.00E-01 |
| 206 | SM(d17:1/24:1) | -0.87 | (-1.32--0.41) | **2.01E-04** | **6.11E-04** | -0.13 | (-0.57-0.31) | 5.57E-01 | 6.17E-01 | -0.47 | (-1.01-0.1) | 1.02E-01 | 1.63E-01 | 0.60 | (-0.74-1.96) | 3.83E-01 | 5.29E-01 |
| 207 | SM(d18:2/23:0) | -0.93 | (-1.36--0.5) | **2.33E-05** | **8.74E-05** | -0.51 | (-0.9--0.11) | **1.15E-02** | **1.91E-02** | -0.01 | (-0.68-0.7) | 9.74E-01 | 9.79E-01 | 0.72 | (-0.51-1.96) | 2.51E-01 | 4.04E-01 |
| 208 | SM(d18:1/24:0) | -1.46 | (-1.92--1.01) | **6.70E-10** | **7.31E-09** | 0.04 | (-0.38-0.47) | 8.44E-01 | 8.73E-01 | -0.25 | (-0.81-0.35) | 4.12E-01 | 5.14E-01 | -0.50 | (-1.63-0.63) | 3.83E-01 | 5.29E-01 |
| 209 | SM(d18:1/24:1) | -0.55 | (-1.08--0.01) | **4.44E-02** | 7.06E-02 | -0.10 | (-0.61-0.42) | 7.12E-01 | 7.55E-01 | 0.21 | (-0.37-0.84) | 4.83E-01 | 5.80E-01 | -0.12 | (-1.44-1.22) | 8.57E-01 | 9.18E-01 |
| 210 | SM(d18:2/24:0) | -1.48 | (-1.89--1.07) | **6.63E-12** | **1.45E-10** | -0.85 | (-1.23--0.46) | **1.89E-05** | **5.27E-05** | -0.11 | (-0.67-0.48) | 7.00E-01 | 7.76E-01 | -0.49 | (-1.53-0.56) | 3.54E-01 | 5.07E-01 |
| 211 | SM(43:1) | -1.62 | (-2.09--1.14) | **7.20E-11** | **1.19E-09** | -0.14 | (-0.59-0.31) | 5.37E-01 | 5.96E-01 | -1.04 | (-1.74--0.27) | **9.36E-03** | **1.94E-02** | -1.37 | (-2.82-0.1) | 6.74E-02 | 1.53E-01 |
| 212 | SM(44:1) | -1.59 | (-2.01--1.17) | **4.74E-13** | **1.42E-11** | -0.38 | (-0.76-0.01) | 5.52E-02 | 7.89E-02 | -0.49 | (-1-0.04) | 7.11E-02 | 1.21E-01 | -2.17 | (-3.24--1.08) | **1.19E-04** | **1.17E-03** |
| 213 | SM(44:2) | -1.13 | (-1.63--0.63) | **1.09E-05** | **4.45E-05** | -0.45 | (-0.9-0.01) | 5.39E-02 | 7.74E-02 | -0.23 | (-0.75-0.32) | 4.07E-01 | 5.08E-01 | -1.24 | (-2.48-0.02) | 5.39E-02 | 1.31E-01 |
| 214 | SM(44:3) (a) | -1.42 | (-1.88--0.95) | **4.02E-09** | **3.21E-08** | -1.26 | (-1.69--0.84) | **1.15E-08** | **5.25E-08** | -0.41 | (-0.9-0.11) | 1.18E-01 | 1.85E-01 | -0.55 | (-1.66-0.56) | 3.30E-01 | 4.88E-01 |
| 215 | LPC(14:0) [sn2] | -0.72 | (-1.5-0.06) | 6.98E-02 | 1.02E-01 | 1.18 | (0.41-1.96) | **2.81E-03** | **5.40E-03** | 1.75 | (1.25-2.26) | **6.35E-13** | **4.36E-12** | 4.86 | (2.45-7.33) | **7.65E-05** | **8.95E-04** |
| 216 | LPC(14:0) [sn1] | -0.42 | (-1.17-0.34) | 2.76E-01 | 3.40E-01 | 1.53 | (0.78-2.3) | **7.13E-05** | **1.84E-04** | 1.62 | (1.14-2.11) | **5.17E-12** | **3.06E-11** | 4.70 | (2.35-7.09) | **8.28E-05** | **9.46E-04** |
| 217 | LPC(15:0) [sn2] | -1.50 | (-2.11--0.88) | **2.82E-06** | **1.29E-05** | -0.45 | (-1.04-0.14) | 1.37E-01 | 1.79E-01 | 1.04 | (0.46-1.66) | **3.62E-04** | **9.45E-04** | 1.83 | (-0.4-4.11) | 1.09E-01 | 2.13E-01 |
| 218 | LPC(15:0) [sn1] | -1.28 | (-1.83--0.72) | **8.57E-06** | **3.58E-05** | -0.43 | (-0.97-0.12) | 1.27E-01 | 1.67E-01 | 0.76 | (0.24-1.32) | **4.17E-03** | **9.24E-03** | 1.40 | (-0.68-3.52) | 1.87E-01 | 3.18E-01 |
| 219 | LPC(16:0) [sn2] | -0.01 | (-0.46-0.43) | 9.50E-01 | 9.56E-01 | -0.61 | (-1.03--0.19) | **4.76E-03** | **8.73E-03** | 1.73 | (1.26-2.21) | **7.55E-14** | **5.75E-13** | 0.68 | (-0.95-2.34) | 4.13E-01 | 5.58E-01 |
| 220 | LPC(16:0) [sn1] | 0.26 | (-0.27-0.8) | 3.31E-01 | 3.96E-01 | -0.69 | (-1.19--0.19) | **7.24E-03** | **1.26E-02** | 1.95 | (1.38-2.55) | **2.32E-12** | **1.53E-11** | 0.54 | (-1.35-2.47) | 5.78E-01 | 7.05E-01 |
| 221 | LPC(16:1) [sn2] | 0.17 | (-0.55-0.9) | 6.35E-01 | 6.82E-01 | 0.95 | (0.37-1.53) | **1.29E-03** | **2.69E-03** | 4.00 | (3.27-4.78) | **3.37E-31** | **8.10E-29** | 5.09 | (3.19-7.03) | **1.66E-07** | **1.14E-05** |
| 222 | LPC(16:1) [sn1] | 0.34 | (-0.29-0.97) | 2.93E-01 | 3.56E-01 | 0.87 | (0.34-1.4) | **1.33E-03** | **2.76E-03** | 3.89 | (3.2-4.61) | **6.09E-33** | **2.92E-30** | 3.93 | (2.1-5.79) | **2.67E-05** | **4.62E-04** |
| 223 | LPC(15-MHDA) [sn2] | -1.34 | (-2.09--0.59) | **5.17E-04** | **1.39E-03** | -1.80 | (-2.53--1.05) | **2.97E-06** | **9.21E-06** | 0.88 | (0-1.85) | 5.11E-02 | 9.02E-02 | -0.29 | (-2.99-2.49) | 8.37E-01 | 8.99E-01 |
| 224 | LPC(15-MHDA) [sn1] / LPC(17:0) [sn2] | -1.22 | (-1.86--0.59) | **1.83E-04** | **5.61E-04** | -1.69 | (-2.31--1.07) | **1.33E-07** | **5.00E-07** | 0.53 | (-0.15-1.25) | 1.30E-01 | 2.01E-01 | -0.87 | (-3.13-1.44) | 4.56E-01 | 6.00E-01 |
| 225 | LPC(17:0) [sn1] | -0.89 | (-1.39--0.39) | **5.04E-04** | **1.36E-03** | -1.40 | (-1.9--0.89) | **8.98E-08** | **3.53E-07** | 0.17 | (-0.3-0.67) | 4.88E-01 | 5.80E-01 | -0.97 | (-2.82-0.91) | 3.09E-01 | 4.67E-01 |
| 226 | LPC(17:1) [sn1] (a) / LPC(17:1) [sn2] (b) | -0.95 | (-1.52--0.37) | **1.28E-03** | **3.14E-03** | -0.68 | (-1.27--0.08) | **2.64E-02** | **4.06E-02** | 1.72 | (1.09-2.4) | **3.10E-08** | **1.29E-07** | 0.85 | (-1.18-2.93) | 4.14E-01 | 5.58E-01 |
| 227 | LPC(18:0) [sn2] | -0.05 | (-0.52-0.42) | 8.27E-01 | 8.53E-01 | -0.52 | (-1--0.04) | **3.29E-02** | **4.94E-02** | 0.99 | (0.53-1.46) | **1.70E-05** | **5.36E-05** | 0.72 | (-1.04-2.51) | 4.23E-01 | 5.69E-01 |
| 228 | LPC(18:0) [sn1] | -0.03 | (-0.51-0.44) | 8.92E-01 | 9.08E-01 | -0.55 | (-1.05--0.05) | **3.14E-02** | **4.75E-02** | 0.99 | (0.52-1.48) | **2.53E-05** | **7.78E-05** | 0.77 | (-1.17-2.74) | 4.39E-01 | 5.82E-01 |
| 229 | LPC(18:1) [sn2] | -0.59 | (-1.11--0.07) | **2.69E-02** | **4.57E-02** | -1.43 | (-2--0.86) | **1.28E-06** | **4.15E-06** | 3.61 | (2.91-4.34) | **4.61E-28** | **5.53E-26** | -0.89 | (-2.69-0.94) | 3.36E-01 | 4.95E-01 |
| 230 | LPC(18:1) [sn1] | -0.50 | (-0.97--0.04) | **3.47E-02** | 5.77E-02 | -1.38 | (-1.88--0.87) | **1.29E-07** | **4.88E-07** | 3.36 | (2.74-4.01) | **4.86E-30** | **7.77E-28** | -1.16 | (-2.75-0.46) | 1.60E-01 | 2.91E-01 |
| 231 | LPC(18:2) [sn2] | -0.64 | (-1.13--0.15) | **1.04E-02** | **1.97E-02** | -2.17 | (-2.7--1.63) | **2.57E-14** | **3.16E-13** | 2.93 | (2.22-3.69) | **4.36E-18** | **6.75E-17** | -1.10 | (-2.7-0.54) | 1.86E-01 | 3.18E-01 |
| 232 | LPC(18:2) [sn1] | -0.41 | (-0.96-0.14) | 1.45E-01 | 1.96E-01 | -2.32 | (-2.91--1.74) | **5.71E-14** | **6.85E-13** | 2.94 | (2.22-3.7) | **9.01E-18** | **1.35E-16** | -1.59 | (-3.34-0.19) | 8.00E-02 | 1.73E-01 |
| 233 | LPC(18:3) [sn2] (a) | -1.54 | (-2.23--0.85) | **1.68E-05** | **6.51E-05** | -0.86 | (-1.59--0.12) | **2.29E-02** | **3.56E-02** | 2.31 | (1.65-3.01) | **5.03E-13** | **3.55E-12** | 0.51 | (-1.64-2.71) | 6.43E-01 | 7.48E-01 |
| 234 | LPC(18:3) [sn1] (a)/LPC(18:3) [sn2] (b) | -0.95 | (-1.67--0.23) | **9.74E-03** | **1.88E-02** | -0.22 | (-0.97-0.54) | 5.67E-01 | 6.27E-01 | 4.43 | (3.51-5.42) | **1.76E-25** | **1.41E-23** | 1.18 | (-1.15-3.57) | 3.23E-01 | 4.84E-01 |
| 235 | LPC(18:3) (a) [sn1] [104_sn1] | -0.94 | (-1.68--0.2) | **1.34E-02** | **2.43E-02** | -0.40 | (-1.16-0.36) | 3.02E-01 | 3.59E-01 | 4.09 | (3.21-5.02) | **1.27E-23** | **6.77E-22** | 1.30 | (-1.04-3.69) | 2.77E-01 | 4.39E-01 |
| 236 | LPC(19:0) [sn1] (a) / LPC(19:0) [sn2] (b) | -1.56 | (-2.02--1.09) | **1.61E-10** | **2.49E-09** | -2.28 | (-2.79--1.76) | **6.69E-17** | **1.61E-15** | -0.24 | (-0.87-0.42) | 4.64E-01 | 5.64E-01 | -2.26 | (-3.96--0.53) | **1.08E-02** | **3.85E-02** |
| 237 | LPC(20:0) [sn1] | -1.59 | (-2.03--1.14) | **1.26E-11** | **2.64E-10** | -2.10 | (-2.58--1.62) | **1.05E-16** | **2.41E-15** | 0.30 | (-0.12-0.74) | 1.68E-01 | 2.51E-01 | -3.35 | (-4.85--1.82) | **2.48E-05** | **4.62E-04** |
| 238 | LPC(20:1) [sn2] | -0.60 | (-1.07--0.14) | **1.07E-02** | **2.02E-02** | -1.58 | (-2.1--1.06) | **5.88E-09** | **2.79E-08** | 1.44 | (0.92-1.98) | **1.54E-08** | **6.60E-08** | -1.99 | (-3.63--0.33) | **1.93E-02** | 6.09E-02 |
| 239 | LPC(20:1) [sn1] | -0.62 | (-1.08--0.15) | **9.49E-03** | **1.84E-02** | -1.86 | (-2.38--1.33) | **1.33E-11** | **9.82E-11** | 1.75 | (1.27-2.25) | **1.55E-13** | **1.12E-12** | -2.32 | (-3.93--0.7) | **5.46E-03** | **2.19E-02** |
| 240 | LPC(20:2) [sn2] | -0.91 | (-1.41--0.4) | **4.81E-04** | **1.32E-03** | -0.80 | (-1.3--0.31) | **1.50E-03** | **3.03E-03** | 3.83 | (3.05-4.65) | **3.73E-26** | **3.58E-24** | -0.47 | (-2.07-1.15) | 5.65E-01 | 6.94E-01 |
| 241 | LPC(20:2) [sn1] | -0.63 | (-1.11--0.16) | **9.14E-03** | **1.78E-02** | -0.94 | (-1.4--0.46) | **1.11E-04** | **2.77E-04** | 3.35 | (2.66-4.06) | **4.61E-25** | **3.16E-23** | -0.77 | (-2.38-0.86) | 3.50E-01 | 5.06E-01 |
| 242 | LPC(20:3) [sn2] | 1.05 | (0.4-1.7) | **1.48E-03** | **3.59E-03** | 1.19 | (0.56-1.83) | **2.39E-04** | **5.67E-04** | 2.58 | (1.91-3.3) | **1.79E-15** | **1.72E-14** | 2.63 | (0.79-4.5) | **4.98E-03** | **2.10E-02** |
| 243 | LPC(20:3) [sn1] | 1.28 | (0.59-1.97) | **2.78E-04** | **8.08E-04** | 0.91 | (0.26-1.56) | **5.99E-03** | **1.06E-02** | 3.27 | (2.53-4.05) | **7.83E-21** | **2.50E-19** | 2.43 | (0.45-4.44) | **1.61E-02** | 5.28E-02 |
| 244 | LPC(20:3) [104_sn1] | 1.42 | (0.72-2.12) | **7.46E-05** | **2.47E-04** | 0.93 | (0.28-1.58) | **4.92E-03** | **8.95E-03** | 3.22 | (2.49-3.99) | **9.49E-21** | **2.85E-19** | 2.53 | (0.51-4.6) | **1.42E-02** | **4.91E-02** |
| 245 | LPC(20:4) [sn2] | 0.85 | (0.33-1.37) | **1.45E-03** | **3.53E-03** | -0.79 | (-1.3--0.28) | **2.64E-03** | **5.13E-03** | 2.39 | (1.75-3.06) | **6.31E-15** | **5.61E-14** | 1.51 | (-0.14-3.19) | 7.23E-02 | 1.59E-01 |
| 246 | LPC(20:4) [sn1] | 1.36 | (0.79-1.93) | **3.16E-06** | **1.42E-05** | -1.16 | (-1.71--0.61) | **4.15E-05** | **1.10E-04** | 2.66 | (1.98-3.38) | **2.33E-16** | **2.60E-15** | 1.04 | (-0.76-2.87) | 2.58E-01 | 4.13E-01 |
| 247 | LPC(20:5) [sn2] | -0.04 | (-1.13-1.06) | 9.43E-01 | 9.53E-01 | -0.23 | (-1.33-0.88) | 6.78E-01 | 7.23E-01 | 0.45 | (-0.63-1.66) | 4.27E-01 | 5.27E-01 | 4.34 | (1.51-7.25) | **2.58E-03** | **1.24E-02** |
| 248 | LPC(20:5) [sn1] | 0.24 | (-0.88-1.36) | 6.78E-01 | 7.25E-01 | -0.27 | (-1.38-0.86) | 6.41E-01 | 6.91E-01 | 0.62 | (-0.47-1.83) | 2.74E-01 | 3.66E-01 | 4.41 | (1.48-7.43) | **3.09E-03** | **1.43E-02** |
| 249 | LPC(22:4) [sn2] | -0.46 | (-1.09-0.17) | 1.50E-01 | 2.02E-01 | -0.27 | (-0.86-0.32) | 3.74E-01 | 4.29E-01 | 2.50 | (1.81-3.24) | **7.33E-14** | **5.68E-13** | -0.12 | (-1.97-1.77) | 8.99E-01 | 9.51E-01 |
| 250 | LPC(22:4) [sn1] | 0.02 | (-0.59-0.64) | 9.45E-01 | 9.53E-01 | -0.55 | (-1.14-0.05) | 7.14E-02 | 9.97E-02 | 2.70 | (2.04-3.39) | **9.68E-18** | **1.37E-16** | 0.72 | (-1.19-2.67) | 4.63E-01 | 6.06E-01 |
| 251 | LPC(22:5) [sn2] (n3) | -0.98 | (-1.6--0.35) | **2.21E-03** | **5.12E-03** | -0.74 | (-1.34--0.14) | **1.67E-02** | **2.68E-02** | 1.73 | (0.89-2.63) | **2.63E-05** | **8.04E-05** | 0.81 | (-0.85-2.5) | 3.39E-01 | 4.95E-01 |
| 252 | LPC(22:5) [sn1] (n3)/LPC(22:5) [sn2] (n6) | -0.84 | (-1.42--0.26) | **4.63E-03** | **9.97E-03** | -0.82 | (-1.43--0.21) | **8.19E-03** | **1.42E-02** | 2.12 | (1.41-2.88) | **6.79E-10** | **3.29E-09** | 0.43 | (-1.27-2.16) | 6.21E-01 | 7.41E-01 |

| | | | | | | | | | | | | | | | | | |
|---|---|---|---|---|---|---|---|---|---|---|---|---|---|---|---|---|---|
| 253 | LPC(22:5) [sn1] (n6) | -0.53 | (-1.3-0.25) | 1.80E-01 | 2.35E-01 | -1.14 | (-1.82--0.47) | **9.90E-04** | **2.10E-03** | 4.48 | (3.33-5.74) | **1.81E-17** | **2.48E-16** | -0.41 | (-2.62-1.85) | 7.17E-01 | 7.98E-01 |
| 254 | LPC(22:5) (n3) [sn1] [104_sn1] | -0.74 | (-1.42--0.05) | **3.52E-02** | 5.83E-02 | -0.95 | (-1.59--0.31) | **4.00E-03** | **7.50E-03** | 1.03 | (0.27-1.84) | **7.20E-03** | **1.54E-02** | 0.80 | (-0.97-2.6) | 3.79E-01 | 5.29E-01 |
| 255 | LPC(22:6) [sn2] | -0.66 | (-1.28--0.03) | **3.88E-02** | 6.33E-02 | -1.09 | (-1.7--0.47) | **5.66E-04** | **1.25E-03** | 1.62 | (0.94-2.33) | **1.13E-06** | **3.92E-06** | 0.47 | (-1.44-2.43) | 6.30E-01 | 7.41E-01 |
| 256 | LPC(22:6) [sn1] | -0.13 | (-0.79-0.53) | 6.95E-01 | 7.40E-01 | -1.17 | (-1.8--0.53) | **3.39E-04** | **7.95E-04** | 2.08 | (1.36-2.84) | **2.30E-09** | **1.04E-08** | 0.38 | (-1.62-2.43) | 7.09E-01 | 7.93E-01 |
| 257 | LPC(O-16:0) | -0.32 | (-0.76-0.13) | 1.63E-01 | 2.17E-01 | -0.94 | (-1.4--0.47) | **9.91E-05** | **2.49E-04** | 1.73 | (1.23-2.26) | **3.01E-12** | **1.87E-11** | 0.40 | (-1.39-2.22) | 6.66E-01 | 7.58E-01 |
| 258 | LPC(O-18:0) | -1.05 | (-1.59--0.5) | **1.83E-04** | **5.61E-04** | -1.53 | (-2.07--0.99) | **5.02E-08** | **2.13E-07** | 0.28 | (-0.28-0.88) | 3.35E-01 | 4.33E-01 | -1.09 | (-3.06-0.93) | 2.87E-01 | 4.45E-01 |
| 259 | LPC(O-18:1) | 0.52 | (0.07-0.97) | **2.38E-02** | **4.13E-02** | -0.80 | (-1.25--0.35) | **5.53E-04** | **1.23E-03** | 1.82 | (1.3-2.37) | **7.59E-13** | **5.13E-12** | -0.28 | (-2.05-1.54) | 7.63E-01 | 8.42E-01 |
| 260 | LPC(O-20:0) | -1.70 | (-2.21--1.19) | **1.86E-10** | **2.70E-09** | -2.23 | (-2.77--1.69) | **4.19E-15** | **6.29E-14** | 0.04 | (-0.78-0.93) | 9.25E-01 | 9.42E-01 | -2.32 | (-4.14--0.47) | **1.46E-02** | **4.96E-02** |
| 261 | LPC(O-20:1) | 0.35 | (-0.5-1.21) | 4.19E-01 | 4.80E-01 | -0.87 | (-1.46--0.26) | **4.90E-03** | **8.94E-03** | 2.68 | (1.69-3.77) | **1.85E-08** | **7.79E-08** | 0.12 | (-2.19-2.48) | 9.22E-01 | 9.64E-01 |
| 262 | LPC(O-22:0) | -2.40 | (-2.88--1.91) | **1.69E-20** | **2.02E-18** | -1.43 | (-1.85--1) | **1.03E-10** | **6.79E-10** | 0.22 | (-0.26-0.72) | 3.73E-01 | 4.73E-01 | -1.92 | (-3.1--0.72) | **1.79E-03** | **9.15E-03** |
| 263 | LPC(O-22:1) | -1.62 | (-2.15--1.1) | **2.73E-09** | **2.33E-08** | -2.10 | (-2.63--1.56) | **9.90E-14** | **1.16E-12** | 1.29 | (0.72-1.9) | **6.07E-06** | **1.97E-05** | -1.92 | (-3.6--0.2) | **2.86E-02** | 8.18E-02 |
| 264 | LPC(O-24:0) | -3.04 | (-3.62--2.45) | **1.56E-22** | **7.48E-20** | -1.17 | (-1.62--0.71) | **7.24E-07** | **2.45E-06** | -0.48 | (-0.98-0.06) | 7.85E-02 | 1.31E-01 | -1.49 | (-2.7--0.26) | **1.81E-02** | 5.76E-02 |
| 265 | LPC(O-24:1) | -2.43 | (-2.98--1.87) | **1.19E-16** | **8.16E-15** | -1.37 | (-1.84--0.9) | **1.75E-08** | **7.86E-08** | 0.47 | (-0.04-1.01) | 7.26E-02 | 1.23E-01 | -0.57 | (-1.98-0.86) | 4.30E-01 | 5.74E-01 |
| 266 | LPC(O-24:2) | -1.94 | (-2.53--1.34) | **3.86E-10** | **4.64E-09** | -2.07 | (-2.6--1.53) | **1.68E-13** | **1.79E-12** | 0.20 | (-0.32-0.75) | 4.61E-01 | 5.61E-01 | -1.62 | (-3.28-0.07) | 6.00E-02 | 1.43E-01 |
| 267 | LPC(P-16:0) | 0.43 | (-0.05-0.9) | 7.81E-02 | 1.12E-01 | -1.04 | (-1.51--0.56) | **2.15E-05** | **5.96E-05** | 1.13 | (0.65-1.63) | **2.63E-06** | **8.83E-06** | 0.40 | (-1.33-2.16) | 6.53E-01 | 7.51E-01 |
| 268 | LPC(P-18:0) | 0.41 | (-0.04-0.87) | 7.48E-02 | 1.09E-01 | -0.79 | (-1.24--0.34) | **6.48E-04** | **1.41E-03** | 1.76 | (1.24-2.29) | **2.96E-12** | **1.87E-11** | -0.43 | (-2.19-1.36) | 6.34E-01 | 7.42E-01 |
| 269 | LPC(P-18:1) | 0.95 | (0.36-1.54) | **1.57E-03** | **3.78E-03** | -0.88 | (-1.4--0.35) | **1.14E-03** | **2.40E-03** | 1.25 | (0.59-1.95) | **1.32E-04** | **3.64E-04** | 0.83 | (-1.2-2.9) | 4.25E-01 | 5.70E-01 |
| 270 | LPC(P-20:0) | -0.73 | (-1.3--0.15) | **1.34E-02** | **2.43E-02** | -1.56 | (-2.13--0.99) | **1.23E-07** | **4.68E-07** | 2.02 | (1.36-2.71) | **3.08E-10** | **1.54E-09** | -1.83 | (-3.76-0.15) | 6.94E-02 | 1.56E-01 |
| 271 | LPE(16:0) [sn2] | -0.58 | (-1.09--0.08) | **2.45E-02** | **4.23E-02** | -1.40 | (-1.94--0.85) | **8.49E-07** | **2.81E-06** | 2.24 | (1.6-2.92) | **6.15E-13** | **4.28E-12** | -1.78 | (-3.44--0.09) | **3.95E-02** | 1.05E-01 |
| 272 | LPE(16:0) [sn1] | -0.47 | (-0.95-0.01) | 5.74E-02 | 8.74E-02 | -1.36 | (-1.89--0.84) | **4.93E-07** | **1.73E-06** | 2.24 | (1.64-2.88) | **1.97E-14** | **1.60E-13** | -1.41 | (-2.97-0.19) | 8.30E-02 | 1.76E-01 |
| 273 | LPE(18:0) [sn2] | 0.18 | (-0.3-0.66) | 4.70E-01 | 5.28E-01 | -0.80 | (-1.27--0.33) | **9.36E-04** | **2.01E-03** | 1.40 | (0.87-1.96) | **9.59E-08** | **3.71E-07** | -1.19 | (-2.65-0.28) | 1.12E-01 | 2.14E-01 |
| 274 | LPE(18:0) [sn1] | 0.40 | (-0.05-0.85) | 7.97E-02 | 1.14E-01 | -0.85 | (-1.33--0.37) | **5.55E-04** | **1.23E-03** | 1.56 | (1.03-2.12) | **2.13E-09** | **9.83E-09** | -0.83 | (-2.34-0.7) | 2.86E-01 | 4.45E-01 |
| 275 | LPE(18:1) [sn2] | -0.78 | (-1.33--0.24) | **5.15E-03** | **1.10E-02** | -0.57 | (-1.23-0.09) | 8.95E-02 | 1.22E-01 | 2.54 | (1.88-3.24) | **1.28E-15** | **1.28E-14** | -1.28 | (-3.21-0.69) | 2.01E-01 | 3.37E-01 |
| 276 | LPE(18:1) [sn1] | -0.74 | (-1.28--0.2) | **7.22E-03** | **1.46E-02** | -0.63 | (-1.3-0.04) | 6.44E-02 | 9.03E-02 | 2.94 | (2.26-3.65) | **1.02E-19** | **2.33E-18** | -0.98 | (-2.95-1.04) | 3.39E-01 | 4.95E-01 |
| 277 | LPE(18:2) [sn2] | -0.93 | (-1.59--0.27) | **6.21E-03** | **1.29E-02** | -0.58 | (-1.27-0.12) | 1.03E-01 | 1.38E-01 | 3.65 | (2.71-4.66) | **9.44E-17** | **1.10E-15** | -1.63 | (-3.55-0.33) | 1.03E-01 | 2.07E-01 |
| 278 | LPE(18:2) [sn1] | -0.96 | (-1.63--0.29) | **5.28E-03** | **1.12E-02** | -0.59 | (-1.27-0.09) | 8.83E-02 | 1.21E-01 | 3.74 | (2.85-4.69) | **1.99E-19** | **4.16E-18** | -1.59 | (-3.53-0.38) | 1.12E-01 | 2.14E-01 |
| 279 | LPE(20:4) [sn1] | 0.57 | (0.03-1.11) | **3.84E-02** | 6.29E-02 | -0.63 | (-1.14--0.11) | **1.73E-02** | **2.76E-02** | 3.15 | (2.41-3.95) | **4.99E-19** | **9.21E-18** | -1.23 | (-2.61-0.18) | 8.67E-02 | 1.82E-01 |
| 280 | LPE(22:6) [sn2] | -0.37 | (-0.96-0.22) | 2.19E-01 | 2.77E-01 | -0.37 | (-0.97-0.22) | 2.16E-01 | 2.73E-01 | 2.12 | (1.4-2.88) | **9.96E-10** | **4.73E-09** | -0.60 | (-2.39-1.23) | 5.19E-01 | 6.50E-01 |
| 281 | LPE(22:6) [sn1] | -0.32 | (-0.95-0.31) | 3.16E-01 | 3.83E-01 | -0.51 | (-1.08-0.06) | 8.10E-02 | 1.12E-01 | 2.35 | (1.63-3.11) | **9.70E-12** | **5.68E-11** | -0.45 | (-2.17-1.29) | 6.09E-01 | 7.29E-01 |
| 282 | PC(28:0) | -2.72 | (-3.8--1.63) | **1.51E-06** | **7.19E-06** | 0.17 | (-0.93-1.29) | 7.62E-01 | 8.00E-01 | -0.29 | (-0.88-0.34) | 3.63E-01 | 4.65E-01 | 2.70 | (-0.18-5.66) | 6.63E-02 | 1.52E-01 |
| 283 | PC(14:0_16:0) | -2.18 | (-2.89--1.47) | **4.19E-09** | **3.29E-08** | 0.08 | (-0.63-0.79) | 8.31E-01 | 8.63E-01 | -0.19 | (-0.85-0.52) | 5.89E-01 | 6.74E-01 | 0.35 | (-1.54-2.28) | 7.18E-01 | 7.98E-01 |
| 284 | PC(31:0) (a) | -3.27 | (-4.32--2.21) | **3.45E-09** | **2.85E-08** | 0.93 | (-0.13-2) | 8.46E-02 | 1.16E-01 | -0.55 | (-1.4-0.4) | 2.47E-01 | 3.40E-01 | 1.80 | (-1.79-5.52) | 3.29E-01 | 4.88E-01 |
| 285 | PC(31:0) (b) | -2.25 | (-2.86--1.64) | **2.46E-12** | **6.57E-11** | -0.26 | (-0.91-0.39) | 4.28E-01 | 4.87E-01 | -1.00 | (-1.85--0.06) | **3.79E-02** | 6.87E-02 | -0.24 | (-2.42-1.99) | 8.32E-01 | 8.97E-01 |
| 286 | PC(31:1) | -2.56 | (-3.45--1.67) | **3.05E-08** | **2.06E-07** | 1.45 | (0.56-2.34) | **1.35E-03** | **2.79E-03** | -0.29 | (-1.06-0.54) | 4.80E-01 | 5.78E-01 | 3.29 | (0.61-6.04) | **1.58E-02** | 5.23E-02 |
| 287 | PC(16:0_16:0) | -0.68 | (-1.18--0.17) | **8.41E-03** | **1.65E-02** | -0.23 | (-0.72-0.26) | 3.50E-01 | 4.04E-01 | -0.57 | (-1.19-0.09) | 9.09E-02 | 1.49E-01 | -1.29 | (-2.54--0.03) | **4.48E-02** | 1.14E-01 |
| 288 | PC(32:1) | -0.87 | (-1.69--0.04) | **4.11E-02** | 6.65E-02 | 2.58 | (1.77-3.39) | **4.13E-10** | **2.39E-09** | 0.94 | (0.11-1.83) | **2.49E-02** | **4.70E-02** | 3.56 | (1.41-5.75) | **1.12E-03** | **6.20E-03** |
| 289 | PC(32:2) | -1.59 | (-2.26--0.92) | **4.67E-06** | **2.00E-05** | 0.85 | (0.03-1.67) | **4.18E-02** | 6.19E-02 | 0.08 | (-0.57-0.78) | 8.09E-01 | 8.65E-01 | 2.57 | (0.47-4.71) | **1.64E-02** | 5.36E-02 |
| 290 | PC(33:0) (a) | -2.28 | (-3.03--1.52) | **7.38E-09** | **5.29E-08** | 0.04 | (-0.76-0.84) | 9.31E-01 | 9.49E-01 | -0.79 | (-1.65-0.15) | 9.71E-02 | 1.58E-01 | 0.06 | (-2.65-2.83) | 9.68E-01 | 9.86E-01 |
| 291 | PC(33:0) (b) | -1.78 | (-2.34--1.22) | **1.18E-09** | **1.18E-08** | -0.47 | (-1.06-0.13) | 1.25E-01 | 1.64E-01 | -1.45 | (-2.17--0.66) | **5.09E-04** | **1.30E-03** | -1.02 | (-2.85-0.84) | 2.79E-01 | 4.39E-01 |
| 292 | PC(33:1) | -2.44 | (-3.14--1.73) | **5.10E-11** | **8.75E-10** | 0.47 | (-0.25-1.2) | 2.03E-01 | 2.57E-01 | -0.58 | (-1.18-0.07) | 7.75E-02 | 1.30E-01 | 0.03 | (-2.09-2.19) | 9.82E-01 | 9.92E-01 |
| 293 | PC(33:2) | -2.27 | (-2.89--1.64) | **6.00E-12** | **1.37E-10** | -0.81 | (-1.42--0.19) | **1.07E-02** | **1.78E-02** | -1.25 | (-1.87--0.6) | **3.06E-04** | **8.08E-04** | -0.85 | (-2.83-1.16) | 4.02E-01 | 5.53E-01 |
| 294 | PC(16:0_18:0) | -0.99 | (-1.41--0.57) | **4.55E-06** | **1.97E-05** | -0.29 | (-0.73-0.16) | 2.08E-01 | 2.63E-01 | -0.26 | (-0.82-0.33) | 3.83E-01 | 4.82E-01 | -0.81 | (-2.02-0.42) | 1.95E-01 | 3.30E-01 |
| 295 | PC(16:0_18:1) | -1.09 | (-1.64--0.53) | **1.37E-04** | **4.33E-04** | 0.44 | (-0.22-1.09) | 1.91E-01 | 2.42E-01 | 0.08 | (-0.6-0.82) | 8.21E-01 | 8.74E-01 | -0.54 | (-2.16-1.12) | 5.22E-01 | 6.50E-01 |
| 296 | PC(16:0_18:2) | -0.34 | (-0.64--0.04) | **2.86E-02** | **4.82E-02** | -0.56 | (-0.98--0.15) | **7.64E-03** | **1.33E-02** | -0.59 | (-1.39-0.29) | 1.81E-01 | 2.64E-01 | -0.98 | (-2.32-0.37) | 1.54E-01 | 2.81E-01 |
| 297 | PC(16:1_18:2) | -0.87 | (-1.5--0.23) | **7.57E-03** | **1.51E-02** | -0.04 | (-0.63-0.57) | 9.08E-01 | 9.27E-01 | -0.75 | (-1.37--0.09) | **2.77E-02** | 5.15E-02 | 1.83 | (0.15-3.54) | **3.28E-02** | 9.16E-02 |
| 298 | PC(16:0_18:3) (a) | -1.78 | (-2.43--1.13) | **1.51E-07** | **9.27E-07** | 0.34 | (-0.34-1.03) | 3.27E-01 | 3.83E-01 | 0.05 | (-0.73-0.9) | 8.95E-01 | 9.24E-01 | 0.46 | (-1.47-2.43) | 6.39E-01 | 7.45E-01 |
| 299 | PC(14:0_20:4) | -0.73 | (-1.49-0.05) | 6.64E-02 | 9.86E-02 | 1.14 | (0.28-2.01) | **9.67E-03** | **1.64E-02** | -0.15 | (-0.74-0.48) | 6.36E-01 | 7.20E-01 | 4.75 | (2.3-7.26) | **1.38E-04** | **1.25E-03** |
| 300 | PC(34:5) | -1.19 | (-2.36-0) | **4.95E-02** | 7.71E-02 | 1.64 | (0.4-2.9) | **9.61E-03** | **1.64E-02** | -1.14 | (-1.93--0.28) | **1.10E-02** | **2.23E-02** | 7.01 | (3.69-10.43) | **3.00E-05** | **4.64E-04** |
| 301 | PC(15-MHDA_18:1) | -2.51 | (-3.31--1.72) | **1.39E-09** | **1.34E-08** | -0.81 | (-1.58--0.03) | **4.30E-02** | 6.35E-02 | -1.26 | (-1.99--0.45) | **2.77E-03** | **6.37E-03** | -1.45 | (-4-1.16) | 2.72E-01 | 4.33E-01 |
| 302 | PC(17:0_18:1) | -1.99 | (-2.49--1.47) | **1.15E-13** | **3.93E-12** | -0.24 | (-0.76-0.29) | 3.78E-01 | 4.33E-01 | -1.06 | (-1.56--0.54) | **1.14E-04** | **3.25E-04** | -1.23 | (-2.85-0.43) | 1.45E-01 | 2.67E-01 |
| 303 | PC(15-MHDA_18:2) | -1.86 | (-2.67--1.05) | **9.80E-06** | **4.05E-05** | -1.20 | (-1.96--0.42) | **2.53E-03** | **4.94E-03** | -1.39 | (-2.07--0.66) | **3.55E-04** | **9.32E-04** | -1.77 | (-4.23-0.75) | 1.66E-01 | 2.99E-01 |
| 304 | PC(17:0_18:2) | -1.54 | (-2.08--1) | **3.86E-08** | **2.57E-07** | -1.17 | (-1.68--0.65) | **1.30E-05** | **3.74E-05** | -1.28 | (-1.84--0.68) | **5.78E-05** | **1.69E-04** | -2.03 | (-3.62--0.41) | **1.44E-02** | **4.92E-02** |

| | | | | | | | | | | | | | | | | | |
|---|---|---|---|---|---|---|---|---|---|---|---|---|---|---|---|---|---|
| 305 | PC(17:1_18:2) | -1.21 | (-1.75--0.67) | **1.36E-05** | **5.30E-05** | 0.46 | (-0.14-1.06) | 1.30E-01 | 1.70E-01 | -0.72 | (-1.27--0.14) | **1.54E-02** | **3.03E-02** | 1.20 | (-0.55-2.98) | 1.80E-01 | 3.14E-01 |
| 306 | PC(15:0_20:3) | -0.57 | (-0.83--0.31) | **1.77E-05** | **6.79E-05** | -0.19 | (-0.52-0.15) | 2.73E-01 | 3.29E-01 | -0.02 | (-0.98-1.04) | 9.65E-01 | 9.75E-01 | -0.51 | (-1.62-0.63) | 3.79E-01 | 5.29E-01 |
| 307 | PC(15:0_20:4) | -1.28 | (-1.97--0.58) | **3.68E-04** | **1.06E-03** | -0.54 | (-1.19-0.12) | 1.08E-01 | 1.44E-01 | -1.32 | (-1.88--0.72) | **3.80E-05** | **1.13E-04** | 1.28 | (-0.87-3.47) | 2.45E-01 | 3.97E-01 |
| 308 | PC(35:5) | -1.57 | (-2.8--0.32) | **1.38E-02** | **2.50E-02** | 0.66 | (-0.56-1.9) | 2.89E-01 | 3.45E-01 | -2.33 | (-3.15--1.42) | **4.33E-06** | **1.41E-05** | 4.17 | (0.8-7.65) | **1.51E-02** | 5.03E-02 |
| 309 | PC 36:0 | 0.97 | (0.36-1.59) | **2.00E-03** | **4.66E-03** | 3.16 | (2.47-3.84) | **7.15E-19** | **3.43E-17** | 0.25 | (-0.29-0.82) | 3.65E-01 | 4.66E-01 | 1.52 | (-0.21-3.29) | 8.57E-02 | 1.80E-01 |
| 310 | PC(18:0_18:1) | -1.34 | (-1.89--0.78) | **3.35E-06** | **1.49E-05** | 0.98 | (0.36-1.59) | **1.86E-03** | **3.74E-03** | 0.47 | (-0.1-1.07) | 1.09E-01 | 1.73E-01 | 1.07 | (-0.46-2.62) | 1.71E-01 | 3.05E-01 |
| 311 | PC(18:0_18:2) | -0.61 | (-1.15--0.08) | **2.54E-02** | **4.35E-02** | -0.06 | (-0.62-0.5) | 8.30E-01 | 8.63E-01 | -0.46 | (-1.09-0.21) | 1.76E-01 | 2.58E-01 | 0.10 | (-1.42-1.65) | 8.94E-01 | 9.48E-01 |
| 312 | PC(16:0_20:3) (a) | -1.23 | (-1.82--0.64) | **5.06E-05** | **1.75E-04** | -1.61 | (-2.18--1.04) | **5.63E-08** | **2.33E-07** | -0.39 | (-1.07-0.34) | 2.85E-01 | 3.78E-01 | -2.86 | (-4.42--1.29) | **4.47E-04** | **3.02E-03** |
| 313 | PC(16:0_20:3) (b) | 0.67 | (-0.08-1.43) | 8.02E-02 | 1.14E-01 | 1.95 | (1.24-2.66) | **7.67E-08** | **3.09E-07** | 0.32 | (-0.31-0.98) | 3.33E-01 | 4.32E-01 | 1.74 | (0.01-3.51) | **4.89E-02** | 1.23E-01 |
| 314 | PC(18:2_18:2) | -2.57 | (-3.28--1.87) | **3.69E-12** | **9.32E-11** | -2.49 | (-3.22--1.75) | **1.24E-10** | **7.80E-10** | 0.36 | (-0.55-1.37) | 4.49E-01 | 5.50E-01 | -3.04 | (-5.24--0.78) | **8.73E-03** | **3.24E-02** |
| 315 | PC(16:1_20:4) | -0.42 | (-1.09-0.27) | 2.30E-01 | 2.89E-01 | 0.56 | (-0.07-1.2) | 8.11E-02 | 1.12E-01 | 0.04 | (-0.59-0.7) | 9.13E-01 | 9.34E-01 | 3.69 | (1.89-5.53) | **5.90E-05** | **7.86E-04** |
| 316 | PC(16:0_20:5) | -0.26 | (-1.46-0.96) | 6.74E-01 | 7.22E-01 | 0.55 | (-0.59-1.69) | 3.45E-01 | 4.00E-01 | -1.86 | (-2.69--0.94) | **1.83E-04** | **4.92E-04** | 3.86 | (0.8-7.01) | **1.32E-02** | **4.65E-02** |
| 317 | PC(36:6) | -2.22 | (-3.03--1.4) | **1.62E-07** | **9.83E-07** | 0.53 | (-0.37-1.43) | 2.52E-01 | 3.06E-01 | -0.83 | (-1.42--0.2) | **1.07E-02** | **2.19E-02** | 4.41 | (1.91-6.97) | **5.17E-04** | **3.45E-03** |
| 318 | PC(15-MHDA_20:4) | -0.68 | (-1.46-0.11) | 9.03E-02 | 1.28E-01 | -0.47 | (-1.22-0.29) | 2.23E-01 | 2.78E-01 | -1.62 | (-2.28--0.91) | **2.18E-05** | **6.78E-05** | 0.08 | (-2.33-2.55) | 9.47E-01 | 9.76E-01 |
| 319 | PC(17:0_20:4) | -0.33 | (-0.97-0.32) | 3.20E-01 | 3.85E-01 | -0.68 | (-1.27--0.09) | **2.42E-02** | **3.73E-02** | -1.70 | (-2.2--1.18) | **4.15E-09** | **1.86E-08** | -0.10 | (-1.92-1.75) | 9.13E-01 | 9.57E-01 |
| 320 | PC(15:0_22:6) | -2.68 | (-3.42--1.94) | **3.93E-12** | **9.43E-11** | -0.93 | (-1.62--0.24) | **8.87E-03** | **1.52E-02** | -1.88 | (-2.45--1.28) | **1.58E-08** | **6.70E-08** | 0.59 | (-1.64-2.87) | 6.05E-01 | 7.26E-01 |
| 321 | PC(38:2) | -1.13 | (-1.58--0.67) | **1.45E-06** | **6.94E-06** | 0.67 | (0.19-1.14) | **5.76E-03** | **1.03E-02** | 0.92 | (0.32-1.55) | **2.19E-03** | **5.11E-03** | 1.05 | (-0.09-2.21) | 7.19E-02 | 1.59E-01 |
| 322 | PC(18:0_20:3) | 1.23 | (0.54-1.93) | **5.29E-04** | **1.41E-03** | 3.69 | (2.93-4.46) | **1.54E-20** | **1.06E-18** | 0.12 | (-0.44-0.71) | 6.79E-01 | 7.58E-01 | 5.35 | (3.52-7.21) | **1.22E-08** | **1.46E-06** |
| 323 | PC(18:1_20:3) | -0.59 | (-1.2-0.03) | 6.21E-02 | 9.35E-02 | 0.96 | (0.37-1.55) | **1.39E-03** | **2.85E-03** | 0.23 | (-0.38-0.88) | 4.60E-01 | 5.61E-01 | 0.85 | (-0.73-2.46) | 2.91E-01 | 4.50E-01 |
| 324 | PC(38:4) (b) | -0.97 | (-1.66--0.28) | **5.93E-03** | **1.24E-02** | 1.29 | (0.57-2.02) | **4.90E-04** | **1.11E-03** | -0.11 | (-0.83-0.67) | 7.73E-01 | 8.40E-01 | 1.67 | (-0.2-3.57) | 8.07E-02 | 1.74E-01 |
| 325 | PC(18:0_20:4) | 0.98 | (0.35-1.61) | **2.29E-03** | **5.29E-03** | 0.43 | (-0.17-1.03) | 1.56E-01 | 2.01E-01 | -1.07 | (-1.65--0.46) | **8.92E-04** | **2.21E-03** | 1.69 | (-0.04-3.44) | 5.52E-02 | 1.34E-01 |
| 326 | PC(38:5) (a) | -1.09 | (-1.59--0.58) | **2.73E-05** | **9.86E-05** | -0.22 | (-0.71-0.28) | 3.89E-01 | 4.45E-01 | -1.17 | (-1.68--0.63) | **3.91E-05** | **1.15E-04** | 0.06 | (-1.45-1.59) | 9.39E-01 | 9.74E-01 |
| 327 | PC(38:5) (b) | -1.24 | (-1.97--0.51) | **9.15E-04** | **2.31E-03** | 0.54 | (-0.23-1.31) | 1.72E-01 | 2.20E-01 | 0.11 | (-0.54-0.82) | 7.40E-01 | 8.15E-01 | 1.61 | (-0.44-3.69) | 1.23E-01 | 2.33E-01 |
| 328 | PC(38:6) (a) | -1.01 | (-1.61--0.42) | **9.39E-04** | **2.36E-03** | -1.09 | (-1.67--0.5) | **2.90E-04** | **6.81E-04** | 0.02 | (-0.69-0.79) | 9.50E-01 | 9.64E-01 | 0.16 | (-1.65-1.99) | 8.64E-01 | 9.24E-01 |
| 329 | PC(16:0_22:6) | -1.01 | (-1.64--0.37) | **1.94E-03** | **4.55E-03** | -0.37 | (-1.07-0.34) | 3.10E-01 | 3.67E-01 | -1.25 | (-1.93--0.5) | **1.40E-03** | **3.31E-03** | 0.13 | (-1.97-2.26) | 9.07E-01 | 9.57E-01 |
| 330 | PC(18:2_20:5) | -1.76 | (-2.73--0.78) | **4.83E-04** | **1.32E-03** | -0.89 | (-1.88-0.12) | 8.29E-02 | 1.15E-01 | -0.64 | (-1.41-0.21) | 1.36E-01 | 2.09E-01 | 2.51 | (-0.16-5.25) | 6.60E-02 | 1.52E-01 |
| 331 | PC(16:1_22:6) | -1.64 | (-2.33--0.96) | **3.49E-06** | **1.54E-05** | -0.66 | (-1.32-0.01) | 5.51E-02 | 7.89E-02 | -1.04 | (-1.56--0.48) | **4.09E-04** | **1.06E-03** | 1.62 | (-0.35-3.62) | 1.08E-01 | 2.12E-01 |
| 332 | PC(38:7)(c) | -1.58 | (-2.17--0.99) | **2.34E-07** | **1.32E-06** | -0.64 | (-1.21--0.06) | **2.93E-02** | **4.47E-02** | -0.88 | (-1.37--0.37) | **1.02E-03** | **2.45E-03** | -0.03 | (-1.69-1.66) | 9.73E-01 | 9.88E-01 |
| 333 | PC(39:5)(a) | -2.53 | (-3.35--1.7) | **4.56E-09** | **3.53E-08** | -1.82 | (-2.59--1.05) | **4.96E-06** | **1.51E-05** | -1.98 | (-2.62--1.29) | **2.49E-07** | **9.28E-07** | -0.54 | (-2.96-1.95) | 6.68E-01 | 7.58E-01 |
| 334 | PC(39:5)(b) | -2.31 | (-3.04--1.58) | **1.68E-09** | **1.56E-08** | -0.86 | (-1.58--0.13) | **2.03E-02** | **3.21E-02** | -1.95 | (-2.55--1.29) | **7.31E-08** | **2.90E-07** | -0.94 | (-3.24-1.42) | 4.31E-01 | 5.75E-01 |
| 335 | PC(15-MHDA_22:6) | -2.51 | (-3.31--1.7) | **2.77E-09** | **2.33E-08** | -1.62 | (-2.33--0.91) | **1.10E-05** | **3.17E-05** | -2.07 | (-2.73--1.36) | **1.67E-07** | **6.31E-07** | -1.16 | (-3.44-1.17) | 3.25E-01 | 4.86E-01 |
| 336 | PC(17:0_22:6) | -2.00 | (-2.74--1.26) | **1.87E-07** | **1.08E-06** | -1.05 | (-1.79--0.31) | **5.44E-03** | **9.82E-03** | -2.24 | (-2.78--1.65) | **1.65E-11** | **9.21E-11** | -0.48 | (-2.73-1.83) | 6.82E-01 | 7.69E-01 |
| 337 | PC(18:0_22:5) (n6) | -1.32 | (-2.07--0.57) | **6.32E-04** | **1.67E-03** | 1.17 | (0.37-1.97) | **3.94E-03** | **7.42E-03** | -1.24 | (-2.36-0.06) | 6.01E-02 | 1.04E-01 | 2.25 | (0.3-4.25) | **2.39E-02** | 7.17E-02 |
| 338 | PC(18:0_22:6) | -0.61 | (-1.32-0.1) | 9.21E-02 | 1.30E-01 | 1.19 | (0.44-1.95) | **1.98E-03** | **3.94E-03** | -1.44 | (-2.09--0.74) | **1.21E-04** | **3.40E-04** | 3.01 | (0.9-5.18) | **5.22E-03** | **2.12E-02** |
| 339 | PC(40:7) (a) | -1.36 | (-1.99--0.73) | **2.64E-05** | **9.76E-05** | 0.02 | (-0.65-0.69) | 9.56E-01 | 9.69E-01 | 0.94 | (0-1.97) | **4.91E-02** | 8.72E-02 | 1.35 | (-0.64-3.39) | 1.84E-01 | 3.18E-01 |
| 340 | PC(18:1_22:6) (a) | -1.70 | (-2.24--1.16) | **1.69E-09** | **1.56E-08** | -0.86 | (-1.44--0.27) | **4.10E-03** | **7.62E-03** | -0.45 | (-1.13-0.27) | 2.15E-01 | 3.04E-01 | -1.64 | (-3.29-0.03) | 5.37E-02 | 1.31E-01 |
| 341 | PC(40:8) | -1.77 | (-2.3--1.24) | **1.86E-10** | **2.70E-09** | -1.88 | (-2.39--1.37) | **2.15E-12** | **1.91E-11** | 0.83 | (0.2-1.5) | **9.19E-03** | **1.92E-02** | -0.97 | (-2.47-0.55) | 2.09E-01 | 3.48E-01 |
| 342 | PC(44:12) | -2.40 | (-3.44--1.34) | **1.14E-05** | **4.60E-05** | -2.61 | (-3.74--1.46) | **1.06E-05** | **3.09E-05** | -1.02 | (-1.76--0.22) | **1.39E-02** | **2.75E-02** | -0.64 | (-3.66-2.47) | 6.82E-01 | 7.69E-01 |
| 343 | PC(O-16:0/16:0) | -0.78 | (-1.21--0.35) | **3.82E-04** | **1.09E-03** | -0.75 | (-1.18--0.33) | **5.02E-04** | **1.13E-03** | 0.45 | (-0.19-1.14) | 1.70E-01 | 2.52E-01 | -1.08 | (-2.23-0.07) | 6.58E-02 | 1.52E-01 |
| 344 | PC(O-32:1) | -1.01 | (-1.56--0.45) | **4.20E-04** | **1.18E-03** | -0.53 | (-1.07-0.02) | 5.89E-02 | 8.34E-02 | 0.73 | (0-1.52) | 5.05E-02 | 8.95E-02 | -0.34 | (-1.82-1.16) | 6.50E-01 | 7.50E-01 |
| 345 | PC(O-34:1) | -0.84 | (-1.27--0.41) | **1.54E-04** | **4.84E-04** | -1.21 | (-1.62--0.79) | **2.44E-08** | **1.08E-07** | 0.68 | (0.02-1.38) | **4.18E-02** | 7.52E-02 | -2.24 | (-3.3--1.18) | **4.84E-05** | **7.04E-04** |
| 346 | PC(O-34:2) | -2.02 | (-2.62--1.41) | **2.10E-10** | **2.80E-09** | -1.75 | (-2.35--1.15) | **2.36E-08** | **1.05E-07** | -0.06 | (-0.77-0.69) | 8.63E-01 | 9.01E-01 | -1.81 | (-3.58-0) | **4.98E-02** | 1.24E-01 |
| 347 | PC(O-34:4) | -0.88 | (-1.84-0.1) | 7.86E-02 | 1.13E-01 | -0.61 | (-1.64-0.42) | 2.46E-01 | 3.02E-01 | -0.01 | (-0.7-0.73) | 9.69E-01 | 9.77E-01 | 1.29 | (-1.45-4.11) | 3.58E-01 | 5.10E-01 |
| 348 | PC(O-35:4) | -0.89 | (-1.73--0.04) | **4.09E-02** | 6.65E-02 | -0.70 | (-1.63-0.24) | 1.44E-01 | 1.86E-01 | -0.49 | (-1.13-0.19) | 1.52E-01 | 2.29E-01 | 0.45 | (-2.21-3.19) | 7.41E-01 | 8.21E-01 |
| 349 | PC(O-36:0) | -1.30 | (-1.8--0.81) | **3.88E-07** | **2.12E-06** | -0.84 | (-1.33--0.35) | **9.07E-04** | **1.95E-03** | 0.14 | (-0.67-1.02) | 7.52E-01 | 8.24E-01 | -2.41 | (-3.76--1.03) | **6.74E-04** | **4.23E-03** |
| 350 | PC(O-18:0/18:1) | -1.51 | (-1.98--1.03) | **1.00E-09** | **1.02E-08** | -1.48 | (-1.92--1.04) | **1.07E-10** | **6.87E-10** | 1.31 | (0.62-2.05) | **1.35E-04** | **3.71E-04** | -3.04 | (-4.22--1.85) | **1.05E-06** | **3.88E-05** |
| 351 | PC(O-18:1/18:1) | -1.20 | (-1.66--0.74) | **4.16E-07** | **2.24E-06** | -1.71 | (-2.13--1.29) | **1.10E-14** | **1.42E-13** | 0.71 | (0.11-1.35) | **2.01E-02** | **3.86E-02** | -1.86 | (-2.95--0.75) | **1.12E-03** | **6.20E-03** |
| 352 | PC(O-18:0/18:2) | -2.36 | (-2.92--1.79) | **3.60E-15** | **1.73E-13** | -1.89 | (-2.38--1.39) | **4.43E-13** | **4.43E-12** | 0.44 | (-0.18-1.11) | 1.67E-01 | 2.51E-01 | -2.63 | (-4.12--1.12) | **7.48E-04** | **4.54E-03** |
| 353 | PC(O-18:1/18:2) | -1.41 | (-1.96--0.86) | **7.43E-07** | **3.71E-06** | -1.34 | (-1.85--0.82) | **6.84E-07** | **2.35E-06** | -0.06 | (-0.7-0.63) | 8.59E-01 | 9.01E-01 | -1.45 | (-2.94-0.06) | 5.92E-02 | 1.43E-01 |
| 354 | PC(O-16:0/20:3) | -0.51 | (-1.03-0.02) | 6.07E-02 | 9.23E-02 | 0.21 | (-0.32-0.76) | 4.34E-01 | 4.92E-01 | 0.39 | (-0.29-1.12) | 2.68E-01 | 3.62E-01 | 0.94 | (-0.55-2.45) | 2.17E-01 | 3.61E-01 |
| 355 | PC(O-16:0/20:4) | -0.15 | (-0.68-0.38) | 5.74E-01 | 6.26E-01 | -0.43 | (-0.95-0.09) | 1.04E-01 | 1.39E-01 | -0.15 | (-0.73-0.46) | 6.14E-01 | 6.98E-01 | 0.53 | (-0.97-2.05) | 4.89E-01 | 6.31E-01 |
| 356 | PC(O-36:5) | -1.11 | (-2.09--0.12) | **2.87E-02** | **4.82E-02** | -0.38 | (-1.4-0.66) | 4.72E-01 | 5.31E-01 | -1.06 | (-1.73--0.34) | **4.52E-03** | **9.91E-03** | 2.54 | (-0.25-5.41) | 7.43E-02 | 1.63E-01 |

| | | | | | | | | | | | | | | | | | |
|---|---|---|---|---|---|---|---|---|---|---|---|---|---|---|---|---|---|
| 357 | PC(O-18:0/20:4) | -0.40 | (-0.91-0.12) | 1.28E-01 | 1.76E-01 | -0.72 | (-1.18--0.24) | **3.06E-03** | **5.83E-03** | -0.31 | (-0.85-0.26) | 2.75E-01 | 3.66E-01 | -1.33 | (-2.77-0.14) | 7.48E-02 | 1.63E-01 |
| 358 | PC(O-38:5) | -0.01 | (-0.47-0.46) | 9.81E-01 | 9.83E-01 | -0.63 | (-1.12--0.13) | **1.31E-02** | **2.16E-02** | -0.16 | (-0.83-0.55) | 6.44E-01 | 7.28E-01 | 0.05 | (-1.34-1.45) | 9.46E-01 | 9.76E-01 |
| 359 | PC(O-16:0/22:6) | -0.23 | (-0.91-0.45) | 5.00E-01 | 5.53E-01 | -0.30 | (-1-0.4) | 3.99E-01 | 4.55E-01 | -0.81 | (-1.4--0.17) | **1.38E-02** | **2.75E-02** | 1.12 | (-0.78-3.07) | 2.49E-01 | 4.02E-01 |
| 360 | PC(O-40:5) | -0.80 | (-1.3--0.3) | **1.67E-03** | **3.98E-03** | -0.88 | (-1.34--0.42) | **1.90E-04** | **4.60E-04** | -0.16 | (-0.68-0.39) | 5.65E-01 | 6.53E-01 | -0.79 | (-2.12-0.56) | 2.48E-01 | 4.01E-01 |
| 361 | PC(O-18:0/22:6) | -0.69 | (-1.25--0.12) | **1.77E-02** | **3.15E-02** | -1.09 | (-1.63--0.55) | **9.15E-05** | **2.31E-04** | -0.93 | (-1.43--0.39) | **9.93E-04** | **2.41E-03** | -1.23 | (-2.72-0.29) | 1.11E-01 | 2.14E-01 |
| 362 | PC(O-40:7) (a) | -0.06 | (-0.71-0.59) | 8.54E-01 | 8.76E-01 | -0.80 | (-1.41--0.18) | **1.10E-02** | **1.83E-02** | -0.71 | (-1.28--0.11) | **2.17E-02** | **4.11E-02** | -0.07 | (-1.89-1.79) | 9.42E-01 | 9.74E-01 |
| 363 | PC(P-16:0/14:0) | -1.27 | (-1.86--0.68) | **2.71E-05** | **9.86E-05** | -1.18 | (-1.79--0.56) | **1.98E-04** | **4.77E-04** | -0.35 | (-1.06-0.41) | 3.52E-01 | 4.52E-01 | -0.01 | (-1.86-1.89) | 9.95E-01 | 9.99E-01 |
| 364 | PC(P-16:0/16:0) | -0.71 | (-1.21--0.2) | **6.69E-03** | **1.37E-02** | -1.03 | (-1.54--0.53) | **6.60E-05** | **1.72E-04** | -0.52 | (-1.15-0.16) | 1.33E-01 | 2.06E-01 | -1.25 | (-2.56-0.08) | 6.48E-02 | 1.50E-01 |
| 365 | PC(P-16:0/16:1) | -0.71 | (-1.31--0.11) | **2.01E-02** | **3.52E-02** | -0.83 | (-1.39--0.26) | **4.66E-03** | **8.63E-03** | -0.04 | (-0.71-0.69) | 9.17E-01 | 9.36E-01 | 0.80 | (-0.71-2.32) | 3.00E-01 | 4.58E-01 |
| 366 | PC(P-16:0/18:0) | -1.20 | (-1.59--0.81) | **3.58E-09** | **2.91E-08** | -1.47 | (-1.84--1.09) | **1.05E-13** | **1.20E-12** | -0.24 | (-0.76-0.32) | 4.00E-01 | 5.02E-01 | -2.11 | (-3.09--1.13) | **3.73E-05** | **5.60E-04** |
| 367 | PC(P-16:0/18:1) | -1.21 | (-1.75--0.67) | **1.36E-05** | **5.30E-05** | -2.19 | (-2.7--1.69) | **2.94E-16** | **6.14E-15** | -0.28 | (-0.86-0.34) | 3.73E-01 | 4.73E-01 | -1.99 | (-3.17--0.79) | **1.27E-03** | **6.69E-03** |
| 368 | PC(P-16:0/18:2) | -1.10 | (-1.68--0.5) | **3.02E-04** | **8.73E-04** | -2.03 | (-2.57--1.48) | **1.39E-12** | **1.28E-11** | -1.30 | (-1.9--0.66) | **1.43E-04** | **3.91E-04** | -1.51 | (-3.09-0.09) | 6.46E-02 | 1.50E-01 |
| 369 | PC(P-16:0/18:3) | -1.64 | (-2.27--1.01) | **5.09E-07** | **2.66E-06** | -1.42 | (-1.99--0.84) | **2.28E-06** | **7.17E-06** | -0.48 | (-1.01-0.08) | 9.39E-02 | 1.53E-01 | -0.44 | (-2.21-1.35) | 6.24E-01 | 7.41E-01 |
| 370 | PC(P-35:2) (a) | -2.26 | (-3.13--1.39) | **5.70E-07** | **2.91E-06** | -1.87 | (-2.71--1.02) | **1.87E-05** | **5.25E-05** | -0.45 | (-1.13-0.28) | 2.20E-01 | 3.08E-01 | -2.39 | (-4.91-0.2) | 7.04E-02 | 1.58E-01 |
| 371 | PC(P-35:2) (b) | -1.83 | (-2.55--1.1) | **1.07E-06** | **5.20E-06** | -2.14 | (-2.83--1.45) | **2.27E-09** | **1.13E-08** | -1.45 | (-2.06--0.79) | **3.89E-05** | **1.15E-04** | -2.72 | (-4.69--0.7) | **8.59E-03** | **3.22E-02** |
| 372 | PC(P-15:0/20:4) (b) | -0.66 | (-1.34-0.01) | 5.51E-02 | 8.45E-02 | -0.44 | (-1.19-0.32) | 2.53E-01 | 3.07E-01 | -0.76 | (-1.47-0.01) | 5.19E-02 | 9.09E-02 | 0.40 | (-1.76-2.61) | 7.16E-01 | 7.98E-01 |
| 373 | PC(P-18:1/18:1) | -1.20 | (-1.72--0.68) | **7.54E-06** | **3.17E-05** | -1.58 | (-2.08--1.09) | **7.85E-10** | **4.18E-09** | 1.22 | (0.39-2.12) | **3.50E-03** | **7.92E-03** | -0.48 | (-1.71-0.76) | 4.45E-01 | 5.88E-01 |
| 374 | PC(P-18:0/18:2) | -1.93 | (-2.51--1.36) | **1.35E-10** | **2.16E-09** | -2.58 | (-3.1--2.06) | **1.54E-20** | **1.06E-18** | -0.65 | (-1.19--0.07) | **2.78E-02** | 5.15E-02 | -2.97 | (-4.48--1.45) | **1.69E-04** | **1.48E-03** |
| 375 | PC(P-16:0/20:4) | 0.43 | (-0.18-1.05) | 1.67E-01 | 2.22E-01 | -0.65 | (-1.18--0.11) | **1.81E-02** | **2.88E-02** | -1.10 | (-1.65--0.5) | **4.44E-04** | **1.14E-03** | 0.30 | (-1.26-1.89) | 7.03E-01 | 7.91E-01 |
| 376 | PC(P-16:0/20:5) | -0.52 | (-1.56-0.53) | 3.29E-01 | 3.95E-01 | -0.31 | (-1.37-0.77) | 5.75E-01 | 6.33E-01 | -1.95 | (-2.64--1.2) | **2.21E-06** | **7.46E-06** | 3.00 | (0.12-5.97) | **4.13E-02** | 1.10E-01 |
| 377 | PC(P-17:0/20:4) (a) | -1.01 | (-1.8--0.22) | **1.29E-02** | **2.38E-02** | -0.63 | (-1.41-0.16) | 1.19E-01 | 1.57E-01 | -1.57 | (-2.22--0.86) | **3.59E-05** | **1.08E-04** | -0.85 | (-3.43-1.79) | 5.22E-01 | 6.50E-01 |
| 378 | PC(P-17:0/20:4) (b) | -0.32 | (-1.05-0.42) | 3.96E-01 | 4.59E-01 | -1.00 | (-1.68--0.31) | **4.69E-03** | **8.66E-03** | -1.89 | (-2.47--1.27) | **4.13E-08** | **1.68E-07** | -1.03 | (-3.11-1.1) | 3.39E-01 | 4.95E-01 |
| 379 | PC(P-18:0/20:4) | -0.09 | (-0.67-0.5) | 7.64E-01 | 8.02E-01 | -1.20 | (-1.71--0.68) | **7.83E-06** | **2.32E-05** | -1.09 | (-1.6--0.56) | **1.16E-04** | **3.27E-04** | -0.77 | (-2.4-0.88) | 3.56E-01 | 5.09E-01 |
| 380 | PC(P-38:5) (a) | 0.24 | (-0.36-0.84) | 4.31E-01 | 4.89E-01 | -0.55 | (-1.09--0.01) | **4.46E-02** | 6.57E-02 | -1.21 | (-1.75--0.63) | **8.09E-05** | **2.33E-04** | 0.72 | (-0.88-2.35) | 3.77E-01 | 5.29E-01 |
| 381 | PC(P-38:5) (b) | -0.85 | (-1.43--0.27) | **4.44E-03** | **9.70E-03** | -1.10 | (-1.67--0.53) | **1.76E-04** | **4.33E-04** | -0.87 | (-1.5--0.2) | **1.16E-02** | **2.34E-02** | -0.41 | (-2.09-1.31) | 6.40E-01 | 7.45E-01 |
| 382 | PC(P-16:0/22:6) | -0.29 | (-1.01-0.43) | 4.26E-01 | 4.86E-01 | -0.95 | (-1.62--0.28) | **5.79E-03** | **1.03E-02** | -1.52 | (-2.09--0.91) | **3.95E-06** | **1.31E-05** | 0.28 | (-1.67-2.27) | 7.79E-01 | 8.55E-01 |
| 383 | PC(P-20:0/20:4) | -0.43 | (-1.04-0.17) | 1.58E-01 | 2.12E-01 | -1.30 | (-1.83--0.77) | **1.74E-06** | **5.57E-06** | -0.86 | (-1.45--0.23) | **7.93E-03** | **1.68E-02** | -1.69 | (-3.16--0.2) | **2.69E-02** | 7.82E-02 |
| 384 | PC(P-40:5) (b) | -0.80 | (-1.37--0.24) | **5.56E-03** | **1.18E-02** | -1.41 | (-1.93--0.9) | **1.16E-07** | **4.47E-07** | -0.80 | (-1.31--0.26) | **4.26E-03** | **9.37E-03** | -1.36 | (-2.75-0.05) | 5.94E-02 | 1.43E-01 |
| 385 | PC(P-18:0/22:6) | -0.68 | (-1.37-0) | 5.13E-02 | 7.95E-02 | -2.07 | (-2.72--1.42) | **9.79E-10** | **5.11E-09** | -1.57 | (-2.08--1.02) | **1.26E-07** | **4.78E-07** | -1.63 | (-3.5-0.27) | 9.23E-02 | 1.88E-01 |
| 386 | PE(16:0_16:1) | -0.54 | (-1.69-0.63) | 3.62E-01 | 4.26E-01 | 3.03 | (1.74-4.33) | **3.84E-06** | **1.18E-05** | 0.44 | (-0.57-1.57) | 4.07E-01 | 5.08E-01 | 4.35 | (0.88-7.95) | **1.38E-02** | **4.78E-02** |
| 387 | PE(16:0_18:1) | -0.10 | (-0.87-0.67) | 7.94E-01 | 8.25E-01 | 1.21 | (0.29-2.14) | **1.01E-02** | **1.71E-02** | -0.60 | (-1.29-0.14) | 1.11E-01 | 1.76E-01 | 0.32 | (-2.24-2.94) | 8.11E-01 | 8.80E-01 |
| 388 | PE(16:0_18:2) | -0.32 | (-1.18-0.55) | 4.71E-01 | 5.29E-01 | 0.55 | (-0.44-1.56) | 2.76E-01 | 3.31E-01 | -0.68 | (-1.39-0.1) | 8.58E-02 | 1.42E-01 | -0.95 | (-3.68-1.86) | 5.03E-01 | 6.36E-01 |
| 389 | PE(16:1_18:2) | -0.95 | (-1.96-0.08) | 6.93E-02 | 1.01E-01 | 0.53 | (-0.35-1.43) | 2.39E-01 | 2.94E-01 | 0.38 | (-0.4-1.22) | 3.50E-01 | 4.50E-01 | 0.00 | (-2.28-2.33) | 1.00E+00 | 1.00E+00 |
| 390 | PE(16:0_18:3) (a) | -1.62 | (-2.53--0.69) | **6.80E-04** | **1.77E-03** | 0.67 | (-0.33-1.68) | 1.90E-01 | 2.42E-01 | 0.07 | (-0.71-0.9) | 8.73E-01 | 9.06E-01 | -0.67 | (-3.48-2.22) | 6.45E-01 | 7.48E-01 |
| 391 | PE(16:0_18:3) (b) | -0.42 | (-1.33-0.49) | 3.65E-01 | 4.27E-01 | 1.99 | (0.89-3.11) | **3.82E-04** | **8.81E-04** | 0.56 | (-0.31-1.5) | 2.15E-01 | 3.04E-01 | 1.71 | (-1.39-4.9) | 2.82E-01 | 4.41E-01 |
| 392 | PE(17:0_18:2) | -1.21 | (-1.9--0.52) | **6.65E-04** | **1.74E-03** | 0.07 | (-0.73-0.89) | 8.59E-01 | 8.83E-01 | -0.85 | (-1.35--0.31) | **2.33E-03** | **5.41E-03** | -1.18 | (-3.43-1.13) | 3.14E-01 | 4.72E-01 |
| 393 | PE(18:0_18:1) | 0.43 | (-0.21-1.08) | 1.90E-01 | 2.46E-01 | 1.56 | (0.67-2.45) | **5.73E-04** | **1.26E-03** | -0.09 | (-0.87-0.75) | 8.21E-01 | 8.74E-01 | 2.66 | (0.19-5.19) | **3.45E-02** | 9.47E-02 |
| 394 | PE(18:1_18:1) | -0.40 | (-1.06-0.27) | 2.42E-01 | 3.03E-01 | 0.28 | (-0.49-1.05) | 4.78E-01 | 5.36E-01 | -0.01 | (-1.07-1.17) | 9.83E-01 | 9.83E-01 | 0.26 | (-2.06-2.63) | 8.29E-01 | 8.96E-01 |
| 395 | PE(18:0_18:2) | 0.50 | (-0.21-1.22) | 1.70E-01 | 2.25E-01 | 1.26 | (0.38-2.15) | **5.10E-03** | **9.24E-03** | -0.50 | (-1.17-0.22) | 1.68E-01 | 2.51E-01 | 1.32 | (-1.13-3.82) | 2.93E-01 | 4.50E-01 |
| 396 | PE(18:1_18:2) | -0.77 | (-1.58-0.04) | 6.38E-02 | 9.54E-02 | -0.81 | (-1.77-0.15) | 9.83E-02 | 1.33E-01 | -0.26 | (-1.01-0.56) | 5.29E-01 | 6.21E-01 | -2.14 | (-4.71-0.49) | 1.10E-01 | 2.14E-01 |
| 397 | PE(16:0_20:3) | -0.37 | (-1.21-0.47) | 3.85E-01 | 4.48E-01 | 0.97 | (-0.01-1.97) | 5.21E-02 | 7.54E-02 | -0.31 | (-1.06-0.5) | 4.42E-01 | 5.42E-01 | -0.12 | (-2.72-2.55) | 9.29E-01 | 9.67E-01 |
| 398 | PE(16:0_20:4) | 0.75 | (0.01-1.5) | **4.65E-02** | 7.31E-02 | -0.44 | (-1.27-0.4) | 3.04E-01 | 3.60E-01 | -0.34 | (-0.96-0.33) | 3.14E-01 | 4.11E-01 | -1.51 | (-3.89-0.92) | 2.19E-01 | 3.63E-01 |
| 399 | PE(16:1_20:4) | -0.18 | (-1.08-0.73) | 6.98E-01 | 7.41E-01 | -0.08 | (-0.84-0.69) | 8.40E-01 | 8.71E-01 | 0.73 | (-0.09-1.62) | 8.04E-02 | 1.34E-01 | 1.01 | (-1.06-3.12) | 3.42E-01 | 4.98E-01 |
| 400 | PE(16:0_20:5) | -0.47 | (-1.52-0.58) | 3.80E-01 | 4.43E-01 | 0.27 | (-0.78-1.33) | 6.16E-01 | 6.70E-01 | -0.82 | (-1.58-0.01) | 5.18E-02 | 9.09E-02 | 1.62 | (-1.04-4.36) | 2.34E-01 | 3.82E-01 |
| 401 | PE(17:0_20:4) | -0.03 | (-0.7-0.65) | 9.34E-01 | 9.46E-01 | -0.05 | (-0.82-0.72) | 8.96E-01 | 9.17E-01 | -1.71 | (-2.3--1.07) | **8.04E-07** | **2.86E-06** | -1.45 | (-3.59-0.74) | 1.91E-01 | 3.24E-01 |
| 402 | PE(18:0_20:3) (a) | 1.68 | (0.87-2.5) | **4.94E-05** | **1.72E-04** | 2.90 | (1.96-3.84) | **1.44E-09** | **7.41E-09** | -0.61 | (-1.25-0.09) | 8.50E-02 | 1.41E-01 | 3.60 | (1.22-6.04) | **3.02E-03** | **1.41E-02** |
| 403 | PE(18:0_20:4) | 2.04 | (1.41-2.67) | **2.05E-10** | **2.80E-09** | 0.25 | (-0.47-0.96) | 5.01E-01 | 5.59E-01 | -0.70 | (-1.29--0.07) | **3.03E-02** | 5.55E-02 | 0.55 | (-1.48-2.63) | 5.96E-01 | 7.19E-01 |
| 404 | PE(38:5) (a) | 0.01 | (-0.72-0.75) | 9.72E-01 | 9.76E-01 | -0.72 | (-1.56-0.13) | 9.55E-02 | 1.30E-01 | -0.63 | (-1.26-0.04) | 6.53E-02 | 1.12E-01 | -1.61 | (-3.89-0.73) | 1.76E-01 | 3.07E-01 |
| 405 | PE(38:5) (b) | -0.87 | (-1.73--0.01) | **4.69E-02** | 7.36E-02 | 0.00 | (-0.94-0.95) | 9.99E-01 | 9.99E-01 | 0.39 | (-0.63-1.53) | 4.68E-01 | 5.68E-01 | 0.76 | (-2.02-3.62) | 5.95E-01 | 7.19E-01 |
| 406 | PE(16:0_22:6) | 0.00 | (-0.73-0.74) | 9.96E-01 | 9.96E-01 | -0.56 | (-1.51-0.4) | 2.52E-01 | 3.06E-01 | -1.58 | (-2.23--0.88) | **2.50E-05** | **7.75E-05** | -0.46 | (-3.22-2.37) | 7.47E-01 | 8.26E-01 |
| 407 | PE(17:0_22:6) | -0.21 | (-0.95-0.55) | 5.87E-01 | 6.38E-01 | -0.18 | (-0.98-0.62) | 6.55E-01 | 7.03E-01 | -1.52 | (-2.1--0.89) | **7.66E-06** | **2.47E-05** | -0.10 | (-2.25-2.09) | 9.25E-01 | 9.66E-01 |
| 408 | PE(18:0_22:4) | 0.52 | (-0.35-1.4) | 2.43E-01 | 3.03E-01 | 1.66 | (0.84-2.48) | **6.84E-05** | **1.77E-04** | -0.22 | (-1.01-0.65) | 6.13E-01 | 6.98E-01 | 2.27 | (0.13-4.45) | **3.74E-02** | 1.00E-01 |

| | | | | | | | | | | | | | | | | | |
|---|---|---|---|---|---|---|---|---|---|---|---|---|---|---|---|---|---|
| 409 | PE(18:0_22:5) (n3) | 0.45 | (-0.26-1.16) | 2.14E-01 | 2.72E-01 | 1.79 | (0.94-2.64) | **3.74E-05** | **9.97E-05** | -1.09 | (-1.86--0.24) | **1.27E-02** | **2.54E-02** | 2.54 | (0.19-4.95) | **3.42E-02** | 9.47E-02 |
| 410 | PE(18:0_22:5) (n6) | 0.40 | (-0.53-1.33) | 4.00E-01 | 4.62E-01 | 0.72 | (-0.23-1.68) | 1.39E-01 | 1.80E-01 | -0.46 | (-1.41-0.59) | 3.78E-01 | 4.77E-01 | 2.62 | (0.04-5.27) | **4.66E-02** | 1.18E-01 |
| 411 | PE(18:0_22:6) | 1.34 | (0.55-2.14) | **8.34E-04** | **2.13E-03** | 1.67 | (0.73-2.63) | **5.37E-04** | **1.20E-03** | -1.61 | (-2.34--0.8) | **2.02E-04** | **5.43E-04** | 3.81 | (1.08-6.61) | **6.06E-03** | **2.36E-02** |
| 412 | PE(40:7) | -0.40 | (-1.25-0.44) | 3.48E-01 | 4.14E-01 | -1.43 | (-2.41--0.44) | **4.72E-03** | **8.68E-03** | -1.29 | (-1.94--0.58) | **5.38E-04** | **1.36E-03** | -1.14 | (-3.97-1.76) | 4.36E-01 | 5.79E-01 |
| 413 | PE(O-34:1) | -1.42 | (-2--0.83) | **2.58E-06** | **1.19E-05** | -1.43 | (-2.04--0.82) | **6.33E-06** | **1.91E-05** | 0.33 | (-0.78-1.57) | 5.73E-01 | 6.59E-01 | -1.48 | (-3.2-0.26) | 9.51E-02 | 1.93E-01 |
| 414 | PE(O-16:0/18:2) | -2.74 | (-3.5--1.96) | **1.39E-11** | **2.79E-10** | -2.48 | (-3.27--1.68) | **2.70E-09** | **1.32E-08** | -0.23 | (-2-1.95) | 8.24E-01 | 8.75E-01 | -0.34 | (-2.54-1.91) | 7.64E-01 | 8.42E-01 |
| 415 | PE(O-18:1/18:2) | -1.56 | (-2.3--0.82) | **4.20E-05** | **1.47E-04** | -1.10 | (-1.81--0.38) | **2.71E-03** | **5.25E-03** | 0.06 | (-0.88-1.09) | 9.08E-01 | 9.31E-01 | -0.25 | (-2.27-1.81) | 8.10E-01 | 8.80E-01 |
| 416 | PE(O-16:0/20:4) | -1.39 | (-2.12--0.65) | **2.38E-04** | **7.02E-04** | -1.38 | (-2.13--0.61) | **4.51E-04** | **1.03E-03** | -0.81 | (-1.62-0.07) | 6.94E-02 | 1.18E-01 | 1.19 | (-0.97-3.4) | 2.82E-01 | 4.41E-01 |
| 417 | PE(O-36:5) | -2.34 | (-3.42--1.25) | **3.37E-05** | **1.20E-04** | -1.17 | (-2.3--0.03) | **4.50E-02** | 6.59E-02 | -0.42 | (-1.08-0.29) | 2.38E-01 | 3.31E-01 | 2.28 | (-0.48-5.12) | 1.06E-01 | 2.10E-01 |
| 418 | PE(O-16:0/22:4) | -0.63 | (-1.21--0.05) | **3.37E-02** | 5.62E-02 | -0.79 | (-1.37--0.22) | **7.21E-03** | **1.26E-02** | 0.56 | (-0.18-1.36) | 1.43E-01 | 2.19E-01 | 0.37 | (-1.29-2.06) | 6.60E-01 | 7.54E-01 |
| 419 | PE(O-18:0/20:4) | -1.81 | (-2.63--0.99) | **2.02E-05** | **7.69E-05** | -1.76 | (-2.55--0.97) | **1.81E-05** | **5.11E-05** | -0.17 | (-0.95-0.69) | 6.94E-01 | 7.73E-01 | 0.60 | (-1.82-3.08) | 6.30E-01 | 7.41E-01 |
| 420 | PE(O-38:5) (a) | -1.44 | (-2.16--0.71) | **1.14E-04** | **3.65E-04** | -0.89 | (-1.6--0.18) | **1.37E-02** | **2.25E-02** | -0.10 | (-0.78-0.64) | 7.90E-01 | 8.51E-01 | 1.52 | (-0.73-3.83) | 1.86E-01 | 3.18E-01 |
| 421 | PE(O-38:5) (b) | -1.98 | (-2.64--1.31) | **9.88E-09** | **6.87E-08** | -1.06 | (-1.81--0.29) | **6.73E-03** | **1.19E-02** | 0.16 | (-0.75-1.17) | 7.38E-01 | 8.15E-01 | 0.24 | (-1.68-2.19) | 8.07E-01 | 8.80E-01 |
| 422 | PE(O-16:0/22:6) | -1.03 | (-1.66--0.4) | **1.52E-03** | **3.67E-03** | -0.83 | (-1.53--0.13) | **2.05E-02** | **3.21E-02** | 0.55 | (-0.23-1.38) | 1.70E-01 | 2.52E-01 | 0.77 | (-1.03-2.6) | 4.02E-01 | 5.53E-01 |
| 423 | PE(O-18:0/22:6) | -1.54 | (-2.14--0.93) | **1.02E-06** | **5.01E-06** | -1.41 | (-2.04--0.77) | **1.78E-05** | **5.04E-05** | 0.01 | (-0.62-0.68) | 9.77E-01 | 9.79E-01 | -0.02 | (-1.8-1.8) | 9.85E-01 | 9.93E-01 |
| 424 | PE(O-18:1/22:6) | -1.16 | (-1.76--0.56) | **1.73E-04** | **5.35E-04** | -1.29 | (-1.9--0.67) | **5.13E-05** | **1.35E-04** | 0.08 | (-0.49-0.68) | 7.96E-01 | 8.54E-01 | 0.41 | (-1.42-2.28) | 6.60E-01 | 7.54E-01 |
| 425 | PE(P-16:0/18:1) | -1.04 | (-1.56--0.52) | **1.03E-04** | **3.35E-04** | -0.62 | (-1.13--0.11) | **1.76E-02** | **2.81E-02** | 1.69 | (0.98-2.44) | **1.11E-06** | **3.90E-06** | -0.19 | (-1.6-1.25) | 7.98E-01 | 8.73E-01 |
| 426 | PE(P-16:0/18:2) | -1.53 | (-2.2--0.85) | **1.07E-05** | **4.41E-05** | -0.73 | (-1.4--0.05) | **3.44E-02** | 5.14E-02 | 0.14 | (-0.73-1.1) | 7.59E-01 | 8.29E-01 | 0.55 | (-1.4-2.55) | 5.80E-01 | 7.05E-01 |
| 427 | PE(P-16:0/20:3) | -0.36 | (-1.08-0.37) | 3.33E-01 | 3.97E-01 | 1.10 | (0.38-1.82) | **2.78E-03** | **5.35E-03** | 0.18 | (-0.65-1.09) | 6.78E-01 | 7.58E-01 | 4.14 | (2.26-6.06) | **1.60E-05** | **3.36E-04** |
| 428 | PE(P-16:0/20:4) | 0.15 | (-0.58-0.88) | 6.90E-01 | 7.36E-01 | 0.18 | (-0.55-0.9) | 6.32E-01 | 6.83E-01 | -0.23 | (-0.99-0.59) | 5.66E-01 | 6.53E-01 | 3.45 | (1.36-5.57) | **1.16E-03** | **6.34E-03** |
| 429 | PE(P-16:0/20:5) | -1.00 | (-2.32-0.34) | 1.42E-01 | 1.93E-01 | 0.17 | (-1.12-1.48) | 7.96E-01 | 8.34E-01 | -0.81 | (-2.51-1.27) | 4.15E-01 | 5.15E-01 | 4.78 | (1.15-8.54) | **9.55E-03** | **3.47E-02** |
| 430 | PE(P-16:0/22:4) | 0.38 | (-0.18-0.94) | 1.87E-01 | 2.43E-01 | -0.41 | (-0.96-0.13) | 1.38E-01 | 1.79E-01 | 0.23 | (-0.47-0.98) | 5.32E-01 | 6.23E-01 | -0.44 | (-1.83-0.97) | 5.37E-01 | 6.65E-01 |
| 431 | PE(P-16:0/22:5) (n3) | -0.62 | (-1.21--0.02) | **4.12E-02** | 6.65E-02 | 0.30 | (-0.26-0.87) | 2.97E-01 | 3.54E-01 | -0.48 | (-1.19-0.28) | 2.09E-01 | 2.98E-01 | 1.24 | (-0.52-3.03) | 1.69E-01 | 3.02E-01 |
| 432 | PE(P-16:0/22:5) (n6) | -0.76 | (-1.4--0.12) | **2.07E-02** | **3.60E-02** | -0.37 | (-0.95-0.22) | 2.21E-01 | 2.77E-01 | 0.91 | (0.1-1.79) | **2.66E-02** | **4.98E-02** | 0.45 | (-1.46-2.39) | 6.48E-01 | 7.49E-01 |
| 433 | PE(P-16:0/22:6) | 0.09 | (-0.49-0.67) | 7.67E-01 | 8.02E-01 | -0.02 | (-0.62-0.58) | 9.45E-01 | 9.61E-01 | 0.17 | (-0.5-0.88) | 6.33E-01 | 7.19E-01 | 1.74 | (0.13-3.38) | **3.45E-02** | 9.47E-02 |
| 434 | PE(P-17:0/20:4) (a) | -0.28 | (-1.14-0.59) | 5.22E-01 | 5.75E-01 | -0.20 | (-1.09-0.7) | 6.62E-01 | 7.10E-01 | -0.39 | (-0.98-0.23) | 2.11E-01 | 2.99E-01 | 1.34 | (-1-3.73) | 2.64E-01 | 4.21E-01 |
| 435 | PE(P-17:0/20:4) (b) | -0.21 | (-1.01-0.6) | 6.15E-01 | 6.63E-01 | -0.27 | (-1.05-0.52) | 5.04E-01 | 5.62E-01 | -0.96 | (-1.57--0.3) | **4.73E-03** | **1.03E-02** | 1.16 | (-0.96-3.33) | 2.86E-01 | 4.45E-01 |
| 436 | PE(P-17:0/22:6) (a) | -1.24 | (-2.03--0.44) | **2.33E-03** | **5.35E-03** | -0.51 | (-1.32-0.31) | 2.22E-01 | 2.77E-01 | -0.27 | (-1.02-0.55) | 5.11E-01 | 6.02E-01 | 0.05 | (-2.29-2.45) | 9.65E-01 | 9.85E-01 |
| 437 | PE(P-17:0/22:6) (b) | -0.75 | (-1.46--0.03) | **4.23E-02** | 6.81E-02 | -0.90 | (-1.67--0.13) | **2.28E-02** | **3.55E-02** | -0.44 | (-1.11-0.27) | 2.18E-01 | 3.06E-01 | -0.50 | (-2.48-1.53) | 6.27E-01 | 7.41E-01 |
| 438 | PE(P-18:0/18:1) | -1.25 | (-1.84--0.66) | **3.98E-05** | **1.40E-04** | -1.01 | (-1.6--0.41) | **9.72E-04** | **2.07E-03** | 1.01 | (0.24-1.85) | **9.34E-03** | **1.94E-02** | -0.71 | (-2.29-0.88) | 3.78E-01 | 5.29E-01 |
| 439 | PE(P-18:0/18:2) | -1.83 | (-2.48--1.19) | **4.04E-08** | **2.66E-07** | -1.53 | (-2.15--0.91) | **1.62E-06** | **5.22E-06** | -0.15 | (-1-0.78) | 7.43E-01 | 8.16E-01 | -0.69 | (-2.59-1.24) | 4.78E-01 | 6.22E-01 |
| 440 | PE(P-18:0/20:3) | -0.92 | (-1.55--0.28) | **4.71E-03** | **1.01E-02** | -0.15 | (-0.76-0.46) | 6.31E-01 | 6.83E-01 | -0.34 | (-0.91-0.26) | 2.56E-01 | 3.51E-01 | 2.28 | (0.68-3.89) | **5.08E-03** | **2.10E-02** |
| 441 | PE(P-18:0/20:4) | -0.19 | (-0.95-0.58) | 6.33E-01 | 6.81E-01 | -0.83 | (-1.51--0.15) | **1.66E-02** | **2.68E-02** | -0.65 | (-1.33-0.08) | 7.77E-02 | 1.30E-01 | 1.63 | (-0.45-3.75) | 1.24E-01 | 2.33E-01 |
| 442 | PE(P-18:0/20:5) | -1.63 | (-2.99--0.26) | **2.00E-02** | **3.52E-02** | -0.80 | (-2.1-0.52) | 2.34E-01 | 2.91E-01 | -1.97 | (-3.3--0.38) | **1.77E-02** | **3.42E-02** | 3.62 | (0.14-7.22) | **4.16E-02** | 1.10E-01 |
| 443 | PE(P-18:0/22:4) | -0.72 | (-1.31--0.13) | **1.74E-02** | **3.11E-02** | -0.98 | (-1.57--0.38) | **1.48E-03** | **3.02E-03** | -0.28 | (-1.04-0.54) | 4.88E-01 | 5.80E-01 | -1.74 | (-3.12--0.34) | **1.51E-02** | 5.03E-02 |
| 444 | PE(P-18:0/22:5) (n3) | -0.81 | (-1.4--0.21) | **8.18E-03** | **1.61E-02** | -0.47 | (-1-0.07) | 8.56E-02 | 1.17E-01 | -0.41 | (-1.01-0.23) | 2.02E-01 | 2.89E-01 | 0.50 | (-1.13-2.16) | 5.49E-01 | 6.78E-01 |
| 445 | PE(P-18:0/22:5) (n6) | -1.29 | (-1.87--0.72) | **1.35E-05** | **5.30E-05** | -1.25 | (-1.83--0.67) | **3.16E-05** | **8.48E-05** | 0.36 | (-0.27-1.04) | 2.67E-01 | 3.62E-01 | -0.44 | (-2.36-1.51) | 6.54E-01 | 7.51E-01 |
| 446 | PE(P-18:0/22:6) | -0.79 | (-1.43--0.14) | **1.80E-02** | **3.18E-02** | -1.21 | (-1.87--0.55) | **3.71E-04** | **8.65E-04** | -0.06 | (-0.69-0.62) | 8.57E-01 | 9.01E-01 | 0.00 | (-1.8-1.82) | 9.98E-01 | 1.00E+00 |
| 447 | PE(P-18:1/18:1) | -0.90 | (-1.45--0.35) | **1.43E-03** | **3.50E-03** | -0.83 | (-1.38--0.28) | **3.06E-03** | **5.83E-03** | 0.36 | (-0.26-1.02) | 2.58E-01 | 3.53E-01 | -0.47 | (-1.9-0.98) | 5.21E-01 | 6.50E-01 |
| 448 | PE(P-18:1/18:2) | -1.86 | (-2.56--1.16) | **3.27E-07** | **1.80E-06** | -1.22 | (-1.9--0.53) | **5.96E-04** | **1.30E-03** | -0.63 | (-1.73-0.61) | 3.01E-01 | 3.97E-01 | -0.76 | (-2.77-1.29) | 4.62E-01 | 6.05E-01 |
| 449 | PE(P-18:1/20:4) | -0.06 | (-0.81-0.71) | 8.85E-01 | 9.02E-01 | -0.43 | (-1.13-0.27) | 2.28E-01 | 2.83E-01 | -0.55 | (-1.23-0.19) | 1.40E-01 | 2.14E-01 | 1.88 | (-0.24-4.04) | 8.28E-02 | 1.76E-01 |
| 450 | PE(P-18:1/20:5) | -1.57 | (-2.88--0.25) | **2.01E-02** | **3.52E-02** | -0.22 | (-1.51-1.08) | 7.37E-01 | 7.75E-01 | -0.82 | (-2.78-1.69) | 4.88E-01 | 5.80E-01 | 3.76 | (0.1-7.55) | **4.42E-02** | 1.14E-01 |
| 451 | PE(P-18:1/22:4) | -0.28 | (-0.92-0.37) | 4.02E-01 | 4.62E-01 | -0.99 | (-1.63--0.35) | **2.44E-03** | **4.77E-03** | 0.33 | (-0.55-1.3) | 4.70E-01 | 5.68E-01 | -1.15 | (-2.57-0.28) | 1.14E-01 | 2.16E-01 |
| 452 | PE(P-18:1/22:5) (a) | -0.71 | (-1.33--0.09) | **2.54E-02** | **4.35E-02** | -0.39 | (-0.96-0.18) | 1.83E-01 | 2.33E-01 | -0.38 | (-1-0.29) | 2.55E-01 | 3.51E-01 | 0.91 | (-0.84-2.69) | 3.07E-01 | 4.66E-01 |
| 453 | PE(P-18:1/22:6) (a) | -0.50 | (-1.16-0.15) | 1.32E-01 | 1.81E-01 | -0.66 | (-1.33-0.02) | 5.55E-02 | 7.90E-02 | -0.05 | (-0.73-0.67) | 8.79E-01 | 9.09E-01 | 0.85 | (-1.04-2.77) | 3.80E-01 | 5.29E-01 |
| 454 | PE(P-20:0/18:1) | -1.27 | (-1.89--0.65) | **7.01E-05** | **2.35E-04** | -1.14 | (-1.71--0.57) | **1.01E-04** | **2.52E-04** | 0.52 | (-1.11-2.44) | 5.56E-01 | 6.45E-01 | -1.09 | (-2.82-0.67) | 2.22E-01 | 3.67E-01 |
| 455 | PE(P-20:0/18:2) | -1.81 | (-2.36--1.25) | **3.40E-10** | **4.18E-09** | -1.98 | (-2.5--1.45) | **1.04E-12** | **9.79E-12** | -0.03 | (-0.46-0.42) | 9.01E-01 | 9.28E-01 | -2.72 | (-4.31--1.11) | **1.09E-03** | **6.15E-03** |
| 456 | PE(P-20:0/20:4) | -0.17 | (-0.76-0.43) | 5.85E-01 | 6.36E-01 | -1.62 | (-2.15--1.08) | **6.16E-09** | **2.90E-08** | -0.61 | (-1.46-0.34) | 2.00E-01 | 2.87E-01 | -0.21 | (-1.81-1.41) | 7.97E-01 | 8.73E-01 |
| 457 | PE(P-20:0/22:6) | -0.78 | (-1.31--0.26) | **3.71E-03** | **8.27E-03** | -1.89 | (-2.39--1.4) | **3.83E-13** | **3.92E-12** | -0.70 | (-1.25--0.11) | **2.13E-02** | **4.05E-02** | -1.79 | (-3.28--0.27) | **2.13E-02** | 6.63E-02 |
| 458 | PE(P-20:1/20:4) | -0.41 | (-1.28-0.47) | 3.64E-01 | 4.27E-01 | -1.06 | (-2.02--0.09) | **3.22E-02** | **4.84E-02** | -0.82 | (-1.84-0.32) | 1.51E-01 | 2.29E-01 | 0.93 | (-1.28-3.19) | 4.10E-01 | 5.58E-01 |
| 459 | PG(34:2) | -1.07 | (-1.82--0.31) | **6.14E-03** | **1.28E-02** | 1.17 | (0.21-2.14) | **1.72E-02** | **2.76E-02** | -0.67 | (-1.86-0.7) | 3.23E-01 | 4.21E-01 | 2.03 | (-0.75-4.88) | 1.53E-01 | 2.80E-01 |
| 460 | PG(36:1) | -0.28 | (-0.94-0.38) | 4.00E-01 | 4.62E-01 | 2.52 | (1.6-3.45) | **8.86E-08** | **3.52E-07** | -1.38 | (-2.5--0.09) | **3.65E-02** | 6.63E-02 | 2.63 | (-0.01-5.33) | 5.10E-02 | 1.27E-01 |

| | | | | | | | | | | | | | | | | | |
|---|---|---|---|---|---|---|---|---|---|---|---|---|---|---|---|---|---|
| 461 | PI(16:0/16:0) | -2.44 | (-3.37--1.51) | **4.46E-07** | **2.38E-06** | 1.75 | (0.7-2.81) | **1.12E-03** | **2.36E-03** | -0.09 | (-1.03-0.93) | 8.50E-01 | 8.99E-01 | 3.09 | (0.46-5.79) | **2.10E-02** | 6.58E-02 |
| 462 | PI(16:0_16:1) | -1.73 | (-2.79--0.66) | **1.69E-03** | **4.01E-03** | 3.85 | (2.77-4.94) | **3.76E-12** | **3.11E-11** | 0.72 | (-0.12-1.62) | 9.40E-02 | 1.53E-01 | 8.42 | (5.56-11.34) | **6.42E-09** | **1.03E-06** |
| 463 | PI(34:0) | -2.66 | (-3.58--1.72) | **4.56E-08** | **2.96E-07** | 1.08 | (0-2.17) | **4.90E-02** | 7.13E-02 | -1.86 | (-2.99--0.54) | **7.41E-03** | **1.58E-02** | -0.02 | (-3.05-3.11) | 9.90E-01 | 9.96E-01 |
| 464 | PI(34:1) | -2.01 | (-2.68--1.34) | **8.14E-09** | **5.75E-08** | 0.20 | (-0.53-0.93) | 5.92E-01 | 6.47E-01 | -0.19 | (-0.81-0.47) | 5.56E-01 | 6.45E-01 | 0.45 | (-1.4-2.34) | 6.33E-01 | 7.42E-01 |
| 465 | PI(17:0_18:1) | -2.37 | (-2.95--1.79) | **6.39E-15** | **2.79E-13** | 0.44 | (-0.29-1.18) | 2.37E-01 | 2.93E-01 | -1.30 | (-2.26--0.22) | **1.94E-02** | **3.75E-02** | -0.77 | (-2.94-1.45) | 4.91E-01 | 6.31E-01 |
| 466 | PI(17:0_18:2) | -2.70 | (-3.33--2.07) | **4.72E-16** | **2.83E-14** | 0.69 | (-0.03-1.41) | 6.05E-02 | 8.54E-02 | -1.91 | (-2.87--0.83) | **9.67E-04** | **2.37E-03** | 0.15 | (-1.85-2.18) | 8.85E-01 | 9.42E-01 |
| 467 | PI(18:0_18:1) | -1.94 | (-2.65--1.23) | **1.16E-07** | **7.23E-07** | -0.74 | (-1.49-0.01) | 5.26E-02 | 7.58E-02 | -1.18 | (-1.84--0.46) | **1.71E-03** | **4.02E-03** | -2.01 | (-4.12-0.14) | 6.70E-02 | 1.52E-01 |
| 468 | PI(36:2) | -1.93 | (-2.51--1.34) | **2.53E-10** | **3.28E-09** | -0.30 | (-0.88-0.3) | 3.26E-01 | 3.83E-01 | -1.22 | (-1.78--0.62) | **1.28E-04** | **3.56E-04** | -1.49 | (-3.08-0.13) | 7.12E-02 | 1.58E-01 |
| 469 | PI(18:1_18:2) | -2.66 | (-3.29--2.03) | **1.11E-15** | **5.91E-14** | -1.29 | (-1.97--0.61) | **2.16E-04** | **5.17E-04** | -1.47 | (-1.98--0.93) | **4.93E-07** | **1.78E-06** | -3.22 | (-4.99--1.41) | **5.62E-04** | **3.65E-03** |
| 470 | PI(16:0_20:3) (a) | -2.16 | (-2.82--1.5) | **3.96E-10** | **4.64E-09** | 0.93 | (0.22-1.65) | **1.07E-02** | **1.78E-02** | 0.22 | (-0.31-0.79) | 4.19E-01 | 5.18E-01 | -0.04 | (-1.72-1.66) | 9.62E-01 | 9.85E-01 |
| 471 | PI(16:0_20:3) (b) | -2.46 | (-3.2--1.72) | **1.99E-10** | **2.80E-09** | 0.86 | (0.13-1.6) | **2.06E-02** | **3.21E-02** | 2.99 | (1.94-4.14) | **2.09E-09** | **9.74E-09** | 0.67 | (-1.27-2.64) | 5.00E-01 | 6.34E-01 |
| 472 | PI(16:0_20:4) | -1.45 | (-2.03--0.86) | **1.88E-06** | **8.78E-06** | 0.99 | (0.36-1.62) | **2.13E-03** | **4.20E-03** | 0.12 | (-0.42-0.7) | 6.69E-01 | 7.50E-01 | 1.36 | (-0.32-3.07) | 1.12E-01 | 2.14E-01 |
| 473 | PI(18:0_20:2) | -2.66 | (-3.36--1.95) | **6.56E-13** | **1.85E-11** | -0.07 | (-0.8-0.66) | 8.50E-01 | 8.75E-01 | 2.45 | (1.2-3.85) | **5.83E-05** | **1.70E-04** | -2.45 | (-4.37--0.49) | **1.48E-02** | 5.00E-02 |
| 474 | PI(18:0_20:3) (a) | -0.92 | (-1.42--0.41) | **4.19E-04** | **1.18E-03** | -0.19 | (-0.7-0.31) | 4.57E-01 | 5.16E-01 | -0.28 | (-0.75-0.22) | 2.65E-01 | 3.60E-01 | -1.02 | (-2.35-0.32) | 1.33E-01 | 2.48E-01 |
| 475 | PI(18:0_20:4) | -0.79 | (-1.31--0.28) | **2.68E-03** | **6.09E-03** | -0.13 | (-0.63-0.37) | 6.01E-01 | 6.56E-01 | -0.59 | (-1.01--0.14) | **1.08E-02** | **2.19E-02** | -1.18 | (-2.55-0.2) | 9.21E-02 | 1.88E-01 |
| 476 | PI(38:5) (a) | -2.43 | (-2.97--1.88) | **5.10E-17** | **4.08E-15** | -0.27 | (-0.84-0.3) | 3.46E-01 | 4.00E-01 | -0.59 | (-1.16-0.02) | 5.72E-02 | 9.95E-02 | -1.02 | (-2.47-0.45) | 1.73E-01 | 3.06E-01 |
| 477 | PI(38:6) | -2.63 | (-3.27--1.98) | **1.30E-14** | **5.19E-13** | -0.89 | (-1.56--0.21) | **1.01E-02** | **1.70E-02** | -0.92 | (-1.44--0.37) | **1.31E-03** | **3.11E-03** | -0.06 | (-1.76-1.66) | 9.42E-01 | 9.74E-01 |
| 478 | PI(18:0_22:4) | -1.24 | (-1.89--0.58) | **2.27E-04** | **6.78E-04** | 0.25 | (-0.38-0.9) | 4.35E-01 | 4.92E-01 | 1.20 | (0.52-1.92) | **4.03E-04** | **1.05E-03** | -0.81 | (-2.49-0.89) | 3.47E-01 | 5.03E-01 |
| 479 | PI(18:0_22:5) (n3) | -1.73 | (-2.4--1.06) | **5.69E-07** | **2.91E-06** | 0.86 | (0.2-1.52) | **1.06E-02** | **1.78E-02** | -1.24 | (-1.88--0.56) | **5.87E-04** | **1.47E-03** | 0.63 | (-1.16-2.45) | 4.93E-01 | 6.31E-01 |
| 480 | PI(18:0_22:6) | -2.03 | (-2.68--1.38) | **2.31E-09** | **2.05E-08** | -0.63 | (-1.3-0.05) | 6.94E-02 | 9.71E-02 | -1.11 | (-1.63--0.55) | **1.59E-04** | **4.30E-04** | -0.13 | (-1.98-1.75) | 8.89E-01 | 9.44E-01 |

**Supplementary Table 4a.** The overlapping lipid species between the ppBMI and postnatal BMI studies. Related to Figure 3.

| | ppBMI (antenatal) | | | | Adult BMI (postnatal) | | | | |
|---|---|---|---|---|---|---|---|---|---|
| Lipid Species | %change in lipid concentration per unit BMI | 95%CI | p-value | p-value(BH) | %change in lipid concentration per unit BMI | 95%CI | p-value | p-value(BH) | Trend |
| CE(16:1) | -1.06 | (-1.82--0.29) | **7.23E-03** | **1.46E-02** | 2.17 | (1.53-2.82) | **6.89E-11** | **4.60E-10** | opposite |
| CE(18:3) | -1.93 | (-2.54--1.32) | **1.28E-09** | **1.25E-08** | 0.72 | (0.11-1.32) | **2.04E-02** | **3.21E-02** | opposite |
| CE(20:1) | -0.83 | (-1.29--0.37) | **4.07E-04** | **1.15E-03** | -0.70 | (-1.17--0.22) | **4.06E-03** | **7.58E-03** | same |
| CE(22:5) (n6) | -1.88 | (-2.46--1.29) | **7.49E-10** | **7.92E-09** | -0.68 | (-1.1--0.26) | **1.77E-03** | **3.56E-03** | same |
| DG(16:0_18:1) | 1.33 | (0.59-2.07) | **4.36E-04** | **1.22E-03** | 4.26 | (3.33-5.21) | **1.32E-18** | **4.86E-17** | same |
| DG(16:0_18:2) | 0.92 | (0.21-1.65) | **1.15E-02** | **2.16E-02** | 3.62 | (2.72-4.52) | **5.53E-15** | **7.59E-14** | same |
| DG(18:0_18:1) | 1.36 | (0.69-2.04) | **7.91E-05** | **2.60E-04** | 4.41 | (3.51-5.31) | **4.70E-21** | **4.51E-19** | same |
| DG(18:1_18:1) | 1.37 | (0.64-2.1) | **2.23E-04** | **6.69E-04** | 2.93 | (2.08-3.77) | **1.44E-11** | **1.03E-10** | same |
| DG(18:0_18:2) | 0.99 | (0.23-1.76) | **1.04E-02** | **1.97E-02** | 4.40 | (3.41-5.39) | **7.51E-18** | **2.57E-16** | same |
| DG(18:1_18:2) | 0.95 | (0.22-1.68) | **1.02E-02** | **1.97E-02** | 2.26 | (1.45-3.08) | **5.91E-08** | **2.42E-07** | same |
| DG(16:0_20:4) | 1.19 | (0.26-2.12) | **1.22E-02** | **2.27E-02** | 4.27 | (3.25-5.3) | **4.79E-16** | **9.59E-15** | same |
| DG(18:1_20:4) | 1.14 | (0.4-1.88) | **2.42E-03** | **5.53E-03** | 2.47 | (1.71-3.22) | **1.81E-10** | **1.10E-09** | same |
| DG(18:1_22:6) | 1.53 | (0.47-2.61) | **4.63E-03** | **9.97E-03** | 3.18 | (2.04-4.34) | **5.26E-08** | **2.22E-07** | same |
| DG(18:2_22:6) | 1.52 | (0.42-2.63) | **6.55E-03** | **1.35E-02** | 3.05 | (1.87-4.24) | **3.60E-07** | **1.28E-06** | same |
| TG(48:0) [NL-18:0] | -1.56 | (-2.43--0.68) | **5.76E-04** | **1.53E-03** | 4.34 | (2.99-5.71) | **3.54E-10** | **2.07E-09** | opposite |
| TG(48:2) [NL-14:1] | -1.20 | (-2.25--0.13) | **2.78E-02** | **4.70E-02** | 4.79 | (3.42-6.18) | **7.51E-12** | **5.72E-11** | opposite |
| TG(48:2) [NL-18:2] | -1.16 | (-2.2--0.12) | **2.91E-02** | **4.87E-02** | 5.21 | (3.68-6.77) | **2.87E-11** | **2.03E-10** | opposite |
| TG(48:3) [NL-14:0] | -1.51 | (-2.64--0.37) | **9.55E-03** | **1.85E-02** | 4.32 | (2.9-5.76) | **2.31E-09** | **1.14E-08** | opposite |
| TG(48:3) [NL-18:3] | -1.81 | (-3.07--0.53) | **5.76E-03** | **1.21E-02** | 4.99 | (3.41-6.59) | **5.23E-10** | **2.92E-09** | opposite |
| TG(50:1) [NL-16:0] | 0.83 | (0.22-1.43) | **7.27E-03** | **1.46E-02** | 4.34 | (3.39-5.3) | **1.03E-18** | **4.12E-17** | same |
| TG(50:1) [NL-18:1] | 0.76 | (0.18-1.34) | **1.03E-02** | **1.97E-02** | 4.10 | (3.16-5.05) | **4.06E-17** | **1.08E-15** | same |
| TG(50:2) [NL-14:0] | -0.65 | (-1.21--0.07) | **2.69E-02** | **4.57E-02** | 2.80 | (1.76-3.86) | **1.50E-07** | **5.54E-07** | opposite |
| TG(50:2) [NL-18:2] | 0.66 | (0.08-1.24) | **2.58E-02** | **4.41E-02** | 3.72 | (2.82-4.62) | **8.45E-16** | **1.53E-14** | same |
| TG(52:1) [NL-18:0] | 1.38 | (0.59-2.18) | **6.58E-04** | **1.73E-03** | 5.69 | (4.53-6.86) | **2.15E-21** | **2.58E-19** | same |
| TG(52:1) [NL-18:1] | 1.45 | (0.69-2.23) | **2.04E-04** | **6.17E-04** | 5.86 | (4.73-7) | **1.79E-23** | **8.60E-21** | same |
| TG(52:2) [NL-16:0] | 0.80 | (0.4-1.21) | **1.01E-04** | **3.29E-04** | 2.18 | (1.57-2.79) | **3.69E-12** | **3.11E-11** | same |
| TG(52:3) [NL-18:2] | 0.76 | (0.32-1.2) | **8.08E-04** | **2.08E-03** | 1.84 | (1.22-2.48) | **1.11E-08** | **5.14E-08** | same |
| TG(52:4) [NL-18:2] | 1.20 | (0.44-1.96) | **1.92E-03** | **4.52E-03** | 2.29 | (1.29-3.3) | **7.06E-06** | **2.12E-05** | same |
| TG(53:2) [NL-18:1] | 0.88 | (0.2-1.56) | **1.08E-02** | **2.03E-02** | 2.66 | (1.85-3.47) | **1.63E-10** | **1.00E-09** | same |
| TG(54:1) [NL-18:1] | 0.99 | (0.32-1.66) | **3.57E-03** | **8.00E-03** | 4.45 | (3.57-5.34) | **4.54E-22** | **7.26E-20** | same |
| TG(54:2) [NL-18:0] | 1.41 | (0.72-2.1) | **6.55E-05** | **2.23E-04** | 4.24 | (3.35-5.13) | **2.72E-20** | **1.63E-18** | same |
| TG(54:3) [NL-18:2] | 0.72 | (0.14-1.3) | **1.43E-02** | **2.59E-02** | 2.23 | (1.57-2.89) | **5.13E-11** | **3.51E-10** | same |
| TG(54:5) [NL-20:4] | 1.04 | (0.38-1.71) | **1.91E-03** | **4.51E-03** | 3.37 | (2.55-4.2) | **2.12E-15** | **3.39E-14** | same |
| TG(54:6) [NL-20:4] | 1.16 | (0.33-2) | **6.24E-03** | **1.29E-02** | 3.69 | (2.71-4.68) | **2.17E-13** | **2.27E-12** | same |
| TG(54:6) [NL-22:6] | 1.79 | (0.57-3.02) | **4.10E-03** | **9.06E-03** | 5.76 | (4.33-7.22) | **4.96E-15** | **7.00E-14** | same |
| TG(56:6) [NL-20:4] | 1.34 | (0.81-1.87) | **7.12E-07** | **3.60E-06** | 1.36 | (0.85-1.88) | **2.38E-07** | **8.65E-07** | same |
| TG(56:7) [NL-20:4] | 1.02 | (0.35-1.7) | **2.84E-03** | **6.40E-03** | 1.88 | (1.14-2.62) | **6.63E-07** | **2.29E-06** | same |
| TG(56:7) [NL-22:6] | 1.93 | (0.84-3.04) | **4.97E-04** | **1.35E-03** | 3.96 | (2.65-5.28) | **3.33E-09** | **1.61E-08** | same |
| TG(56:8) [NL-22:6] | 1.76 | (0.54-2.99) | **4.49E-03** | **9.76E-03** | 3.97 | (2.64-5.31) | **4.96E-09** | **2.38E-08** | same |
| TG(58:8) [NL-22:6] | 1.46 | (0.38-2.56) | **7.98E-03** | **1.58E-02** | 2.80 | (1.58-4.04) | **7.21E-06** | **2.15E-05** | same |
| TG(O-54:2) [NL-18:1] | 0.85 | (0.18-1.53) | **1.33E-02** | **2.43E-02** | 2.63 | (1.82-3.44) | **2.48E-10** | **1.47E-09** | same |
| dhCer(d18:0/22:0) | 1.14 | (0.5-1.79) | **4.77E-04** | **1.32E-03** | 1.96 | (1.35-2.58) | **4.98E-10** | **2.81E-09** | same |
| dhCer(d18:0/24:1) | 1.70 | (0.99-2.41) | **3.01E-06** | **1.36E-05** | 1.49 | (0.78-2.2) | **4.22E-05** | **1.11E-04** | same |
| Cer(d16:1/22:0) | -0.73 | (-1.33--0.13) | **1.79E-02** | **3.18E-02** | 1.20 | (0.56-1.84) | **2.31E-04** | **5.51E-04** | opposite |
| Cer(d16:1/23:0) | -0.99 | (-1.66--0.32) | **3.93E-03** | **8.74E-03** | 0.76 | (0.1-1.42) | **2.38E-02** | **3.69E-02** | opposite |
| Cer(d18:1/18:0) | 1.97 | (1.31-2.64) | **7.34E-09** | **5.29E-08** | 2.07 | (1.45-2.7) | **1.07E-10** | **6.87E-10** | same |
| Cer(d18:2/24:0) | -1.25 | (-1.79--0.69) | **1.15E-05** | **4.61E-05** | -0.68 | (-1.22--0.14) | **1.42E-02** | **2.32E-02** | same |
| Cer(d19:1/24:0) | -1.22 | (-2.12--0.32) | **8.01E-03** | **1.58E-02** | 1.08 | (0.23-1.93) | **1.29E-02** | **2.13E-02** | opposite |
| Cer(m18:0/24:1) | 1.15 | (0.26-2.04) | **1.11E-02** | **2.08E-02** | 2.84 | (2-3.68) | **3.94E-11** | **2.74E-10** | same |
| Cer(m18:1/24:1) | 1.37 | (0.47-2.27) | **2.70E-03** | **6.11E-03** | 2.73 | (1.88-3.59) | **4.23E-10** | **2.42E-09** | same |
| GM3(d18:1/18:0) | -1.16 | (-1.7--0.62) | **2.65E-05** | **9.76E-05** | -0.70 | (-1.2--0.2) | **5.94E-03** | **1.06E-02** | same |
| GM3(d18:1/24:0) | -3.17 | (-3.78--2.56) | **5.91E-22** | **9.47E-20** | -1.92 | (-2.46--1.39) | **6.41E-12** | **5.13E-11** | same |
| GM3(d18:1/24:1) | -1.77 | (-2.36--1.17) | **1.05E-08** | **7.23E-08** | -1.28 | (-1.87--0.68) | **2.89E-05** | **7.79E-05** | same |
| HexCer(d18:1/18:0) | -1.56 | (-2.15--0.97) | **3.24E-07** | **1.80E-06** | -1.40 | (-2.04--0.75) | **2.70E-05** | **7.38E-05** | same |
| HexCer(d18:1/20:0) | -1.84 | (-2.41--1.27) | **4.40E-10** | **5.02E-09** | -1.29 | (-1.81--0.76) | **2.18E-06** | **6.88E-06** | same |

| | | | | | | | | | |
|---|---|---|---|---|---|---|---|---|---|
| HexCer(d18:1/22:0) | -1.93 | (-2.43--1.44) | **1.05E-13** | **3.87E-12** | -0.92 | (-1.41--0.43) | **2.61E-04** | **6.17E-04** | same |
| HexCer(d18:1/24:0) | -2.60 | (-3.1--2.09) | **5.92E-22** | **9.47E-20** | -1.34 | (-1.83--0.85) | **1.19E-07** | **4.57E-07** | same |
| HexCer(d18:1/24:1) | -1.68 | (-2.24--1.13) | **5.30E-09** | **4.04E-08** | -1.23 | (-1.76--0.69) | **8.47E-06** | **2.48E-05** | same |
| Hex2Cer(d18:1/16:0) | -1.58 | (-2.1--1.05) | **6.78E-09** | **5.09E-08** | -1.17 | (-1.7--0.63) | **2.30E-05** | **6.34E-05** | same |
| Hex2Cer(d18:1/22:0) | -1.73 | (-2.29--1.17) | **2.30E-09** | **2.05E-08** | -1.43 | (-2--0.87) | **8.84E-07** | **2.91E-06** | same |
| Hex2Cer(d18:1/24:1) | -1.66 | (-2.25--1.07) | **4.96E-08** | **3.13E-07** | -1.91 | (-2.5--1.32) | **5.82E-10** | **3.21E-09** | same |
| Hex3Cer(d18:1/16:0) | -1.04 | (-1.45--0.62) | **1.58E-06** | **7.44E-06** | -1.03 | (-1.43--0.63) | **6.36E-07** | **2.21E-06** | same |
| Hex3Cer(d18:1/22:0) | -1.54 | (-2.14--0.93) | **9.28E-07** | **4.59E-06** | -2.07 | (-2.63--1.51) | **2.59E-12** | **2.22E-11** | same |
| Hex3Cer(d18:1/24:0) | -1.57 | (-2.15--0.99) | **1.64E-07** | **9.85E-07** | -1.90 | (-2.42--1.37) | **6.58E-12** | **5.18E-11** | same |
| Hex3Cer(d18:1/24:1) | -1.14 | (-1.69--0.58) | **6.61E-05** | **2.23E-04** | -1.42 | (-1.92--0.93) | **3.28E-08** | **1.42E-07** | same |
| SM(d18:0/14:0) | -0.89 | (-1.42--0.36) | **1.08E-03** | **2.70E-03** | 0.73 | (0.18-1.28) | **9.62E-03** | **1.64E-02** | opposite |
| SM(d18:1/14:0)/SM(d16:1/16:0) | -1.03 | (-1.5--0.55) | **2.66E-05** | **9.76E-05** | 0.72 | (0.27-1.18) | **1.93E-03** | **3.86E-03** | opposite |
| SM(d18:1/16:0) | -1.46 | (-1.99--0.92) | **1.89E-07** | **1.08E-06** | -0.89 | (-1.41--0.37) | **9.07E-04** | **1.95E-03** | same |
| SM(35:2) (b) | -1.53 | (-2.12--0.94) | **4.86E-07** | **2.56E-06** | -0.79 | (-1.37--0.2) | **8.48E-03** | **1.46E-02** | same |
| SM(37:2) | -1.21 | (-1.81--0.6) | **1.07E-04** | **3.46E-04** | -1.13 | (-1.68--0.57) | **7.90E-05** | **2.02E-04** | same |
| SM(d18:1/20:0)/SM(d16:1/22:0) | -0.75 | (-1.17--0.33) | **4.82E-04** | **1.32E-03** | 0.43 | (0.04-0.83) | **2.93E-02** | **4.47E-02** | opposite |
| SM(38:3) (b) | -0.77 | (-1.33--0.22) | **6.71E-03** | **1.37E-02** | -0.67 | (-1.28--0.07) | **3.00E-02** | **4.55E-02** | same |
| SM(d18:1/22:0)/SM(d16:1/24:0) | -0.59 | (-1.06--0.13) | **1.28E-02** | **2.38E-02** | 0.71 | (0.26-1.16) | **2.08E-03** | **4.13E-03** | opposite |
| SM(d16:1/24:1) | -0.70 | (-1.21--0.19) | **7.53E-03** | **1.51E-02** | 0.55 | (0.05-1.05) | **3.15E-02** | **4.75E-02** | opposite |
| SM(d18:2/23:0) | -0.93 | (-1.36--0.5) | **2.33E-05** | **8.74E-05** | -0.51 | (-0.9--0.11) | **1.15E-02** | **1.91E-02** | same |
| SM(d18:2/24:0) | -1.48 | (-1.89--1.07) | **6.63E-12** | **1.45E-10** | -0.85 | (-1.23--0.46) | **1.89E-05** | **5.27E-05** | same |
| SM(44:3) (a) | -1.42 | (-1.88--0.95) | **4.02E-09** | **3.21E-08** | -1.26 | (-1.69--0.84) | **1.15E-08** | **5.25E-08** | same |
| LPC(15-MHDA) [sn2] | -1.34 | (-2.09--0.59) | **5.17E-04** | **1.39E-03** | -1.80 | (-2.53--1.05) | **2.97E-06** | **9.21E-06** | same |
| LPC(15-MHDA) [sn1] / LPC(17:0) [sn2] | -1.22 | (-1.86--0.59) | **1.83E-04** | **5.61E-04** | -1.69 | (-2.31--1.07) | **1.33E-07** | **5.00E-07** | same |
| LPC(17:0) [sn1] | -0.89 | (-1.39--0.39) | **5.04E-04** | **1.36E-03** | -1.40 | (-1.9--0.89) | **8.98E-08** | **3.53E-07** | same |
| LPC(17:1) [sn1] (a) / LPC(17:1) [sn2] (b) | -0.95 | (-1.52--0.37) | **1.28E-03** | **3.14E-03** | -0.68 | (-1.27--0.08) | **2.64E-02** | **4.06E-02** | same |
| LPC(18:1) [sn2] | -0.59 | (-1.11--0.07) | **2.69E-02** | **4.57E-02** | -1.43 | (-2--0.86) | **1.28E-06** | **4.15E-06** | same |
| LPC(18:2) [sn2] | -0.64 | (-1.13--0.15) | **1.04E-02** | **1.97E-02** | -2.17 | (-2.7--1.63) | **2.57E-14** | **3.16E-13** | same |
| LPC(18:3) [sn2] (a) | -1.54 | (-2.23--0.85) | **1.68E-05** | **6.51E-05** | -0.86 | (-1.59--0.12) | **2.29E-02** | **3.56E-02** | same |
| LPC(19:0) [sn1] (a) / LPC(19:0) [sn2] (b) | -1.56 | (-2.02--1.09) | **1.61E-10** | **2.49E-09** | -2.28 | (-2.79--1.76) | **6.69E-17** | **1.61E-15** | same |
| LPC(20:0) [sn1] | -1.59 | (-2.03--1.14) | **1.26E-11** | **2.64E-10** | -2.10 | (-2.58--1.62) | **1.05E-16** | **2.41E-15** | same |
| LPC(20:1) [sn2] | -0.60 | (-1.07--0.14) | **1.07E-02** | **2.02E-02** | -1.58 | (-2.1--1.06) | **5.88E-09** | **2.79E-08** | same |
| LPC(20:1) [sn1] | -0.62 | (-1.08--0.15) | **9.49E-03** | **1.84E-02** | -1.86 | (-2.38--1.33) | **1.33E-11** | **9.82E-11** | same |
| LPC(20:2) [sn2] | -0.91 | (-1.41--0.4) | **4.81E-04** | **1.32E-03** | -0.80 | (-1.3--0.31) | **1.50E-03** | **3.03E-03** | same |
| LPC(20:2) [sn1] | -0.63 | (-1.11--0.16) | **9.14E-03** | **1.78E-02** | -0.94 | (-1.4--0.46) | **1.11E-04** | **2.77E-04** | same |
| LPC(20:3) [sn2] | 1.05 | (0.4-1.7) | **1.48E-03** | **3.59E-03** | 1.19 | (0.56-1.83) | **2.39E-04** | **5.67E-04** | same |
| LPC(20:3) [sn1] | 1.28 | (0.59-1.97) | **2.78E-04** | **8.08E-04** | 0.91 | (0.26-1.56) | **5.99E-03** | **1.06E-02** | same |
| LPC(20:3) [104_sn1] | 1.42 | (0.72-2.12) | **7.46E-05** | **2.47E-04** | 0.93 | (0.28-1.58) | **4.92E-03** | **8.95E-03** | same |
| LPC(20:4) [sn2] | 0.85 | (0.33-1.37) | **1.45E-03** | **3.53E-03** | -0.79 | (-1.3--0.28) | **2.64E-03** | **5.13E-03** | opposite |
| LPC(20:4) [sn1] | 1.36 | (0.79-1.93) | **3.16E-06** | **1.42E-05** | -1.16 | (-1.71--0.61) | **4.15E-05** | **1.10E-04** | opposite |
| LPC(22:5) [sn2] (n3) | -0.98 | (-1.6--0.35) | **2.21E-03** | **5.12E-03** | -0.74 | (-1.34--0.14) | **1.67E-02** | **2.68E-02** | same |
| LPC(22:5) [sn1] (n3)/LPC(22:5) [sn2] (n6) | -0.84 | (-1.42--0.26) | **4.63E-03** | **9.97E-03** | -0.82 | (-1.43--0.21) | **8.19E-03** | **1.42E-02** | same |
| LPC(O-18:0) | -1.05 | (-1.59--0.5) | **1.83E-04** | **5.61E-04** | -1.53 | (-2.07--0.99) | **5.02E-08** | **2.13E-07** | same |
| LPC(O-18:1) | 0.52 | (0.07-0.97) | **2.38E-02** | **4.13E-02** | -0.80 | (-1.25--0.35) | **5.53E-04** | **1.23E-03** | opposite |
| LPC(O-20:0) | -1.70 | (-2.21--1.19) | **1.86E-10** | **2.70E-09** | -2.23 | (-2.77--1.69) | **4.19E-15** | **6.29E-14** | same |
| LPC(O-22:0) | -2.40 | (-2.88--1.91) | **1.69E-20** | **2.02E-18** | -1.43 | (-1.85--1) | **1.03E-10** | **6.79E-10** | same |
| LPC(O-22:1) | -1.62 | (-2.15--1.1) | **2.73E-09** | **2.33E-08** | -2.10 | (-2.63--1.56) | **9.90E-14** | **1.16E-12** | same |
| LPC(O-24:0) | -3.04 | (-3.62--2.45) | **1.56E-22** | **7.48E-20** | -1.17 | (-1.62--0.71) | **7.24E-07** | **2.45E-06** | same |
| LPC(O-24:1) | -2.43 | (-2.98--1.87) | **1.19E-16** | **8.16E-15** | -1.37 | (-1.84--0.9) | **1.75E-08** | **7.86E-08** | same |
| LPC(O-24:2) | -1.94 | (-2.53--1.34) | **3.86E-10** | **4.64E-09** | -2.07 | (-2.6--1.53) | **1.68E-13** | **1.79E-12** | same |
| LPC(P-18:1) | 0.95 | (0.36-1.54) | **1.57E-03** | **3.78E-03** | -0.88 | (-1.4--0.35) | **1.14E-03** | **2.40E-03** | opposite |
| LPC(P-20:0) | -0.73 | (-1.3--0.15) | **1.34E-02** | **2.43E-02** | -1.56 | (-2.13--0.99) | **1.23E-07** | **4.68E-07** | same |
| LPE(16:0) [sn2] | -0.58 | (-1.09--0.08) | **2.45E-02** | **4.23E-02** | -1.40 | (-1.94--0.85) | **8.49E-07** | **2.81E-06** | same |
| PC(31:1) | -2.56 | (-3.45--1.67) | **3.05E-08** | **2.06E-07** | 1.45 | (0.56-2.34) | **1.35E-03** | **2.79E-03** | opposite |
| PC(33:2) | -2.27 | (-2.89--1.64) | **6.00E-12** | **1.37E-10** | -0.81 | (-1.42--0.19) | **1.07E-02** | **1.78E-02** | same |
| PC(16:0_18:2) | -0.34 | (-0.64--0.04) | **2.86E-02** | **4.82E-02** | -0.56 | (-0.98--0.15) | **7.64E-03** | **1.33E-02** | same |
| PC(15-MHDA_18:2) | -1.86 | (-2.67--1.05) | **9.80E-06** | **4.05E-05** | -1.20 | (-1.96--0.42) | **2.53E-03** | **4.94E-03** | same |
| PC(17:0_18:2) | -1.54 | (-2.08--1) | **3.86E-08** | **2.57E-07** | -1.17 | (-1.68--0.65) | **1.30E-05** | **3.74E-05** | same |
| PC 36:0 | 0.97 | (0.36-1.59) | **2.00E-03** | **4.66E-03** | 3.16 | (2.47-3.84) | **7.15E-19** | **3.43E-17** | same |
| PC(18:0_18:1) | -1.34 | (-1.89--0.78) | **3.35E-06** | **1.49E-05** | 0.98 | (0.36-1.59) | **1.86E-03** | **3.74E-03** | opposite |

| | | | | | | | | | |
|---|---|---|---|---|---|---|---|---|---|
| PC(16:0_20:3) (a) | -1.23 | (-1.82--0.64) | **5.06E-05** | **1.75E-04** | -1.61 | (-2.18--1.04) | **5.63E-08** | **2.33E-07** | same |
| PC(18:2_18:2) | -2.57 | (-3.28--1.87) | **3.69E-12** | **9.32E-11** | -2.49 | (-3.22--1.75) | **1.24E-10** | **7.80E-10** | same |
| PC(15:0_22:6) | -2.68 | (-3.42--1.94) | **3.93E-12** | **9.43E-11** | -0.93 | (-1.62--0.24) | **8.87E-03** | **1.52E-02** | same |
| PC(38:2) | -1.13 | (-1.58--0.67) | **1.45E-06** | **6.94E-06** | 0.67 | (0.19-1.14) | **5.76E-03** | **1.03E-02** | opposite |
| PC(18:0_20:3) | 1.23 | (0.54-1.93) | **5.29E-04** | **1.41E-03** | 3.69 | (2.93-4.46) | **1.54E-20** | **1.06E-18** | same |
| PC(38:4) (b) | -0.97 | (-1.66--0.28) | **5.93E-03** | **1.24E-02** | 1.29 | (0.57-2.02) | **4.90E-04** | **1.11E-03** | opposite |
| PC(38:6) (a) | -1.01 | (-1.61--0.42) | **9.39E-04** | **2.36E-03** | -1.09 | (-1.67--0.5) | **2.90E-04** | **6.81E-04** | same |
| PC(38:7)(c) | -1.58 | (-2.17--0.99) | **2.34E-07** | **1.32E-06** | -0.64 | (-1.21--0.06) | **2.93E-02** | **4.47E-02** | same |
| PC(39:5)(a) | -2.53 | (-3.35--1.7) | **4.56E-09** | **3.53E-08** | -1.82 | (-2.59--1.05) | **4.96E-06** | **1.51E-05** | same |
| PC(39:5)(b) | -2.31 | (-3.04--1.58) | **1.68E-09** | **1.56E-08** | -0.86 | (-1.58--0.13) | **2.03E-02** | **3.21E-02** | same |
| PC(15-MHDA_22:6) | -2.51 | (-3.31--1.7) | **2.77E-09** | **2.33E-08** | -1.62 | (-2.33--0.91) | **1.10E-05** | **3.17E-05** | same |
| PC(17:0_22:6) | -2.00 | (-2.74--1.26) | **1.87E-07** | **1.08E-06** | -1.05 | (-1.79--0.31) | **5.44E-03** | **9.82E-03** | same |
| PC(18:0_22:5) (n6) | -1.32 | (-2.07--0.57) | **6.32E-04** | **1.67E-03** | 1.17 | (0.37-1.97) | **3.94E-03** | **7.42E-03** | opposite |
| PC(18:1_22:6) (a) | -1.70 | (-2.24--1.16) | **1.69E-09** | **1.56E-08** | -0.86 | (-1.44--0.27) | **4.10E-03** | **7.62E-03** | same |
| PC(40:8) | -1.77 | (-2.3--1.24) | **1.86E-10** | **2.70E-09** | -1.88 | (-2.39--1.37) | **2.15E-12** | **1.91E-11** | same |
| PC(44:12) | -2.40 | (-3.44--1.34) | **1.14E-05** | **4.60E-05** | -2.61 | (-3.74--1.46) | **1.06E-05** | **3.09E-05** | same |
| PC(O-16:0/16:0) | -0.78 | (-1.21--0.35) | **3.82E-04** | **1.09E-03** | -0.75 | (-1.18--0.33) | **5.02E-04** | **1.13E-03** | same |
| PC(O-34:1) | -0.84 | (-1.27--0.41) | **1.54E-04** | **4.84E-04** | -1.21 | (-1.62--0.79) | **2.44E-08** | **1.08E-07** | same |
| PC(O-34:2) | -2.02 | (-2.62--1.41) | **2.10E-10** | **2.80E-09** | -1.75 | (-2.35--1.15) | **2.36E-08** | **1.05E-07** | same |
| PC(O-36:0) | -1.30 | (-1.8--0.81) | **3.88E-07** | **2.12E-06** | -0.84 | (-1.33--0.35) | **9.07E-04** | **1.95E-03** | same |
| PC(O-18:0/18:1) | -1.51 | (-1.98--1.03) | **1.00E-09** | **1.02E-08** | -1.48 | (-1.92--1.04) | **1.07E-10** | **6.87E-10** | same |
| PC(O-18:1/18:1) | -1.20 | (-1.66--0.74) | **4.16E-07** | **2.24E-06** | -1.71 | (-2.13--1.29) | **1.10E-14** | **1.42E-13** | same |
| PC(O-18:0/18:2) | -2.36 | (-2.92--1.79) | **3.60E-15** | **1.73E-13** | -1.89 | (-2.38--1.39) | **4.43E-13** | **4.43E-12** | same |
| PC(O-18:1/18:2) | -1.41 | (-1.96--0.86) | **7.43E-07** | **3.71E-06** | -1.34 | (-1.85--0.82) | **6.84E-07** | **2.35E-06** | same |
| PC(O-40:5) | -0.80 | (-1.3--0.3) | **1.67E-03** | **3.98E-03** | -0.88 | (-1.34--0.42) | **1.90E-04** | **4.60E-04** | same |
| PC(O-18:0/22:6) | -0.69 | (-1.25--0.12) | **1.77E-02** | **3.15E-02** | -1.09 | (-1.63--0.55) | **9.15E-05** | **2.31E-04** | same |
| PC(P-16:0/14:0) | -1.27 | (-1.86--0.68) | **2.71E-05** | **9.86E-05** | -1.18 | (-1.79--0.56) | **1.98E-04** | **4.77E-04** | same |
| PC(P-16:0/16:0) | -0.71 | (-1.21--0.2) | **6.69E-03** | **1.37E-02** | -1.03 | (-1.54--0.53) | **6.60E-05** | **1.72E-04** | same |
| PC(P-16:0/16:1) | -0.71 | (-1.31--0.11) | **2.01E-02** | **3.52E-02** | -0.83 | (-1.39--0.26) | **4.66E-03** | **8.63E-03** | same |
| PC(P-16:0/18:0) | -1.20 | (-1.59--0.81) | **3.58E-09** | **2.91E-08** | -1.47 | (-1.84--1.09) | **1.05E-13** | **1.20E-12** | same |
| PC(P-16:0/18:1) | -1.21 | (-1.75--0.67) | **1.36E-05** | **5.30E-05** | -2.19 | (-2.7--1.69) | **2.94E-16** | **6.14E-15** | same |
| PC(P-16:0/18:2) | -1.10 | (-1.68--0.5) | **3.02E-04** | **8.73E-04** | -2.03 | (-2.57--1.48) | **1.39E-12** | **1.28E-11** | same |
| PC(P-16:0/18:3) | -1.64 | (-2.27--1.01) | **5.09E-07** | **2.66E-06** | -1.42 | (-1.99--0.84) | **2.28E-06** | **7.17E-06** | same |
| PC(P-35:2) (a) | -2.26 | (-3.13--1.39) | **5.70E-07** | **2.91E-06** | -1.87 | (-2.71--1.02) | **1.87E-05** | **5.25E-05** | same |
| PC(P-35:2) (b) | -1.83 | (-2.55--1.1) | **1.07E-06** | **5.20E-06** | -2.14 | (-2.83--1.45) | **2.27E-09** | **1.13E-08** | same |
| PC(P-18:1/18:1) | -1.20 | (-1.72--0.68) | **7.54E-06** | **3.17E-05** | -1.58 | (-2.08--1.09) | **7.85E-10** | **4.18E-09** | same |
| PC(P-18:0/18:2) | -1.93 | (-2.51--1.36) | **1.35E-10** | **2.16E-09** | -2.58 | (-3.1--2.06) | **1.54E-20** | **1.06E-18** | same |
| PC(P-38:5) (b) | -0.85 | (-1.43--0.27) | **4.44E-03** | **9.70E-03** | -1.10 | (-1.67--0.53) | **1.76E-04** | **4.33E-04** | same |
| PC(P-40:5) (b) | -0.80 | (-1.37--0.24) | **5.56E-03** | **1.18E-02** | -1.41 | (-1.93--0.9) | **1.16E-07** | **4.47E-07** | same |
| PE(18:0_20:3) (a) | 1.68 | (0.87-2.5) | **4.94E-05** | **1.72E-04** | 2.90 | (1.96-3.84) | **1.44E-09** | **7.41E-09** | same |
| PE(18:0_22:6) | 1.34 | (0.55-2.14) | **8.34E-04** | **2.13E-03** | 1.67 | (0.73-2.63) | **5.37E-04** | **1.20E-03** | same |
| PE(O-34:1) | -1.42 | (-2--0.83) | **2.58E-06** | **1.19E-05** | -1.43 | (-2.04--0.82) | **6.33E-06** | **1.91E-05** | same |
| PE(O-16:0/18:2) | -2.74 | (-3.5--1.96) | **1.39E-11** | **2.79E-10** | -2.48 | (-3.27--1.68) | **2.70E-09** | **1.32E-08** | same |
| PE(O-18:1/18:2) | -1.56 | (-2.3--0.82) | **4.20E-05** | **1.47E-04** | -1.10 | (-1.81--0.38) | **2.71E-03** | **5.25E-03** | same |
| PE(O-16:0/20:4) | -1.39 | (-2.12--0.65) | **2.38E-04** | **7.02E-04** | -1.38 | (-2.13--0.61) | **4.51E-04** | **1.03E-03** | same |
| PE(O-18:0/20:4) | -1.81 | (-2.63--0.99) | **2.02E-05** | **7.69E-05** | -1.76 | (-2.55--0.97) | **1.81E-05** | **5.11E-05** | same |
| PE(O-38:5) (a) | -1.44 | (-2.16--0.71) | **1.14E-04** | **3.65E-04** | -0.89 | (-1.6--0.18) | **1.37E-02** | **2.25E-02** | same |
| PE(O-38:5) (b) | -1.98 | (-2.64--1.31) | **9.88E-09** | **6.87E-08** | -1.06 | (-1.81--0.29) | **6.73E-03** | **1.19E-02** | same |
| PE(O-16:0/22:6) | -1.03 | (-1.66--0.4) | **1.52E-03** | **3.67E-03** | -0.83 | (-1.53--0.13) | **2.05E-02** | **3.21E-02** | same |
| PE(O-18:0/22:6) | -1.54 | (-2.14--0.93) | **1.02E-06** | **5.01E-06** | -1.41 | (-2.04--0.77) | **1.78E-05** | **5.04E-05** | same |
| PE(O-18:1/22:6) | -1.16 | (-1.76--0.56) | **1.73E-04** | **5.35E-04** | -1.29 | (-1.9--0.67) | **5.13E-05** | **1.35E-04** | same |
| PE(P-16:0/18:1) | -1.04 | (-1.56--0.52) | **1.03E-04** | **3.35E-04** | -0.62 | (-1.13--0.11) | **1.76E-02** | **2.81E-02** | same |
| PE(P-18:0/18:1) | -1.25 | (-1.84--0.66) | **3.98E-05** | **1.40E-04** | -1.01 | (-1.6--0.41) | **9.72E-04** | **2.07E-03** | same |
| PE(P-18:0/18:2) | -1.83 | (-2.48--1.19) | **4.04E-08** | **2.66E-07** | -1.53 | (-2.15--0.91) | **1.62E-06** | **5.22E-06** | same |
| PE(P-18:0/22:4) | -0.72 | (-1.31--0.13) | **1.74E-02** | **3.11E-02** | -0.98 | (-1.57--0.38) | **1.48E-03** | **3.02E-03** | same |
| PE(P-18:0/22:5) (n6) | -1.29 | (-1.87--0.72) | **1.35E-05** | **5.30E-05** | -1.25 | (-1.83--0.67) | **3.16E-05** | **8.48E-05** | same |
| PE(P-18:0/22:6) | -0.79 | (-1.43--0.14) | **1.80E-02** | **3.18E-02** | -1.21 | (-1.87--0.55) | **3.71E-04** | **8.65E-04** | same |
| PE(P-18:1/18:1) | -0.90 | (-1.45--0.35) | **1.43E-03** | **3.50E-03** | -0.83 | (-1.38--0.28) | **3.06E-03** | **5.83E-03** | same |
| PE(P-18:1/18:2) | -1.86 | (-2.56--1.16) | **3.27E-07** | **1.80E-06** | -1.22 | (-1.9--0.53) | **5.96E-04** | **1.30E-03** | same |
| PE(P-20:0/18:1) | -1.27 | (-1.89--0.65) | **7.01E-05** | **2.35E-04** | -1.14 | (-1.71--0.57) | **1.01E-04** | **2.52E-04** | same |

| | | | | | | | | | |
|---|---|---|---|---|---|---|---|---|---|
| PE(P-20:0/18:2) | -1.81 | (-2.36--1.25) | **3.40E-10** | **4.18E-09** | -1.98 | (-2.5--1.45) | **1.04E-12** | **9.79E-12** | same |
| PE(P-20:0/22:6) | -0.78 | (-1.31--0.26) | **3.71E-03** | **8.27E-03** | -1.89 | (-2.39--1.4) | **3.83E-13** | **3.92E-12** | same |
| PG(34:2) | -1.07 | (-1.82--0.31) | **6.14E-03** | **1.28E-02** | 1.17 | (0.21-2.14) | **1.72E-02** | **2.76E-02** | opposite |
| PI(16:0/16:0) | -2.44 | (-3.37--1.51) | **4.46E-07** | **2.38E-06** | 1.75 | (0.7-2.81) | **1.12E-03** | **2.36E-03** | opposite |
| PI(16:0_16:1) | -1.73 | (-2.79--0.66) | **1.69E-03** | **4.01E-03** | 3.85 | (2.77-4.94) | **3.76E-12** | **3.11E-11** | opposite |
| PI(18:1_18:2) | -2.66 | (-3.29--2.03) | **1.11E-15** | **5.91E-14** | -1.29 | (-1.97--0.61) | **2.16E-04** | **5.17E-04** | same |
| PI(16:0_20:3) (a) | -2.16 | (-2.82--1.5) | **3.96E-10** | **4.64E-09** | 0.93 | (0.22-1.65) | **1.07E-02** | **1.78E-02** | opposite |
| PI(16:0_20:3) (b) | -2.46 | (-3.2--1.72) | **1.99E-10** | **2.80E-09** | 0.86 | (0.13-1.6) | **2.06E-02** | **3.21E-02** | opposite |
| PI(16:0_20:4) | -1.45 | (-2.03--0.86) | **1.88E-06** | **8.78E-06** | 0.99 | (0.36-1.62) | **2.13E-03** | **4.20E-03** | opposite |
| PI(38:6) | -2.63 | (-3.27--1.98) | **1.30E-14** | **5.19E-13** | -0.89 | (-1.56--0.21) | **1.01E-02** | **1.70E-02** | same |
| PI(18:0_22:5) (n3) | -1.73 | (-2.4--1.06) | **5.69E-07** | **2.91E-06** | 0.86 | (0.2-1.52) | **1.06E-02** | **1.78E-02** | opposite |

**Supplementary Table 4b.** The overlapping lipid species between the ppBMI and birth weight studies. Related to Figure 3.

| Lipid Sepcies | ppBMI (antenatal) | | | | Birth Weight (cord blood) | | | | Trend |
|---|---|---|---|---|---|---|---|---|---|
| | %change in lipid concentration per unit BMI | 95%CI | p-value | p-value(BH) | %change in lipid concentration per 100 grams | 95%CI | p-value | p-value(BH) | |
| AC(14:1) | 1.38 | (0.22-2.56) | **1.98E-02** | **3.50E-02** | -1.08 | (-1.73--0.37) | **3.63E-03** | **8.13E-03** | opposite |
| AC(16:1) | 1.19 | (0.37-2.02) | **4.39E-03** | **9.66E-03** | -1.00 | (-1.64--0.31) | **5.17E-03** | **1.12E-02** | opposite |
| CE(16:1) | -1.06 | (-1.82--0.29) | **7.23E-03** | **1.46E-02** | 0.98 | (0.31-1.69) | **3.55E-03** | **8.00E-03** | opposite |
| CE(20:2) | -1.33 | (-1.74--0.91) | **7.59E-10** | **7.92E-09** | 0.79 | (0.16-1.46) | **1.36E-02** | **2.71E-02** | opposite |
| CE(20:5) | -1.71 | (-3.04--0.36) | **1.31E-02** | **2.41E-02** | -2.42 | (-3.39--1.3) | **9.07E-05** | **2.59E-04** | same |
| CE(22:6) | -1.55 | (-2.27--0.82) | **3.29E-05** | **1.18E-04** | -1.97 | (-2.73--1.12) | **1.95E-05** | **6.12E-05** | same |
| CE(24:5) | -2.01 | (-2.72--1.3) | **4.73E-08** | **3.03E-07** | -2.88 | (-3.68--1.98) | **3.40E-08** | **1.41E-07** | same |
| CE(24:6) | -2.55 | (-3.28--1.82) | **2.50E-11** | **4.44E-10** | -1.14 | (-2.05--0.14) | **2.66E-02** | **4.98E-02** | same |
| DG(16:0_18:1) | 1.33 | (0.59-2.07) | **4.36E-04** | **1.22E-03** | -1.91 | (-2.57--1.19) | **1.35E-06** | **4.62E-06** | opposite |
| DG(16:0_18:2) | 0.92 | (0.21-1.65) | **1.15E-02** | **2.16E-02** | -3.07 | (-3.76--2.31) | **1.09E-11** | **6.22E-11** | opposite |
| DG(18:0_18:1) | 1.36 | (0.69-2.04) | **7.91E-05** | **2.60E-04** | -1.29 | (-1.99--0.53) | **1.23E-03** | **2.94E-03** | opposite |
| DG(18:1_18:1) | 1.37 | (0.64-2.1) | **2.23E-04** | **6.69E-04** | -1.85 | (-2.5--1.16) | **1.20E-06** | **4.16E-06** | opposite |
| DG(18:0_18:2) | 0.99 | (0.23-1.76) | **1.04E-02** | **1.97E-02** | -1.62 | (-2.52--0.61) | **2.43E-03** | **5.62E-03** | opposite |
| DG(18:1_18:2) | 0.95 | (0.22-1.68) | **1.02E-02** | **1.97E-02** | -3.15 | (-3.76--2.48) | **8.45E-15** | **7.12E-14** | opposite |
| DG(16:0_20:4) | 1.19 | (0.26-2.12) | **1.22E-02** | **2.27E-02** | -2.06 | (-2.83--1.21) | **9.20E-06** | **2.95E-05** | opposite |
| DG(18:1_20:4) | 1.14 | (0.4-1.88) | **2.42E-03** | **5.53E-03** | -2.06 | (-2.71--1.36) | **1.08E-07** | **4.14E-07** | opposite |
| DG(18:1_22:6) | 1.53 | (0.47-2.61) | **4.63E-03** | **9.97E-03** | -3.49 | (-4.15--2.76) | **7.44E-15** | **6.49E-14** | opposite |
| DG(18:2_22:6) | 1.52 | (0.42-2.63) | **6.55E-03** | **1.35E-02** | -4.00 | (-4.74--3.16) | **9.00E-14** | **6.75E-13** | opposite |
| TG(48:0) [NL-18:0] | -1.56 | (-2.43--0.68) | **5.76E-04** | **1.53E-03** | -1.44 | (-2.28--0.51) | **3.14E-03** | **7.14E-03** | same |
| TG(48:2) [NL-14:1] | -1.20 | (-2.25--0.13) | **2.78E-02** | **4.70E-02** | -3.06 | (-3.81--2.23) | **4.50E-10** | **2.23E-09** | same |
| TG(48:2) [NL-18:2] | -1.16 | (-2.2--0.12) | **2.91E-02** | **4.87E-02** | -4.14 | (-4.86--3.32) | **5.42E-15** | **4.91E-14** | same |
| TG(48:3) [NL-14:0] | -1.51 | (-2.64--0.37) | **9.55E-03** | **1.85E-02** | -4.04 | (-4.73--3.26) | **1.02E-15** | **1.04E-14** | same |
| TG(48:3) [NL-18:3] | -1.81 | (-3.07--0.53) | **5.76E-03** | **1.21E-02** | -3.23 | (-3.91--2.48) | **8.75E-13** | **5.83E-12** | same |
| TG(50:1) [NL-16:0] | 0.83 | (0.22-1.43) | **7.27E-03** | **1.46E-02** | -2.49 | (-3.25--1.64) | **2.17E-07** | **8.15E-07** | opposite |
| TG(50:1) [NL-18:1] | 0.76 | (0.18-1.34) | **1.03E-02** | **1.97E-02** | -2.47 | (-3.24--1.62) | **2.61E-07** | **9.63E-07** | opposite |
| TG(50:2) [NL-14:0] | -0.65 | (-1.21--0.07) | **2.69E-02** | **4.57E-02** | -3.34 | (-4.1--2.48) | **1.16E-10** | **6.01E-10** | same |
| TG(50:2) [NL-18:2] | 0.66 | (0.08-1.24) | **2.58E-02** | **4.41E-02** | -4.17 | (-4.92--3.31) | **5.26E-14** | **4.14E-13** | opposite |
| TG(52:1) [NL-18:0] | 1.38 | (0.59-2.18) | **6.58E-04** | **1.73E-03** | -1.86 | (-2.76--0.83) | **6.81E-04** | **1.69E-03** | opposite |
| TG(52:1) [NL-18:1] | 1.45 | (0.69-2.23) | **2.04E-04** | **6.17E-04** | -1.80 | (-2.71--0.77) | **9.96E-04** | **2.41E-03** | opposite |
| TG(52:2) [NL-16:0] | 0.80 | (0.4-1.21) | **1.01E-04** | **3.29E-04** | -2.79 | (-3.44--2.07) | **3.12E-11** | **1.68E-10** | opposite |
| TG(52:3) [NL-18:2] | 0.76 | (0.32-1.2) | **8.08E-04** | **2.08E-03** | -4.40 | (-5.07--3.64) | **3.04E-18** | **4.87E-17** | opposite |
| TG(52:4) [NL-18:2] | 1.20 | (0.44-1.96) | **1.92E-03** | **4.52E-03** | -4.97 | (-5.63--4.21) | **2.07E-20** | **5.23E-19** | opposite |
| TG(53:2) [NL-18:1] | 0.88 | (0.2-1.56) | **1.08E-02** | **2.03E-02** | -2.69 | (-3.33--1.98) | **5.26E-11** | **2.81E-10** | opposite |
| TG(54:1) [NL-18:1] | 0.99 | (0.32-1.66) | **3.57E-03** | **8.00E-03** | -1.64 | (-2.56--0.59) | **2.89E-03** | **6.61E-03** | opposite |
| TG(54:2) [NL-18:0] | 1.41 | (0.72-2.1) | **6.55E-05** | **2.23E-04** | -1.85 | (-2.69--0.91) | **2.40E-04** | **6.39E-04** | opposite |
| TG(54:3) [NL-18:2] | 0.72 | (0.14-1.3) | **1.43E-02** | **2.59E-02** | -3.06 | (-3.75--2.3) | **1.28E-11** | **7.23E-11** | opposite |
| TG(54:5) [NL-20:4] | 1.04 | (0.38-1.71) | **1.91E-03** | **4.51E-03** | -3.73 | (-4.49--2.86) | **4.40E-12** | **2.64E-11** | opposite |
| TG(54:6) [NL-20:4] | 1.16 | (0.33-2) | **6.24E-03** | **1.29E-02** | -4.11 | (-4.83--3.28) | **1.19E-14** | **9.81E-14** | opposite |
| TG(54:6) [NL-22:6] | 1.79 | (0.57-3.02) | **4.10E-03** | **9.06E-03** | -5.16 | (-5.83--4.38) | **1.98E-20** | **5.23E-19** | opposite |
| TG(56:6) [NL-20:4] | 1.34 | (0.81-1.87) | **7.12E-07** | **3.60E-06** | -3.00 | (-3.77--2.15) | **1.89E-09** | **8.92E-09** | opposite |
| TG(56:7) [NL-20:4] | 1.02 | (0.35-1.7) | **2.84E-03** | **6.40E-03** | -3.66 | (-4.41--2.81) | **3.38E-12** | **2.08E-11** | opposite |
| TG(56:7) [NL-22:6] | 1.93 | (0.84-3.04) | **4.97E-04** | **1.35E-03** | -5.07 | (-5.72--4.33) | **1.85E-21** | **7.12E-20** | opposite |
| TG(56:8) [NL-22:6] | 1.76 | (0.54-2.99) | **4.49E-03** | **9.76E-03** | -5.36 | (-5.97--4.65) | **1.69E-24** | **1.02E-22** | opposite |
| TG(58:8) [NL-22:6] | 1.46 | (0.38-2.56) | **7.98E-03** | **1.58E-02** | -4.50 | (-5.16--3.76) | **3.67E-19** | **7.05E-18** | opposite |
| TG(O-54:2) [NL-18:1] | 0.85 | (0.18-1.53) | **1.33E-02** | **2.43E-02** | -2.66 | (-3.31--1.95) | **1.06E-10** | **5.53E-10** | opposite |
| Cer(d16:1/22:0) | -0.73 | (-1.33--0.13) | **1.79E-02** | **3.18E-02** | -1.03 | (-1.75--0.25) | **1.07E-02** | **2.19E-02** | same |
| Cer(d17:1/24:0) | -2.19 | (-2.76--1.61) | **4.35E-13** | **1.39E-11** | -1.07 | (-1.78--0.28) | **8.53E-03** | **1.79E-02** | same |
| Cer(d19:1/24:0) | -1.22 | (-2.12--0.32) | **8.01E-03** | **1.58E-02** | -1.23 | (-2.03--0.34) | **7.85E-03** | **1.67E-02** | same |
| Cer(m18:0/24:1) | 1.15 | (0.26-2.04) | **1.11E-02** | **2.08E-02** | -1.46 | (-2.42--0.37) | **9.81E-03** | **2.02E-02** | opposite |
| Cer(m18:1/24:1) | 1.37 | (0.47-2.27) | **2.70E-03** | **6.11E-03** | -2.16 | (-2.91--1.32) | **3.30E-06** | **1.10E-05** | opposite |
| GM3(d18:1/18:0) | -1.16 | (-1.7--0.62) | **2.65E-05** | **9.76E-05** | -0.78 | (-1.37--0.16) | **1.53E-02** | **3.01E-02** | same |
| SM(d17:1/14:0) | -1.88 | (-2.45--1.3) | **3.06E-10** | **3.86E-09** | -0.98 | (-1.49--0.44) | **5.42E-04** | **1.36E-03** | same |
| SM(d17:1/16:0) | -1.40 | (-1.85--0.95) | **2.57E-09** | **2.25E-08** | -0.98 | (-1.47--0.47) | **2.86E-04** | **7.59E-04** | same |
| SM(37:1) | -0.75 | (-1.3--0.2) | **7.20E-03** | **1.46E-02** | -0.82 | (-1.45--0.15) | **1.76E-02** | **3.42E-02** | same |
| SM(37:2) | -1.21 | (-1.81--0.6) | **1.07E-04** | **3.46E-04** | -0.76 | (-1.35--0.12) | **2.06E-02** | **3.93E-02** | same |
| SM(40:3) (b) | -0.91 | (-1.46--0.36) | **1.23E-03** | **3.06E-03** | -0.89 | (-1.5--0.24) | **8.17E-03** | **1.72E-02** | same |

| | | | | | | | | | |
|---|---|---|---|---|---|---|---|---|---|
| SM(43:1) | -1.62 | (-2.09--1.14) | **7.20E-11** | **1.19E-09** | -1.04 | (-1.74--0.27) | **9.36E-03** | **1.94E-02** | same |
| LPC(15:0) [sn2] | -1.50 | (-2.11--0.88) | **2.82E-06** | **1.29E-05** | 1.04 | (0.46-1.66) | **3.62E-04** | **9.45E-04** | opposite |
| LPC(15:0) [sn1] | -1.28 | (-1.83--0.72) | **8.57E-06** | **3.58E-05** | 0.76 | (0.24-1.32) | **4.17E-03** | **9.24E-03** | opposite |
| LPC(17:1) [sn1] (a) / LPC(17:1) [sn2] (b) | -0.95 | (-1.52--0.37) | **1.28E-03** | **3.14E-03** | 1.72 | (1.09-2.4) | **3.10E-08** | **1.29E-07** | opposite |
| LPC(18:1) [sn2] | -0.59 | (-1.11--0.07) | **2.69E-02** | **4.57E-02** | 3.61 | (2.91-4.34) | **4.61E-28** | **5.53E-26** | opposite |
| LPC(18:2) [sn2] | -0.64 | (-1.13--0.15) | **1.04E-02** | **1.97E-02** | 2.93 | (2.22-3.69) | **4.36E-18** | **6.75E-17** | opposite |
| LPC(18:3) [sn2] (a) | -1.54 | (-2.23--0.85) | **1.68E-05** | **6.51E-05** | 2.31 | (1.65-3.01) | **5.03E-13** | **3.55E-12** | opposite |
| LPC(18:3) [sn1] (a)/LPC(18:3) [sn2] (b) | -0.95 | (-1.67--0.23) | **9.74E-03** | **1.88E-02** | 4.43 | (3.51-5.42) | **1.76E-25** | **1.41E-23** | opposite |
| LPC(18:3) (a) [sn1] [104_sn1] | -0.94 | (-1.68--0.2) | **1.34E-02** | **2.43E-02** | 4.09 | (3.21-5.02) | **1.27E-23** | **6.77E-22** | opposite |
| LPC(20:1) [sn2] | -0.60 | (-1.07--0.14) | **1.07E-02** | **2.02E-02** | 1.44 | (0.92-1.98) | **1.54E-08** | **6.60E-08** | opposite |
| LPC(20:1) [sn1] | -0.62 | (-1.08--0.15) | **9.49E-03** | **1.84E-02** | 1.75 | (1.27-2.25) | **1.55E-13** | **1.12E-12** | opposite |
| LPC(20:2) [sn2] | -0.91 | (-1.41--0.4) | **4.81E-04** | **1.32E-03** | 3.83 | (3.05-4.65) | **3.73E-26** | **3.58E-24** | opposite |
| LPC(20:2) [sn1] | -0.63 | (-1.11--0.16) | **9.14E-03** | **1.78E-02** | 3.35 | (2.66-4.06) | **4.61E-25** | **3.16E-23** | opposite |
| LPC(20:3) [sn2] | 1.05 | (0.4-1.7) | **1.48E-03** | **3.59E-03** | 2.58 | (1.91-3.3) | **1.79E-15** | **1.72E-14** | same |
| LPC(20:3) [sn1] | 1.28 | (0.59-1.97) | **2.78E-04** | **8.08E-04** | 3.27 | (2.53-4.05) | **7.83E-21** | **2.50E-19** | same |
| LPC(20:3) [104_sn1] | 1.42 | (0.72-2.12) | **7.46E-05** | **2.47E-04** | 3.22 | (2.49-3.99) | **9.49E-21** | **2.85E-19** | same |
| LPC(20:4) [sn2] | 0.85 | (0.33-1.37) | **1.45E-03** | **3.53E-03** | 2.39 | (1.75-3.06) | **6.31E-15** | **5.61E-14** | same |
| LPC(20:4) [sn1] | 1.36 | (0.79-1.93) | **3.16E-06** | **1.42E-05** | 2.66 | (1.98-3.38) | **2.33E-16** | **2.60E-15** | same |
| LPC(22:5) [sn2] (n3) | -0.98 | (-1.6--0.35) | **2.21E-03** | **5.12E-03** | 1.73 | (0.89-2.63) | **2.63E-05** | **8.04E-05** | opposite |
| LPC(22:5) [sn1] (n3)/LPC(22:5) [sn2] (n6) | -0.84 | (-1.42--0.26) | **4.63E-03** | **9.97E-03** | 2.12 | (1.41-2.88) | **6.79E-10** | **3.29E-09** | opposite |
| LPC(O-18:1) | 0.52 | (0.07-0.97) | **2.38E-02** | **4.13E-02** | 1.82 | (1.3-2.37) | **7.59E-13** | **5.13E-12** | same |
| LPC(O-22:1) | -1.62 | (-2.15--1.1) | **2.73E-09** | **2.33E-08** | 1.29 | (0.72-1.9) | **6.07E-06** | **1.97E-05** | opposite |
| LPC(P-18:1) | 0.95 | (0.36-1.54) | **1.57E-03** | **3.78E-03** | 1.25 | (0.59-1.95) | **1.32E-04** | **3.64E-04** | same |
| LPC(P-20:0) | -0.73 | (-1.3--0.15) | **1.34E-02** | **2.43E-02** | 2.02 | (1.36-2.71) | **3.08E-10** | **1.54E-09** | opposite |
| LPE(16:0) [sn2] | -0.58 | (-1.09--0.08) | **2.45E-02** | **4.23E-02** | 2.24 | (1.6-2.92) | **6.15E-13** | **4.28E-12** | opposite |
| LPE(18:1) [sn2] | -0.78 | (-1.33--0.24) | **5.15E-03** | **1.10E-02** | 2.54 | (1.88-3.24) | **1.28E-15** | **1.28E-14** | opposite |
| LPE(18:1) [sn1] | -0.74 | (-1.28--0.2) | **7.22E-03** | **1.46E-02** | 2.94 | (2.26-3.65) | **1.02E-19** | **2.33E-18** | opposite |
| LPE(18:2) [sn2] | -0.93 | (-1.59--0.27) | **6.21E-03** | **1.29E-02** | 3.65 | (2.71-4.66) | **9.44E-17** | **1.10E-15** | opposite |
| LPE(18:2) [sn1] | -0.96 | (-1.63--0.29) | **5.28E-03** | **1.12E-02** | 3.74 | (2.85-4.69) | **1.99E-19** | **4.16E-18** | opposite |
| PC(33:0) (b) | -1.78 | (-2.34--1.22) | **1.18E-09** | **1.18E-08** | -1.45 | (-2.17--0.66) | **5.09E-04** | **1.30E-03** | same |
| PC(33:2) | -2.27 | (-2.89--1.64) | **6.00E-12** | **1.37E-10** | -1.25 | (-1.87--0.6) | **3.06E-04** | **8.08E-04** | same |
| PC(15-MHDA_18:1) | -2.51 | (-3.31--1.72) | **1.39E-09** | **1.34E-08** | -1.26 | (-1.99--0.45) | **2.77E-03** | **6.37E-03** | same |
| PC(17:0_18:1) | -1.99 | (-2.49--1.47) | **1.15E-13** | **3.93E-12** | -1.06 | (-1.56--0.54) | **1.14E-04** | **3.25E-04** | same |
| PC(15-MHDA_18:2) | -1.86 | (-2.67--1.05) | **9.80E-06** | **4.05E-05** | -1.39 | (-2.07--0.66) | **3.55E-04** | **9.32E-04** | same |
| PC(17:0_18:2) | -1.54 | (-2.08--1) | **3.86E-08** | **2.57E-07** | -1.28 | (-1.84--0.68) | **5.78E-05** | **1.69E-04** | same |
| PC(17:1_18:2) | -1.21 | (-1.75--0.67) | **1.36E-05** | **5.30E-05** | -0.72 | (-1.27--0.14) | **1.54E-02** | **3.03E-02** | same |
| PC(15:0_20:4) | -1.28 | (-1.97--0.58) | **3.68E-04** | **1.06E-03** | -1.32 | (-1.88--0.72) | **3.80E-05** | **1.13E-04** | same |
| PC(35:5) | -1.57 | (-2.8--0.32) | **1.38E-02** | **2.50E-02** | -2.33 | (-3.15--1.42) | **4.33E-06** | **1.41E-05** | same |
| PC(36:6) | -2.22 | (-3.03--1.4) | **1.62E-07** | **9.83E-07** | -0.83 | (-1.42--0.2) | **1.07E-02** | **2.19E-02** | same |
| PC(15:0_22:6) | -2.68 | (-3.42--1.94) | **3.93E-12** | **9.43E-11** | -1.88 | (-2.45--1.28) | **1.58E-08** | **6.70E-08** | same |
| PC(38:2) | -1.13 | (-1.58--0.67) | **1.45E-06** | **6.94E-06** | 0.92 | (0.32-1.55) | **2.19E-03** | **5.11E-03** | opposite |
| PC(18:0_20:4) | 0.98 | (0.35-1.61) | **2.29E-03** | **5.29E-03** | -1.07 | (-1.65--0.46) | **8.92E-04** | **2.21E-03** | opposite |
| PC(38:5) (a) | -1.09 | (-1.59--0.58) | **2.73E-05** | **9.86E-05** | -1.17 | (-1.68--0.63) | **3.91E-05** | **1.15E-04** | same |
| PC(16:0_22:6) | -1.01 | (-1.64--0.37) | **1.94E-03** | **4.55E-03** | -1.25 | (-1.93--0.5) | **1.40E-03** | **3.31E-03** | same |
| PC(16:1_22:6) | -1.64 | (-2.33--0.96) | **3.49E-06** | **1.54E-05** | -1.04 | (-1.56--0.48) | **4.09E-04** | **1.06E-03** | same |
| PC(38:7)(c) | -1.58 | (-2.17--0.99) | **2.34E-07** | **1.32E-06** | -0.88 | (-1.37--0.37) | **1.02E-03** | **2.45E-03** | same |
| PC(39:5)(a) | -2.53 | (-3.35--1.7) | **4.56E-09** | **3.53E-08** | -1.98 | (-2.62--1.29) | **2.49E-07** | **9.28E-07** | same |
| PC(39:5)(b) | -2.31 | (-3.04--1.58) | **1.68E-09** | **1.56E-08** | -1.95 | (-2.55--1.29) | **7.31E-08** | **2.90E-07** | same |
| PC(15-MHDA_22:6) | -2.51 | (-3.31--1.7) | **2.77E-09** | **2.33E-08** | -2.07 | (-2.73--1.36) | **1.67E-07** | **6.31E-07** | same |
| PC(17:0_22:6) | -2.00 | (-2.74--1.26) | **1.87E-07** | **1.08E-06** | -2.24 | (-2.78--1.65) | **1.65E-11** | **9.21E-11** | same |
| PC(40:8) | -1.77 | (-2.3--1.24) | **1.86E-10** | **2.70E-09** | 0.83 | (0.2-1.5) | **9.19E-03** | **1.92E-02** | opposite |
| PC(44:12) | -2.40 | (-3.44--1.34) | **1.14E-05** | **4.60E-05** | -1.02 | (-1.76--0.22) | **1.39E-02** | **2.75E-02** | same |
| PC(O-18:0/18:1) | -1.51 | (-1.98--1.03) | **1.00E-09** | **1.02E-08** | 1.31 | (0.62-2.05) | **1.35E-04** | **3.71E-04** | opposite |
| PC(O-18:1/18:1) | -1.20 | (-1.66--0.74) | **4.16E-07** | **2.24E-06** | 0.71 | (0.11-1.35) | **2.01E-02** | **3.86E-02** | opposite |
| PC(O-36:5) | -1.11 | (-2.09--0.12) | **2.87E-02** | **4.82E-02** | -1.06 | (-1.73--0.34) | **4.52E-03** | **9.91E-03** | same |
| PC(O-18:0/22:6) | -0.69 | (-1.25--0.12) | **1.77E-02** | **3.15E-02** | -0.93 | (-1.43--0.39) | **9.93E-04** | **2.41E-03** | same |
| PC(P-16:0/18:2) | -1.10 | (-1.68--0.5) | **3.02E-04** | **8.73E-04** | -1.30 | (-1.9--0.66) | **1.43E-04** | **3.91E-04** | same |
| PC(P-35:2) (b) | -1.83 | (-2.55--1.1) | **1.07E-06** | **5.20E-06** | -1.45 | (-2.06--0.79) | **3.89E-05** | **1.15E-04** | same |
| PC(P-18:1/18:1) | -1.20 | (-1.72--0.68) | **7.54E-06** | **3.17E-05** | 1.22 | (0.39-2.12) | **3.50E-03** | **7.92E-03** | opposite |
| PC(P-17:0/20:4) (a) | -1.01 | (-1.8--0.22) | **1.29E-02** | **2.38E-02** | -1.57 | (-2.22--0.86) | **3.59E-05** | **1.08E-04** | same |
| PC(P-38:5) (b) | -0.85 | (-1.43--0.27) | **4.44E-03** | **9.70E-03** | -0.87 | (-1.5--0.2) | **1.16E-02** | **2.34E-02** | same |

| | | | | | | | | | |
|---|---|---|---|---|---|---|---|---|---|
| PC(P-40:5) (b) | -0.80 | (-1.37--0.24) | **5.56E-03** | **1.18E-02** | -0.80 | (-1.31--0.26) | **4.26E-03** | **9.37E-03** | same |
| PE(17:0_18:2) | -1.21 | (-1.9--0.52) | **6.65E-04** | **1.74E-03** | -0.85 | (-1.35--0.31) | **2.33E-03** | **5.41E-03** | same |
| PE(18:0_22:6) | 1.34 | (0.55-2.14) | **8.34E-04** | **2.13E-03** | -1.61 | (-2.34--0.8) | **2.02E-04** | **5.43E-04** | opposite |
| PE(P-16:0/18:1) | -1.04 | (-1.56--0.52) | **1.03E-04** | **3.35E-04** | 1.69 | (0.98-2.44) | **1.11E-06** | **3.90E-06** | opposite |
| PE(P-16:0/22:5) (n6) | -0.76 | (-1.4--0.12) | **2.07E-02** | **3.60E-02** | 0.91 | (0.1-1.79) | **2.66E-02** | **4.98E-02** | opposite |
| PE(P-18:0/18:1) | -1.25 | (-1.84--0.66) | **3.98E-05** | **1.40E-04** | 1.01 | (0.24-1.85) | **9.34E-03** | **1.94E-02** | opposite |
| PE(P-18:0/20:5) | -1.63 | (-2.99--0.26) | **2.00E-02** | **3.52E-02** | -1.97 | (-3.3--0.38) | **1.77E-02** | **3.42E-02** | same |
| PE(P-20:0/22:6) | -0.78 | (-1.31--0.26) | **3.71E-03** | **8.27E-03** | -0.70 | (-1.25--0.11) | **2.13E-02** | **4.05E-02** | same |
| PI(34:0) | -2.66 | (-3.58--1.72) | **4.56E-08** | **2.96E-07** | -1.86 | (-2.99--0.54) | **7.41E-03** | **1.58E-02** | same |
| PI(17:0_18:1) | -2.37 | (-2.95--1.79) | **6.39E-15** | **2.79E-13** | -1.30 | (-2.26--0.22) | **1.94E-02** | **3.75E-02** | same |
| PI(17:0_18:2) | -2.70 | (-3.33--2.07) | **4.72E-16** | **2.83E-14** | -1.91 | (-2.87--0.83) | **9.67E-04** | **2.37E-03** | same |
| PI(18:0_18:1) | -1.94 | (-2.65--1.23) | **1.16E-07** | **7.23E-07** | -1.18 | (-1.84--0.46) | **1.71E-03** | **4.02E-03** | same |
| PI(36:2) | -1.93 | (-2.51--1.34) | **2.53E-10** | **3.28E-09** | -1.22 | (-1.78--0.62) | **1.28E-04** | **3.56E-04** | same |
| PI(18:1_18:2) | -2.66 | (-3.29--2.03) | **1.11E-15** | **5.91E-14** | -1.47 | (-1.98--0.93) | **4.93E-07** | **1.78E-06** | same |
| PI(16:0_20:3) (b) | -2.46 | (-3.2--1.72) | **1.99E-10** | **2.80E-09** | 2.99 | (1.94-4.14) | **2.09E-09** | **9.74E-09** | opposite |
| PI(18:0_20:2) | -2.66 | (-3.36--1.95) | **6.56E-13** | **1.85E-11** | 2.45 | (1.2-3.85) | **5.83E-05** | **1.70E-04** | opposite |
| PI(18:0_20:4) | -0.79 | (-1.31--0.28) | **2.68E-03** | **6.09E-03** | -0.59 | (-1.01--0.14) | **1.08E-02** | **2.19E-02** | same |
| PI(38:6) | -2.63 | (-3.27--1.98) | **1.30E-14** | **5.19E-13** | -0.92 | (-1.44--0.37) | **1.31E-03** | **3.11E-03** | same |
| PI(18:0_22:4) | -1.24 | (-1.89--0.58) | **2.27E-04** | **6.78E-04** | 1.20 | (0.52-1.92) | **4.03E-04** | **1.05E-03** | opposite |
| PI(18:0_22:5) (n3) | -1.73 | (-2.4--1.06) | **5.69E-07** | **2.91E-06** | -1.24 | (-1.88--0.56) | **5.87E-04** | **1.47E-03** | same |
| PI(18:0_22:6) | -2.03 | (-2.68--1.38) | **2.31E-09** | **2.05E-08** | -1.11 | (-1.63--0.55) | **1.59E-04** | **4.30E-04** | same |

**Supplementary Table 4c.** The overlapping lipid species between the birth weight and 6-year-old child BMI studies. Related to Figure 3.

| Lipid Species | Birth Weight (cord blood) | | | | Child BMI (six-year-old) | | | | Trend |
|---|---|---|---|---|---|---|---|---|---|
| | %change in lipid concentration per 100 grams | 95%CI | p-value | p-value(BH) | %change in lipid concentration per unit BMI | 95%CI | p-value | p-value(BH) | Trend |
| AC(18:0) | -0.78 | (-1.3--0.24) | **5.68E-03** | **1.22E-02** | -1.98 | (-3.33--0.62) | **4.60E-03** | **1.97E-02** | same |
| CE(16:1) | 0.98 | (0.31-1.69) | **3.55E-03** | **8.00E-03** | 5.77 | (4.04-7.53) | **9.25E-11** | **2.22E-08** | same |
| DG(16:0_16:1) | -1.32 | (-2.11--0.45) | **3.64E-03** | **8.13E-03** | 6.21 | (3.58-8.89) | **3.25E-06** | **8.67E-05** | opposite |
| DG(14:0_18:2) | -3.08 | (-4.04--1.96) | **1.68E-06** | **5.70E-06** | 6.70 | (3.02-10.51) | **3.23E-04** | **2.28E-03** | opposite |
| DG(16:0_18:1) | -1.91 | (-2.57--1.19) | **1.35E-06** | **4.62E-06** | 3.90 | (1.38-6.49) | **2.40E-03** | **1.18E-02** | opposite |
| DG(16:1_18:1) | -2.08 | (-2.77--1.32) | **7.91E-07** | **2.84E-06** | 6.17 | (3.26-9.17) | **2.92E-05** | **4.64E-04** | opposite |
| DG(16:0_18:2) | -3.07 | (-3.76--2.31) | **1.09E-11** | **6.22E-11** | 4.00 | (1.57-6.48) | **1.20E-03** | **6.49E-03** | opposite |
| DG(18:0_18:1) | -1.29 | (-1.99--0.53) | **1.23E-03** | **2.94E-03** | 4.06 | (1.71-6.46) | **6.87E-04** | **4.23E-03** | opposite |
| DG(18:0_18:2) | -1.62 | (-2.52--0.61) | **2.43E-03** | **5.62E-03** | 3.76 | (1.3-6.29) | **2.68E-03** | **1.27E-02** | opposite |
| DG(16:0_20:4) | -2.06 | (-2.83--1.21) | **9.20E-06** | **2.95E-05** | 4.97 | (2.34-7.66) | **1.98E-04** | **1.61E-03** | opposite |
| DG(18:1_20:4) | -2.06 | (-2.71--1.36) | **1.08E-07** | **4.14E-07** | 3.90 | (1.68-6.16) | **5.47E-04** | **3.60E-03** | opposite |
| DG(16:0_22:5) | -1.95 | (-2.72--1.09) | **2.99E-05** | **9.02E-05** | 4.60 | (2.03-7.23) | **4.22E-04** | **2.89E-03** | opposite |
| DG(18:2_20:4) | -3.14 | (-3.87--2.33) | **7.74E-11** | **4.08E-10** | 4.23 | (1.78-6.73) | **6.78E-04** | **4.23E-03** | opposite |
| DG(16:0_22:6) | -4.09 | (-4.76--3.33) | **6.80E-17** | **8.37E-16** | 5.50 | (2.13-8.99) | **1.31E-03** | **6.84E-03** | opposite |
| DG(18:1_22:6) | -3.49 | (-4.15--2.76) | **7.44E-15** | **6.49E-14** | 4.14 | (1.32-7.04) | **3.88E-03** | **1.73E-02** | opposite |
| DG(18:2_22:6) | -4.00 | (-4.74--3.16) | **9.00E-14** | **6.75E-13** | 4.79 | (1.45-8.24) | **4.80E-03** | **2.04E-02** | opposite |
| TG(48:0) [NL-18:0] | -1.44 | (-2.28--0.51) | **3.14E-03** | **7.14E-03** | 7.45 | (3.74-11.29) | **7.09E-05** | **8.54E-04** | opposite |
| TG(48:1) [NL-16:1] | -2.43 | (-3.32--1.41) | **1.61E-05** | **5.11E-05** | 9.58 | (5.88-13.41) | **2.99E-07** | **1.32E-05** | opposite |
| TG(48:1) [NL-18:1] | -3.15 | (-3.94--2.26) | **2.16E-09** | **9.86E-09** | 8.20 | (3.97-12.59) | **1.20E-04** | **1.17E-03** | opposite |
| TG(48:2) [NL-14:0] | -3.76 | (-4.46--2.97) | **4.13E-14** | **3.31E-13** | 8.73 | (4.41-13.22) | **6.10E-05** | **7.91E-04** | opposite |
| TG(48:2) [NL-14:1] | -3.06 | (-3.81--2.23) | **4.50E-10** | **2.23E-09** | 8.90 | (4.81-13.16) | **1.61E-05** | **3.36E-04** | opposite |
| TG(48:2) [NL-16:1] | -2.85 | (-3.65--1.96) | **3.48E-08** | **1.43E-07** | 10.74 | (6.57-15.07) | **3.03E-07** | **1.32E-05** | opposite |
| TG(48:2) [NL-18:2] | -4.14 | (-4.86--3.32) | **5.42E-15** | **4.91E-14** | 8.58 | (4.3-13.04) | **7.11E-05** | **8.54E-04** | opposite |
| TG(48:3) [NL-14:0] | -4.04 | (-4.73--3.26) | **1.02E-15** | **1.04E-14** | 9.53 | (5.17-14.08) | **1.43E-05** | **3.28E-04** | opposite |
| TG(48:3) [NL-16:1] | -2.89 | (-3.59--2.12) | **1.90E-10** | **9.60E-10** | 8.59 | (4.99-12.31) | **2.28E-06** | **6.49E-05** | opposite |
| TG(48:3) [NL-18:3] | -3.23 | (-3.91--2.48) | **8.75E-13** | **5.83E-12** | 8.18 | (3.69-12.86) | **3.09E-04** | **2.27E-03** | opposite |
| TG(49:1) [NL-17:1] | -2.19 | (-2.84--1.47) | **5.29E-08** | **2.14E-07** | 4.94 | (2.47-7.47) | **8.54E-05** | **9.54E-04** | opposite |
| TG(50:1) [NL-14:0] | -1.46 | (-2.35--0.47) | **4.76E-03** | **1.03E-02** | 6.89 | (3.47-10.42) | **7.00E-05** | **8.54E-04** | opposite |
| TG(50:1) [NL-16:0] | -2.49 | (-3.25--1.64) | **2.17E-07** | **8.15E-07** | 4.63 | (2.1-7.22) | **3.13E-04** | **2.27E-03** | opposite |
| TG(50:1) [NL-18:1] | -2.47 | (-3.24--1.62) | **2.61E-07** | **9.63E-07** | 4.17 | (1.73-6.68) | **7.99E-04** | **4.74E-03** | opposite |
| TG(50:2) [NL-14:0] | -3.34 | (-4.1--2.48) | **1.16E-10** | **6.01E-10** | 4.36 | (1.14-7.69) | **7.83E-03** | **3.01E-02** | opposite |
| TG(50:2) [NL-16:1] | -3.01 | (-3.81--2.11) | **1.08E-08** | **4.67E-08** | 6.45 | (3.42-9.58) | **2.79E-05** | **4.62E-04** | opposite |
| TG(50:2) [NL-18:1] | -3.12 | (-3.91--2.21) | **4.54E-09** | **2.00E-08** | 6.24 | (3.2-9.38) | **5.20E-05** | **7.33E-04** | opposite |
| TG(50:2) [NL-18:2] | -4.17 | (-4.92--3.31) | **5.26E-14** | **4.14E-13** | 4.53 | (2.11-7) | **2.30E-04** | **1.78E-03** | opposite |
| TG(50:3) [NL-14:0] | -4.25 | (-4.95--3.45) | **4.04E-16** | **4.31E-15** | 5.47 | (1.88-9.18) | **2.69E-03** | **1.27E-02** | opposite |
| TG(50:3) [NL-14:1] | -2.39 | (-3.17--1.51) | **1.11E-06** | **3.90E-06** | 4.17 | (1.33-7.09) | **3.92E-03** | **1.73E-02** | opposite |
| TG(50:3) [NL-16:1] | -3.88 | (-4.58--3.09) | **8.11E-15** | **6.95E-14** | 8.28 | (4.83-11.85) | **2.15E-06** | **6.49E-05** | opposite |
| TG(50:3) [NL-18:2] | -4.37 | (-5.06--3.59) | **3.53E-17** | **4.70E-16** | 7.14 | (3.78-10.61) | **2.72E-05** | **4.62E-04** | opposite |
| TG(50:3) [NL-18:3] | -3.81 | (-4.54--2.98) | **2.31E-13** | **1.65E-12** | 6.76 | (3.2-10.44) | **1.78E-04** | **1.52E-03** | opposite |
| TG(50:4) [NL-14:0] | -4.31 | (-4.97--3.57) | **2.52E-18** | **4.32E-17** | 5.22 | (1.61-8.96) | **4.43E-03** | **1.92E-02** | opposite |
| TG(50:4) [NL-18:3] | -3.93 | (-4.56--3.22) | **9.72E-18** | **1.37E-16** | 7.93 | (4.25-11.74) | **2.01E-05** | **4.02E-04** | opposite |
| TG(50:4) [NL-20:4] | -3.52 | (-4.34--2.57) | **6.23E-10** | **3.05E-09** | 10.00 | (5.69-14.48) | **3.96E-06** | **1.00E-04** | opposite |
| TG(51:1) [NL-17:0] | -2.72 | (-3.44--1.91) | **4.45E-09** | **1.98E-08** | 4.90 | (2-7.89) | **8.93E-04** | **5.23E-03** | opposite |
| TG(51:2) [NL-17:0] | -3.40 | (-4.11--2.6) | **2.54E-12** | **1.63E-11** | 5.48 | (2.52-8.52) | **2.67E-04** | **2.03E-03** | opposite |
| TG(51:2) [NL-17:1] | -2.89 | (-3.53--2.19) | **2.44E-12** | **1.58E-11** | 4.90 | (2.22-7.65) | **3.17E-04** | **2.27E-03** | opposite |
| TG(52:1) [NL-18:0] | -1.86 | (-2.76--0.83) | **6.81E-04** | **1.69E-03** | 5.79 | (2.8-8.86) | **1.32E-04** | **1.22E-03** | opposite |
| TG(52:1) [NL-18:1] | -1.80 | (-2.71--0.77) | **9.96E-04** | **2.41E-03** | 5.71 | (2.83-8.68) | **9.59E-05** | **1.00E-03** | opposite |
| TG(52:2) [NL-18:2] | -2.75 | (-3.58--1.81) | **3.20E-07** | **1.17E-06** | 4.45 | (2.1-6.85) | **1.98E-04** | **1.61E-03** | opposite |
| TG(52:3) [NL-16:1] | -3.37 | (-4.1--2.56) | **1.00E-11** | **5.79E-11** | 3.39 | (0.97-5.88) | **5.95E-03** | **2.36E-02** | opposite |
| TG(52:4) [NL-16:1] | -4.52 | (-5.17--3.78) | **1.14E-19** | **2.48E-18** | 4.16 | (1.08-7.34) | **7.91E-03** | **3.01E-02** | opposite |
| TG(52:4) [NL-18:3] | -4.11 | (-4.78--3.35) | **9.43E-17** | **1.10E-15** | 4.40 | (1.43-7.46) | **3.60E-03** | **1.63E-02** | opposite |
| TG(52:5) [NL-18:3] | -4.16 | (-4.81--3.43) | **2.87E-18** | **4.75E-17** | 5.19 | (2-8.47) | **1.36E-03** | **6.99E-03** | opposite |
| TG(52:5) [NL-20:4] | -3.63 | (-4.4--2.74) | **2.19E-11** | **1.19E-10** | 10.03 | (6.39-13.8) | **4.75E-08** | **4.56E-06** | opposite |

| | | | | | | | | | |
|---|---|---|---|---|---|---|---|---|---|
| TG(52:5) [NL-20:5] | -4.43 | (-5.13--3.64) | **4.66E-17** | **5.88E-16** | 7.78 | (4.53-11.13) | **2.30E-06** | **6.49E-05** | opposite |
| TG(54:1) [NL-18:1] | -1.64 | (-2.56--0.59) | **2.89E-03** | **6.61E-03** | 3.54 | (1.13-6.02) | **3.96E-03** | **1.73E-02** | opposite |
| TG(54:2) [NL-18:0] | -1.85 | (-2.69--0.91) | **2.40E-04** | **6.39E-04** | 3.30 | (0.94-5.71) | **6.02E-03** | **2.36E-02** | opposite |
| TG(54:3) [NL-18:2] | -3.06 | (-3.75--2.3) | **1.28E-11** | **7.23E-11** | 2.32 | (0.48-4.19) | **1.32E-02** | **4.65E-02** | opposite |
| TG(54:4) [NL-20:3] | -2.73 | (-3.52--1.84) | **8.04E-08** | **3.16E-07** | 4.94 | (2.41-7.54) | **1.24E-04** | **1.19E-03** | opposite |
| TG(54:5) [NL-20:4] | -3.73 | (-4.49--2.86) | **4.40E-12** | **2.64E-11** | 5.84 | (3.43-8.3) | **1.90E-06** | **6.49E-05** | opposite |
| TG(54:6) [NL-20:4] | -4.11 | (-4.83--3.28) | **1.19E-14** | **9.81E-14** | 7.52 | (4.68-10.44) | **1.91E-07** | **1.15E-05** | opposite |
| TG(54:6) [NL-20:5] | -5.04 | (-5.69--4.28) | **7.74E-21** | **2.50E-19** | 8.39 | (5.04-11.85) | **7.54E-07** | **3.02E-05** | opposite |
| TG(54:6) [NL-22:6] | -5.16 | (-5.83--4.38) | **1.98E-20** | **5.23E-19** | 8.20 | (4.48-12.06) | **1.29E-05** | **3.10E-04** | opposite |
| TG(54:7) [NL-20:5] | -4.97 | (-5.63--4.22) | **1.23E-20** | **3.49E-19** | 8.46 | (5.22-11.8) | **2.49E-07** | **1.32E-05** | opposite |
| TG(54:7) [NL-22:6] | -4.92 | (-5.56--4.18) | **1.93E-21** | **7.12E-20** | 10.42 | (6.51-14.47) | **1.23E-07** | **9.83E-06** | opposite |
| TG(56:6) [NL-20:4] | -3.00 | (-3.77--2.15) | **1.89E-09** | **8.92E-09** | 3.72 | (1.91-5.57) | **5.78E-05** | **7.86E-04** | opposite |
| TG(56:6) [NL-22:5] | -3.83 | (-4.52--3.06) | **4.63E-15** | **4.36E-14** | 3.96 | (1.82-6.14) | **2.87E-04** | **2.15E-03** | opposite |
| TG(56:7) [NL-20:4] | -3.66 | (-4.41--2.81) | **3.38E-12** | **2.08E-11** | 4.16 | (1.73-6.64) | **7.60E-04** | **4.56E-03** | opposite |
| TG(56:7) [NL-20:5] | -3.79 | (-4.51--2.97) | **1.22E-13** | **9.04E-13** | 5.58 | (2.64-8.6) | **1.82E-04** | **1.54E-03** | opposite |
| TG(56:7) [NL-22:5] | -4.29 | (-4.93--3.57) | **2.43E-19** | **4.87E-18** | 5.15 | (2.46-7.91) | **1.64E-04** | **1.46E-03** | opposite |
| TG(56:7) [NL-22:6] | -5.07 | (-5.72--4.33) | **1.85E-21** | **7.12E-20** | 6.45 | (3.01-10) | **2.16E-04** | **1.70E-03** | opposite |
| TG(56:8) [NL-20:4] | -2.95 | (-3.72--2.08) | **5.67E-09** | **2.47E-08** | 5.12 | (2.54-7.75) | **9.23E-05** | **1.00E-03** | opposite |
| TG(56:8) [NL-20:5] | -4.13 | (-4.79--3.37) | **3.84E-17** | **4.98E-16** | 5.03 | (2.01-8.13) | **1.03E-03** | **5.94E-03** | opposite |
| TG(56:8) [NL-22:6] | -5.36 | (-5.97--4.65) | **1.69E-24** | **1.02E-22** | 7.57 | (4.02-11.24) | **2.51E-05** | **4.62E-04** | opposite |
| TG(56:9) [NL-22:6] | -4.88 | (-5.5--4.17) | **1.57E-22** | **7.54E-21** | 6.66 | (3.29-10.14) | **9.49E-05** | **1.00E-03** | opposite |
| TG(58:10) [NL-22:6] | -4.48 | (-5.11--3.76) | **3.35E-20** | **8.05E-19** | 5.17 | (1.89-8.56) | **1.90E-03** | **9.52E-03** | opposite |
| TG(58:8) [NL-22:6] | -4.50 | (-5.16--3.76) | **3.67E-19** | **7.05E-18** | 4.73 | (1.39-8.18) | **5.34E-03** | **2.16E-02** | opposite |
| TG(58:9) [NL-22:6] | -4.76 | (-5.39--4.06) | **4.43E-22** | **1.93E-20** | 4.51 | (1.17-7.97) | **7.98E-03** | **3.02E-02** | opposite |
| TG(O-50:1) [NL-17:1] | -2.21 | (-2.88--1.47) | **9.38E-08** | **3.66E-07** | 4.99 | (2.47-7.57) | **1.01E-04** | **1.03E-03** | opposite |
| TG(O-52:2) [NL-17:1] | -2.91 | (-3.55--2.2) | **4.20E-12** | **2.55E-11** | 4.89 | (2.19-7.66) | **3.63E-04** | **2.52E-03** | opposite |
| TG(O-52:2) [NL-18:1] | -1.84 | (-2.72--0.85) | **5.37E-04** | **1.36E-03** | 5.04 | (2-8.16) | **1.08E-03** | **6.15E-03** | opposite |
| Cer(m18:0/22:0) | -1.25 | (-2.16--0.23) | **1.75E-02** | **3.42E-02** | 2.43 | (0.5-4.39) | **1.34E-02** | **4.68E-02** | opposite |
| Cer(m18:0/24:1) | -1.46 | (-2.42--0.37) | **9.81E-03** | **2.02E-02** | 4.35 | (2.12-6.63) | **1.28E-04** | **1.20E-03** | opposite |
| LPC(14:0) [sn2] | 1.75 | (1.25-2.26) | **6.35E-13** | **4.36E-12** | 4.86 | (2.45-7.33) | **7.65E-05** | **8.95E-04** | same |
| LPC(14:0) [sn1] | 1.62 | (1.14-2.11) | **5.17E-12** | **3.06E-11** | 4.70 | (2.35-7.09) | **8.28E-05** | **9.46E-04** | same |
| LPC(16:1) [sn2] | 4.00 | (3.27-4.78) | **3.37E-31** | **8.10E-29** | 5.09 | (3.19-7.03) | **1.66E-07** | **1.14E-05** | same |
| LPC(16:1) [sn1] | 3.89 | (3.2-4.61) | **6.09E-33** | **2.92E-30** | 3.93 | (2.1-5.79) | **2.67E-05** | **4.62E-04** | same |
| LPC(20:1) [sn1] | 1.75 | (1.27-2.25) | **1.55E-13** | **1.12E-12** | -2.32 | (-3.93--0.7) | **5.46E-03** | **2.19E-02** | opposite |
| LPC(20:3) [sn2] | 2.58 | (1.91-3.3) | **1.79E-15** | **1.72E-14** | 2.63 | (0.79-4.5) | **4.98E-03** | **2.10E-02** | same |
| LPC(20:3) [104_sn1] | 3.22 | (2.49-3.99) | **9.49E-21** | **2.85E-19** | 2.53 | (0.51-4.6) | **1.42E-02** | **4.91E-02** | same |
| PC(32:1) | 0.94 | (0.11-1.83) | **2.49E-02** | **4.70E-02** | 3.56 | (1.41-5.75) | **1.12E-03** | **6.20E-03** | same |
| PC(34:5) | -1.14 | (-1.93--0.28) | **1.10E-02** | **2.23E-02** | 7.01 | (3.69-10.43) | **3.00E-05** | **4.64E-04** | opposite |
| PC(17:0_18:2) | -1.28 | (-1.84--0.68) | **5.78E-05** | **1.69E-04** | -2.03 | (-3.62--0.41) | **1.44E-02** | **4.92E-02** | same |
| PC(16:0_20:5) | -1.86 | (-2.69--0.94) | **1.83E-04** | **4.92E-04** | 3.86 | (0.8-7.01) | **1.32E-02** | **4.65E-02** | opposite |
| PC(36:6) | -0.83 | (-1.42--0.2) | **1.07E-02** | **2.19E-02** | 4.41 | (1.91-6.97) | **5.17E-04** | **3.45E-03** | opposite |
| PC(18:0_22:6) | -1.44 | (-2.09--0.74) | **1.21E-04** | **3.40E-04** | 3.01 | (0.9-5.18) | **5.22E-03** | **2.12E-02** | opposite |
| PC(O-18:0/18:1) | 1.31 | (0.62-2.05) | **1.35E-04** | **3.71E-04** | -3.04 | (-4.22--1.85) | **1.05E-06** | **3.88E-05** | opposite |
| PC(O-18:1/18:1) | 0.71 | (0.11-1.35) | **2.01E-02** | **3.86E-02** | -1.86 | (-2.95--0.75) | **1.12E-03** | **6.20E-03** | opposite |
| PC(P-35:2) (b) | -1.45 | (-2.06--0.79) | **3.89E-05** | **1.15E-04** | -2.72 | (-4.69--0.7) | **8.59E-03** | **3.22E-02** | same |
| PE(18:0_22:6) | -1.61 | (-2.34--0.8) | **2.02E-04** | **5.43E-04** | 3.81 | (1.08-6.61) | **6.06E-03** | **2.36E-02** | opposite |
| PI(18:1_18:2) | -1.47 | (-1.98--0.93) | **4.93E-07** | **1.78E-06** | -3.22 | (-4.99--1.41) | **5.62E-04** | **3.65E-03** | same |

**Supplementary Table 4d.** The overlapping lipid species between the 6-year-old child BMI and adult (postnatal mother) BMI studies. Related to Figure 3.

| Lipid Species | Child BMI (six-year-old) | | | | Adult BMI (postnatal) | | | | |
|---|---|---|---|---|---|---|---|---|---|
| | %change in lipid concentration per unit BMI | 95%CI | p-value | p-value(BH) | %change in lipid concentration per unit BMI | 95%CI | p-value | p-value(BH) | Trend |
| AC(18:0) | -1.98 | (-3.33--0.62) | **4.60E-03** | **1.97E-02** | -0.68 | (-1.13--0.23) | **3.16E-03** | **6.00E-03** | same |
| CE(16:1) | 5.77 | (4.04-7.53) | **9.25E-11** | **2.22E-08** | 2.17 | (1.53-2.82) | **6.89E-11** | **4.60E-10** | same |
| CE(18:3) | 2.78 | (0.7-4.91) | **8.76E-03** | **3.24E-02** | 0.72 | (0.11-1.32) | **2.04E-02** | **3.21E-02** | same |
| DG(16:0_16:1) | 6.21 | (3.58-8.89) | **3.25E-06** | **8.67E-05** | 4.43 | (3.46-5.4) | **9.58E-19** | **4.12E-17** | same |
| DG(14:0_18:2) | 6.70 | (3.02-10.51) | **3.23E-04** | **2.28E-03** | 4.19 | (2.86-5.54) | **7.61E-10** | **4.11E-09** | same |
| DG(16:0_18:1) | 3.90 | (1.38-6.49) | **2.40E-03** | **1.18E-02** | 4.26 | (3.33-5.21) | **1.32E-18** | **4.86E-17** | same |
| DG(16:1_18:1) | 6.17 | (3.26-9.17) | **2.92E-05** | **4.64E-04** | 4.04 | (3.07-5.03) | **1.05E-15** | **1.80E-14** | same |
| DG(16:0_18:2) | 4.00 | (1.57-6.48) | **1.20E-03** | **6.49E-03** | 3.62 | (2.72-4.52) | **5.53E-15** | **7.59E-14** | same |
| DG(18:0_18:1) | 4.06 | (1.71-6.46) | **6.87E-04** | **4.23E-03** | 4.41 | (3.51-5.31) | **4.70E-21** | **4.51E-19** | same |
| DG(18:0_18:2) | 3.76 | (1.3-6.29) | **2.68E-03** | **1.27E-02** | 4.40 | (3.41-5.39) | **7.51E-18** | **2.57E-16** | same |
| DG(16:0_20:4) | 4.97 | (2.34-7.66) | **1.98E-04** | **1.61E-03** | 4.27 | (3.25-5.3) | **4.79E-16** | **9.59E-15** | same |
| DG(18:1_20:4) | 3.90 | (1.68-6.16) | **5.47E-04** | **3.60E-03** | 2.47 | (1.71-3.22) | **1.81E-10** | **1.10E-09** | same |
| DG(16:0_22:5) | 4.60 | (2.03-7.23) | **4.22E-04** | **2.89E-03** | 2.42 | (1.46-3.38) | **7.01E-07** | **2.39E-06** | same |
| DG(18:2_20:4) | 4.23 | (1.78-6.73) | **6.78E-04** | **4.23E-03** | 2.42 | (1.55-3.3) | **5.34E-08** | **2.23E-07** | same |
| DG(16:0_22:6) | 5.50 | (2.13-8.99) | **1.31E-03** | **6.84E-03** | 4.15 | (2.87-5.45) | **2.35E-10** | **1.41E-09** | same |
| DG(18:1_22:6) | 4.14 | (1.32-7.04) | **3.88E-03** | **1.73E-02** | 3.18 | (2.04-4.34) | **5.26E-08** | **2.22E-07** | same |
| DG(18:2_22:6) | 4.79 | (1.45-8.24) | **4.80E-03** | **2.04E-02** | 3.05 | (1.87-4.24) | **3.60E-07** | **1.28E-06** | same |
| TG(48:0) [NL-18:0] | 7.45 | (3.74-11.29) | **7.09E-05** | **8.54E-04** | 4.34 | (2.99-5.71) | **3.54E-10** | **2.07E-09** | same |
| TG(48:1) [NL-16:1] | 9.58 | (5.88-13.41) | **2.99E-07** | **1.32E-05** | 6.64 | (5.32-7.97) | **1.51E-22** | **3.62E-20** | same |
| TG(48:1) [NL-18:1] | 8.20 | (3.97-12.59) | **1.20E-04** | **1.17E-03** | 5.21 | (3.76-6.69) | **2.44E-12** | **2.13E-11** | same |
| TG(48:2) [NL-14:0] | 8.73 | (4.41-13.22) | **6.10E-05** | **7.91E-04** | 5.21 | (3.73-6.72) | **5.81E-12** | **4.73E-11** | same |
| TG(48:2) [NL-14:1] | 8.90 | (4.81-13.16) | **1.61E-05** | **3.36E-04** | 4.79 | (3.42-6.18) | **7.51E-12** | **5.72E-11** | same |
| TG(48:2) [NL-16:1] | 10.74 | (6.57-15.07) | **3.03E-07** | **1.32E-05** | 5.71 | (4.32-7.12) | **1.20E-15** | **1.99E-14** | same |
| TG(48:2) [NL-18:2] | 8.58 | (4.3-13.04) | **7.11E-05** | **8.54E-04** | 5.21 | (3.68-6.77) | **2.87E-11** | **2.03E-10** | same |
| TG(48:3) [NL-14:0] | 9.53 | (5.17-14.08) | **1.43E-05** | **3.28E-04** | 4.32 | (2.9-5.76) | **2.31E-09** | **1.14E-08** | same |
| TG(48:3) [NL-16:1] | 8.59 | (4.99-12.31) | **2.28E-06** | **6.49E-05** | 4.00 | (2.79-5.24) | **1.30E-10** | **8.10E-10** | same |
| TG(48:3) [NL-18:3] | 8.18 | (3.69-12.86) | **3.09E-04** | **2.27E-03** | 4.99 | (3.41-6.59) | **5.23E-10** | **2.92E-09** | same |
| TG(49:1) [NL-17:1] | 4.94 | (2.47-7.47) | **8.54E-05** | **9.54E-04** | 4.04 | (3.11-4.97) | **2.94E-17** | **8.82E-16** | same |
| TG(50:1) [NL-14:0] | 6.89 | (3.47-10.42) | **7.00E-05** | **8.54E-04** | 4.78 | (3.52-6.05) | **1.11E-13** | **1.24E-12** | same |
| TG(50:1) [NL-16:0] | 4.63 | (2.1-7.22) | **3.13E-04** | **2.27E-03** | 4.34 | (3.39-5.3) | **1.03E-18** | **4.12E-17** | same |
| TG(50:1) [NL-18:1] | 4.17 | (1.73-6.68) | **7.99E-04** | **4.74E-03** | 4.10 | (3.16-5.05) | **4.06E-17** | **1.08E-15** | same |
| TG(50:2) [NL-14:0] | 4.36 | (1.14-7.69) | **7.83E-03** | **3.01E-02** | 2.80 | (1.76-3.86) | **1.50E-07** | **5.54E-07** | same |
| TG(50:2) [NL-16:1] | 6.45 | (3.42-9.58) | **2.79E-05** | **4.62E-04** | 4.04 | (3.07-5.02) | **6.67E-16** | **1.28E-14** | same |
| TG(50:2) [NL-18:1] | 6.24 | (3.2-9.38) | **5.20E-05** | **7.33E-04** | 3.59 | (2.65-4.54) | **1.19E-13** | **1.30E-12** | same |
| TG(50:2) [NL-18:2] | 4.53 | (2.11-7) | **2.30E-04** | **1.78E-03** | 3.72 | (2.82-4.62) | **8.45E-16** | **1.53E-14** | same |
| TG(50:3) [NL-14:0] | 5.47 | (1.88-9.18) | **2.69E-03** | **1.27E-02** | 3.13 | (1.93-4.34) | **2.88E-07** | **1.04E-06** | same |
| TG(50:3) [NL-14:1] | 4.17 | (1.33-7.09) | **3.92E-03** | **1.73E-02** | 2.66 | (1.69-3.63) | **7.85E-08** | **3.14E-07** | same |
| TG(50:3) [NL-16:1] | 8.28 | (4.83-11.85) | **2.15E-06** | **6.49E-05** | 4.38 | (3.28-5.5) | **1.32E-14** | **1.67E-13** | same |
| TG(50:3) [NL-18:2] | 7.14 | (3.78-10.61) | **2.72E-05** | **4.62E-04** | 3.72 | (2.65-4.81) | **1.38E-11** | **1.00E-10** | same |
| TG(50:3) [NL-18:3] | 6.76 | (3.2-10.44) | **1.78E-04** | **1.52E-03** | 5.18 | (3.92-6.44) | **8.59E-16** | **1.53E-14** | same |
| TG(50:4) [NL-14:0] | 5.22 | (1.61-8.96) | **4.43E-03** | **1.92E-02** | 2.75 | (1.47-4.04) | **2.32E-05** | **6.36E-05** | same |
| TG(50:4) [NL-18:3] | 7.93 | (4.25-11.74) | **2.01E-05** | **4.02E-04** | 4.14 | (2.91-5.38) | **5.20E-11** | **3.51E-10** | same |
| TG(50:4) [NL-20:4] | 10.00 | (5.69-14.48) | **3.96E-06** | **1.00E-04** | 5.64 | (4.02-7.29) | **8.80E-12** | **6.60E-11** | same |
| TG(51:1) [NL-17:0] | 4.90 | (2-7.89) | **8.93E-04** | **5.23E-03** | 4.54 | (3.5-5.59) | **2.82E-17** | **8.82E-16** | same |
| TG(51:2) [NL-17:0] | 5.48 | (2.52-8.52) | **2.67E-04** | **2.03E-03** | 4.07 | (3.06-5.09) | **6.13E-15** | **8.18E-14** | same |
| TG(51:2) [NL-17:1] | 4.90 | (2.22-7.65) | **3.17E-04** | **2.27E-03** | 3.80 | (2.86-4.74) | **3.86E-15** | **5.98E-14** | same |
| TG(52:1) [NL-18:0] | 5.79 | (2.8-8.86) | **1.32E-04** | **1.22E-03** | 5.69 | (4.53-6.86) | **2.15E-21** | **2.58E-19** | same |
| TG(52:1) [NL-18:1] | 5.71 | (2.83-8.68) | **9.59E-05** | **1.00E-03** | 5.86 | (4.73-7) | **1.79E-23** | **8.60E-21** | same |
| TG(52:2) [NL-18:2] | 4.45 | (2.1-6.85) | **1.98E-04** | **1.61E-03** | 4.17 | (3.27-5.07) | **4.36E-19** | **2.33E-17** | same |
| TG(52:3) [NL-16:1] | 3.39 | (0.97-5.88) | **5.95E-03** | **2.36E-02** | 1.94 | (1.21-2.66) | **1.73E-07** | **6.33E-07** | same |
| TG(52:4) [NL-16:1] | 4.16 | (1.08-7.34) | **7.91E-03** | **3.01E-02** | 2.22 | (1.27-3.18) | **4.67E-06** | **1.43E-05** | same |
| TG(52:4) [NL-18:3] | 4.40 | (1.43-7.46) | **3.60E-03** | **1.63E-02** | 3.19 | (2.17-4.21) | **8.03E-10** | **4.24E-09** | same |

| | | | | | | | | | |
|---|---|---|---|---|---|---|---|---|---|
| TG(52:5) [NL-18:3] | 5.19 | (2-8.47) | **1.36E-03** | **6.99E-03** | 2.89 | (1.78-4.01) | **3.35E-07** | **1.20E-06** | same |
| TG(52:5) [NL-20:4] | 10.03 | (6.39-13.8) | **4.75E-08** | **4.56E-06** | 5.46 | (4.2-6.74) | **3.97E-17** | **1.08E-15** | same |
| TG(52:5) [NL-20:5] | 7.78 | (4.53-11.13) | **2.30E-06** | **6.49E-05** | 5.06 | (3.69-6.44) | **4.80E-13** | **4.70E-12** | same |
| TG(54:1) [NL-18:1] | 3.54 | (1.13-6.02) | **3.96E-03** | **1.73E-02** | 4.45 | (3.57-5.34) | **4.54E-22** | **7.26E-20** | same |
| TG(54:2) [NL-18:0] | 3.30 | (0.94-5.71) | **6.02E-03** | **2.36E-02** | 4.24 | (3.35-5.13) | **2.72E-20** | **1.63E-18** | same |
| TG(54:3) [NL-18:2] | 2.32 | (0.48-4.19) | **1.32E-02** | **4.65E-02** | 2.23 | (1.57-2.89) | **5.13E-11** | **3.51E-10** | same |
| TG(54:4) [NL-20:3] | 4.94 | (2.41-7.54) | **1.24E-04** | **1.19E-03** | 3.70 | (2.83-4.58) | **2.18E-16** | **4.75E-15** | same |
| TG(54:5) [NL-20:4] | 5.84 | (3.43-8.3) | **1.90E-06** | **6.49E-05** | 3.37 | (2.55-4.2) | **2.12E-15** | **3.39E-14** | same |
| TG(54:6) [NL-20:4] | 7.52 | (4.68-10.44) | **1.91E-07** | **1.15E-05** | 3.69 | (2.71-4.68) | **2.17E-13** | **2.27E-12** | same |
| TG(54:6) [NL-20:5] | 8.39 | (5.04-11.85) | **7.54E-07** | **3.02E-05** | 3.82 | (2.52-5.13) | **7.53E-09** | **3.51E-08** | same |
| TG(54:6) [NL-22:6] | 8.20 | (4.48-12.06) | **1.29E-05** | **3.10E-04** | 5.76 | (4.33-7.22) | **4.96E-15** | **7.00E-14** | same |
| TG(54:7) [NL-20:5] | 8.46 | (5.22-11.8) | **2.49E-07** | **1.32E-05** | 3.45 | (2.17-4.75) | **1.37E-07** | **5.08E-07** | same |
| TG(54:7) [NL-22:6] | 10.42 | (6.51-14.47) | **1.23E-07** | **9.83E-06** | 5.09 | (3.68-6.52) | **1.79E-12** | **1.62E-11** | same |
| TG(56:6) [NL-20:4] | 3.72 | (1.91-5.57) | **5.78E-05** | **7.86E-04** | 1.36 | (0.85-1.88) | **2.38E-07** | **8.65E-07** | same |
| TG(56:6) [NL-22:5] | 3.96 | (1.82-6.14) | **2.87E-04** | **2.15E-03** | 1.99 | (1.2-2.77) | **7.33E-07** | **2.46E-06** | same |
| TG(56:7) [NL-20:4] | 4.16 | (1.73-6.64) | **7.60E-04** | **4.56E-03** | 1.88 | (1.14-2.62) | **6.63E-07** | **2.29E-06** | same |
| TG(56:7) [NL-20:5] | 5.58 | (2.64-8.6) | **1.82E-04** | **1.54E-03** | 2.47 | (1.35-3.61) | **1.56E-05** | **4.46E-05** | same |
| TG(56:7) [NL-22:5] | 5.15 | (2.46-7.91) | **1.64E-04** | **1.46E-03** | 2.55 | (1.61-3.5) | **1.14E-07** | **4.45E-07** | same |
| TG(56:7) [NL-22:6] | 6.45 | (3.01-10) | **2.16E-04** | **1.70E-03** | 3.96 | (2.65-5.28) | **3.33E-09** | **1.61E-08** | same |
| TG(56:8) [NL-20:4] | 5.12 | (2.54-7.75) | **9.23E-05** | **1.00E-03** | 1.88 | (1-2.77) | **2.85E-05** | **7.72E-05** | same |
| TG(56:8) [NL-20:5] | 5.03 | (2.01-8.13) | **1.03E-03** | **5.94E-03** | 1.59 | (0.43-2.77) | **7.20E-03** | **1.26E-02** | same |
| TG(56:8) [NL-22:6] | 7.57 | (4.02-11.24) | **2.51E-05** | **4.62E-04** | 3.97 | (2.64-5.31) | **4.96E-09** | **2.38E-08** | same |
| TG(56:9) [NL-22:6] | 6.66 | (3.29-10.14) | **9.49E-05** | **1.00E-03** | 3.07 | (1.89-4.27) | **3.82E-07** | **1.35E-06** | same |
| TG(58:10) [NL-22:6] | 5.17 | (1.89-8.56) | **1.90E-03** | **9.52E-03** | 2.08 | (0.92-3.25) | **4.34E-04** | **9.92E-04** | same |
| TG(58:8) [NL-22:6] | 4.73 | (1.39-8.18) | **5.34E-03** | **2.16E-02** | 2.80 | (1.58-4.04) | **7.21E-06** | **2.15E-05** | same |
| TG(58:9) [NL-22:6] | 4.51 | (1.17-7.97) | **7.98E-03** | **3.02E-02** | 2.17 | (0.97-3.38) | **3.75E-04** | **8.69E-04** | same |
| TG(O-50:1) [NL-17:1] | 4.99 | (2.47-7.57) | **1.01E-04** | **1.03E-03** | 4.08 | (3.14-5.03) | **4.97E-17** | **1.26E-15** | same |
| TG(O-52:2) [NL-17:1] | 4.89 | (2.19-7.66) | **3.63E-04** | **2.52E-03** | 3.82 | (2.87-4.77) | **4.43E-15** | **6.45E-14** | same |
| TG(O-52:2) [NL-18:1] | 5.04 | (2-8.16) | **1.08E-03** | **6.15E-03** | 2.26 | (1.27-3.27) | **8.29E-06** | **2.44E-05** | same |
| Cer(m18:0/22:0) | 2.43 | (0.5-4.39) | **1.34E-02** | **4.68E-02** | 2.95 | (2.15-3.76) | **8.19E-13** | **7.86E-12** | same |
| Cer(m18:0/24:1) | 4.35 | (2.12-6.63) | **1.28E-04** | **1.20E-03** | 2.84 | (2-3.68) | **3.94E-11** | **2.74E-10** | same |
| GM3(d18:1/24:0) | -2.41 | (-3.83--0.96) | **1.23E-03** | **6.55E-03** | -1.92 | (-2.46--1.39) | **6.41E-12** | **5.13E-11** | same |
| HexCer(d18:1/20:0) | -2.39 | (-4.02--0.73) | **5.05E-03** | **2.10E-02** | -1.29 | (-1.81--0.76) | **2.18E-06** | **6.88E-06** | same |
| HexCer(d18:1/22:0) | -2.35 | (-3.8--0.88) | **1.85E-03** | **9.34E-03** | -0.92 | (-1.41--0.43) | **2.61E-04** | **6.17E-04** | same |
| HexCer(d18:1/24:0) | -1.92 | (-3.37--0.46) | **1.03E-02** | **3.73E-02** | -1.34 | (-1.83--0.85) | **1.19E-07** | **4.57E-07** | same |
| Hex2Cer(d18:1/22:0) | -2.48 | (-4.09--0.83) | **3.43E-03** | **1.57E-02** | -1.43 | (-2--0.87) | **8.84E-07** | **2.91E-06** | same |
| Hex2Cer(d18:1/24:1) | -2.62 | (-4.35--0.85) | **3.91E-03** | **1.73E-02** | -1.91 | (-2.5--1.32) | **5.82E-10** | **3.21E-09** | same |
| Hex3Cer(d18:1/24:0) | -2.28 | (-3.7--0.83) | **2.21E-03** | **1.09E-02** | -1.90 | (-2.42--1.37) | **6.58E-12** | **5.18E-11** | same |
| SM(d18:1/14:0)/SM(d16:1/16:0) | 2.11 | (0.57-3.68) | **7.37E-03** | **2.85E-02** | 0.72 | (0.27-1.18) | **1.93E-03** | **3.86E-03** | same |
| SM(d18:2/14:0) | 6.29 | (4.83-7.77) | **2.07E-16** | **9.93E-14** | 1.49 | (1.01-1.98) | **2.12E-09** | **1.07E-08** | same |
| SM(34:3) | 2.92 | (1.38-4.49) | **2.12E-04** | **1.70E-03** | 0.97 | (0.49-1.46) | **8.92E-05** | **2.27E-04** | same |
| SM(38:3) (a) | 1.93 | (0.68-3.2) | **2.56E-03** | **1.24E-02** | 0.56 | (0.14-0.98) | **8.55E-03** | **1.47E-02** | same |
| SM(d16:1/24:1) | 2.62 | (1.12-4.15) | **6.11E-04** | **3.91E-03** | 0.55 | (0.05-1.05) | **3.15E-02** | **4.75E-02** | same |
| LPC(14:0) [sn2] | 4.86 | (2.45-7.33) | **7.65E-05** | **8.95E-04** | 1.18 | (0.41-1.96) | **2.81E-03** | **5.40E-03** | same |
| LPC(14:0) [sn1] | 4.70 | (2.35-7.09) | **8.28E-05** | **9.46E-04** | 1.53 | (0.78-2.3) | **7.13E-05** | **1.84E-04** | same |
| LPC(16:1) [sn2] | 5.09 | (3.19-7.03) | **1.66E-07** | **1.14E-05** | 0.95 | (0.37-1.53) | **1.29E-03** | **2.69E-03** | same |
| LPC(16:1) [sn1] | 3.93 | (2.1-5.79) | **2.67E-05** | **4.62E-04** | 0.87 | (0.34-1.4) | **1.33E-03** | **2.76E-03** | same |
| LPC(19:0) [sn1] (a) / LPC(19:0) [sn2] (b) | -2.26 | (-3.96--0.53) | **1.08E-02** | **3.85E-02** | -2.28 | (-2.79--1.76) | **6.69E-17** | **1.61E-15** | same |
| LPC(20:0) [sn1] | -3.35 | (-4.85--1.82) | **2.48E-05** | **4.62E-04** | -2.10 | (-2.58--1.62) | **1.05E-16** | **2.41E-15** | same |
| LPC(20:1) [sn1] | -2.32 | (-3.93--0.7) | **5.46E-03** | **2.19E-02** | -1.86 | (-2.38--1.33) | **1.33E-11** | **9.82E-11** | same |
| LPC(20:3) [sn2] | 2.63 | (0.79-4.5) | **4.98E-03** | **2.10E-02** | 1.19 | (0.56-1.83) | **2.39E-04** | **5.67E-04** | same |
| LPC(20:3) [104_sn1] | 2.53 | (0.51-4.6) | **1.42E-02** | **4.91E-02** | 0.93 | (0.28-1.58) | **4.92E-03** | **8.95E-03** | same |
| LPC(O-20:0) | -2.32 | (-4.14--0.47) | **1.46E-02** | **4.96E-02** | -2.23 | (-2.77--1.69) | **4.19E-15** | **6.29E-14** | same |
| LPC(O-22:0) | -1.92 | (-3.1--0.72) | **1.79E-03** | **9.15E-03** | -1.43 | (-1.85--1) | **1.03E-10** | **6.79E-10** | same |
| PC(32:1) | 3.56 | (1.41-5.75) | **1.12E-03** | **6.20E-03** | 2.58 | (1.77-3.39) | **4.13E-10** | **2.39E-09** | same |
| PC(14:0_20:4) | 4.75 | (2.3-7.26) | **1.38E-04** | **1.25E-03** | 1.14 | (0.28-2.01) | **9.67E-03** | **1.64E-02** | same |
| PC(34:5) | 7.01 | (3.69-10.43) | **3.00E-05** | **4.64E-04** | 1.64 | (0.4-2.9) | **9.61E-03** | **1.64E-02** | same |
| PC(17:0_18:2) | -2.03 | (-3.62--0.41) | **1.44E-02** | **4.92E-02** | -1.17 | (-1.68--0.65) | **1.30E-05** | **3.74E-05** | same |

| | | | | | | | | | |
|---|---|---|---|---|---|---|---|---|---|
| PC(16:0_20:3) (a) | -2.86 | (-4.42--1.29) | **4.47E-04** | **3.02E-03** | -1.61 | (-2.18--1.04) | **5.63E-08** | **2.33E-07** | same |
| PC(18:2_18:2) | -3.04 | (-5.24--0.78) | **8.73E-03** | **3.24E-02** | -2.49 | (-3.22--1.75) | **1.24E-10** | **7.80E-10** | same |
| PC(18:0_20:3) | 5.35 | (3.52-7.21) | **1.22E-08** | **1.46E-06** | 3.69 | (2.93-4.46) | **1.54E-20** | **1.06E-18** | same |
| PC(18:0_22:6) | 3.01 | (0.9-5.18) | **5.22E-03** | **2.12E-02** | 1.19 | (0.44-1.95) | **1.98E-03** | **3.94E-03** | same |
| PC(O-34:1) | -2.24 | (-3.3--1.18) | **4.84E-05** | **7.04E-04** | -1.21 | (-1.62--0.79) | **2.44E-08** | **1.08E-07** | same |
| PC(O-36:0) | -2.41 | (-3.76--1.03) | **6.74E-04** | **4.23E-03** | -0.84 | (-1.33--0.35) | **9.07E-04** | **1.95E-03** | same |
| PC(O-18:0/18:1) | -3.04 | (-4.22--1.85) | **1.05E-06** | **3.88E-05** | -1.48 | (-1.92--1.04) | **1.07E-10** | **6.87E-10** | same |
| PC(O-18:1/18:1) | -1.86 | (-2.95--0.75) | **1.12E-03** | **6.20E-03** | -1.71 | (-2.13--1.29) | **1.10E-14** | **1.42E-13** | same |
| PC(O-18:0/18:2) | -2.63 | (-4.12--1.12) | **7.48E-04** | **4.54E-03** | -1.89 | (-2.38--1.39) | **4.43E-13** | **4.43E-12** | same |
| PC(P-16:0/18:0) | -2.11 | (-3.09--1.13) | **3.73E-05** | **5.60E-04** | -1.47 | (-1.84--1.09) | **1.05E-13** | **1.20E-12** | same |
| PC(P-16:0/18:1) | -1.99 | (-3.17--0.79) | **1.27E-03** | **6.69E-03** | -2.19 | (-2.7--1.69) | **2.94E-16** | **6.14E-15** | same |
| PC(P-35:2) (b) | -2.72 | (-4.69--0.7) | **8.59E-03** | **3.22E-02** | -2.14 | (-2.83--1.45) | **2.27E-09** | **1.13E-08** | same |
| PC(P-18:0/18:2) | -2.97 | (-4.48--1.45) | **1.69E-04** | **1.48E-03** | -2.58 | (-3.1--2.06) | **1.54E-20** | **1.06E-18** | same |
| PE(16:0_16:1) | 4.35 | (0.88-7.95) | **1.38E-02** | **4.78E-02** | 3.03 | (1.74-4.33) | **3.84E-06** | **1.18E-05** | same |
| PE(18:0_20:3) (a) | 3.60 | (1.22-6.04) | **3.02E-03** | **1.41E-02** | 2.90 | (1.96-3.84) | **1.44E-09** | **7.41E-09** | same |
| PE(18:0_22:6) | 3.81 | (1.08-6.61) | **6.06E-03** | **2.36E-02** | 1.67 | (0.73-2.63) | **5.37E-04** | **1.20E-03** | same |
| PE(P-16:0/20:3) | 4.14 | (2.26-6.06) | **1.60E-05** | **3.36E-04** | 1.10 | (0.38-1.82) | **2.78E-03** | **5.35E-03** | same |
| PE(P-20:0/18:2) | -2.72 | (-4.31--1.11) | **1.09E-03** | **6.15E-03** | -1.98 | (-2.5--1.45) | **1.04E-12** | **9.79E-12** | same |
| PI(16:0_16:1) | 8.42 | (5.56-11.34) | **6.42E-09** | **1.03E-06** | 3.85 | (2.77-4.94) | **3.76E-12** | **3.11E-11** | same |
| PI(18:1_18:2) | -3.22 | (-4.99--1.41) | **5.62E-04** | **3.65E-03** | -1.29 | (-1.97--0.61) | **2.16E-04** | **5.17E-04** | same |

**Supplementary Table 4e.** Overlapping lipid species in the four lipid-adiposity association studies. Related to Figure 3.

| | **ppBMI (antenatal)** | | | | **Adult BMI (postnatal)** | | | | **Birth Weight (cord blood)** | | | | **Child BMI (six-year-old)** | | | | |
|---|---|---|---|---|---|---|---|---|---|---|---|---|---|---|---|---|---|
| Lipid Species | %change in lipid concentration per unit BMI | 95%CI | p-value | p-value(BH) | %change in lipid concentration per unit BMI | 95%CI | p-value | p-value(BH) | %change in lipid concentration per 100 grams | 95%CI | p-value | p-value(BH) | %change in lipid concentration per unit BMI | 95%CI | p-value | p-value(BH) | Trend |
| CE(16:1) | -1.06 | (-1.82--0.29) | **7.23E-03** | **1.46E-02** | 2.17 | (1.53-2.82) | **6.89E-11** | **4.60E-10** | 0.98 | (0.31-1.69) | **3.55E-03** | **8.00E-03** | 5.77 | (4.04-7.53) | **9.25E-11** | **2.22E-08** | different |
| DG(16:0_18:1) | 1.33 | (0.59-2.07) | **4.36E-04** | **1.22E-03** | 4.26 | (3.33-5.21) | **1.32E-18** | **4.86E-17** | -1.91 | (-2.57--1.19) | **1.35E-06** | **4.62E-06** | 3.90 | (1.38-6.49) | **2.40E-03** | **1.18E-02** | different |
| DG(16:0_18:2) | 0.92 | (0.21-1.65) | **1.15E-02** | **2.16E-02** | 3.62 | (2.72-4.52) | **5.53E-15** | **7.59E-14** | -3.07 | (-3.76--2.31) | **1.09E-11** | **6.22E-11** | 4.00 | (1.57-6.48) | **1.20E-03** | **6.49E-03** | different |
| DG(18:0_18:1) | 1.36 | (0.69-2.04) | **7.91E-05** | **2.60E-04** | 4.41 | (3.51-5.31) | **4.70E-21** | **4.51E-19** | -1.29 | (-1.99--0.53) | **1.23E-03** | **2.94E-03** | 4.06 | (1.71-6.46) | **6.87E-04** | **4.23E-03** | different |
| DG(18:0_18:2) | 0.99 | (0.23-1.76) | **1.04E-02** | **1.97E-02** | 4.40 | (3.41-5.39) | **7.51E-18** | **2.57E-16** | -1.62 | (-2.52--0.61) | **2.43E-03** | **5.62E-03** | 3.76 | (1.3-6.29) | **2.68E-03** | **1.27E-02** | different |
| DG(16:0_20:4) | 1.19 | (0.26-2.12) | **1.22E-02** | **2.27E-02** | 4.27 | (3.25-5.3) | **4.79E-16** | **9.59E-15** | -2.06 | (-2.83--1.21) | **9.20E-06** | **2.95E-05** | 4.97 | (2.34-7.66) | **1.98E-04** | **1.61E-03** | different |
| DG(18:1_20:4) | 1.14 | (0.4-1.88) | **2.42E-03** | **5.53E-03** | 2.47 | (1.71-3.22) | **1.81E-10** | **1.10E-09** | -2.06 | (-2.71--1.36) | **1.08E-07** | **4.14E-07** | 3.90 | (1.68-6.16) | **5.47E-04** | **3.60E-03** | different |
| DG(18:1_22:6) | 1.53 | (0.47-2.61) | **4.63E-03** | **9.97E-03** | 3.18 | (2.04-4.34) | **5.26E-08** | **2.22E-07** | -3.49 | (-4.15--2.76) | **7.44E-15** | **6.49E-14** | 4.14 | (1.32-7.04) | **3.88E-03** | **1.73E-02** | different |
| DG(18:2_22:6) | 1.52 | (0.42-2.63) | **6.55E-03** | **1.35E-02** | 3.05 | (1.87-4.24) | **3.60E-07** | **1.28E-06** | -4.00 | (-4.74--3.16) | **9.00E-14** | **6.75E-13** | 4.79 | (1.45-8.24) | **4.80E-03** | **2.04E-02** | different |
| TG(48:0) [NL-18:0] | -1.56 | (-2.43--0.68) | **5.76E-04** | **1.53E-03** | 4.34 | (2.99-5.71) | **3.54E-10** | **2.07E-09** | -1.44 | (-2.28--0.51) | **3.14E-03** | **7.14E-03** | 7.45 | (3.74-11.29) | **7.09E-05** | **8.54E-04** | different |
| TG(48:2) [NL-14:1] | -1.20 | (-2.25--0.13) | **2.78E-02** | **4.70E-02** | 4.79 | (3.42-6.18) | **7.51E-12** | **5.72E-11** | -3.06 | (-3.81--2.23) | **4.50E-10** | **2.23E-09** | 8.90 | (4.81-13.16) | **1.61E-05** | **3.36E-04** | different |
| TG(48:2) [NL-18:2] | -1.16 | (-2.2--0.12) | **2.91E-02** | **4.87E-02** | 5.21 | (3.68-6.77) | **2.87E-11** | **2.03E-10** | -4.14 | (-4.86--3.32) | **5.42E-15** | **4.91E-14** | 8.58 | (4.3-13.04) | **7.11E-05** | **8.54E-04** | different |
| TG(48:3) [NL-14:0] | -1.51 | (-2.64--0.37) | **9.55E-03** | **1.85E-02** | 4.32 | (2.9-5.76) | **2.31E-09** | **1.14E-08** | -4.04 | (-4.73--3.26) | **1.02E-15** | **1.04E-14** | 9.53 | (5.17-14.08) | **1.43E-05** | **3.28E-04** | different |
| TG(48:3) [NL-18:3] | -1.81 | (-3.07--0.53) | **5.76E-03** | **1.21E-02** | 4.99 | (3.41-6.59) | **5.23E-10** | **2.92E-09** | -3.23 | (-3.91--2.48) | **8.75E-13** | **5.83E-12** | 8.18 | (3.69-12.86) | **3.09E-04** | **2.27E-03** | different |
| TG(50:1) [NL-16:0] | 0.83 | (0.22-1.43) | **7.27E-03** | **1.46E-02** | 4.34 | (3.39-5.3) | **1.03E-18** | **4.12E-17** | -2.49 | (-3.25--1.64) | **2.17E-07** | **8.15E-07** | 4.63 | (2.1-7.22) | **3.13E-04** | **2.27E-03** | different |
| TG(50:1) [NL-18:1] | 0.76 | (0.18-1.34) | **1.03E-02** | **1.97E-02** | 4.10 | (3.16-5.05) | **4.06E-17** | **1.08E-15** | -2.47 | (-3.24--1.62) | **2.61E-07** | **9.63E-07** | 4.17 | (1.73-6.68) | **7.99E-04** | **4.74E-03** | different |
| TG(50:2) [NL-14:0] | -0.65 | (-1.21--0.07) | **2.69E-02** | **4.57E-02** | 2.80 | (1.76-3.86) | **1.50E-07** | **5.54E-07** | -3.34 | (-4.1--2.48) | **1.16E-10** | **6.01E-10** | 4.36 | (1.14-7.69) | **7.83E-03** | **3.01E-02** | different |
| TG(50:2) [NL-18:2] | 0.66 | (0.08-1.24) | **2.58E-02** | **4.41E-02** | 3.72 | (2.82-4.62) | **8.45E-16** | **1.53E-14** | -4.17 | (-4.92--3.31) | **5.26E-14** | **4.14E-13** | 4.53 | (2.11-7) | **2.30E-04** | **1.78E-03** | different |
| TG(52:1) [NL-18:0] | 1.38 | (0.59-2.18) | **6.58E-04** | **1.73E-03** | 5.69 | (4.53-6.86) | **2.15E-21** | **2.58E-19** | -1.86 | (-2.76--0.83) | **6.81E-04** | **1.69E-03** | 5.79 | (2.8-8.86) | **1.32E-04** | **1.22E-03** | different |
| TG(52:1) [NL-18:1] | 1.45 | (0.69-2.23) | **2.04E-04** | **6.17E-04** | 5.86 | (4.73-7) | **1.79E-23** | **8.60E-21** | -1.80 | (-2.71--0.77) | **9.96E-04** | **2.41E-03** | 5.71 | (2.83-8.68) | **9.59E-05** | **1.00E-03** | different |
| TG(54:1) [NL-18:1] | 0.99 | (0.32-1.66) | **3.57E-03** | **8.00E-03** | 4.45 | (3.57-5.34) | **4.54E-22** | **7.26E-20** | -1.64 | (-2.56--0.59) | **2.89E-03** | **6.61E-03** | 3.54 | (1.13-6.02) | **3.96E-03** | **1.73E-02** | different |
| TG(54:2) [NL-18:0] | 1.41 | (0.72-2.1) | **6.55E-05** | **2.23E-04** | 4.24 | (3.35-5.13) | **2.72E-20** | **1.63E-18** | -1.85 | (-2.69--0.91) | **2.40E-04** | **6.39E-04** | 3.30 | (0.94-5.71) | **6.02E-03** | **2.36E-02** | different |
| TG(54:3) [NL-18:2] | 0.72 | (0.14-1.3) | **1.43E-02** | **2.59E-02** | 2.23 | (1.57-2.89) | **5.13E-11** | **3.51E-10** | -3.06 | (-3.75--2.3) | **1.28E-11** | **7.23E-11** | 2.32 | (0.48-4.19) | **1.32E-02** | **4.65E-02** | different |
| TG(54:5) [NL-20:4] | 1.04 | (0.38-1.71) | **1.91E-03** | **4.51E-03** | 3.37 | (2.55-4.2) | **2.12E-15** | **3.39E-14** | -3.73 | (-4.49--2.86) | **4.40E-12** | **2.64E-11** | 5.84 | (3.43-8.3) | **1.90E-06** | **6.49E-05** | different |
| TG(54:6) [NL-20:4] | 1.16 | (0.33-2) | **6.24E-03** | **1.29E-02** | 3.69 | (2.71-4.68) | **2.17E-13** | **2.27E-12** | -4.11 | (-4.83--3.28) | **1.19E-14** | **9.81E-14** | 7.52 | (4.68-10.44) | **1.91E-07** | **1.15E-05** | different |
| TG(54:6) [NL-22:6] | 1.79 | (0.57-3.02) | **4.10E-03** | **9.06E-03** | 5.76 | (4.33-7.22) | **4.96E-15** | **7.00E-14** | -5.16 | (-5.83--4.38) | **1.98E-20** | **5.23E-19** | 8.20 | (4.48-12.06) | **1.29E-05** | **3.10E-04** | different |
| TG(56:6) [NL-20:4] | 1.34 | (0.81-1.87) | **7.12E-07** | **3.60E-06** | 1.36 | (0.85-1.88) | **2.38E-07** | **8.65E-07** | -3.00 | (-3.77--2.15) | **1.89E-09** | **8.92E-09** | 3.72 | (1.91-5.57) | **5.78E-05** | **7.86E-04** | different |
| TG(56:7) [NL-20:4] | 1.02 | (0.35-1.7) | **2.84E-03** | **6.40E-03** | 1.88 | (1.14-2.62) | **6.63E-07** | **2.29E-06** | -3.66 | (-4.41--2.81) | **3.38E-12** | **2.08E-11** | 4.16 | (1.73-6.64) | **7.60E-04** | **4.56E-03** | different |
| TG(56:7) [NL-22:6] | 1.93 | (0.84-3.04) | **4.97E-04** | **1.35E-03** | 3.96 | (2.65-5.28) | **3.33E-09** | **1.61E-08** | -5.07 | (-5.72--4.33) | **1.85E-21** | **7.12E-20** | 6.45 | (3.01-10) | **2.16E-04** | **1.70E-03** | different |
| TG(56:8) [NL-22:6] | 1.76 | (0.54-2.99) | **4.49E-03** | **9.76E-03** | 3.97 | (2.64-5.31) | **4.96E-09** | **2.38E-08** | -5.36 | (-5.97--4.65) | **1.69E-24** | **1.02E-22** | 7.57 | (4.02-11.24) | **2.51E-05** | **4.62E-04** | different |
| TG(58:8) [NL-22:6] | 1.46 | (0.38-2.56) | **7.98E-03** | **1.58E-02** | 2.80 | (1.58-4.04) | **7.21E-06** | **2.15E-05** | -4.50 | (-5.16--3.76) | **3.67E-19** | **7.05E-18** | 4.73 | (1.39-8.18) | **5.34E-03** | **2.16E-02** | different |
| Cer(m18:0/24:1) | 1.15 | (0.26-2.04) | **1.11E-02** | **2.08E-02** | 2.84 | (2-3.68) | **3.94E-11** | **2.74E-10** | -1.46 | (-2.42--0.37) | **9.81E-03** | **2.02E-02** | 4.35 | (2.12-6.63) | **1.28E-04** | **1.20E-03** | different |
| LPC(20:1) [sn1] | -0.62 | (-1.08--0.15) | **9.49E-03** | **1.84E-02** | -1.86 | (-2.38--1.33) | **1.33E-11** | **9.82E-11** | 1.75 | (1.27-2.25) | **1.55E-13** | **1.12E-12** | -2.32 | (-3.93--0.7) | **5.46E-03** | **2.19E-02** | different |
| LPC(20:3) [sn2] | 1.05 | (0.4-1.7) | **1.48E-03** | **3.59E-03** | 1.19 | (0.56-1.83) | **2.39E-04** | **5.67E-04** | 2.58 | (1.91-3.3) | **1.79E-15** | **1.72E-14** | 2.63 | (0.79-4.5) | **4.98E-03** | **2.10E-02** | same |
| LPC(20:3) [104_sn1] | 1.42 | (0.72-2.12) | **7.46E-05** | **2.47E-04** | 0.93 | (0.28-1.58) | **4.92E-03** | **8.95E-03** | 3.22 | (2.49-3.99) | **9.49E-21** | **2.85E-19** | 2.53 | (0.51-4.6) | **1.42E-02** | **4.91E-02** | same |
| PC(17:0_18:2) | -1.54 | (-2.08--1) | **3.86E-08** | **2.57E-07** | -1.17 | (-1.68--0.65) | **1.30E-05** | **3.74E-05** | -1.28 | (-1.84--0.68) | **5.78E-05** | **1.69E-04** | -2.03 | (-3.62--0.41) | **1.44E-02** | **4.92E-02** | same |
| PC(O-18:0/18:1) | -1.51 | (-1.98--1.03) | **1.00E-09** | **1.02E-08** | -1.48 | (-1.92--1.04) | **1.07E-10** | **6.87E-10** | 1.31 | (0.62-2.05) | **1.35E-04** | **3.71E-04** | -3.04 | (-4.22--1.85) | **1.05E-06** | **3.88E-05** | different |
| PC(O-18:1/18:1) | -1.20 | (-1.66--0.74) | **4.16E-07** | **2.24E-06** | -1.71 | (-2.13--1.29) | **1.10E-14** | **1.42E-13** | 0.71 | (0.11-1.35) | **2.01E-02** | **3.86E-02** | -1.86 | (-2.95--0.75) | **1.12E-03** | **6.20E-03** | different |
| PC(P-35:2) (b) | -1.83 | (-2.55--1.1) | **1.07E-06** | **5.20E-06** | -2.14 | (-2.83--1.45) | **2.27E-09** | **1.13E-08** | -1.45 | (-2.06--0.79) | **3.89E-05** | **1.15E-04** | -2.72 | (-4.69--0.7) | **8.59E-03** | **3.22E-02** | same |

| | | | | | | | | | | | | | | | | | |
|---|---|---|---|---|---|---|---|---|---|---|---|---|---|---|---|---|---|
| PE(18:0_22:6) | 1.34 | (0.55-2.14) | **8.34E-04** | **2.13E-03** | 1.67 | (0.73-2.63) | **5.37E-04** | **1.20E-03** | -1.61 | (-2.34--0.8) | **2.02E-04** | **5.43E-04** | 3.81 | (1.08-6.61) | **6.06E-03** | **2.36E-02** | different |
| PI(18:1_18:2) | -2.66 | (-3.29--2.03) | **1.11E-15** | **5.91E-14** | -1.29 | (-1.97--0.61) | **2.16E-04** | **5.17E-04** | -1.47 | (-1.98--0.93) | **4.93E-07** | **1.78E-06** | -3.22 | (-4.99--1.41) | **5.62E-04** | **3.65E-03** | same |

**Supplementary Table 5a**. Sex difference in association studies with birth weight and child BMI in terms of class levels. Related to Suplementry Figure 4.

| | **Birth Weight (cord blood) - Male** adjusted for ethnicity, maternal age, maternal education level, pre-pregnancy BMI, total gestational weight gain, gestational age and parity | | | | **Birth Weight (cord blood) -Female** adjusted for ethnicity, maternal age, maternal education level, pre-pregnancy BMI, total gestational weight gain, gestational age and parity | | | | **Child BMI (six-year-old) - Male** adjusted for ethnicity and maternal education level | | | | **Child BMI (six-year-old) -Female** adjusted for ethnicity and maternal education level | | | |
|---|---|---|---|---|---|---|---|---|---|---|---|---|---|---|---|---|
| Lipid Class | %change in lipid concentration per 100 grams | 95%CI | p-value | p-value(BH) | %change in lipid concentration per 100 grams | 95%CI | p-value | p-value(BH) | %change in lipid concentration per unit BMI | 95%CI | p-value | p-value(BH) | %change in lipid concentration per unit BMI | 95%CI | p-value | p-value(BH) |
| AC | -0.87 | (-1.54--0.15) | **1.90E-02** | **3.96E-02** | -0.06 | (-0.91- 0.87) | 8.95E-01 | 9.66E-01 | -0.13 | (-2.08-1.86) | 8.98E-01 | 9.35E-01 | -1.79 | (-4.19- 0.68) | 1.53E-01 | 4.77E-01 |
| CE | -0.44 | (-0.99- 0.14) | 1.36E-01 | 2.12E-01 | -0.06 | (-0.79- 0.74) | 8.84E-01 | 9.66E-01 | 0.36 | (-0.87-1.61) | 5.63E-01 | 7.19E-01 | 0.71 | (-0.70- 2.13) | 3.22E-01 | 5.75E-01 |
| DG | -2.70 | (-3.37--1.97) | **3.67E-10** | **2.29E-09** | -1.42 | (-2.40--0.32) | **1.34E-02** | **4.17E-02** | 1.75 | (-1.03-4.60) | 2.18E-01 | 6.07E-01 | 4.47 | ( 1.03- 8.02) | **1.08E-02** | 9.02E-02 |
| TG | -3.85 | (-4.60--3.01) | **9.57E-13** | **1.20E-11** | -2.25 | (-3.47--0.80) | **3.73E-03** | **1.33E-02** | 3.11 | ( 0.51-5.78) | **1.89E-02** | 4.73E-01 | 4.46 | ( 1.23- 7.80) | **6.78E-03** | 8.48E-02 |
| TG(O) | -3.00 | (-3.77--2.14) | **4.15E-09** | **2.08E-08** | -2.25 | (-3.29--1.04) | **6.13E-04** | **2.55E-03** | 2.70 | (-0.12-5.59) | 6.05E-02 | 6.07E-01 | 3.59 | (-0.10- 7.40) | 5.63E-02 | 2.81E-01 |
| dhCer | -0.17 | (-1.00- 0.74) | 7.03E-01 | 8.36E-01 | -0.41 | (-1.38- 0.67) | 4.37E-01 | 8.87E-01 | 0.94 | (-1.19-3.12) | 3.88E-01 | 6.47E-01 | 0.67 | (-1.69- 3.08) | 5.80E-01 | 7.94E-01 |
| Cer | -0.63 | (-1.30- 0.08) | 8.20E-02 | 1.46E-01 | -0.01 | (-0.90- 0.97) | 9.81E-01 | 9.81E-01 | 0.60 | (-1.14-2.36) | 5.02E-01 | 6.97E-01 | 0.48 | (-1.32- 2.33) | 6.00E-01 | 7.94E-01 |
| DeoxyCer | -2.13 | (-2.82--1.37) | **5.38E-07** | **2.24E-06** | -1.97 | (-2.84--0.99) | **2.03E-04** | **1.02E-03** | 1.15 | (-0.99-3.33) | 2.95E-01 | 6.20E-01 | 2.73 | (-0.07- 5.61) | 5.59E-02 | 2.81E-01 |
| GM3 | -0.41 | (-1.25- 0.50) | 3.60E-01 | 4.91E-01 | -0.17 | (-1.25- 1.05) | 7.78E-01 | 9.66E-01 | -1.01 | (-2.59-0.59) | 2.14E-01 | 6.07E-01 | -1.94 | (-4.02- 0.18) | 7.18E-02 | 2.96E-01 |
| HexCer | 0.33 | (-0.63- 1.39) | 5.16E-01 | 6.44E-01 | 0.06 | (-1.03- 1.29) | 9.15E-01 | 9.66E-01 | -0.89 | (-2.66-0.91) | 3.29E-01 | 6.28E-01 | -2.84 | (-4.69--0.95) | **3.67E-03** | 8.48E-02 |
| Hex2Cer | -0.41 | (-1.26- 0.52) | 3.74E-01 | 4.91E-01 | -0.41 | (-1.46- 0.77) | 4.82E-01 | 8.87E-01 | -1.03 | (-3.16-1.16) | 3.52E-01 | 6.28E-01 | -1.47 | (-4.02- 1.15) | 2.68E-01 | 5.15E-01 |
| Hex3Cer | -0.39 | (-1.12- 0.40) | 3.27E-01 | 4.81E-01 | -0.35 | (-1.29- 0.70) | 4.99E-01 | 8.87E-01 | -1.00 | (-2.45-0.47) | 1.78E-01 | 6.07E-01 | -1.19 | (-2.97- 0.63) | 1.98E-01 | 4.78E-01 |
| SM | -0.09 | (-0.77- 0.64) | 8.03E-01 | 8.36E-01 | -0.05 | (-0.90- 0.88) | 9.10E-01 | 9.66E-01 | -0.57 | (-2.14-1.03) | 4.81E-01 | 6.97E-01 | -0.38 | (-2.32- 1.59) | 6.99E-01 | 7.94E-01 |
| LPC | 2.24 | ( 1.55- 2.97) | **2.84E-11** | **2.37E-10** | 2.30 | ( 1.49- 3.17) | **5.89E-09** | **7.37E-08** | -0.86 | (-3.09-1.41) | 4.52E-01 | 6.97E-01 | 1.77 | (-0.96- 4.58) | 2.05E-01 | 4.78E-01 |
| LPC(O) | 1.34 | ( 0.75- 1.96) | **5.46E-06** | **1.71E-05** | 1.33 | ( 0.66- 2.05) | **7.05E-05** | **4.41E-04** | -1.44 | (-3.53-0.70) | 1.84E-01 | 6.07E-01 | 1.07 | (-1.41- 3.61) | 3.99E-01 | 6.66E-01 |
| LPC(P) | 1.51 | ( 0.88- 2.17) | **1.27E-06** | **4.54E-06** | 1.44 | ( 0.73- 2.19) | **4.20E-05** | **3.50E-04** | -1.31 | (-3.56-0.99) | 2.60E-01 | 6.20E-01 | 1.59 | (-1.04- 4.29) | 2.36E-01 | 4.91E-01 |
| LPE | 2.54 | ( 1.82- 3.30) | **3.66E-13** | **9.16E-12** | 2.67 | ( 1.78- 3.62) | **5.31E-10** | **1.33E-08** | -1.66 | (-3.45-0.17) | 7.42E-02 | 6.07E-01 | -0.61 | (-2.99- 1.82) | 6.18E-01 | 7.94E-01 |
| PC | -0.56 | (-1.19- 0.12) | 1.04E-01 | 1.74E-01 | -0.42 | (-1.24- 0.47) | 3.44E-01 | 8.60E-01 | -0.21 | (-1.73-1.33) | 7.82E-01 | 8.51E-01 | -0.20 | (-2.08- 1.73) | 8.39E-01 | 9.03E-01 |
| PC(O) | -0.10 | (-0.84- 0.69) | 7.92E-01 | 8.36E-01 | -0.04 | (-0.94- 0.94) | 9.28E-01 | 9.66E-01 | 0.02 | (-1.71-1.78) | 9.82E-01 | 9.82E-01 | -0.05 | (-2.12- 2.06) | 9.61E-01 | 9.61E-01 |
| PC(P) | -1.00 | (-1.68--0.27) | **8.01E-03** | **2.00E-02** | -0.71 | (-1.57- 0.24) | 1.35E-01 | 3.76E-01 | -0.32 | (-2.14-1.53) | 7.30E-01 | 8.29E-01 | -0.50 | (-2.70- 1.75) | 6.59E-01 | 7.94E-01 |
| PE | -1.12 | (-1.79--0.39) | **3.12E-03** | **8.68E-03** | -0.45 | (-1.43- 0.63) | 3.94E-01 | 8.87E-01 | 0.74 | (-2.19-3.76) | 6.24E-01 | 7.42E-01 | -0.76 | (-4.33- 2.94) | 6.81E-01 | 7.94E-01 |
| PE(O) | -0.08 | (-0.84- 0.74) | 8.36E-01 | 8.36E-01 | 0.06 | (-0.88- 1.10) | 9.06E-01 | 9.66E-01 | 0.68 | (-1.70-3.13) | 5.75E-01 | 7.19E-01 | 1.01 | (-1.85- 3.96) | 4.91E-01 | 7.67E-01 |
| PE(P) | -0.11 | (-0.83- 0.66) | 7.63E-01 | 8.36E-01 | 0.20 | (-0.78- 1.29) | 7.03E-01 | 9.66E-01 | 1.42 | (-0.67-3.54) | 1.83E-01 | 6.07E-01 | 1.57 | (-0.89- 4.09) | 2.11E-01 | 4.78E-01 |
| PG | -1.47 | (-2.50--0.31) | **1.49E-02** | **3.39E-02** | -0.23 | (-1.74- 1.56) | 7.86E-01 | 9.66E-01 | 1.59 | (-1.39-4.66) | 2.98E-01 | 6.20E-01 | 3.25 | (-0.42- 7.06) | 8.29E-02 | 2.96E-01 |
| PI | -0.58 | (-1.07--0.06) | **2.80E-02** | 5.38E-02 | -0.23 | (-0.90- 0.50) | 5.32E-01 | 8.87E-01 | -1.23 | (-2.80-0.37) | 1.30E-01 | 6.07E-01 | -0.18 | (-2.27- 1.95) | 8.67E-01 | 9.03E-01 |

**Supplementary Table 5b**. Sex difference in association studies with birth weight and child BMI in terms of species levels. Related to Supplementry Figure 4.

| Studies | | **Birth Weight (cord blood) - Male** | | | | **Birth Weight (cord blood) - Female** | | | | **Child BMI (six-year-old) - Male** | | | | **Child BMI (six-year-old) - Female** | | | |
|---|---|---|---|---|---|---|---|---|---|---|---|---|---|---|---|---|---|
| Covariates | | adjusted for ethnicity, maternal age, maternal education level, pre-pregnancy BMI, total gestational weight gain, gestational age and parity | | | | adjusted for ethnicity, maternal age, maternal education level, pre-pregnancy BMI, total gestational weight gain, gestational age and parity | | | | adjusted for ethnicity and maternal education level | | | | adjusted for ethnicity and maternal education level | | | |
| No. | Lipid Species | %change in lipid concentration per 100 grams | 95%CI | p-value | p-value(BH) | %change in lipid concentration per 100 grams | 95%CI | p-value | p-value(BH) | %change in lipid concentration per unit BMI | 95%CI | p-value | p-value(BH) | %change in lipid concentration per unit BMI | 95%CI | p-value | p-value(BH) |
| 1 | AC(12:0) | -1.55 | (-2.29--0.75) | **2.80E-04** | **8.56E-04** | -1.39 | (-2.33--0.33) | **1.19E-02** | **4.10E-02** | -0.10 | (-3.26- 3.15) | 9.49E-01 | 9.83E-01 | -2.70 | (-6.60- 1.35) | 1.87E-01 | 3.70E-01 |
| 2 | AC(13:0) | -1.05 | (-2.40- 0.54) | 1.83E-01 | 2.90E-01 | -1.14 | (-2.80- 0.89) | 2.49E-01 | 4.38E-01 | 3.09 | (-1.44- 7.82) | 1.83E-01 | 4.24E-01 | -1.31 | (-6.81- 4.51) | 6.50E-01 | 8.08E-01 |
| 3 | AC(14:0) | -1.18 | (-1.87--0.42) | **3.07E-03** | **8.02E-03** | -0.61 | (-1.54- 0.43) | 2.43E-01 | 4.31E-01 | 0.19 | (-2.16- 2.60) | 8.75E-01 | 9.47E-01 | -2.93 | (-5.92- 0.14) | 6.12E-02 | 1.88E-01 |
| 4 | AC(14:1) | -0.99 | (-1.84--0.05) | **3.96E-02** | 7.73E-02 | -1.10 | (-2.13- 0.06) | 6.17E-02 | 1.56E-01 | -0.67 | (-4.11- 2.88) | 7.04E-01 | 8.83E-01 | -3.22 | (-7.33- 1.06) | 1.37E-01 | 3.14E-01 |
| 5 | AC(14:2) | -1.15 | (-1.83--0.41) | **2.88E-03** | **7.61E-03** | -0.59 | (-1.49- 0.41) | 2.40E-01 | 4.30E-01 | -0.20 | (-3.28- 2.98) | 9.02E-01 | 9.64E-01 | -2.92 | (-6.94- 1.28) | 1.69E-01 | 3.54E-01 |
| 6 | AC(16:0) | -0.83 | (-1.49--0.11) | **2.41E-02** | 5.01E-02 | 0.25 | (-0.63- 1.20) | 5.89E-01 | 7.63E-01 | 0.21 | (-1.33- 1.76) | 7.92E-01 | 9.16E-01 | -1.11 | (-3.00- 0.81) | 2.53E-01 | 4.44E-01 |
| 7 | AC(16:1) | -1.15 | (-1.98--0.25) | **1.40E-02** | **3.08E-02** | -0.62 | (-1.64- 0.54) | 2.82E-01 | 4.75E-01 | 0.97 | (-1.63- 3.64) | 4.67E-01 | 7.19E-01 | -1.48 | (-4.69- 1.83) | 3.73E-01 | 5.66E-01 |
| 8 | AC(18:0) | -1.07 | (-1.70--0.38) | **2.94E-03** | **7.72E-03** | -0.25 | (-1.12- 0.69) | 5.85E-01 | 7.61E-01 | -1.51 | (-3.37- 0.38) | 1.16E-01 | 3.23E-01 | -2.29 | (-4.23--0.31) | **2.42E-02** | 1.13E-01 |
| 9 | AC(18:1) | -0.19 | (-1.08- 0.78) | 6.86E-01 | 7.66E-01 | 0.76 | (-0.31- 1.94) | 1.69E-01 | 3.37E-01 | -0.29 | (-2.16- 1.62) | 7.66E-01 | 9.12E-01 | -0.68 | (-2.84- 1.53) | 5.43E-01 | 7.15E-01 |
| 10 | AC(18:2) | -1.15 | (-1.90--0.32) | **7.70E-03** | **1.82E-02** | 0.80 | (-0.23- 1.94) | 1.33E-01 | 2.80E-01 | 0.07 | (-1.58- 1.75) | 9.34E-01 | 9.76E-01 | -0.58 | (-2.82- 1.71) | 6.12E-01 | 7.74E-01 |
| 11 | CE(14:0) | -0.36 | (-1.04- 0.36) | 3.17E-01 | 4.49E-01 | -0.68 | (-1.50- 0.21) | 1.30E-01 | 2.75E-01 | 2.68 | ( 0.50- 4.90) | **1.59E-02** | 9.00E-02 | 1.44 | (-1.34- 4.29) | 3.10E-01 | 5.08E-01 |
| 12 | CE(16:1) | 1.66 | ( 0.73- 2.66) | **3.09E-04** | **9.28E-04** | 0.19 | (-0.76- 1.24) | 7.07E-01 | 8.47E-01 | 5.39 | ( 3.07- 7.76) | **6.25E-06** | **1.50E-03** | 6.14 | ( 3.51- 8.85) | **6.52E-06** | **1.04E-03** |
| 13 | CE(18:0) | 0.37 | (-0.68- 1.54) | 5.01E-01 | 6.25E-01 | 0.32 | (-0.81- 1.60) | 5.90E-01 | 7.63E-01 | -0.70 | (-2.78- 1.44) | 5.18E-01 | 7.51E-01 | 2.88 | (-0.05- 5.89) | 5.41E-02 | 1.76E-01 |
| 14 | CE(18:1) | 0.38 | (-0.21- 1.02) | 2.13E-01 | 3.28E-01 | -0.15 | (-0.78- 0.51) | 6.38E-01 | 8.04E-01 | 0.10 | (-0.92- 1.13) | 8.48E-01 | 9.40E-01 | 0.71 | (-0.63- 2.07) | 2.96E-01 | 4.94E-01 |
| 15 | CE(18:2) | -0.77 | (-1.37--0.12) | **2.02E-02** | **4.22E-02** | 0.13 | (-0.70- 1.04) | 7.60E-01 | 8.85E-01 | 0.13 | (-1.29- 1.57) | 8.59E-01 | 9.40E-01 | 0.65 | (-0.78- 2.09) | 3.74E-01 | 5.66E-01 |
| 16 | CE(18:3) | 0.10 | (-0.98- 1.31) | 8.60E-01 | 9.13E-01 | 0.68 | (-0.71- 2.27) | 3.52E-01 | 5.54E-01 | 2.38 | (-0.29- 5.11) | 8.07E-02 | 2.53E-01 | 3.38 | (-0.03- 6.90) | 5.17E-02 | 1.71E-01 |
| 17 | CE(20:1) | 1.28 | ( 0.11- 2.58) | **3.06E-02** | 6.24E-02 | -0.37 | (-1.47- 0.88) | 5.44E-01 | 7.32E-01 | -1.79 | (-3.52--0.02) | **4.70E-02** | 1.97E-01 | -0.44 | (-2.40- 1.55) | 6.58E-01 | 8.10E-01 |
| 18 | CE(20:2) | 1.17 | ( 0.30- 2.12) | **7.56E-03** | **1.80E-02** | 0.23 | (-0.68- 1.22) | 6.38E-01 | 8.04E-01 | 0.00 | (-1.42- 1.44) | 9.99E-01 | 9.99E-01 | 0.94 | (-0.79- 2.70) | 2.86E-01 | 4.81E-01 |
| 19 | CE(20:4) | -0.38 | (-1.07- 0.37) | 3.11E-01 | 4.44E-01 | 0.18 | (-0.76- 1.21) | 7.14E-01 | 8.47E-01 | 0.77 | (-1.10- 2.67) | 4.20E-01 | 6.78E-01 | -0.10 | (-2.65- 2.52) | 9.42E-01 | 9.75E-01 |
| 20 | CE(20:5) | -2.79 | (-4.02--1.30) | **6.99E-04** | **2.00E-03** | -1.80 | (-3.35- 0.12) | 6.47E-02 | 1.61E-01 | 4.14 | ( 0.08- 8.37) | **4.58E-02** | 1.95E-01 | 3.58 | (-2.12- 9.61) | 2.21E-01 | 4.16E-01 |
| 21 | CE(22:5) (n6) | 0.42 | (-0.56- 1.51) | 4.11E-01 | 5.44E-01 | 0.44 | (-0.74- 1.76) | 4.80E-01 | 6.66E-01 | 0.91 | (-0.98- 2.84) | 3.44E-01 | 6.22E-01 | 1.14 | (-0.95- 3.27) | 2.87E-01 | 4.81E-01 |
| 22 | CE(22:6) | -2.22 | (-3.18--1.13) | **1.99E-04** | **6.27E-04** | -1.50 | (-2.75--0.05) | **4.32E-02** | 1.18E-01 | 2.64 | (-0.21- 5.56) | 6.93E-02 | 2.38E-01 | 1.44 | (-1.89- 4.88) | 3.98E-01 | 5.90E-01 |
| 23 | CE(24:5) | -2.66 | (-3.77--1.34) | **2.75E-04** | **8.45E-04** | -3.18 | (-4.29--1.85) | **3.17E-05** | **3.05E-04** | -2.06 | (-4.70- 0.66) | 1.35E-01 | 3.48E-01 | -0.11 | (-2.85- 2.70) | 9.37E-01 | 9.75E-01 |
| 24 | CE(24:6) | -0.72 | (-1.96- 0.72) | 3.11E-01 | 4.44E-01 | -1.59 | (-2.88--0.06) | **4.25E-02** | 1.16E-01 | -0.14 | (-2.51- 2.29) | 9.08E-01 | 9.66E-01 | 2.09 | (-1.39- 5.69) | 2.41E-01 | 4.32E-01 |
| 25 | DG(16:0_16:1) | -1.88 | (-2.82--0.81) | **9.94E-04** | **2.76E-03** | -0.43 | (-1.79- 1.15) | 5.71E-01 | 7.51E-01 | 5.60 | ( 2.49- 8.79) | **4.07E-04** | **1.50E-02** | 7.08 | ( 2.48-11.88) | **2.47E-03** | **2.97E-02** |
| 26 | DG(14:0_18:2) | -3.22 | (-4.38--1.82) | **5.89E-05** | **2.02E-04** | -2.69 | (-4.28--0.66) | **1.23E-02** | **4.20E-02** | 4.96 | ( 0.30- 9.84) | **3.66E-02** | 1.64E-01 | 9.36 | ( 3.33-15.75) | **2.21E-03** | **2.72E-02** |
| 27 | DG(16:0_18:1) | -2.54 | (-3.32--1.67) | **3.15E-07** | **1.41E-06** | -0.88 | (-2.05- 0.45) | 1.84E-01 | 3.57E-01 | 2.76 | (-0.43- 6.05) | 9.02E-02 | 2.70E-01 | 5.44 | ( 1.29- 9.76) | **1.01E-02** | 6.90E-02 |
| 28 | DG(16:1_18:1) | -2.51 | (-3.34--1.57) | **2.13E-06** | **8.66E-06** | -1.33 | (-2.53- 0.06) | 5.94E-02 | 1.53E-01 | 5.07 | ( 1.31- 8.98) | **8.12E-03** | 6.04E-02 | 7.86 | ( 3.21-12.72) | **8.87E-04** | **1.90E-02** |
| 29 | DG(16:0_18:2) | -3.79 | (-4.59--2.88) | **3.80E-11** | **3.04E-10** | -1.96 | (-3.17--0.53) | **9.02E-03** | **3.16E-02** | 2.63 | (-0.54- 5.90) | 1.04E-01 | 2.94E-01 | 5.97 | ( 2.18- 9.90) | **2.02E-03** | **2.63E-02** |
| 30 | DG(18:0_18:1) | -1.73 | (-2.56--0.81) | **4.38E-04** | **1.29E-03** | -0.61 | (-1.82- 0.78) | 3.72E-01 | 5.75E-01 | 2.91 | ( 0.07- 5.84) | **4.47E-02** | 1.93E-01 | 5.77 | ( 1.69-10.00) | **5.45E-03** | **4.59E-02** |
| 31 | DG(18:1_18:1) | -2.22 | (-3.01--1.33) | **6.57E-06** | **2.50E-05** | -1.24 | (-2.31--0.02) | **4.73E-02** | 1.27E-01 | 1.16 | (-1.81- 4.23) | 4.45E-01 | 6.97E-01 | 3.38 | (-0.31- 7.20) | 7.26E-02 | 2.10E-01 |
| 32 | DG(18:0_18:2) | -2.55 | (-3.58--1.36) | **1.20E-04** | **3.97E-04** | -0.19 | (-1.81- 1.76) | 8.38E-01 | 9.33E-01 | 2.56 | (-0.52- 5.74) | 1.03E-01 | 2.93E-01 | 5.71 | ( 1.57-10.02) | **6.78E-03** | 5.51E-02 |
| 33 | DG(18:1_18:2) | -3.75 | (-4.45--2.96) | **9.13E-14** | **1.14E-12** | -2.19 | (-3.30--0.90) | **1.57E-03** | **6.92E-03** | 0.82 | (-2.10- 3.83) | 5.85E-01 | 8.05E-01 | 3.57 | ( 0.14- 7.13) | **4.14E-02** | 1.54E-01 |
| 34 | DG(18:2_18:2) | -4.90 | (-5.67--3.99) | **1.12E-14** | **1.85E-13** | -2.89 | (-4.13--1.39) | **5.37E-04** | **2.93E-03** | 1.30 | (-2.03- 4.75) | 4.46E-01 | 6.97E-01 | 4.10 | ( 0.00- 8.37) | **4.99E-02** | 1.71E-01 |
| 35 | DG(18:1_18:3) | -3.19 | (-4.08--2.15) | **1.73E-07** | **8.05E-07** | -1.52 | (-2.89- 0.12) | 6.76E-02 | 1.67E-01 | 1.95 | (-1.75- 5.79) | 3.04E-01 | 5.75E-01 | 5.41 | ( 1.17- 9.83) | **1.23E-02** | 7.46E-02 |
| 36 | DG(16:0_20:4) | -2.73 | (-3.63--1.69) | **3.52E-06** | **1.39E-05** | -0.95 | (-2.29- 0.63) | 2.23E-01 | 4.08E-01 | 4.44 | ( 1.01- 7.98) | **1.10E-02** | 6.76E-02 | 6.02 | ( 1.85-10.35) | **4.58E-03** | **4.18E-02** |
| 37 | DG(18:1_20:3) | -0.41 | (-1.44- 0.75) | 4.76E-01 | 6.03E-01 | 0.00 | (-1.21- 1.38) | 9.99E-01 | 9.99E-01 | -0.05 | (-3.17- 3.18) | 9.77E-01 | 9.94E-01 | 3.24 | (-0.57- 7.20) | 9.57E-02 | 2.50E-01 |
| 38 | DG(18:1_20:4) | -2.44 | (-3.23--1.56) | **9.71E-07** | **4.02E-06** | -1.43 | (-2.51--0.20) | **2.48E-02** | 7.31E-02 | 3.22 | ( 0.37- 6.14) | **2.67E-02** | 1.29E-01 | 5.18 | ( 1.62- 8.87) | **4.35E-03** | **4.17E-02** |
| 39 | DG(16:0_22:5) | -2.24 | (-3.18--1.17) | **1.36E-04** | **4.40E-04** | -1.33 | (-2.64- 0.22) | 8.92E-02 | 2.04E-01 | 2.97 | (-0.34- 6.39) | 7.91E-02 | 2.50E-01 | 7.25 | ( 3.11-11.56) | **5.90E-04** | **1.57E-02** |
| 40 | DG(18:2_20:4) | -3.51 | (-4.37--2.51) | **7.48E-09** | **4.23E-08** | -2.60 | (-3.82--1.13) | **1.21E-03** | **5.67E-03** | 3.79 | ( 0.56- 7.13) | **2.13E-02** | 1.10E-01 | 5.07 | ( 1.23- 9.06) | **9.64E-03** | 6.72E-02 |
| 41 | DG(16:0_22:6) | -4.54 | (-5.36--3.59) | **1.43E-12** | **1.43E-11** | -3.35 | (-4.46--2.00) | **1.79E-05** | **1.87E-04** | 5.24 | ( 0.67-10.01) | **2.42E-02** | 1.21E-01 | 5.67 | ( 0.73-10.85) | **2.43E-02** | 1.13E-01 |
| 42 | DG(18:1_22:5) | -1.93 | (-2.92--0.80) | **1.41E-03** | **3.82E-03** | -1.94 | (-3.11--0.58) | **6.78E-03** | **2.58E-02** | 0.81 | (-2.44- 4.17) | 6.28E-01 | 8.41E-01 | 2.39 | (-0.96- 5.86) | 1.63E-01 | 3.44E-01 |
| 43 | DG(18:1_22:6) | -3.76 | (-4.58--2.81) | **1.91E-10** | **1.37E-09** | -3.07 | (-4.12--1.81) | **1.92E-05** | **1.92E-04** | 3.21 | (-0.40- 6.95) | 8.20E-02 | 2.56E-01 | 5.85 | ( 1.28-10.63) | **1.20E-02** | 7.38E-02 |
| 44 | DG(18:2_22:6) | -4.58 | (-5.42--3.59) | **5.13E-12** | **4.56E-11** | -3.08 | (-4.42--1.43) | **7.99E-04** | **4.00E-03** | 3.42 | (-0.90- 7.93) | 1.21E-01 | 3.31E-01 | 7.28 | ( 1.88-12.98) | **8.03E-03** | 6.02E-02 |
| 45 | TG(48:0) [NL-18:0] | -2.22 | (-3.17--1.13) | **1.87E-04** | **5.94E-04** | -0.01 | (-1.56- 1.82) | 9.88E-01 | 9.94E-01 | 7.53 | ( 2.92-12.35) | **1.30E-03** | **2.49E-02** | 7.33 | ( 1.10-13.95) | **2.08E-02** | 1.05E-01 |

| | | | | | | | | | | | | | | | | | |
|---|---|---|---|---|---|---|---|---|---|---|---|---|---|---|---|---|---|
| 46 | TG(48:1) [NL-16:1] | -3.12 | (-4.11--1.98) | **2.70E-06** | **1.09E-05** | -1.26 | (-2.92- 0.79) | 2.11E-01 | 3.96E-01 | 9.50 | ( 4.81-14.39) | **6.51E-05** | **4.46E-03** | 9.88 | ( 3.77-16.34) | **1.40E-03** | **2.49E-02** |
| 47 | TG(48:1) [NL-18:1] | -3.72 | (-4.59--2.71) | **2.17E-09** | **1.37E-08** | -2.13 | (-3.61--0.31) | **2.43E-02** | 7.22E-02 | 7.66 | ( 2.32-13.28) | **4.71E-03** | **4.82E-02** | 9.11 | ( 2.17-16.51) | **9.66E-03** | 6.72E-02 |
| 48 | TG(48:2) [NL-14:0] | -4.21 | (-5.01--3.29) | **2.40E-12** | **2.31E-11** | -2.93 | (-4.22--1.36) | **7.95E-04** | **4.00E-03** | 7.99 | ( 2.52-13.76) | **3.97E-03** | **4.82E-02** | 10.05 | ( 3.00-17.58) | **4.88E-03** | **4.18E-02** |
| 49 | TG(48:2) [NL-14:1] | -3.30 | (-4.17--2.29) | **4.21E-08** | **2.06E-07** | -2.60 | (-3.88--1.04) | **2.05E-03** | **8.87E-03** | 9.57 | ( 4.45-14.93) | **2.21E-04** | **9.89E-03** | 7.84 | ( 1.05-15.09) | **2.33E-02** | 1.11E-01 |
| 50 | TG(48:2) [NL-16:1] | -3.18 | (-4.11--2.10) | **5.04E-07** | **2.18E-06** | -2.21 | (-3.60--0.51) | **1.32E-02** | **4.40E-02** | 10.34 | ( 5.02-15.92) | **1.21E-04** | **6.46E-03** | 11.55 | ( 4.79-18.75) | **7.22E-04** | **1.76E-02** |
| 51 | TG(48:2) [NL-18:2] | -4.78 | (-5.58--3.84) | **1.42E-13** | **1.66E-12** | -2.99 | (-4.34--1.31) | **1.26E-03** | **5.81E-03** | 7.95 | ( 2.40-13.79) | **4.73E-03** | **4.82E-02** | 9.77 | ( 2.92-17.08) | **4.87E-03** | **4.18E-02** |
| 52 | TG(48:3) [NL-14:0] | -4.43 | (-5.23--3.51) | **5.69E-13** | **6.35E-12** | -3.39 | (-4.62--1.87) | **1.03E-04** | **8.22E-04** | 8.83 | ( 3.18-14.78) | **2.02E-03** | **3.23E-02** | 11.17 | ( 4.21-18.60) | **1.49E-03** | **2.56E-02** |
| 53 | TG(48:3) [NL-16:1] | -3.11 | (-3.96--2.15) | **4.59E-08** | **2.22E-07** | -2.48 | (-3.66--1.08) | **1.12E-03** | **5.34E-03** | 8.05 | ( 3.43-12.88) | **5.94E-04** | **1.59E-02** | 9.57 | ( 3.77-15.69) | **1.13E-03** | **2.08E-02** |
| 54 | TG(48:3) [NL-18:3] | -3.79 | (-4.56--2.91) | **8.55E-12** | **7.33E-11** | -2.31 | (-3.52--0.87) | **2.79E-03** | **1.16E-02** | 8.11 | ( 2.36-14.17) | **5.40E-03** | **4.82E-02** | 8.74 | ( 1.44-16.57) | **1.85E-02** | 9.75E-02 |
| 55 | TG(49:1) [NL-17:1] | -2.57 | (-3.37--1.67) | **5.47E-07** | **2.34E-06** | -1.52 | (-2.61--0.27) | **1.89E-02** | 5.89E-02 | 4.87 | ( 1.86- 7.96) | **1.50E-03** | **2.77E-02** | 5.00 | ( 0.73- 9.45) | **2.16E-02** | 1.06E-01 |
| 56 | TG(50:1) [NL-14:0] | -2.17 | (-3.16--1.03) | **4.53E-04** | **1.32E-03** | -0.23 | (-1.87- 1.74) | 8.05E-01 | 9.14E-01 | 6.29 | ( 2.13-10.62) | **2.96E-03** | **3.98E-02** | 7.98 | ( 2.08-14.21) | **7.72E-03** | 5.98E-02 |
| 57 | TG(50:1) [NL-16:0] | -3.30 | (-4.13--2.35) | **6.72E-09** | **3.84E-08** | -1.09 | (-2.55- 0.65) | 2.04E-01 | 3.85E-01 | 4.46 | ( 1.31- 7.70) | **5.42E-03** | **4.82E-02** | 4.85 | ( 0.60- 9.29) | **2.53E-02** | 1.16E-01 |
| 58 | TG(50:1) [NL-18:1] | -3.08 | (-3.94--2.09) | **1.04E-07** | **4.95E-07** | -1.46 | (-2.85- 0.21) | 8.37E-02 | 1.93E-01 | 4.11 | ( 1.07- 7.24) | **7.95E-03** | 6.04E-02 | 4.47 | ( 0.34- 8.77) | **3.39E-02** | 1.35E-01 |
| 59 | TG(50:2) [NL-14:0] | -3.73 | (-4.62--2.69) | **5.05E-09** | **2.99E-08** | -2.59 | (-3.95--0.93) | **3.78E-03** | **1.54E-02** | 3.78 | (-0.34- 8.07) | 7.26E-02 | 2.45E-01 | 5.30 | ( 0.07-10.80) | **4.68E-02** | 1.67E-01 |
| 60 | TG(50:2) [NL-16:1] | -3.52 | (-4.43--2.47) | **3.08E-08** | **1.62E-07** | -2.09 | (-3.54--0.30) | **2.41E-02** | 7.22E-02 | 5.88 | ( 1.92-10.00) | **3.51E-03** | **4.51E-02** | 7.56 | ( 2.75-12.60) | **1.98E-03** | **2.63E-02** |
| 61 | TG(50:2) [NL-18:1] | -3.55 | (-4.47--2.48) | **3.91E-08** | **1.95E-07** | -2.31 | (-3.74--0.56) | **1.23E-02** | **4.20E-02** | 5.45 | ( 1.48- 9.58) | **7.00E-03** | 5.60E-02 | 7.61 | ( 2.81-12.63) | **1.82E-03** | **2.63E-02** |
| 62 | TG(50:2) [NL-18:2] | -5.03 | (-5.82--4.09) | **2.94E-14** | **3.92E-13** | -2.65 | (-4.14--0.79) | **7.57E-03** | **2.82E-02** | 4.85 | ( 1.65- 8.16) | **2.98E-03** | **3.98E-02** | 4.31 | ( 0.54- 8.22) | **2.48E-02** | 1.14E-01 |
| 63 | TG(50:3) [NL-14:0] | -4.63 | (-5.44--3.68) | **6.63E-13** | **7.24E-12** | -3.57 | (-4.82--2.03) | **6.73E-05** | **5.57E-04** | 4.38 | (-0.26- 9.22) | 6.42E-02 | 2.28E-01 | 7.21 | ( 1.46-13.28) | **1.37E-02** | 7.82E-02 |
| 64 | TG(50:3) [NL-14:1] | -2.50 | (-3.48--1.38) | **6.15E-05** | **2.09E-04** | -2.01 | (-3.30--0.46) | **1.32E-02** | **4.40E-02** | 4.51 | ( 0.99- 8.16) | **1.20E-02** | 7.12E-02 | 3.82 | (-0.97- 8.83) | 1.19E-01 | 2.83E-01 |
| 65 | TG(50:3) [NL-16:1] | -4.37 | (-5.18--3.44) | **1.66E-12** | **1.63E-11** | -3.01 | (-4.27--1.47) | **4.48E-04** | **2.57E-03** | 7.59 | ( 3.01-12.37) | **1.10E-03** | **2.30E-02** | 9.80 | ( 4.47-15.41) | **2.95E-04** | **1.09E-02** |
| 66 | TG(50:3) [NL-18:2] | -5.01 | (-5.76--4.13) | **1.19E-15** | **2.49E-14** | -3.22 | (-4.54--1.59) | **4.43E-04** | **2.57E-03** | 6.26 | ( 1.84-10.89) | **5.41E-03** | **4.82E-02** | 8.76 | ( 3.53-14.26) | **9.67E-04** | **1.93E-02** |
| 67 | TG(50:3) [NL-18:3] | -4.66 | (-5.43--3.77) | **2.57E-14** | **3.53E-13** | -2.28 | (-3.75--0.47) | **1.62E-02** | 5.20E-02 | 5.99 | ( 1.39-10.81) | **1.05E-02** | 6.73E-02 | 8.51 | ( 2.79-14.55) | **3.40E-03** | **3.55E-02** |
| 68 | TG(50:4) [NL-14:0] | -4.84 | (-5.57--3.97) | **1.09E-15** | **2.48E-14** | -3.48 | (-4.68--2.00) | **4.78E-05** | **4.17E-04** | 4.58 | (-0.16- 9.55) | 5.83E-02 | 2.17E-01 | 6.38 | ( 0.70-12.37) | **2.74E-02** | 1.19E-01 |
| 69 | TG(50:4) [NL-18:3] | -4.57 | (-5.27--3.76) | **1.60E-16** | **7.66E-15** | -2.79 | (-4.01--1.33) | **5.52E-04** | **2.97E-03** | 6.89 | ( 2.13-11.88) | **4.40E-03** | **4.82E-02** | 10.01 | ( 4.14-16.21) | **7.70E-04** | **1.76E-02** |
| 70 | TG(50:4) [NL-20:4] | -4.27 | (-5.17--3.19) | **7.40E-10** | **5.00E-09** | -2.20 | (-3.75--0.28) | **2.72E-02** | 7.78E-02 | 9.73 | ( 4.17-15.58) | **5.40E-04** | **1.59E-02** | 10.87 | ( 3.92-18.28) | **1.97E-03** | **2.63E-02** |
| 71 | TG(51:1) [NL-17:0] | -3.15 | (-4.00--2.18) | **3.94E-08** | **1.95E-07** | -2.01 | (-3.26--0.52) | **1.01E-02** | **3.51E-02** | 4.79 | ( 1.36- 8.34) | **6.09E-03** | 5.13E-02 | 4.99 | (-0.16-10.40) | 5.75E-02 | 1.83E-01 |
| 72 | TG(51:2) [NL-15:0] | -3.51 | (-4.41--2.46) | **3.27E-08** | **1.70E-07** | -2.96 | (-4.18--1.48) | **3.55E-04** | **2.21E-03** | 2.33 | (-1.42- 6.22) | 2.26E-01 | 4.75E-01 | 3.68 | (-0.75- 8.31) | 1.04E-01 | 2.65E-01 |
| 73 | TG(51:2) [NL-17:0] | -3.68 | (-4.53--2.70) | **1.23E-09** | **8.23E-09** | -2.87 | (-4.09--1.40) | **4.50E-04** | **2.57E-03** | 4.92 | ( 1.21- 8.76) | **9.19E-03** | 6.39E-02 | 6.43 | ( 1.49-11.61) | **1.05E-02** | 6.90E-02 |
| 74 | TG(51:2) [NL-17:1] | -3.12 | (-3.90--2.23) | **3.49E-09** | **2.15E-08** | -2.44 | (-3.49--1.22) | **2.81E-04** | **1.88E-03** | 4.39 | ( 1.04- 7.85) | **1.01E-02** | 6.71E-02 | 5.72 | ( 1.25-10.39) | **1.20E-02** | 7.38E-02 |
| 75 | TG(52:1) [NL-18:0] | -2.68 | (-3.68--1.52) | **3.70E-05** | **1.31E-04** | -0.47 | (-2.16- 1.59) | 6.29E-01 | 7.96E-01 | 5.25 | ( 1.63- 8.99) | **4.38E-03** | **4.82E-02** | 6.56 | ( 1.39-12.00) | **1.26E-02** | 7.47E-02 |
| 76 | TG(52:1) [NL-18:1] | -2.69 | (-3.69--1.55) | **3.02E-05** | **1.08E-04** | -0.30 | (-2.03- 1.79) | 7.57E-01 | 8.85E-01 | 4.81 | ( 1.41- 8.33) | **5.55E-03** | **4.85E-02** | 7.03 | ( 1.90-12.41) | **7.00E-03** | 5.51E-02 |
| 77 | TG(52:2) [NL-16:0] | -3.28 | (-4.05--2.41) | **5.03E-10** | **3.45E-09** | -1.92 | (-3.07--0.58) | **6.63E-03** | **2.55E-02** | 1.44 | (-0.82- 3.75) | 2.13E-01 | 4.62E-01 | 2.66 | (-0.05- 5.43) | 5.40E-02 | 1.76E-01 |
| 78 | TG(52:2) [NL-18:2] | -3.74 | (-4.64--2.69) | **7.94E-09** | **4.43E-08** | -1.05 | (-2.65- 0.89) | 2.67E-01 | 4.62E-01 | 3.99 | ( 0.99- 7.08) | **9.19E-03** | 6.39E-02 | 5.33 | ( 1.49- 9.32) | **6.52E-03** | 5.40E-02 |
| 79 | TG(52:3) [NL-16:1] | -3.72 | (-4.60--2.69) | **4.42E-09** | **2.65E-08** | -2.79 | (-4.02--1.31) | **6.50E-04** | **3.46E-03** | 3.16 | (-0.06- 6.48) | 5.47E-02 | 2.12E-01 | 3.77 | ( 0.03- 7.65) | **4.81E-02** | 1.69E-01 |
| 80 | TG(52:3) [NL-18:2] | -5.00 | (-5.73--4.14) | **2.68E-16** | **1.07E-14** | -3.38 | (-4.64--1.82) | **1.47E-04** | **1.14E-03** | 1.36 | (-1.28- 4.06) | 3.14E-01 | 5.90E-01 | 2.73 | (-0.16- 5.70) | 6.39E-02 | 1.90E-01 |
| 81 | TG(52:4) [NL-16:1] | -4.96 | (-5.71--4.08) | **1.14E-15** | **2.48E-14** | -3.77 | (-4.93--2.34) | **9.19E-06** | **1.05E-04** | 3.59 | (-0.53- 7.88) | 8.81E-02 | 2.66E-01 | 5.26 | ( 0.57-10.17) | **2.77E-02** | 1.19E-01 |
| 82 | TG(52:4) [NL-18:2] | -5.66 | (-6.38--4.80) | **9.57E-18** | **7.65E-16** | -3.76 | (-5.02--2.17) | **5.65E-05** | **4.76E-04** | 3.00 | (-0.87- 7.02) | 1.29E-01 | 3.47E-01 | 4.86 | ( 0.11- 9.85) | **4.50E-02** | 1.62E-01 |
| 83 | TG(52:4) [NL-18:3] | -4.75 | (-5.48--3.90) | **1.12E-15** | **2.48E-14** | -2.97 | (-4.26--1.38) | **7.42E-04** | **3.87E-03** | 3.67 | (-0.16- 7.65) | 6.01E-02 | 2.20E-01 | 5.99 | ( 1.18-11.03) | **1.44E-02** | 8.16E-02 |
| 84 | TG(52:5) [NL-18:3] | -4.81 | (-5.54--3.97) | **3.12E-16** | **1.07E-14** | -3.09 | (-4.29--1.63) | **1.81E-04** | **1.32E-03** | 3.83 | (-0.26- 8.10) | 6.66E-02 | 2.33E-01 | 7.81 | ( 2.65-13.23) | **2.90E-03** | **3.19E-02** |
| 85 | TG(52:5) [NL-20:4] | -4.28 | (-5.14--3.27) | **7.13E-11** | **5.43E-10** | -2.50 | (-3.94--0.70) | **8.74E-03** | **3.13E-02** | 9.64 | ( 4.84-14.66) | **7.49E-05** | **4.49E-03** | 11.13 | ( 5.45-17.11) | **1.09E-04** | **5.25E-03** |
| 86 | TG(52:5) [NL-20:5] | -5.06 | (-5.85--4.13) | **2.07E-14** | **3.01E-13** | -3.43 | (-4.69--1.88) | **1.15E-04** | **9.08E-04** | 7.27 | ( 2.99-11.74) | **8.36E-04** | **2.04E-02** | 9.30 | ( 4.19-14.65) | **3.35E-04** | **1.14E-02** |
| 87 | TG(53:2) [NL-18:1] | -2.88 | (-3.68--1.99) | **2.98E-08** | **1.60E-07** | -2.34 | (-3.40--1.11) | **4.81E-04** | **2.71E-03** | 1.38 | (-1.60- 4.45) | 3.67E-01 | 6.37E-01 | 2.62 | (-1.14- 6.53) | 1.73E-01 | 3.57E-01 |
| 88 | TG(54:1) [NL-18:1] | -1.99 | (-3.09--0.72) | **3.33E-03** | **8.56E-03** | -1.10 | (-2.67- 0.82) | 2.41E-01 | 4.30E-01 | 2.86 | (-0.08- 5.88) | 5.64E-02 | 2.17E-01 | 4.57 | ( 0.39- 8.92) | **3.20E-02** | 1.31E-01 |
| 89 | TG(54:2) [NL-18:0] | -2.33 | (-3.31--1.21) | **1.61E-04** | **5.18E-04** | -1.11 | (-2.57- 0.64) | 1.98E-01 | 3.76E-01 | 2.69 | (-0.17- 5.64) | 6.56E-02 | 2.32E-01 | 4.23 | ( 0.14- 8.48) | **4.26E-02** | 1.57E-01 |
| 90 | TG(54:2) [NL-20:1] | -2.03 | (-3.00--0.91) | **7.29E-04** | **2.06E-03** | -1.43 | (-2.77- 0.16) | 7.48E-02 | 1.78E-01 | 0.69 | (-2.44- 3.92) | 6.69E-01 | 8.54E-01 | 3.09 | (-0.72- 7.05) | 1.12E-01 | 2.75E-01 |
| 91 | TG(54:3) [NL-18:1] | -2.50 | (-3.37--1.52) | **5.92E-06** | **2.29E-05** | -1.73 | (-2.89--0.38) | **1.40E-02** | **4.62E-02** | -0.38 | (-2.60- 1.88) | 7.35E-01 | 8.86E-01 | 1.10 | (-1.75- 4.04) | 4.51E-01 | 6.33E-01 |
| 92 | TG(54:3) [NL-18:2] | -3.84 | (-4.60--2.97) | **3.74E-12** | **3.39E-11** | -1.87 | (-3.14--0.36) | **1.73E-02** | 5.50E-02 | 2.13 | (-0.23- 4.54) | 7.65E-02 | 2.45E-01 | 2.55 | (-0.44- 5.63) | 9.51E-02 | 2.50E-01 |
| 93 | TG(54:4) [NL-18:2] | -4.54 | (-5.34--3.59) | **7.27E-13** | **7.59E-12** | -3.20 | (-4.49--1.62) | **3.40E-04** | **2.19E-03** | 0.22 | (-2.47- 2.98) | 8.76E-01 | 9.47E-01 | 1.43 | (-2.12- 5.10) | 4.33E-01 | 6.15E-01 |
| 94 | TG(54:4) [NL-20:3] | -3.15 | (-4.06--2.09) | **3.70E-07** | **1.63E-06** | -1.95 | (-3.36--0.24) | **2.72E-02** | 7.78E-02 | 4.06 | ( 0.73- 7.50) | **1.67E-02** | 9.22E-02 | 6.70 | ( 2.74-10.82) | **9.11E-04** | **1.90E-02** |
| 95 | TG(54:5) [NL-18:3] | -4.38 | (-5.12--3.54) | **1.18E-14** | **1.85E-13** | -2.71 | (-3.93--1.24) | **8.01E-04** | **4.00E-03** | 1.63 | (-2.12- 5.52) | 3.98E-01 | 6.68E-01 | 5.33 | ( 0.39-10.50) | **3.41E-02** | 1.35E-01 |
| 96 | TG(54:5) [NL-20:4] | -4.47 | (-5.29--3.51) | **2.78E-12** | **2.56E-11** | -2.41 | (-3.89--0.57) | **1.28E-02** | **4.34E-02** | 5.48 | ( 2.17- 8.90) | **1.17E-03** | **2.34E-02** | 6.77 | ( 3.23-10.43) | **1.81E-04** | **7.88E-03** |
| 97 | TG(54:6) [NL-18:3] | -4.46 | (-5.21--3.59) | **2.51E-14** | **3.53E-13** | -2.83 | (-4.04--1.36) | **5.10E-04** | **2.82E-03** | 1.25 | (-2.82- 5.50) | 5.50E-01 | 7.82E-01 | 6.09 | ( 0.82-11.62) | **2.31E-02** | 1.11E-01 |
| 98 | TG(54:6) [NL-20:4] | -4.82 | (-5.61--3.88) | **9.29E-14** | **1.14E-12** | -2.87 | (-4.25--1.16) | **2.15E-03** | **9.22E-03** | 6.68 | ( 2.90-10.60) | **5.14E-04** | **1.59E-02** | 9.25 | ( 4.84-13.84) | **3.85E-05** | **3.08E-03** |

| | | | | | | | | | | | | | | | | | |
|---|---|---|---|---|---|---|---|---|---|---|---|---|---|---|---|---|---|
| 99 | TG(54:6) [NL-20:5] | -5.63 | (-6.35--4.75) | **2.45E-17** | **1.47E-15** | -3.99 | (-5.22--2.43) | **1.89E-05** | **1.92E-04** | 7.67 | ( 3.27-12.26) | **5.97E-04** | **1.59E-02** | 10.66 | ( 5.44-16.14) | **5.87E-05** | **4.03E-03** |
| 100 | TG(54:6) [NL-22:6] | -5.67 | (-6.44--4.73) | **1.49E-15** | **2.97E-14** | -4.31 | (-5.49--2.82) | **2.99E-06** | **4.09E-05** | 8.61 | ( 3.84-13.59) | **3.69E-04** | **1.47E-02** | 7.77 | ( 1.81-14.07) | **1.02E-02** | 6.90E-02 |
| 101 | TG(54:7) [NL-20:5] | -5.55 | (-6.29--4.66) | **9.60E-17** | **5.12E-15** | -4.07 | (-5.26--2.58) | **6.79E-06** | **8.14E-05** | 7.69 | ( 3.39-12.17) | **4.37E-04** | **1.50E-02** | 10.93 | ( 5.96-16.14) | **1.59E-05** | **1.53E-03** |
| 102 | TG(54:7) [NL-22:6] | -5.27 | (-6.06--4.34) | **7.77E-15** | **1.33E-13** | -4.34 | (-5.40--3.02) | **1.65E-07** | **3.61E-06** | 10.45 | ( 5.29-15.86) | **6.36E-05** | **4.46E-03** | 10.52 | ( 4.54-16.84) | **5.16E-04** | **1.46E-02** |
| 103 | TG(56:6) [NL-20:4] | -3.58 | (-4.43--2.61) | **1.98E-09** | **1.28E-08** | -2.08 | (-3.48--0.39) | **1.84E-02** | 5.77E-02 | 3.64 | ( 1.29- 6.06) | **2.50E-03** | **3.63E-02** | 4.30 | ( 1.39- 7.28) | **3.75E-03** | **3.83E-02** |
| 104 | TG(56:6) [NL-22:5] | -4.23 | (-5.02--3.33) | **1.07E-12** | **1.09E-11** | -3.05 | (-4.30--1.53) | **3.52E-04** | **2.21E-03** | 3.90 | ( 0.93- 6.97) | **1.01E-02** | 6.71E-02 | 4.13 | ( 1.07- 7.29) | **8.24E-03** | 6.02E-02 |
| 105 | TG(56:7) [NL-20:4] | -4.36 | (-5.20--3.38) | **1.38E-11** | **1.17E-10** | -2.47 | (-3.86--0.76) | **6.63E-03** | **2.55E-02** | 3.96 | ( 0.81- 7.21) | **1.35E-02** | 7.74E-02 | 4.93 | ( 1.03- 8.99) | **1.32E-02** | 7.63E-02 |
| 106 | TG(56:7) [NL-20:5] | -4.42 | (-5.25--3.44) | **7.75E-12** | **6.76E-11** | -2.75 | (-4.04--1.18) | **1.37E-03** | **6.19E-03** | 5.08 | ( 1.17- 9.15) | **1.08E-02** | 6.76E-02 | 7.20 | ( 2.68-11.92) | **1.74E-03** | **2.63E-02** |
| 107 | TG(56:7) [NL-22:5] | -4.79 | (-5.52--3.93) | **8.16E-16** | **2.06E-14** | -3.45 | (-4.60--2.05) | **2.32E-05** | **2.27E-04** | 4.85 | ( 1.24- 8.58) | **8.32E-03** | 6.05E-02 | 5.93 | ( 1.83-10.19) | **4.51E-03** | **4.18E-02** |
| 108 | TG(56:7) [NL-22:6] | -5.45 | (-6.23--4.55) | **3.63E-15** | **6.46E-14** | -4.45 | (-5.55--3.08) | **2.88E-07** | **5.53E-06** | 6.41 | ( 1.90-11.11) | **5.16E-03** | **4.82E-02** | 6.57 | ( 1.26-12.16) | **1.50E-02** | 8.40E-02 |
| 109 | TG(56:8) [NL-20:4] | -3.75 | (-4.62--2.74) | **2.02E-09** | **1.29E-08** | -1.57 | (-3.00- 0.14) | 7.05E-02 | 1.72E-01 | 4.51 | ( 1.22- 7.90) | **7.15E-03** | 5.62E-02 | 6.47 | ( 2.25-10.87) | **2.61E-03** | **3.06E-02** |
| 110 | TG(56:8) [NL-20:5] | -4.81 | (-5.55--3.94) | **2.16E-15** | **4.14E-14** | -3.07 | (-4.30--1.59) | **2.42E-04** | **1.69E-03** | 4.10 | ( 0.03- 8.35) | **4.84E-02** | 1.98E-01 | 7.48 | ( 2.98-12.19) | **1.11E-03** | **2.08E-02** |
| 111 | TG(56:8) [NL-22:6] | -5.84 | (-6.57--4.96) | **2.22E-17** | **1.47E-15** | -4.61 | (-5.66--3.32) | **3.86E-08** | **1.42E-06** | 7.11 | ( 2.31-12.15) | **3.57E-03** | **4.51E-02** | 8.30 | ( 3.04-13.83) | **1.89E-03** | **2.63E-02** |
| 112 | TG(56:9) [NL-22:6] | -5.30 | (-6.04--4.42) | **2.03E-16** | **8.86E-15** | -4.24 | (-5.30--2.94) | **1.95E-07** | **3.90E-06** | 5.85 | ( 1.37-10.53) | **1.02E-02** | 6.71E-02 | 8.01 | ( 2.89-13.38) | **2.08E-03** | **2.63E-02** |
| 113 | TG(58:10) [NL-22:6] | -5.02 | (-5.75--4.16) | **2.94E-16** | **1.07E-14** | -3.63 | (-4.76--2.26) | **7.78E-06** | **9.11E-05** | 5.10 | ( 0.69- 9.69) | **2.31E-02** | 1.17E-01 | 5.55 | ( 0.57-10.77) | **2.86E-02** | 1.21E-01 |
| 114 | TG(58:8) [NL-22:6] | -4.95 | (-5.76--4.00) | **1.12E-13** | **1.34E-12** | -3.76 | (-4.86--2.43) | **2.50E-06** | **3.64E-05** | 4.39 | (-0.08- 9.06) | 5.42E-02 | 2.11E-01 | 5.12 | ( 0.12-10.38) | **4.49E-02** | 1.62E-01 |
| 115 | TG(58:9) [NL-22:6] | -5.22 | (-5.96--4.33) | **4.35E-16** | **1.33E-14** | -4.05 | (-5.11--2.76) | **3.75E-07** | **6.67E-06** | 3.42 | (-0.99- 8.03) | 1.29E-01 | 3.47E-01 | 6.25 | ( 1.09-11.68) | **1.73E-02** | 9.33E-02 |
| 116 | TG(O-50:1) [NL-16:0] | -0.91 | (-1.82- 0.11) | 7.75E-02 | 1.42E-01 | -0.15 | (-1.24- 1.08) | 8.04E-01 | 9.14E-01 | 1.93 | (-0.91- 4.85) | 1.83E-01 | 4.24E-01 | 2.23 | (-1.09- 5.66) | 1.89E-01 | 3.72E-01 |
| 117 | TG(O-50:1) [NL-17:1] | -2.62 | (-3.44--1.70) | **5.75E-07** | **2.44E-06** | -1.48 | (-2.61--0.18) | **2.71E-02** | 7.78E-02 | 4.94 | ( 1.86- 8.11) | **1.64E-03** | **2.81E-02** | 5.04 | ( 0.70- 9.57) | **2.28E-02** | 1.11E-01 |
| 118 | TG(O-50:1) [NL-18:1] | -1.31 | (-2.20--0.31) | **1.16E-02** | **2.57E-02** | -0.47 | (-1.58- 0.78) | 4.41E-01 | 6.29E-01 | 3.27 | ( 0.25- 6.37) | **3.36E-02** | 1.52E-01 | 2.73 | (-1.04- 6.65) | 1.57E-01 | 3.40E-01 |
| 119 | TG(O-50:2) [NL-18:2] | -0.54 | (-1.53- 0.56) | 3.20E-01 | 4.52E-01 | -0.24 | (-1.39- 1.07) | 7.04E-01 | 8.47E-01 | 3.60 | ( 0.90- 6.36) | **8.82E-03** | 6.32E-02 | 2.19 | (-1.06- 5.55) | 1.86E-01 | 3.70E-01 |
| 120 | TG(O-52:2) [NL-16:0] | -1.09 | (-2.30- 0.31) | 1.20E-01 | 2.02E-01 | -1.08 | (-2.40- 0.48) | 1.64E-01 | 3.31E-01 | 4.14 | ( 0.42- 8.00) | **2.91E-02** | 1.40E-01 | 1.81 | (-2.69- 6.51) | 4.35E-01 | 6.16E-01 |
| 121 | TG(O-52:2) [NL-17:1] | -3.15 | (-3.95--2.26) | **3.90E-09** | **2.37E-08** | -2.42 | (-3.49--1.17) | **4.01E-04** | **2.41E-03** | 4.41 | ( 1.03- 7.90) | **1.05E-02** | 6.73E-02 | 5.69 | ( 1.18-10.39) | **1.32E-02** | 7.63E-02 |
| 122 | TG(O-52:2) [NL-18:1] | -1.39 | (-2.63- 0.06) | 5.90E-02 | 1.12E-01 | -2.16 | (-3.40--0.70) | **5.47E-03** | **2.17E-02** | 5.31 | ( 1.22- 9.58) | **1.09E-02** | 6.76E-02 | 4.33 | (-0.27- 9.15) | 6.54E-02 | 1.93E-01 |
| 123 | TG(O-54:2) [NL-18:1] | -2.88 | (-3.68--1.99) | **3.31E-08** | **1.71E-07** | -2.28 | (-3.35--1.03) | **7.73E-04** | **3.99E-03** | 1.34 | (-1.63- 4.40) | 3.79E-01 | 6.53E-01 | 2.58 | (-1.17- 6.48) | 1.79E-01 | 3.65E-01 |
| 124 | TG(O-54:4) [NL-17:1] | -4.46 | (-5.28--3.50) | **2.66E-12** | **2.51E-11** | -3.21 | (-4.43--1.71) | **1.55E-04** | **1.18E-03** | 2.45 | (-1.04- 6.06) | 1.69E-01 | 4.07E-01 | 3.54 | (-0.34- 7.58) | 7.38E-02 | 2.11E-01 |
| 125 | TG(O-54:4) [NL-18:2] | -5.23 | (-5.97--4.34) | **4.44E-16** | **1.33E-14** | -3.45 | (-4.69--1.92) | **9.27E-05** | **7.54E-04** | 2.63 | (-1.04- 6.44) | 1.61E-01 | 3.95E-01 | 2.86 | (-1.47- 7.39) | 1.97E-01 | 3.82E-01 |
| 126 | dhCer(d18:0/22:0) | 0.02 | (-0.87- 1.00) | 9.65E-01 | 9.88E-01 | -0.15 | (-1.17- 0.99) | 7.89E-01 | 9.01E-01 | 1.33 | (-1.12- 3.85) | 2.87E-01 | 5.55E-01 | 1.21 | (-1.38- 3.86) | 3.60E-01 | 5.57E-01 |
| 127 | dhCer(d18:0/24:0) | -0.64 | (-1.46- 0.27) | 1.63E-01 | 2.64E-01 | -0.26 | (-1.36- 0.98) | 6.69E-01 | 8.33E-01 | 0.07 | (-2.14- 2.33) | 9.50E-01 | 9.83E-01 | -0.13 | (-2.50- 2.29) | 9.13E-01 | 9.74E-01 |
| 128 | dhCer(d18:0/24:1) | 0.39 | (-0.80- 1.73) | 5.33E-01 | 6.57E-01 | -0.93 | (-2.05- 0.35) | 1.48E-01 | 3.06E-01 | 1.80 | (-0.84- 4.51) | 1.82E-01 | 4.24E-01 | 1.38 | (-1.82- 4.67) | 4.00E-01 | 5.90E-01 |
| 129 | Cer(d16:1/22:0) | -1.09 | (-1.99--0.09) | **3.43E-02** | 6.86E-02 | -0.93 | (-2.09- 0.40) | 1.61E-01 | 3.27E-01 | 1.94 | (-0.68- 4.63) | 1.46E-01 | 3.64E-01 | 3.14 | ( 0.04- 6.34) | **4.68E-02** | 1.67E-01 |
| 130 | Cer(d16:1/23:0) | -0.97 | (-2.54- 0.93) | 2.96E-01 | 4.29E-01 | -1.00 | (-2.68- 1.06) | 3.14E-01 | 5.08E-01 | 1.84 | (-1.15- 4.91) | 2.29E-01 | 4.78E-01 | 3.13 | (-0.35- 6.73) | 7.82E-02 | 2.18E-01 |
| 131 | Cer(d16:1/24:0) | -0.99 | (-1.94- 0.07) | 6.68E-02 | 1.23E-01 | -0.55 | (-1.64- 0.67) | 3.60E-01 | 5.61E-01 | 1.20 | (-1.61- 4.08) | 4.04E-01 | 6.75E-01 | 3.27 | ( 0.08- 6.55) | **4.43E-02** | 1.62E-01 |
| 132 | Cer(d16:1/24:1) | -0.69 | (-1.81- 0.58) | 2.72E-01 | 4.00E-01 | -0.12 | (-1.50- 1.48) | 8.74E-01 | 9.49E-01 | 1.16 | (-1.12- 3.50) | 3.18E-01 | 5.94E-01 | 3.31 | ( 0.23- 6.48) | **3.49E-02** | 1.37E-01 |
| 133 | Cer(d17:1/22:0) | -0.81 | (-2.00- 0.56) | 2.33E-01 | 3.51E-01 | -0.84 | (-2.19- 0.74) | 2.77E-01 | 4.73E-01 | 0.81 | (-1.73- 3.42) | 5.32E-01 | 7.65E-01 | 1.34 | (-1.57- 4.33) | 3.68E-01 | 5.64E-01 |
| 134 | Cer(d17:1/23:0) | -1.93 | (-3.39--0.16) | **3.39E-02** | 6.81E-02 | -0.26 | (-2.64- 2.89) | 8.52E-01 | 9.38E-01 | 1.79 | (-1.17- 4.84) | 2.37E-01 | 4.91E-01 | 2.09 | (-1.37- 5.66) | 2.37E-01 | 4.31E-01 |
| 135 | Cer(d17:1/24:0) | -1.25 | (-2.17--0.23) | **1.80E-02** | **3.85E-02** | -0.76 | (-1.90- 0.54) | 2.40E-01 | 4.30E-01 | 0.35 | (-2.27- 3.03) | 7.96E-01 | 9.16E-01 | -0.11 | (-2.85- 2.70) | 9.35E-01 | 9.75E-01 |
| 136 | Cer(d17:1/24:1) | -1.39 | (-2.39--0.25) | **1.80E-02** | **3.85E-02** | -0.09 | (-1.43- 1.46) | 9.03E-01 | 9.63E-01 | 1.34 | (-1.18- 3.92) | 3.00E-01 | 5.71E-01 | 0.84 | (-2.17- 3.95) | 5.84E-01 | 7.50E-01 |
| 137 | Cer(d18:1/16:0) | -0.44 | (-1.18- 0.36) | 2.71E-01 | 3.99E-01 | -0.54 | (-1.45- 0.47) | 2.81E-01 | 4.75E-01 | -0.57 | (-2.30- 1.18) | 5.17E-01 | 7.51E-01 | -0.07 | (-2.07- 1.97) | 9.45E-01 | 9.75E-01 |
| 138 | Cer(d18:1/18:0) | -0.77 | (-1.59- 0.14) | 9.51E-02 | 1.69E-01 | -0.64 | (-1.70- 0.54) | 2.73E-01 | 4.70E-01 | 1.53 | (-0.75- 3.85) | 1.88E-01 | 4.29E-01 | 2.70 | (-0.06- 5.53) | 5.50E-02 | 1.77E-01 |
| 139 | Cer(d18:1/20:0) | -0.75 | (-1.57- 0.14) | 9.73E-02 | 1.72E-01 | -0.73 | (-1.72- 0.37) | 1.83E-01 | 3.55E-01 | 1.44 | (-0.66- 3.58) | 1.79E-01 | 4.20E-01 | 0.36 | (-2.00- 2.78) | 7.67E-01 | 8.89E-01 |
| 140 | Cer(d18:1/22:0) | -0.33 | (-1.07- 0.49) | 4.19E-01 | 5.53E-01 | 0.31 | (-0.67- 1.40) | 5.47E-01 | 7.34E-01 | 1.19 | (-0.79- 3.21) | 2.38E-01 | 4.91E-01 | 0.69 | (-1.37- 2.80) | 5.11E-01 | 6.95E-01 |
| 141 | Cer(d18:1/23:0) | -0.69 | (-1.59- 0.30) | 1.63E-01 | 2.64E-01 | 0.13 | (-1.16- 1.61) | 8.51E-01 | 9.38E-01 | 0.86 | (-1.19- 2.95) | 4.11E-01 | 6.75E-01 | 1.89 | (-0.27- 4.09) | 8.62E-02 | 2.35E-01 |
| 142 | Cer(d18:1/24:0) | -0.72 | (-1.46- 0.09) | 7.91E-02 | 1.43E-01 | 0.16 | (-0.84- 1.26) | 7.67E-01 | 8.85E-01 | 0.25 | (-1.65- 2.18) | 7.97E-01 | 9.16E-01 | 0.12 | (-1.72- 2.00) | 8.98E-01 | 9.70E-01 |
| 143 | Cer(d18:1/24:1) | -0.42 | (-1.22- 0.46) | 3.41E-01 | 4.71E-01 | -0.10 | (-1.17- 1.09) | 8.57E-01 | 9.41E-01 | 1.32 | (-0.57- 3.25) | 1.72E-01 | 4.11E-01 | -0.14 | (-2.18- 1.95) | 8.97E-01 | 9.70E-01 |
| 144 | Cer(d18:2/16:0) | -0.65 | (-1.77- 0.63) | 3.06E-01 | 4.40E-01 | 1.39 | (-0.18- 3.22) | 8.54E-02 | 1.96E-01 | 0.26 | (-2.08- 2.66) | 8.28E-01 | 9.33E-01 | 1.02 | (-1.93- 4.07) | 4.98E-01 | 6.83E-01 |
| 145 | Cer(d18:2/22:0) | -0.81 | (-1.69- 0.17) | 1.01E-01 | 1.75E-01 | 0.56 | (-0.60- 1.86) | 3.58E-01 | 5.60E-01 | 0.29 | (-1.78- 2.39) | 7.86E-01 | 9.16E-01 | 0.85 | (-1.81- 3.58) | 5.33E-01 | 7.11E-01 |
| 146 | Cer(d18:2/23:0) | -0.88 | (-2.01- 0.41) | 1.70E-01 | 2.72E-01 | 0.56 | (-1.06- 2.47) | 5.20E-01 | 7.08E-01 | 0.99 | (-1.30- 3.32) | 3.98E-01 | 6.68E-01 | 1.80 | (-0.89- 4.57) | 1.90E-01 | 3.72E-01 |
| 147 | Cer(d18:2/24:0) | -0.98 | (-1.86--0.00) | **4.96E-02** | 9.48E-02 | -0.45 | (-1.55- 0.79) | 4.58E-01 | 6.47E-01 | -0.88 | (-2.89- 1.17) | 3.97E-01 | 6.68E-01 | 0.09 | (-2.34- 2.58) | 9.40E-01 | 9.75E-01 |
| 148 | Cer(d18:2/24:1) | 0.03 | (-0.91- 1.07) | 9.55E-01 | 9.79E-01 | 0.03 | (-1.14- 1.35) | 9.64E-01 | 9.86E-01 | 0.50 | (-1.53- 2.57) | 6.31E-01 | 8.42E-01 | 0.92 | (-1.53- 3.43) | 4.63E-01 | 6.46E-01 |
| 149 | Cer(d19:1/24:0) | -0.93 | (-2.01- 0.30) | 1.33E-01 | 2.22E-01 | -1.46 | (-2.65--0.06) | **4.13E-02** | 1.14E-01 | 0.49 | (-3.24- 4.36) | 7.99E-01 | 9.16E-01 | -0.26 | (-4.45- 4.12) | 9.05E-01 | 9.71E-01 |
| 150 | Cer(d19:1/24:1) | -0.55 | (-1.88- 1.01) | 4.67E-01 | 5.93E-01 | -2.07 | (-3.42--0.45) | **1.49E-02** | **4.87E-02** | 1.52 | (-2.52- 5.72) | 4.65E-01 | 7.18E-01 | 0.78 | (-4.13- 5.94) | 7.59E-01 | 8.85E-01 |
| 151 | Cer(m18:0/22:0) | -0.67 | (-1.89- 0.74) | 3.33E-01 | 4.65E-01 | -1.77 | (-3.11--0.16) | **3.30E-02** | 9.22E-02 | 1.99 | (-0.60- 4.65) | 1.32E-01 | 3.48E-01 | 3.74 | ( 0.81- 6.75) | **1.24E-02** | 7.46E-02 |

| | | | | | | | | | | | | | | | | | |
|---|---|---|---|---|---|---|---|---|---|---|---|---|---|---|---|---|---|
| 152 | Cer(m18:0/23:0) | -3.06 | (-4.80--0.73) | **1.34E-02** | **2.97E-02** | 1.54 | (-1.71- 6.07) | 3.94E-01 | 5.91E-01 | 2.29 | (-0.70- 5.38) | 1.34E-01 | 3.48E-01 | 3.57 | (-0.39- 7.68) | 7.77E-02 | 2.18E-01 |
| 153 | Cer(m18:0/24:0) | -1.48 | (-2.39--0.46) | **5.63E-03** | **1.38E-02** | -0.56 | (-1.79- 0.87) | 4.22E-01 | 6.18E-01 | 2.03 | (-0.41- 4.53) | 1.03E-01 | 2.93E-01 | 2.54 | (-0.33- 5.50) | 8.29E-02 | 2.28E-01 |
| 154 | Cer(m18:0/24:1) | -1.01 | (-2.26- 0.45) | 1.64E-01 | 2.64E-01 | -2.11 | (-3.51--0.40) | **1.82E-02** | 5.75E-02 | 3.88 | ( 0.85- 7.01) | **1.20E-02** | 7.12E-02 | 5.40 | ( 2.05- 8.86) | **1.59E-03** | **2.62E-02** |
| 155 | Cer(m18:1/22:0) | -2.74 | (-3.55--1.84) | **1.44E-07** | **6.76E-07** | -2.05 | (-3.15--0.77) | **2.79E-03** | **1.16E-02** | 0.15 | (-2.59- 2.96) | 9.16E-01 | 9.69E-01 | 3.27 | (-0.61- 7.30) | 9.93E-02 | 2.58E-01 |
| 156 | Cer(m18:1/23:0) | -3.02 | (-4.50--1.15) | **3.15E-03** | **8.17E-03** | -3.90 | (-5.70--1.34) | **5.88E-03** | **2.30E-02** | 2.36 | (-0.98- 5.81) | 1.67E-01 | 4.06E-01 | 1.68 | (-2.29- 5.82) | 4.09E-01 | 5.97E-01 |
| 157 | Cer(m18:1/24:0) | -2.64 | (-3.45--1.74) | **3.32E-07** | **1.48E-06** | -2.36 | (-3.41--1.14) | **4.08E-04** | **2.42E-03** | -0.49 | (-3.07- 2.17) | 7.14E-01 | 8.85E-01 | 1.01 | (-2.29- 4.43) | 5.50E-01 | 7.24E-01 |
| 158 | Cer(m18:1/24:1) | -1.76 | (-2.83--0.52) | **6.74E-03** | **1.62E-02** | -2.53 | (-3.57--1.31) | **1.76E-04** | **1.31E-03** | 0.60 | (-2.10- 3.38) | 6.63E-01 | 8.54E-01 | 3.93 | ( 0.25- 7.74) | **3.63E-02** | 1.40E-01 |
| 159 | GM3(d18:1/18:0) | -0.87 | (-1.62--0.06) | **3.50E-02** | 6.96E-02 | -0.55 | (-1.50- 0.50) | 2.92E-01 | 4.89E-01 | -0.52 | (-2.29- 1.28) | 5.66E-01 | 7.93E-01 | -2.21 | (-4.46- 0.10) | 6.06E-02 | 1.88E-01 |
| 160 | GM3(d18:1/22:0) | -0.04 | (-1.21- 1.29) | 9.51E-01 | 9.77E-01 | 0.35 | (-1.27- 2.26) | 6.93E-01 | 8.47E-01 | -0.44 | (-2.44- 1.60) | 6.69E-01 | 8.54E-01 | -0.29 | (-2.92- 2.41) | 8.30E-01 | 9.23E-01 |
| 161 | GM3(d18:1/24:0) | -0.66 | (-1.64- 0.44) | 2.28E-01 | 3.47E-01 | -0.16 | (-1.35- 1.20) | 8.09E-01 | 9.16E-01 | -2.09 | (-3.98--0.17) | **3.36E-02** | 1.52E-01 | -2.80 | (-5.04--0.50) | **1.76E-02** | 9.38E-02 |
| 162 | GM3(d18:1/24:1) | -0.02 | (-1.04- 1.13) | 9.75E-01 | 9.89E-01 | -0.14 | (-1.40- 1.30) | 8.41E-01 | 9.33E-01 | -1.02 | (-2.87- 0.87) | 2.87E-01 | 5.55E-01 | -2.51 | (-4.97- 0.01) | 5.06E-02 | 1.71E-01 |
| 163 | HexCer(d16:1/22:0) | -0.85 | (-2.39- 0.99) | 3.41E-01 | 4.71E-01 | -0.92 | (-2.88- 1.59) | 4.37E-01 | 6.27E-01 | -0.57 | (-3.29- 2.23) | 6.85E-01 | 8.62E-01 | -0.95 | (-3.91- 2.11) | 5.37E-01 | 7.12E-01 |
| 164 | HexCer(d16:1/24:0) | -0.91 | (-2.25- 0.66) | 2.40E-01 | 3.60E-01 | -0.76 | (-2.59- 1.53) | 4.84E-01 | 6.67E-01 | 0.97 | (-1.90- 3.91) | 5.11E-01 | 7.47E-01 | -0.57 | (-3.45- 2.41) | 7.04E-01 | 8.48E-01 |
| 165 | HexCer(d18:1/16:0) | 0.24 | (-0.82- 1.43) | 6.66E-01 | 7.48E-01 | -0.25 | (-1.48- 1.15) | 7.06E-01 | 8.47E-01 | -0.79 | (-2.70- 1.17) | 4.26E-01 | 6.81E-01 | -2.68 | (-4.63--0.68) | **9.26E-03** | 6.64E-02 |
| 166 | HexCer(d18:1/18:0) | -0.03 | (-1.30- 1.43) | 9.71E-01 | 9.89E-01 | 0.61 | (-1.01- 2.53) | 4.82E-01 | 6.66E-01 | -1.68 | (-3.97- 0.67) | 1.58E-01 | 3.89E-01 | -1.86 | (-4.53- 0.89) | 1.83E-01 | 3.70E-01 |
| 167 | HexCer(d18:1/20:0) | -0.11 | (-1.28- 1.23) | 8.67E-01 | 9.19E-01 | 0.21 | (-1.24- 1.90) | 7.93E-01 | 9.04E-01 | -1.76 | (-4.03- 0.56) | 1.34E-01 | 3.48E-01 | -3.20 | (-5.58--0.76) | **1.08E-02** | 6.90E-02 |
| 168 | HexCer(d18:1/22:0) | 0.30 | (-0.68- 1.38) | 5.64E-01 | 6.80E-01 | 0.84 | (-0.34- 2.16) | 1.71E-01 | 3.39E-01 | -1.61 | (-3.63- 0.44) | 1.23E-01 | 3.33E-01 | -3.03 | (-5.05--0.96) | **4.64E-03** | **4.18E-02** |
| 169 | HexCer(d18:1/24:0) | 0.22 | (-0.78- 1.32) | 6.80E-01 | 7.61E-01 | -0.07 | (-1.18- 1.18) | 9.11E-01 | 9.69E-01 | -0.95 | (-2.87- 1.00) | 3.34E-01 | 6.14E-01 | -2.91 | (-5.07--0.70) | **1.06E-02** | 6.90E-02 |
| 170 | HexCer(d18:1/24:1) | 0.81 | (-0.21- 1.94) | 1.22E-01 | 2.06E-01 | 0.21 | (-0.92- 1.48) | 7.26E-01 | 8.57E-01 | -0.23 | (-2.22- 1.80) | 8.23E-01 | 9.32E-01 | -2.92 | (-5.12--0.66) | **1.18E-02** | 7.38E-02 |
| 171 | Hex2Cer(d16:1/16:0) | -1.29 | (-2.52- 0.13) | 7.36E-02 | 1.35E-01 | -0.64 | (-2.19- 1.22) | 4.72E-01 | 6.59E-01 | 2.53 | (-0.25- 5.39) | 7.44E-02 | 2.45E-01 | 2.38 | (-1.06- 5.93) | 1.75E-01 | 3.60E-01 |
| 172 | Hex2Cer(d18:1/16:0) | -0.46 | (-1.32- 0.49) | 3.33E-01 | 4.65E-01 | -0.50 | (-1.58- 0.71) | 3.99E-01 | 5.96E-01 | -0.92 | (-3.16- 1.37) | 4.25E-01 | 6.81E-01 | -1.58 | (-4.25- 1.16) | 2.54E-01 | 4.44E-01 |
| 173 | Hex2Cer(d18:1/22:0) | -0.10 | (-1.22- 1.18) | 8.76E-01 | 9.20E-01 | -0.27 | (-1.55- 1.20) | 7.03E-01 | 8.47E-01 | -2.30 | (-4.36--0.19) | **3.26E-02** | 1.52E-01 | -2.58 | (-5.23- 0.15) | 6.38E-02 | 1.90E-01 |
| 174 | Hex2Cer(d18:1/24:1) | -0.09 | (-1.11- 1.05) | 8.75E-01 | 9.20E-01 | -0.15 | (-1.36- 1.22) | 8.19E-01 | 9.20E-01 | -2.61 | (-4.86--0.32) | **2.59E-02** | 1.27E-01 | -1.91 | (-4.63- 0.88) | 1.76E-01 | 3.60E-01 |
| 175 | Hex2Cer(d18:2/16:0) | -0.69 | (-1.70- 0.44) | 2.18E-01 | 3.34E-01 | 0.03 | (-1.19- 1.43) | 9.59E-01 | 9.85E-01 | 1.38 | (-0.81- 3.62) | 2.18E-01 | 4.66E-01 | 0.15 | (-2.47- 2.85) | 9.10E-01 | 9.73E-01 |
| 176 | Hex3Cer(d18:1/16:0) | -0.37 | (-1.11- 0.44) | 3.60E-01 | 4.87E-01 | -0.54 | (-1.47- 0.49) | 2.94E-01 | 4.89E-01 | -0.68 | (-2.09- 0.76) | 3.51E-01 | 6.24E-01 | -0.66 | (-2.52- 1.24) | 4.90E-01 | 6.78E-01 |
| 177 | Hex3Cer(d18:1/22:0) | 0.32 | (-0.86- 1.65) | 6.14E-01 | 7.17E-01 | 0.05 | (-1.47- 1.84) | 9.52E-01 | 9.82E-01 | -1.58 | (-3.65- 0.54) | 1.43E-01 | 3.60E-01 | -1.47 | (-3.84- 0.95) | 2.30E-01 | 4.27E-01 |
| 178 | Hex3Cer(d18:1/24:0) | -0.65 | (-1.60- 0.41) | 2.20E-01 | 3.36E-01 | 0.07 | (-1.17- 1.48) | 9.21E-01 | 9.70E-01 | -2.36 | (-4.26--0.41) | **1.79E-02** | 9.75E-02 | -1.74 | (-3.95- 0.52) | 1.29E-01 | 3.01E-01 |
| 179 | Hex3Cer(d18:1/24:1) | -0.48 | (-1.41- 0.55) | 3.47E-01 | 4.75E-01 | -0.50 | (-1.68- 0.86) | 4.50E-01 | 6.38E-01 | -0.34 | (-2.19- 1.54) | 7.17E-01 | 8.85E-01 | -1.89 | (-4.13- 0.39) | 1.03E-01 | 2.64E-01 |
| 180 | SM(d17:1/14:0) | -0.65 | (-1.33- 0.08) | 7.79E-02 | 1.42E-01 | -1.33 | (-2.10--0.48) | **2.83E-03** | **1.17E-02** | 1.99 | (-0.50- 4.54) | 1.17E-01 | 3.25E-01 | 3.07 | (-0.05- 6.29) | 5.35E-02 | 1.76E-01 |
| 181 | SM(d18:0/14:0) | 0.28 | (-0.43- 1.04) | 4.48E-01 | 5.75E-01 | -0.91 | (-1.64--0.11) | **2.70E-02** | 7.78E-02 | 1.29 | (-0.76- 3.38) | 2.18E-01 | 4.66E-01 | 2.32 | (-0.21- 4.91) | 7.26E-02 | 2.10E-01 |
| 182 | SM(d18:1/14:0)/SM(d16:1/16:0) | -0.27 | (-0.92- 0.43) | 4.36E-01 | 5.66E-01 | -0.84 | (-1.61- 0.00) | 5.12E-02 | 1.36E-01 | 1.74 | (-0.25- 3.78) | 8.64E-02 | 2.66E-01 | 2.96 | ( 0.46- 5.53) | **2.05E-02** | 1.05E-01 |
| 183 | SM(d18:2/14:0) | 0.33 | (-0.35- 1.06) | 3.50E-01 | 4.79E-01 | -0.17 | (-1.01- 0.75) | 7.02E-01 | 8.47E-01 | 5.30 | ( 3.41- 7.21) | **6.74E-08** | **3.24E-05** | 8.07 | ( 5.72-10.47) | **1.01E-10** | **4.83E-08** |
| 184 | SM(d17:1/16:0) | -0.92 | (-1.54--0.24) | **8.73E-03** | **2.03E-02** | -1.01 | (-1.78--0.17) | **2.03E-02** | 6.25E-02 | -0.43 | (-2.40- 1.58) | 6.71E-01 | 8.54E-01 | -0.16 | (-2.61- 2.36) | 9.01E-01 | 9.70E-01 |
| 185 | SM(d18:1/16:0) | -0.23 | (-0.98- 0.58) | 5.66E-01 | 6.80E-01 | -0.43 | (-1.33- 0.56) | 3.78E-01 | 5.75E-01 | -1.51 | (-3.36- 0.38) | 1.15E-01 | 3.23E-01 | -1.47 | (-3.73- 0.85) | 2.12E-01 | 4.01E-01 |
| 186 | SM(d18:2/16:0) | 0.19 | (-0.46- 0.89) | 5.74E-01 | 6.85E-01 | 0.00 | (-0.81- 0.87) | 9.97E-01 | 9.99E-01 | 0.75 | (-0.62- 2.14) | 2.84E-01 | 5.53E-01 | 2.14 | ( 0.41- 3.90) | **1.57E-02** | 8.55E-02 |
| 187 | SM(34:3) | 0.48 | (-0.25- 1.26) | 1.99E-01 | 3.12E-01 | -0.26 | (-1.10- 0.67) | 5.76E-01 | 7.52E-01 | 1.94 | (-0.06- 3.98) | 5.77E-02 | 2.17E-01 | 4.73 | ( 2.25- 7.28) | **2.10E-04** | **8.39E-03** |
| 188 | SM(d18:2/17:0) | -0.74 | (-1.40--0.04) | **3.75E-02** | 7.37E-02 | -1.05 | (-1.83--0.21) | **1.60E-02** | 5.15E-02 | 0.98 | (-0.78- 2.78) | 2.76E-01 | 5.46E-01 | 1.61 | (-0.69- 3.97) | 1.69E-01 | 3.54E-01 |
| 189 | SM(35:2) (b) | -0.24 | (-1.22- 0.85) | 6.50E-01 | 7.36E-01 | -1.13 | (-2.18- 0.06) | 6.18E-02 | 1.56E-01 | -1.15 | (-3.40- 1.15) | 3.21E-01 | 5.98E-01 | -0.73 | (-3.52- 2.14) | 6.13E-01 | 7.74E-01 |
| 190 | SM(d18:1/18:0)/SM(d16:1/20:0) | -0.21 | (-0.94- 0.58) | 5.87E-01 | 6.98E-01 | 0.20 | (-0.73- 1.23) | 6.83E-01 | 8.39E-01 | 0.13 | (-1.70- 1.99) | 8.93E-01 | 9.59E-01 | 0.94 | (-1.33- 3.26) | 4.19E-01 | 6.04E-01 |
| 191 | SM(d18:2/18:1) | 0.37 | (-0.40- 1.19) | 3.56E-01 | 4.85E-01 | -0.44 | (-1.33- 0.54) | 3.66E-01 | 5.68E-01 | 0.92 | (-0.67- 2.54) | 2.55E-01 | 5.14E-01 | 2.18 | ( 0.24- 4.16) | **2.81E-02** | 1.19E-01 |
| 192 | SM(37:1) | -0.91 | (-1.71--0.04) | **4.10E-02** | 7.91E-02 | -0.63 | (-1.64- 0.49) | 2.56E-01 | 4.48E-01 | 0.13 | (-1.90- 2.20) | 9.02E-01 | 9.64E-01 | 0.37 | (-2.29- 3.10) | 7.87E-01 | 8.98E-01 |
| 193 | SM(37:2) | -0.70 | (-1.47- 0.14) | 9.82E-02 | 1.73E-01 | -0.74 | (-1.67- 0.30) | 1.58E-01 | 3.22E-01 | -0.55 | (-2.77- 1.72) | 6.29E-01 | 8.41E-01 | -0.33 | (-2.97- 2.39) | 8.10E-01 | 9.17E-01 |
| 194 | SM(d18:1/20:0)/SM(d16:1/22:0) | 0.06 | (-0.64- 0.80) | 8.74E-01 | 9.20E-01 | 0.62 | (-0.26- 1.58) | 1.71E-01 | 3.39E-01 | 0.37 | (-1.21- 1.98) | 6.44E-01 | 8.52E-01 | 0.99 | (-1.07- 3.09) | 3.45E-01 | 5.41E-01 |
| 195 | SM(d18:2/20:0) | 0.26 | (-0.65- 1.26) | 5.87E-01 | 6.98E-01 | 0.45 | (-0.74- 1.80) | 4.74E-01 | 6.60E-01 | 0.22 | (-1.74- 2.22) | 8.26E-01 | 9.33E-01 | 1.04 | (-1.41- 3.55) | 4.06E-01 | 5.96E-01 |
| 196 | SM(38:3) (a) | -0.18 | (-0.91- 0.60) | 6.35E-01 | 7.30E-01 | -0.96 | (-1.76--0.08) | **3.34E-02** | 9.26E-02 | 1.86 | ( 0.24- 3.52) | **2.49E-02** | 1.23E-01 | 2.01 | ( 0.01- 4.05) | **4.88E-02** | 1.69E-01 |
| 197 | SM(38:3) (b) | 0.11 | (-0.83- 1.14) | 8.31E-01 | 8.90E-01 | -0.21 | (-1.28- 0.99) | 7.14E-01 | 8.47E-01 | -0.30 | (-2.65- 2.11) | 8.03E-01 | 9.16E-01 | 1.95 | (-0.75- 4.73) | 1.57E-01 | 3.40E-01 |
| 198 | SM(d16:1/23:0)/SM(d17:1/22:0) | -0.46 | (-1.12- 0.26) | 2.03E-01 | 3.16E-01 | -0.19 | (-1.02- 0.71) | 6.62E-01 | 8.28E-01 | 1.00 | (-1.03- 3.07) | 3.36E-01 | 6.14E-01 | 2.18 | (-0.31- 4.73) | 8.66E-02 | 2.35E-01 |
| 199 | SM(d18:1/22:0)/SM(d16:1/24:0) | 0.22 | (-0.51- 1.00) | 5.64E-01 | 6.80E-01 | 0.71 | (-0.20- 1.70) | 1.32E-01 | 2.79E-01 | 0.47 | (-1.22- 2.20) | 5.84E-01 | 8.05E-01 | 0.50 | (-1.47- 2.52) | 6.18E-01 | 7.76E-01 |
| 200 | SM(d16:1/24:1) | 0.13 | (-0.66- 0.98) | 7.60E-01 | 8.22E-01 | -0.42 | (-1.32- 0.59) | 4.01E-01 | 5.96E-01 | 2.65 | ( 0.81- 4.53) | **4.85E-03** | **4.82E-02** | 2.77 | ( 0.21- 5.39) | **3.39E-02** | 1.35E-01 |
| 201 | SM(d18:2/22:0) | 0.08 | (-0.66- 0.87) | 8.37E-01 | 8.90E-01 | 0.91 | (-0.11- 2.04) | 8.05E-02 | 1.87E-01 | -0.02 | (-1.56- 1.54) | 9.78E-01 | 9.94E-01 | 1.34 | (-0.63- 3.35) | 1.83E-01 | 3.70E-01 |
| 202 | SM(40:3) (a) | -0.26 | (-1.01- 0.56) | 5.27E-01 | 6.50E-01 | -0.61 | (-1.49- 0.36) | 2.06E-01 | 3.88E-01 | 1.80 | ( 0.02- 3.60) | **4.73E-02** | 1.97E-01 | 1.75 | (-0.50- 4.05) | 1.26E-01 | 2.98E-01 |
| 203 | SM(40:3) (b) | -0.72 | (-1.50- 0.12) | 9.23E-02 | 1.65E-01 | -0.99 | (-1.97- 0.10) | 7.25E-02 | 1.74E-01 | 0.58 | (-1.73- 2.95) | 6.23E-01 | 8.38E-01 | 1.69 | (-1.18- 4.63) | 2.49E-01 | 4.40E-01 |
| 204 | SM(41:0) | -0.29 | (-1.44- 1.02) | 6.51E-01 | 7.36E-01 | -0.66 | (-1.99- 0.90) | 3.87E-01 | 5.82E-01 | 0.08 | (-1.58- 1.76) | 9.27E-01 | 9.71E-01 | 1.03 | (-0.99- 3.08) | 3.18E-01 | 5.17E-01 |

| | | | | | | | | | | | | | | | | | |
|---|---|---|---|---|---|---|---|---|---|---|---|---|---|---|---|---|---|
| 205 | SM(41:1) | -0.33 | (-1.14- 0.56) | 4.52E-01 | 5.78E-01 | 0.08 | (-0.95- 1.23) | 8.82E-01 | 9.56E-01 | 0.36 | (-1.33- 2.08) | 6.74E-01 | 8.56E-01 | 0.83 | (-1.16- 2.85) | 4.15E-01 | 6.02E-01 |
| 206 | SM(d17:1/24:1) | -0.21 | (-0.92- 0.54) | 5.67E-01 | 6.80E-01 | -0.74 | (-1.58- 0.19) | 1.14E-01 | 2.46E-01 | 0.68 | (-1.01- 2.40) | 4.31E-01 | 6.87E-01 | 0.56 | (-1.67- 2.84) | 6.24E-01 | 7.82E-01 |
| 207 | SM(d18:2/23:0) | -0.18 | (-1.01- 0.73) | 6.93E-01 | 7.70E-01 | 0.34 | (-0.76- 1.58) | 5.55E-01 | 7.37E-01 | 0.38 | (-1.20- 1.97) | 6.40E-01 | 8.51E-01 | 1.43 | (-0.56- 3.45) | 1.59E-01 | 3.41E-01 |
| 208 | SM(d18:1/24:0) | -0.43 | (-1.12- 0.32) | 2.55E-01 | 3.81E-01 | 0.15 | (-0.79- 1.19) | 7.63E-01 | 8.85E-01 | -0.52 | (-2.00- 0.99) | 4.96E-01 | 7.35E-01 | -0.33 | (-2.11- 1.48) | 7.15E-01 | 8.49E-01 |
| 209 | SM(d18:1/24:1) | 0.49 | (-0.30- 1.35) | 2.29E-01 | 3.48E-01 | -0.14 | (-1.03- 0.82) | 7.61E-01 | 8.85E-01 | 0.19 | (-1.56- 1.97) | 8.32E-01 | 9.35E-01 | -0.55 | (-2.58- 1.52) | 5.97E-01 | 7.62E-01 |
| 210 | SM(d18:2/24:0) | -0.39 | (-1.07- 0.34) | 2.84E-01 | 4.15E-01 | 0.39 | (-0.55- 1.42) | 4.26E-01 | 6.21E-01 | -0.61 | (-1.92- 0.72) | 3.68E-01 | 6.37E-01 | -0.11 | (-1.83- 1.64) | 9.00E-01 | 9.70E-01 |
| 211 | SM(43:1) | -0.63 | (-1.58- 0.43) | 2.32E-01 | 3.51E-01 | -1.42 | (-2.48--0.21) | **2.33E-02** | 7.02E-02 | -1.48 | (-3.37- 0.45) | 1.31E-01 | 3.48E-01 | -1.13 | (-3.45- 1.25) | 3.48E-01 | 5.43E-01 |
| 212 | SM(44:1) | -0.46 | (-1.10- 0.22) | 1.78E-01 | 2.83E-01 | -0.37 | (-1.19- 0.53) | 4.11E-01 | 6.07E-01 | -2.46 | (-3.86--1.04) | **8.52E-04** | **2.04E-02** | -1.52 | (-3.23- 0.23) | 8.83E-02 | 2.37E-01 |
| 213 | SM(44:2) | -0.01 | (-0.68- 0.72) | 9.84E-01 | 9.92E-01 | -0.42 | (-1.25- 0.48) | 3.42E-01 | 5.43E-01 | -1.16 | (-2.80- 0.50) | 1.69E-01 | 4.07E-01 | -1.20 | (-3.16- 0.80) | 2.36E-01 | 4.31E-01 |
| 214 | SM(44:3) (a) | -0.37 | (-1.00- 0.30) | 2.67E-01 | 3.95E-01 | -0.38 | (-1.16- 0.48) | 3.75E-01 | 5.75E-01 | -0.41 | (-1.83- 1.02) | 5.69E-01 | 7.94E-01 | -0.71 | (-2.52- 1.13) | 4.46E-01 | 6.28E-01 |
| 215 | LPC(14:0) [sn2] | 2.00 | ( 1.33- 2.71) | **1.28E-09** | **8.39E-09** | 1.49 | ( 0.75- 2.28) | **5.65E-05** | **4.76E-04** | 4.13 | ( 1.16- 7.19) | **6.34E-03** | 5.25E-02 | 6.01 | ( 1.90-10.30) | **4.15E-03** | **4.07E-02** |
| 216 | LPC(14:0) [sn1] | 1.76 | ( 1.12- 2.43) | **2.26E-08** | **1.23E-07** | 1.49 | ( 0.76- 2.27) | **4.09E-05** | **3.65E-04** | 3.98 | ( 1.07- 6.96) | **7.29E-03** | 5.64E-02 | 5.83 | ( 1.87- 9.93) | **3.84E-03** | **3.84E-02** |
| 217 | LPC(15:0) [sn2] | 1.45 | ( 0.66- 2.30) | **2.29E-04** | **7.18E-04** | 0.60 | (-0.26- 1.53) | 1.75E-01 | 3.44E-01 | 0.77 | (-2.04- 3.65) | 5.95E-01 | 8.10E-01 | 3.15 | (-0.54- 6.98) | 9.44E-02 | 2.49E-01 |
| 218 | LPC(15:0) [sn1] | 0.93 | ( 0.23- 1.69) | **8.96E-03** | **2.07E-02** | 0.67 | (-0.13- 1.53) | 1.01E-01 | 2.26E-01 | 0.40 | (-2.22- 3.08) | 7.69E-01 | 9.12E-01 | 2.66 | (-0.77- 6.21) | 1.29E-01 | 3.01E-01 |
| 219 | LPC(16:0) [sn2] | 1.54 | ( 0.94- 2.17) | **1.98E-07** | **9.14E-07** | 2.07 | ( 1.31- 2.88) | **2.86E-08** | **1.15E-06** | -0.59 | (-2.69- 1.54) | 5.82E-01 | 8.05E-01 | 2.50 | (-0.13- 5.20) | 6.20E-02 | 1.88E-01 |
| 220 | LPC(16:0) [sn1] | 1.85 | ( 1.12- 2.64) | **3.00E-07** | **1.36E-06** | 2.15 | ( 1.25- 3.13) | **1.15E-06** | **1.79E-05** | -0.69 | (-3.16- 1.83) | 5.85E-01 | 8.05E-01 | 2.18 | (-0.81- 5.25) | 1.53E-01 | 3.40E-01 |
| 221 | LPC(16:1) [sn2] | 4.38 | ( 3.37- 5.47) | **4.90E-20** | **1.18E-17** | 3.52 | ( 2.46- 4.66) | **2.65E-12** | **1.27E-09** | 4.20 | ( 1.71- 6.76) | **9.57E-04** | **2.14E-02** | 6.65 | ( 3.67- 9.72) | **1.49E-05** | **1.53E-03** |
| 222 | LPC(16:1) [sn1] | 4.38 | ( 3.43- 5.40) | **2.15E-22** | **1.03E-19** | 3.29 | ( 2.29- 4.37) | **6.50E-12** | **1.56E-09** | 3.09 | ( 0.68- 5.56) | **1.19E-02** | 7.12E-02 | 5.26 | ( 2.38- 8.22) | **3.56E-04** | **1.14E-02** |
| 223 | LPC(15-MHDA) [sn2] | 1.64 | ( 0.37- 3.07) | **1.03E-02** | **2.35E-02** | -0.03 | (-1.21- 1.32) | 9.65E-01 | 9.86E-01 | -1.63 | (-5.04- 1.91) | 3.61E-01 | 6.36E-01 | 1.64 | (-2.82- 6.29) | 4.75E-01 | 6.59E-01 |
| 224 | LPC(15-MHDA) [sn1] / LPC(17:0) [sn2] | 0.99 | ( 0.04- 2.03) | **4.04E-02** | 7.83E-02 | 0.02 | (-0.93- 1.06) | 9.72E-01 | 9.87E-01 | -2.02 | (-4.94- 0.99) | 1.85E-01 | 4.25E-01 | 0.77 | (-2.85- 4.52) | 6.80E-01 | 8.31E-01 |
| 225 | LPC(17:0) [sn1] | 0.13 | (-0.49- 0.79) | 6.88E-01 | 7.66E-01 | 0.34 | (-0.40- 1.14) | 3.77E-01 | 5.75E-01 | -2.13 | (-4.56- 0.35) | 9.18E-02 | 2.70E-01 | 0.61 | (-2.30- 3.60) | 6.84E-01 | 8.33E-01 |
| 226 | LPC(17:1) [sn1] (a) / LPC(17:1) [sn2] (b) | 2.44 | ( 1.55- 3.41) | **2.06E-08** | **1.14E-07** | 0.87 | (-0.01- 1.82) | 5.23E-02 | 1.38E-01 | 0.00 | (-2.59- 2.66) | 9.99E-01 | 9.99E-01 | 2.13 | (-1.21- 5.59) | 2.12E-01 | 4.01E-01 |
| 227 | LPC(18:0) [sn2] | 0.72 | ( 0.16- 1.31) | **1.09E-02** | **2.43E-02** | 1.42 | ( 0.64- 2.25) | **2.43E-04** | **1.69E-03** | -0.75 | (-3.04- 1.59) | 5.25E-01 | 7.57E-01 | 2.97 | ( 0.20- 5.81) | **3.56E-02** | 1.38E-01 |
| 228 | LPC(18:0) [sn1] | 0.81 | ( 0.23- 1.43) | **5.76E-03** | **1.40E-02** | 1.29 | ( 0.50- 2.13) | **1.03E-03** | **4.99E-03** | -0.72 | (-3.29- 1.92) | 5.87E-01 | 8.05E-01 | 2.97 | (-0.01- 6.04) | 5.08E-02 | 1.71E-01 |
| 229 | LPC(18:1) [sn2] | 3.96 | ( 3.00- 4.99) | **4.21E-18** | **4.04E-16** | 3.16 | ( 2.16- 4.24) | **4.94E-11** | **5.23E-09** | -1.92 | (-4.17- 0.37) | 9.93E-02 | 2.86E-01 | 0.79 | (-2.24- 3.90) | 6.13E-01 | 7.74E-01 |
| 230 | LPC(18:1) [sn1] | 3.71 | ( 2.86- 4.62) | **1.22E-19** | **1.96E-17** | 2.91 | ( 2.01- 3.87) | **2.21E-11** | **3.54E-09** | -1.93 | (-4.00- 0.19) | 7.43E-02 | 2.45E-01 | 0.02 | (-2.53- 2.65) | 9.86E-01 | 9.91E-01 |
| 231 | LPC(18:2) [sn2] | 2.88 | ( 1.95- 3.88) | **1.19E-10** | **8.82E-10** | 2.95 | ( 1.85- 4.14) | **2.38E-08** | **1.04E-06** | -1.36 | (-3.44- 0.77) | 2.09E-01 | 4.58E-01 | -0.57 | (-3.16- 2.10) | 6.71E-01 | 8.24E-01 |
| 232 | LPC(18:2) [sn1] | 2.93 | ( 1.99- 3.95) | **9.62E-11** | **7.21E-10** | 2.87 | ( 1.77- 4.07) | **6.68E-08** | **2.14E-06** | -1.55 | (-3.85- 0.81) | 1.95E-01 | 4.38E-01 | -1.69 | (-4.46- 1.16) | 2.41E-01 | 4.32E-01 |
| 233 | LPC(18:3) [sn2] (a) | 2.28 | ( 1.42- 3.20) | **5.55E-08** | **2.67E-07** | 2.29 | ( 1.27- 3.40) | **4.21E-06** | **5.32E-05** | 0.04 | (-2.69- 2.85) | 9.76E-01 | 9.94E-01 | 1.46 | (-2.11- 5.16) | 4.26E-01 | 6.08E-01 |
| 234 | LPC(18:3) [sn1] (a)/LPC(18:3) [sn2] (b) | 4.69 | ( 3.44- 6.07) | **7.42E-16** | **2.04E-14** | 4.00 | ( 2.66- 5.48) | **2.33E-10** | **1.87E-08** | 0.54 | (-2.41- 3.59) | 7.20E-01 | 8.86E-01 | 2.51 | (-1.38- 6.54) | 2.07E-01 | 3.98E-01 |
| 235 | LPC(18:3) (a) [sn1] [104_sn1] | 4.40 | ( 3.21- 5.70) | **2.66E-15** | **4.91E-14** | 3.62 | ( 2.33- 5.04) | **2.99E-09** | **1.60E-07** | 0.80 | (-2.16- 3.84) | 5.99E-01 | 8.11E-01 | 2.36 | (-1.54- 6.40) | 2.37E-01 | 4.31E-01 |
| 236 | LPC(19:0) [sn1] (a) / LPC(19:0) [sn2] (b) | 0.02 | (-0.83- 0.94) | 9.73E-01 | 9.89E-01 | -0.49 | (-1.41- 0.52) | 3.25E-01 | 5.20E-01 | -3.27 | (-5.49--0.99) | **5.33E-03** | **4.82E-02** | -0.95 | (-3.63- 1.80) | 4.92E-01 | 6.78E-01 |
| 237 | LPC(20:0) [sn1] | 0.27 | (-0.27- 0.83) | 3.36E-01 | 4.67E-01 | 0.39 | (-0.28- 1.12) | 2.61E-01 | 4.51E-01 | -4.17 | (-6.10--2.20) | **5.50E-05** | **4.46E-03** | -2.07 | (-4.53- 0.45) | 1.06E-01 | 2.68E-01 |
| 238 | LPC(20:1) [sn2] | 1.66 | ( 0.97- 2.38) | **8.89E-07** | **3.74E-06** | 1.10 | ( 0.31- 1.95) | **5.90E-03** | **2.30E-02** | -3.18 | (-5.31--1.00) | **4.74E-03** | **4.82E-02** | -0.02 | (-2.60- 2.63) | 9.88E-01 | 9.91E-01 |
| 239 | LPC(20:1) [sn1] | 2.06 | ( 1.41- 2.75) | **1.46E-10** | **1.06E-09** | 1.32 | ( 0.60- 2.09) | **2.38E-04** | **1.69E-03** | -3.45 | (-5.56--1.28) | **2.11E-03** | **3.27E-02** | -0.70 | (-3.17- 1.83) | 5.79E-01 | 7.46E-01 |
| 240 | LPC(20:2) [sn2] | 3.91 | ( 2.89- 5.02) | **7.66E-16** | **2.04E-14** | 3.66 | ( 2.49- 4.95) | **5.44E-11** | **5.23E-09** | -1.54 | (-3.61- 0.58) | 1.52E-01 | 3.75E-01 | 1.28 | (-1.25- 3.88) | 3.22E-01 | 5.19E-01 |
| 241 | LPC(20:2) [sn1] | 3.72 | ( 2.82- 4.68) | **3.71E-18** | **4.04E-16** | 2.88 | ( 1.83- 4.03) | **1.33E-08** | **6.37E-07** | -1.87 | (-3.97- 0.28) | 8.69E-02 | 2.66E-01 | 0.84 | (-1.68- 3.42) | 5.16E-01 | 6.98E-01 |
| 242 | LPC(20:3) [sn2] | 2.83 | ( 1.94- 3.80) | **5.17E-11** | **4.07E-10** | 2.33 | ( 1.30- 3.47) | **4.18E-06** | **5.32E-05** | 1.02 | (-1.28- 3.37) | 3.84E-01 | 6.55E-01 | 5.44 | ( 2.42- 8.55) | **4.27E-04** | **1.28E-02** |
| 243 | LPC(20:3) [sn1] | 3.53 | ( 2.56- 4.59) | **3.33E-14** | **4.32E-13** | 2.97 | ( 1.84- 4.21) | **4.25E-08** | **1.46E-06** | 0.88 | (-1.71- 3.53) | 5.07E-01 | 7.44E-01 | 4.99 | ( 1.89- 8.19) | **1.64E-03** | **2.62E-02** |
| 244 | LPC(20:3) [104_sn1] | 3.58 | ( 2.60- 4.65) | **2.02E-14** | **3.01E-13** | 2.80 | ( 1.72- 3.99) | **9.31E-08** | **2.74E-06** | 0.99 | (-1.63- 3.68) | 4.62E-01 | 7.17E-01 | 5.08 | ( 1.85- 8.41) | **2.07E-03** | **2.63E-02** |
| 245 | LPC(20:4) [sn2] | 2.46 | ( 1.61- 3.37) | **2.40E-09** | **1.50E-08** | 2.31 | ( 1.35- 3.35) | **8.21E-07** | **1.36E-05** | 0.06 | (-2.09- 2.24) | 9.59E-01 | 9.88E-01 | 4.00 | ( 1.39- 6.67) | **2.70E-03** | **3.09E-02** |
| 246 | LPC(20:4) [sn1] | 2.74 | ( 1.83- 3.72) | **4.87E-10** | **3.39E-09** | 2.59 | ( 1.58- 3.70) | **1.46E-07** | **3.50E-06** | 0.00 | (-2.41- 2.46) | 9.97E-01 | 9.99E-01 | 2.65 | (-0.11- 5.48) | 5.97E-02 | 1.87E-01 |
| 247 | LPC(20:5) [sn2] | 0.31 | (-1.09- 1.93) | 6.80E-01 | 7.61E-01 | 0.70 | (-0.99- 2.71) | 4.37E-01 | 6.27E-01 | 3.35 | (-0.26- 7.09) | 6.93E-02 | 2.38E-01 | 6.65 | ( 2.03-11.49) | **4.73E-03** | **4.18E-02** |
| 248 | LPC(20:5) [sn1] | 0.69 | (-0.74- 2.35) | 3.63E-01 | 4.90E-01 | 0.57 | (-1.07- 2.51) | 5.17E-01 | 7.05E-01 | 3.54 | (-0.28- 7.50) | 6.95E-02 | 2.38E-01 | 6.45 | ( 1.75-11.36) | **6.99E-03** | 5.51E-02 |
| 249 | LPC(22:4) [sn2] | 2.45 | ( 1.53- 3.43) | **3.64E-08** | **1.86E-07** | 2.66 | ( 1.58- 3.85) | **4.14E-07** | **7.09E-06** | -1.43 | (-3.81- 1.01) | 2.46E-01 | 5.03E-01 | 2.15 | (-0.87- 5.25) | 1.63E-01 | 3.44E-01 |
| 250 | LPC(22:4) [sn1] | 2.78 | ( 1.91- 3.70) | **3.03E-11** | **2.47E-10** | 2.61 | ( 1.60- 3.72) | **1.03E-07** | **2.74E-06** | -0.19 | (-2.72- 2.40) | 8.82E-01 | 9.49E-01 | 2.15 | (-0.85- 5.23) | 1.60E-01 | 3.41E-01 |
| 251 | LPC(22:5) [sn2] (n3) | 1.51 | ( 0.45- 2.67) | **4.32E-03** | **1.09E-02** | 2.18 | ( 0.81- 3.71) | **1.22E-03** | **5.68E-03** | -0.01 | (-2.16- 2.18) | 9.91E-01 | 9.99E-01 | 2.49 | (-0.14- 5.18) | 6.37E-02 | 1.90E-01 |
| 252 | LPC(22:5) [sn1] (n3)/LPC(22:5) [sn2] (n6) | 1.94 | ( 1.04- 2.92) | **1.19E-05** | **4.49E-05** | 2.46 | ( 1.30- 3.73) | **1.26E-05** | **1.37E-04** | -0.32 | (-2.56- 1.98) | 7.84E-01 | 9.16E-01 | 2.00 | (-0.65- 4.72) | 1.39E-01 | 3.14E-01 |
| 253 | LPC(22:5) [sn1] (n6) | 4.71 | ( 3.19- 6.41) | **2.08E-11** | **1.72E-10** | 4.30 | ( 2.54- 6.32) | **1.85E-07** | **3.86E-06** | -1.41 | (-4.32- 1.58) | 3.50E-01 | 6.24E-01 | 1.02 | (-2.44- 4.60) | 5.67E-01 | 7.36E-01 |
| 254 | LPC(22:5) (n3) [sn1] [104_sn1] | 0.84 | (-0.13- 1.89) | 9.17E-02 | 1.64E-01 | 1.38 | ( 0.17- 2.74) | **2.44E-02** | 7.22E-02 | 0.33 | (-1.98- 2.68) | 7.83E-01 | 9.16E-01 | 1.91 | (-0.91- 4.81) | 1.85E-01 | 3.70E-01 |
| 255 | LPC(22:6) [sn2] | 1.49 | ( 0.60- 2.45) | **7.81E-04** | **2.19E-03** | 1.86 | ( 0.82- 3.00) | **3.09E-04** | **2.03E-03** | -0.09 | (-2.54- 2.41) | 9.42E-01 | 9.79E-01 | 1.58 | (-1.47- 4.73) | 3.10E-01 | 5.08E-01 |
| 256 | LPC(22:6) [sn1] | 2.03 | ( 1.07- 3.07) | **1.50E-05** | **5.60E-05** | 2.19 | ( 1.10- 3.38) | **4.10E-05** | **3.65E-04** | 0.27 | (-2.39- 3.01) | 8.41E-01 | 9.40E-01 | 0.63 | (-2.40- 3.75) | 6.86E-01 | 8.34E-01 |
| 257 | LPC(O-16:0) | 1.64 | ( 0.97- 2.36) | **8.96E-07** | **3.74E-06** | 1.94 | ( 1.17- 2.76) | **3.11E-07** | **5.74E-06** | -0.82 | (-3.14- 1.56) | 4.94E-01 | 7.35E-01 | 2.27 | (-0.57- 5.20) | 1.18E-01 | 2.83E-01 |

| | | | | | | | | | | | | | | | | | |
|---|---|---|---|---|---|---|---|---|---|---|---|---|---|---|---|---|---|
| 258 | LPC(O-18:0) | 0.55 | (-0.23- 1.38) | 1.71E-01 | 2.74E-01 | 0.05 | (-0.77- 0.94) | 9.15E-01 | 9.70E-01 | -2.38 | (-4.94- 0.25) | 7.56E-02 | 2.45E-01 | 0.76 | (-2.38- 3.99) | 6.38E-01 | 7.96E-01 |
| 259 | LPC(O-18:1) | 1.90 | ( 1.20- 2.64) | **3.74E-08** | **1.89E-07** | 1.74 | ( 0.96- 2.57) | **6.71E-06** | **8.14E-05** | -1.36 | (-3.72- 1.05) | 2.64E-01 | 5.28E-01 | 1.34 | (-1.42- 4.18) | 3.41E-01 | 5.39E-01 |
| 260 | LPC(O-20:0) | 0.44 | (-0.66- 1.67) | 4.43E-01 | 5.70E-01 | -0.41 | (-1.60- 0.96) | 5.39E-01 | 7.27E-01 | -3.78 | (-6.12--1.38) | **2.35E-03** | **3.53E-02** | -0.31 | (-3.22- 2.69) | 8.39E-01 | 9.30E-01 |
| 261 | LPC(O-20:1) | 2.44 | ( 1.22- 3.79) | **4.14E-05** | **1.45E-04** | 3.29 | ( 1.62- 5.21) | **3.99E-05** | **3.65E-04** | -0.97 | (-3.77- 1.91) | 5.01E-01 | 7.38E-01 | 1.82 | (-2.15- 5.96) | 3.71E-01 | 5.66E-01 |
| 262 | LPC(O-22:0) | 0.23 | (-0.41- 0.91) | 4.88E-01 | 6.13E-01 | 0.22 | (-0.50- 1.00) | 5.57E-01 | 7.37E-01 | -1.81 | (-3.29--0.31) | **1.87E-02** | 1.01E-01 | -1.92 | (-3.89- 0.09) | 6.11E-02 | 1.88E-01 |
| 263 | LPC(O-22:1) | 1.62 | ( 0.86- 2.44) | **1.60E-05** | **5.90E-05** | 0.80 | (-0.07- 1.73) | 7.14E-02 | 1.73E-01 | -2.10 | (-4.24- 0.10) | 6.10E-02 | 2.20E-01 | -1.54 | (-4.28- 1.28) | 2.80E-01 | 4.80E-01 |
| 264 | LPC(O-24:0) | -0.48 | (-1.17- 0.26) | 1.94E-01 | 3.06E-01 | -0.38 | (-1.13- 0.45) | 3.58E-01 | 5.60E-01 | -1.06 | (-2.58- 0.49) | 1.77E-01 | 4.20E-01 | -1.85 | (-3.86- 0.21) | 7.72E-02 | 2.18E-01 |
| 265 | LPC(O-24:1) | 0.86 | ( 0.15- 1.61) | **1.61E-02** | **3.54E-02** | -0.04 | (-0.79- 0.78) | 9.26E-01 | 9.70E-01 | -0.68 | (-2.41- 1.09) | 4.48E-01 | 6.98E-01 | -0.18 | (-2.56- 2.26) | 8.84E-01 | 9.67E-01 |
| 266 | LPC(O-24:2) | 0.25 | (-0.43- 0.97) | 4.82E-01 | 6.07E-01 | 0.13 | (-0.69- 1.03) | 7.58E-01 | 8.85E-01 | -1.61 | (-3.70- 0.53) | 1.38E-01 | 3.52E-01 | -1.36 | (-4.11- 1.47) | 3.41E-01 | 5.39E-01 |
| 267 | LPC(P-16:0) | 1.02 | ( 0.39- 1.69) | **1.27E-03** | **3.51E-03** | 1.30 | ( 0.55- 2.10) | **5.10E-04** | **2.82E-03** | -0.77 | (-3.08- 1.60) | 5.20E-01 | 7.52E-01 | 2.16 | (-0.52- 4.90) | 1.14E-01 | 2.80E-01 |
| 268 | LPC(P-18:0) | 1.89 | ( 1.20- 2.63) | **3.00E-08** | **1.60E-07** | 1.58 | ( 0.81- 2.39) | **3.27E-05** | **3.08E-04** | -1.58 | (-3.89- 0.78) | 1.86E-01 | 4.25E-01 | 1.26 | (-1.51- 4.10) | 3.75E-01 | 5.66E-01 |
| 269 | LPC(P-18:1) | 1.24 | ( 0.40- 2.14) | **3.19E-03** | **8.23E-03** | 1.39 | ( 0.34- 2.56) | **8.86E-03** | **3.15E-02** | -0.72 | (-3.39- 2.02) | 6.00E-01 | 8.11E-01 | 3.19 | (-0.01- 6.49) | 5.06E-02 | 1.71E-01 |
| 270 | LPC(P-20:0) | 2.59 | ( 1.67- 3.57) | **5.48E-09** | **3.21E-08** | 1.26 | ( 0.33- 2.29) | **7.45E-03** | **2.80E-02** | -2.47 | (-4.95- 0.07) | 5.69E-02 | 2.17E-01 | -0.91 | (-4.05- 2.34) | 5.76E-01 | 7.45E-01 |
| 271 | LPE(16:0) [sn2] | 1.63 | ( 0.86- 2.47) | **2.17E-05** | **7.90E-05** | 3.34 | ( 2.23- 4.55) | **2.77E-10** | **1.90E-08** | -2.11 | (-4.27- 0.11) | 6.24E-02 | 2.23E-01 | -1.46 | (-4.09- 1.23) | 2.83E-01 | 4.80E-01 |
| 272 | LPE(16:0) [sn1] | 1.81 | ( 1.08- 2.59) | **4.95E-07** | **2.16E-06** | 3.02 | ( 1.99- 4.14) | **1.07E-09** | **6.42E-08** | -1.76 | (-3.77- 0.28) | 9.04E-02 | 2.70E-01 | -1.01 | (-3.55- 1.61) | 4.44E-01 | 6.26E-01 |
| 273 | LPE(18:0) [sn2] | 1.12 | ( 0.47- 1.82) | **6.66E-04** | **1.93E-03** | 1.94 | ( 1.04- 2.90) | **1.05E-05** | **1.17E-04** | -1.99 | (-3.76--0.19) | **3.03E-02** | 1.43E-01 | 0.07 | (-2.43- 2.63) | 9.57E-01 | 9.75E-01 |
| 274 | LPE(18:0) [sn1] | 1.17 | ( 0.55- 1.83) | **1.80E-04** | **5.75E-04** | 2.23 | ( 1.30- 3.23) | **9.65E-07** | **1.54E-05** | -1.75 | (-3.65- 0.18) | 7.51E-02 | 2.45E-01 | 0.39 | (-2.11- 2.96) | 7.61E-01 | 8.85E-01 |
| 275 | LPE(18:1) [sn2] | 2.78 | ( 1.90- 3.72) | **5.35E-11** | **4.14E-10** | 2.29 | ( 1.28- 3.39) | **3.51E-06** | **4.67E-05** | -2.39 | (-4.81- 0.08) | 5.79E-02 | 2.17E-01 | 0.11 | (-3.12- 3.44) | 9.48E-01 | 9.75E-01 |
| 276 | LPE(18:1) [sn1] | 3.45 | ( 2.51- 4.46) | **1.19E-14** | **1.85E-13** | 2.27 | ( 1.30- 3.34) | **1.89E-06** | **2.84E-05** | -1.74 | (-4.22- 0.81) | 1.77E-01 | 4.20E-01 | -0.13 | (-3.42- 3.27) | 9.40E-01 | 9.75E-01 |
| 277 | LPE(18:2) [sn2] | 3.67 | ( 2.45- 5.01) | **1.97E-10** | **1.39E-09** | 3.61 | ( 2.16- 5.24) | **1.53E-07** | **3.51E-06** | -1.63 | (-4.13- 0.93) | 2.08E-01 | 4.58E-01 | -1.73 | (-4.83- 1.46) | 2.82E-01 | 4.80E-01 |
| 278 | LPE(18:2) [sn1] | 3.94 | ( 2.79- 5.19) | **3.20E-13** | **3.65E-12** | 3.44 | ( 2.08- 4.96) | **1.00E-07** | **2.74E-06** | -1.56 | (-4.10- 1.05) | 2.37E-01 | 4.91E-01 | -1.71 | (-4.76- 1.45) | 2.83E-01 | 4.80E-01 |
| 279 | LPE(20:4) [sn1] | 3.50 | ( 2.47- 4.61) | **6.96E-13** | **7.43E-12** | 2.76 | ( 1.68- 3.95) | **1.42E-07** | **3.50E-06** | -1.81 | (-3.60- 0.02) | 5.20E-02 | 2.08E-01 | -0.42 | (-2.66- 1.88) | 7.16E-01 | 8.49E-01 |
| 280 | LPE(22:6) [sn2] | 2.13 | ( 1.18- 3.15) | **4.65E-06** | **1.82E-05** | 2.27 | ( 1.19- 3.46) | **1.76E-05** | **1.87E-04** | -0.99 | (-3.33- 1.40) | 4.11E-01 | 6.75E-01 | 0.12 | (-2.65- 2.98) | 9.32E-01 | 9.75E-01 |
| 281 | LPE(22:6) [sn1] | 2.40 | ( 1.44- 3.44) | **2.49E-07** | **1.14E-06** | 2.46 | ( 1.38- 3.64) | **2.68E-06** | **3.78E-05** | -0.35 | (-2.65- 2.00) | 7.66E-01 | 9.12E-01 | -0.55 | (-3.14- 2.11) | 6.79E-01 | 8.31E-01 |
| 282 | PC(28:0) | 0.04 | (-0.73- 0.87) | 9.20E-01 | 9.58E-01 | -0.63 | (-1.55- 0.40) | 2.22E-01 | 4.08E-01 | 3.28 | (-0.56- 7.26) | 9.45E-02 | 2.76E-01 | 2.22 | (-2.23- 6.88) | 3.32E-01 | 5.31E-01 |
| 283 | PC(14:0_16:0) | 0.24 | (-0.64- 1.20) | 6.04E-01 | 7.09E-01 | -0.67 | (-1.66- 0.44) | 2.28E-01 | 4.13E-01 | 0.90 | (-1.55- 3.41) | 4.73E-01 | 7.26E-01 | -0.10 | (-3.15- 3.05) | 9.52E-01 | 9.75E-01 |
| 284 | PC(31:0) (a) | 0.40 | (-0.78- 1.72) | 5.25E-01 | 6.50E-01 | -1.56 | (-2.77--0.15) | **3.13E-02** | 8.83E-02 | 2.46 | (-2.13- 7.26) | 2.97E-01 | 5.68E-01 | 0.72 | (-5.13- 6.93) | 8.13E-01 | 9.17E-01 |
| 285 | PC(31:0) (b) | -0.86 | (-1.93- 0.36) | 1.58E-01 | 2.57E-01 | -1.02 | (-2.37- 0.58) | 1.97E-01 | 3.76E-01 | 0.17 | (-2.57- 2.99) | 9.05E-01 | 9.65E-01 | -0.52 | (-4.15- 3.26) | 7.84E-01 | 8.98E-01 |
| 286 | PC(31:1) | 0.49 | (-0.57- 1.68) | 3.78E-01 | 5.07E-01 | -1.21 | (-2.28- 0.02) | 5.28E-02 | 1.38E-01 | 4.35 | ( 0.93- 7.88) | **1.26E-02** | 7.37E-02 | 2.19 | (-2.18- 6.76) | 3.28E-01 | 5.27E-01 |
| 287 | PC(16:0_16:0) | -0.25 | (-1.08- 0.66) | 5.73E-01 | 6.85E-01 | -0.92 | (-1.86- 0.12) | 7.96E-02 | 1.86E-01 | -0.81 | (-2.48- 0.89) | 3.45E-01 | 6.22E-01 | -1.74 | (-3.64- 0.21) | 7.91E-02 | 2.19E-01 |
| 288 | PC(32:1) | 1.65 | ( 0.51- 2.90) | **3.70E-03** | **9.44E-03** | 0.08 | (-1.10- 1.42) | 8.96E-01 | 9.63E-01 | 4.07 | ( 1.14- 7.08) | **6.52E-03** | 5.30E-02 | 3.13 | (-0.08- 6.44) | 5.60E-02 | 1.79E-01 |
| 289 | PC(32:2) | 0.46 | (-0.41- 1.41) | 3.04E-01 | 4.38E-01 | -0.31 | (-1.30- 0.78) | 5.58E-01 | 7.37E-01 | 3.33 | ( 0.60- 6.13) | **1.67E-02** | 9.22E-02 | 1.85 | (-1.51- 5.33) | 2.82E-01 | 4.80E-01 |
| 290 | PC(33:0) (a) | 0.02 | (-1.20- 1.41) | 9.76E-01 | 9.89E-01 | -1.71 | (-2.86--0.37) | **1.43E-02** | **4.71E-02** | 0.13 | (-3.32- 3.70) | 9.41E-01 | 9.79E-01 | 0.08 | (-4.35- 4.73) | 9.71E-01 | 9.85E-01 |
| 291 | PC(33:0) (b) | -1.04 | (-1.98- 0.01) | 5.31E-02 | 1.01E-01 | -1.91 | (-2.99--0.66) | **4.05E-03** | **1.62E-02** | -0.53 | (-2.90- 1.90) | 6.64E-01 | 8.54E-01 | -1.35 | (-4.28- 1.67) | 3.76E-01 | 5.66E-01 |
| 292 | PC(33:1) | -0.17 | (-0.98- 0.72) | 7.03E-01 | 7.75E-01 | -1.00 | (-1.89--0.02) | **4.68E-02** | 1.26E-01 | 0.38 | (-2.33- 3.18) | 7.83E-01 | 9.16E-01 | -0.22 | (-3.69- 3.37) | 9.01E-01 | 9.70E-01 |
| 293 | PC(33:2) | -1.09 | (-1.89--0.20) | **1.71E-02** | **3.71E-02** | -1.43 | (-2.35--0.38) | **8.68E-03** | **3.13E-02** | -0.35 | (-2.89- 2.27) | 7.91E-01 | 9.16E-01 | -1.50 | (-4.70- 1.81) | 3.68E-01 | 5.64E-01 |
| 294 | PC(16:0_18:0) | -0.19 | (-0.92- 0.60) | 6.24E-01 | 7.22E-01 | -0.32 | (-1.20- 0.64) | 4.99E-01 | 6.87E-01 | -1.01 | (-2.60- 0.62) | 2.22E-01 | 4.72E-01 | -0.37 | (-2.29- 1.58) | 7.05E-01 | 8.48E-01 |
| 295 | PC(16:0_18:1) | 0.36 | (-0.56- 1.37) | 4.58E-01 | 5.83E-01 | -0.22 | (-1.25- 0.93) | 6.92E-01 | 8.47E-01 | -0.48 | (-2.56- 1.65) | 6.56E-01 | 8.54E-01 | -0.18 | (-2.81- 2.53) | 8.96E-01 | 9.70E-01 |
| 296 | PC(16:0_18:2) | -0.89 | (-1.86- 0.20) | 1.05E-01 | 1.82E-01 | -0.07 | (-1.40- 1.47) | 9.22E-01 | 9.70E-01 | -0.76 | (-2.59- 1.11) | 4.23E-01 | 6.81E-01 | -1.00 | (-2.95- 1.00) | 3.22E-01 | 5.19E-01 |
| 297 | PC(16:1_18:2) | -0.55 | (-1.35- 0.33) | 2.14E-01 | 3.29E-01 | -0.99 | (-1.94- 0.08) | 6.84E-02 | 1.68E-01 | 2.00 | (-0.20- 4.25) | 7.55E-02 | 2.45E-01 | 2.02 | (-0.64- 4.74) | 1.36E-01 | 3.13E-01 |
| 298 | PC(16:0_18:3) (a) | 0.01 | (-1.00- 1.15) | 9.78E-01 | 9.89E-01 | 0.12 | (-1.08- 1.49) | 8.48E-01 | 9.37E-01 | 0.53 | (-1.95- 3.07) | 6.78E-01 | 8.57E-01 | 1.02 | (-2.12- 4.25) | 5.26E-01 | 7.05E-01 |
| 299 | PC(14:0_20:4) | 0.19 | (-0.59- 1.04) | 6.39E-01 | 7.32E-01 | -0.43 | (-1.33- 0.57) | 3.85E-01 | 5.82E-01 | 4.71 | ( 1.62- 7.89) | **2.76E-03** | **3.89E-02** | 5.46 | ( 1.40- 9.68) | **8.28E-03** | 6.02E-02 |
| 300 | PC(34:5) | -0.95 | (-2.00- 0.25) | 1.16E-01 | 1.97E-01 | -1.30 | (-2.48- 0.07) | 6.15E-02 | 1.56E-01 | 7.08 | ( 2.85-11.48) | **9.79E-04** | **2.14E-02** | 8.14 | ( 2.75-13.81) | **2.92E-03** | **3.19E-02** |
| 301 | PC(15-MHDA_18:1) | -0.60 | (-1.64- 0.56) | 2.93E-01 | 4.26E-01 | -1.99 | (-3.01--0.83) | **1.43E-03** | **6.39E-03** | -1.31 | (-4.61- 2.10) | 4.45E-01 | 6.97E-01 | -1.46 | (-5.59- 2.85) | 4.97E-01 | 6.83E-01 |
| 302 | PC(17:0_18:1) | -0.78 | (-1.44--0.07) | **3.11E-02** | 6.30E-02 | -1.36 | (-2.10--0.54) | **1.62E-03** | **7.06E-03** | -1.11 | (-3.18- 1.00) | 2.97E-01 | 5.68E-01 | -1.12 | (-3.82- 1.65) | 4.21E-01 | 6.04E-01 |
| 303 | PC(15-MHDA_18:2) | -0.90 | (-1.82- 0.13) | 8.58E-02 | 1.55E-01 | -1.96 | (-2.93--0.84) | **1.05E-03** | **5.02E-03** | -1.52 | (-4.66- 1.73) | 3.53E-01 | 6.25E-01 | -1.83 | (-5.87- 2.38) | 3.86E-01 | 5.75E-01 |
| 304 | PC(17:0_18:2) | -1.11 | (-1.84--0.32) | **6.91E-03** | **1.65E-02** | -1.46 | (-2.33--0.50) | **3.88E-03** | **1.57E-02** | -1.88 | (-3.90- 0.18) | 7.26E-02 | 2.45E-01 | -1.95 | (-4.60- 0.77) | 1.56E-01 | 3.40E-01 |
| 305 | PC(17:1_18:2) | -0.36 | (-1.09- 0.43) | 3.57E-01 | 4.85E-01 | -1.03 | (-1.85--0.13) | **2.64E-02** | 7.72E-02 | 0.98 | (-1.24- 3.25) | 3.86E-01 | 6.55E-01 | 1.74 | (-1.16- 4.73) | 2.40E-01 | 4.32E-01 |
| 306 | PC(15:0_20:3) | 0.05 | (-1.17- 1.44) | 9.41E-01 | 9.73E-01 | 0.07 | (-1.47- 1.87) | 9.37E-01 | 9.74E-01 | -0.26 | (-1.73- 1.23) | 7.25E-01 | 8.86E-01 | -0.56 | (-2.32- 1.24) | 5.38E-01 | 7.12E-01 |
| 307 | PC(15:0_20:4) | -0.90 | (-1.67--0.06) | **3.60E-02** | 7.11E-02 | -1.77 | (-2.58--0.87) | **2.74E-04** | **1.88E-03** | 0.98 | (-1.73- 3.77) | 4.80E-01 | 7.29E-01 | 1.92 | (-1.65- 5.63) | 2.93E-01 | 4.90E-01 |
| 308 | PC(35:5) | -1.91 | (-3.02--0.63) | **4.88E-03** | **1.21E-02** | -2.80 | (-3.98--1.39) | **3.66E-04** | **2.23E-03** | 4.57 | ( 0.35- 8.97) | **3.35E-02** | 1.52E-01 | 4.26 | (-1.36-10.19) | 1.39E-01 | 3.14E-01 |
| 309 | PC 36:0 | 0.35 | (-0.33- 1.08) | 3.17E-01 | 4.49E-01 | 0.18 | (-0.69- 1.12) | 7.01E-01 | 8.47E-01 | 0.86 | (-1.56- 3.34) | 4.88E-01 | 7.35E-01 | 2.54 | ( 0.02- 5.12) | **4.82E-02** | 1.69E-01 |
| 310 | PC(18:0_18:1) | 0.64 | (-0.11- 1.46) | 9.55E-02 | 1.69E-01 | 0.21 | (-0.66- 1.15) | 6.48E-01 | 8.15E-01 | 1.00 | (-0.81- 2.85) | 2.78E-01 | 5.47E-01 | 1.51 | (-1.20- 4.30) | 2.75E-01 | 4.77E-01 |

| | | | | | | | | | | | | | | | | | |
|---|---|---|---|---|---|---|---|---|---|---|---|---|---|---|---|---|---|
| 311 | PC(18:0_18:2) | -0.59 | (-1.39- 0.29) | 1.81E-01 | 2.88E-01 | -0.31 | (-1.31- 0.82) | 5.76E-01 | 7.52E-01 | 0.44 | (-1.52- 2.44) | 6.60E-01 | 8.54E-01 | -0.02 | (-2.47- 2.49) | 9.85E-01 | 9.91E-01 |
| 312 | PC(16:0_20:3) (a) | -0.19 | (-1.10- 0.81) | 7.00E-01 | 7.74E-01 | -0.66 | (-1.68- 0.49) | 2.49E-01 | 4.38E-01 | -2.82 | (-4.70--0.90) | **4.32E-03** | **4.82E-02** | -2.58 | (-5.25- 0.16) | 6.50E-02 | 1.92E-01 |
| 313 | PC(16:0_20:3) (b) | 0.42 | (-0.38- 1.29) | 3.15E-01 | 4.49E-01 | 0.31 | (-0.70- 1.43) | 5.58E-01 | 7.37E-01 | 0.96 | (-1.17- 3.13) | 3.79E-01 | 6.53E-01 | 3.32 | ( 0.37- 6.35) | **2.75E-02** | 1.19E-01 |
| 314 | PC(18:2_18:2) | 0.50 | (-0.69- 1.84) | 4.28E-01 | 5.57E-01 | 0.15 | (-1.24- 1.77) | 8.39E-01 | 9.33E-01 | -2.39 | (-5.31- 0.62) | 1.18E-01 | 3.26E-01 | -3.83 | (-7.27--0.26) | **3.57E-02** | 1.38E-01 |
| 315 | PC(16:1_20:4) | 0.45 | (-0.39- 1.36) | 3.02E-01 | 4.37E-01 | -0.40 | (-1.33- 0.64) | 4.35E-01 | 6.27E-01 | 3.02 | ( 0.78- 5.31) | **8.18E-03** | 6.04E-02 | 5.24 | ( 2.20- 8.37) | **7.43E-04** | **1.76E-02** |
| 316 | PC(16:0_20:5) | -1.88 | (-2.96--0.63) | **4.43E-03** | **1.10E-02** | -1.79 | (-3.07--0.27) | **2.27E-02** | 6.89E-02 | 3.68 | (-0.38- 7.91) | 7.61E-02 | 2.45E-01 | 5.20 | ( 0.48-10.14) | **3.08E-02** | 1.29E-01 |
| 317 | PC(36:6) | -0.64 | (-1.42- 0.21) | 1.35E-01 | 2.25E-01 | -0.93 | (-1.85- 0.09) | 7.21E-02 | 1.74E-01 | 5.11 | ( 1.94- 8.37) | **1.56E-03** | **2.77E-02** | 3.99 | (-0.02- 8.16) | 5.13E-02 | 1.71E-01 |
| 318 | PC(15-MHDA_20:4) | -1.12 | (-2.00--0.14) | **2.66E-02** | 5.51E-02 | -2.12 | (-3.08--1.03) | **3.59E-04** | **2.21E-03** | -0.66 | (-3.70- 2.48) | 6.76E-01 | 8.56E-01 | 1.62 | (-2.39- 5.79) | 4.32E-01 | 6.15E-01 |
| 319 | PC(17:0_20:4) | -1.63 | (-2.29--0.92) | **2.58E-05** | **9.30E-05** | -1.66 | (-2.41--0.84) | **1.77E-04** | **1.31E-03** | -0.55 | (-2.87- 1.83) | 6.45E-01 | 8.52E-01 | 0.96 | (-2.03- 4.04) | 5.30E-01 | 7.09E-01 |
| 320 | PC(15:0_22:6) | -1.99 | (-2.70--1.21) | **3.93E-06** | **1.55E-05** | -1.58 | (-2.48--0.57) | **3.04E-03** | **1.25E-02** | 0.74 | (-2.10- 3.66) | 6.11E-01 | 8.24E-01 | 0.52 | (-3.08- 4.26) | 7.80E-01 | 8.97E-01 |
| 321 | PC(38:2) | 0.98 | ( 0.23- 1.78) | **9.39E-03** | **2.16E-02** | 0.87 | (-0.10- 1.93) | 7.94E-02 | 1.86E-01 | 0.64 | (-0.76- 2.06) | 3.67E-01 | 6.37E-01 | 1.98 | ( 0.01- 3.98) | **4.89E-02** | 1.69E-01 |
| 322 | PC(18:0_20:3) | 0.19 | (-0.51- 0.93) | 6.09E-01 | 7.13E-01 | 0.11 | (-0.80- 1.11) | 8.24E-01 | 9.22E-01 | 4.07 | ( 1.92- 6.27) | **2.27E-04** | **9.89E-03** | 7.65 | ( 4.41-11.00) | **4.56E-06** | **1.04E-03** |
| 323 | PC(18:1_20:3) | 0.62 | (-0.22- 1.52) | 1.51E-01 | 2.47E-01 | -0.17 | (-1.07- 0.82) | 7.27E-01 | 8.57E-01 | 0.17 | (-1.75- 2.14) | 8.59E-01 | 9.40E-01 | 2.37 | (-0.35- 5.16) | 8.77E-02 | 2.37E-01 |
| 324 | PC(38:4) (b) | -0.05 | (-0.99- 0.99) | 9.22E-01 | 9.58E-01 | -0.06 | (-1.18- 1.21) | 9.26E-01 | 9.70E-01 | 0.85 | (-1.51- 3.27) | 4.82E-01 | 7.29E-01 | 3.49 | ( 0.40- 6.67) | **2.66E-02** | 1.19E-01 |
| 325 | PC(18:0_20:4) | -1.14 | (-1.87--0.34) | **5.99E-03** | **1.45E-02** | -0.86 | (-1.79- 0.17) | 9.85E-02 | 2.23E-01 | 1.09 | (-1.09- 3.30) | 3.28E-01 | 6.08E-01 | 3.01 | ( 0.14- 5.97) | **3.98E-02** | 1.49E-01 |
| 326 | PC(38:5) (a) | -1.04 | (-1.68--0.35) | **3.90E-03** | **9.89E-03** | -1.25 | (-2.06--0.35) | **7.37E-03** | **2.79E-02** | -0.36 | (-2.25- 1.57) | 7.14E-01 | 8.85E-01 | 1.30 | (-1.17- 3.83) | 3.02E-01 | 5.02E-01 |
| 327 | PC(38:5) (b) | 0.14 | (-0.70- 1.05) | 7.52E-01 | 8.15E-01 | 0.22 | (-0.84- 1.40) | 6.95E-01 | 8.47E-01 | 1.13 | (-1.57- 3.91) | 4.14E-01 | 6.76E-01 | 2.83 | (-0.37- 6.14) | 8.28E-02 | 2.28E-01 |
| 328 | PC(38:6) (a) | 0.03 | (-0.91- 1.07) | 9.51E-01 | 9.77E-01 | -0.01 | (-1.09- 1.19) | 9.80E-01 | 9.90E-01 | -0.19 | (-2.41- 2.08) | 8.66E-01 | 9.44E-01 | 1.00 | (-2.07- 4.17) | 5.25E-01 | 7.05E-01 |
| 329 | PC(16:0_22:6) | -1.37 | (-2.26--0.37) | **8.36E-03** | **1.96E-02** | -0.95 | (-2.03- 0.27) | 1.20E-01 | 2.58E-01 | 0.50 | (-2.15- 3.22) | 7.15E-01 | 8.85E-01 | -0.43 | (-3.81- 3.07) | 8.07E-01 | 9.17E-01 |
| 330 | PC(18:2_20:5) | -0.59 | (-1.59- 0.53) | 2.87E-01 | 4.18E-01 | -0.70 | (-1.90- 0.68) | 3.05E-01 | 4.98E-01 | 2.05 | (-1.40- 5.62) | 2.46E-01 | 5.03E-01 | 3.99 | (-0.32- 8.48) | 6.95E-02 | 2.03E-01 |
| 331 | PC(16:1_22:6) | -0.81 | (-1.50--0.06) | **3.54E-02** | 7.02E-02 | -1.22 | (-2.03--0.33) | **8.62E-03** | **3.13E-02** | 1.62 | (-0.94- 4.23) | 2.15E-01 | 4.66E-01 | 2.07 | (-1.01- 5.25) | 1.89E-01 | 3.72E-01 |
| 332 | PC(38:7)(c) | -0.92 | (-1.54--0.25) | **7.98E-03** | **1.88E-02** | -0.71 | (-1.49- 0.14) | 1.00E-01 | 2.25E-01 | 0.28 | (-1.86- 2.47) | 7.99E-01 | 9.16E-01 | 0.02 | (-2.61- 2.71) | 9.91E-01 | 9.91E-01 |
| 333 | PC(39:5)(a) | -1.75 | (-2.57--0.83) | **3.70E-04** | **1.10E-03** | -2.10 | (-3.10--0.95) | **7.13E-04** | **3.76E-03** | -0.44 | (-3.46- 2.68) | 7.80E-01 | 9.16E-01 | -0.32 | (-4.36- 3.90) | 8.80E-01 | 9.66E-01 |
| 334 | PC(39:5)(b) | -1.89 | (-2.68--1.01) | **8.04E-05** | **2.72E-04** | -1.82 | (-2.76--0.76) | **1.36E-03** | **6.19E-03** | -1.18 | (-4.17- 1.89) | 4.44E-01 | 6.97E-01 | -0.17 | (-3.85- 3.64) | 9.28E-01 | 9.75E-01 |
| 335 | PC(15-MHDA_22:6) | -1.84 | (-2.69--0.89) | **3.31E-04** | **9.86E-04** | -2.26 | (-3.27--1.11) | **3.42E-04** | **2.19E-03** | -1.04 | (-3.88- 1.88) | 4.77E-01 | 7.28E-01 | -1.22 | (-5.04- 2.75) | 5.38E-01 | 7.12E-01 |
| 336 | PC(17:0_22:6) | -2.46 | (-3.13--1.72) | **6.30E-09** | **3.64E-08** | -1.74 | (-2.64--0.71) | **1.44E-03** | **6.39E-03** | -0.28 | (-3.17- 2.70) | 8.50E-01 | 9.40E-01 | -0.45 | (-4.08- 3.32) | 8.12E-01 | 9.17E-01 |
| 337 | PC(18:0_22:5) (n6) | -1.73 | (-3.03--0.17) | **3.09E-02** | 6.28E-02 | -0.35 | (-2.32- 2.14) | 7.63E-01 | 8.85E-01 | 1.64 | (-0.82- 4.17) | 1.92E-01 | 4.33E-01 | 3.63 | ( 0.39- 6.98) | **2.81E-02** | 1.19E-01 |
| 338 | PC(18:0_22:6) | -1.78 | (-2.56--0.92) | **1.28E-04** | **4.22E-04** | -0.83 | (-1.94- 0.44) | 1.91E-01 | 3.68E-01 | 3.12 | ( 0.46- 5.86) | **2.16E-02** | 1.10E-01 | 3.17 | (-0.31- 6.78) | 7.42E-02 | 2.11E-01 |
| 339 | PC(40:7) (a) | 1.11 | (-0.16- 2.53) | 8.81E-02 | 1.58E-01 | 0.82 | (-0.56- 2.41) | 2.57E-01 | 4.48E-01 | 0.56 | (-1.86- 3.03) | 6.53E-01 | 8.54E-01 | 2.78 | (-0.68- 6.36) | 1.16E-01 | 2.80E-01 |
| 340 | PC(18:1_22:6) (a) | -0.29 | (-1.18- 0.70) | 5.54E-01 | 6.72E-01 | -0.54 | (-1.59- 0.63) | 3.50E-01 | 5.53E-01 | -1.88 | (-4.02- 0.31) | 9.10E-02 | 2.70E-01 | -0.98 | (-3.53- 1.65) | 4.59E-01 | 6.42E-01 |
| 341 | PC(40:8) | 1.00 | ( 0.16- 1.92) | **1.88E-02** | **3.97E-02** | 0.64 | (-0.31- 1.69) | 1.96E-01 | 3.76E-01 | -1.34 | (-3.23- 0.58) | 1.69E-01 | 4.07E-01 | -0.28 | (-2.77- 2.28) | 8.27E-01 | 9.21E-01 |
| 342 | PC(44:12) | -1.20 | (-2.14--0.13) | **2.87E-02** | 5.89E-02 | -0.64 | (-1.81- 0.70) | 3.31E-01 | 5.28E-01 | -0.51 | (-4.52- 3.66) | 8.05E-01 | 9.16E-01 | -0.87 | (-5.51- 4.00) | 7.20E-01 | 8.50E-01 |
| 343 | PC(O-16:0/16:0) | 0.50 | (-0.35- 1.42) | 2.60E-01 | 3.86E-01 | 0.45 | (-0.53- 1.54) | 3.79E-01 | 5.75E-01 | -0.62 | (-2.10- 0.89) | 4.16E-01 | 6.76E-01 | -1.46 | (-3.27- 0.38) | 1.18E-01 | 2.83E-01 |
| 344 | PC(O-32:1) | 1.20 | ( 0.21- 2.28) | **1.64E-02** | **3.57E-02** | 0.22 | (-0.88- 1.45) | 7.10E-01 | 8.47E-01 | 0.19 | (-1.85- 2.27) | 8.56E-01 | 9.40E-01 | -0.57 | (-2.70- 1.59) | 5.99E-01 | 7.62E-01 |
| 345 | PC(O-34:1) | 1.05 | ( 0.18- 2.01) | **1.75E-02** | **3.77E-02** | 0.22 | (-0.78- 1.33) | 6.75E-01 | 8.33E-01 | -1.96 | (-3.30--0.60) | **5.04E-03** | **4.82E-02** | -2.29 | (-4.00--0.54) | **1.07E-02** | 6.90E-02 |
| 346 | PC(O-34:2) | -0.17 | (-1.05- 0.80) | 7.20E-01 | 7.86E-01 | 0.13 | (-1.02- 1.42) | 8.35E-01 | 9.32E-01 | -1.11 | (-3.41- 1.23) | 3.47E-01 | 6.22E-01 | -2.03 | (-4.76- 0.79) | 1.55E-01 | 3.40E-01 |
| 347 | PC(O-34:4) | 0.43 | (-0.54- 1.50) | 3.97E-01 | 5.27E-01 | -0.50 | (-1.45- 0.56) | 3.42E-01 | 5.43E-01 | 0.50 | (-3.15- 4.29) | 7.91E-01 | 9.16E-01 | 3.00 | (-1.17- 7.35) | 1.60E-01 | 3.41E-01 |
| 348 | PC(O-35:4) | -0.23 | (-1.06- 0.67) | 6.00E-01 | 7.06E-01 | -0.74 | (-1.72- 0.35) | 1.74E-01 | 3.43E-01 | 0.10 | (-3.29- 3.62) | 9.52E-01 | 9.83E-01 | 1.61 | (-2.69- 6.09) | 4.67E-01 | 6.50E-01 |
| 349 | PC(O-36:0) | 0.42 | (-0.64- 1.60) | 4.53E-01 | 5.79E-01 | -0.03 | (-1.28- 1.41) | 9.70E-01 | 9.87E-01 | -2.10 | (-3.83--0.34) | **1.97E-02** | 1.05E-01 | -2.53 | (-4.73--0.28) | **2.79E-02** | 1.19E-01 |
| 350 | PC(O-18:0/18:1) | 1.55 | ( 0.63- 2.54) | **7.13E-04** | **2.02E-03** | 1.06 | ( 0.01- 2.22) | **4.81E-02** | 1.28E-01 | -3.34 | (-4.86--1.79) | **3.82E-05** | **4.46E-03** | -2.26 | (-4.14--0.36) | **2.06E-02** | 1.05E-01 |
| 351 | PC(O-18:1/18:1) | 1.04 | ( 0.25- 1.90) | **9.51E-03** | **2.17E-02** | 0.26 | (-0.66- 1.27) | 5.93E-01 | 7.64E-01 | -1.48 | (-2.89--0.04) | **4.38E-02** | 1.91E-01 | -1.92 | (-3.64--0.17) | **3.20E-02** | 1.31E-01 |
| 352 | PC(O-18:0/18:2) | 0.61 | (-0.21- 1.50) | 1.51E-01 | 2.47E-01 | 0.26 | (-0.70- 1.32) | 6.13E-01 | 7.83E-01 | -1.98 | (-3.91--0.02) | **4.82E-02** | 1.98E-01 | -2.99 | (-5.34--0.59) | **1.54E-02** | 8.50E-02 |
| 353 | PC(O-18:1/18:2) | -0.04 | (-0.87- 0.86) | 9.25E-01 | 9.59E-01 | -0.07 | (-1.09- 1.07) | 9.02E-01 | 9.63E-01 | -0.65 | (-2.52- 1.25) | 4.99E-01 | 7.37E-01 | -1.92 | (-4.31- 0.53) | 1.23E-01 | 2.91E-01 |
| 354 | PC(O-16:0/20:3) | 0.28 | (-0.58- 1.21) | 5.36E-01 | 6.58E-01 | 0.66 | (-0.46- 1.90) | 2.59E-01 | 4.50E-01 | 0.68 | (-1.19- 2.59) | 4.76E-01 | 7.28E-01 | 1.95 | (-0.44- 4.40) | 1.10E-01 | 2.72E-01 |
| 355 | PC(O-16:0/20:4) | -0.18 | (-0.93- 0.63) | 6.49E-01 | 7.36E-01 | -0.04 | (-0.94- 0.96) | 9.37E-01 | 9.74E-01 | 0.58 | (-1.38- 2.58) | 5.63E-01 | 7.90E-01 | 0.96 | (-1.34- 3.32) | 4.15E-01 | 6.02E-01 |
| 356 | PC(O-36:5) | -1.35 | (-2.17--0.45) | **4.42E-03** | **1.10E-02** | -0.51 | (-1.62- 0.74) | 4.05E-01 | 6.00E-01 | 2.46 | (-1.21- 6.27) | 1.90E-01 | 4.31E-01 | 3.61 | (-0.71- 8.11) | 1.02E-01 | 2.63E-01 |
| 357 | PC(O-18:0/20:4) | -0.31 | (-1.01- 0.46) | 4.20E-01 | 5.53E-01 | -0.28 | (-1.10- 0.63) | 5.35E-01 | 7.24E-01 | -1.24 | (-3.20- 0.75) | 2.18E-01 | 4.66E-01 | -1.13 | (-3.29- 1.07) | 3.09E-01 | 5.08E-01 |
| 358 | PC(O-38:5) | -0.03 | (-0.89- 0.90) | 9.43E-01 | 9.74E-01 | -0.24 | (-1.30- 0.94) | 6.73E-01 | 8.33E-01 | 0.43 | (-1.39- 2.28) | 6.46E-01 | 8.52E-01 | 0.07 | (-2.04- 2.23) | 9.46E-01 | 9.75E-01 |
| 359 | PC(O-16:0/22:6) | -1.00 | (-1.77--0.16) | **2.02E-02** | **4.22E-02** | -0.42 | (-1.37- 0.63) | 4.15E-01 | 6.09E-01 | 1.60 | (-0.89- 4.16) | 2.08E-01 | 4.58E-01 | 0.86 | (-2.08- 3.89) | 5.67E-01 | 7.36E-01 |
| 360 | PC(O-40:5) | 0.01 | (-0.67- 0.74) | 9.79E-01 | 9.89E-01 | -0.37 | (-1.18- 0.51) | 4.00E-01 | 5.96E-01 | -0.29 | (-2.02- 1.48) | 7.47E-01 | 8.99E-01 | -1.20 | (-3.28- 0.92) | 2.62E-01 | 4.57E-01 |
| 361 | PC(O-18:0/22:6) | -1.03 | (-1.69--0.32) | **5.47E-03** | **1.35E-02** | -0.70 | (-1.51- 0.18) | 1.13E-01 | 2.46E-01 | -0.55 | (-2.51- 1.44) | 5.83E-01 | 8.05E-01 | -2.02 | (-4.34- 0.35) | 9.42E-02 | 2.49E-01 |
| 362 | PC(O-40:7) (a) | -0.64 | (-1.39- 0.17) | 1.19E-01 | 2.01E-01 | -0.78 | (-1.65- 0.19) | 1.10E-01 | 2.42E-01 | 0.74 | (-1.65- 3.18) | 5.47E-01 | 7.80E-01 | -0.84 | (-3.61- 2.01) | 5.55E-01 | 7.26E-01 |
| 363 | PC(P-16:0/14:0) | -0.22 | (-1.11- 0.76) | 6.52E-01 | 7.36E-01 | -0.42 | (-1.55- 0.86) | 5.01E-01 | 6.87E-01 | 0.13 | (-2.37- 2.70) | 9.18E-01 | 9.69E-01 | 0.13 | (-2.71- 3.06) | 9.28E-01 | 9.75E-01 |

| | | | | | | | | | | | | | | | | | |
|---|---|---|---|---|---|---|---|---|---|---|---|---|---|---|---|---|---|
| 364 | PC(P-16:0/16:0) | -0.52 | (-1.34- 0.38) | 2.45E-01 | 3.66E-01 | -0.43 | (-1.43- 0.69) | 4.40E-01 | 6.28E-01 | -0.86 | (-2.63- 0.95) | 3.47E-01 | 6.22E-01 | -1.52 | (-3.47- 0.47) | 1.32E-01 | 3.06E-01 |
| 365 | PC(P-16:0/16:1) | 0.15 | (-0.73- 1.13) | 7.42E-01 | 8.06E-01 | -0.21 | (-1.26- 0.97) | 7.15E-01 | 8.47E-01 | 1.14 | (-0.90- 3.22) | 2.73E-01 | 5.44E-01 | 0.76 | (-1.48- 3.04) | 5.08E-01 | 6.94E-01 |
| 366 | PC(P-16:0/18:0) | -0.30 | (-0.98- 0.44) | 4.20E-01 | 5.53E-01 | -0.05 | (-0.89- 0.85) | 9.03E-01 | 9.63E-01 | -2.07 | (-3.33--0.79) | **1.71E-03** | **2.82E-02** | -1.93 | (-3.50--0.32) | **1.93E-02** | 1.01E-01 |
| 367 | PC(P-16:0/18:1) | -0.19 | (-0.93- 0.62) | 6.41E-01 | 7.32E-01 | -0.32 | (-1.24- 0.70) | 5.22E-01 | 7.08E-01 | -1.85 | (-3.38--0.29) | **2.02E-02** | 1.07E-01 | -1.85 | (-3.75- 0.09) | 6.13E-02 | 1.88E-01 |
| 368 | PC(P-16:0/18:2) | -1.57 | (-2.30--0.77) | **2.57E-04** | **7.96E-04** | -0.83 | (-1.84- 0.31) | 1.45E-01 | 3.00E-01 | -0.81 | (-2.86- 1.28) | 4.43E-01 | 6.97E-01 | -1.85 | (-4.29- 0.66) | 1.47E-01 | 3.29E-01 |
| 369 | PC(P-16:0/18:3) | -0.54 | (-1.23- 0.21) | 1.55E-01 | 2.53E-01 | -0.35 | (-1.17- 0.55) | 4.32E-01 | 6.27E-01 | 0.37 | (-1.97- 2.75) | 7.60E-01 | 9.12E-01 | -0.76 | (-3.42- 1.97) | 5.78E-01 | 7.46E-01 |
| 370 | PC(P-35:2) (a) | -0.18 | (-1.06- 0.78) | 6.99E-01 | 7.74E-01 | -0.79 | (-1.84- 0.39) | 1.81E-01 | 3.53E-01 | -1.40 | (-4.65- 1.97) | 4.09E-01 | 6.75E-01 | -3.01 | (-7.03- 1.18) | 1.55E-01 | 3.40E-01 |
| 371 | PC(P-35:2) (b) | -1.75 | (-2.52--0.90) | **1.33E-04** | **4.34E-04** | -1.04 | (-2.03- 0.07) | 6.41E-02 | 1.60E-01 | -2.17 | (-4.70- 0.42) | 9.96E-02 | 2.86E-01 | -2.64 | (-5.78- 0.61) | 1.10E-01 | 2.72E-01 |
| 372 | PC(P-15:0/20:4) (b) | -0.96 | (-1.85- 0.03) | 5.62E-02 | 1.07E-01 | -0.31 | (-1.45- 0.99) | 6.27E-01 | 7.96E-01 | 0.16 | (-2.68- 3.08) | 9.11E-01 | 9.68E-01 | 1.40 | (-1.99- 4.90) | 4.21E-01 | 6.04E-01 |
| 373 | PC(P-18:1/18:1) | 1.43 | ( 0.35- 2.63) | **8.79E-03** | **2.04E-02** | 0.97 | (-0.31- 2.42) | 1.45E-01 | 3.00E-01 | -0.57 | (-2.19- 1.08) | 4.92E-01 | 7.35E-01 | 0.21 | (-1.67- 2.13) | 8.24E-01 | 9.21E-01 |
| 374 | PC(P-18:0/18:2) | -0.60 | (-1.28- 0.14) | 1.07E-01 | 1.84E-01 | -0.67 | (-1.55- 0.30) | 1.67E-01 | 3.35E-01 | -2.43 | (-4.45--0.37) | **2.12E-02** | 1.10E-01 | -3.16 | (-5.41--0.85) | **7.88E-03** | 6.01E-02 |
| 375 | PC(P-16:0/20:4) | -1.23 | (-1.93--0.46) | **2.23E-03** | **5.95E-03** | -0.83 | (-1.73- 0.17) | 9.99E-02 | 2.25E-01 | 0.38 | (-1.65- 2.45) | 7.13E-01 | 8.85E-01 | 0.56 | (-1.91- 3.10) | 6.57E-01 | 8.10E-01 |
| 376 | PC(P-16:0/20:5) | -2.29 | (-3.14--1.33) | **1.72E-05** | **6.30E-05** | -1.43 | (-2.56--0.13) | **3.29E-02** | 9.22E-02 | 3.50 | (-0.16- 7.30) | 6.11E-02 | 2.20E-01 | 3.19 | (-1.51- 8.11) | 1.85E-01 | 3.70E-01 |
| 377 | PC(P-17:0/20:4) (a) | -1.29 | (-2.15--0.32) | **1.04E-02** | **2.36E-02** | -1.79 | (-2.78--0.66) | **2.76E-03** | **1.16E-02** | -0.74 | (-3.98- 2.60) | 6.57E-01 | 8.54E-01 | -0.35 | (-4.60- 4.09) | 8.75E-01 | 9.63E-01 |
| 378 | PC(P-17:0/20:4) (b) | -2.12 | (-2.86--1.31) | **2.76E-06** | **1.10E-05** | -1.46 | (-2.40--0.41) | **7.72E-03** | **2.83E-02** | -1.20 | (-3.85- 1.54) | 3.85E-01 | 6.55E-01 | -0.10 | (-3.46- 3.38) | 9.55E-01 | 9.75E-01 |
| 379 | PC(P-18:0/20:4) | -1.18 | (-1.84--0.48) | **1.45E-03** | **3.91E-03** | -0.88 | (-1.68- 0.01) | 5.25E-02 | 1.38E-01 | -0.99 | (-3.10- 1.16) | 3.62E-01 | 6.36E-01 | -0.03 | (-2.62- 2.64) | 9.84E-01 | 9.91E-01 |
| 380 | PC(P-38:5) (a) | -1.28 | (-1.98--0.52) | **1.39E-03** | **3.78E-03** | -1.06 | (-1.92--0.10) | **3.11E-02** | 8.83E-02 | 0.64 | (-1.40- 2.73) | 5.38E-01 | 7.71E-01 | 1.49 | (-1.04- 4.08) | 2.49E-01 | 4.40E-01 |
| 381 | PC(P-38:5) (b) | -0.85 | (-1.67- 0.04) | 6.12E-02 | 1.15E-01 | -0.77 | (-1.74- 0.31) | 1.54E-01 | 3.17E-01 | -0.39 | (-2.60- 1.87) | 7.31E-01 | 8.86E-01 | -0.02 | (-2.65- 2.68) | 9.89E-01 | 9.91E-01 |
| 382 | PC(P-16:0/22:6) | -1.66 | (-2.39--0.87) | **9.95E-05** | **3.34E-04** | -1.21 | (-2.13--0.19) | **2.17E-02** | 6.63E-02 | 0.84 | (-1.69- 3.44) | 5.18E-01 | 7.51E-01 | -0.10 | (-3.19- 3.08) | 9.47E-01 | 9.75E-01 |
| 383 | PC(P-20:0/20:4) | -1.01 | (-1.76--0.19) | **1.72E-02** | **3.71E-02** | -0.60 | (-1.54- 0.45) | 2.49E-01 | 4.38E-01 | -1.46 | (-3.34- 0.46) | 1.35E-01 | 3.48E-01 | -2.02 | (-4.42- 0.44) | 1.06E-01 | 2.68E-01 |
| 384 | PC(P-40:5) (b) | -0.93 | (-1.59--0.23) | **1.07E-02** | **2.42E-02** | -0.55 | (-1.36- 0.35) | 2.24E-01 | 4.09E-01 | -1.16 | (-3.01- 0.73) | 2.27E-01 | 4.75E-01 | -1.23 | (-3.35- 0.93) | 2.59E-01 | 4.51E-01 |
| 385 | PC(P-18:0/22:6) | -1.73 | (-2.38--1.03) | **6.54E-06** | **2.50E-05** | -1.24 | (-2.06--0.33) | **8.94E-03** | **3.15E-02** | -1.02 | (-3.45- 1.47) | 4.16E-01 | 6.76E-01 | -2.19 | (-5.18- 0.88) | 1.59E-01 | 3.41E-01 |
| 386 | PE(16:0_16:1) | 0.79 | (-0.57- 2.35) | 2.67E-01 | 3.95E-01 | 0.10 | (-1.39- 1.84) | 9.03E-01 | 9.63E-01 | 4.60 | (-0.05- 9.48) | 5.25E-02 | 2.08E-01 | 4.25 | (-1.08- 9.86) | 1.19E-01 | 2.83E-01 |
| 387 | PE(16:0_18:1) | -0.45 | (-1.29- 0.47) | 3.22E-01 | 4.53E-01 | -0.64 | (-1.79- 0.67) | 3.23E-01 | 5.18E-01 | 0.33 | (-2.93- 3.71) | 8.43E-01 | 9.40E-01 | 0.13 | (-4.04- 4.47) | 9.54E-01 | 9.75E-01 |
| 388 | PE(16:0_18:2) | -1.02 | (-1.89--0.05) | **3.98E-02** | 7.73E-02 | -0.08 | (-1.29- 1.29) | 8.97E-01 | 9.63E-01 | -0.28 | (-3.93- 3.51) | 8.82E-01 | 9.49E-01 | -2.10 | (-6.28- 2.26) | 3.37E-01 | 5.37E-01 |
| 389 | PE(16:1_18:2) | 0.47 | (-0.56- 1.61) | 3.84E-01 | 5.13E-01 | 0.32 | (-0.87- 1.66) | 6.12E-01 | 7.83E-01 | 0.31 | (-2.73- 3.43) | 8.45E-01 | 9.40E-01 | -0.52 | (-4.03- 3.13) | 7.76E-01 | 8.97E-01 |
| 390 | PE(16:0_18:3) (a) | 0.26 | (-0.79- 1.42) | 6.46E-01 | 7.36E-01 | -0.14 | (-1.30- 1.16) | 8.18E-01 | 9.20E-01 | -0.70 | (-4.39- 3.13) | 7.15E-01 | 8.85E-01 | -0.85 | (-5.27- 3.77) | 7.11E-01 | 8.49E-01 |
| 391 | PE(16:0_18:3) (b) | 0.43 | (-0.64- 1.64) | 4.43E-01 | 5.70E-01 | 0.90 | (-0.50- 2.52) | 2.20E-01 | 4.08E-01 | 1.62 | (-2.42- 5.83) | 4.36E-01 | 6.92E-01 | 2.25 | (-2.68- 7.43) | 3.75E-01 | 5.66E-01 |
| 392 | PE(17:0_18:2) | -0.71 | (-1.41- 0.04) | 6.27E-02 | 1.17E-01 | -1.00 | (-1.74--0.20) | **1.57E-02** | 5.09E-02 | -1.54 | (-4.48- 1.50) | 3.15E-01 | 5.90E-01 | -0.65 | (-4.27- 3.11) | 7.31E-01 | 8.58E-01 |
| 393 | PE(18:0_18:1) | -0.15 | (-1.11- 0.91) | 7.66E-01 | 8.26E-01 | 0.04 | (-1.23- 1.48) | 9.57E-01 | 9.85E-01 | 2.46 | (-0.64- 5.67) | 1.21E-01 | 3.31E-01 | 2.72 | (-1.38- 6.99) | 1.95E-01 | 3.79E-01 |
| 394 | PE(18:1_18:1) | 0.51 | (-0.94- 2.19) | 5.12E-01 | 6.35E-01 | -0.76 | (-2.24- 1.00) | 3.73E-01 | 5.75E-01 | 0.25 | (-2.69- 3.28) | 8.68E-01 | 9.44E-01 | -0.29 | (-4.08- 3.66) | 8.84E-01 | 9.67E-01 |
| 395 | PE(18:0_18:2) | -0.99 | (-1.81--0.09) | **3.15E-02** | 6.35E-02 | 0.27 | (-0.86- 1.55) | 6.50E-01 | 8.15E-01 | 1.99 | (-1.21- 5.30) | 2.25E-01 | 4.75E-01 | 0.21 | (-3.66- 4.24) | 9.16E-01 | 9.75E-01 |
| 396 | PE(18:1_18:2) | -0.38 | (-1.38- 0.74) | 4.92E-01 | 6.17E-01 | -0.10 | (-1.25- 1.20) | 8.73E-01 | 9.49E-01 | -1.86 | (-5.14- 1.53) | 2.76E-01 | 5.46E-01 | -2.88 | (-7.07- 1.50) | 1.93E-01 | 3.76E-01 |
| 397 | PE(16:0_20:3) | -0.36 | (-1.29- 0.67) | 4.82E-01 | 6.07E-01 | -0.07 | (-1.29- 1.32) | 9.17E-01 | 9.70E-01 | 0.39 | (-2.93- 3.82) | 8.20E-01 | 9.30E-01 | -0.99 | (-5.22- 3.42) | 6.51E-01 | 8.08E-01 |
| 398 | PE(16:0_20:4) | -0.38 | (-1.16- 0.46) | 3.60E-01 | 4.87E-01 | -0.10 | (-1.13- 1.06) | 8.64E-01 | 9.47E-01 | -0.96 | (-4.10- 2.27) | 5.53E-01 | 7.83E-01 | -2.43 | (-6.13- 1.42) | 2.11E-01 | 4.01E-01 |
| 399 | PE(16:1_20:4) | 0.97 | (-0.10- 2.15) | 7.71E-02 | 1.41E-01 | 0.33 | (-0.91- 1.74) | 6.20E-01 | 7.90E-01 | 1.50 | (-1.26- 4.33) | 2.88E-01 | 5.55E-01 | 0.27 | (-2.93- 3.58) | 8.69E-01 | 9.59E-01 |
| 400 | PE(16:0_20:5) | -0.90 | (-1.87- 0.18) | 9.99E-02 | 1.74E-01 | -0.71 | (-1.91- 0.68) | 3.00E-01 | 4.96E-01 | 2.40 | (-1.13- 6.06) | 1.84E-01 | 4.24E-01 | 0.83 | (-3.33- 5.17) | 6.99E-01 | 8.47E-01 |
| 401 | PE(17:0_20:4) | -1.83 | (-2.56--1.02) | **3.39E-05** | **1.20E-04** | -1.32 | (-2.30--0.23) | **1.97E-02** | 6.11E-02 | -1.06 | (-3.85- 1.81) | 4.64E-01 | 7.18E-01 | -1.91 | (-5.34- 1.64) | 2.84E-01 | 4.81E-01 |
| 402 | PE(18:0_20:3) (a) | -0.67 | (-1.50- 0.24) | 1.45E-01 | 2.40E-01 | -0.43 | (-1.44- 0.71) | 4.44E-01 | 6.30E-01 | 3.27 | ( 0.31- 6.33) | **3.03E-02** | 1.43E-01 | 4.21 | ( 0.20- 8.38) | **3.97E-02** | 1.49E-01 |
| 403 | PE(18:0_20:4) | -0.88 | (-1.60--0.10) | **2.71E-02** | 5.58E-02 | -0.29 | (-1.29- 0.83) | 5.94E-01 | 7.64E-01 | 0.78 | (-1.83- 3.47) | 5.60E-01 | 7.88E-01 | 0.12 | (-3.17- 3.52) | 9.45E-01 | 9.75E-01 |
| 404 | PE(38:5) (a) | -0.56 | (-1.35- 0.31) | 1.98E-01 | 3.10E-01 | -0.58 | (-1.59- 0.56) | 3.03E-01 | 4.98E-01 | -1.44 | (-4.36- 1.57) | 3.43E-01 | 6.22E-01 | -1.69 | (-5.42- 2.19) | 3.87E-01 | 5.75E-01 |
| 405 | PE(38:5) (b) | 0.37 | (-0.91- 1.83) | 5.91E-01 | 7.01E-01 | 0.64 | (-1.02- 2.60) | 4.71E-01 | 6.59E-01 | 1.89 | (-1.70- 5.62) | 3.04E-01 | 5.75E-01 | -0.63 | (-5.08- 4.03) | 7.87E-01 | 8.98E-01 |
| 406 | PE(16:0_22:6) | -1.91 | (-2.69--1.05) | **4.47E-05** | **1.55E-04** | -0.94 | (-2.05- 0.32) | 1.36E-01 | 2.85E-01 | 0.66 | (-3.01- 4.47) | 7.26E-01 | 8.86E-01 | -2.20 | (-6.35- 2.14) | 3.13E-01 | 5.12E-01 |
| 407 | PE(17:0_22:6) | -1.98 | (-2.73--1.15) | **1.42E-05** | **5.32E-05** | -0.85 | (-1.77- 0.18) | 1.03E-01 | 2.28E-01 | 0.36 | (-2.40- 3.19) | 8.02E-01 | 9.16E-01 | -0.67 | (-4.13- 2.91) | 7.08E-01 | 8.49E-01 |
| 408 | PE(18:0_22:4) | -0.28 | (-1.26- 0.80) | 5.95E-01 | 7.04E-01 | 0.09 | (-1.24- 1.62) | 8.99E-01 | 9.63E-01 | 2.66 | (-0.04- 5.42) | 5.34E-02 | 2.10E-01 | 1.85 | (-1.70- 5.52) | 3.08E-01 | 5.08E-01 |
| 409 | PE(18:0_22:5) (n3) | -1.83 | (-2.72--0.83) | **6.62E-04** | **1.93E-03** | 0.21 | (-1.17- 1.82) | 7.76E-01 | 8.91E-01 | 2.74 | (-0.41- 5.99) | 8.81E-02 | 2.66E-01 | 2.13 | (-1.43- 5.82) | 2.42E-01 | 4.32E-01 |
| 410 | PE(18:0_22:5) (n6) | -0.53 | (-1.72- 0.84) | 4.28E-01 | 5.57E-01 | -0.19 | (-1.72- 1.62) | 8.24E-01 | 9.22E-01 | 2.97 | (-0.29- 6.33) | 7.46E-02 | 2.45E-01 | 2.09 | (-2.17- 6.54) | 3.39E-01 | 5.39E-01 |
| 411 | PE(18:0_22:6) | -2.15 | (-3.01--1.18) | **5.23E-05** | **1.81E-04** | -0.65 | (-1.96- 0.87) | 3.78E-01 | 5.75E-01 | 5.09 | ( 1.50- 8.81) | **5.41E-03** | **4.82E-02** | 1.93 | (-2.24- 6.29) | 3.66E-01 | 5.64E-01 |
| 412 | PE(40:7) | -1.27 | (-2.08--0.38) | **6.54E-03** | **1.58E-02** | -1.18 | (-2.25- 0.04) | 5.83E-02 | 1.51E-01 | -0.15 | (-3.95- 3.80) | 9.40E-01 | 9.79E-01 | -2.69 | (-6.91- 1.73) | 2.28E-01 | 4.25E-01 |
| 413 | PE(O-34:1) | 0.79 | (-0.74- 2.57) | 3.30E-01 | 4.63E-01 | -0.03 | (-1.61- 1.85) | 9.72E-01 | 9.87E-01 | -1.06 | (-3.39- 1.34) | 3.81E-01 | 6.53E-01 | -1.66 | (-4.16- 0.90) | 2.00E-01 | 3.86E-01 |
| 414 | PE(O-16:0/18:2) | 0.71 | (-1.89- 4.14) | 6.29E-01 | 7.25E-01 | -1.25 | (-3.52- 1.82) | 3.82E-01 | 5.79E-01 | 0.01 | (-2.86- 2.98) | 9.92E-01 | 9.99E-01 | -0.32 | (-3.72- 3.21) | 8.56E-01 | 9.46E-01 |
| 415 | PE(O-18:1/18:2) | 0.01 | (-1.19- 1.38) | 9.88E-01 | 9.94E-01 | 0.40 | (-1.06- 2.09) | 6.10E-01 | 7.83E-01 | -0.25 | (-2.78- 2.35) | 8.50E-01 | 9.40E-01 | -0.08 | (-3.36- 3.31) | 9.61E-01 | 9.77E-01 |
| 416 | PE(O-16:0/20:4) | -0.90 | (-1.91- 0.22) | 1.11E-01 | 1.90E-01 | -0.59 | (-1.88- 0.90) | 4.14E-01 | 6.09E-01 | 0.96 | (-1.80- 3.80) | 4.96E-01 | 7.35E-01 | 1.90 | (-1.53- 5.44) | 2.79E-01 | 4.80E-01 |

| | | | | | | | | | | | | | | | | | |
|---|---|---|---|---|---|---|---|---|---|---|---|---|---|---|---|---|---|
| 417 | PE(O-36:5) | 0.01 | (-0.91- 1.01) | 9.92E-01 | 9.94E-01 | -0.85 | (-1.81- 0.21) | 1.12E-01 | 2.46E-01 | 2.66 | (-0.91- 6.35) | 1.45E-01 | 3.62E-01 | 2.08 | (-2.32- 6.67) | 3.58E-01 | 5.55E-01 |
| 418 | PE(O-16:0/22:4) | 0.38 | (-0.55- 1.39) | 4.38E-01 | 5.67E-01 | 0.93 | (-0.28- 2.29) | 1.38E-01 | 2.88E-01 | 0.12 | (-2.09- 2.37) | 9.17E-01 | 9.69E-01 | 1.03 | (-1.48- 3.61) | 4.21E-01 | 6.04E-01 |
| 419 | PE(O-18:0/20:4) | -0.07 | (-1.13- 1.12) | 9.01E-01 | 9.42E-01 | -0.18 | (-1.33- 1.12) | 7.72E-01 | 8.89E-01 | 0.55 | (-2.54- 3.75) | 7.29E-01 | 8.86E-01 | 0.47 | (-3.42- 4.52) | 8.14E-01 | 9.17E-01 |
| 420 | PE(O-38:5) (a) | -0.22 | (-1.04- 0.68) | 6.21E-01 | 7.22E-01 | 0.18 | (-0.97- 1.48) | 7.67E-01 | 8.85E-01 | 1.87 | (-1.06- 4.88) | 2.11E-01 | 4.61E-01 | 1.50 | (-2.01- 5.14) | 4.05E-01 | 5.96E-01 |
| 421 | PE(O-38:5) (b) | 0.13 | (-1.02- 1.42) | 8.35E-01 | 8.90E-01 | 0.33 | (-1.14- 2.03) | 6.80E-01 | 8.36E-01 | 0.01 | (-2.45- 2.53) | 9.96E-01 | 9.99E-01 | 1.01 | (-2.04- 4.14) | 5.18E-01 | 6.99E-01 |
| 422 | PE(O-16:0/22:6) | 0.42 | (-0.58- 1.53) | 4.23E-01 | 5.55E-01 | 0.75 | (-0.45- 2.09) | 2.29E-01 | 4.13E-01 | 0.63 | (-1.70- 3.02) | 5.96E-01 | 8.10E-01 | 1.25 | (-1.57- 4.14) | 3.87E-01 | 5.75E-01 |
| 423 | PE(O-18:0/22:6) | 0.07 | (-0.76- 0.97) | 8.76E-01 | 9.20E-01 | -0.08 | (-1.02- 0.96) | 8.72E-01 | 9.49E-01 | -0.06 | (-2.41- 2.36) | 9.63E-01 | 9.89E-01 | 0.11 | (-2.69- 2.99) | 9.37E-01 | 9.75E-01 |
| 424 | PE(O-18:1/22:6) | 0.14 | (-0.60- 0.95) | 7.11E-01 | 7.81E-01 | -0.02 | (-0.89- 0.93) | 9.60E-01 | 9.85E-01 | 0.35 | (-2.00- 2.75) | 7.74E-01 | 9.15E-01 | 0.88 | (-2.07- 3.92) | 5.59E-01 | 7.30E-01 |
| 425 | PE(P-16:0/18:1) | 1.66 | ( 0.80- 2.59) | **1.05E-04** | **3.49E-04** | 1.93 | ( 0.75- 3.24) | **9.42E-04** | **4.66E-03** | 0.01 | (-1.84- 1.90) | 9.89E-01 | 9.99E-01 | 0.14 | (-2.05- 2.39) | 8.99E-01 | 9.70E-01 |
| 426 | PE(P-16:0/18:2) | -0.25 | (-1.28- 0.91) | 6.61E-01 | 7.44E-01 | 1.02 | (-0.50- 2.78) | 1.98E-01 | 3.76E-01 | 0.74 | (-1.96- 3.52) | 5.93E-01 | 8.10E-01 | 0.95 | (-1.86- 3.85) | 5.09E-01 | 6.94E-01 |
| 427 | PE(P-16:0/20:3) | 0.50 | (-0.57- 1.70) | 3.73E-01 | 5.01E-01 | -0.19 | (-1.48- 1.30) | 7.89E-01 | 9.01E-01 | 3.59 | ( 1.12- 6.12) | **4.43E-03** | **4.82E-02** | 5.78 | ( 2.90- 8.75) | **9.29E-05** | **5.07E-03** |
| 428 | PE(P-16:0/20:4) | -0.26 | (-1.20- 0.79) | 6.16E-01 | 7.18E-01 | 0.05 | (-1.18- 1.44) | 9.44E-01 | 9.77E-01 | 3.59 | ( 0.88- 6.37) | **9.38E-03** | 6.43E-02 | 3.85 | ( 0.58- 7.23) | **2.10E-02** | 1.05E-01 |
| 429 | PE(P-16:0/20:5) | -1.60 | (-3.57- 0.99) | 2.03E-01 | 3.16E-01 | 0.39 | (-2.49- 4.38) | 8.17E-01 | 9.20E-01 | 4.73 | ( 0.09- 9.58) | **4.55E-02** | 1.95E-01 | 6.37 | ( 0.53-12.55) | **3.23E-02** | 1.31E-01 |
| 430 | PE(P-16:0/22:4) | 0.00 | (-0.85- 0.92) | 9.94E-01 | 9.94E-01 | 0.64 | (-0.55- 1.99) | 3.04E-01 | 4.98E-01 | -0.42 | (-2.25- 1.44) | 6.51E-01 | 8.54E-01 | -0.13 | (-2.28- 2.06) | 9.06E-01 | 9.71E-01 |
| 431 | PE(P-16:0/22:5) (n3) | -0.80 | (-1.66- 0.16) | 1.00E-01 | 1.74E-01 | 0.02 | (-1.17- 1.37) | 9.77E-01 | 9.89E-01 | 1.33 | (-1.08- 3.79) | 2.80E-01 | 5.49E-01 | 1.88 | (-0.66- 4.48) | 1.48E-01 | 3.30E-01 |
| 432 | PE(P-16:0/22:5) (n6) | 0.83 | (-0.20- 1.97) | 1.17E-01 | 1.98E-01 | 1.20 | (-0.11- 2.67) | 7.31E-02 | 1.75E-01 | 0.48 | (-1.92- 2.93) | 6.99E-01 | 8.78E-01 | 0.38 | (-2.79- 3.64) | 8.17E-01 | 9.18E-01 |
| 433 | PE(P-16:0/22:6) | 0.10 | (-0.76- 1.03) | 8.33E-01 | 8.90E-01 | 0.33 | (-0.73- 1.51) | 5.59E-01 | 7.37E-01 | 1.78 | (-0.26- 3.87) | 8.76E-02 | 2.66E-01 | 2.09 | (-0.51- 4.76) | 1.16E-01 | 2.80E-01 |
| 434 | PE(P-17:0/20:4) (a) | 0.00 | (-0.79- 0.85) | 9.93E-01 | 9.94E-01 | -0.81 | (-1.69- 0.16) | 9.79E-02 | 2.23E-01 | 1.04 | (-1.91- 4.07) | 4.92E-01 | 7.35E-01 | 2.47 | (-1.39- 6.47) | 2.11E-01 | 4.01E-01 |
| 435 | PE(P-17:0/20:4) (b) | -0.88 | (-1.67--0.01) | **4.77E-02** | 9.16E-02 | -0.93 | (-1.87- 0.13) | 8.27E-02 | 1.92E-01 | 0.60 | (-2.14- 3.41) | 6.71E-01 | 8.54E-01 | 2.56 | (-0.82- 6.05) | 1.38E-01 | 3.14E-01 |
| 436 | PE(P-17:0/22:6) (a) | 0.21 | (-0.82- 1.35) | 7.04E-01 | 7.75E-01 | -0.67 | (-1.78- 0.59) | 2.81E-01 | 4.75E-01 | -0.06 | (-2.93- 2.91) | 9.70E-01 | 9.93E-01 | 0.58 | (-3.42- 4.74) | 7.79E-01 | 8.97E-01 |
| 437 | PE(P-17:0/22:6) (b) | -0.45 | (-1.31- 0.49) | 3.35E-01 | 4.66E-01 | -0.44 | (-1.45- 0.70) | 4.35E-01 | 6.27E-01 | -0.78 | (-3.27- 1.78) | 5.45E-01 | 7.79E-01 | 0.75 | (-2.50- 4.11) | 6.53E-01 | 8.08E-01 |
| 438 | PE(P-18:0/18:1) | 1.12 | ( 0.18- 2.15) | **1.87E-02** | **3.97E-02** | 0.90 | (-0.38- 2.35) | 1.74E-01 | 3.43E-01 | -0.49 | (-2.61- 1.68) | 6.54E-01 | 8.54E-01 | -0.45 | (-2.80- 1.96) | 7.13E-01 | 8.49E-01 |
| 439 | PE(P-18:0/18:2) | -0.21 | (-1.33- 1.07) | 7.39E-01 | 8.04E-01 | 0.01 | (-1.27- 1.48) | 9.87E-01 | 9.94E-01 | 0.06 | (-2.65- 2.85) | 9.65E-01 | 9.90E-01 | -1.23 | (-3.78- 1.38) | 3.49E-01 | 5.44E-01 |
| 440 | PE(P-18:0/20:3) | -0.23 | (-0.93- 0.53) | 5.41E-01 | 6.61E-01 | -0.36 | (-1.26- 0.65) | 4.71E-01 | 6.59E-01 | 2.17 | ( 0.07- 4.32) | **4.30E-02** | 1.90E-01 | 2.87 | ( 0.42- 5.38) | **2.15E-02** | 1.06E-01 |
| 441 | PE(P-18:0/20:4) | -0.94 | (-1.74--0.05) | **3.86E-02** | 7.57E-02 | -0.04 | (-1.20- 1.26) | 9.44E-01 | 9.77E-01 | 1.77 | (-0.98- 4.60) | 2.08E-01 | 4.58E-01 | 1.93 | (-1.25- 5.22) | 2.34E-01 | 4.31E-01 |
| 442 | PE(P-18:0/20:5) | -1.78 | (-3.49- 0.38) | 9.86E-02 | 1.73E-01 | -2.61 | (-4.50--0.08) | **4.40E-02** | 1.19E-01 | 3.74 | (-0.66- 8.34) | 9.63E-02 | 2.80E-01 | 4.66 | (-1.03-10.67) | 1.10E-01 | 2.72E-01 |
| 443 | PE(P-18:0/22:4) | -0.27 | (-1.20- 0.77) | 5.97E-01 | 7.05E-01 | -0.11 | (-1.36- 1.32) | 8.71E-01 | 9.49E-01 | -2.03 | (-3.90--0.13) | **3.69E-02** | 1.64E-01 | -1.02 | (-3.10- 1.11) | 3.44E-01 | 5.41E-01 |
| 444 | PE(P-18:0/22:5) (n3) | -0.49 | (-1.25- 0.33) | 2.35E-01 | 3.53E-01 | -0.16 | (-1.14- 0.93) | 7.63E-01 | 8.85E-01 | 0.33 | (-1.82- 2.53) | 7.64E-01 | 9.12E-01 | 1.53 | (-0.94- 4.05) | 2.25E-01 | 4.23E-01 |
| 445 | PE(P-18:0/22:5) (n6) | 0.64 | (-0.18- 1.54) | 1.30E-01 | 2.18E-01 | 0.04 | (-0.93- 1.12) | 9.35E-01 | 9.74E-01 | -0.23 | (-2.75- 2.35) | 8.59E-01 | 9.40E-01 | -0.79 | (-3.81- 2.33) | 6.15E-01 | 7.74E-01 |
| 446 | PE(P-18:0/22:6) | -0.15 | (-0.92- 0.69) | 7.20E-01 | 7.86E-01 | 0.22 | (-0.84- 1.40) | 7.02E-01 | 8.47E-01 | 0.34 | (-1.95- 2.69) | 7.70E-01 | 9.12E-01 | -0.12 | (-3.02- 2.87) | 9.37E-01 | 9.75E-01 |
| 447 | PE(P-18:1/18:1) | 0.19 | (-0.59- 1.03) | 6.40E-01 | 7.32E-01 | 0.64 | (-0.37- 1.75) | 2.22E-01 | 4.08E-01 | -0.33 | (-2.23- 1.61) | 7.35E-01 | 8.86E-01 | -0.08 | (-2.25- 2.14) | 9.45E-01 | 9.75E-01 |
| 448 | PE(P-18:1/18:2) | -0.56 | (-2.03- 1.17) | 4.98E-01 | 6.23E-01 | -0.93 | (-2.49- 0.94) | 3.05E-01 | 4.98E-01 | 0.13 | (-2.62- 2.96) | 9.25E-01 | 9.71E-01 | -1.32 | (-4.23- 1.68) | 3.83E-01 | 5.75E-01 |
| 449 | PE(P-18:1/20:4) | -0.39 | (-1.23- 0.53) | 3.93E-01 | 5.24E-01 | -0.65 | (-1.76- 0.61) | 2.95E-01 | 4.89E-01 | 2.07 | (-0.63- 4.85) | 1.33E-01 | 3.48E-01 | 2.43 | (-0.92- 5.90) | 1.56E-01 | 3.40E-01 |
| 450 | PE(P-18:1/20:5) | -0.76 | (-3.25- 2.65) | 6.22E-01 | 7.22E-01 | -0.79 | (-3.72- 3.50) | 6.71E-01 | 8.33E-01 | 3.61 | (-1.12- 8.58) | 1.36E-01 | 3.49E-01 | 5.56 | (-0.22-11.68) | 5.93E-02 | 1.87E-01 |
| 451 | PE(P-18:1/22:4) | 0.36 | (-0.76- 1.62) | 5.42E-01 | 6.61E-01 | 0.50 | (-0.91- 2.14) | 5.05E-01 | 6.90E-01 | -1.12 | (-3.01- 0.81) | 2.51E-01 | 5.11E-01 | -0.94 | (-3.14- 1.31) | 4.08E-01 | 5.97E-01 |
| 452 | PE(P-18:1/22:5) (a) | -0.78 | (-1.53- 0.04) | 6.26E-02 | 1.17E-01 | 0.34 | (-0.73- 1.54) | 5.51E-01 | 7.36E-01 | 0.91 | (-1.26- 3.13) | 4.11E-01 | 6.75E-01 | 1.77 | (-1.11- 4.73) | 2.30E-01 | 4.27E-01 |
| 453 | PE(P-18:1/22:6) (a) | -0.06 | (-0.93- 0.90) | 9.03E-01 | 9.42E-01 | -0.06 | (-1.11- 1.12) | 9.21E-01 | 9.70E-01 | 1.17 | (-1.22- 3.62) | 3.36E-01 | 6.14E-01 | 0.93 | (-2.13- 4.09) | 5.52E-01 | 7.24E-01 |
| 454 | PE(P-20:0/18:1) | 0.93 | (-1.24- 3.64) | 4.28E-01 | 5.57E-01 | 0.11 | (-2.22- 3.14) | 9.36E-01 | 9.74E-01 | -0.45 | (-2.94- 2.10) | 7.24E-01 | 8.86E-01 | -2.01 | (-4.37- 0.42) | 1.03E-01 | 2.64E-01 |
| 455 | PE(P-20:0/18:2) | 0.18 | (-0.40- 0.80) | 5.41E-01 | 6.61E-01 | -0.35 | (-0.99- 0.34) | 3.14E-01 | 5.08E-01 | -2.04 | (-4.27- 0.24) | 7.90E-02 | 2.50E-01 | -3.60 | (-5.89--1.25) | **3.06E-03** | **3.26E-02** |
| 456 | PE(P-20:0/20:4) | -0.37 | (-1.48- 0.88) | 5.46E-01 | 6.63E-01 | -0.76 | (-2.09- 0.78) | 3.14E-01 | 5.08E-01 | 0.02 | (-2.11- 2.19) | 9.85E-01 | 9.99E-01 | -0.42 | (-2.89- 2.11) | 7.43E-01 | 8.69E-01 |
| 457 | PE(P-20:0/22:6) | -0.58 | (-1.31- 0.21) | 1.46E-01 | 2.41E-01 | -0.75 | (-1.60- 0.19) | 1.14E-01 | 2.46E-01 | -0.94 | (-2.82- 0.98) | 3.33E-01 | 6.14E-01 | -2.82 | (-5.23--0.34) | **2.62E-02** | 1.18E-01 |
| 458 | PE(P-20:1/20:4) | -0.65 | (-1.86- 0.75) | 3.44E-01 | 4.74E-01 | -0.72 | (-2.42- 1.35) | 4.65E-01 | 6.55E-01 | 1.61 | (-1.14- 4.43) | 2.53E-01 | 5.13E-01 | 0.42 | (-3.19- 4.17) | 8.21E-01 | 9.21E-01 |
| 459 | PG(34:2) | -1.94 | (-3.27--0.34) | **2.00E-02** | **4.21E-02** | 1.64 | (-0.60- 4.42) | 1.64E-01 | 3.31E-01 | 1.09 | (-2.53- 4.83) | 5.59E-01 | 7.88E-01 | 3.09 | (-1.33- 7.70) | 1.72E-01 | 3.57E-01 |
| 460 | PG(36:1) | -1.48 | (-2.82- 0.11) | 6.66E-02 | 1.23E-01 | -1.21 | (-3.06- 1.12) | 2.82E-01 | 4.75E-01 | 1.93 | (-1.44- 5.41) | 2.63E-01 | 5.28E-01 | 3.92 | (-0.38- 8.40) | 7.44E-02 | 2.11E-01 |
| 461 | PI(16:0/16:0) | -0.39 | (-1.45- 0.82) | 5.12E-01 | 6.35E-01 | 0.49 | (-1.14- 2.43) | 5.76E-01 | 7.52E-01 | 3.35 | (-0.03- 6.83) | 5.17E-02 | 2.08E-01 | 2.54 | (-1.69- 6.94) | 2.42E-01 | 4.32E-01 |
| 462 | PI(16:0_16:1) | 0.67 | (-0.36- 1.82) | 2.09E-01 | 3.23E-01 | 0.86 | (-0.50- 2.41) | 2.25E-01 | 4.09E-01 | 8.10 | ( 4.43-11.90) | **1.52E-05** | **2.43E-03** | 9.16 | ( 4.55-13.96) | **9.51E-05** | **5.07E-03** |
| 463 | PI(34:0) | -1.69 | (-3.19- 0.12) | 6.60E-02 | 1.23E-01 | -2.02 | (-3.70- 0.11) | 6.18E-02 | 1.56E-01 | 1.71 | (-2.41- 5.99) | 4.19E-01 | 6.78E-01 | -2.77 | (-7.20- 1.87) | 2.36E-01 | 4.31E-01 |
| 464 | PI(34:1) | -0.15 | (-0.92- 0.69) | 7.20E-01 | 7.86E-01 | -0.23 | (-1.24- 0.89) | 6.73E-01 | 8.33E-01 | 0.34 | (-2.02- 2.76) | 7.78E-01 | 9.16E-01 | 0.56 | (-2.48- 3.69) | 7.21E-01 | 8.50E-01 |
| 465 | PI(17:0_18:1) | -1.02 | (-2.19- 0.33) | 1.31E-01 | 2.18E-01 | -1.70 | (-3.22- 0.15) | 6.97E-02 | 1.71E-01 | -1.17 | (-3.90- 1.63) | 4.06E-01 | 6.75E-01 | -0.41 | (-4.02- 3.34) | 8.27E-01 | 9.21E-01 |
| 466 | PI(17:0_18:2) | -2.29 | (-3.38--1.03) | **8.51E-04** | **2.37E-03** | -1.03 | (-2.76- 1.10) | 3.16E-01 | 5.08E-01 | 0.58 | (-2.09- 3.34) | 6.71E-01 | 8.54E-01 | -0.59 | (-3.65- 2.57) | 7.09E-01 | 8.49E-01 |
| 467 | PI(18:0_18:1) | -1.36 | (-2.17--0.46) | **4.04E-03** | **1.02E-02** | -0.99 | (-2.08- 0.24) | 1.10E-01 | 2.42E-01 | -2.06 | (-4.76- 0.71) | 1.43E-01 | 3.60E-01 | -1.79 | (-5.23- 1.78) | 3.19E-01 | 5.17E-01 |
| 468 | PI(36:2) | -1.48 | (-2.19--0.71) | **3.09E-04** | **9.28E-04** | -0.79 | (-1.70- 0.23) | 1.26E-01 | 2.69E-01 | -1.43 | (-3.58- 0.77) | 2.01E-01 | 4.48E-01 | -1.68 | (-4.10- 0.80) | 1.81E-01 | 3.68E-01 |
| 469 | PI(18:1_18:2) | -1.40 | (-2.07--0.67) | **2.93E-04** | **8.89E-04** | -1.51 | (-2.29--0.65) | **9.89E-04** | **4.85E-03** | -3.25 | (-5.48--0.96) | **5.81E-03** | **4.98E-02** | -3.22 | (-6.17--0.17) | **3.86E-02** | 1.47E-01 |

| | | | | | | | | | | | | | | | | | |
|---|---|---|---|---|---|---|---|---|---|---|---|---|---|---|---|---|---|
| 470 | PI(16:0_20:3) (a) | 0.08 | (-0.59- 0.78) | 8.27E-01 | 8.88E-01 | 0.57 | (-0.33- 1.56) | 2.20E-01 | 4.08E-01 | -0.18 | (-2.18- 1.86) | 8.60E-01 | 9.40E-01 | 0.42 | (-2.53- 3.46) | 7.81E-01 | 8.97E-01 |
| 471 | PI(16:0_20:3) (b) | 3.31 | ( 1.88- 4.92) | **1.16E-06** | **4.74E-06** | 2.73 | ( 1.18- 4.48) | **2.82E-04** | **1.88E-03** | -0.43 | (-2.81- 2.02) | 7.29E-01 | 8.86E-01 | 2.82 | (-0.48- 6.22) | 9.39E-02 | 2.49E-01 |
| 472 | PI(16:0_20:4) | -0.09 | (-0.75- 0.63) | 8.04E-01 | 8.65E-01 | 0.52 | (-0.40- 1.51) | 2.77E-01 | 4.73E-01 | 0.95 | (-1.10- 3.03) | 3.65E-01 | 6.37E-01 | 2.24 | (-0.68- 5.23) | 1.32E-01 | 3.06E-01 |
| 473 | PI(18:0_20:2) | 3.06 | ( 1.34- 5.04) | **2.33E-04** | **7.25E-04** | 1.63 | (-0.16- 3.76) | 7.72E-02 | 1.82E-01 | -3.21 | (-5.66--0.70) | **1.28E-02** | 7.41E-02 | -1.07 | (-4.23- 2.20) | 5.15E-01 | 6.98E-01 |
| 474 | PI(18:0_20:3) (a) | -0.40 | (-0.97- 0.20) | 1.87E-01 | 2.95E-01 | 0.00 | (-0.81- 0.87) | 9.93E-01 | 9.97E-01 | -1.35 | (-3.12- 0.45) | 1.39E-01 | 3.52E-01 | -0.37 | (-2.42- 1.73) | 7.28E-01 | 8.57E-01 |
| 475 | PI(18:0_20:4) | -0.73 | (-1.25--0.17) | **1.08E-02** | **2.42E-02** | -0.31 | (-1.02- 0.47) | 4.28E-01 | 6.22E-01 | -1.81 | (-3.59- 0.00) | 5.03E-02 | 2.05E-01 | 0.04 | (-2.13- 2.25) | 9.73E-01 | 9.85E-01 |
| 476 | PI(38:5) (a) | -0.06 | (-0.87- 0.82) | 8.84E-01 | 9.27E-01 | -1.22 | (-2.02--0.34) | **7.69E-03** | **2.83E-02** | -1.78 | (-3.60- 0.08) | 6.07E-02 | 2.20E-01 | 0.38 | (-2.04- 2.86) | 7.58E-01 | 8.85E-01 |
| 477 | PI(38:6) | -1.14 | (-1.80--0.42) | **2.42E-03** | **6.43E-03** | -0.49 | (-1.33- 0.44) | 2.92E-01 | 4.89E-01 | 0.21 | (-1.99- 2.46) | 8.54E-01 | 9.40E-01 | -0.66 | (-3.33- 2.08) | 6.31E-01 | 7.89E-01 |
| 478 | PI(18:0_22:4) | 1.38 | ( 0.50- 2.34) | **1.81E-03** | **4.87E-03** | 1.01 | (-0.05- 2.18) | 6.34E-02 | 1.59E-01 | -0.76 | (-2.93- 1.45) | 4.94E-01 | 7.35E-01 | -0.75 | (-3.46- 2.04) | 5.93E-01 | 7.59E-01 |
| 479 | PI(18:0_22:5) (n3) | -1.48 | (-2.27--0.61) | **1.28E-03** | **3.52E-03** | -0.73 | (-1.79- 0.47) | 2.22E-01 | 4.08E-01 | 0.11 | (-2.14- 2.41) | 9.25E-01 | 9.71E-01 | 1.75 | (-1.21- 4.81) | 2.48E-01 | 4.40E-01 |
| 480 | PI(18:0_22:6) | -1.27 | (-1.93--0.56) | **6.94E-04** | **2.00E-03** | -0.74 | (-1.60- 0.21) | 1.22E-01 | 2.61E-01 | 0.19 | (-2.12- 2.56) | 8.70E-01 | 9.45E-01 | -0.59 | (-3.61- 2.52) | 7.04E-01 | 8.48E-01 |

**Supplementary Table 6.** Association studies with pre-pregnancy BMI and birth weight in Barwon Infant Study (BIS) and the meta-analysis results (GUSTO and BIS). Related to Figure 4 and Supplementary Figure 5.

| | BIS - ppBMI (antenatal) adjusted for ethnicity, maternal age, maternal education level and gestational weight gain | | | | | Meta Analysis of ppBMI study (GUSTO and BIS) | | | | BIS - Birth Weight (cord blood) adjusted for sex, ethnicity, maternal age, maternal education level, pre-pregnancy BMI, gestational age and parity | | | | | Meta Analysis of BW study (GUSTO and BIS) | | | |
|---|---|---|---|---|---|---|---|---|---|---|---|---|---|---|---|---|---|---|
| Lipid Species | %change in lipid concentration per unit BMI | 95%CI | p-value | p-value(BH) | Comparison between GUSTO and BIS | Overall effect size (%change) | Overall 95%CI | Overall p-value | Overall p-value(BH) | %change in lipid concentration per 100 grams | 95%CI | p-value | p-value(BH) | Comparison between GUSTO and BIS | Overall Effect Size (%change) | Overall 95%CI | Overall p-value | Overall p-value(BH) |
| AC(12:0) | 3.36 | (1.75-4.99) | 4.02E-05 | 2.06E-04 | BIS only | 1.23 | (0.34-2.11) | 6.89E-03 | 1.11E-02 | -1.93 | (-2.73--1.13) | 2.89E-06 | 1.63E-05 | GUSTO and BIS | -1.69 | (-2.17--1.21) | 6.51E-12 | 2.70E-11 |
| AC(13:0) | 1.81 | (0.09-3.56) | 3.97E-02 | 7.21E-02 | notSig | 1.51 | (0.45-2.57) | 5.44E-03 | 9.06E-03 | -1.04 | (-1.89--0.2) | 1.59E-02 | 3.46E-02 | BIS only | -1.09 | (-1.76--0.41) | 1.66E-03 | 3.23E-03 |
| AC(14:0) | 1.10 | (0.07-2.14) | 3.63E-02 | 6.69E-02 | notSig | 0.57 | (-0.01-1.15) | 5.28E-02 | 7.22E-02 | -1.47 | (-2.33--0.61) | 8.38E-04 | 2.64E-03 | GUSTO and BIS | -1.17 | (-1.65--0.68) | 2.12E-06 | 5.68E-06 |
| AC(14:1) | 5.56 | (3.78-7.37) | 7.46E-10 | 2.16E-08 | GUSTO and BIS | 2.62 | (1.64-3.6) | 1.57E-07 | 6.30E-07 | -1.48 | (-2.34--0.63) | 7.02E-04 | 2.25E-03 | GUSTO and BIS | -1.23 | (-1.77--0.7) | 5.89E-06 | 1.52E-05 |
| AC(14:2) | 4.48 | (2.74-6.25) | 3.73E-07 | 3.93E-06 | BIS only | 2.02 | (1.08-2.96) | 2.51E-05 | 6.69E-05 | -1.38 | (-2.05--0.71) | 5.56E-05 | 2.26E-04 | GUSTO and BIS | -1.13 | (-1.56--0.69) | 3.41E-07 | 9.70E-07 |
| AC(16:0) | 1.08 | (0.31-1.85) | 6.12E-03 | 1.49E-02 | BIS only | 0.56 | (0.13-0.99) | 1.07E-02 | 1.66E-02 | -0.36 | (-0.98-0.27) | 2.63E-01 | 3.63E-01 | notSig | -0.42 | (-0.83--0.01) | 4.43E-02 | 6.64E-02 |
| AC(16:1) | 3.86 | (2.73-5) | 2.01E-11 | 1.87E-09 | GUSTO and BIS | 2.12 | (1.45-2.78) | 5.13E-10 | 2.94E-09 | -1.29 | (-2.29--0.3) | 1.08E-02 | 2.50E-02 | GUSTO and BIS | -1.09 | (-1.65--0.54) | 1.08E-04 | 2.49E-04 |
| AC(18:0) | -0.02 | (-0.89-0.85) | 9.57E-01 | 9.66E-01 | notSig | -0.06 | (-0.51-0.39) | 7.85E-01 | 8.17E-01 | -0.32 | (-1.03-0.39) | 3.76E-01 | 4.82E-01 | GUSTO only | -0.62 | (-1.04--0.19) | 4.54E-03 | 8.40E-03 |
| AC(18:1) | 2.21 | (1.23-3.21) | 1.07E-05 | 6.72E-05 | GUSTO and BIS | 1.64 | (1.07-2.22) | 1.74E-08 | 7.98E-08 | -0.02 | (-0.96-0.92) | 9.62E-01 | 9.66E-01 | notSig | 0.05 | (-0.51-0.62) | 8.52E-01 | 8.86E-01 |
| AC(18:2) | -0.10 | (-0.96-0.78) | 8.30E-01 | 8.71E-01 | GUSTO only | 0.58 | (0.04-1.11) | 3.55E-02 | 4.97E-02 | -0.86 | (-1.83-0.12) | 8.58E-02 | 1.42E-01 | notSig | -0.55 | (-1.09--0.02) | 4.16E-02 | 6.31E-02 |
| CE(14:0) | -3.91 | (-4.95--2.86) | 1.52E-12 | 3.69E-10 | GUSTO and BIS | -2.36 | (-2.84--1.87) | 2.35E-21 | 9.90E-20 | 0.46 | (-0.09-1.02) | 9.98E-02 | 1.62E-01 | notSig | -0.04 | (-0.42-0.35) | 8.54E-01 | 8.86E-01 |
| CE(16:1) | -1.32 | (-2.52--0.11) | 3.33E-02 | 6.31E-02 | GUSTO only | -1.13 | (-1.78--0.49) | 6.00E-04 | 1.16E-03 | 2.15 | (1.5-2.81) | 2.31E-10 | 2.75E-09 | GUSTO and BIS | 1.59 | (1.12-2.07) | 5.07E-11 | 1.88E-10 |
| CE(18:0) | -0.68 | (-1.39-0.05) | 6.63E-02 | 1.13E-01 | GUSTO only | -0.97 | (-1.39--0.56) | 4.91E-06 | 1.44E-05 | 1.13 | (0.4-1.85) | 2.42E-03 | 6.75E-03 | BIS only | 0.74 | (0.2-1.28) | 7.07E-03 | 1.25E-02 |
| CE(18:1) | -0.17 | (-0.61-0.28) | 4.52E-01 | 5.34E-01 | GUSTO only | -0.68 | (-0.93--0.43) | 1.48E-07 | 6.13E-07 | 0.94 | (0.41-1.46) | 4.81E-04 | 1.60E-03 | BIS only | 0.48 | (0.14-0.81) | 5.70E-03 | 1.03E-02 |
| CE(18:2) | -0.53 | (-0.92--0.15) | 6.72E-03 | 1.62E-02 | GUSTO and BIS | -0.70 | (-0.95--0.44) | 1.33E-07 | 5.65E-07 | 0.14 | (-0.48-0.76) | 6.58E-01 | 7.31E-01 | notSig | -0.18 | (-0.58-0.21) | 3.58E-01 | 4.15E-01 |
| CE(18:3) | -4.43 | (-5.6--3.24) | 1.59E-12 | 3.69E-10 | GUSTO and BIS | -2.46 | (-3--1.91) | 6.77E-19 | 1.85E-17 | 2.20 | (1.43-2.97) | 2.92E-08 | 2.38E-07 | BIS only | 1.40 | (0.81-1.98) | 2.73E-06 | 7.24E-06 |
| CE(20:1) | -0.30 | (-0.98-0.38) | 3.84E-01 | 4.73E-01 | GUSTO only | -0.67 | (-1.05--0.29) | 5.79E-04 | 1.13E-03 | 1.11 | (0.33-1.9) | 5.51E-03 | 1.42E-02 | BIS only | 0.86 | (0.28-1.43) | 3.60E-03 | 6.75E-03 |
| CE(20:2) | -1.31 | (-1.91--0.72) | 1.87E-05 | 1.08E-04 | GUSTO and BIS | -1.32 | (-1.66--0.98) | 2.52E-14 | 2.93E-13 | 1.17 | (0.57-1.78) | 1.47E-04 | 5.52E-04 | GUSTO and BIS | 1.00 | (0.55-1.44) | 1.04E-05 | 2.60E-05 |
| CE(20:4) | -1.38 | (-2.49--0.27) | 1.51E-02 | 3.24E-02 | BIS only | -0.84 | (-1.39--0.29) | 2.95E-03 | 5.14E-03 | 1.21 | (0.43-1.98) | 2.38E-03 | 6.68E-03 | BIS only | 0.32 | (-0.14-0.79) | 1.72E-01 | 2.22E-01 |
| CE(20:5) | -2.86 | (-4.45--1.24) | 6.17E-04 | 2.19E-03 | GUSTO and BIS | -2.18 | (-3.21--1.15) | 3.28E-05 | 8.36E-05 | 1.01 | (-0.11-2.13) | 7.69E-02 | 1.31E-01 | GUSTO only | -0.82 | (-1.58--0.05) | 3.66E-02 | 5.67E-02 |
| CE(22:5) (n6) | -2.13 | (-3.07--1.17) | 1.65E-05 | 9.70E-05 | GUSTO and BIS | -1.95 | (-2.44--1.45) | 1.99E-14 | 2.44E-13 | 1.56 | (0.94-2.17) | 8.10E-07 | 5.01E-06 | BIS only | 1.13 | (0.65-1.61) | 4.76E-06 | 1.24E-05 |
| CE(22:6) | -1.77 | (-2.92--0.61) | 2.93E-03 | 8.08E-03 | GUSTO and BIS | -1.61 | (-2.22--1) | 2.48E-07 | 9.75E-07 | 0.14 | (-0.48-0.76) | 6.58E-01 | 7.31E-01 | GUSTO only | -0.65 | (-1.14--0.15) | 9.99E-03 | 1.71E-02 |
| CE(24:5) | 0.29 | (-0.71-1.3) | 5.69E-01 | 6.48E-01 | GUSTO only | -1.24 | (-1.82--0.67) | 2.48E-05 | 6.69E-05 | -1.43 | (-2.54--0.31) | 1.21E-02 | 2.75E-02 | GUSTO and BIS | -2.34 | (-3.02--1.67) | 1.14E-11 | 4.55E-11 |
| CE(24:6) | -3.03 | (-4.14--1.91) | 1.92E-07 | 2.41E-06 | GUSTO and BIS | -2.69 | (-3.3--2.08) | 4.80E-18 | 1.24E-16 | -0.37 | (-1.22-0.47) | 3.85E-01 | 4.90E-01 | GUSTO only | -0.71 | (-1.34--0.08) | 2.72E-02 | 4.35E-02 |
| DG(16:0_16:1) | 1.88 | (0.13-3.66) | 3.49E-02 | 6.55E-02 | notSig | 0.96 | (0.07-1.85) | 3.54E-02 | 4.97E-02 | -1.05 | (-1.86--0.24) | 1.15E-02 | 2.65E-02 | GUSTO and BIS | -1.18 | (-1.76--0.6) | 6.52E-05 | 1.54E-04 |
| DG(14:0_18:2) | -1.79 | (-3.27--0.29) | 1.99E-02 | 4.02E-02 | BIS only | -1.27 | (-2.1--0.43) | 2.89E-03 | 5.05E-03 | -2.05 | (-3.02--1.08) | 3.47E-05 | 1.49E-04 | GUSTO and BIS | -2.53 | (-3.23--1.82) | 2.49E-12 | 1.06E-11 |
| DG(16:0_18:1) | 2.82 | (1.49-4.16) | 2.90E-05 | 1.57E-04 | GUSTO and BIS | 1.68 | (1.03-2.33) | 3.83E-07 | 1.42E-06 | -1.34 | (-2.33--0.36) | 7.64E-03 | 1.88E-02 | GUSTO and BIS | -1.72 | (-2.29--1.16) | 2.38E-09 | 8.12E-09 |
| DG(16:1_18:1) | 3.07 | (1.7-4.47) | 1.09E-05 | 6.72E-05 | BIS only | 1.43 | (0.69-2.17) | 1.38E-04 | 3.16E-04 | -0.86 | (-1.72--0.01) | 4.75E-02 | 9.00E-02 | GUSTO only | -1.57 | (-2.12--1.01) | 2.91E-08 | 8.99E-08 |
| DG(16:0_18:2) | 1.91 | (0.58-3.25) | 4.66E-03 | 1.21E-02 | GUSTO and BIS | 1.15 | (0.51-1.78) | 3.88E-04 | 7.93E-04 | -2.81 | (-3.89--1.73) | 4.47E-07 | 3.01E-06 | GUSTO and BIS | -2.99 | (-3.59--2.39) | 1.76E-22 | 1.36E-21 |
| DG(18:0_18:1) | 2.39 | (1.17-3.63) | 1.27E-04 | 5.45E-04 | GUSTO and BIS | 1.60 | (1.01-2.19) | 1.29E-07 | 5.53E-07 | -0.66 | (-1.5-0.18) | 1.25E-01 | 1.95E-01 | GUSTO only | -1.02 | (-1.57--0.47) | 2.89E-04 | 6.35E-04 |
| DG(18:1_18:1) | 3.44 | (2.11-4.79) | 3.65E-07 | 3.93E-06 | GUSTO and BIS | 1.84 | (1.2-2.48) | 1.70E-08 | 7.89E-08 | -0.91 | (-1.93-0.11) | 8.07E-02 | 1.35E-01 | GUSTO only | -1.57 | (-2.13--1.01) | 3.94E-08 | 1.20E-07 |
| DG(18:0_18:2) | 1.65 | (0.26-3.06) | 1.98E-02 | 4.01E-02 | GUSTO and BIS | 1.14 | (0.47-1.81) | 8.15E-04 | 1.54E-03 | -1.86 | (-2.84--0.87) | 2.22E-04 | 8.05E-04 | GUSTO and BIS | -1.74 | (-2.42--1.05) | 7.02E-07 | 1.93E-06 |
| DG(18:1_18:2) | 2.55 | (1.26-3.86) | 1.00E-04 | 4.47E-04 | GUSTO and BIS | 1.33 | (0.7-1.97) | 3.83E-05 | 9.51E-05 | -2.31 | (-3.35--1.27) | 1.45E-05 | 6.95E-05 | GUSTO and BIS | -2.92 | (-3.47--2.37) | 9.92E-26 | 9.21E-25 |
| DG(18:2_18:2) | 1.55 | (-0.09-3.22) | 6.41E-02 | 1.11E-01 | notSig | 1.04 | (0.23-1.86) | 1.19E-02 | 1.84E-02 | -3.01 | (-4.18--1.84) | 5.35E-07 | 3.54E-06 | GUSTO and BIS | -3.84 | (-4.46--3.23) | 1.99E-34 | 2.98E-33 |
| DG(18:1_18:3) | 0.76 | (-0.53-2.06) | 2.50E-01 | 3.37E-01 | notSig | 0.47 | (-0.2-1.14) | 1.67E-01 | 2.06E-01 | -1.91 | (-2.84--0.98) | 6.56E-05 | 2.65E-04 | GUSTO and BIS | -2.32 | (-2.93--1.71) | 1.08E-13 | 5.07E-13 |
| DG(16:0_20:4) | 2.13 | (0.73-3.55) | 2.73E-03 | 7.58E-03 | GUSTO and BIS | 1.47 | (0.7-2.25) | 2.00E-04 | 4.39E-04 | -2.23 | (-3.31--1.16) | 5.00E-05 | 2.07E-04 | GUSTO and BIS | -2.12 | (-2.77--1.48) | 1.10E-10 | 3.94E-10 |
| DG(18:1_20:3) | 1.68 | (0.53-2.84) | 4.11E-03 | 1.09E-02 | BIS only | 0.80 | (0.17-1.44) | 1.29E-02 | 1.96E-02 | 0.37 | (-0.43-1.17) | 3.59E-01 | 4.65E-01 | notSig | 0.04 | (-0.53-0.61) | 8.84E-01 | 9.07E-01 |
| DG(18:1_20:4) | 3.07 | (1.99-4.16) | 2.60E-08 | 4.17E-07 | GUSTO and BIS | 1.75 | (1.14-2.36) | 2.01E-08 | 9.07E-08 | -1.70 | (-2.54--0.86) | 7.91E-05 | 3.06E-04 | GUSTO and BIS | -1.92 | (-2.45--1.4) | 6.80E-13 | 2.98E-12 |
| DG(16:0_22:5) | 2.32 | (0.94-3.72) | 9.73E-04 | 3.07E-03 | BIS only | 1.08 | (0.33-1.82) | 4.65E-03 | 7.82E-03 | -4.31 | (-5.44--3.18) | 2.22E-13 | 4.12E-12 | GUSTO and BIS | -2.76 | (-3.42--2.09) | 3.35E-16 | 1.85E-15 |
| DG(18:2_20:4) | 2.45 | (1.24-3.67) | 6.69E-05 | 3.11E-04 | BIS only | 1.35 | (0.65-2.04) | 1.50E-04 | 3.41E-04 | -3.12 | (-4.09--2.15) | 4.51E-10 | 4.77E-09 | GUSTO and BIS | -3.13 | (-3.74--2.53) | 2.07E-24 | 1.78E-23 |
| DG(16:0_22:6) | 3.43 | (1.88-5.01) | 1.30E-05 | 7.85E-05 | BIS only | 1.94 | (1.03-2.86) | 3.23E-05 | 8.28E-05 | -4.45 | (-5.52--3.39) | 1.00E-15 | 2.45E-14 | GUSTO and BIS | -4.20 | (-4.79--3.61) | 5.66E-44 | 1.75E-42 |
| DG(18:1_22:5) | 3.06 | (1.95-4.17) | 5.46E-08 | 7.45E-07 | BIS only | 1.43 | (0.76-2.09) | 2.68E-05 | 7.06E-05 | -1.58 | (-2.32--0.84) | 3.41E-05 | 1.48E-04 | GUSTO and BIS | -1.73 | (-2.27--1.19) | 4.74E-10 | 1.68E-09 |
| DG(18:1_22:6) | 4.34 | (2.91-5.79) | 2.49E-09 | 5.50E-08 | GUSTO and BIS | 2.53 | (1.67-3.39) | 7.83E-09 | 3.79E-08 | -3.89 | (-4.84--2.93) | 4.72E-15 | 1.04E-13 | GUSTO and BIS | -3.63 | (-4.19--3.07) | 4.57E-37 | 9.64E-36 |
| DG(18:2_22:6) | 3.34 | (1.77-4.94) | 2.92E-05 | 1.57E-04 | GUSTO and BIS | 2.12 | (1.21-3.02) | 4.73E-06 | 1.41E-05 | -5.08 | (-6.21--3.96) | 4.34E-18 | 2.01E-16 | GUSTO and BIS | -4.36 | (-5.01--3.71) | 7.07E-40 | 1.64E-38 |
| TG(48:0) [NL-18:0] | -3.52 | (-5.19--1.83) | 5.76E-05 | 2.73E-04 | GUSTO and BIS | -1.98 | (-2.76--1.2) | 6.07E-07 | 2.13E-06 | 0.07 | (-0.69-0.84) | 8.51E-01 | 8.91E-01 | GUSTO only | -0.58 | (-1.16-0) | 5.07E-02 | 7.47E-02 |
| TG(48:1) [NL-16:1] | -0.31 | (-1.92-1.33) | 7.10E-01 | 7.74E-01 | notSig | 0.24 | (-0.69-1.17) | 6.09E-01 | 6.62E-01 | -0.31 | (-1.24-0.62) | 5.11E-01 | 6.06E-01 | GUSTO only | -1.34 | (-2.01--0.68) | 7.78E-05 | 1.82E-04 |
| TG(48:1) [NL-18:1] | -2.33 | (-3.29--1.37) | 3.02E-06 | 2.46E-05 | BIS only | -1.40 | (-2.05--0.75) | 2.50E-05 | 6.69E-05 | -0.80 | (-1.74-0.14) | 9.49E-02 | 1.54E-01 | GUSTO only | -2.11 | (-2.73--1.48) | 4.31E-11 | 1.61E-10 |

| | | | | | | | | | | | | | | | | | | |
|---|---|---|---|---|---|---|---|---|---|---|---|---|---|---|---|---|---|---|
| TG(48:2) [NL-14:0] | -1.38 | (-2.47--0.28) | **1.44E-02** | **3.12E-02** | BIS only | -1.15 | (-1.9--0.39) | **2.83E-03** | **4.98E-03** | -2.11 | (-3.22--1) | **1.96E-04** | **7.20E-04** | GUSTO and BIS | -3.24 | (-3.86--2.62) | **1.29E-24** | **1.13E-23** |
| TG(48:2) [NL-14:1] | -1.37 | (-2.85-0.13) | 7.37E-02 | 1.22E-01 | GUSTO only | -1.26 | (-2.12--0.39) | **4.40E-03** | **7.46E-03** | -1.65 | (-2.91--0.39) | **1.01E-02** | **2.37E-02** | GUSTO and BIS | -2.67 | (-3.33--2) | **5.20E-15** | **2.65E-14** |
| TG(48:2) [NL-16:1] | 0.72 | (-0.78-2.25) | 3.48E-01 | 4.37E-01 | notSig | 0.21 | (-0.78-1.2) | 6.73E-01 | 7.19E-01 | -0.87 | (-1.76-0.03) | 5.73E-02 | 1.05E-01 | GUSTO only | -1.92 | (-2.53--1.3) | **9.98E-10** | **3.43E-09** |
| TG(48:2) [NL-18:2] | -2.43 | (-3.5--1.36) | **1.21E-05** | **7.41E-05** | GUSTO and BIS | -1.78 | (-2.52--1.03) | **2.94E-06** | **9.16E-06** | -2.40 | (-3.4--1.39) | **3.60E-06** | **1.99E-05** | GUSTO and BIS | -3.50 | (-4.11--2.89) | **3.79E-29** | **4.19E-28** |
| TG(48:3) [NL-14:0] | -1.75 | (-3.43--0.04) | **4.49E-02** | 8.05E-02 | GUSTO only | -1.58 | (-2.53--0.64) | **9.75E-04** | **1.81E-03** | -3.53 | (-4.8--2.25) | **7.61E-08** | **5.88E-07** | GUSTO and BIS | -3.91 | (-4.55--3.27) | **3.03E-33** | **4.26E-32** |
| TG(48:3) [NL-16:1] | 0.38 | (-1.25-2.05) | 6.47E-01 | 7.13E-01 | notSig | -0.64 | (-1.59-0.31) | 1.86E-01 | 2.27E-01 | -2.17 | (-3.11--1.23) | **6.87E-06** | **3.58E-05** | GUSTO and BIS | -2.62 | (-3.2--2.04) | **9.32E-19** | **5.61E-18** |
| TG(48:3) [NL-18:3] | -3.12 | (-4.76--1.46) | **2.71E-04** | **1.06E-03** | GUSTO and BIS | -2.30 | (-3.3--1.29) | **7.64E-06** | **2.19E-05** | -2.68 | (-3.82--1.55) | **4.18E-06** | **2.23E-05** | GUSTO and BIS | -3.08 | (-3.68--2.47) | **1.61E-23** | **1.31E-22** |
| TG(49:1) [NL-17:1] | 0.59 | (-0.99-2.2) | 4.67E-01 | 5.50E-01 | notSig | 0.39 | (-0.37-1.16) | 3.14E-01 | 3.62E-01 | -1.18 | (-2.06--0.3) | **8.39E-03** | **2.04E-02** | GUSTO and BIS | -1.80 | (-2.34--1.26) | **6.71E-11** | **2.43E-10** |
| TG(50:1) [NL-14:0] | -2.79 | (-3.99--1.57) | **9.51E-06** | **6.23E-05** | BIS only | -1.37 | (-2.01--0.73) | **3.05E-05** | **7.90E-05** | -0.33 | (-1.16-0.51) | 4.43E-01 | 5.41E-01 | GUSTO only | -0.83 | (-1.46--0.21) | **9.17E-03** | **1.59E-02** |
| TG(50:1) [NL-16:0] | 0.08 | (-0.72-0.89) | 8.39E-01 | 8.75E-01 | GUSTO only | 0.56 | (0.07-1.04) | **2.39E-02** | **3.43E-02** | -0.91 | (-1.67--0.15) | **1.93E-02** | **4.12E-02** | GUSTO and BIS | -1.65 | (-2.2--1.1) | **4.95E-09** | **1.64E-08** |
| TG(50:1) [NL-18:1] | 0.05 | (-0.66-0.76) | 8.98E-01 | 9.20E-01 | GUSTO only | 0.47 | (0.02-0.92) | **3.88E-02** | 5.39E-02 | -0.86 | (-1.52--0.21) | **1.01E-02** | **2.37E-02** | GUSTO and BIS | -1.50 | (-2.01--0.99) | **7.44E-09** | **2.40E-08** |
| TG(50:2) [NL-14:0] | -0.97 | (-1.73--0.2) | **1.41E-02** | **3.07E-02** | GUSTO and BIS | -0.76 | (-1.22--0.3) | **1.14E-03** | **2.08E-03** | -0.48 | (-1.4-0.45) | 3.11E-01 | 4.14E-01 | GUSTO only | -2.09 | (-2.7--1.48) | **1.95E-11** | **7.56E-11** |
| TG(50:2) [NL-16:1] | 1.29 | (0.46-2.13) | **2.41E-03** | **6.77E-03** | BIS only | 0.92 | (0.38-1.47) | **9.27E-04** | **1.73E-03** | -0.44 | (-1.23-0.34) | 2.68E-01 | 3.67E-01 | GUSTO only | -1.62 | (-2.2--1.05) | **3.43E-08** | **1.05E-07** |
| TG(50:2) [NL-18:1] | 0.16 | (-0.42-0.75) | 5.85E-01 | 6.61E-01 | notSig | 0.23 | (-0.19-0.66) | 2.82E-01 | 3.33E-01 | -0.42 | (-1.11-0.27) | 2.33E-01 | 3.33E-01 | GUSTO only | -1.50 | (-2.03--0.96) | **4.88E-08** | **1.45E-07** |
| TG(50:2) [NL-18:2] | -0.43 | (-1.17-0.32) | 2.61E-01 | 3.50E-01 | GUSTO only | 0.25 | (-0.21-0.7) | 2.91E-01 | 3.40E-01 | -2.06 | (-2.84--1.27) | **3.16E-07** | **2.19E-06** | GUSTO and BIS | -3.08 | (-3.64--2.52) | **4.68E-27** | **4.72E-26** |
| TG(50:3) [NL-14:0] | -1.31 | (-2.05--0.58) | **5.22E-04** | **1.92E-03** | BIS only | -0.91 | (-1.43--0.39) | **5.80E-04** | **1.13E-03** | -2.92 | (-4.09--1.74) | **1.49E-06** | **8.86E-06** | GUSTO and BIS | -3.87 | (-4.5--3.23) | **4.82E-33** | **6.39E-32** |
| TG(50:3) [NL-14:1] | -0.54 | (-1.57-0.5) | 3.05E-01 | 3.97E-01 | notSig | -0.61 | (-1.17--0.06) | **3.04E-02** | **4.30E-02** | -1.34 | (-2.49--0.18) | **2.31E-02** | **4.77E-02** | GUSTO and BIS | -2.03 | (-2.7--1.35) | **3.68E-09** | **1.23E-08** |
| TG(50:3) [NL-16:1] | 1.82 | (0.94-2.72) | **5.63E-05** | **2.69E-04** | BIS only | 1.17 | (0.55-1.78) | **1.83E-04** | **4.07E-04** | -1.95 | (-2.85--1.05) | **2.21E-05** | **1.01E-04** | GUSTO and BIS | -3.10 | (-3.67--2.52) | **3.03E-26** | **2.99E-25** |
| TG(50:3) [NL-18:2] | 0.36 | (-0.27-0.99) | 2.69E-01 | 3.60E-01 | notSig | 0.32 | (-0.16-0.81) | 1.85E-01 | 2.26E-01 | -2.52 | (-3.45--1.6) | **1.12E-07** | **8.36E-07** | GUSTO and BIS | -3.65 | (-4.23--3.08) | **1.24E-35** | **2.13E-34** |
| TG(50:3) [NL-18:3] | -1.20 | (-2.3--0.08) | **3.50E-02** | 6.56E-02 | notSig | -0.41 | (-1.12-0.3) | 2.57E-01 | 3.06E-01 | -2.39 | (-3.36--1.42) | **1.62E-06** | **9.51E-06** | GUSTO and BIS | -3.25 | (-3.86--2.64) | **1.05E-25** | **9.52E-25** |
| TG(50:4) [NL-14:0] | -1.26 | (-2.67-0.17) | 8.43E-02 | 1.37E-01 | notSig | -1.00 | (-1.83--0.18) | **1.73E-02** | **2.55E-02** | -4.67 | (-6.01--3.34) | **1.27E-11** | **1.73E-10** | GUSTO and BIS | -4.39 | (-5.01--3.77) | **1.13E-43** | **3.28E-42** |
| TG(50:4) [NL-18:3] | -0.36 | (-1.65-0.94) | 5.84E-01 | 6.61E-01 | notSig | -0.58 | (-1.39-0.23) | 1.62E-01 | 2.01E-01 | -3.25 | (-4.4--2.11) | **3.61E-08** | **2.84E-07** | GUSTO and BIS | -3.75 | (-4.33--3.17) | **1.06E-36** | **2.05E-35** |
| TG(50:4) [NL-20:4] | -1.30 | (-2.92-0.35) | 1.22E-01 | 1.84E-01 | notSig | -1.02 | (-2.07-0.04) | 5.92E-02 | 8.06E-02 | -2.53 | (-3.68--1.38) | **1.81E-05** | **8.42E-05** | GUSTO and BIS | -3.15 | (-3.85--2.45) | **1.04E-18** | **6.18E-18** |
| TG(51:1) [NL-17:0] | 1.56 | (0.29-2.83) | **1.55E-02** | **3.29E-02** | BIS only | 0.90 | (0.25-1.54) | **6.39E-03** | **1.04E-02** | -1.72 | (-2.74--0.71) | **8.91E-04** | **2.77E-03** | GUSTO and BIS | -2.36 | (-2.97--1.75) | **3.56E-14** | **1.72E-13** |
| TG(51:2) [NL-15:0] | 1.27 | (0.22-2.32) | **1.79E-02** | **3.72E-02** | BIS only | -0.02 | (-0.58-0.54) | 9.37E-01 | 9.47E-01 | -2.20 | (-3.35--1.06) | **1.71E-04** | **6.36E-04** | GUSTO and BIS | -3.00 | (-3.64--2.35) | **9.85E-20** | **6.38E-19** |
| TG(51:2) [NL-17:0] | 2.24 | (1.27-3.23) | **6.14E-06** | **4.43E-05** | BIS only | 0.99 | (0.42-1.55) | **5.85E-04** | **1.14E-03** | -1.83 | (-2.83--0.84) | **3.12E-04** | **1.11E-03** | GUSTO and BIS | -2.83 | (-3.43--2.23) | **3.31E-20** | **2.19E-19** |
| TG(51:2) [NL-17:1] | 2.25 | (1.27-3.24) | **6.21E-06** | **4.43E-05** | BIS only | 0.95 | (0.4-1.5) | **6.95E-04** | **1.33E-03** | -1.84 | (-2.84--0.84) | **3.17E-04** | **1.11E-03** | GUSTO and BIS | -2.57 | (-3.12--2.01) | **1.23E-19** | **7.83E-19** |
| TG(52:1) [NL-18:0] | 1.13 | (0.07-2.21) | **3.75E-02** | 6.89E-02 | GUSTO only | 1.29 | (0.65-1.93) | **7.59E-05** | **1.81E-04** | -0.34 | (-1.34-0.67) | 5.14E-01 | 6.06E-01 | GUSTO only | -1.13 | (-1.83--0.43) | **1.48E-03** | **2.90E-03** |
| TG(52:1) [NL-18:1] | 1.62 | (0.43-2.82) | **7.55E-03** | **1.79E-02** | GUSTO and BIS | 1.50 | (0.85-2.15) | **5.43E-06** | **1.57E-05** | -0.56 | (-1.66-0.55) | 3.24E-01 | 4.28E-01 | GUSTO only | -1.26 | (-1.99--0.53) | **6.97E-04** | **1.45E-03** |
| TG(52:2) [NL-16:0] | 1.13 | (0.49-1.77) | **5.41E-04** | **1.97E-03** | GUSTO and BIS | 0.90 | (0.55-1.24) | **2.89E-07** | **1.12E-06** | -0.99 | (-1.78--0.2) | **1.40E-02** | **3.09E-02** | GUSTO and BIS | -2.02 | (-2.53--1.5) | **2.34E-14** | **1.14E-13** |
| TG(52:2) [NL-18:2] | -0.46 | (-1.3-0.38) | 2.83E-01 | 3.74E-01 | notSig | 0.14 | (-0.35-0.64) | 5.74E-01 | 6.31E-01 | -1.63 | (-2.53--0.73) | **3.91E-04** | **1.33E-03** | GUSTO and BIS | -2.20 | (-2.83--1.56) | **9.40E-12** | **3.86E-11** |
| TG(52:3) [NL-16:1] | 1.90 | (1.05-2.75) | **1.03E-05** | **6.59E-05** | BIS only | 0.86 | (0.39-1.34) | **3.56E-04** | **7.37E-04** | -0.57 | (-1.42-0.28) | 1.90E-01 | 2.80E-01 | GUSTO only | -2.11 | (-2.68--1.54) | **5.56E-13** | **2.45E-12** |
| TG(52:3) [NL-18:2] | 0.51 | (-0.04-1.07) | 7.11E-02 | 1.19E-01 | GUSTO only | 0.66 | (0.32-1.01) | **1.84E-04** | **4.07E-04** | -2.39 | (-3.27--1.51) | **1.37E-07** | **9.90E-07** | GUSTO and BIS | -3.61 | (-4.16--3.05) | **2.57E-37** | **5.69E-36** |
| TG(52:4) [NL-16:1] | 2.54 | (1.64-3.44) | **2.78E-08** | **4.30E-07** | BIS only | 1.42 | (0.85-1.98) | **9.55E-07** | **3.10E-06** | -3.04 | (-4.19--1.89) | **2.88E-07** | **2.02E-06** | GUSTO and BIS | -4.12 | (-4.72--3.53) | **2.39E-42** | **6.16E-41** |
| TG(52:4) [NL-18:2] | 1.00 | (0.17-1.84) | **1.88E-02** | **3.85E-02** | GUSTO and BIS | 1.11 | (0.55-1.67) | **1.13E-04** | **2.62E-04** | -3.62 | (-4.64--2.59) | **1.11E-11** | **1.61E-10** | GUSTO and BIS | -4.53 | (-5.12--3.95) | **3.66E-52** | **1.70E-50** |
| TG(52:4) [NL-18:3] | -0.10 | (-0.85-0.65) | 7.92E-01 | 8.41E-01 | notSig | 0.20 | (-0.32-0.71) | 4.51E-01 | 5.09E-01 | -2.88 | (-4--1.75) | **6.99E-07** | **4.44E-06** | GUSTO and BIS | -3.75 | (-4.36--3.15) | **6.14E-34** | **8.90E-33** |
| TG(52:5) [NL-18:3] | 0.20 | (-1.12-1.53) | 7.71E-01 | 8.21E-01 | notSig | 0.13 | (-0.67-0.93) | 7.44E-01 | 7.84E-01 | -4.40 | (-5.64--3.16) | **8.60E-12** | **1.33E-10** | GUSTO and BIS | -4.22 | (-4.82--3.62) | **6.84E-43** | **1.87E-41** |
| TG(52:5) [NL-20:4] | 1.26 | (-0.1-2.63) | 6.85E-02 | 1.16E-01 | notSig | 0.82 | (-0.08-1.72) | 7.36E-02 | 9.81E-02 | -2.37 | (-3.41--1.33) | **8.78E-06** | **4.35E-05** | GUSTO and BIS | -3.14 | (-3.78--2.49) | **2.43E-21** | **1.73E-20** |
| TG(52:5) [NL-20:5] | -0.16 | (-1.67-1.38) | 8.41E-01 | 8.75E-01 | notSig | 0.43 | (-0.55-1.41) | 3.92E-01 | 4.47E-01 | -4.88 | (-6.11--3.66) | **1.65E-14** | **3.33E-13** | GUSTO and BIS | -4.56 | (-5.19--3.92) | **7.69E-45** | **2.74E-43** |
| TG(53:2) [NL-18:1] | 2.37 | (1.44-3.31) | **5.97E-07** | **5.90E-06** | GUSTO and BIS | 1.39 | (0.84-1.94) | **6.65E-07** | **2.26E-06** | -1.82 | (-2.78--0.86) | **2.02E-04** | **7.36E-04** | GUSTO and BIS | -2.40 | (-2.95--1.85) | **1.36E-17** | **7.80E-17** |
| TG(54:1) [NL-18:1] | 0.32 | (-1.39-2.06) | 7.16E-01 | 7.77E-01 | GUSTO only | 0.90 | (0.28-1.53) | **4.52E-03** | **7.62E-03** | 0.19 | (-0.84-1.21) | 7.22E-01 | 7.92E-01 | GUSTO only | -0.76 | (-1.47--0.05) | **3.56E-02** | 5.52E-02 |
| TG(54:2) [NL-18:0] | 0.95 | (-0.15-2.06) | 9.07E-02 | 1.46E-01 | GUSTO only | 1.28 | (0.69-1.87) | **1.97E-05** | **5.44E-05** | -0.31 | (-1.36-0.73) | 5.58E-01 | 6.40E-01 | GUSTO only | -1.21 | (-1.88--0.53) | **4.79E-04** | **1.03E-03** |
| TG(54:2) [NL-20:1] | 2.17 | (1.15-3.19) | **2.60E-05** | **1.44E-04** | BIS only | 1.07 | (0.46-1.69) | **6.58E-04** | **1.27E-03** | -0.36 | (-1.53-0.81) | 5.45E-01 | 6.32E-01 | GUSTO only | -1.33 | (-2.01--0.64) | **1.40E-04** | **3.20E-04** |
| TG(54:3) [NL-18:1] | 0.68 | (-0.2-1.58) | 1.31E-01 | 1.96E-01 | notSig | 0.60 | (0.13-1.08) | **1.27E-02** | **1.94E-02** | -0.07 | (-0.75-0.61) | 8.34E-01 | 8.82E-01 | GUSTO only | -1.04 | (-1.54--0.54) | **4.79E-05** | **1.15E-04** |
| TG(54:3) [NL-18:2] | 0.43 | (-0.44-1.31) | 3.33E-01 | 4.21E-01 | GUSTO only | 0.63 | (0.15-1.11) | **1.02E-02** | **1.59E-02** | -1.84 | (-2.74--0.93) | **7.18E-05** | **2.85E-04** | GUSTO and BIS | -2.58 | (-3.15--2.02) | **3.03E-19** | **1.88E-18** |
| TG(54:4) [NL-18:2] | 0.45 | (-0.55-1.45) | 3.81E-01 | 4.70E-01 | notSig | 0.38 | (-0.16-0.91) | 1.66E-01 | 2.05E-01 | -2.10 | (-3.02--1.18) | **8.74E-06** | **4.35E-05** | GUSTO and BIS | -3.25 | (-3.84--2.67) | **1.13E-27** | **1.19E-26** |
| TG(54:4) [NL-20:3] | 0.90 | (0.07-1.73) | **3.33E-02** | 6.31E-02 | notSig | 0.74 | (0.21-1.27) | **5.86E-03** | **9.61E-03** | -1.21 | (-2.13--0.3) | **9.51E-03** | **2.26E-02** | GUSTO and BIS | -2.04 | (-2.65--1.42) | **1.13E-10** | **4.04E-10** |
| TG(54:5) [NL-18:3] | 0.00 | (-1.21-1.22) | 9.98E-01 | 9.98E-01 | notSig | 0.04 | (-0.66-0.75) | 9.02E-01 | 9.21E-01 | -2.32 | (-3.36--1.27) | **1.47E-05** | **6.95E-05** | GUSTO and BIS | -3.32 | (-3.9--2.74) | **2.65E-29** | **3.00E-28** |
| TG(54:5) [NL-20:4] | 1.44 | (0.64-2.25) | **4.00E-04** | **1.51E-03** | GUSTO and BIS | 1.21 | (0.7-1.72) | **3.66E-06** | **1.11E-05** | -2.32 | (-3.25--1.38) | **1.46E-06** | **8.79E-06** | GUSTO and BIS | -3.12 | (-3.73--2.5) | **2.94E-23** | **2.35E-22** |
| TG(54:6) [NL-18:3] | 0.13 | (-1.5-1.79) | 8.78E-01 | 9.01E-01 | notSig | -0.23 | (-1.18-0.72) | 6.39E-01 | 6.89E-01 | -3.59 | (-4.73--2.45) | **1.06E-09** | **1.03E-08** | GUSTO and BIS | -3.78 | (-4.38--3.19) | **2.63E-35** | **4.21E-34** |
| TG(54:6) [NL-20:4] | 1.79 | (0.88-2.71) | **1.16E-04** | **5.05E-04** | GUSTO and BIS | 1.45 | (0.83-2.06) | **4.34E-06** | **1.30E-05** | -3.12 | (-4.1--2.14) | **7.87E-10** | **7.77E-09** | GUSTO and BIS | -3.73 | (-4.34--3.12) | **3.88E-33** | **5.29E-32** |
| TG(54:6) [NL-20:5] | 1.12 | (-0.09-2.35) | 6.96E-02 | 1.17E-01 | notSig | 0.97 | (0.14-1.81) | **2.18E-02** | **3.15E-02** | -4.78 | (-5.95--3.6) | **6.83E-15** | **1.44E-13** | GUSTO and BIS | -4.97 | (-5.57--4.36) | **3.94E-58** | **2.28E-56** |
| TG(54:6) [NL-22:6] | 1.92 | (0.5-3.36) | **7.92E-03** | **1.85E-02** | GUSTO and BIS | 1.84 | (0.91-2.78) | **1.06E-04** | **2.47E-04** | -4.45 | (-5.51--3.38) | **1.07E-15** | **2.49E-14** | GUSTO and BIS | -4.93 | (-5.53--4.34) | **6.08E-59** | **4.03E-57** |
| TG(54:7) [NL-20:5] | 2.11 | (0.44-3.82) | **1.34E-02** | **2.95E-02** | BIS only | 1.07 | (0.05-2.1) | **4.02E-02** | 5.57E-02 | -6.33 | (-7.66--5) | **1.20E-19** | **7.13E-18** | GUSTO and BIS | -5.27 | (-5.89--4.65) | **1.35E-61** | **1.57E-59** |
| TG(54:7) [NL-22:6] | 3.03 | (1.47-4.61) | **1.30E-04** | **5.49E-04** | BIS only | 1.84 | (0.82-2.85) | **3.90E-04** | **7.93E-04** | -5.23 | (-6.42--4.05) | **3.09E-17** | **1.10E-15** | GUSTO and BIS | -5.00 | (-5.59--4.4) | **5.92E-61** | **4.58E-59** |

| | | | | | | | | | | | | | | | | | | |
|---|---|---|---|---|---|---|---|---|---|---|---|---|---|---|---|---|---|---|
| TG(56:6) [NL-20:4] | 1.52 | (0.58-2.46) | **1.53E-03** | **4.58E-03** | GUSTO and BIS | 1.38 | (0.92-1.84) | **4.37E-09** | **2.23E-08** | -1.04 | (-1.92--0.15) | **2.18E-02** | **4.56E-02** | GUSTO and BIS | -2.11 | (-2.71--1.52) | **3.91E-12** | **1.64E-11** |
| TG(56:7) [NL-20:4] | 1.01 | (0.1-1.92) | **2.88E-02** | 5.56E-02 | GUSTO only | 1.02 | (0.48-1.56) | **2.28E-04** | **4.92E-04** | -2.52 | (-3.61--1.44) | **6.15E-06** | **3.24E-05** | GUSTO and BIS | -3.26 | (-3.91--2.62) | **3.97E-23** | **3.12E-22** |
| TG(56:7) [NL-20:5] | 0.93 | (-0.31-2.19) | 1.42E-01 | 2.11E-01 | notSig | 0.63 | (-0.2-1.46) | 1.39E-01 | 1.74E-01 | -3.67 | (-4.79--2.56) | **2.02E-10** | **2.47E-09** | GUSTO and BIS | -3.75 | (-4.38--3.12) | **3.53E-31** | **4.31E-30** |
| TG(56:7) [NL-22:6] | 3.23 | (2.1-4.37) | **2.13E-08** | **3.81E-07** | GUSTO and BIS | 2.56 | (1.77-3.35) | **2.14E-10** | **1.36E-09** | -4.32 | (-5.32--3.31) | **1.93E-16** | **5.98E-15** | GUSTO and BIS | -4.83 | (-5.4--4.25) | **2.44E-61** | **2.27E-59** |
| TG(56:8) [NL-20:4] | 0.85 | (-0.38-2.09) | 1.78E-01 | 2.55E-01 | notSig | 0.82 | (0.07-1.56) | **3.19E-02** | **4.49E-02** | -3.04 | (-4.15--1.94) | **8.83E-08** | **6.72E-07** | GUSTO and BIS | -2.98 | (-3.64--2.32) | **7.27E-19** | **4.44E-18** |
| TG(56:8) [NL-20:5] | 1.07 | (-0.46-2.61) | 1.71E-01 | 2.47E-01 | notSig | 0.54 | (-0.43-1.51) | 2.78E-01 | 3.29E-01 | -5.25 | (-6.49--4.01) | **5.15E-16** | **1.41E-14** | GUSTO and BIS | -4.40 | (-5.02--3.79) | **1.62E-44** | **5.38E-43** |
| TG(56:8) [NL-22:6] | 3.61 | (2.28-4.97) | **1.02E-07** | **1.31E-06** | GUSTO and BIS | 2.60 | (1.7-3.51) | **1.75E-08** | **7.98E-08** | -5.76 | (-6.86--4.65) | **6.44E-23** | **1.19E-20** | GUSTO and BIS | -5.46 | (-6.03--4.9) | **9.51E-80** | **4.41E-77** |
| TG(56:9) [NL-22:6] | 2.32 | (0.63-4.04) | **7.16E-03** | **1.70E-02** | BIS only | 1.29 | (0.31-2.27) | **9.72E-03** | **1.52E-02** | -6.30 | (-7.52--5.09) | **7.68E-23** | **1.19E-20** | GUSTO and BIS | -5.21 | (-5.79--4.62) | **1.65E-68** | **3.83E-66** |
| TG(58:10) [NL-22:6] | 1.99 | (0.3-3.71) | **2.10E-02** | **4.18E-02** | BIS only | 1.29 | (0.32-2.26) | **9.36E-03** | **1.47E-02** | -5.80 | (-6.99--4.61) | **1.65E-20** | **1.27E-18** | GUSTO and BIS | -4.80 | (-5.39--4.21) | **1.63E-57** | **8.42E-56** |
| TG(58:8) [NL-22:6] | 2.68 | (1.4-3.98) | **4.04E-05** | **2.06E-04** | GUSTO and BIS | 1.97 | (1.14-2.8) | **3.53E-06** | **1.08E-05** | -4.00 | (-5.01--2.99) | **2.81E-14** | **5.43E-13** | GUSTO and BIS | -4.34 | (-4.92--3.76) | **2.48E-49** | **1.04E-47** |
| TG(58:9) [NL-22:6] | 2.27 | (0.81-3.74) | **2.19E-03** | **6.26E-03** | BIS only | 1.62 | (0.72-2.53) | **4.53E-04** | **9.06E-04** | -5.43 | (-6.62--4.24) | **2.33E-18** | **1.20E-16** | GUSTO and BIS | -4.92 | (-5.5--4.34) | **4.05E-62** | **6.26E-60** |
| TG(O-50:1) [NL-16:0] | -0.17 | (-1.21-0.88) | 7.51E-01 | 8.08E-01 | notSig | -0.19 | (-0.71-0.32) | 4.60E-01 | 5.19E-01 | -0.74 | (-1.49-0.02) | 5.49E-02 | 1.01E-01 | notSig | -0.66 | (-1.19--0.13) | **1.39E-02** | **2.33E-02** |
| TG(O-50:1) [NL-17:1] | -0.65 | (-1.62-0.34) | 1.97E-01 | 2.76E-01 | notSig | -0.10 | (-0.75-0.55) | 7.61E-01 | 7.99E-01 | -0.61 | (-1.19--0.03) | **3.78E-02** | 7.30E-02 | GUSTO only | -1.25 | (-1.69--0.8) | **4.46E-08** | **1.33E-07** |
| TG(O-50:1) [NL-18:1] | -0.83 | (-1.86-0.21) | 1.16E-01 | 1.79E-01 | notSig | -0.68 | (-1.23--0.12) | **1.70E-02** | **2.53E-02** | -0.86 | (-1.65--0.07) | **3.23E-02** | 6.40E-02 | GUSTO only | -0.93 | (-1.46--0.39) | **7.04E-04** | **1.46E-03** |
| TG(O-50:2) [NL-18:2] | -0.25 | (-1.58-1.1) | 7.14E-01 | 7.76E-01 | notSig | -0.10 | (-0.63-0.43) | 7.25E-01 | 7.70E-01 | -0.57 | (-1.52-0.38) | 2.42E-01 | 3.43E-01 | notSig | -0.50 | (-1.1-0.1) | 1.05E-01 | 1.44E-01 |
| TG(O-52:2) [NL-16:0] | 0.73 | (-0.37-1.85) | 1.91E-01 | 2.71E-01 | notSig | 0.12 | (-0.54-0.77) | 7.27E-01 | 7.70E-01 | -0.04 | (-1.03-0.96) | 9.42E-01 | 9.54E-01 | GUSTO only | -0.62 | (-1.31-0.07) | 7.77E-02 | 1.09E-01 |
| TG(O-52:2) [NL-17:1] | 0.64 | (-0.33-1.62) | 1.99E-01 | 2.78E-01 | notSig | 0.45 | (-0.1-1) | 1.12E-01 | 1.43E-01 | -0.56 | (-1.27-0.14) | 1.18E-01 | 1.86E-01 | GUSTO only | -1.78 | (-2.27--1.29) | **9.10E-13** | **3.95E-12** |
| TG(O-52:2) [NL-18:1] | 0.72 | (-0.36-1.81) | 1.93E-01 | 2.72E-01 | notSig | -0.02 | (-0.68-0.64) | 9.47E-01 | 9.55E-01 | -0.58 | (-1.36-0.19) | 1.38E-01 | 2.12E-01 | GUSTO only | -1.09 | (-1.69--0.49) | **3.34E-04** | **7.27E-04** |
| TG(O-54:2) [NL-18:1] | -0.29 | (-1.33-0.77) | 5.90E-01 | 6.64E-01 | GUSTO only | 0.52 | (-0.05-1.08) | 7.46E-02 | 9.89E-02 | -0.40 | (-1.24-0.45) | 3.57E-01 | 4.63E-01 | GUSTO only | -1.77 | (-2.29--1.24) | **5.95E-11** | **2.17E-10** |
| TG(O-54:4) [NL-17:1] | -0.64 | (-1.98-0.72) | 3.54E-01 | 4.42E-01 | notSig | -0.09 | (-0.76-0.57) | 7.81E-01 | 8.15E-01 | -1.75 | (-3.65-0.14) | 7.01E-02 | 1.23E-01 | GUSTO only | -3.73 | (-4.42--3.04) | **3.57E-26** | **3.45E-25** |
| TG(O-54:4) [NL-18:2] | 1.17 | (-0.64-3.02) | 2.07E-01 | 2.89E-01 | notSig | 0.25 | (-0.49-0.98) | 5.12E-01 | 5.72E-01 | -1.24 | (-2.74-0.27) | 1.07E-01 | 1.70E-01 | GUSTO only | -3.95 | (-4.6--3.31) | **4.99E-33** | **6.43E-32** |
| dhCer(d18:0/22:0) | 0.06 | (-0.7-0.82) | 8.75E-01 | 9.01E-01 | GUSTO only | 0.69 | (0.2-1.18) | **5.76E-03** | **9.49E-03** | 1.24 | (0.7-1.79) | **9.64E-06** | **4.66E-05** | BIS only | 0.74 | (0.31-1.17) | **6.87E-04** | **1.44E-03** |
| dhCer(d18:0/24:0) | -0.60 | (-1.34-0.15) | 1.18E-01 | 1.81E-01 | notSig | -0.10 | (-0.57-0.36) | 6.64E-01 | 7.13E-01 | 0.80 | (0.17-1.43) | **1.31E-02** | **2.90E-02** | BIS only | 0.19 | (-0.27-0.66) | 4.21E-01 | 4.79E-01 |
| dhCer(d18:0/24:1) | 2.18 | (1.31-3.04) | **7.33E-07** | **6.77E-06** | GUSTO and BIS | 1.89 | (1.34-2.44) | **1.64E-11** | **1.28E-10** | 1.21 | (0.52-1.9) | **5.91E-04** | **1.93E-03** | BIS only | 0.63 | (0.1-1.17) | **2.11E-02** | **3.44E-02** |
| Cer(d16:1/22:0) | -0.82 | (-1.81-0.17) | 1.04E-01 | 1.63E-01 | GUSTO only | -0.75 | (-1.27--0.24) | **3.98E-03** | **6.81E-03** | -0.13 | (-0.89-0.63) | 7.36E-01 | 8.01E-01 | GUSTO only | -0.59 | (-1.12--0.05) | **3.11E-02** | **4.91E-02** |
| Cer(d16:1/23:0) | -1.81 | (-2.87--0.73) | **1.07E-03** | **3.34E-03** | GUSTO and BIS | -1.22 | (-1.79--0.65) | **2.51E-05** | **6.69E-05** | -0.46 | (-1.25-0.33) | 2.49E-01 | 3.49E-01 | notSig | -0.64 | (-1.31-0.03) | 6.07E-02 | 8.70E-02 |
| Cer(d16:1/24:0) | -2.27 | (-3.24--1.28) | **8.83E-06** | **5.94E-05** | GUSTO and BIS | -2.10 | (-2.63--1.57) | **6.69E-15** | **8.63E-14** | -0.23 | (-0.91-0.45) | 5.09E-01 | 6.06E-01 | notSig | -0.48 | (-0.98-0.02) | 5.95E-02 | 8.54E-02 |
| Cer(d16:1/24:1) | 0.99 | (0.06-1.93) | **3.63E-02** | 6.69E-02 | notSig | 0.03 | (-0.49-0.55) | 9.04E-01 | 9.21E-01 | 0.32 | (-0.44-1.09) | 4.05E-01 | 5.07E-01 | notSig | -0.02 | (-0.61-0.56) | 9.37E-01 | 9.45E-01 |
| Cer(d17:1/22:0) | 0.01 | (-0.89-0.93) | 9.77E-01 | 9.82E-01 | GUSTO only | -0.83 | (-1.36--0.3) | **2.04E-03** | **3.64E-03** | -0.16 | (-0.8-0.49) | 6.35E-01 | 7.17E-01 | notSig | -0.37 | (-0.9-0.17) | 1.79E-01 | 2.29E-01 |
| Cer(d17:1/23:0) | -0.62 | (-1.66-0.42) | 2.39E-01 | 3.26E-01 | GUSTO only | -1.09 | (-1.65--0.52) | **1.72E-04** | **3.87E-04** | -0.57 | (-1.37-0.23) | 1.63E-01 | 2.46E-01 | notSig | -0.74 | (-1.44--0.04) | **3.75E-02** | 5.75E-02 |
| Cer(d17:1/24:0) | -0.87 | (-1.76-0.03) | 5.68E-02 | 1.00E-01 | GUSTO only | -1.80 | (-2.28--1.32) | **2.78E-13** | **2.67E-12** | -0.51 | (-1.11-0.08) | 8.98E-02 | 1.47E-01 | GUSTO only | -0.73 | (-1.19--0.26) | **2.23E-03** | **4.27E-03** |
| Cer(d17:1/24:1) | 1.23 | (0.34-2.12) | **6.90E-03** | **1.66E-02** | GUSTO and BIS | -0.32 | (-0.83-0.19) | 2.18E-01 | 2.64E-01 | -0.05 | (-0.77-0.66) | 8.80E-01 | 9.07E-01 | notSig | -0.41 | (-0.96-0.13) | 1.38E-01 | 1.83E-01 |
| Cer(d18:1/16:0) | -0.12 | (-0.75-0.5) | 7.00E-01 | 7.64E-01 | GUSTO only | -0.60 | (-0.97--0.23) | **1.51E-03** | **2.73E-03** | -0.25 | (-0.99-0.49) | 5.04E-01 | 6.06E-01 | notSig | -0.41 | (-0.87-0.05) | 8.23E-02 | 1.15E-01 |
| Cer(d18:1/18:0) | 2.86 | (1.89-3.83) | **6.98E-09** | **1.35E-07** | GUSTO and BIS | 2.26 | (1.71-2.81) | **9.19E-16** | **1.47E-14** | -0.23 | (-0.99-0.53) | 5.53E-01 | 6.37E-01 | notSig | -0.48 | (-0.99-0.03) | 6.59E-02 | 9.35E-02 |
| Cer(d18:1/20:0) | 1.49 | (0.58-2.41) | **1.28E-03** | **3.87E-03** | BIS only | 0.58 | (0.1-1.06) | **1.82E-02** | **2.66E-02** | 0.09 | (-0.64-0.82) | 8.06E-01 | 8.64E-01 | notSig | -0.37 | (-0.85-0.12) | 1.42E-01 | 1.88E-01 |
| Cer(d18:1/22:0) | 0.32 | (-0.42-1.08) | 3.94E-01 | 4.79E-01 | notSig | 0.04 | (-0.37-0.44) | 8.64E-01 | 8.85E-01 | 0.80 | (0.08-1.51) | **2.95E-02** | 5.92E-02 | notSig | 0.29 | (-0.18-0.76) | 2.26E-01 | 2.80E-01 |
| Cer(d18:1/23:0) | -0.19 | (-0.98-0.6) | 6.37E-01 | 7.06E-01 | notSig | -0.34 | (-0.75-0.07) | 1.04E-01 | 1.35E-01 | -0.29 | (-1.19-0.61) | 5.26E-01 | 6.17E-01 | notSig | -0.36 | (-0.95-0.23) | 2.33E-01 | 2.87E-01 |
| Cer(d18:1/24:0) | -0.56 | (-1.26-0.15) | 1.25E-01 | 1.88E-01 | GUSTO only | -0.83 | (-1.22--0.44) | **2.71E-05** | **7.11E-05** | 0.52 | (-0.1-1.15) | 1.02E-01 | 1.65E-01 | notSig | 0.06 | (-0.38-0.5) | 7.89E-01 | 8.30E-01 |
| Cer(d18:1/24:1) | 2.15 | (1.4-2.9) | **2.24E-08** | **3.83E-07** | BIS only | 0.77 | (0.33-1.2) | **5.12E-04** | **1.01E-03** | 0.71 | (-0.04-1.47) | 6.48E-02 | 1.15E-01 | notSig | 0.14 | (-0.36-0.64) | 5.90E-01 | 6.48E-01 |
| Cer(d18:2/16:0) | 0.45 | (-0.15-1.05) | 1.39E-01 | 2.08E-01 | notSig | 0.05 | (-0.39-0.48) | 8.32E-01 | 8.60E-01 | 0.42 | (-0.32-1.15) | 2.63E-01 | 3.63E-01 | notSig | 0.32 | (-0.26-0.91) | 2.82E-01 | 3.40E-01 |
| Cer(d18:2/22:0) | 0.75 | (-0.07-1.58) | 7.26E-02 | 1.21E-01 | notSig | 0.00 | (-0.47-0.47) | 1.00E+00 | 1.00E+00 | 0.74 | (0.08-1.4) | **2.84E-02** | 5.73E-02 | notSig | 0.29 | (-0.2-0.78) | 2.46E-01 | 3.00E-01 |
| Cer(d18:2/23:0) | 0.31 | (-0.57-1.19) | 4.93E-01 | 5.72E-01 | notSig | -0.29 | (-0.78-0.2) | 2.51E-01 | 3.00E-01 | -0.14 | (-0.98-0.69) | 7.37E-01 | 8.01E-01 | notSig | -0.25 | (-0.89-0.39) | 4.43E-01 | 4.99E-01 |
| Cer(d18:2/24:0) | -0.71 | (-1.51-0.1) | 8.56E-02 | 1.39E-01 | GUSTO only | -1.08 | (-1.53--0.62) | **3.50E-06** | **1.08E-05** | 0.40 | (-0.28-1.09) | 2.49E-01 | 3.49E-01 | notSig | -0.18 | (-0.68-0.31) | 4.63E-01 | 5.15E-01 |
| Cer(d18:2/24:1) | 2.56 | (1.72-3.4) | **2.06E-09** | **4.77E-08** | BIS only | 0.79 | (0.32-1.27) | **1.05E-03** | **1.94E-03** | 0.98 | (0.28-1.68) | **6.30E-03** | **1.57E-02** | BIS only | 0.52 | (0.01-1.04) | **4.71E-02** | 7.03E-02 |
| Cer(d19:1/24:0) | 0.85 | (-0.11-1.81) | 8.16E-02 | 1.33E-01 | GUSTO only | -0.26 | (-0.91-0.4) | 4.43E-01 | 5.03E-01 | -0.06 | (-0.7-0.58) | 8.49E-01 | 8.91E-01 | GUSTO only | -0.48 | (-0.99-0.03) | 6.26E-02 | 8.94E-02 |
| Cer(d19:1/24:1) | 3.55 | (2.42-4.7) | **8.62E-10** | **2.34E-08** | BIS only | 1.45 | (0.68-2.22) | **2.30E-04** | **4.93E-04** | 0.19 | (-0.62-0.99) | 6.53E-01 | 7.31E-01 | notSig | -0.35 | (-0.99-0.29) | 2.80E-01 | 3.38E-01 |
| Cer(m18:0/22:0) | -0.83 | (-2.06-0.41) | 1.89E-01 | 2.68E-01 | notSig | -0.06 | (-0.72-0.6) | 8.57E-01 | 8.82E-01 | -0.33 | (-1.13-0.48) | 4.25E-01 | 5.26E-01 | GUSTO only | -0.70 | (-1.32--0.09) | **2.55E-02** | **4.10E-02** |
| Cer(m18:0/23:0) | -1.40 | (-2.61--0.17) | **2.60E-02** | 5.09E-02 | notSig | -0.20 | (-0.91-0.51) | 5.80E-01 | 6.36E-01 | -1.06 | (-1.98--0.13) | **2.54E-02** | 5.17E-02 | notSig | -1.11 | (-1.94--0.28) | **8.89E-03** | **1.55E-02** |
| Cer(m18:0/24:0) | -2.13 | (-3.31--0.92) | **5.83E-04** | **2.10E-03** | BIS only | -1.07 | (-1.73--0.42) | **1.33E-03** | **2.43E-03** | -0.62 | (-1.4-0.16) | 1.19E-01 | 1.86E-01 | GUSTO only | -0.91 | (-1.46--0.37) | **1.08E-03** | **2.20E-03** |
| Cer(m18:0/24:1) | 0.66 | (-0.57-1.9) | 2.96E-01 | 3.87E-01 | GUSTO only | 0.98 | (0.26-1.7) | **7.82E-03** | **1.25E-02** | -0.53 | (-1.45-0.39) | 2.60E-01 | 3.63E-01 | GUSTO only | -0.95 | (-1.63--0.26) | **6.81E-03** | **1.21E-02** |
| Cer(m18:1/22:0) | 0.04 | (-1.12-1.21) | 9.50E-01 | 9.64E-01 | notSig | 0.43 | (-0.26-1.11) | 2.26E-01 | 2.72E-01 | -0.75 | (-1.54-0.05) | 6.51E-02 | 1.15E-01 | GUSTO only | -1.76 | (-2.28--1.24) | **3.29E-11** | **1.24E-10** |
| Cer(m18:1/23:0) | -0.58 | (-1.76-0.62) | 3.41E-01 | 4.30E-01 | notSig | -0.10 | (-0.83-0.62) | 7.75E-01 | 8.10E-01 | -1.59 | (-2.57--0.61) | **1.46E-03** | **4.34E-03** | GUSTO and BIS | -2.20 | (-3--1.41) | **5.32E-08** | **1.57E-07** |
| Cer(m18:1/24:0) | -1.07 | (-2.13-0.01) | 5.17E-02 | 9.19E-02 | notSig | -0.65 | (-1.28--0.02) | **4.47E-02** | 6.18E-02 | -0.93 | (-1.66--0.2) | **1.21E-02** | **2.75E-02** | GUSTO and BIS | -1.82 | (-2.32--1.33) | **4.92E-13** | **2.24E-12** |
| Cer(m18:1/24:1) | 1.69 | (0.52-2.88) | **4.75E-03** | **1.22E-02** | GUSTO and BIS | 1.49 | (0.77-2.2) | **4.63E-05** | **1.14E-04** | -0.84 | (-1.68--0.01) | **4.75E-02** | 9.00E-02 | GUSTO only | -1.53 | (-2.1--0.95) | **2.07E-07** | **5.95E-07** |
| GM3(d18:1/18:0) | -0.02 | (-0.71-0.69) | 9.64E-01 | 9.70E-01 | GUSTO only | -0.74 | (-1.16--0.31) | **6.75E-04** | **1.29E-03** | -0.21 | (-0.72-0.3) | 4.26E-01 | 5.26E-01 | GUSTO only | -0.44 | (-0.83--0.05) | **2.56E-02** | **4.11E-02** |

| | | | | | | | | | | | | | | | | | | |
|---|---|---|---|---|---|---|---|---|---|---|---|---|---|---|---|---|---|---|
| GM3(d18:1/22:0) | -0.33 | (-1.15-0.5) | 4.39E-01 | 5.25E-01 | GUSTO only | -1.50 | (-2--1.01) | **2.53E-09** | **1.37E-08** | 0.64 | (0.01-1.28) | **4.78E-02** | 9.02E-02 | notSig | 0.49 | (-0.05-1.03) | 7.66E-02 | 1.07E-01 |
| GM3(d18:1/24:0) | -1.84 | (-2.77--0.89) | **1.55E-04** | **6.37E-04** | GUSTO and BIS | -2.77 | (-3.29--2.26) | **4.58E-26** | **4.25E-24** | 0.66 | (-0.03-1.34) | 6.13E-02 | 1.11E-01 | notSig | 0.17 | (-0.35-0.69) | 5.23E-01 | 5.78E-01 |
| GM3(d18:1/24:1) | 0.52 | (-0.34-1.39) | 2.33E-01 | 3.19E-01 | GUSTO only | -1.03 | (-1.52--0.54) | **3.56E-05** | **8.91E-05** | 0.63 | (-0.08-1.33) | 8.00E-02 | 1.35E-01 | notSig | 0.33 | (-0.2-0.87) | 2.25E-01 | 2.80E-01 |
| HexCer(d16:1/22:0) | -1.53 | (-2.4--0.64) | **7.70E-04** | **2.57E-03** | GUSTO and BIS | -1.66 | (-2.18--1.14) | **2.92E-10** | **1.78E-09** | 0.04 | (-0.72-0.8) | 9.15E-01 | 9.33E-01 | notSig | -0.14 | (-0.8-0.52) | 6.77E-01 | 7.26E-01 |
| HexCer(d16:1/24:0) | -2.20 | (-3.04--1.34) | **6.57E-07** | **6.22E-06** | GUSTO and BIS | -2.85 | (-3.36--2.33) | **4.63E-27** | **5.37E-25** | -0.22 | (-0.88-0.43) | 5.02E-01 | 6.05E-01 | notSig | -0.37 | (-0.94-0.2) | 2.04E-01 | 2.58E-01 |
| HexCer(d18:1/16:0) | -0.69 | (-1.48-0.12) | 9.39E-02 | 1.50E-01 | GUSTO only | -1.15 | (-1.56--0.73) | **6.18E-08** | **2.68E-07** | 0.77 | (-0.04-1.59) | 6.35E-02 | 1.14E-01 | notSig | 0.38 | (-0.2-0.97) | 2.00E-01 | 2.54E-01 |
| HexCer(d18:1/18:0) | -0.24 | (-1.08-0.6) | 5.71E-01 | 6.49E-01 | GUSTO only | -1.12 | (-1.61--0.64) | **4.85E-06** | **1.43E-05** | 0.52 | (-0.31-1.36) | 2.17E-01 | 3.13E-01 | notSig | 0.43 | (-0.22-1.09) | 1.95E-01 | 2.48E-01 |
| HexCer(d18:1/20:0) | -0.19 | (-1-0.62) | 6.41E-01 | 7.08E-01 | GUSTO only | -1.30 | (-1.77--0.84) | **3.73E-08** | **1.65E-07** | 0.82 | (0.13-1.52) | **2.02E-02** | **4.27E-02** | BIS only | 0.56 | (0-1.13) | 5.02E-02 | 7.41E-02 |
| HexCer(d18:1/22:0) | -0.93 | (-1.75--0.11) | **2.73E-02** | 5.29E-02 | GUSTO only | -1.67 | (-2.09--1.24) | **1.38E-14** | **1.73E-13** | 0.58 | (-0.08-1.25) | 8.55E-02 | 1.42E-01 | notSig | 0.55 | (0.04-1.05) | **3.43E-02** | 5.36E-02 |
| HexCer(d18:1/24:0) | -1.88 | (-2.65--1.1) | **3.16E-06** | **2.48E-05** | GUSTO and BIS | -2.38 | (-2.81--1.96) | **2.66E-28** | **4.11E-26** | 0.43 | (-0.2-1.06) | 1.81E-01 | 2.70E-01 | notSig | 0.28 | (-0.2-0.77) | 2.53E-01 | 3.08E-01 |
| HexCer(d18:1/24:1) | 0.03 | (-0.79-0.85) | 9.46E-01 | 9.64E-01 | GUSTO only | -1.14 | (-1.6--0.69) | **1.02E-06** | **3.25E-06** | 0.95 | (0.32-1.58) | **3.02E-03** | **8.10E-03** | BIS only | 0.80 | (0.31-1.3) | **1.40E-03** | **2.76E-03** |
| Hex2Cer(d16:1/16:0) | -1.54 | (-2.44--0.63) | **9.65E-04** | **3.07E-03** | GUSTO and BIS | -1.38 | (-1.89--0.86) | **1.56E-07** | **6.30E-07** | -0.60 | (-1.45-0.24) | 1.62E-01 | 2.46E-01 | notSig | -0.78 | (-1.43--0.13) | **1.95E-02** | **3.22E-02** |
| Hex2Cer(d18:1/16:0) | -1.32 | (-2.07--0.57) | **6.43E-04** | **2.23E-03** | GUSTO and BIS | -1.49 | (-1.92--1.06) | **9.38E-12** | **7.64E-11** | 0.04 | (-0.72-0.81) | 9.10E-01 | 9.30E-01 | notSig | -0.27 | (-0.79-0.25) | 3.08E-01 | 3.66E-01 |
| Hex2Cer(d18:1/22:0) | -1.01 | (-1.72--0.3) | **5.74E-03** | **1.42E-02** | GUSTO and BIS | -1.46 | (-1.9--1.02) | **7.22E-11** | **4.93E-10** | 0.31 | (-0.35-0.98) | 3.56E-01 | 4.63E-01 | notSig | 0.13 | (-0.41-0.66) | 6.38E-01 | 6.88E-01 |
| Hex2Cer(d18:1/24:1) | -0.33 | (-1.2-0.54) | 4.53E-01 | 5.34E-01 | GUSTO only | -1.25 | (-1.74--0.76) | **5.00E-07** | **1.79E-06** | 0.64 | (-0.13-1.41) | 1.05E-01 | 1.69E-01 | notSig | 0.27 | (-0.29-0.83) | 3.45E-01 | 4.03E-01 |
| Hex2Cer(d18:2/16:0) | -0.40 | (-1.18-0.39) | 3.22E-01 | 4.14E-01 | GUSTO only | -0.58 | (-1.03--0.13) | **1.10E-02** | **1.70E-02** | 0.11 | (-0.58-0.79) | 7.57E-01 | 8.19E-01 | notSig | -0.14 | (-0.66-0.38) | 5.94E-01 | 6.52E-01 |
| Hex3Cer(d18:1/16:0) | -0.40 | (-0.95-0.16) | 1.58E-01 | 2.31E-01 | GUSTO only | -0.81 | (-1.14--0.47) | **2.22E-06** | **6.97E-06** | 0.01 | (-0.48-0.49) | 9.79E-01 | 9.79E-01 | notSig | -0.19 | (-0.56-0.19) | 3.31E-01 | 3.90E-01 |
| Hex3Cer(d18:1/22:0) | -1.08 | (-1.75--0.4) | **1.90E-03** | **5.59E-03** | GUSTO and BIS | -1.33 | (-1.78--0.88) | **6.73E-09** | **3.32E-08** | 0.01 | (-0.65-0.67) | 9.71E-01 | 9.73E-01 | notSig | 0.05 | (-0.5-0.6) | 8.47E-01 | 8.83E-01 |
| Hex3Cer(d18:1/24:0) | -1.93 | (-2.71--1.15) | **1.79E-06** | **1.49E-05** | GUSTO and BIS | -1.70 | (-2.16--1.23) | **7.77E-13** | **7.07E-12** | -0.22 | (-0.88-0.44) | 5.11E-01 | 6.06E-01 | notSig | -0.32 | (-0.82-0.19) | 2.19E-01 | 2.73E-01 |
| Hex3Cer(d18:1/24:1) | -0.24 | (-0.86-0.38) | 4.48E-01 | 5.34E-01 | GUSTO only | -0.74 | (-1.15--0.33) | **4.43E-04** | **8.90E-04** | 0.24 | (-0.35-0.84) | 4.19E-01 | 5.20E-01 | notSig | -0.05 | (-0.52-0.42) | 8.32E-01 | 8.71E-01 |
| SM(d17:1/14:0) | -1.42 | (-2.13--0.7) | **1.29E-04** | **5.48E-04** | GUSTO and BIS | -1.70 | (-2.15--1.25) | **9.43E-14** | **9.72E-13** | -0.42 | (-1.05-0.22) | 1.98E-01 | 2.88E-01 | GUSTO only | -0.75 | (-1.15--0.35) | **2.76E-04** | **6.11E-04** |
| SM(d18:0/14:0) | -1.55 | (-2.31--0.78) | **8.10E-05** | **3.72E-04** | GUSTO and BIS | -1.11 | (-1.54--0.67) | **6.32E-07** | **2.19E-06** | 0.89 | (0.31-1.46) | **2.43E-03** | **6.76E-03** | BIS only | 0.26 | (-0.12-0.65) | 1.80E-01 | 2.30E-01 |
| SM(d18:1/14:0/SM(d16:1/16:0) | -1.10 | (-1.73--0.47) | **6.97E-04** | **2.36E-03** | GUSTO and BIS | -1.05 | (-1.43--0.67) | **5.15E-08** | **2.26E-07** | 0.08 | (-0.57-0.72) | 8.17E-01 | 8.70E-01 | notSig | -0.31 | (-0.71-0.09) | 1.27E-01 | 1.72E-01 |
| SM(d18:2/14:0) | -0.28 | (-0.89-0.34) | 3.76E-01 | 4.67E-01 | notSig | -0.01 | (-0.4-0.38) | 9.66E-01 | 9.72E-01 | 0.75 | (0.16-1.34) | **1.23E-02** | **2.76E-02** | BIS only | 0.39 | (-0.01-0.79) | 5.77E-02 | 8.32E-02 |
| SM(d17:1/16:0) | -0.17 | (-0.81-0.47) | 5.97E-01 | 6.69E-01 | GUSTO only | -0.99 | (-1.37--0.62) | **1.39E-07** | **5.86E-07** | -0.43 | (-1.11-0.25) | 2.17E-01 | 3.13E-01 | GUSTO only | -0.79 | (-1.19--0.38) | **1.33E-04** | **3.06E-04** |
| SM(d18:1/16:0) | -0.07 | (-0.49-0.35) | 7.40E-01 | 8.01E-01 | GUSTO only | -0.60 | (-0.93--0.26) | **4.33E-04** | **8.74E-04** | 0.33 | (-0.35-1.01) | 3.45E-01 | 4.51E-01 | notSig | -0.05 | (-0.5-0.4) | 8.28E-01 | 8.69E-01 |
| SM(d18:2/16:0) | 0.87 | (0.23-1.52) | **7.72E-03** | **1.82E-02** | BIS only | 0.30 | (-0.04-0.64) | 8.61E-02 | 1.13E-01 | 0.84 | (0.21-1.47) | **9.23E-03** | **2.21E-02** | BIS only | 0.38 | (-0.02-0.78) | 6.45E-02 | 9.18E-02 |
| SM(34:3) | 0.77 | (0.21-1.33) | **7.00E-03** | **1.67E-02** | BIS only | 0.44 | (0.08-0.8) | **1.79E-02** | **2.64E-02** | 0.62 | (0.04-1.2) | **3.50E-02** | 6.90E-02 | notSig | 0.37 | (-0.03-0.78) | 7.18E-02 | 1.01E-01 |
| SM(d18:2/17:0) | 1.10 | (0.37-1.83) | **2.94E-03** | **8.08E-03** | BIS only | 0.06 | (-0.33-0.45) | 7.71E-01 | 8.08E-01 | -0.28 | (-1-0.44) | 4.40E-01 | 5.40E-01 | GUSTO only | -0.70 | (-1.12--0.28) | **1.01E-03** | **2.05E-03** |
| SM(35:2) (b) | -0.08 | (-0.77-0.61) | 8.19E-01 | 8.62E-01 | GUSTO only | -0.92 | (-1.37--0.47) | **5.41E-05** | **1.31E-04** | 0.45 | (-0.67-1.57) | 4.34E-01 | 5.34E-01 | notSig | -0.33 | (-0.95-0.29) | 2.97E-01 | 3.57E-01 |
| SM(d18:1/18:0/SM(d16:1/20:0) | 1.36 | (0.58-2.14) | **6.33E-04** | **2.21E-03** | BIS only | 0.58 | (0.15-1) | **7.96E-03** | **1.27E-02** | 0.86 | (0.16-1.57) | **1.66E-02** | **3.61E-02** | BIS only | 0.33 | (-0.13-0.78) | 1.56E-01 | 2.03E-01 |
| SM(d18:2/18:1) | 1.51 | (0.81-2.22) | **2.36E-05** | **1.33E-04** | BIS only | 0.73 | (0.33-1.12) | **2.99E-04** | **6.28E-04** | 0.51 | (-0.23-1.24) | 1.77E-01 | 2.65E-01 | notSig | 0.20 | (-0.27-0.66) | 4.07E-01 | 4.65E-01 |
| SM(37:1) | 0.72 | (0.06-1.39) | **3.23E-02** | 6.17E-02 | GUSTO only | -0.16 | (-0.58-0.26) | 4.63E-01 | 5.19E-01 | -0.24 | (-0.98-0.51) | 5.31E-01 | 6.18E-01 | GUSTO only | -0.57 | (-1.06--0.08) | **2.27E-02** | **3.69E-02** |
| SM(37:2) | 1.13 | (0.41-1.86) | **2.12E-03** | **6.10E-03** | GUSTO and BIS | -0.25 | (-0.71-0.22) | 2.99E-01 | 3.49E-01 | -0.03 | (-0.78-0.72) | 9.37E-01 | 9.51E-01 | GUSTO only | -0.46 | (-0.94-0.01) | 5.63E-02 | 8.17E-02 |
| SM(d18:1/20:0)/SM(d16:1/22:0) | -0.52 | (-1.21-0.19) | 1.49E-01 | 2.20E-01 | GUSTO only | -0.69 | (-1.05--0.33) | **1.72E-04** | **3.87E-04** | 0.95 | (0.29-1.6) | **4.51E-03** | **1.18E-02** | BIS only | 0.56 | (0.14-0.98) | **9.74E-03** | **1.67E-02** |
| SM(d18:2/20:0) | 0.62 | (0.02-1.23) | **4.38E-02** | 7.91E-02 | notSig | 0.04 | (-0.35-0.43) | 8.44E-01 | 8.70E-01 | 1.12 | (0.5-1.74) | **3.98E-04** | **1.35E-03** | BIS only | 0.80 | (0.32-1.28) | **1.13E-03** | **2.26E-03** |
| SM(38:3) (a) | 0.95 | (0.22-1.68) | **1.03E-02** | **2.35E-02** | BIS only | 0.63 | (0.23-1.04) | **2.25E-03** | **4.00E-03** | 0.02 | (-0.59-0.63) | 9.52E-01 | 9.60E-01 | notSig | -0.27 | (-0.69-0.14) | 1.91E-01 | 2.44E-01 |
| SM(38:3) (b) | 0.82 | (0.14-1.51) | **1.79E-02** | **3.72E-02** | GUSTO and BIS | -0.14 | (-0.57-0.29) | 5.33E-01 | 5.95E-01 | 0.49 | (-0.19-1.16) | 1.57E-01 | 2.39E-01 | notSig | 0.23 | (-0.27-0.73) | 3.61E-01 | 4.17E-01 |
| SM(d16:1/23:0)/SM(d17:1/22:0) | -0.81 | (-1.51--0.11) | **2.40E-02** | **4.72E-02** | GUSTO and BIS | -1.00 | (-1.39--0.61) | **4.10E-07** | **1.50E-06** | -0.17 | (-0.93-0.59) | 6.58E-01 | 7.31E-01 | notSig | -0.30 | (-0.74-0.13) | 1.70E-01 | 2.20E-01 |
| SM(d18:1/22:0)/SM(d16:1/24:0) | -0.35 | (-0.91-0.21) | 2.15E-01 | 2.98E-01 | GUSTO only | -0.50 | (-0.85--0.14) | **6.67E-03** | **1.08E-02** | 0.93 | (0.32-1.54) | **2.75E-03** | **7.50E-03** | BIS only | 0.65 | (0.23-1.07) | **2.40E-03** | **4.59E-03** |
| SM(d16:1/24:1) | 0.46 | (-0.31-1.23) | 2.43E-01 | 3.30E-01 | GUSTO only | -0.35 | (-0.77-0.08) | 1.10E-01 | 1.40E-01 | 0.95 | (0.18-1.72) | **1.57E-02** | **3.43E-02** | BIS only | 0.29 | (-0.19-0.77) | 2.30E-01 | 2.84E-01 |
| SM(d18:2/22:0) | -0.11 | (-0.77-0.56) | 7.48E-01 | 8.08E-01 | GUSTO only | -0.57 | (-0.95--0.19) | **3.04E-03** | **5.26E-03** | 1.12 | (0.41-1.82) | **2.03E-03** | **5.84E-03** | BIS only | 0.70 | (0.24-1.17) | **3.20E-03** | **6.06E-03** |
| SM(40:3) (a) | 0.84 | (0.17-1.52) | **1.46E-02** | **3.14E-02** | BIS only | 0.20 | (-0.19-0.6) | 3.15E-01 | 3.62E-01 | 0.55 | (-0.11-1.21) | 1.03E-01 | 1.66E-01 | notSig | 0.00 | (-0.44-0.44) | 9.98E-01 | 9.99E-01 |
| SM(40:3) (b) | 0.28 | (-0.35-0.91) | 3.88E-01 | 4.75E-01 | GUSTO only | -0.40 | (-0.81-0.02) | 5.95E-02 | 8.08E-02 | -0.34 | (-1.01-0.33) | 3.14E-01 | 4.17E-01 | GUSTO only | -0.64 | (-1.1--0.18) | **6.75E-03** | **1.20E-02** |
| SM(41:0) | -0.44 | (-1.14-0.28) | 2.30E-01 | 3.17E-01 | notSig | -0.42 | (-0.79--0.06) | **2.26E-02** | **3.26E-02** | 0.07 | (-0.52-0.66) | 8.09E-01 | 8.65E-01 | notSig | -0.10 | (-0.59-0.4) | 6.99E-01 | 7.46E-01 |
| SM(d17:1/24:1) | 1.08 | (0.45-1.71) | **7.28E-04** | **2.45E-03** | GUSTO and BIS | -0.20 | (-0.57-0.17) | 2.90E-01 | 3.40E-01 | -0.05 | (-0.72-0.61) | 8.76E-01 | 9.06E-01 | notSig | -0.30 | (-0.73-0.13) | 1.66E-01 | 2.16E-01 |
| SM(d18:2/23:0) | -0.44 | (-1.06-0.18) | 1.65E-01 | 2.40E-01 | GUSTO only | -0.77 | (-1.13--0.42) | **1.63E-05** | **4.54E-05** | 0.12 | (-0.67-0.9) | 7.74E-01 | 8.31E-01 | notSig | 0.04 | (-0.48-0.56) | 8.69E-01 | 8.96E-01 |
| SM(d18:1/24:0) | -0.89 | (-1.51--0.26) | **5.55E-03** | **1.38E-02** | GUSTO and BIS | -1.26 | (-1.63--0.9) | **1.68E-11** | **1.28E-10** | 0.41 | (-0.15-0.98) | 1.51E-01 | 2.31E-01 | notSig | 0.09 | (-0.31-0.5) | 6.52E-01 | 7.01E-01 |
| SM(d18:1/24:1) | 1.63 | (0.99-2.28) | **7.44E-07** | **6.77E-06** | BIS only | 0.34 | (-0.08-0.75) | 1.10E-01 | 1.40E-01 | 1.08 | (0.42-1.74) | **1.41E-03** | **4.28E-03** | BIS only | 0.61 | (0.16-1.05) | **7.53E-03** | **1.32E-02** |
| SM(d18:2/24:0) | -0.90 | (-1.47--0.32) | **2.29E-03** | **6.51E-03** | GUSTO and BIS | -1.28 | (-1.61--0.95) | **6.62E-14** | **7.15E-13** | 0.33 | (-0.26-0.91) | 2.72E-01 | 3.72E-01 | notSig | 0.10 | (-0.31-0.51) | 6.21E-01 | 6.75E-01 |
| SM(43:1) | -0.30 | (-0.92-0.33) | 3.52E-01 | 4.40E-01 | GUSTO only | -1.13 | (-1.51--0.75) | **4.63E-09** | **2.33E-08** | -0.82 | (-1.44--0.2) | **9.72E-03** | **2.30E-02** | GUSTO and BIS | -0.91 | (-1.39--0.43) | **1.78E-04** | **4.01E-04** |
| SM(44:1) | -0.70 | (-1.29--0.11) | **2.04E-02** | **4.08E-02** | GUSTO and BIS | -1.29 | (-1.63--0.95) | **1.41E-13** | **1.39E-12** | -0.28 | (-0.86-0.29) | 3.34E-01 | 4.37E-01 | notSig | -0.40 | (-0.78--0.01) | **4.33E-02** | 6.53E-02 |
| SM(44:2) | 1.12 | (0.47-1.76) | **6.60E-04** | **2.25E-03** | GUSTO and BIS | -0.29 | (-0.68-0.1) | 1.49E-01 | 1.86E-01 | 0.42 | (-0.11-0.94) | 1.20E-01 | 1.86E-01 | notSig | 0.10 | (-0.27-0.47) | 6.00E-01 | 6.55E-01 |
| LPC(14:0) [sn2] | -2.10 | (-3.31--0.87) | **8.49E-04** | **2.79E-03** | BIS only | -1.12 | (-1.78--0.47) | **7.98E-04** | **1.51E-03** | 1.68 | (0.89-2.47) | **3.30E-05** | **1.46E-04** | GUSTO and BIS | 1.73 | (1.3-2.15) | **1.87E-15** | **9.76E-15** |
| LPC(14:0) [sn1] | -1.99 | (-3.28--0.68) | **3.01E-03** | **8.16E-03** | BIS only | -0.81 | (-1.46--0.16) | **1.44E-02** | **2.17E-02** | 1.55 | (0.71-2.39) | **2.92E-04** | **1.04E-03** | GUSTO and BIS | 1.60 | (1.18-2.02) | **8.50E-14** | **4.07E-13** |

| | | | | | | | | | | | | | | | | | | |
|---|---|---|---|---|---|---|---|---|---|---|---|---|---|---|---|---|---|---|
| LPC(15:0) [sn2] | -0.51 | (-1.43-0.42) | 2.83E-01 | 3.74E-01 | GUSTO only | -1.19 | (-1.71--0.68) | **5.38E-06** | **1.57E-05** | 0.55 | (-0.08-1.19) | 8.88E-02 | 1.46E-01 | GUSTO only | 0.81 | (0.37-1.25) | **2.70E-04** | **5.99E-04** |
| LPC(15:0) [sn1] | -0.33 | (-1.25-0.6) | 4.82E-01 | 5.65E-01 | GUSTO only | -1.03 | (-1.5--0.55) | **2.44E-05** | **6.69E-05** | 0.56 | (-0.05-1.17) | 7.21E-02 | 1.25E-01 | GUSTO only | 0.67 | (0.27-1.08) | **1.11E-03** | **2.24E-03** |
| LPC(16:0) [sn2] | 0.65 | (-0.51-1.83) | 2.72E-01 | 3.62E-01 | notSig | 0.07 | (-0.35-0.49) | 7.40E-01 | 7.82E-01 | 1.29 | (0.69-1.9) | **3.05E-05** | **1.36E-04** | GUSTO and BIS | 1.56 | (1.18-1.93) | **4.22E-16** | **2.30E-15** |
| LPC(16:0) [sn1] | 0.53 | (-0.46-1.54) | 2.93E-01 | 3.85E-01 | notSig | 0.32 | (-0.15-0.79) | 1.77E-01 | 2.18E-01 | 1.48 | (0.9-2.07) | **6.48E-07** | **4.18E-06** | GUSTO and BIS | 1.71 | (1.3-2.13) | **3.21E-16** | **1.80E-15** |
| LPC(16:1) [sn2] | 0.61 | (-0.5-1.73) | 2.85E-01 | 3.76E-01 | notSig | 0.30 | (-0.3-0.91) | 3.28E-01 | 3.76E-01 | 2.89 | (1.95-3.83) | **2.73E-09** | **2.53E-08** | GUSTO and BIS | 3.57 | (2.98-4.16) | **1.72E-32** | **2.16E-31** |
| LPC(16:1) [sn1] | 0.73 | (-0.29-1.76) | 1.59E-01 | 2.32E-01 | notSig | 0.45 | (-0.09-0.98) | 1.03E-01 | 1.34E-01 | 2.96 | (2.03-3.89) | **7.73E-10** | **7.77E-09** | GUSTO and BIS | 3.55 | (2.98-4.11) | **5.45E-35** | **8.44E-34** |
| LPC(15-MHDA) [sn2] | 0.45 | (-0.61-1.51) | 4.06E-01 | 4.92E-01 | GUSTO only | -0.74 | (-1.35--0.13) | **1.72E-02** | **2.55E-02** | -0.05 | (-0.69-0.59) | 8.75E-01 | 9.06E-01 | notSig | 0.25 | (-0.28-0.78) | 3.50E-01 | 4.08E-01 |
| LPC(15-MHDA) [sn1] / LPC(17:0) [sn2] | 0.45 | (-0.55-1.46) | 3.78E-01 | 4.68E-01 | GUSTO only | -0.75 | (-1.29--0.21) | **6.36E-03** | **1.04E-02** | -0.22 | (-0.8-0.37) | 4.69E-01 | 5.67E-01 | notSig | 0.09 | (-0.36-0.54) | 6.96E-01 | 7.44E-01 |
| LPC(17:0) [sn1] | 0.29 | (-0.69-1.29) | 5.63E-01 | 6.43E-01 | GUSTO only | -0.65 | (-1.1--0.21) | **4.11E-03** | **7.01E-03** | -0.29 | (-0.84-0.26) | 2.96E-01 | 3.98E-01 | notSig | -0.03 | (-0.4-0.33) | 8.60E-01 | 8.91E-01 |
| LPC(17:1) [sn1] (a) / LPC(17:1) [sn2] (b) | 0.15 | (-0.79-1.09) | 7.59E-01 | 8.13E-01 | GUSTO only | -0.65 | (-1.14--0.16) | **8.94E-03** | **1.41E-02** | 1.44 | (0.79-2.09) | **1.69E-05** | **7.90E-05** | GUSTO and BIS | 1.58 | (1.12-2.04) | **1.91E-11** | **7.44E-11** |
| LPC(18:0) [sn2] | -0.33 | (-1.33-0.68) | 5.17E-01 | 5.94E-01 | notSig | -0.10 | (-0.53-0.32) | 6.37E-01 | 6.89E-01 | 0.89 | (0.35-1.44) | **1.42E-03** | **4.29E-03** | GUSTO and BIS | 0.95 | (0.59-1.3) | **1.84E-07** | **5.35E-07** |
| LPC(18:0) [sn1] | -0.35 | (-1.32-0.63) | 4.84E-01 | 5.66E-01 | notSig | -0.09 | (-0.52-0.33) | 6.69E-01 | 7.16E-01 | 0.85 | (0.33-1.38) | **1.46E-03** | **4.34E-03** | GUSTO and BIS | 0.93 | (0.57-1.28) | **2.85E-07** | **8.18E-07** |
| LPC(18:1) [sn2] | -0.39 | (-1.26-0.5) | 3.91E-01 | 4.77E-01 | GUSTO only | -0.54 | (-0.99--0.09) | **1.89E-02** | **2.75E-02** | 2.84 | (2.25-3.44) | **1.23E-19** | **7.13E-18** | GUSTO and BIS | 3.16 | (2.7-3.62) | **1.57E-41** | **3.82E-40** |
| LPC(18:1) [sn1] | -0.58 | (-1.49-0.33) | 2.12E-01 | 2.94E-01 | notSig | -0.52 | (-0.93--0.1) | **1.41E-02** | **2.14E-02** | 2.62 | (2.09-3.14) | **3.10E-21** | **3.59E-19** | GUSTO and BIS | 2.92 | (2.52-3.33) | **2.29E-45** | **8.85E-44** |
| LPC(18:2) [sn2] | -1.39 | (-2.25--0.52) | **1.86E-03** | **5.49E-03** | GUSTO and BIS | -0.82 | (-1.25--0.4) | **1.53E-04** | **3.47E-04** | 1.91 | (1.24-2.58) | **3.10E-08** | **2.48E-07** | GUSTO and BIS | 2.38 | (1.88-2.87) | **4.09E-21** | **2.88E-20** |
| LPC(18:2) [sn1] | -2.03 | (-3.05--1.01) | **1.15E-04** | **5.02E-04** | BIS only | -0.77 | (-1.26--0.29) | **1.68E-03** | **3.01E-03** | 2.34 | (1.63-3.05) | **1.96E-10** | **2.46E-09** | GUSTO and BIS | 2.63 | (2.11-3.14) | **1.05E-23** | **8.71E-23** |
| LPC(18:3) [sn2] (a) | -2.89 | (-3.97--1.8) | **3.23E-07** | **3.56E-06** | GUSTO and BIS | -1.93 | (-2.52--1.35) | **8.77E-11** | **5.81E-10** | 1.44 | (0.78-2.1) | **1.95E-05** | **8.96E-05** | GUSTO and BIS | 1.86 | (1.39-2.33) | **1.30E-14** | **6.55E-14** |
| LPC(18:3) [sn1] (a)/LPC(18:3) [sn2] (b) | -3.07 | (-4.21--1.91) | **3.18E-07** | **3.56E-06** | GUSTO and BIS | -1.54 | (-2.15--0.94) | **6.54E-07** | **2.25E-06** | 2.68 | (1.97-3.39) | **2.83E-13** | **4.86E-12** | GUSTO and BIS | 3.30 | (2.73-3.87) | **4.86E-30** | **5.64E-29** |
| LPC(18:3) (a) [sn1] [104_sn1] | -3.24 | (-4.4--2.07) | **1.01E-07** | **1.31E-06** | GUSTO and BIS | -1.61 | (-2.24--0.98) | **4.79E-07** | **1.74E-06** | 2.20 | (1.47-2.94) | **5.05E-09** | **4.51E-08** | GUSTO and BIS | 2.94 | (2.37-3.51) | **4.30E-24** | **3.63E-23** |
| LPC(19:0) [sn1] (a) / LPC(19:0) [sn2] (b) | -0.92 | (-1.88-0.05) | 6.18E-02 | 1.08E-01 | GUSTO only | -1.44 | (-1.86--1.02) | **2.14E-11** | **1.58E-10** | -0.48 | (-1.35-0.39) | 2.78E-01 | 3.80E-01 | notSig | -0.33 | (-0.84-0.19) | 2.15E-01 | 2.69E-01 |
| LPC(20:0) [sn1] | -1.33 | (-2.16--0.49) | **1.99E-03** | **5.76E-03** | GUSTO and BIS | -1.53 | (-1.92--1.13) | **3.37E-14** | **3.72E-13** | 0.46 | (-0.04-0.95) | 7.18E-02 | 1.25E-01 | notSig | 0.36 | (0.04-0.69) | **2.75E-02** | **4.39E-02** |
| LPC(20:1) [sn2] | -0.35 | (-1.31-0.61) | 4.75E-01 | 5.57E-01 | GUSTO only | -0.56 | (-0.97--0.14) | **8.81E-03** | **1.40E-02** | 1.81 | (1.3-2.32) | **9.50E-12** | **1.42E-10** | GUSTO and BIS | 1.63 | (1.26-2) | **3.73E-18** | **2.19E-17** |
| LPC(20:1) [sn1] | -0.33 | (-1.29-0.64) | 5.00E-01 | 5.78E-01 | GUSTO only | -0.56 | (-0.98--0.15) | **8.30E-03** | **1.32E-02** | 1.58 | (1.09-2.08) | **5.70E-10** | **5.87E-09** | GUSTO and BIS | 1.67 | (1.32-2.02) | **8.39E-21** | **5.73E-20** |
| LPC(20:2) [sn2] | -1.74 | (-2.63--0.84) | **1.77E-04** | **7.21E-04** | GUSTO and BIS | -1.11 | (-1.55--0.67) | **8.29E-07** | **2.77E-06** | 2.44 | (1.88-2.99) | **5.34E-17** | **1.77E-15** | GUSTO and BIS | 2.89 | (2.44-3.35) | **1.77E-35** | **2.94E-34** |
| LPC(20:2) [sn1] | -1.58 | (-2.61--0.53) | **3.24E-03** | **8.75E-03** | GUSTO and BIS | -0.80 | (-1.23--0.36) | **3.01E-04** | **6.28E-04** | 2.42 | (1.89-2.96) | **8.39E-18** | **3.54E-16** | GUSTO and BIS | 2.77 | (2.34-3.2) | **7.23E-37** | **1.46E-35** |
| LPC(20:3) [sn2] | -0.47 | (-1.38-0.45) | 3.11E-01 | 4.03E-01 | GUSTO only | 0.54 | (0.01-1.07) | **4.59E-02** | 6.33E-02 | 2.08 | (1.43-2.72) | **4.52E-10** | **4.77E-09** | GUSTO and BIS | 2.31 | (1.84-2.78) | **1.04E-21** | **7.53E-21** |
| LPC(20:3) [sn1] | -0.76 | (-1.88-0.37) | 1.87E-01 | 2.68E-01 | GUSTO only | 0.72 | (0.13-1.31) | **1.68E-02** | **2.50E-02** | 2.31 | (1.62-3) | **1.12E-10** | **1.44E-09** | GUSTO and BIS | 2.74 | (2.23-3.26) | **8.02E-26** | **7.59E-25** |
| LPC(20:4) [sn2] | 1.07 | (0.19-1.95) | **1.72E-02** | **3.61E-02** | GUSTO and BIS | 0.91 | (0.46-1.36) | **8.01E-05** | **1.90E-04** | 1.48 | (0.77-2.19) | **4.40E-05** | **1.84E-04** | GUSTO and BIS | 1.97 | (1.49-2.45) | **7.76E-16** | **4.09E-15** |
| LPC(20:4) [sn1] | 1.02 | (-0.05-2.11) | 6.15E-02 | 1.08E-01 | GUSTO only | 1.29 | (0.78-1.79) | **6.11E-07** | **2.13E-06** | 1.68 | (0.89-2.48) | **3.65E-05** | **1.55E-04** | GUSTO and BIS | 2.24 | (1.71-2.76) | **5.90E-17** | **3.34E-16** |
| LPC(20:5) [sn2] | -0.56 | (-1.89-0.78) | 4.12E-01 | 4.96E-01 | notSig | -0.25 | (-1.09-0.6) | 5.65E-01 | 6.24E-01 | 0.81 | (-0.06-1.68) | 6.85E-02 | 1.20E-01 | notSig | 0.68 | (-0.01-1.37) | 5.49E-02 | 7.99E-02 |
| LPC(20:5) [sn1] | -0.74 | (-2.22-0.76) | 3.33E-01 | 4.21E-01 | notSig | -0.12 | (-1.01-0.78) | 8.01E-01 | 8.31E-01 | 1.19 | (0.3-2.09) | **8.96E-03** | **2.16E-02** | BIS only | 0.98 | (0.27-1.68) | **6.65E-03** | **1.19E-02** |
| LPC(22:4) [sn2] | -1.14 | (-2.14--0.13) | **2.65E-02** | 5.17E-02 | notSig | -0.66 | (-1.19--0.12) | **1.61E-02** | **2.41E-02** | 1.74 | (1.19-2.3) | **1.31E-09** | **1.25E-08** | GUSTO and BIS | 2.03 | (1.59-2.47) | **1.55E-19** | **9.70E-19** |
| LPC(22:4) [sn1] | -1.20 | (-2.44-0.06) | 6.15E-02 | 1.08E-01 | notSig | -0.22 | (-0.77-0.34) | 4.42E-01 | 5.03E-01 | 1.82 | (1.21-2.43) | **7.00E-09** | **6.01E-08** | GUSTO and BIS | 2.22 | (1.76-2.67) | **8.20E-22** | **6.04E-21** |
| LPC(22:5) [sn2] (n3) | -1.45 | (-2.43--0.45) | **4.51E-03** | **1.18E-02** | GUSTO and BIS | -1.11 | (-1.64--0.58) | **3.57E-05** | **8.91E-05** | 0.85 | (0.22-1.49) | **8.51E-03** | **2.06E-02** | GUSTO and BIS | 1.16 | (0.65-1.67) | **9.48E-06** | **2.40E-05** |
| LPC(22:5) [sn1] (n3)/LPC(22:5) [sn2] (n6) | -1.50 | (-2.61--0.38) | **8.68E-03** | **2.00E-02** | GUSTO and BIS | -0.98 | (-1.5--0.47) | **1.77E-04** | **3.96E-04** | 1.76 | (1.17-2.35) | **6.68E-09** | **5.84E-08** | GUSTO and BIS | 1.90 | (1.45-2.36) | **4.33E-16** | **2.34E-15** |
| LPC(22:5) [sn1] (n6) | -1.52 | (-2.78--0.24) | **2.03E-02** | **4.08E-02** | BIS only | -0.80 | (-1.46--0.14) | **1.81E-02** | **2.65E-02** | 3.06 | (2.26-3.87) | **2.41E-13** | **4.31E-12** | GUSTO and BIS | 3.50 | (2.83-4.17) | **1.21E-24** | **1.08E-23** |
| LPC(22:5) (n3) [sn1] [104_sn1] | -1.52 | (-2.73--0.29) | **1.54E-02** | **3.28E-02** | BIS only | -0.92 | (-1.52--0.33) | **2.38E-03** | **4.21E-03** | 1.05 | (0.38-1.71) | **2.10E-03** | **6.02E-03** | GUSTO and BIS | 1.04 | (0.53-1.54) | **6.15E-05** | **1.46E-04** |
| LPC(22:6) [sn2] | -0.08 | (-0.94-0.8) | 8.63E-01 | 8.94E-01 | notSig | -0.46 | (-0.97-0.05) | 7.43E-02 | 9.88E-02 | 1.13 | (0.57-1.69) | **7.67E-05** | **2.99E-04** | GUSTO and BIS | 1.32 | (0.89-1.76) | **2.75E-09** | **9.31E-09** |
| LPC(22:6) [sn1] | -0.27 | (-1.38-0.85) | 6.34E-01 | 7.04E-01 | notSig | -0.17 | (-0.73-0.4) | 5.62E-01 | 6.23E-01 | 1.43 | (0.8-2.06) | **9.31E-06** | **4.55E-05** | GUSTO and BIS | 1.70 | (1.22-2.18) | **3.29E-12** | **1.39E-11** |
| LPC(O-16:0) | -0.71 | (-1.72-0.31) | 1.72E-01 | 2.47E-01 | notSig | -0.38 | (-0.78-0.03) | 6.76E-02 | 9.07E-02 | 1.22 | (0.7-1.73) | **3.81E-06** | **2.05E-05** | GUSTO and BIS | 1.47 | (1.11-1.84) | **2.33E-15** | **1.20E-14** |
| LPC(O-18:0) | -1.41 | (-2.38--0.42) | **5.26E-03** | **1.33E-02** | GUSTO and BIS | -1.13 | (-1.61--0.66) | **3.08E-06** | **9.53E-06** | 0.93 | (0.4-1.46) | **6.44E-04** | **2.08E-03** | BIS only | 0.63 | (0.24-1.02) | **1.54E-03** | **3.02E-03** |
| LPC(O-18:1) | 0.37 | (-0.67-1.41) | 4.87E-01 | 5.68E-01 | GUSTO only | 0.49 | (0.08-0.91) | **1.91E-02** | **2.76E-02** | 1.64 | (1.13-2.15) | **4.02E-10** | **4.44E-09** | GUSTO and BIS | 1.73 | (1.36-2.1) | **2.94E-20** | **1.98E-19** |
| LPC(O-20:0) | -2.20 | (-3.04--1.36) | **4.96E-07** | **5.01E-06** | GUSTO and BIS | -1.84 | (-2.27--1.4) | **1.96E-16** | **3.63E-15** | 0.94 | (0.46-1.42) | **1.27E-04** | **4.82E-04** | BIS only | 0.73 | (0.31-1.14) | **6.82E-04** | **1.43E-03** |
| LPC(O-20:1) | -0.83 | (-1.85-0.21) | 1.16E-01 | 1.79E-01 | notSig | -0.13 | (-0.79-0.53) | 6.96E-01 | 7.43E-01 | 1.51 | (1-2.02) | **8.84E-09** | **7.46E-08** | GUSTO and BIS | 1.74 | (1.28-2.19) | **1.00E-13** | **4.74E-13** |
| LPC(O-22:0) | -2.49 | (-3.23--1.75) | **1.17E-10** | **5.76E-09** | GUSTO and BIS | -2.43 | (-2.83--2.02) | **1.03E-31** | **2.39E-29** | 0.42 | (0-0.85) | 5.17E-02 | 9.67E-02 | notSig | 0.34 | (0.01-0.66) | **4.09E-02** | 6.23E-02 |
| LPC(O-22:1) | -1.58 | (-2.47--0.68) | **6.18E-04** | **2.19E-03** | GUSTO and BIS | -1.61 | (-2.06--1.16) | **2.91E-12** | **2.50E-11** | 1.50 | (1.01-1.99) | **2.80E-09** | **2.55E-08** | GUSTO and BIS | 1.41 | (1.04-1.79) | **1.82E-13** | **8.37E-13** |
| LPC(O-24:0) | -3.03 | (-3.89--2.17) | **1.85E-11** | **1.87E-09** | GUSTO and BIS | -3.04 | (-3.52--2.55) | **3.77E-35** | **1.75E-32** | 0.04 | (-0.5-0.57) | 8.87E-01 | 9.11E-01 | notSig | -0.23 | (-0.6-0.15) | 2.33E-01 | 2.87E-01 |
| LPC(O-24:1) | -1.89 | (-2.69--1.08) | **6.47E-06** | **4.55E-05** | GUSTO and BIS | -2.25 | (-2.71--1.8) | **3.96E-22** | **2.04E-20** | 1.07 | (0.61-1.54) | **7.35E-06** | **3.79E-05** | BIS only | 0.81 | (0.46-1.16) | **5.66E-06** | **1.47E-05** |
| LPC(O-24:2) | -2.24 | (-3.13--1.33) | **1.70E-06** | **1.43E-05** | GUSTO and BIS | -2.03 | (-2.53--1.53) | **9.99E-16** | **1.55E-14** | 0.15 | (-0.33-0.62) | 5.47E-01 | 6.32E-01 | notSig | 0.17 | (-0.19-0.53) | 3.51E-01 | 4.08E-01 |
| LPC(P-16:0) | -0.11 | (-1.16-0.96) | 8.40E-01 | 8.75E-01 | notSig | 0.34 | (-0.1-0.77) | 1.28E-01 | 1.62E-01 | 0.86 | (0.38-1.34) | **4.97E-04** | **1.65E-03** | GUSTO and BIS | 0.99 | (0.64-1.33) | **1.78E-08** | **5.55E-08** |
| LPC(P-18:0) | -0.84 | (-1.88-0.22) | 1.20E-01 | 1.83E-01 | notSig | 0.22 | (-0.2-0.63) | 3.14E-01 | 3.62E-01 | 0.72 | (0.2-1.24) | **6.31E-03** | **1.57E-02** | GUSTO and BIS | 1.23 | (0.86-1.6) | **5.74E-11** | **2.11E-10** |
| LPC(P-18:1) | 0.22 | (-0.87-1.33) | 6.94E-01 | 7.59E-01 | GUSTO only | 0.79 | (0.27-1.31) | **3.03E-03** | **5.26E-03** | 1.12 | (0.59-1.64) | **3.33E-05** | **1.46E-04** | GUSTO and BIS | 1.17 | (0.75-1.58) | **3.54E-08** | **1.08E-07** |
| LPC(P-20:0) | -1.26 | (-1.97--0.56) | **5.13E-04** | **1.91E-03** | GUSTO and BIS | -0.94 | (-1.39--0.5) | **3.48E-05** | **8.84E-05** | 0.62 | (0.18-1.06) | **6.04E-03** | **1.53E-02** | GUSTO and BIS | 1.03 | (0.66-1.4) | **4.35E-08** | **1.31E-07** |
| LPE(16:0) [sn2] | 0.33 | (-0.67-1.35) | 5.16E-01 | 5.94E-01 | GUSTO only | -0.40 | (-0.85-0.05) | 8.42E-02 | 1.11E-01 | 1.64 | (1.17-2.12) | **2.77E-11** | **3.67E-10** | GUSTO and BIS | 1.85 | (1.46-2.23) | **8.01E-21** | **5.55E-20** |
| LPE(16:0) [sn1] | 0.24 | (-0.73-1.23) | 6.26E-01 | 6.98E-01 | notSig | -0.33 | (-0.77-0.1) | 1.35E-01 | 1.70E-01 | 1.61 | (1.15-2.07) | **1.20E-11** | **1.69E-10** | GUSTO and BIS | 1.83 | (1.46-2.2) | **2.07E-22** | **1.57E-21** |

| | | | | | | | | | | | | | | | | | | |
|---|---|---|---|---|---|---|---|---|---|---|---|---|---|---|---|---|---|---|
| LPE(18:0) [sn2] | 0.52 | (-0.52-1.56) | 3.27E-01 | 4.19E-01 | notSig | 0.24 | (-0.2-0.67) | 2.87E-01 | 3.38E-01 | 1.26 | (0.78-1.74) | **2.72E-07** | **1.94E-06** | GUSTO and BIS | 1.32 | (0.96-1.68) | **5.11E-13** | **2.30E-12** |
| LPE(18:0) [sn1] | 0.74 | (-0.28-1.76) | 1.55E-01 | 2.27E-01 | notSig | 0.46 | (0.04-0.87) | **2.99E-02** | **4.24E-02** | 1.21 | (0.72-1.69) | **1.18E-06** | **7.21E-06** | GUSTO and BIS | 1.36 | (1-1.72) | **1.32E-13** | **6.11E-13** |
| LPE(18:1) [sn2] | -1.14 | (-2.04--0.24) | **1.36E-02** | **2.98E-02** | GUSTO and BIS | -0.88 | (-1.35--0.41) | **2.22E-04** | **4.84E-04** | 2.73 | (2.2-3.25) | **5.46E-23** | **1.19E-20** | GUSTO and BIS | 2.66 | (2.24-3.07) | **4.00E-36** | **7.14E-35** |
| LPE(18:1) [sn1] | -1.23 | (-2.15--0.3) | **9.49E-03** | **2.17E-02** | GUSTO and BIS | -0.87 | (-1.33--0.4) | **2.63E-04** | **5.57E-04** | 2.60 | (2.07-3.13) | **1.36E-20** | **1.27E-18** | GUSTO and BIS | 2.72 | (2.3-3.15) | **1.57E-36** | **2.91E-35** |
| LPE(18:2) [sn2] | -2.31 | (-3.32--1.3) | **1.04E-05** | **6.59E-05** | GUSTO and BIS | -1.35 | (-1.9--0.79) | **1.93E-06** | **6.11E-06** | 3.21 | (2.46-3.97) | **3.01E-16** | **8.74E-15** | GUSTO and BIS | 3.37 | (2.78-3.97) | **1.44E-28** | **1.55E-27** |
| LPE(18:2) [sn1] | -2.50 | (-3.58--1.41) | **9.53E-06** | **6.23E-05** | GUSTO and BIS | -1.39 | (-1.96--0.82) | **1.94E-06** | **6.11E-06** | 3.28 | (2.49-4.06) | **9.07E-16** | **2.34E-14** | GUSTO and BIS | 3.47 | (2.87-4.07) | **3.74E-30** | **4.45E-29** |
| LPE(20:4) [sn1] | 0.57 | (-0.4-1.55) | 2.47E-01 | 3.35E-01 | notSig | 0.57 | (0.1-1.04) | **1.80E-02** | **2.65E-02** | 2.92 | (2.13-3.71) | **8.91E-13** | **1.42E-11** | GUSTO and BIS | 3.04 | (2.49-3.59) | **2.81E-27** | **2.90E-26** |
| LPE(22:6) [sn2] | 1.32 | (0.22-2.44) | **1.86E-02** | **3.84E-02** | BIS only | 0.00 | (-0.52-0.52) | 9.92E-01 | 9.94E-01 | 1.76 | (0.86-2.66) | **1.31E-04** | **4.94E-04** | GUSTO and BIS | 1.97 | (1.4-2.54) | **1.14E-11** | **4.55E-11** |
| LPE(22:6) [sn1] | 0.85 | (-0.22-1.93) | 1.21E-01 | 1.83E-01 | notSig | -0.02 | (-0.57-0.52) | 9.31E-01 | 9.44E-01 | 1.91 | (1.07-2.75) | **8.81E-06** | **4.35E-05** | GUSTO and BIS | 2.16 | (1.6-2.71) | **2.23E-14** | **1.10E-13** |
| PC(28:0) | -4.68 | (-6.07--3.26) | **3.29E-10** | **1.22E-08** | GUSTO and BIS | -3.45 | (-4.31--2.59) | **3.66E-15** | **5.14E-14** | 0.39 | (-0.36-1.14) | 3.06E-01 | 4.09E-01 | notSig | -0.02 | (-0.49-0.46) | 9.45E-01 | 9.49E-01 |
| PC(14:0_16:0) | -2.51 | (-3.49--1.53) | **8.07E-07** | **7.20E-06** | GUSTO and BIS | -2.30 | (-2.87--1.72) | **5.36E-15** | **7.31E-14** | 0.19 | (-0.59-0.98) | 6.28E-01 | 7.10E-01 | notSig | -0.03 | (-0.54-0.49) | 9.24E-01 | 9.36E-01 |
| PC(31:0) (a) | -2.80 | (-3.99--1.6) | **6.84E-06** | **4.67E-05** | GUSTO and BIS | -3.07 | (-3.86--2.27) | **3.15E-14** | **3.57E-13** | -0.64 | (-1.49-0.2) | 1.35E-01 | 2.09E-01 | notSig | -0.60 | (-1.21-0.02) | 5.72E-02 | 8.27E-02 |
| PC(31:0) (b) | -1.34 | (-2.09--0.59) | **5.43E-04** | **1.97E-03** | GUSTO and BIS | -1.89 | (-2.37--1.41) | **6.56E-15** | **8.63E-14** | -0.67 | (-1.4-0.06) | 7.16E-02 | 1.25E-01 | notSig | -0.80 | (-1.37--0.24) | **5.49E-03** | **1.01E-02** |
| PC(16:0_16:0) | -0.40 | (-1.07-0.28) | 2.51E-01 | 3.37E-01 | GUSTO only | -0.58 | (-0.98--0.18) | **4.92E-03** | **8.24E-03** | -0.14 | (-0.9-0.62) | 7.13E-01 | 7.84E-01 | notSig | -0.39 | (-0.88-0.1) | 1.18E-01 | 1.60E-01 |
| PC(32:1) | -0.42 | (-1.51-0.68) | 4.52E-01 | 5.34E-01 | notSig | -0.71 | (-1.37--0.04) | **3.65E-02** | 5.10E-02 | 1.46 | (0.74-2.19) | **7.35E-05** | **2.89E-04** | GUSTO and BIS | 1.25 | (0.7-1.8) | **9.54E-06** | **2.41E-05** |
| PC(32:2) | -2.17 | (-3.15--1.17) | **2.37E-05** | **1.33E-04** | GUSTO and BIS | -1.77 | (-2.33--1.22) | **4.02E-10** | **2.39E-09** | 0.62 | (-0.29-1.54) | 1.82E-01 | 2.71E-01 | notSig | 0.27 | (-0.27-0.82) | 3.24E-01 | 3.84E-01 |
| PC(33:0) (a) | -0.43 | (-1.29-0.43) | 3.28E-01 | 4.19E-01 | GUSTO only | -1.47 | (-2.04--0.91) | **3.59E-07** | **1.37E-06** | -0.80 | (-1.6-0.01) | 5.16E-02 | 9.67E-02 | notSig | -0.79 | (-1.39--0.2) | **9.27E-03** | **1.61E-02** |
| PC(33:0) (b) | -0.60 | (-1.25-0.05) | 6.96E-02 | 1.17E-01 | GUSTO only | -1.28 | (-1.7--0.85) | **3.92E-09** | **2.04E-08** | -0.88 | (-1.6--0.16) | **1.73E-02** | **3.74E-02** | GUSTO and BIS | -1.15 | (-1.67--0.63) | **1.49E-05** | **3.72E-05** |
| PC(33:1) | -0.46 | (-1.33-0.43) | 3.10E-01 | 4.02E-01 | GUSTO only | -1.66 | (-2.21--1.11) | **3.79E-09** | **2.00E-08** | 0.09 | (-0.77-0.95) | 8.36E-01 | 8.82E-01 | notSig | -0.35 | (-0.85-0.16) | 1.78E-01 | 2.28E-01 |
| PC(33:2) | -0.83 | (-1.53--0.12) | **2.21E-02** | **4.38E-02** | GUSTO and BIS | -1.63 | (-2.1--1.16) | **9.66E-12** | **7.73E-11** | -0.84 | (-1.79-0.1) | 8.06E-02 | 1.35E-01 | GUSTO only | -1.13 | (-1.65--0.6) | **2.78E-05** | **6.79E-05** |
| PC(16:0_18:0) | -1.04 | (-1.6--0.48) | **3.21E-04** | **1.22E-03** | GUSTO and BIS | -1.01 | (-1.34--0.67) | **3.98E-09** | **2.05E-08** | 0.07 | (-0.52-0.66) | 8.24E-01 | 8.75E-01 | notSig | -0.10 | (-0.51-0.31) | 6.35E-01 | 6.86E-01 |
| PC(16:0_18:1) | 0.19 | (-0.19-0.57) | 3.31E-01 | 4.21E-01 | GUSTO only | -0.22 | (-0.54-0.09) | 1.65E-01 | 2.05E-01 | 0.90 | (0.36-1.44) | **1.07E-03** | **3.29E-03** | BIS only | 0.60 | (0.17-1.03) | **6.00E-03** | **1.08E-02** |
| PC(16:0_18:2) | -0.04 | (-0.34-0.25) | 7.70E-01 | 8.21E-01 | GUSTO only | -0.19 | (-0.4-0.02) | 8.34E-02 | 1.10E-01 | 0.20 | (-0.4-0.8) | 5.12E-01 | 6.06E-01 | notSig | -0.07 | (-0.56-0.42) | 7.88E-01 | 8.30E-01 |
| PC(16:1_18:2) | -0.45 | (-1.31-0.41) | 2.99E-01 | 3.90E-01 | GUSTO only | -0.72 | (-1.23--0.21) | **5.46E-03** | **9.06E-03** | 0.21 | (-0.68-1.1) | 6.43E-01 | 7.22E-01 | notSig | -0.42 | (-0.94-0.1) | 1.12E-01 | 1.53E-01 |
| PC(16:0_18:3) (a) | -1.99 | (-2.94--1.04) | **5.17E-05** | **2.50E-04** | GUSTO and BIS | -1.85 | (-2.39--1.31) | **1.69E-11** | **1.28E-10** | -0.34 | (-1.4-0.72) | 5.30E-01 | 6.18E-01 | notSig | -0.09 | (-0.74-0.56) | 7.81E-01 | 8.25E-01 |
| PC(14:0_20:4) | -1.32 | (-2.33--0.29) | **1.21E-02** | **2.69E-02** | BIS only | -0.94 | (-1.56--0.32) | **2.75E-03** | **4.84E-03** | 0.74 | (-0.04-1.53) | 6.46E-02 | 1.15E-01 | notSig | 0.18 | (-0.3-0.66) | 4.51E-01 | 5.05E-01 |
| PC(34:5) | -2.45 | (-3.87--1.01) | **9.36E-04** | **3.00E-03** | BIS only | -1.70 | (-2.61--0.79) | **2.50E-04** | **5.34E-04** | -0.53 | (-1.48-0.41) | 2.66E-01 | 3.66E-01 | GUSTO only | -0.88 | (-1.5--0.26) | **5.70E-03** | **1.03E-02** |
| PC(15-MHDA_18:1) | 0.22 | (-0.59-1.04) | 5.93E-01 | 6.67E-01 | GUSTO only | -1.18 | (-1.75--0.62) | **4.51E-05** | **1.11E-04** | -0.73 | (-1.55-0.1) | 8.35E-02 | 1.39E-01 | GUSTO only | -1.01 | (-1.57--0.45) | **4.28E-04** | **9.24E-04** |
| PC(17:0_18:1) | 0.02 | (-0.75-0.8) | 9.58E-01 | 9.66E-01 | GUSTO only | -1.38 | (-1.8--0.95) | **2.17E-10** | **1.36E-09** | -0.48 | (-1.2-0.25) | 1.96E-01 | 2.87E-01 | GUSTO only | -0.87 | (-1.28--0.45) | **4.19E-05** | **1.01E-04** |
| PC(15-MHDA_18:2) | 0.01 | (-0.9-0.92) | 9.86E-01 | 9.88E-01 | GUSTO only | -1.03 | (-1.64--0.43) | **8.42E-04** | **1.58E-03** | -1.27 | (-2.34--0.2) | **2.01E-02** | **4.26E-02** | GUSTO and BIS | -1.35 | (-1.94--0.76) | **6.66E-06** | **1.71E-05** |
| PC(17:0_18:2) | -0.35 | (-1.19-0.49) | 4.12E-01 | 4.96E-01 | GUSTO only | -1.20 | (-1.65--0.74) | **2.56E-07** | **9.97E-07** | -0.94 | (-1.91-0.04) | 5.99E-02 | 1.09E-01 | GUSTO only | -1.19 | (-1.69--0.69) | **2.85E-06** | **7.51E-06** |
| PC(17:1_18:2) | -0.82 | (-1.51--0.14) | **1.89E-02** | **3.86E-02** | GUSTO and BIS | -1.06 | (-1.49--0.64) | **8.76E-07** | **2.88E-06** | -0.75 | (-1.59-0.08) | 7.59E-02 | 1.31E-01 | GUSTO only | -0.73 | (-1.2--0.27) | **2.08E-03** | **4.03E-03** |
| PC(15:0_20:3) | -0.99 | (-1.64--0.34) | **3.00E-03** | **8.16E-03** | GUSTO and BIS | -0.63 | (-0.87--0.39) | **2.98E-07** | **1.14E-06** | -0.32 | (-1.04-0.4) | 3.82E-01 | 4.87E-01 | notSig | -0.22 | (-0.81-0.37) | 4.61E-01 | 5.14E-01 |
| PC(15:0_20:4) | 0.76 | (-0.05-1.58) | 6.46E-02 | 1.11E-01 | GUSTO only | -0.41 | (-0.94-0.12) | 1.26E-01 | 1.59E-01 | -0.49 | (-1.2-0.22) | 1.72E-01 | 2.59E-01 | GUSTO only | -0.99 | (-1.44--0.54) | **1.70E-05** | **4.19E-05** |
| PC(35:5) | -0.98 | (-2.36-0.41) | 1.66E-01 | 2.40E-01 | GUSTO only | -1.31 | (-2.23--0.39) | **5.47E-03** | **9.06E-03** | -1.77 | (-2.84--0.71) | **1.13E-03** | **3.45E-03** | GUSTO and BIS | -2.11 | (-2.78--1.44) | **7.15E-10** | **2.50E-09** |
| PC(18:0_18:1) | -1.19 | (-1.88--0.5) | **8.08E-04** | **2.68E-03** | GUSTO and BIS | -1.28 | (-1.71--0.85) | **7.15E-09** | **3.49E-08** | 1.22 | (0.63-1.81) | **5.51E-05** | **2.26E-04** | BIS only | 0.84 | (0.42-1.26) | **7.52E-05** | **1.77E-04** |
| PC(18:0_18:2) | -0.64 | (-1.03--0.25) | **1.46E-03** | **4.41E-03** | GUSTO and BIS | -0.63 | (-0.95--0.31) | **9.38E-05** | **2.21E-04** | 0.28 | (-0.39-0.95) | 4.10E-01 | 5.10E-01 | notSig | -0.10 | (-0.57-0.37) | 6.78E-01 | 7.26E-01 |
| PC(16:0_20:3) (a) | -0.46 | (-1.03-0.11) | 1.14E-01 | 1.76E-01 | GUSTO only | -0.83 | (-1.24--0.42) | **6.66E-05** | **1.59E-04** | 0.95 | (0.33-1.57) | **2.84E-03** | **7.66E-03** | BIS only | 0.36 | (-0.1-0.83) | 1.27E-01 | 1.71E-01 |
| PC(16:0_20:3) (b) | -1.26 | (-2.19--0.32) | **8.79E-03** | **2.02E-02** | BIS only | -0.09 | (-0.68-0.5) | 7.59E-01 | 7.99E-01 | 3.62 | (2.8-4.44) | **2.32E-17** | **8.96E-16** | BIS only | 1.59 | (1.08-2.1) | **8.24E-10** | **2.85E-09** |
| PC(18:2_18:2) | -2.91 | (-3.7--2.11) | **3.37E-12** | **5.22E-10** | GUSTO and BIS | -2.72 | (-3.25--2.19) | **5.05E-24** | **2.93E-22** | 0.49 | (-0.43-1.4) | 2.97E-01 | 3.99E-01 | notSig | 0.43 | (-0.23-1.09) | 2.06E-01 | 2.59E-01 |
| PC(16:1_20:4) | 0.28 | (-0.51-1.08) | 4.90E-01 | 5.70E-01 | notSig | -0.12 | (-0.64-0.39) | 6.39E-01 | 6.89E-01 | 0.98 | (0.35-1.61) | **2.30E-03** | **6.52E-03** | BIS only | 0.52 | (0.07-0.97) | **2.38E-02** | **3.85E-02** |
| PC(16:0_20:5) | 0.04 | (-1.11-1.2) | 9.49E-01 | 9.64E-01 | notSig | -0.10 | (-0.94-0.73) | 8.07E-01 | 8.36E-01 | -0.81 | (-1.71-0.09) | 7.88E-02 | 1.34E-01 | GUSTO only | -1.35 | (-1.98--0.72) | **2.54E-05** | **6.23E-05** |
| PC(15-MHDA_20:4) | 1.05 | (0.36-1.74) | **2.70E-03** | **7.53E-03** | BIS only | 0.30 | (-0.22-0.81) | 2.60E-01 | 3.08E-01 | -1.15 | (-1.81--0.5) | **5.34E-04** | **1.76E-03** | GUSTO and BIS | -1.38 | (-1.85--0.91) | **9.74E-09** | **3.05E-08** |
| PC(17:0_20:4) | 1.30 | (0.57-2.04) | **5.05E-04** | **1.89E-03** | BIS only | 0.38 | (-0.1-0.86) | 1.25E-01 | 1.59E-01 | -1.35 | (-1.9--0.8) | **1.68E-06** | **9.75E-06** | GUSTO and BIS | -1.54 | (-1.92--1.17) | **6.96E-16** | **3.71E-15** |
| PC(15:0_22:6) | -0.74 | (-1.59-0.12) | 9.04E-02 | 1.46E-01 | GUSTO only | -1.86 | (-2.41--1.3) | **6.72E-11** | **4.65E-10** | -1.40 | (-2.16--0.63) | **3.66E-04** | **1.26E-03** | GUSTO and BIS | -1.71 | (-2.17--1.24) | **5.20E-13** | **2.32E-12** |
| PC(38:2) | -1.40 | (-1.99--0.81) | **4.63E-06** | **3.52E-05** | GUSTO and BIS | -1.23 | (-1.59--0.87) | **2.15E-11** | **1.58E-10** | 1.28 | (0.78-1.78) | **5.73E-07** | **3.75E-06** | GUSTO and BIS | 1.14 | (0.75-1.52) | **8.45E-09** | **2.69E-08** |
| PC(18:0_20:3) | -0.59 | (-1.34-0.17) | 1.26E-01 | 1.89E-01 | GUSTO only | 0.39 | (-0.12-0.9) | 1.37E-01 | 1.73E-01 | 0.56 | (0.08-1.05) | **2.18E-02** | **4.56E-02** | BIS only | 0.38 | (0.01-0.75) | **4.33E-02** | 6.53E-02 |
| PC(18:1_20:3) | -0.77 | (-1.5--0.04) | **3.98E-02** | 7.21E-02 | notSig | -0.66 | (-1.13--0.19) | **5.77E-03** | **9.49E-03** | 1.35 | (0.71-2) | **4.08E-05** | **1.72E-04** | BIS only | 0.78 | (0.33-1.23) | **6.66E-04** | **1.40E-03** |
| PC(38:4) (b) | -1.31 | (-2.05--0.56) | **6.54E-04** | **2.25E-03** | GUSTO and BIS | -1.13 | (-1.63--0.62) | **1.26E-05** | **3.52E-05** | 0.55 | (0.05-1.06) | **3.07E-02** | 6.14E-02 | notSig | 0.35 | (-0.07-0.77) | 1.02E-01 | 1.40E-01 |
| PC(18:0_20:4) | 1.26 | (0.63-1.91) | **1.01E-04** | **4.47E-04** | GUSTO and BIS | 1.12 | (0.67-1.57) | **1.02E-06** | **3.25E-06** | -0.34 | (-0.8-0.12) | 1.49E-01 | 2.28E-01 | GUSTO only | -0.61 | (-0.98--0.25) | **9.84E-04** | **2.02E-03** |
| PC(38:5) (a) | 0.26 | (-0.28-0.8) | 3.46E-01 | 4.35E-01 | GUSTO only | -0.47 | (-0.83--0.1) | **1.34E-02** | **2.04E-02** | -0.03 | (-0.53-0.47) | 9.05E-01 | 9.27E-01 | GUSTO only | -0.58 | (-0.94--0.21) | **1.76E-03** | **3.41E-03** |
| PC(38:5) (b) | -0.57 | (-1.24-0.1) | 9.65E-02 | 1.53E-01 | GUSTO only | -0.88 | (-1.37--0.39) | **4.74E-04** | **9.37E-04** | 0.88 | (0.33-1.44) | **1.68E-03** | **4.95E-03** | BIS only | 0.58 | (0.15-1.01) | **7.95E-03** | **1.39E-02** |
| PC(38:6) (a) | -0.86 | (-1.53--0.19) | **1.19E-02** | **2.67E-02** | GUSTO and BIS | -0.95 | (-1.39--0.5) | **3.04E-05** | **7.90E-05** | 0.44 | (-0.31-1.19) | 2.48E-01 | 3.49E-01 | notSig | 0.23 | (-0.3-0.75) | 3.93E-01 | 4.51E-01 |
| PC(16:0_22:6) | 0.05 | (-0.47-0.59) | 8.42E-01 | 8.75E-01 | GUSTO only | -0.38 | (-0.79-0.02) | 6.43E-02 | 8.68E-02 | -0.50 | (-0.95--0.04) | **3.14E-02** | 6.25E-02 | GUSTO only | -0.71 | (-1.09--0.33) | **2.65E-04** | **5.90E-04** |
| PC(18:2_20:5) | -1.97 | (-3.01--0.91) | **2.88E-04** | **1.11E-03** | GUSTO and BIS | -1.86 | (-2.57--1.14) | **3.62E-07** | **1.37E-06** | 0.23 | (-0.63-1.08) | 6.01E-01 | 6.83E-01 | notSig | -0.23 | (-0.82-0.36) | 4.48E-01 | 5.03E-01 |

| | | | | | | | | | | | | | | | | | | |
|---|---|---|---|---|---|---|---|---|---|---|---|---|---|---|---|---|---|---|
| PC(16:1_22:6) | -0.18 | (-0.96-0.61) | 6.57E-01 | 7.22E-01 | GUSTO only | -1.01 | (-1.52--0.49) | **1.26E-04** | **2.90E-04** | -0.19 | (-0.8-0.41) | 5.28E-01 | 6.17E-01 | GUSTO only | -0.66 | (-1.07--0.26) | **1.33E-03** | **2.63E-03** |
| PC(38:7)(c) | 0.06 | (-0.68-0.81) | 8.75E-01 | 9.01E-01 | GUSTO only | -0.95 | (-1.41--0.49) | **5.83E-05** | **1.40E-04** | -0.34 | (-0.94-0.26) | 2.62E-01 | 3.63E-01 | GUSTO only | -0.66 | (-1.04--0.27) | **7.68E-04** | **1.58E-03** |
| PC(39:5)(a) | -0.60 | (-1.43-0.24) | 1.62E-01 | 2.36E-01 | GUSTO only | -1.57 | (-2.16--0.99) | **1.49E-07** | **6.13E-07** | -1.86 | (-2.59--1.13) | **8.03E-07** | **5.01E-06** | GUSTO and BIS | -1.93 | (-2.42--1.43) | **2.05E-14** | **1.02E-13** |
| PC(39:5)(b) | -0.42 | (-1.1-0.27) | 2.33E-01 | 3.19E-01 | GUSTO only | -1.30 | (-1.8--0.8) | **3.85E-07** | **1.42E-06** | -1.07 | (-1.72--0.43) | **1.03E-03** | **3.18E-03** | GUSTO and BIS | -1.52 | (-1.97--1.07) | **3.30E-11** | **1.24E-10** |
| PC(15-MHDA_22:6) | 0.12 | (-0.79-1.04) | 7.99E-01 | 8.45E-01 | GUSTO only | -1.36 | (-1.97--0.76) | **1.06E-05** | **2.99E-05** | -2.37 | (-3.11--1.64) | **3.98E-10** | **4.44E-09** | GUSTO and BIS | -2.21 | (-2.71--1.71) | **4.68E-18** | **2.72E-17** |
| PC(17:0_22:6) | -0.17 | (-0.99-0.65) | 6.75E-01 | 7.41E-01 | GUSTO only | -1.18 | (-1.73--0.63) | **2.49E-05** | **6.69E-05** | -1.94 | (-2.61--1.27) | **1.81E-08** | **1.50E-07** | GUSTO and BIS | -2.12 | (-2.55--1.68) | **6.87E-22** | **5.14E-21** |
| PC(18:0_22:5) (n6) | -1.13 | (-2.15--0.11) | **3.01E-02** | 5.80E-02 | GUSTO only | -1.25 | (-1.86--0.65) | **4.66E-05** | **1.14E-04** | 1.74 | (1.01-2.47) | **3.76E-06** | **2.05E-05** | BIS only | 0.94 | (0.31-1.57) | **3.28E-03** | **6.18E-03** |
| PC(18:0_22:6) | -0.20 | (-0.92-0.52) | 5.77E-01 | 6.54E-01 | notSig | -0.41 | (-0.92-0.09) | 1.11E-01 | 1.42E-01 | -0.82 | (-1.39--0.25) | **4.70E-03** | **1.23E-02** | GUSTO and BIS | -1.08 | (-1.51--0.64) | **1.23E-06** | **3.32E-06** |
| PC(40:7) (a) | -1.26 | (-2.07--0.45) | **2.30E-03** | **6.51E-03** | GUSTO and BIS | -1.33 | (-1.82--0.83) | **1.57E-07** | **6.30E-07** | 1.75 | (1.08-2.42) | **3.58E-07** | **2.44E-06** | BIS only | 1.49 | (0.94-2.05) | **1.16E-07** | **3.41E-07** |
| PC(18:1_22:6) (a) | -0.62 | (-1.26-0.03) | 6.33E-02 | 1.10E-01 | GUSTO only | -1.25 | (-1.67--0.84) | **3.36E-09** | **1.79E-08** | -0.08 | (-0.58-0.42) | 7.45E-01 | 8.07E-01 | notSig | -0.21 | (-0.61-0.2) | 3.18E-01 | 3.77E-01 |
| PC(40:8) | -1.39 | (-2.06--0.72) | **5.91E-05** | **2.77E-04** | GUSTO and BIS | -1.62 | (-2.04--1.2) | **2.37E-14** | **2.82E-13** | 1.18 | (0.54-1.83) | **3.61E-04** | **1.25E-03** | GUSTO and BIS | 1.01 | (0.55-1.47) | **1.69E-05** | **4.19E-05** |
| PC(44:12) | -1.58 | (-2.86--0.27) | **1.81E-02** | **3.76E-02** | GUSTO and BIS | -2.07 | (-2.89--1.25) | **6.67E-07** | **2.26E-06** | -0.84 | (-1.54--0.14) | **1.90E-02** | **4.07E-02** | GUSTO and BIS | -0.92 | (-1.44--0.4) | **4.94E-04** | **1.06E-03** |
| PC(O-16:0/16:0) | -1.28 | (-1.95--0.61) | **2.19E-04** | **8.67E-04** | GUSTO and BIS | -0.93 | (-1.29--0.57) | **5.07E-07** | **1.80E-06** | 0.75 | (0.05-1.44) | **3.63E-02** | 7.11E-02 | notSig | 0.59 | (0.11-1.07) | **1.58E-02** | **2.64E-02** |
| PC(O-32:1) | -1.65 | (-2.49--0.8) | **1.47E-04** | **6.07E-04** | GUSTO and BIS | -1.20 | (-1.66--0.74) | **3.71E-07** | **1.39E-06** | 0.48 | (-0.17-1.14) | 1.49E-01 | 2.29E-01 | notSig | 0.59 | (0.09-1.09) | **2.01E-02** | **3.30E-02** |
| PC(O-34:1) | -1.01 | (-1.65--0.37) | **1.99E-03** | **5.76E-03** | GUSTO and BIS | -0.89 | (-1.25--0.54) | **9.38E-07** | **3.06E-06** | 1.00 | (0.26-1.73) | **7.80E-03** | **1.90E-02** | BIS only | 0.83 | (0.33-1.33) | **1.13E-03** | **2.27E-03** |
| PC(O-34:2) | -2.65 | (-3.43--1.86) | **1.09E-10** | **5.76E-09** | GUSTO and BIS | -2.25 | (-2.73--1.77) | **3.67E-20** | **1.22E-18** | -0.02 | (-0.77-0.73) | 9.57E-01 | 9.63E-01 | notSig | -0.04 | (-0.57-0.48) | 8.71E-01 | 8.97E-01 |
| PC(O-34:4) | -1.10 | (-1.88--0.31) | **6.50E-03** | **1.58E-02** | BIS only | -1.01 | (-1.62--0.4) | **1.19E-03** | **2.18E-03** | 0.08 | (-0.67-0.84) | 8.29E-01 | 8.78E-01 | notSig | 0.03 | (-0.49-0.55) | 9.05E-01 | 9.25E-01 |
| PC(O-35:4) | -0.77 | (-1.6-0.07) | 7.30E-02 | 1.21E-01 | notSig | -0.83 | (-1.42--0.23) | **6.46E-03** | **1.05E-02** | -0.21 | (-0.87-0.44) | 5.24E-01 | 6.16E-01 | notSig | -0.35 | (-0.82-0.11) | 1.37E-01 | 1.82E-01 |
| PC(O-36:0) | -2.06 | (-2.7--1.41) | **9.59E-10** | **2.34E-08** | GUSTO and BIS | -1.58 | (-1.97--1.19) | **3.04E-15** | **4.41E-14** | 0.34 | (-0.29-0.98) | 2.88E-01 | 3.89E-01 | notSig | 0.27 | (-0.24-0.78) | 3.00E-01 | 3.59E-01 |
| PC(O-18:0/18:1) | -2.11 | (-2.85--1.36) | **4.47E-08** | **6.48E-07** | GUSTO and BIS | -1.68 | (-2.08--1.28) | **1.52E-16** | **2.94E-15** | 1.03 | (0.38-1.67) | **1.87E-03** | **5.46E-03** | GUSTO and BIS | 1.16 | (0.68-1.63) | **2.26E-06** | **6.02E-06** |
| PC(O-18:1/18:1) | -1.68 | (-2.58--0.77) | **3.04E-04** | **1.16E-03** | GUSTO and BIS | -1.30 | (-1.71--0.89) | **4.60E-10** | **2.70E-09** | 1.46 | (0.46-2.45) | **4.13E-03** | **1.09E-02** | GUSTO and BIS | 0.92 | (0.39-1.45) | **6.12E-04** | **1.30E-03** |
| PC(O-18:0/18:2) | -3.85 | (-4.96--2.73) | **5.08E-11** | **3.37E-09** | GUSTO and BIS | -2.66 | (-3.17--2.16) | **5.23E-25** | **3.47E-23** | 0.44 | (-0.59-1.47) | 4.01E-01 | 5.07E-01 | notSig | 0.44 | (-0.1-0.99) | 1.12E-01 | 1.53E-01 |
| PC(O-18:1/18:2) | -1.90 | (-2.61--1.19) | **2.28E-07** | **2.75E-06** | GUSTO and BIS | -1.60 | (-2.03--1.16) | **6.19E-13** | **5.74E-12** | 0.47 | (-0.24-1.17) | 1.96E-01 | 2.87E-01 | notSig | 0.19 | (-0.3-0.67) | 4.49E-01 | 5.03E-01 |
| PC(O-16:0/20:3) | -1.54 | (-2.15--0.92) | **1.44E-06** | **1.24E-05** | BIS only | -0.94 | (-1.34--0.54) | **4.09E-06** | **1.23E-05** | 0.89 | (0.31-1.47) | **2.49E-03** | **6.88E-03** | BIS only | 0.69 | (0.24-1.14) | **2.42E-03** | **4.60E-03** |
| PC(O-16:0/20:4) | -0.16 | (-0.81-0.5) | 6.34E-01 | 7.04E-01 | notSig | -0.15 | (-0.57-0.26) | 4.62E-01 | 5.19E-01 | 0.73 | (0.04-1.42) | **3.74E-02** | 7.25E-02 | notSig | 0.23 | (-0.23-0.68) | 3.28E-01 | 3.87E-01 |
| PC(O-36:5) | -1.22 | (-2.18--0.26) | **1.31E-02** | **2.89E-02** | GUSTO and BIS | -1.17 | (-1.86--0.48) | **8.81E-04** | **1.65E-03** | -0.32 | (-1.06-0.43) | 4.03E-01 | 5.07E-01 | GUSTO only | -0.71 | (-1.22--0.21) | **5.66E-03** | **1.03E-02** |
| PC(O-18:0/20:4) | -1.23 | (-1.95--0.51) | **8.74E-04** | **2.86E-03** | BIS only | -0.68 | (-1.1--0.26) | **1.41E-03** | **2.55E-03** | 0.20 | (-0.4-0.8) | 5.12E-01 | 6.06E-01 | notSig | -0.08 | (-0.48-0.33) | 7.13E-01 | 7.58E-01 |
| PC(O-38:5) | 0.23 | (-0.31-0.77) | 4.13E-01 | 4.97E-01 | notSig | 0.09 | (-0.26-0.45) | 6.06E-01 | 6.60E-01 | 0.70 | (0.09-1.32) | **2.50E-02** | 5.11E-02 | notSig | 0.32 | (-0.14-0.78) | 1.71E-01 | 2.21E-01 |
| PC(O-16:0/22:6) | -0.65 | (-1.35-0.05) | 6.81E-02 | 1.15E-01 | notSig | -0.44 | (-0.93-0.05) | 7.84E-02 | 1.04E-01 | -0.15 | (-0.74-0.44) | 6.22E-01 | 7.06E-01 | GUSTO only | -0.46 | (-0.89--0.04) | **3.33E-02** | 5.22E-02 |
| PC(O-40:5) | -0.68 | (-1.28--0.06) | **3.03E-02** | 5.81E-02 | GUSTO only | -0.75 | (-1.14--0.37) | **1.31E-04** | **3.00E-04** | 0.36 | (-0.24-0.95) | 2.38E-01 | 3.39E-01 | notSig | 0.07 | (-0.32-0.47) | 7.16E-01 | 7.60E-01 |
| PC(O-18:0/22:6) | -1.39 | (-2.1--0.66) | **1.87E-04** | **7.56E-04** | GUSTO and BIS | -0.95 | (-1.4--0.51) | **2.61E-05** | **6.91E-05** | -0.49 | (-1.03-0.05) | 7.69E-02 | 1.31E-01 | GUSTO only | -0.71 | (-1.09--0.34) | **1.98E-04** | **4.45E-04** |
| PC(O-40:7) (a) | -0.02 | (-0.71-0.67) | 9.52E-01 | 9.65E-01 | notSig | -0.04 | (-0.52-0.43) | 8.61E-01 | 8.84E-01 | 0.26 | (-0.34-0.86) | 3.91E-01 | 4.95E-01 | GUSTO only | -0.24 | (-0.66-0.18) | 2.69E-01 | 3.26E-01 |
| PC(P-16:0/14:0) | -2.41 | (-3.13--1.69) | **1.37E-10** | **5.76E-09** | GUSTO and BIS | -1.73 | (-2.19--1.28) | **9.24E-14** | **9.72E-13** | 0.37 | (-0.46-1.21) | 3.82E-01 | 4.87E-01 | notSig | -0.04 | (-0.59-0.51) | 8.92E-01 | 9.13E-01 |
| PC(P-16:0/16:0) | -1.04 | (-1.67--0.41) | **1.26E-03** | **3.83E-03** | GUSTO and BIS | -0.84 | (-1.23--0.44) | **3.14E-05** | **8.08E-05** | 0.28 | (-0.48-1.04) | 4.67E-01 | 5.66E-01 | notSig | -0.17 | (-0.67-0.32) | 4.91E-01 | 5.45E-01 |
| PC(P-16:0/16:1) | -0.98 | (-1.64--0.32) | **3.77E-03** | **1.01E-02** | GUSTO and BIS | -0.83 | (-1.28--0.39) | **2.26E-04** | **4.90E-04** | 0.87 | (0.13-1.61) | **2.21E-02** | **4.59E-02** | BIS only | 0.39 | (-0.12-0.9) | 1.34E-01 | 1.80E-01 |
| PC(P-16:0/18:0) | -2.11 | (-3.14--1.07) | **8.57E-05** | **3.90E-04** | GUSTO and BIS | -1.32 | (-1.68--0.95) | **2.10E-12** | **1.84E-11** | -0.19 | (-1.26-0.88) | 7.28E-01 | 7.97E-01 | notSig | -0.23 | (-0.71-0.26) | 3.60E-01 | 4.16E-01 |
| PC(P-16:0/18:1) | -1.46 | (-2.14--0.77) | **3.88E-05** | **2.02E-04** | GUSTO and BIS | -1.31 | (-1.73--0.88) | **1.61E-09** | **8.77E-09** | 0.43 | (-0.26-1.12) | 2.24E-01 | 3.23E-01 | notSig | 0.03 | (-0.43-0.48) | 9.09E-01 | 9.25E-01 |
| PC(P-16:0/18:2) | -1.90 | (-2.69--1.11) | **3.14E-06** | **2.48E-05** | GUSTO and BIS | -1.38 | (-1.86--0.91) | **8.71E-09** | **4.17E-08** | -0.93 | (-1.87-0.02) | 5.42E-02 | 1.00E-01 | GUSTO only | -1.19 | (-1.71--0.67) | **7.54E-06** | **1.92E-05** |
| PC(P-16:0/18:3) | -3.16 | (-4.1--2.22) | **1.35E-10** | **5.76E-09** | GUSTO and BIS | -2.12 | (-2.64--1.59) | **2.34E-15** | **3.50E-14** | -0.88 | (-1.69--0.06) | **3.51E-02** | 6.90E-02 | notSig | -0.60 | (-1.05--0.15) | **9.39E-03** | **1.62E-02** |
| PC(P-35:2)(a) | -2.69 | (-3.8--1.57) | **3.32E-06** | **2.57E-05** | GUSTO and BIS | -2.43 | (-3.11--1.74) | **4.02E-12** | **3.39E-11** | -1.45 | (-2.56--0.33) | **1.10E-02** | **2.53E-02** | BIS only | -0.73 | (-1.33--0.14) | **1.56E-02** | **2.61E-02** |
| PC(P-35:2)(b) | -2.44 | (-3.55--1.32) | **2.37E-05** | **1.33E-04** | GUSTO and BIS | -2.01 | (-2.62--1.4) | **8.01E-11** | **5.39E-10** | -1.49 | (-2.74--0.24) | **1.94E-02** | **4.12E-02** | GUSTO and BIS | -1.46 | (-2.03--0.89) | **4.47E-07** | **1.25E-06** |
| PC(P-15:0/20:4) (b) | -1.21 | (-2.04--0.37) | **4.84E-03** | **1.24E-02** | BIS only | -0.88 | (-1.41--0.35) | **1.03E-03** | **1.91E-03** | -0.56 | (-1.09--0.02) | **4.06E-02** | 7.82E-02 | notSig | -0.63 | (-1.06--0.2) | **4.46E-03** | **8.28E-03** |
| PC(P-18:1/18:1) | -1.46 | (-2.15--0.76) | **4.81E-05** | **2.39E-04** | GUSTO and BIS | -1.29 | (-1.71--0.88) | **1.12E-09** | **6.25E-09** | 2.18 | (1.51-2.84) | **2.66E-10** | **3.09E-09** | GUSTO and BIS | 1.82 | (1.29-2.35) | **1.46E-11** | **5.76E-11** |
| PC(P-18:0/18:2) | -2.51 | (-3.46--1.55) | **4.05E-07** | **4.18E-06** | GUSTO and BIS | -2.09 | (-2.58--1.6) | **1.04E-16** | **2.19E-15** | -0.52 | (-1.46-0.42) | 2.82E-01 | 3.84E-01 | notSig | -0.61 | (-1.09--0.13) | **1.23E-02** | **2.08E-02** |
| PC(P-16:0/20:4) | 0.51 | (-0.25-1.29) | 1.89E-01 | 2.68E-01 | notSig | 0.47 | (-0.02-0.95) | 5.81E-02 | 7.93E-02 | 0.06 | (-0.7-0.82) | 8.76E-01 | 9.06E-01 | GUSTO only | -0.68 | (-1.14--0.22) | **3.87E-03** | **7.23E-03** |
| PC(P-16:0/20:5) | -0.88 | (-1.95-0.2) | 1.10E-01 | 1.71E-01 | notSig | -0.69 | (-1.44-0.05) | 6.92E-02 | 9.26E-02 | -1.12 | (-1.92--0.33) | **5.76E-03** | **1.48E-02** | GUSTO and BIS | -1.58 | (-2.11--1.04) | **6.68E-09** | **2.18E-08** |
| PC(P-17:0/20:4) (a) | -0.26 | (-1.01-0.49) | 4.97E-01 | 5.75E-01 | GUSTO only | -0.62 | (-1.16--0.07) | **2.67E-02** | **3.81E-02** | -1.26 | (-1.93--0.6) | **2.25E-04** | **8.10E-04** | GUSTO and BIS | -1.41 | (-1.89--0.94) | **6.15E-09** | **2.02E-08** |
| PC(P-17:0/20:4) (b) | 0.04 | (-0.72-0.8) | 9.25E-01 | 9.45E-01 | notSig | -0.15 | (-0.67-0.38) | 5.86E-01 | 6.42E-01 | -1.05 | (-1.65--0.45) | **6.31E-04** | **2.05E-03** | GUSTO and BIS | -1.47 | (-1.89--1.04) | **1.23E-11** | **4.90E-11** |
| PC(P-18:0/20:4) | 0.11 | (-0.73-0.95) | 8.04E-01 | 8.48E-01 | notSig | -0.03 | (-0.51-0.45) | 9.16E-01 | 9.31E-01 | -0.30 | (-1.02-0.41) | 4.05E-01 | 5.07E-01 | GUSTO only | -0.82 | (-1.24--0.4) | **1.43E-04** | **3.24E-04** |
| PC(P-38:5) (a) | 0.80 | (0.21-1.39) | **7.78E-03** | **1.82E-02** | BIS only | 0.53 | (0.1-0.95) | **1.44E-02** | **2.17E-02** | -0.16 | (-0.83-0.51) | 6.43E-01 | 7.22E-01 | GUSTO only | -0.78 | (-1.21--0.35) | **4.00E-04** | **8.68E-04** |
| PC(P-38:5) (b) | -0.54 | (-1.26-0.19) | 1.44E-01 | 2.14E-01 | GUSTO only | -0.73 | (-1.18--0.27) | **1.65E-03** | **2.97E-03** | 0.41 | (-0.26-1.09) | 2.28E-01 | 3.26E-01 | GUSTO only | -0.26 | (-0.72-0.21) | 2.83E-01 | 3.40E-01 |
| PC(P-16:0/22:6) | -0.38 | (-1.13-0.37) | 3.15E-01 | 4.05E-01 | notSig | -0.34 | (-0.85-0.18) | 2.03E-01 | 2.47E-01 | -0.67 | (-1.36-0.01) | 5.32E-02 | 9.87E-02 | GUSTO only | -1.16 | (-1.61--0.71) | **3.72E-07** | **1.05E-06** |
| PC(P-20:0/20:4) | -0.55 | (-1.24-0.14) | 1.21E-01 | 1.83E-01 | notSig | -0.48 | (-0.94--0.03) | **3.68E-02** | 5.12E-02 | -0.27 | (-0.8-0.26) | 3.21E-01 | 4.25E-01 | GUSTO only | -0.52 | (-0.92--0.12) | **1.01E-02** | **1.73E-02** |
| PC(P-18:0/22:6) | -0.76 | (-1.56-0.05) | 6.57E-02 | 1.13E-01 | notSig | -0.72 | (-1.24--0.19) | **7.23E-03** | **1.16E-02** | -0.91 | (-1.57--0.25) | **7.08E-03** | **1.75E-02** | GUSTO and BIS | -1.31 | (-1.72--0.9) | **4.85E-10** | **1.70E-09** |
| PE(16:0_16:1) | -0.24 | (-1.79-1.33) | 7.62E-01 | 8.15E-01 | notSig | -0.43 | (-1.36-0.5) | 3.61E-01 | 4.13E-01 | 1.33 | (0.29-2.38) | **1.21E-02** | **2.75E-02** | BIS only | 0.90 | (0.15-1.65) | **1.81E-02** | **3.01E-02** |

| | | | | | | | | | | | | | | | | | | |
|---|---|---|---|---|---|---|---|---|---|---|---|---|---|---|---|---|---|---|
| PE(16:0_18:1) | 0.82 | (-0.18-1.83) | 1.10E-01 | 1.71E-01 | notSig | 0.24 | (-0.37-0.85) | 4.47E-01 | 5.06E-01 | -0.11 | (-1.32-1.11) | 8.64E-01 | 8.98E-01 | notSig | -0.47 | (-1.09-0.15) | 1.34E-01 | 1.80E-01 |
| PE(16:0_18:2) | -0.39 | (-1.39-0.63) | 4.52E-01 | 5.34E-01 | notSig | -0.35 | (-1-0.31) | 3.00E-01 | 3.49E-01 | -0.37 | (-1.48-0.75) | 5.16E-01 | 6.08E-01 | notSig | -0.58 | (-1.2-0.04) | 6.61E-02 | 9.35E-02 |
| PE(16:1_18:2) | -1.14 | (-2.49-0.23) | 1.01E-01 | 1.60E-01 | notSig | -1.02 | (-1.83--0.2) | **1.44E-02** | **2.17E-02** | 0.69 | (-0.43-1.8) | 2.27E-01 | 3.26E-01 | notSig | 0.49 | (-0.17-1.14) | 1.47E-01 | 1.93E-01 |
| PE(16:0_18:3) (a) | -1.97 | (-3.3--0.62) | **4.51E-03** | **1.18E-02** | GUSTO and BIS | -1.73 | (-2.49--0.97) | **8.28E-06** | **2.36E-05** | -0.48 | (-2.12-1.15) | 5.62E-01 | 6.44E-01 | notSig | -0.04 | (-0.77-0.68) | 9.09E-01 | 9.25E-01 |
| PE(16:0_18:3) (b) | -1.64 | (-2.88--0.38) | **1.09E-02** | **2.46E-02** | BIS only | -0.84 | (-1.58--0.11) | **2.48E-02** | **3.55E-02** | 0.89 | (-0.09-1.88) | 7.61E-02 | 1.31E-01 | notSig | 0.71 | (0.04-1.38) | **3.70E-02** | 5.70E-02 |
| PE(17:0_18:2) | -0.60 | (-1.55-0.36) | 2.21E-01 | 3.05E-01 | GUSTO only | -1.00 | (-1.56--0.44) | **4.62E-04** | **9.20E-04** | -1.70 | (-2.81--0.59) | **2.73E-03** | **7.50E-03** | GUSTO and BIS | -1.00 | (-1.47--0.53) | **3.16E-05** | **7.67E-05** |
| PE(18:0_18:1) | -0.39 | (-1.31-0.54) | 4.15E-01 | 4.98E-01 | notSig | 0.16 | (-0.37-0.69) | 5.45E-01 | 6.05E-01 | -0.34 | (-1.5-0.82) | 5.68E-01 | 6.49E-01 | notSig | -0.17 | (-0.84-0.49) | 6.09E-01 | 6.64E-01 |
| PE(18:1_18:1) | -1.29 | (-2.36--0.21) | **1.93E-02** | **3.93E-02** | BIS only | -0.64 | (-1.21--0.08) | **2.56E-02** | **3.65E-02** | 0.12 | (-1.16-1.4) | 8.50E-01 | 8.91E-01 | notSig | 0.05 | (-0.79-0.89) | 9.14E-01 | 9.28E-01 |
| PE(18:0_18:2) | -0.42 | (-1.4-0.57) | 4.00E-01 | 4.86E-01 | notSig | 0.18 | (-0.4-0.76) | 5.42E-01 | 6.03E-01 | -0.59 | (-1.8-0.61) | 3.33E-01 | 4.37E-01 | notSig | -0.52 | (-1.13-0.08) | 8.79E-02 | 1.22E-01 |
| PE(18:1_18:2) | -1.76 | (-2.8--0.71) | **1.07E-03** | **3.34E-03** | BIS only | -1.15 | (-1.79--0.5) | **4.64E-04** | **9.20E-04** | -0.12 | (-1.4-1.16) | 8.56E-01 | 8.94E-01 | notSig | -0.22 | (-0.89-0.45) | 5.24E-01 | 5.78E-01 |
| PE(16:0_20:3) | -0.96 | (-1.91--0.01) | **4.88E-02** | 8.70E-02 | notSig | -0.63 | (-1.26-0) | **4.96E-02** | 6.81E-02 | -0.15 | (-1.12-0.82) | 7.60E-01 | 8.20E-01 | notSig | -0.25 | (-0.85-0.36) | 4.25E-01 | 4.81E-01 |
| PE(16:0_20:4) | 1.51 | (0.59-2.43) | **1.21E-03** | **3.75E-03** | BIS only | 1.05 | (0.47-1.63) | **3.62E-04** | **7.47E-04** | -0.10 | (-0.8-0.59) | 7.69E-01 | 8.28E-01 | notSig | -0.23 | (-0.7-0.24) | 3.43E-01 | 4.02E-01 |
| PE(16:1_20:4) | 0.65 | (-0.52-1.84) | 2.76E-01 | 3.68E-01 | notSig | 0.13 | (-0.59-0.84) | 7.26E-01 | 7.70E-01 | 1.17 | (0.38-1.96) | **3.87E-03** | **1.03E-02** | BIS only | 0.97 | (0.39-1.54) | **1.09E-03** | **2.20E-03** |
| PE(16:0_20:5) | -0.58 | (-1.88-0.74) | 3.86E-01 | 4.74E-01 | notSig | -0.51 | (-1.33-0.3) | 2.19E-01 | 2.65E-01 | -0.96 | (-1.95-0.04) | 5.87E-02 | 1.07E-01 | notSig | -0.87 | (-1.5--0.25) | **5.70E-03** | **1.03E-02** |
| PE(17:0_20:4) | 1.72 | (0.9-2.55) | **3.79E-05** | **2.00E-04** | BIS only | 0.67 | (0.15-1.19) | **1.14E-02** | **1.75E-02** | -1.43 | (-2.16--0.7) | **1.25E-04** | **4.81E-04** | GUSTO and BIS | -1.59 | (-2.06--1.12) | **2.71E-11** | **1.04E-10** |
| PE(18:0_20:3) (a) | 0.32 | (-0.67-1.31) | 5.30E-01 | 6.08E-01 | GUSTO only | 1.13 | (0.5-1.76) | **4.28E-04** | **8.68E-04** | -0.39 | (-1.23-0.45) | 3.66E-01 | 4.72E-01 | notSig | -0.52 | (-1.04-0) | 5.10E-02 | 7.49E-02 |
| PE(18:0_20:4) | 2.38 | (1.53-3.23) | **3.77E-08** | **5.65E-07** | GUSTO and BIS | 2.16 | (1.65-2.66) | **4.96E-17** | **1.09E-15** | -0.57 | (-1.22-0.09) | 9.09E-02 | 1.49E-01 | notSig | -0.64 | (-1.09--0.19) | **5.18E-03** | **9.53E-03** |
| PE(38:5) (a) | 0.92 | (-0.05-1.89) | 6.22E-02 | 1.08E-01 | notSig | 0.34 | (-0.24-0.93) | 2.51E-01 | 3.00E-01 | 0.11 | (-0.83-1.05) | 8.14E-01 | 8.68E-01 | notSig | -0.39 | (-0.93-0.14) | 1.52E-01 | 1.99E-01 |
| PE(38:5) (b) | 0.26 | (-0.78-1.32) | 6.22E-01 | 6.95E-01 | notSig | -0.42 | (-1.08-0.25) | 2.16E-01 | 2.63E-01 | 0.79 | (-0.17-1.74) | 1.07E-01 | 1.70E-01 | notSig | 0.61 | (-0.1-1.33) | 9.38E-02 | 1.30E-01 |
| PE(16:0_22:6) | 1.18 | (0.35-2.02) | **5.47E-03** | **1.37E-02** | BIS only | 0.52 | (-0.04-1.07) | 6.72E-02 | 9.04E-02 | -1.30 | (-2.02--0.58) | **4.24E-04** | **1.43E-03** | GUSTO and BIS | -1.45 | (-1.94--0.96) | **7.06E-09** | **2.29E-08** |
| PE(17:0_22:6) | 1.36 | (0.44-2.29) | **3.86E-03** | **1.03E-02** | BIS only | 0.41 | (-0.17-1) | 1.66E-01 | 2.05E-01 | -2.48 | (-3.39--1.57) | **1.16E-07** | **8.56E-07** | GUSTO and BIS | -1.81 | (-2.32--1.31) | **1.99E-12** | **8.56E-12** |
| PE(18:0_22:4) | 0.13 | (-1.13-1.41) | 8.43E-01 | 8.75E-01 | notSig | 0.39 | (-0.33-1.11) | 2.84E-01 | 3.35E-01 | -0.55 | (-1.49-0.4) | 2.56E-01 | 3.58E-01 | notSig | -0.36 | (-0.98-0.26) | 2.58E-01 | 3.13E-01 |
| PE(18:0_22:5) (n3) | 0.13 | (-0.85-1.12) | 7.97E-01 | 8.44E-01 | notSig | 0.34 | (-0.24-0.92) | 2.48E-01 | 2.97E-01 | -1.32 | (-2.36--0.28) | **1.32E-02** | **2.91E-02** | GUSTO and BIS | -1.17 | (-1.81--0.54) | **3.11E-04** | **6.80E-04** |
| PE(18:0_22:5) (n6) | 1.06 | (-0.23-2.37) | 1.08E-01 | 1.69E-01 | notSig | 0.62 | (-0.13-1.38) | 1.07E-01 | 1.38E-01 | 0.67 | (-0.32-1.66) | 1.85E-01 | 2.74E-01 | notSig | 0.11 | (-0.59-0.81) | 7.56E-01 | 8.01E-01 |
| PE(18:0_22:6) | 2.33 | (1.39-3.27) | **1.15E-06** | **1.01E-05** | GUSTO and BIS | 1.75 | (1.14-2.35) | **1.52E-08** | **7.10E-08** | -1.47 | (-2.33--0.62) | **7.59E-04** | **2.41E-03** | GUSTO and BIS | -1.55 | (-2.12--0.97) | **1.26E-07** | **3.68E-07** |
| PE(O-34:1) | -2.21 | (-2.95--1.45) | **1.58E-08** | **2.93E-07** | GUSTO and BIS | -1.71 | (-2.17--1.25) | **2.82E-13** | **2.67E-12** | 0.48 | (-0.39-1.35) | 2.83E-01 | 3.84E-01 | notSig | 0.42 | (-0.27-1.12) | 2.34E-01 | 2.87E-01 |
| PE(O-16:0/18:2) | -3.11 | (-4.1--2.11) | **2.65E-09** | **5.59E-08** | GUSTO and BIS | -2.88 | (-3.49--2.27) | **2.43E-20** | **8.67E-19** | -0.32 | (-1.36-0.72) | 5.47E-01 | 6.32E-01 | notSig | -0.30 | (-1.22-0.62) | 5.25E-01 | 5.78E-01 |
| PE(O-18:1/18:2) | -2.82 | (-3.81--1.82) | **5.07E-08** | **7.13E-07** | GUSTO and BIS | -2.01 | (-2.6--1.42) | **2.84E-11** | **2.06E-10** | 0.11 | (-1.19-1.42) | 8.63E-01 | 8.98E-01 | notSig | 0.08 | (-0.71-0.86) | 8.45E-01 | 8.83E-01 |
| PE(O-16:0/20:4) | -1.78 | (-2.63--0.93) | **4.81E-05** | **2.39E-04** | GUSTO and BIS | -1.56 | (-2.11--1) | **3.74E-08** | **1.65E-07** | -0.37 | (-1.19-0.45) | 3.71E-01 | 4.77E-01 | notSig | -0.59 | (-1.17-0) | 5.00E-02 | 7.41E-02 |
| PE(O-36:5) | -2.14 | (-3.46--0.8) | **1.81E-03** | **5.38E-03** | GUSTO and BIS | -2.26 | (-3.1--1.42) | **1.40E-07** | **5.87E-07** | -1.46 | (-2.88--0.05) | **4.26E-02** | 8.17E-02 | notSig | -0.62 | (-1.24-0) | **4.89E-02** | 7.27E-02 |
| PE(O-16:0/22:4) | -1.30 | (-2.06--0.54) | **9.30E-04** | **3.00E-03** | BIS only | -0.88 | (-1.34--0.41) | **2.02E-04** | **4.43E-04** | 0.60 | (-0.03-1.24) | 6.39E-02 | 1.14E-01 | notSig | 0.59 | (0.09-1.08) | **1.96E-02** | **3.22E-02** |
| PE(O-18:0/20:4) | -2.73 | (-3.77--1.67) | **6.40E-07** | **6.19E-06** | GUSTO and BIS | -2.16 | (-2.8--1.51) | **6.33E-11** | **4.45E-10** | -0.36 | (-1.27-0.56) | 4.41E-01 | 5.40E-01 | notSig | -0.25 | (-0.86-0.36) | 4.19E-01 | 4.77E-01 |
| PE(O-38:5) (a) | -1.81 | (-2.64--0.96) | **3.03E-05** | **1.62E-04** | GUSTO and BIS | -1.59 | (-2.14--1.05) | **1.09E-08** | **5.15E-08** | 0.15 | (-0.74-1.04) | 7.37E-01 | 8.01E-01 | notSig | 0.00 | (-0.55-0.55) | 9.99E-01 | 9.99E-01 |
| PE(O-38:5) (b) | -2.58 | (-3.46--1.69) | **2.31E-08** | **3.83E-07** | GUSTO and BIS | -2.19 | (-2.72--1.66) | **5.21E-16** | **8.63E-15** | 1.10 | (0.32-1.88) | **5.82E-03** | **1.48E-02** | BIS only | 0.73 | (0.12-1.33) | **1.88E-02** | **3.11E-02** |
| PE(O-16:0/22:6) | -0.91 | (-1.61--0.19) | **1.31E-02** | **2.89E-02** | GUSTO and BIS | -0.98 | (-1.45--0.5) | **5.25E-05** | **1.27E-04** | 1.08 | (0.4-1.77) | **2.01E-03** | **5.82E-03** | BIS only | 0.86 | (0.34-1.38) | **1.29E-03** | **2.57E-03** |
| PE(O-18:0/22:6) | -2.02 | (-2.77--1.26) | **2.50E-07** | **2.90E-06** | GUSTO and BIS | -1.73 | (-2.2--1.25) | **8.23E-13** | **7.34E-12** | 0.67 | (-0.05-1.38) | 6.69E-02 | 1.18E-01 | notSig | 0.31 | (-0.17-0.79) | 2.10E-01 | 2.64E-01 |
| PE(O-18:1/22:6) | -1.03 | (-1.86--0.2) | **1.49E-02** | **3.20E-02** | GUSTO and BIS | -1.12 | (-1.61--0.63) | **6.55E-06** | **1.89E-05** | 0.99 | (0.3-1.68) | **5.18E-03** | **1.34E-02** | BIS only | 0.46 | (0.01-0.91) | **4.42E-02** | 6.63E-02 |
| PE(P-16:0/18:1) | -1.81 | (-2.49--1.14) | **2.31E-07** | **2.75E-06** | GUSTO and BIS | -1.33 | (-1.74--0.92) | **2.70E-10** | **1.67E-09** | 0.95 | (0.21-1.68) | **1.22E-02** | **2.75E-02** | GUSTO and BIS | 1.32 | (0.8-1.84) | **5.85E-07** | **1.62E-06** |
| PE(P-16:0/18:2) | -3.01 | (-3.92--2.09) | **3.43E-10** | **1.22E-08** | GUSTO and BIS | -2.05 | (-2.59--1.51) | **1.25E-13** | **1.27E-12** | -0.23 | (-1.23-0.78) | 6.55E-01 | 7.31E-01 | notSig | -0.03 | (-0.7-0.65) | 9.39E-01 | 9.45E-01 |
| PE(P-16:0/20:4) | -0.54 | (-1.35-0.28) | 1.97E-01 | 2.76E-01 | notSig | -0.16 | (-0.7-0.39) | 5.70E-01 | 6.28E-01 | -0.21 | (-0.98-0.55) | 5.81E-01 | 6.62E-01 | notSig | -0.22 | (-0.77-0.32) | 4.24E-01 | 4.81E-01 |
| PE(P-16:0/20:5) | -1.52 | (-2.9--0.12) | **3.36E-02** | 6.34E-02 | notSig | -1.25 | (-2.21--0.29) | **1.09E-02** | **1.69E-02** | -1.53 | (-2.72--0.33) | **1.24E-02** | **2.78E-02** | BIS only | -1.32 | (-2.33--0.31) | **1.03E-02** | **1.75E-02** |
| PE(P-16:0/22:4) | -0.51 | (-1.12-0.11) | 1.07E-01 | 1.68E-01 | notSig | -0.02 | (-0.44-0.39) | 9.13E-01 | 9.29E-01 | 0.20 | (-0.4-0.8) | 5.10E-01 | 6.06E-01 | notSig | 0.21 | (-0.25-0.67) | 3.69E-01 | 4.25E-01 |
| PE(P-16:0/22:5) (n3) | -0.75 | (-1.45--0.05) | **3.53E-02** | 6.58E-02 | notSig | -0.67 | (-1.13--0.22) | **3.41E-03** | **5.86E-03** | -0.42 | (-1.34-0.49) | 3.64E-01 | 4.71E-01 | notSig | -0.46 | (-1.03-0.11) | 1.17E-01 | 1.59E-01 |
| PE(P-16:0/22:5) (n6) | -1.25 | (-2.1--0.39) | **4.62E-03** | **1.20E-02** | GUSTO and BIS | -0.93 | (-1.45--0.42) | **3.56E-04** | **7.37E-04** | 1.34 | (0.71-1.96) | **2.94E-05** | **1.32E-04** | GUSTO and BIS | 1.19 | (0.69-1.69) | **3.53E-06** | **9.25E-06** |
| PE(P-16:0/22:6) | 0.45 | (-0.17-1.07) | 1.53E-01 | 2.25E-01 | notSig | 0.26 | (-0.17-0.68) | 2.32E-01 | 2.79E-01 | 0.77 | (0.1-1.44) | **2.50E-02** | 5.11E-02 | notSig | 0.48 | (0-0.96) | 5.24E-02 | 7.67E-02 |
| PE(P-17:0/20:4) (a) | -0.84 | (-1.8-0.13) | 8.81E-02 | 1.42E-01 | notSig | -0.53 | (-1.17-0.11) | 1.05E-01 | 1.36E-01 | -1.32 | (-2.15--0.49) | **1.84E-03** | **5.41E-03** | BIS only | -0.72 | (-1.21--0.23) | **4.12E-03** | **7.69E-03** |
| PE(P-17:0/20:4) (b) | -0.50 | (-1.49-0.5) | 3.25E-01 | 4.16E-01 | notSig | -0.32 | (-0.95-0.3) | 3.12E-01 | 3.60E-01 | -1.13 | (-1.94--0.32) | **6.49E-03** | **1.61E-02** | GUSTO and BIS | -1.02 | (-1.52--0.52) | **5.78E-05** | **1.38E-04** |
| PE(P-17:0/22:6) (a) | -0.39 | (-1.14-0.36) | 3.06E-01 | 3.98E-01 | GUSTO only | -0.79 | (-1.34--0.25) | **4.32E-03** | **7.34E-03** | -0.87 | (-1.63--0.11) | **2.46E-02** | 5.07E-02 | notSig | -0.58 | (-1.12--0.03) | **3.76E-02** | 5.75E-02 |
| PE(P-17:0/22:6) (b) | -0.12 | (-0.83-0.61) | 7.54E-01 | 8.10E-01 | notSig | -0.43 | (-0.94-0.08) | 9.62E-02 | 1.25E-01 | -0.68 | (-1.44-0.08) | 8.09E-02 | 1.35E-01 | notSig | -0.55 | (-1.06--0.04) | **3.54E-02** | 5.51E-02 |
| PE(P-18:0/18:1) | -1.84 | (-2.62--1.06) | **5.27E-06** | **3.88E-05** | GUSTO and BIS | -1.47 | (-1.94--1) | **1.05E-09** | **5.94E-09** | 0.19 | (-0.8-1.17) | 7.10E-01 | 7.82E-01 | GUSTO only | 0.68 | (0.06-1.3) | **3.12E-02** | **4.91E-02** |
| PE(P-18:0/18:2) | -3.19 | (-4.16--2.2) | **6.48E-10** | **2.00E-08** | GUSTO and BIS | -2.24 | (-2.78--1.7) | **3.46E-16** | **5.95E-15** | -1.08 | (-2.39-0.23) | 1.07E-01 | 1.70E-01 | notSig | -0.44 | (-1.17-0.29) | 2.40E-01 | 2.94E-01 |
| PE(P-18:0/20:4) | -0.75 | (-1.65-0.15) | 1.03E-01 | 1.63E-01 | notSig | -0.42 | (-1.01-0.16) | 1.55E-01 | 1.93E-01 | -0.57 | (-1.42-0.28) | 1.87E-01 | 2.77E-01 | notSig | -0.62 | (-1.16--0.08) | **2.48E-02** | **4.01E-02** |
| PE(P-18:0/20:5) | -1.77 | (-3.19--0.32) | **1.70E-02** | **3.58E-02** | GUSTO and BIS | -1.70 | (-2.69--0.71) | **7.76E-04** | **1.48E-03** | -1.78 | (-3.05--0.51) | **6.10E-03** | **1.54E-02** | GUSTO and BIS | -1.86 | (-2.82--0.9) | **1.42E-04** | **3.23E-04** |
| PE(P-18:0/22:4) | -0.89 | (-1.55--0.23) | **8.02E-03** | **1.86E-02** | GUSTO and BIS | -0.80 | (-1.24--0.36) | **3.71E-04** | **7.61E-04** | -0.31 | (-1.13-0.51) | 4.52E-01 | 5.49E-01 | notSig | -0.30 | (-0.87-0.27) | 3.05E-01 | 3.63E-01 |
| PE(P-18:0/22:5) (n3) | -1.54 | (-2.33--0.75) | **1.44E-04** | **6.01E-04** | GUSTO and BIS | -1.08 | (-1.55--0.6) | **9.03E-06** | **2.55E-05** | -1.10 | (-2.14--0.07) | **3.66E-02** | 7.14E-02 | notSig | -0.59 | (-1.13--0.06) | **2.86E-02** | **4.54E-02** |

| | | | | | | | | | | | | | | | | | | |
|---|---|---|---|---|---|---|---|---|---|---|---|---|---|---|---|---|---|---|
| PE(P-18:0/22:5) (n6) | -1.95 | (-2.79--1.11) | **6.84E-06** | **4.67E-05** | GUSTO and BIS | -1.51 | (-1.98--1.03) | **5.07E-10** | **2.94E-09** | 0.68 | (0.01-1.36) | **4.72E-02** | 9.00E-02 | notSig | 0.52 | (0.05-0.99) | **3.06E-02** | **4.84E-02** |
| PE(P-18:0/22:6) | -1.05 | (-1.77--0.32) | **4.98E-03** | **1.27E-02** | GUSTO and BIS | -0.90 | (-1.39--0.42) | **2.55E-04** | **5.43E-04** | 0.02 | (-0.7-0.75) | 9.46E-01 | 9.56E-01 | notSig | -0.02 | (-0.51-0.46) | 9.30E-01 | 9.40E-01 |
| PE(P-18:1/18:1) | -1.46 | (-2.18--0.74) | **8.73E-05** | **3.93E-04** | GUSTO and BIS | -1.11 | (-1.55--0.67) | **7.10E-07** | **2.39E-06** | 0.72 | (-0.19-1.62) | 1.20E-01 | 1.86E-01 | notSig | 0.48 | (-0.04-1) | 7.13E-02 | 1.01E-01 |
| PE(P-18:1/22:4) | 0.05 | (-0.54-0.64) | 8.67E-01 | 8.96E-01 | notSig | -0.10 | (-0.54-0.34) | 6.57E-01 | 7.07E-01 | 0.48 | (-0.11-1.07) | 1.14E-01 | 1.80E-01 | notSig | 0.44 | (-0.06-0.93) | 8.65E-02 | 1.20E-01 |
| PE(P-18:1/22:5) (a) | -0.66 | (-1.33-0.02) | 5.54E-02 | 9.80E-02 | GUSTO only | -0.69 | (-1.14--0.23) | **3.15E-03** | **5.42E-03** | -0.42 | (-1.4-0.57) | 4.04E-01 | 5.07E-01 | notSig | -0.39 | (-0.93-0.15) | 1.54E-01 | 2.01E-01 |
| PE(P-18:1/22:6) (a) | 0.34 | (-0.4-1.08) | 3.69E-01 | 4.59E-01 | notSig | -0.13 | (-0.62-0.36) | 5.91E-01 | 6.45E-01 | 0.73 | (0.08-1.38) | **2.74E-02** | 5.55E-02 | notSig | 0.37 | (-0.11-0.85) | 1.29E-01 | 1.74E-01 |
| PE(P-20:0/18:1) | -1.69 | (-2.4--0.97) | **4.75E-06** | **3.55E-05** | GUSTO and BIS | -1.45 | (-1.92--0.98) | **1.21E-09** | **6.70E-09** | -0.28 | (-1-0.45) | 4.50E-01 | 5.48E-01 | notSig | -0.17 | (-0.84-0.5) | 6.29E-01 | 6.82E-01 |
| PE(P-20:0/18:2) | -2.85 | (-3.78--1.9) | **6.34E-09** | **1.28E-07** | GUSTO and BIS | -2.07 | (-2.55--1.6) | **1.23E-17** | **3.00E-16** | -1.15 | (-2.34-0.03) | 5.63E-02 | 1.03E-01 | notSig | -0.16 | (-0.58-0.25) | 4.35E-01 | 4.91E-01 |
| PE(P-20:0/20:4) | -0.84 | (-1.62--0.06) | **3.56E-02** | 6.61E-02 | notSig | -0.42 | (-0.89-0.06) | 8.63E-02 | 1.13E-01 | -0.60 | (-1.34-0.14) | 1.14E-01 | 1.80E-01 | notSig | -0.60 | (-1.17--0.03) | **3.95E-02** | 6.03E-02 |
| PE(P-20:0/22:6) | -0.97 | (-1.7--0.23) | **1.07E-02** | **2.42E-02** | GUSTO and BIS | -0.84 | (-1.27--0.42) | **1.10E-04** | **2.57E-04** | -0.45 | (-1.11-0.2) | 1.76E-01 | 2.65E-01 | GUSTO only | -0.59 | (-1.02--0.16) | **7.17E-03** | **1.26E-02** |
| PE(P-20:1/20:4) | -0.92 | (-1.92-0.09) | 7.48E-02 | 1.23E-01 | notSig | -0.63 | (-1.29-0.03) | 6.27E-02 | 8.48E-02 | 0.15 | (-0.6-0.91) | 6.91E-01 | 7.64E-01 | notSig | -0.17 | (-0.79-0.45) | 5.97E-01 | 6.53E-01 |
| PG(36:1) | 0.58 | (-0.39-1.56) | 2.39E-01 | 3.26E-01 | notSig | -0.01 | (-0.56-0.54) | 9.70E-01 | 9.74E-01 | -0.71 | (-1.79-0.37) | 1.96E-01 | 2.87E-01 | notSig | -1.01 | (-1.81--0.21) | **1.38E-02** | **2.32E-02** |
| PI(16:0/16:0) | -2.67 | (-3.93--1.39) | **5.08E-05** | **2.48E-04** | GUSTO and BIS | -2.52 | (-3.27--1.77) | **4.31E-11** | **3.08E-10** | -1.03 | (-2.08-0.01) | 5.29E-02 | 9.86E-02 | notSig | -0.53 | (-1.25-0.18) | 1.44E-01 | 1.89E-01 |
| PI(16:0_16:1) | -1.35 | (-2.78-0.1) | 6.77E-02 | 1.15E-01 | GUSTO only | -1.60 | (-2.45--0.74) | **2.67E-04** | **5.63E-04** | 0.45 | (-0.31-1.22) | 2.46E-01 | 3.47E-01 | notSig | 0.57 | (-0.01-1.14) | 5.26E-02 | 7.68E-02 |
| PI(34:0) | -1.97 | (-3.12--0.81) | **9.36E-04** | **3.00E-03** | GUSTO and BIS | -2.39 | (-3.11--1.66) | **1.06E-10** | **6.93E-10** | -1.46 | (-2.58--0.34) | **1.06E-02** | **2.46E-02** | GUSTO and BIS | -1.64 | (-2.47--0.81) | **1.00E-04** | **2.33E-04** |
| PI(34:1) | -1.27 | (-2.17--0.37) | **6.11E-03** | **1.49E-02** | GUSTO and BIS | -1.75 | (-2.29--1.21) | **1.84E-10** | **1.19E-09** | -0.40 | (-1.36-0.56) | 4.09E-01 | 5.10E-01 | notSig | -0.26 | (-0.79-0.27) | 3.41E-01 | 4.01E-01 |
| PI(17:0_18:1) | -1.25 | (-2.12--0.38) | **5.16E-03** | **1.31E-02** | GUSTO and BIS | -2.03 | (-2.51--1.55) | **1.31E-16** | **2.63E-15** | -1.48 | (-2.45--0.51) | **2.82E-03** | **7.66E-03** | GUSTO and BIS | -1.40 | (-2.1--0.69) | **9.88E-05** | **2.30E-04** |
| PI(17:0_18:2) | -1.54 | (-2.27--0.8) | **4.83E-05** | **2.39E-04** | GUSTO and BIS | -2.21 | (-2.69--1.73) | **1.21E-19** | **3.74E-18** | -2.27 | (-3.22--1.32) | **3.19E-06** | **1.78E-05** | GUSTO and BIS | -2.10 | (-2.8--1.41) | **2.92E-09** | **9.81E-09** |
| PI(18:0_18:1) | -1.55 | (-2.47--0.61) | **1.23E-03** | **3.77E-03** | GUSTO and BIS | -1.80 | (-2.36--1.24) | **3.52E-10** | **2.12E-09** | -0.70 | (-1.98-0.58) | 2.86E-01 | 3.87E-01 | GUSTO only | -1.07 | (-1.68--0.46) | **5.58E-04** | **1.19E-03** |
| PI(36:2) | -1.10 | (-1.87--0.32) | **6.05E-03** | **1.49E-02** | GUSTO and BIS | -1.63 | (-2.1--1.16) | **8.04E-12** | **6.66E-11** | -1.57 | (-2.58--0.57) | **2.22E-03** | **6.33E-03** | GUSTO and BIS | -1.31 | (-1.81--0.8) | **3.64E-07** | **1.03E-06** |
| PI(18:1_18:2) | -2.97 | (-3.88--2.06) | **5.26E-10** | **1.74E-08** | GUSTO and BIS | -2.76 | (-3.28--2.24) | **1.37E-25** | **1.06E-23** | -1.44 | (-2.14--0.73) | **6.74E-05** | **2.70E-04** | GUSTO and BIS | -1.46 | (-1.88--1.04) | **1.07E-11** | **4.36E-11** |
| PI(16:0_20:3) (a) | -2.61 | (-3.42--1.79) | **9.45E-10** | **2.34E-08** | GUSTO and BIS | -2.34 | (-2.85--1.82) | **4.52E-19** | **1.31E-17** | 0.12 | (-0.47-0.72) | 6.86E-01 | 7.60E-01 | notSig | 0.18 | (-0.23-0.58) | 3.90E-01 | 4.48E-01 |
| PI(16:0_20:3) (b) | -3.52 | (-4.53--2.5) | **4.64E-11** | **3.37E-09** | GUSTO and BIS | -2.83 | (-3.43--2.23) | **1.80E-20** | **6.97E-19** | 2.95 | (2.17-3.74) | **4.47E-13** | **7.41E-12** | GUSTO and BIS | 2.97 | (2.33-3.61) | **9.90E-20** | **6.38E-19** |
| PI(16:0_20:4) | -0.83 | (-1.74-0.09) | 7.58E-02 | 1.24E-01 | GUSTO only | -1.27 | (-1.76--0.77) | **4.90E-07** | **1.76E-06** | -0.04 | (-0.53-0.46) | 8.87E-01 | 9.11E-01 | notSig | 0.03 | (-0.34-0.4) | 8.62E-01 | 8.91E-01 |
| PI(18:0_20:2) | -1.77 | (-2.69--0.85) | **1.90E-04** | **7.62E-04** | GUSTO and BIS | -2.33 | (-2.89--1.77) | **2.72E-16** | **4.86E-15** | 2.04 | (1.15-2.93) | **8.29E-06** | **4.23E-05** | GUSTO and BIS | 2.17 | (1.43-2.91) | **9.52E-09** | **3.01E-08** |
| PI(18:0_20:3) (a) | -1.70 | (-2.59--0.8) | **2.40E-04** | **9.43E-04** | GUSTO and BIS | -1.11 | (-1.55--0.67) | **8.47E-07** | **2.81E-06** | -0.35 | (-1.06-0.35) | 3.27E-01 | 4.32E-01 | notSig | -0.30 | (-0.7-0.1) | 1.38E-01 | 1.83E-01 |
| PI(18:0_20:4) | 0.79 | (0.11-1.46) | **2.22E-02** | **4.38E-02** | GUSTO and BIS | -0.21 | (-0.62-0.2) | 3.10E-01 | 3.59E-01 | -0.38 | (-0.82-0.05) | 8.57E-02 | 1.42E-01 | GUSTO only | -0.48 | (-0.79--0.18) | **2.12E-03** | **4.07E-03** |
| PI(38:5) (a) | -1.71 | (-2.48--0.94) | **1.60E-05** | **9.53E-05** | GUSTO and BIS | -2.19 | (-2.63--1.74) | **6.50E-22** | **3.02E-20** | 0.02 | (-0.51-0.56) | 9.28E-01 | 9.44E-01 | notSig | -0.25 | (-0.65-0.14) | 2.11E-01 | 2.64E-01 |
| PI(38:6) | -1.57 | (-2.46--0.67) | **6.34E-04** | **2.21E-03** | GUSTO and BIS | -2.26 | (-2.78--1.74) | **2.27E-17** | **5.27E-16** | -1.31 | (-1.85--0.77) | **2.00E-06** | **1.15E-05** | GUSTO and BIS | -1.12 | (-1.5--0.74) | **7.75E-09** | **2.48E-08** |
| PI(18:0_22:4) | -0.78 | (-1.53--0.02) | **4.48E-02** | 8.05E-02 | GUSTO only | -1.04 | (-1.53--0.55) | **3.56E-05** | **8.91E-05** | 0.96 | (0.44-1.48) | **3.38E-04** | **1.18E-03** | GUSTO and BIS | 1.04 | (0.62-1.46) | **1.02E-06** | **2.78E-06** |
| PI(18:0_22:5) (n3) | -0.80 | (-1.55--0.04) | **3.81E-02** | 6.96E-02 | GUSTO only | -1.32 | (-1.82--0.82) | **2.13E-07** | **8.44E-07** | -1.29 | (-2.04--0.53) | **8.62E-04** | **2.70E-03** | GUSTO and BIS | -1.26 | (-1.76--0.76) | **6.62E-07** | **1.83E-06** |
| PI(18:0_22:6) | -0.69 | (-1.49-0.12) | 9.45E-02 | 1.51E-01 | GUSTO only | -1.50 | (-2--0.99) | **6.63E-09** | **3.31E-08** | -0.85 | (-1.42--0.27) | **4.13E-03** | **1.09E-02** | GUSTO and BIS | -0.98 | (-1.38--0.59) | **9.58E-07** | **2.61E-06** |
